# Early metazoan origin and multiple losses of a novel clade of RIM pre-synaptic calcium channel scaffolding protein homologues

**DOI:** 10.1101/2020.01.14.906610

**Authors:** Thomas Piekut, Yuen Yan Wong, Sarah E. Walker, Carolyn L. Smith, Julia Gauberg, Alicia N. Harracksingh, Christopher Lowden, Hai-Ying Mary Cheng, Gaynor E. Spencer, Adriano Senatore

**Author notes:** To whom correspondence should be addressed: Adriano Senatore: Department of Biology, University of Toronto Mississauga, 3359 Mississauga rd. Mississauga Ontario, L5L 1C6 Canada. These authors contributed equally to the work (Thomas Piekut, Yuen Yan Wong, Sarah E. Walker, Carolyn L. Smith).

## Abstract

The precise localization of Ca_V_2 voltage-gated calcium channels at the synapse active zone requires various interacting proteins, of which, <u>R</u>ab3 interacting <u>m</u>olecule or RIM is considered particularly important. In vertebrates, RIM interacts with Ca_V_2 channels *in vitro* via a PDZ domain that binds to the extreme C-termini of the channels at acidic ligand motifs of D/E-D/E/H-WC-_COOH_, and knockout of RIM in vertebrates and invertebrates disrupts Ca_V_2 channel synaptic localization and synapse function. Here, we describe a previously uncharacterized clade of RIM proteins bearing homologous domain architectures as known RIM homologues, but some notable differences including key amino acids associated with PDZ domain ligand specificity. This novel RIM emerged near the stem lineage of metazoans and underwent extensive losses, but is retained in select animals including the early-diverging placozoan *Trichoplax adhaerens*, and molluscs. RNA expression and localization studies in *Trichoplax* and the mollusc snail *Lymnaea stagnalis* indicate differential regional/tissue type expression, but overlapping expression in single isolated neurons from *Lymnaea*. Ctenophores, the most early-diverging animals with synapses, are unique among animals with nervous systems in that they lack the canonical RIM, bearing only the newly identified homologue. Through phylogenetic analysis, we find that Ca_V_2 channel D/E-D/E/H-WC-_COOH_ like PDZ ligand motifs were present in the common ancestor of cnidarians and bilaterians, and delineate some deeply conserved C-terminal structures that distinguish Ca_V_1 from Ca_V_2 channels, and Ca_V_1/Ca_V_2 from Ca_V_3 channels.

## Introduction

The tight spatiotemporal regulation of cytoplasmic Ca^2+^ fluxes is integral to ensuring that Ca^2+^-dependent biological processes are effected with fidelity, and preventing the toxicity that arises with prolonged elevated levels of intracellular Ca^2+^ (Clapham 2007). A variety of differentially gated ion channels are the route for Ca^2+^ entry into the cytoplasm. Of these, voltage-gated calcium (Ca_V_) channels mediate Ca^2+^ influx that underlies such fundamental processes as neurotransmitter release (Katz and Miledi 1965) and excitation-contraction coupling (Catterall 2011), and whose dysfunction is causal to variegated pathologies (Adams and Snutch 2007; Simms and Zamponi 2014). All Ca_V_s are defined by a current-conducting α subunit comprised of four homologous domains each containing six transmembrane segments (S1-S6) connected by cytoplasmic linkers. These linkers, along with their cytoplasmic N- and C-termini, are largely disordered in structure (Catterall 2011). The high-voltage activated (HVA) L-type (Ca_V_1.1-1.4), P/Q-type (Ca_V_2.1), N-type (Ca_V_2.2) and R-type (Ca_V_2.3) channels associate with Ca_V_α_2_δ and Ca_V_β ancillary subunits, the latter via the alpha interaction domain (AID) located in the domain I-II linker of the channel α subunit. Calmodulin, a Ca^2+^ sensor important for modulating Ca_V_ channel function, interacts with HVA α subunits at C-terminal IQ motifs (Catterall 2011; Ben-Johny and Yue 2014). Recently, low-voltage activated (LVA) T-type (Ca_V_3) channels have also been shown to interact with the Ca_V_β subunit (Bae, et al. 2010), as well as calmodulin (Chemin, et al. 2017), though they lack AID and IQ motifs. Concomitant to phylogenetic, biophysical and pharmacological distinctions between Ca_V_1-Ca_V_3 channels, it is apparent that distinct sets of interacting proteins are integral to their unique functions in different cell types that appear broadly conserved in the Metazoa (Senatore, et al. 2016).

The presynaptic active zone is the locus for synaptic vesicle exocytosis mediated largely by Ca_V_2 type calcium channels in vertebrate and invertebrate synapses (Figure 1A) (Spafford and Zamponi 2003; Südhof 2012). A leading functional model for Ca_V_2 channel tethering at the active zone involves the tripartite interaction between <u>R</u>ab3 interacting <u>m</u>olecule (RIM), RIM-binding-protein (RIM-BP) and Ca_V_2 channels, thought to ensure that depolarization-induced cytoplasmic Ca^2+^ plumes are close to Ca^2+^ sensors of the exocytotic machinery (Südhof 2012). Specifically, RIM is thought to selectively recruit N- and P/Q-type Ca_V_2 channels in vertebrates, or the single Ca_V_2 channel in invertebrates, via a PDZ (<u>p</u>ost synaptic density 95 protein, *Drosophila* <u>d</u>isc large tumor suppressor, and <u>z</u>onula occludens-1 protein) domain that interacts with amino acid motifs of D/E-D/E/H-WC-_COOH_ located on the extreme C-termini of the calcium channels (Kaeser, et al. 2011; Graf, et al. 2012). Additionally, RIM is involved in priming and docking of synaptic vesicles, binding to the vesicular protein Rab3 with an N-terminal alpha helical structure, and the SNARE-associated protein Munc-13 with an adjacent Zn^2+^-finger domain (Figure 1A) (Wang, et al. 1997; Betz, et al. 2001; Wang, et al. 2001; Fukuda 2003; Dulubova, et al. 2005; Lu, et al. 2006; Quade, et al. 2019). RIM-BP, which bears three Src homology 3 (SH3) protein interaction domains, binds proline-rich motifs in RIMs and the distal Ca_V_2 C-terminus to strengthen the RIM-Ca_V_2 interaction (Wang, et al. 2000; Hibino, et al. 2002; Kaeser, et al. 2011) (Figure 1A). In vertebrates, RIMs have also been shown to suppress Ca_V_2 channel voltage-dependent inactivation to potentiate neurotransmitter release, mediated by an interaction between the RIM C_2_B domain and the Ca_V_β subunit (Kiyonaka, et al. 2007; Uriu, et al. 2010). Vertebrate RIM1 and RIM2 also interact with two separate regions in the N- and P/Q-channel C-terminus (encoded by exons 44 and 47), which for RIM2, further suppresses Ca_V_2 channel voltage-dependent inactivation (Hirano, et al. 2017).

**Figure 1.**
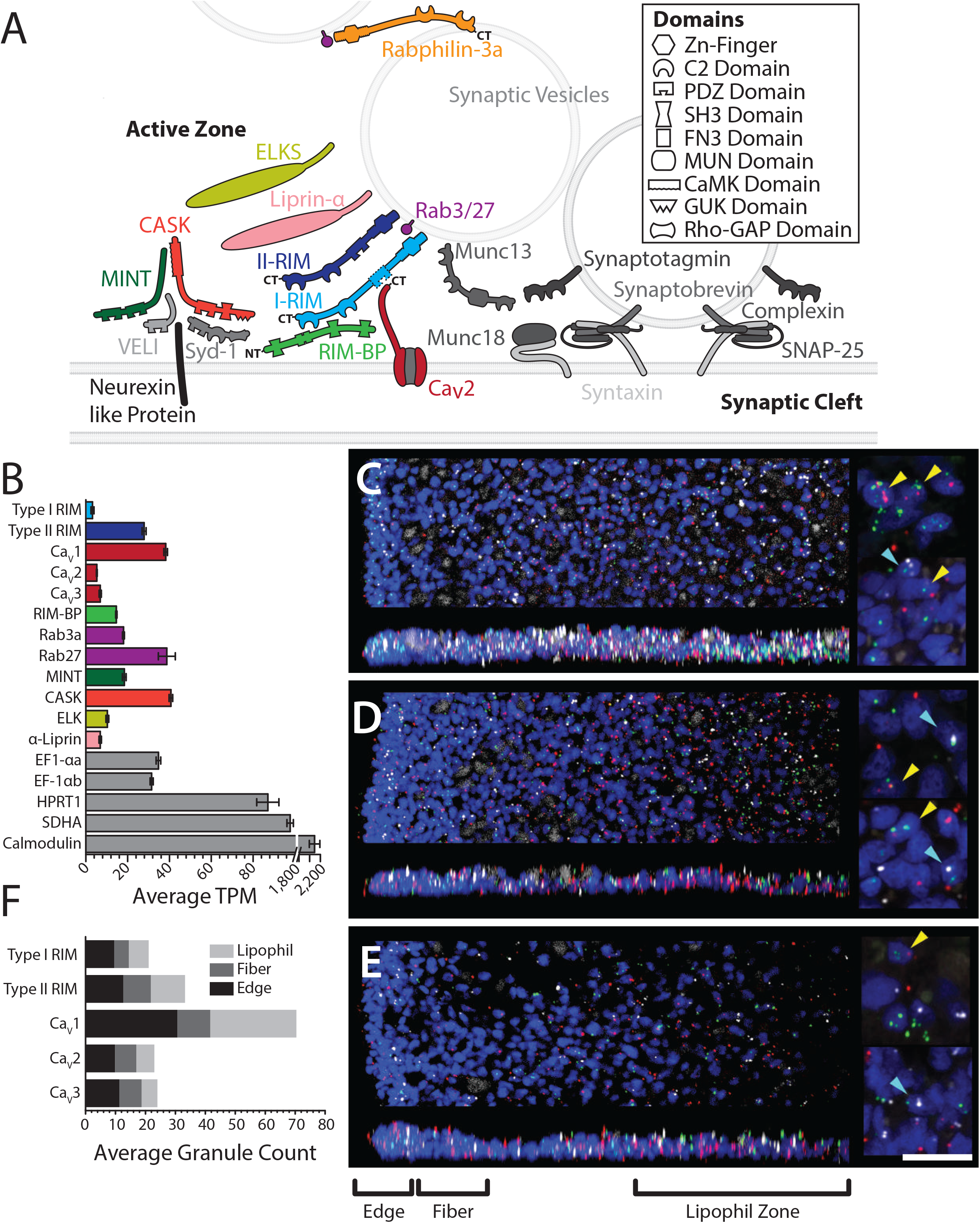
**A)** Schematic of pre-synaptic active zone proteins identified in the transcriptome of *Trichoplax adhaerens* (Wong, et al. 2019). Colored proteins represent key pre-synaptic scaffolding proteins that interact with N-, P/Q-, and R-type Ca_V_2 channels. Presence of InterPro-predicted canonical domain structures for each protein are illustrated. The dashed line denotes the absence of a predicted PDZ domain for the *Trichoplax* I-RIM homologue. **B)** Average transcripts per million (TPM) expression level analysis of the *Trichoplax* whole animal transcriptome (Wong, et al. 2019) reveals expression of a rich set of active zone proteins, plus all three voltage-gated calcium channel paralogues (the color scheme follows that of the Ca_V_-interactome depicted in panel A. The housekeeping genes eukaryotic translation elongation factor-1α (EF-1αa) and EF-1αb, hypoxanthine phosphoribosyltransferase 1 (HPRT1), and succinate dehydrogenase (SDHA) were used as standards for expression level. **C-E)** Fluorescence *in situ* hybridization (FISH) with RNAScope probes for RIM and voltage-gated calcium channel (Ca_V_1, Ca_V_2 or Ca_V_3) genes in wholemounts of *Trichoplax*. **C)** Ca_V_1 (green), II-RIM (red) and I-RIM (white). **D)** Ca_V_2 (green), II-RIM (red) and I-RIM (white). **E)** Ca_V_3 (green), II-RIM (red) and I-RIM (white). Nuclei are blue. The top image in each part shows a horizontal (x,y) projection of a series of optical sections through a region beginning at one edge and extending halfway across the animal, and the lower image shows a vertical (x,z) projection of a 10 µm (C) or 15 µm (D, E) wide strip through the same region. Color-separated images of the same samples are shown in Figure S1. Insets (right) show enlarged views captured with an enhanced resolution detector. In each part, the top inset is a projection of five optical sections (0.185 µm interval) through the most ventral nuclei in a region within the lipophil zone and the lower inset is projection of five optical sections in a region at the edge. Nuclei that are in close apposition to labels for both a Ca_V_ channel and II-RIM or a Ca_V_ channel and I-RIM are indicated (yellow and cyan arrowheads, respectively). Scale, 20 µm (left); 5 µm (right). **F)** Average fluorescent granules for I-RIM, II-RIM and the Ca_V_1-3 channels counted within the edge, fiber zone and lipophil zone regions of fluorescently labeled *Trichoplax* from five (Ca_V_1-3) and seven (I-RIM and II-RIM) separate experiments (i.e. average grains per 10 µm^2^).

Ubiquitously conserved and unique to animals (Paps and Holland 2018), the presence of RIMs has been tied to the origin of animal multicellularity and accords with the underrepresentation of active zone proteins in choanoflagellates, a sister group to the Metazoa otherwise harboring a rich set of synaptic proteins (Burkhardt 2015). RIMs likely also serve non-neuronal functions, given that homologues are found in animals lacking synapses (Paps and Holland 2018), and although predominantly expressed in the nervous systems of vertebrates, RIM expression has been reported in non-nervous tissues (Iezzi, et al. 2000). Unfortunately, a paucity of data exists regarding the conservation of the RIM, RIM-BP and Ca_V_2 interaction among early-diverging metazoans, and more generally, invertebrate phyla, hampering progress in understanding the molecular evolution of the pre-synaptic active zone. Here, via maximum likelihood and Bayesian phylogenetic analyses, we show for the first time that invertebrates possess a novel RIM homologue that emerged near the stem metazoan lineage, and has undergone multiple losses in cnidarians and bilaterians. We demonstrate that this homologue contains, with a single known phyletic exception, a domain architecture akin to that of vertebrate α-RIMs but is significantly shorter and possesses a PDZ domain that differs at amino acid positions associated with ligand specificity compared to previously characterized RIMs. Further, we provide a systematic phylogeny of metazoan voltage-gated calcium channels, complete with annotations of predicted C-terminal PDZ and SH3 domain binding motifs, to evaluate the interactomic potential of distinct Ca_V_ channel clades. On a more granular level, we examine the conservation of short linear motifs in metazoan Ca_V_ channel C-termini, and identify putative structural distinctions between the ancestral Ca_V_1 and Ca_V_2 channels, and between Ca_V_1/Ca_V_2 channels and Ca_V_3 channels.

## Results

### Identification of two RIM homologues in the transcriptome of Trichoplax adhaerens

Previous research has reported that invertebrates possess a single gene encoding the active zone protein RIM (Wang and Südhof 2003; Südhof 2012). Work in our laboratory identified two RIM homologues in the transcriptome of *Trichoplax adhaerens*, a small sea water invertebrate that diverged from other animals roughly 600 million years ago (Dos Reis, et al. 2015), and that lacks a nervous system and synapses (Smith, et al. 2014). One paralogue was found to be considerably longer in protein sequence (2,487 residues) and to lack a predicted PDZ domain for interactions with the C-termini of Ca_V_2 channels (Figure 1A) (Wong, et al. 2019). We decided to name this particular homologue type I RIM (I-RIM), based on similarities with canonical RIM genes described in other animals. The other we named type II RIM (II-RIM), as it is considerably shorter in length (1,098 residues) yet bears the expected domain architecture of an N-terminal Zn^2+^-finger domain flanked by alpha helices, followed by a PDZ domain and two C_2_ domains (Figure 1A, Supplementary file 1). The apparent absence of synapses in *Trichoplax* is not reflected by its expressed gene set, which in addition to RIM includes homologues for key active zone proteins such as SNARE and associated proteins, and scaffolding proteins that interact with Ca_V_2 calcium channels at nerve terminals such as Mint, CASK, Liprin-α, ELKS and RIM-BP (Figure 1A and B). *Trichoplax* is also the most early-diverging animal with homologues for all three metazoan Ca_V_ channel types: Ca_V_1, Ca_V_2 and Ca_V_3 (Senatore, et al. 2012; Moran and Zakon 2014; Senatore, et al. 2016; Smith, et al. 2017).

Fluorescence *in situ* hybridization (FISH) with probes for I-RIM, II-RIM and the Ca_V_1-Ca_V_3 channel mRNAs in wholemounts of *Trichoplax* (Figure 1C-E) confirmed that each gene is expressed. Ca_V_1 expression was more abundant at the edge of the animal and in the central region starting ∼80 µm from the edge compared to the intervening region. Ca_V_2, Ca_V_3, I-RIM and II-RIM had more uniform radial expression patterns (Figures 1C-E, S1A). Label for both RIMs and all three calcium channels was evident near the dorsal and ventral surfaces (Figure 1, vertical projections), suggesting expression in dorsal and ventral epithelial cells. Nuclei that were in close apposition to probes for I-RIM or II-RIM and one of the calcium channels (Ca_V_1, Ca_V_2 or Ca_V_3) were present (Figure 1C-E, right insets), suggesting that RIM proteins are co-expressed with calcium channels. However, only a small number (generally <4) of probe labels were associated with individual nuclei. The small number of probe labels likely indicates that mRNA for these proteins is in low abundance, since much higher probe label densities have been observed by the same *in situ* hybridization technique with probes for highly expressed proteins, such as digestive enzymes (Mayorova, et al. 2019). The higher abundance of Ca_V_1 expressing cells near the edge and within the lipophil zone than in the intervening region is interesting because secretory cells are prevalent near the edge and in the lipophil zone but are rare in the intervening region. Mucocytes are the most prevalent secretory cell type near the edge and can be recognized by staining with a fluorescence conjugated lectin, wheatgerm agglutinin (WGA) (Mayorova, et al. 2019). Combining FISH for calcium channels with WGA staining (Figure S1B) showed that probe labels for Ca_V_1 and Ca_V_2 were often present inside WGA stained mucocytes, whereas only a few mucocytes had Ca_V_3 probe labels in their interiors.

Average counts of fluorescent granules within regions of the animal characterized by distinct cell-type content (i.e. the edge, fiber cell zone and lipophil cell zone (Mayorova, et al. 2019)), revealed that II-RIM is more abundantly expressed than I-RIM overall and in each region (Figure 1F; p-values for Tukey’s tests after one-way ANOVAs: edge <0.05; fiber zone <0.0005; lipophil zone <0.00005; sum of regions <0.00005; ANOVA for separate regions: df = 5, F = 17.1, p = 2.1E-15; ANOVA for regions combined: df = 1, F = 24.5, p = 2.1E-6). Notably, these patterns are consistent with mRNA expression levels measured as average transcripts per million (TPM) in the transcriptome data, where II-RIM is more abundantly expressed than I-RIM at the whole animal level (Figure 1B). Also consistent were the average TPM values and counted granules for *in situ* hybridization of the three *Trichoplax* Ca_V_ channels (Ca_V_1-Ca_V_3), where TPM and granule counts for Ca_V_1 were significantly higher compared to Ca_V_2 and Ca_V_3 within edge and lipophil zones and all three regions combined (i.e. compare Figure 1B and F) (p-values for Tukey’s tests after one-way ANOVAs of granule counts: Ca_V_1 vs. Ca_V_2 edge <0.00005; Ca_V_1 vs. Ca_V_2 lipophil zone <0.00005; Ca_V_1 vs. Ca_V_2 total <0.00005; Ca_V_1 vs. Ca_V_3 edge <0.00005; Ca_V_1 vs. Ca_V_3 lipophil zone <0.00005; Ca_V_1 vs. Ca_V_3 total <0.00005; ANOVA for separate regions: df = 8, F = 86.8, p = 0; ANOVA for regions combined: df = 2, F = 159.0, p = 0;). Indeed, in spite of low-level expression, the consistency between the transcriptome TPM and fluorescent granule count data indicates that fluorescent signals observed in the hybridization experiments reflect true mRNAs expression. Lastly, I-RIM, Ca_V_2 and Ca_V_3 each appear to be enriched in the edge compared to the lipophil zone, with average respective granule count ratios of 1.5, 1.7 and 2.1 (edge/lipophil zone), compared to II-RIM and Ca_V_1 (ratios of 1.0 and 1.1 respectively).

### Invertebrates possess a second RIM homologue with a conserved domain architecture

The existence of two RIM homologues in *Trichoplax* prompted us to determine whether the II-RIM gene is also found in other animals, and whether these are phylogenetically distinct from I-RIMs. Both maximum likelihood and Bayesian protein phylogenetic inference demonstrated that numerous invertebrates possess previously undescribed II-RIMs, which form a sister clade with canonical vertebrate RIM1/RIM2 and invertebrate RIM genes (i.e. I-RIMs) (Figure 2). Vertebrate RIM3 and RIM4 proteins were not included in the phylogeny due to their truncated nature, although they clustered with vertebrate RIM1 and RIM2 sequences in preliminary analyses (not shown), consistent with the notion that these evolved via gene duplication along the vertebrate stem lineage (Wang and Südhof 2003). The related protein, rabphilin 3A, a regulator of synaptic vesicle recruitment that like I-RIMs binds Rab3 in a GTP-dependent manner (Shirataki, et al. 1993; Stahl, et al. 1996; Burns, et al. 1998; Wang, et al. 2001), was used as an outgroup (Figure 2). Excluding sponge homologues, both I and II-RIMs cluster together 99% and 93% of the time, respectively, with the rabphilins as a sister clade. With a few notable exceptions, both I and II-RIMs feature conserved domain architectures comprised of an N-terminal Zn^2+^-finger domain, a PDZ domain, and two C_2_ domains, including the vertebrate homologues RIM1αβ and RIM2α. That I-RIM, II-RIM and rabphilin form distinct clades was also indirectly supported by analysis of average protein length. II-RIM (n = 14, 1,106 aa ± 166 aa) was found to be significantly shorter than I-RIM (n = 33, 1,541 aa ± 344 aa) but significantly longer than rabphilin (n = 28, 664 aa ± 104 aa) (Kruskal-Wallis χ^2^ = 57.656, df = 2, p-value = 3.021E-13 and Dunn’s post-hoc test with the Benjamini-Hochberg adjustment) (Figure 2).

**Figure 2.**
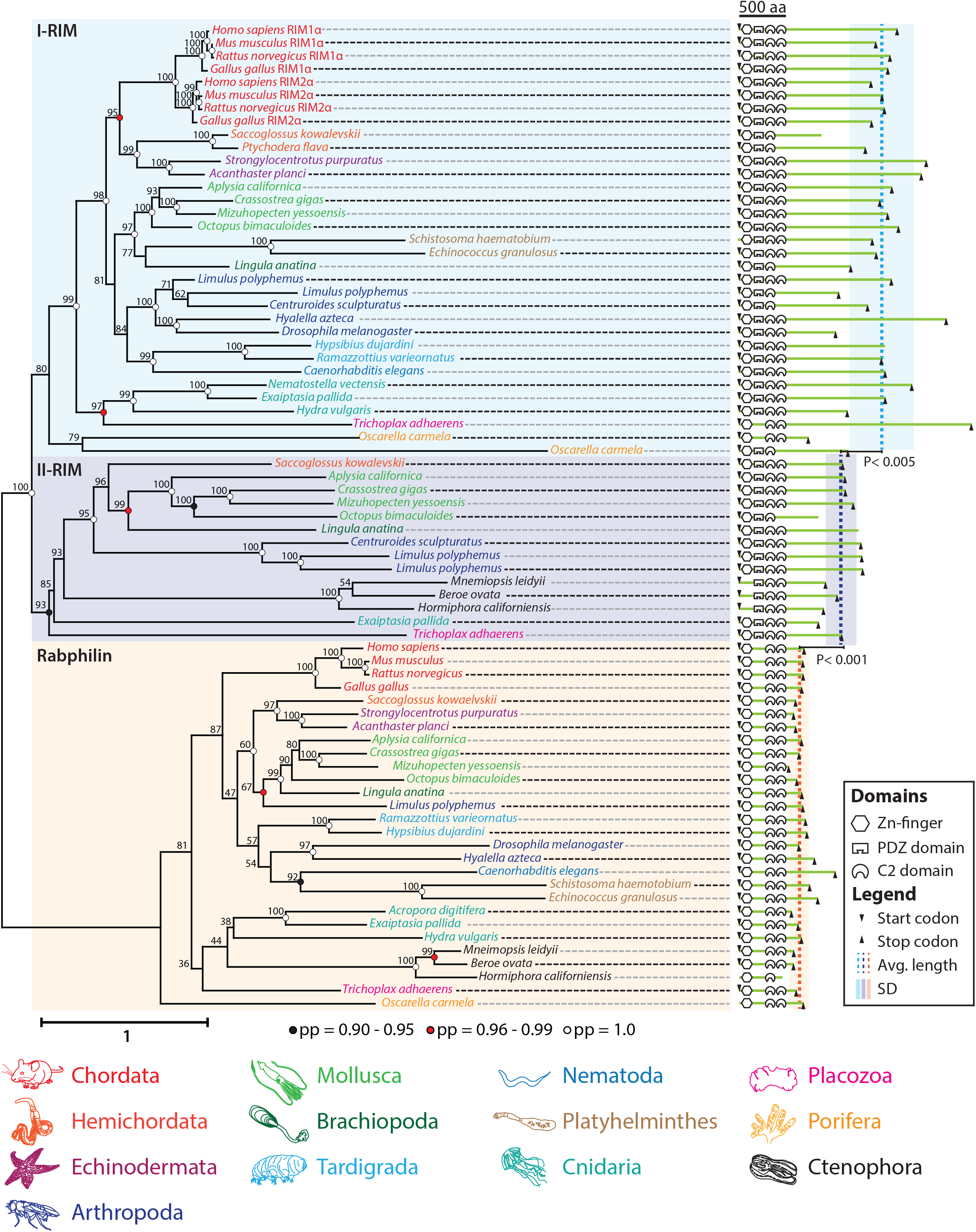
Maximum likelihood phylogenetic analysis reveals a new RIM homologue (II-RIM), present in numerous invertebrate phyla, that forms a separate clade from previously identified RIM proteins (I-RIMs) and rabphilins. Bayesian posterior probabilities indicated on select nodes corroborate maximum likelihood estimation. I-RIM and II-RIM, while significantly different in length (Kruskal Wallis χ^2^ = 57.656, df = 2, p-value = 3.021e-13; p-values for Dunn’s post-hoc test with Benjamini-Hochberg adjustment are shown for pairwise comparisons), possess similar InterProScan-predicted domain architectures, comprised of an N-terminal Zn^2+^-finger domain, a PDZ domain, and two C2 domains. While domain synteny is depicted faithfully, interdomain distances do not accord with the provided scale.

The role of I-RIM in both capacitating Ca^2+^ entry at the pre-synapse through selective recruitment of Ca_V_2s and modulating synaptic vesicle fusion has been documented in rodents (Kaeser, et al. 2011), fruit flies (Graf, et al. 2012; Müller, et al. 2012), and worms (Koushika, et al. 2001; Kushibiki, et al. 2019). The apparent ubiquity of this gene can be contrasted with what appear to be several independent losses of II-RIM in the Bilateria and Cnidaria. We could not identify II-RIM in the fruit fly *Drosophila melanogaster* or the nematode worm *C. elegans*, both of whose genomes and transcriptomes have been subject to considerable annotation. We also failed to identify II-RIM in the tardigrades *Rammazzottius varieornatus* and *Hypsibius dujardini*, which form a sister clade to arthropods. Instead, arthropod chelicerates *Centroides sculpturatus* and *Limulus polyphemus* were found to posses both I and II-RIM genes, but II-RIM was absent in the gene data for the crustacean *Hyalella Azteca*. Thus, it appears as though the ancestral ecdysozoan possessed both RIM genes, and that type II was independently lost in select arthropods (e.g. Mandibulata), nematodes and tardigrades. Further, in deuterostomes the II-RIM gene was not identified in echinoderms *Strongylocentrotus purpuratus* and *Acathaster planci*, but present in gene data for the hemichordate *Saccoglossus kowalevskii*, indicating likely independent losses in Echinodermata and Chordata. We identified I and II-RIM genes in the cnidarian *Exaiptasia pallida*, but only type I in other cnidarians (i.e. *Hydra vulgaris* and *Nematostella vectensis*), suggesting that losses of II-RIM also occurred within Cnidaria, a sister taxa to the bilateria. In contrast to other metazoan phyla, both RIM homologues are highly conserved among molluscs, identified in the gastropods *Aplysia californica* and *Lymnaea stagnalis* (the latter not included in the tree due to fragmentation of assembled mRNA transcripts (Sadamoto, et al. 2012)), the bivalves *Crassostrea gigas* and *Mizuhopecten yessoensis* and the cephalopod *Octopus bimaculoides*. II-RIM was also identified in the brachiopod *Lingula anatina*, suggesting broad conservation among the Lophotrochozoa (Figure 2). Interestingly ctenophores, the most early-diverging animals with synapses, are unique in that they lack I-RIM and possess only II-RIM. Additionally, their II-RIM lacks a Zn^2+^-finger domain broadly conserved in other RIM homologues (Figure 2). This is significant because this particular domain is required for a direct interaction with the protein Munc-13, and hence for RIM to play a role in synaptic vesicle docking and priming (Betz, et al. 2001; Dulubova, et al. 2005; Lu, et al. 2006; Quade, et al. 2019). Nevertheless, ctenophores possess Munc-13 in their genomes (Ryan, et al. 2013; Moroz, et al. 2014). We note that although the absence of identifiable II-RIM genes for various metazoan species could be accounted for by incomplete genome/transcriptome sequencing data, we had little difficulty identifying I-RIM sequences for most of the species included in our analysis.

### I-RIM and II-RIM are differentially expressed in the mollusc snail Lymnaea stagnalis

The obligate retention of RIMs across metazoan genomes (Paps and Holland 2018), coupled with the frequent loss of II-RIM in various clades, suggests that this newly identified gene plays a secondary, redundant role to I-RIMs when both are present in the genome. Nevertheless, our ability to detect II-RIM broadly within Mollusca suggests that they utilize this gene non-redundantly to I-RIM. This compelled us to determine the expression of both genes in different tissues of the pulmonate gastropod *Lymnaea stagnalis* (pond snail). The known anatomy and large neurons of the snail have made it a key model organism for studying the neural correlates of behavior, and much is known about the properties of individual neurons and neural circuits in *Lymnaea* (Syed, et al. 1990; Kemenes and Benjamin 2009). While qPCR experiments of I-RIM revealed relatively high expression in the central nervous system (CNS) of the snail, II-RIM had its highest expression in the prostate, an organ involved in peptidergic secretion and signaling for sexual reproduction (Koene, et al. 2010) (Figure 3A). At lower levels, I-RIM was also detected in the heart, prostate, albumen gland and buccal mass (used for feeding), while II-RIM was detected in the CNS and albumen gland and minimally in buccal mass and heart.

**Figure 3.**
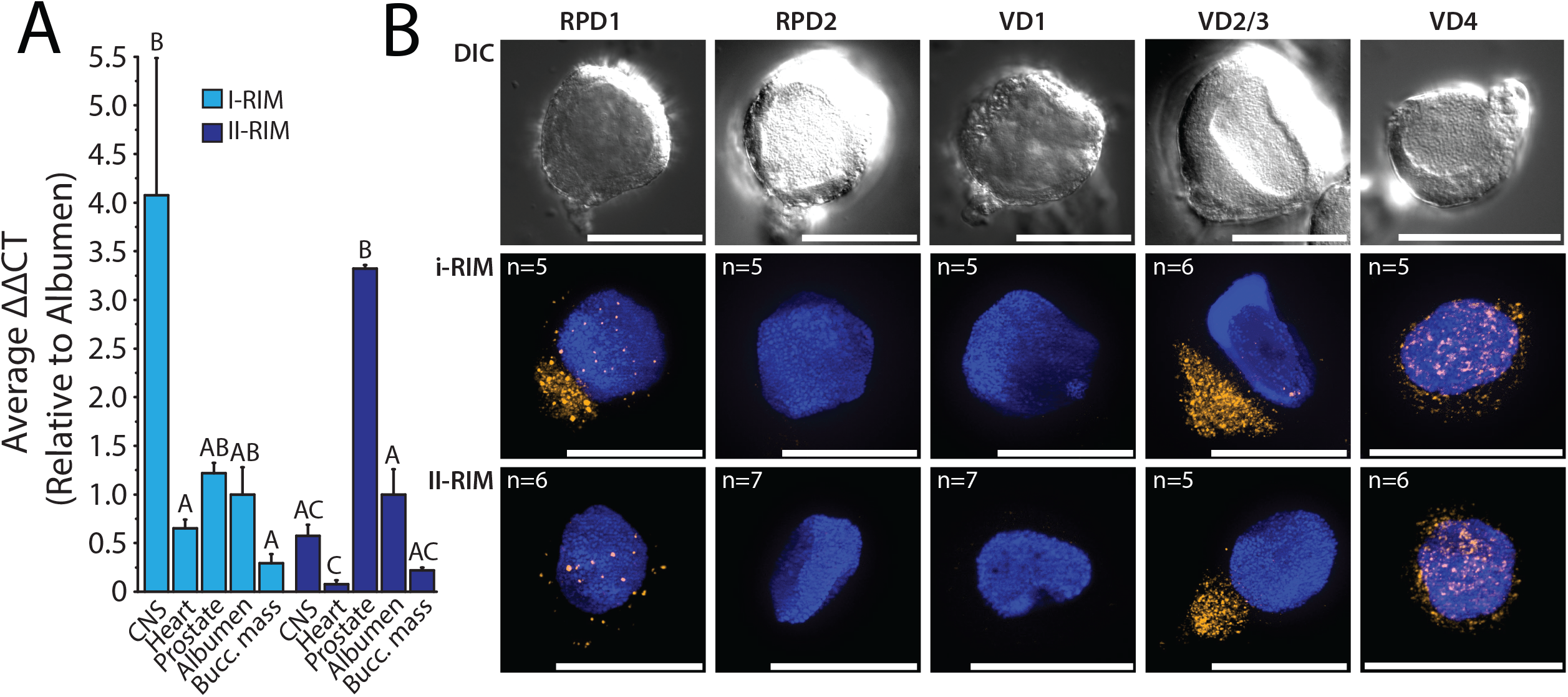
**A)** Quantitative polymerase chain reaction (qPCR) transcript quantification across various tissues of the pond snail *Lymnaea stagnalis* reveals I-RIM expression is most prominent in the CNS, while II-RIM is most abundantly expressed in the prostate gland. **B)** Fluorescent *in situ* hybridization on cultured *Lymnaea* giant neurons from the visceral and right parietal ganglia using gene specific LNA probes indicates co-expression of I-RIM and II-RIM mRNAs in select identified neurons. Differential interference contrast (DIC) and fluorescence channels are shown for right parietal dorsal 1 (RPD1), RPD2, visceral dorsal 1 (VD1), VD2/3 (indistinguishable), and VD4 neurons. Nuclei, labeled with DAPI stain, appear blue. Scale bars denote 50 μm.

Despite the dichotomy seen in *Lymnaea* CNS expression of the two RIMs, the genes exhibited overlapping cellular expression as evidenced by mRNA *in situ* hybridization on isolated and cultured neurons (Figure 3B). Various identifiable neurons chosen to study included: the giant right parietal dorsal 1 (RPD1), RPD2, visceral dorsal 1 (VD1), 2/3 (VD2/3), and 4 (VD4) neurons. Each of these exists as a single cell in each CNS (except for VD2/3, which are analogous neurons and hard to separately identify) and each has been previously subjected to detailed characterization of phenotype, function and morphology (Benjamin and Winlow 1981; Syed, et al. 1990; Beekharry, et al. 2015). While the right parietal ganglion (e.g. A group neurons) are immunoreactive to serotonin (5-HT) (Elekes, et al. 1989), this has not been reported for RPD1, which is thought to express the neuropeptide FMRFamide (Bright, et al. 1993), or RPD2, which harbors a rich peptidome (Jiménez, et al. 2006). Neuropeptide expression has also been reported in the visceral ganglion neurons VD1 (Jimenez et al., 2006) and VD4 (Nesic et al., 1996). VD4 forms reciprocal inhibitory synapses with post-synaptic neuron right pedal dorsal 1 (RPeD1), where interestingly the VD4 neuron is reported to switch transmitters from a FMRFamide-like peptide to acetylcholine, the latter switching the postsynaptic response of RPeD1 to excitatory (Woodin, et al. 2002). RPD1, VD2/3, and VD4 neurons all expressed both RIM homologues, however, RPD1 showed stronger expression of I-RIM than II-RIM, and while both RIM mRNAs clustered in discrete cytoplasmic foci in VD2/3, the expression in VD4 was considerably more diffuse (Figure 3B). Instead, neither RIM homologue was expressed in the electrically-coupled RPD2 and VD1 neurons that innervate organs responsible for cardio-respiratory functions (Bogerd, et al. 1991; Kerkhoven, et al. 1993; Jiménez, et al. 2006; Beekharry, et al. 2015), indicating that these genes are not necessarily ubiquitously/highly expressed in all neurons.

### I-RIM and II-RIM PDZ domains diverged at key loci associated with ligand specificity

As noted, the synaptic interaction between I-RIMs and Ca_V_2 channels is proposed to largely depend on the PDZ domain of RIM binding to D/E-D/E/H-WC-_COOH_ motifs on the extreme C-termini of Ca_V_2 channels (Kaeser, et al. 2011; Graf, et al. 2012; Hirano, et al. 2017). Aiming to infer how I- and II-RIM might compare in mediating PDZ-dependent protein interactions, we aligned representative PDZ domain sequences of both homologues (Figure 4A). The PDZ domains of both RIM types were found to be approximately 100 amino acids long and characterized by predicted stereotypical secondary structures of six beta strands (βA-F), a short alpha helix (αA), and a long alpha helix (αB). Predicted secondary structure was not conserved for the PDZ domain of the sponge *Oscarella carmela*, whose sequence failed to align with other metazoan PDZ domains due to divergence. Globally, the I-RIM PDZ domain shared higher sequence identity (48.6%), compared with PDZ domains of II-RIM (31.5%), particularly towards the N-terminal side upstream of βB. In this region, I-RIMs bear an additional predicted beta strand (β0), consistent with NMR structures (Lu, et al. 2005). Instead, the β0 strand is absent in most other PDZ domains (Lee and Zheng 2010) including II-RIM (Figure 4A). Both I and II-RIM PDZ domains showed high conservation of the ligand carboxylate-accommodating loop of consensus sequence X-φ-G-φ (where φ denotes a hydrophobic amino acid and X any amino acid) (Lee and Zheng 2010), located on the N-terminal side of βB (Figure 4A, black bars). An exception is the *Octopus bimaculoides* I-RIM orthologue, which contains an insert at this key ligand binding locus that was removed from the alignment.

**Figure 4.**
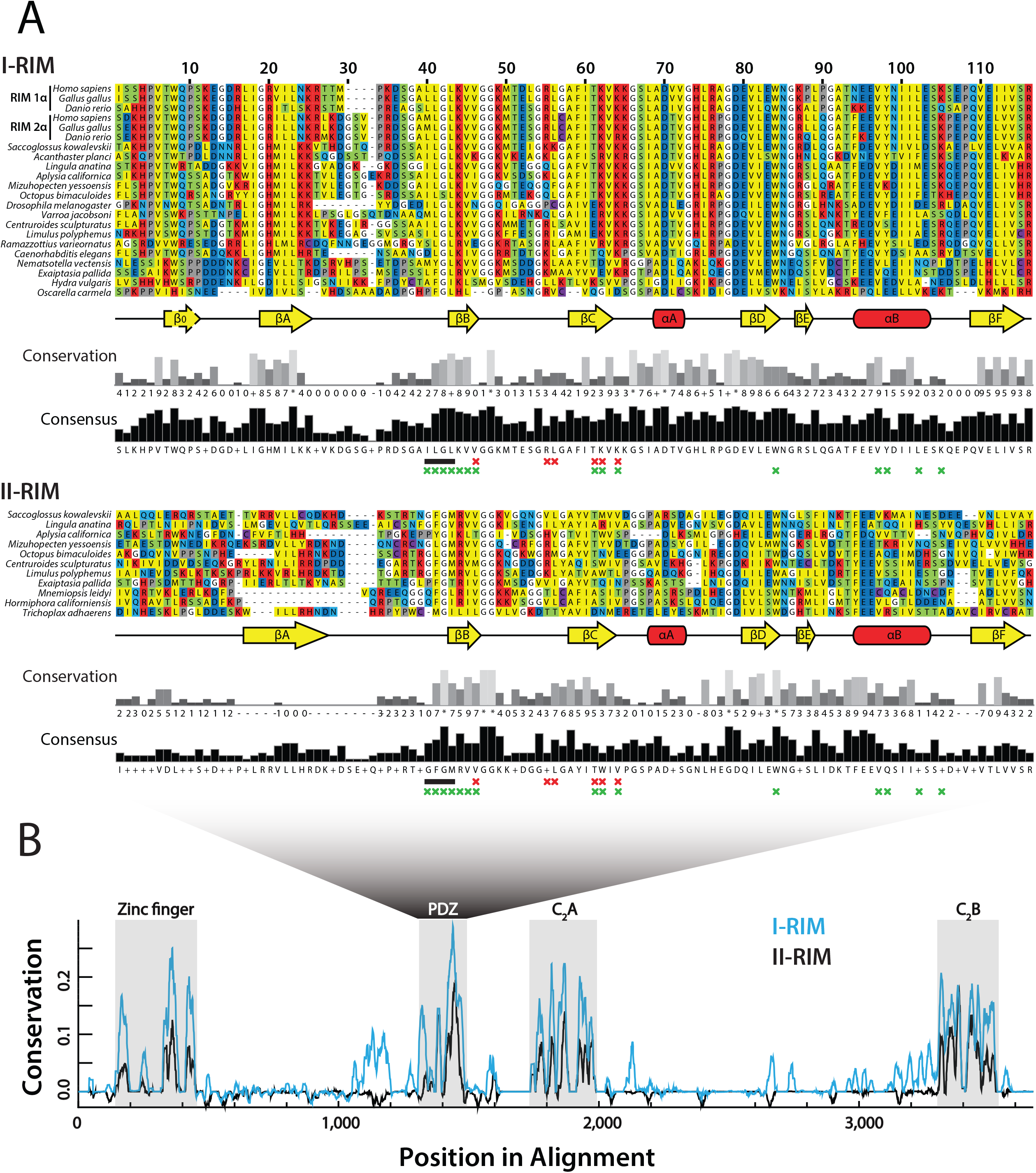
**A)** Multiple sequence alignment of various I-RIM and II-RIM PDZ domains. The secondary structures are defined by six beta strands (βA-F), a short alpha helix (αA), and a long alpha helix (αB). The carboxylate accommodating loop with consensus sequence X-φ-G-φ is indicated by a black bar. Red Xs denote the six residues identified as unequivocal classifiers of distinct PDZ clades (Sakarya, et al. 2009), and green Xs positions of the I-RIM PDZ domain that interact with bound ligands in NMR structures (Lu, et al. 2005). **B)** Graph depicting sequence conservation/entropy of aligned RIM proteins delineates conserved Zn^2+^-finger, PDZ, C_2_A and C_2_B domains, and low homology within inter-domain protein regions.

Early studies on the ligand specificities of different PDZ domains delineated three classes based on C-terminal ligand recognition sequences, the most common being class I PDZ domains (ligand motifs of X-S/T-X-φ-_COOH_, where X denotes any amino acid and φ a hydrophobe), as well as class II and class III domains (respective ligand motifs of X-φ-X-φ-_COOH_ and X-E/D-X-φ_COOH_) (Songyang, et al. 1997; Stricker, et al. 1997; Nourry, et al. 2003). More recently, a comprehensive analysis of PDZ domain ligand specificities through peptide-phage display of over 330 PDZ domains in human and nematode worm expanded the specificity classes of known PDZ domains to 16 distinct ligand classes (Tonikian, et al. 2008). Apparent is that the ligand specificity of type I RIM does not neatly fit within the three or sixteen type classification systems, indicative of a unique specialized selectivity for the Ca_V_2 channel ligand of D/E-D/E/H-WC-_COOH_. By leveraging evolutionary divergence of distinct PDZ domains within the proteomes of numerous animals and closely-related eukaryotes, a study identified six aligned amino acid positions that share high general entropy but low within-clade entropy, representing unequivocal classifiers of the clade to which a given PDZ domain belongs (Sakarya, et al. 2009). Based on NMR structures of I-RIM PDZ domains interacting with C-terminal ligands of ELKS1b and Ca_V_2.1, four of these six amino acids (i.e. βB4, βC4, βC5, and βC-αA-1, where βB4 denotes the fourth residue of the second beta strand) contact the entropic p-1 and p-3 residues of ligands (where p0 denotes the distal-most C-terminal residue), and are involved in ligand selectivity (Lu, et al. 2005; Sakarya, et al. 2009; Kaeser, et al. 2011). In an effort to parse out potential differences in the protein-binding capabilities of type I and II RIMs, we labeled these key residues in our alignment (Figure 4A). Interestingly, βC5, and βC-αA-1 differ between the two RIM homologues: while I-RIM has a highly conserved basic region defined by K46 and K48 (i.e. T<u>K</u>V<u>K</u> motif), II-RIM features hydrophobic W46 and V48 (T<u>W</u>I<u>V</u>). These particular amino acids have been shown to contribute to the binding pocket of I-RIM1, which exhibit shifts in heteronuclear single quantum coherence spectra upon binding to Ca_V_2.1 C-terminal peptides (Kaeser, et al. 2011). Another key site associated with ligand specificity is the consensus amino acid in position αB1 of the PDZ domain, which interacts with p-2 ligand residues (Hung and Sheng 2002). For most RIM PDZ domains examined in this study, this position was occupied by a phenylalanine (F), albeit with considerable variation (Figure 4A). αB1 amino acids that form hydrogen bonds (i.e. Y, N, Q, K, R) preferentially bind hydroxy group-containing serine or threonine p-2 residues of class I ligands (X-<u>S/T</u>-X-φ-_COOH_), while hydrophobic amino acids select for hydrophobic p-2 residues of class II ligands (X-φ -X-φ-_COOH_), and tyrosines interact with acidic p-2 residues of class III ligands (X-[<u>D/E</u>]-X-φ_-COOH_). However, this model is inconsistent with the ligand-binding properties of I-RIM PDZ domains, since both co-immunoprecipitation and NMR experiments have demonstrated that I-RIM1 and I-RIM2 PDZ domains interact with the DDWC-_COOH_ ligand on the Ca_V_2.1 channel C-terminus (Kaeser, et al. 2011; Hirano, et al. 2017), despite I-RIM1 bearing an asparagine in position αB1 consistent with class I ligands, and I-RIM2 bearing a phenylalanine consistent with class II ligands. Thus, this locus might play a minimal role in RIM ligand specificity. In summary, although both type I and II RIM proteins bear canonical PDZ secondary structures, the two homologues have differences at key loci suggesting differences in ligand specificity.

We also compared the Zn^2+^-finger and C_2_ domain sequences conserved between I-RIM and II-RIM (Figure 4B), plus rabphilin (Figure S2A). With few exceptions, the N-termini contained a predicted Zn^2+^-finger domain and α helical structures (αA) involved in Munc-13 and Rab3 binding, respectively (Wang, et al. 1997; Betz, et al. 2001; Wang, et al. 2001; Fukuda 2003; Dulubova, et al. 2005; Lu, et al. 2006; Quade, et al. 2019), two short β strands (βA and βB), and a second α helix (αB) (Figure S2A, Supplementary file 1). Notably, although the molecular determinants for RIM/rabphilin interactions with Munc-13 and Rab3 are considered separate (Ostermeier and Brunger 1999), mutations in the Zn^2+^-finger domain nevertheless disrupt interactions with Rab3 (McKiernan, et al. 1996; Stahl, et al. 1996), indicative of structural interdependence between these two regions. Of the eight Zn^2+^-finger cysteine (C) residues required for Zn^2+^ and Rab3 binding (Stahl, et al. 1996), seven were very highly conserved across most orthologues. Furthermore, the αB helix SGAWFF motif, identified as a Rab complementarity-determining region that confers specificity to select Rab proteins (Ostermeier and Brunger 1999), had deep conservation across rabphilin sequences but was less conserved in I-RIM and II-RIM. The C_2_A and C_2_B domains of all three proteins were characterized by 8 predicted β strands (βA to βH), common to type I C_2_ domains (i.e. synaptotagmin family C_2_ domains), which form an eight-stranded anti-parallel beta sandwich secondary structure (Biadene, et al. 2006). As noted, the RIM C_2_B domain, present in all four vertebrate I-RIM paralogues (RIM1-4), potentiates Ca^2+^ influx through Ca_V_2 channels via an interaction with the Ca_V_β subunit, which attenuates voltage-dependent inactivation to prolong pre-synaptic Ca^2+^ influx (Kiyonaka, et al. 2007; Uriu, et al. 2010; Kaeser, et al. 2012). Although the mechanisms for this protein-protein interaction have not yet been elucidated, it is likely not dependent on Ca^2+^, given that I-RIM C_2_ domains are degenerate in their Ca^2+^-binding capacity, lacking key residues including five aspartates that comprise Ca^2+^ binding sites in related proteins rabphilin and synaptotagmin (Wang, et al. 1997; Ubach, et al. 1998; Coudevylle, et al. 2008). To assess whether the II-RIM C_2_ domains might bind Ca^2+^, we aligned I-RIM, II-RIM and rabphilin C_2_A and C_2_B domains, identifying the five aspartate (D) residues that mediate Ca^2+^ binding (Figure S2B). A high conservation of aspartate in all three proteins was seen exclusively at p110, suggesting that II-RIMs, like I-RIMs, have degenerate C_2_ domains.

### The phylogeny of Ca_V_ channels informs on conserved PDZ and SH3 domain ligand motifs

The reported conservation of D/E-D/E/H-WC-_COOH_ PDZ ligand motifs on the distal C-termini of Ca_V_2 channels from vertebrates (Kaeser, et al. 2011; Gardezi, et al. 2013), fruit flies (Graf, et al. 2012) and molluscs (Spafford, et al. 2003) is notable given that at the phylum-level protein alignments of orthologous Ca_V_ channel intracellular linkers and N- and C-termini tend to show poor sequence homology (Spafford, et al. 2003; Senatore and Spafford 2010; Tyson and Snutch 2013). In addition to binding I-RIMs, the Ca_V_2 PDZ ligand motif also mediates interactions with a PDZ domain of the pre-synaptic scaffolding protein Mint, documented in rodents (Maximov, et al. 1999), chick (Gardezi, et al. 2013) and the gastropod mollusc *Lymnaea stagnalis* (Spafford, et al. 2003). Thus, it seems likely that interactions between Ca_V_2 channels and I-RIM/Mint-1 were present in the last common ancestor of the bilaterians. Nevertheless, a comprehensive analysis of Ca_V_ channel C-terminal sequences within the Metazoa, to explore conservation of C-terminal PDZ ligand motifs, has not been reported. Furthermore, recently sequenced genomes and transcriptomes permit re-exploration of the Ca_V_ channel phylogeny (Moran and Zakon 2014; Senatore, et al. 2016). Hence, using sequences compiled from genomic and transcriptomic databases, we constructed a comprehensive maximum likelihood protein phylogeny of various Ca_V_ α subunits, and aligned their 10 distal-most C-terminal amino acid sequences which would bear putative PDZ ligand motifs (Figure 5A). Rooting the tree with fungal CCH1 Ca_V_ channel homologues (*Saccharomyces cerevisiae* and *Schistosoma pombe*) revealed three distinct clades of metazoan Ca_V_ channels, with LVA Ca_V_3 channels forming a sister clade with the HVA Ca_V_1 and Ca_V_2 channels. Similar to CCH1, Ca_V_ channel homologues from the ciliate *Paramecium tetraurelia*, involved in regulating ciliary beating (Lodh, et al. 2016), form a sister clade with metazoan Ca_V_ channels, while those from the green algae *Chlamydomonas* sp. and *Gonium pectoral* (also involved in regulating ciliary beating (Fujiu, et al. 2009)), formed a sister clade with Ca_V_3 type channels. Our phylogenetic tree is consistent with previous reports that HVA and LVA channels existed in the last common ancestor of animals and choanoflagellates, where the choanoflagellate species *Salpingoecca rosetta* possesses a *bona fide* Ca_V_3 channel homologue, as well a Ca_V_1/2 channel posited to be ancestral to Ca_V_1 and Ca_V_2 channels (Moran and Zakon 2014). Also consistent with previous reports, sponges *Amphimedon queensladica, Haliclona ambioensis* and *Haliclona tubifera* possess single Ca_V_1/2 channel homologues, and lack Ca_V_3 channels, attributed to gene loss. It has been proposed that Ca_V_1 and Ca_V_2 channels emerged via gene duplication from a Ca_V_1/2-like channel, perhaps after sponges diverged from other animals (Moran, et al. 2015). However, here we identify a Ca_V_1 channel homologue in the gene sequences of the sponge *Oscarella carmela*, suggesting instead that this event occurred prior to the divergence of sponges, and in turn, that most lineages of sponges lost Ca_V_1 and Ca_V_2 channels (Figure 5). Indeed, such a scenario would explain the presence of Ca_V_2 channels in the gene sequences of ctenophore species *Mnemiopsis leidyi, Beore ovata* and *Hormiphora californiensis*, which based on the leading species phylogeny, are the most early-diverging group of animals (Ryan, et al. 2013; Moroz, et al. 2014; Whelan, et al. 2017). As such, Ca_V_1/2, Ca_V_1 and Ca_V_2 channels, plus Ca_V_3 channels, might have existed in the common ancestor of all animals, and these were differentially lost such that ctenophores retained only Ca_V_2 channels, and sponges Ca_V_1 or Ca_V_1/2 channels. Instead, the placozoan *Trichoplax adhaerens*, which forms a sister clade with cnidarians and bilaterians, is the most early-diverging animal to possess all three types of metazoan Ca_V_ channels (i.e. Ca_V_1-Ca_V_3), and along with cnidarians and bilaterians, they lack Ca_V_1/2 channels (Figure 5). Also evident are the two rounds of Ca_V_ gene duplication in the stem lineage of vertebrates, resulting in ten vertebrate Ca_V_ channels (i.e. Ca_V_1.1-Ca_V_1.4, Ca_V_2.1-Ca_V_2.3 and Ca_V_3.1-Ca_V_3.3), and independent duplications in Cnidaria resulting in six Ca_V_ channel homologues (Ca_V_1, Ca_V_2a-Ca_V_2c and Ca_V_3a-Ca_V_3b) (Figure 5A) (Jegla, et al. 2009; Moran and Zakon 2014; Moran, et al. 2015).

**Figure 5.**
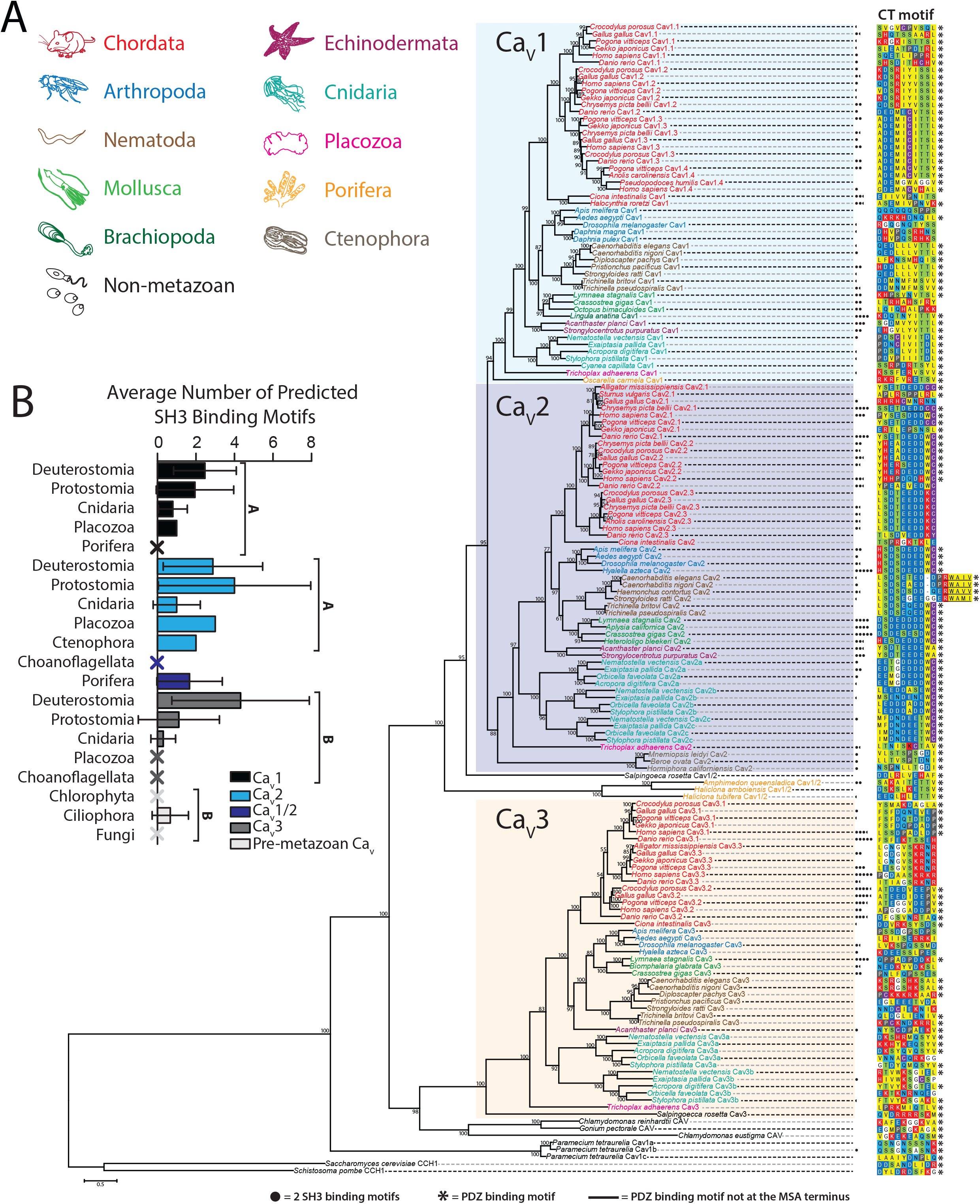
**A)** Maximum likelihood phylogenetic tree of the alpha subunit of metazoan and pre-metazoan voltage-gated calcium channel (Ca_V_) homologues. Bootstrap values for 1,000 ultrafast replicates are indicated on corresponding nodes. The distal 10 amino acids that contain putative PDZ domain ligand motifs are aligned for all sequences. PDZPepInt predictions of PDZ domain binding are annotated by black asterisks. SH3PepInt predictions of C terminal SH3 binding motifs are denoted with filled black circles. **B)** Quantitation of average number of predicted SH3 binding motifs per Ca_V_ paralogue and clade. Ca_V_1 and Ca_V_2 have on average a significantly higher number of SH3 motifs (one way ANOVA and Dun’s post-hoc test with the Benjamini-Hochberg adjustment). Error bars denote standard deviation, and Xs denote zero predictions.

Pursuant to our characterization of the PDZ domains of I-RIM and II-RIM, we examined the sequences of putative Ca_V_ C-terminal PDZ ligands across all homologues (Figure 5). Four amino acids at the extreme C-termini typically participate in the β-strand complementation that mediates PDZ ligand-domain protein interactions (Hung and Sheng 2002), however, at least seven residues upstream of the carboxylate group are known to strengthen this interaction via intermolecular bonds and attractions (Tonikian, et al. 2008; Ernst, et al. 2014). In contrast to the hypervariable sequence that typifies the medial and distal thirds of Ca_V_1 and Ca_V_2 C-termini (Figure 6), our alignment evidences high conservation of the ten most distal amino acids within respective paralogues (Figure 5A). In large part, Ca_V_1 orthologues possess class I PDZ ligands, and Ca_V_2 orthologues possess non-canonical ligands. The Ca_V_1-Ca_V_3 channels from *Trichoplax*, and Ca_V_s from the sponges, all possess class I PDZ ligands, the latter bearing ligand motifs of E-T-S/T-V-_COOH_ identified as a consensus PDZ ligand sequence for Discs large homolog (DLG) scaffolding proteins of both humans and nematode worms (Tonikian, et al. 2008). Importantly, we found motifs similar to D/E-D/E/H-WC-_COOH_ to be conserved in Ca_V_2 channels throughout Bilateria and Cnidaria (Figure 5), but absent in orthologues from *Trichoplax* and ctenophores. Ca_V_3 channels largely lack distal C-terminal conservation across phyla (Figures 5A and 6A). On a more granular level, we observed apparent sequence divergence from a class I PDZ ligand among Ca_V_1 channels in arthropods and Ca_V_1.1 channels in vertebrates, and a conserved hydrophobic insert disrupting the D/E-D/E/H-WC-_COOH_ like motif in nematodes of the clade Rhabditomorpha (*Caenorhabditis, Haemonchus* and *Stronglyloides*), but not Trichinellida. Furthermore, we note the apparent absence of D/E-D/E/H-WC-_COOH_ like motifs in the Ca_V_2 channel of early-diverging chordate *Ciona intestinalis* and avian P/Q-type Ca_V_2.1 channels, the latter concomitant with the reported loss of C-terminal exon 47 (human equivalent) in the gene from *Gallus gallus* (Snidal, et al. 2018). To corroborate our sequence data, we also predicted C-terminal PDZ ligands with the cluster based prediction tool PDZPepInt (Kundu, et al. 2014), which references a set of 226 PDZ domains from humans, mouse, fly and nematode worm, finding that most metazoan and all pre-metazoan Ca_V_s bear predicted PDZ ligands (Figure 5A).

**Figure 6.**
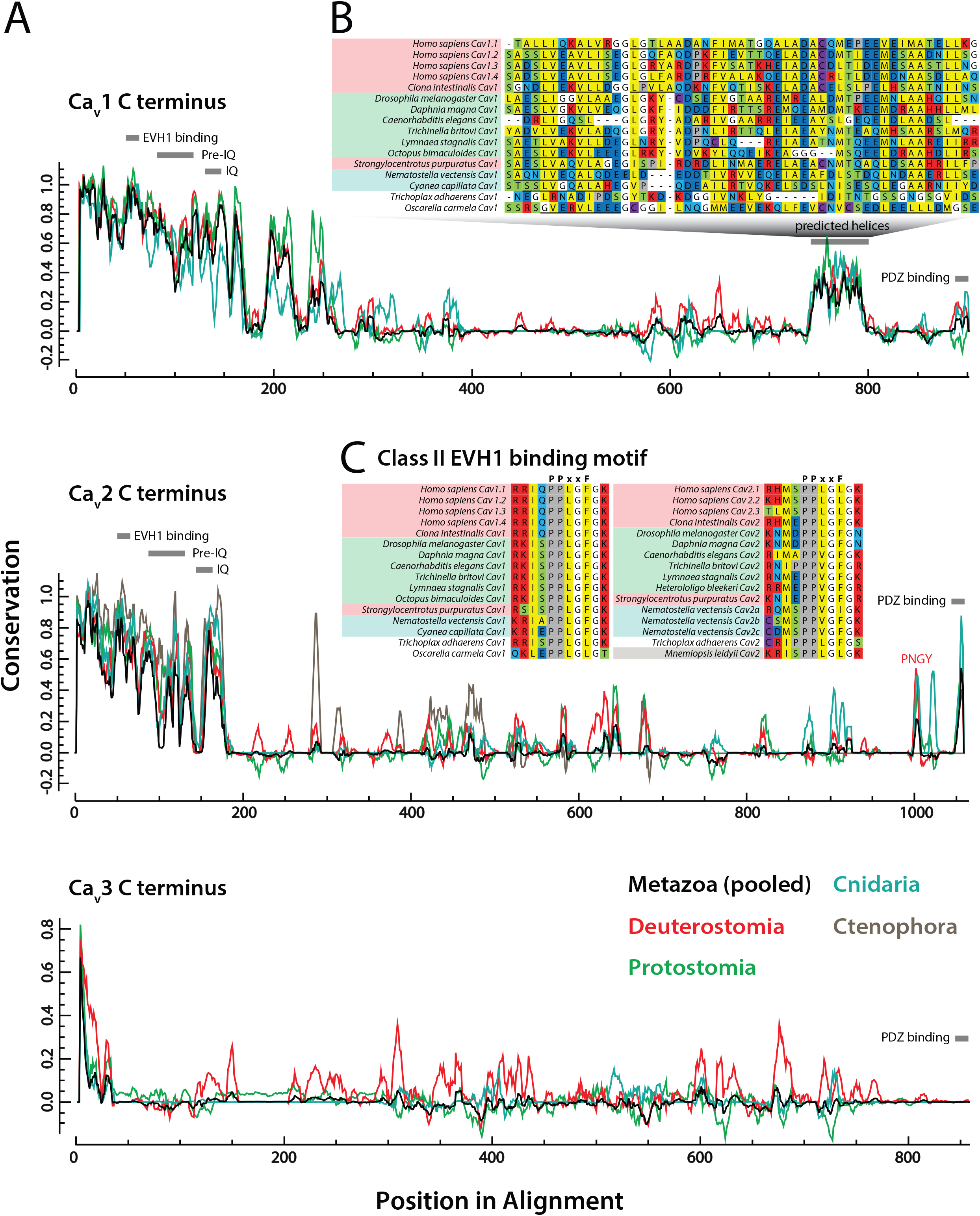
**A)** Graph depicting sequence conservation/entropy of aligned CaV channel C-termini reveals distinct pockets of sequence conservation within the largely disordered protein sequences. Separate traces showing conservation among individual clades were made by pooling sequences constituting a given clade from the original multiple sequence alignment and normalizing to one scale. Previously reported short linear motifs (SLiMs) are indicated with colored bars to denote the degree of conservation among orthologous sequences. **B)** A highly conserved region in the Ca_V_1 distal C-terminus corresponds to a predicted helical structure conserved even in the identified homologue from the sponge *Oscarella carmela*, absent in Ca_V_2, Ca_V_3 and Ca_V_1/2 channels (PSIPRED-predicted α helical secondary structures are denoted with black underlines, and β strands with orange underlines). The Ca_V_1 channel from *Trichoplax adhaerens* uniquely lacked a predicted helix in this region. **C)** Alignment of the identified class II EVH1 domain-binding motif of consensus sequence PPXXF reveals deep conservation of this motif in both Ca_V_1 and Ca_V_2 channels.

To expand our analysis of putative protein-interaction motifs on Ca_V_ channel C-termini, we also predicted Src homology 3 (SH3) domain ligands, present in Ca_V_2 and I-RIM homologues, where the scaffolding protein RIM-BP binds to form a tripartite Ca_V_2/RIM/RIM-BP complex at the synapse active zone (Südhof 2012) (Figure 1A). RIM-BP SH3 domains have also been shown to interact with proline-rich regions on the C-termini of Ca_V_1 channels (Hibino, et al. 2002), however such interactions have not been reported for Ca_V_3 channels, nor has the presence of putative SH3 ligand motifs been studied systematically in Ca_V_ channels from divergent animal phyla. This paucity of data likely stems both from the sequence hypervariability that characterizes the C-termini of Ca_V_s (Figure 6), and the fact that SH3 domains recognize numerous non-canonical binding motifs (Teyra, et al. 2017) making predictions difficult. Nonetheless, we used three independent methods to predict the number of SH3 binding motifs within the C-termini of all examined Ca_V_ homologues (Figures 5 and S3), finding that Ca_V_1 and Ca_V_2 channels contain on average significantly more SH3 motifs than either Ca_V_3 or pre-metazoan Ca_V_ channels (Figure 5B). Intra-channel (within paralogue) inter-clade comparisons revealed a significantly greater number of predicted SH3 motifs among Ca_V_3 channels from deuterostomes compared to protostomes (p=0.0010) or cnidarians (p=0.0013) (Kruskal-Wallis test and Dunn’s post-hoc test with Benjamini-Hochberg adjustment), while inter-clade differences were non-significant for Ca_V_1 and Ca_V_2 paralogues. Interestingly however, we note an enrichment in SH3 motifs among Ca_V_2 channels from molluscs (Figure 5A). Lastly, intra-clade inter-channel comparisons revealed a significantly greater number of SH3 motifs among Ca_V_2 vs. Ca_V_3 channels from protostomes (p=0.00713), but no significant differences between Ca_V_ channel paralogues from deuterostomes or cnidarians.

### Intrinsically disordered Ca_V_ channel C-termini and linker regions are hubs for short linear motifs

The noted sequence entropy within Ca_V_ channel cytoplasmic regions reflects lower evolutionary constraints, perhaps facilitating the emergence of novel motifs with novel interactomic functions in distinct channel clades. Accordingly, channel termini and linkers are important regions for differential Ca_V_ channel modulation by regulatory proteins (Tyson and Snutch 2013). To systematically characterize the cytoplasmic regions of the Ca_V_ channels included in our phylogenetic tree (Figure 5), we first performed a quantitation of the protein sequence length of the N- and C-termini, plus the I-II, II-III and III-IV cytoplasmic linkers (Figure S4). While no significant differences were noted among N-termini lengths between calcium channel paralogs when all clades were pooled (Kruskal-Wallis χ^2^ = 5.9176, df = 3, p-value = 0.1157), both Ca_V_1 and Ca_V_2 had significantly longer C-termini as compared with Ca_V_3 and pre-metazoan channels (Kruskal-Wallis χ^2^ = 45.272, df = 3, p-value = 8.1E-10) (Figure S4). Notably, linkers among Ca_V_3 channels were both significantly longer and more variable than those of Ca_V_1 and Ca_V_2 channels, and particularly so for the I-II and II-III linkers (I-II: Kruskal-Wallis χ^2^ = 113.82, df = 3, p-value < 2.2E-16; II-III: Kruskal-Wallis χ^2^ = 60.884, df = 3, p-value = 3.806E-13; III-IV: Kruskal-Wallis χ^2^ = 140.14, df = 3, p-value < 2.2E-16). Intrachannel (within paralog)-interclade comparisons of termini and linker lengths were also performed. Despite the generally shorter II-III linker in Ca_V_2 channels as compared to Ca_V_3, the deuterostome Ca_V_2 channels have significantly longer linkers relative to those found in protostomes and cnidarians (Kruskal-Wallis χ^2^ = 40.2965, df = 2, p-value = 0). This is consistent with the observation that the II-III linker SYNPRINT motif, involved in interactions between Ca_V_2 channels and exocytotic SNARE proteins, is a feature unique to vertebrate, and perhaps all deuterostome, channels (Spafford, et al. 2003). Lastly, while a significant expansion in length was noted for bilaterian Ca_V_2 C-termini as compared with cnidarian orthologues (Kruskal-Wallis χ^2^ = 22.822, df = 2, p-value = 1.107E-05), deuterostome C-termini were significantly longer than those of protostomes for Ca_V_1 and Ca_V_3 (intra Ca_V_1: Wilcoxon statistic W = 367.5, p-value = 0.03867; intra Ca_V_3 Kruskal-Wallis χ^2^ = 14.265, df = 2, p-value = 0.0007986). Altogether the variability observed in these disordered structures likely reflects differential protein interactions and modulatory capacities for the different Ca_V_ channel types within and across different clades (Tyson and Snutch 2013).

Next, we sought to determine whether we could identify novel short linear motifs (SLiMs), or concomitantly, evidence the lack of known motifs in these disordered protein regions by leveraging sequence conservation analysis (Spafford, et al. 2003). A running window of the sequence conservation of representative bilaterian and cnidarian sequences was generated for all Ca_V_ paralogues, then visualized by pooling respective sequences from the original multiple sequence alignment to allow for identification of clade-specific SLiMs (Figure 6A). Generally, Ca_V_3 channels were found to be more variable than either Ca_V_1 or Ca_V_2 channels, particularly in the proximal third of the C-terminus. Further, we identified an island of conservation amid highly entropic sequences in the distal third of Ca_V_1, found to possess helical character upon PSIPRED secondary structure prediction (Figure 6A and B). This locus has been characterized as a c<u>A</u>MP-dependent protein <u>k</u>inase-<u>a</u>nchoring <u>p</u>rotein 15 (AKAP15) binding domain in Ca_V_1.2 channels, required to effect β-adrenergic receptor mediated increase in calcium current (Hulme, et al. 2003). Additionally, proteolytic cleavage of the distal C-terminus bearing this motif produces an auto-inhibitory peptide that binds a proximal region of the Ca_V_ channel C-terminus to inhibit its activation (Hulme, et al. 2006), and can translocate to the nucleus to act as a transcription factor (Gomez-Ospina, et al. 2006). Here, we show that this helical AKAP15 binding element is conserved across Bilateria and Cnidaria, and exists even in the identified Ca_V_1 channel homologue from the sponge *Oscarella carmela* (Figure 6B), structurally distinguishing it from Ca_V_1/2 channels from other sponge species (Figures 5A and 6A).

Next, we used the motif elicitation tools SIB MyHits (exhaustive database search) (Pagni, et al. 2007) and Multiple Em for Motif Elicitation (MEME) (Bailey, et al. 2009) to identify SLiMs hidden within poorly conserved regions of the Ca_V_ C-termini. While MyHits returned questionable or weak matches, MEME, combined with manual analysis of proline-rich motifs in our Ca_V_ multiple sequence alignment, identified a highly conserved type II *Drosophila* <u>e</u>nabled/<u>v</u>asodilator-stimulated phosphoprotein <u>h</u>omology <u>1</u> (EVH1) domain binding motif within the proximal C-termini of Ca_V_1 channels (consensus of P-P-X-X-F), and Ca_V_2 channels (P-P-X-X-φ; Figure 6C). Like the SH3 domain, EVH1 domains bind proline-rich regions on target proteins with low affinity, and feature prominently in signaling networks and at synapses (Ball, et al. 2002). Notable among EVH1 domain-containing proteins is the post-synaptic scaffolding protein Homer (and related proteins). Homers have been reported to regulate excitation-contraction coupling through a physical interaction with Ca_V_1.2 channels and the ryanodine receptor RyR2 (Huang, et al. 2007), and to mediate the flux of extracellular Ca^2+^ between the plasma membrane and the endoplasmic reticulum though interactions with Ca_V_1.2 and STIM1 (Dionisio, et al. 2015). Notably, it has not yet been determined whether these specific interactions involve the conserved EVH1 binding site identified here for Ca_V_1 channels. Nevertheless, the interaction between Homer and Ca_V_1.2 requires a functional Homer EVH1 domain, since point mutations that disrupt its binding capacity disrupt binding with Ca_V_1.2 channels (Huang, et al. 2007). Much less is known about the potential interaction between Homer and Ca_V_2 channels. In one study, G-protein inhibition of Ca_V_2.2 channels by metabotropic glutamate receptors (mGluRs) was found to be disrupted by select Homer variants (Kammermeier, et al. 2000), suggesting that Homer is either directly modulating mGluR function (Brakeman, et al. 1997), or alternatively, affecting G-protein binding to the Ca_V_ channel. Indeed, here we show that while all human Ca_V_2 subtypes possess a type II EVH1-like motif of P-P-X-X-L, the canonical P-P-X-X-F motif is present in Ca_V_2 channels as early-diverging as *Trichoplax* (Figure 6C), which suggests that EVH1 domain-bearing proteins like Homer may regulate Ca_V_2 channels broadly in the Metazoa.

## Discussion

### On the phylogeny of I-RIM

The multifaceted nature of RIM is still being unraveled more than twenty years after its initial characterization as an effector of Rab3, a neuronal GTP-binding protein that regulates synaptic vesicle fusion (Wang, et al. 1997). RIM protein isoforms that bear N-terminal Zn^2+^ finger domains and flanking alpha helical structures, which bind Munc-13 and Rab3, play important roles in regulating synaptic vesicle priming and docking (Gracheva, et al. 2008; Deng, et al. 2011). Separately, C-terminal regions of RIM interact with Ca_V_2 channels (via the PDZ domain), the Ca_V_β subunit (via the C_2_B domain) and RIM-BP (via a proline-rich motif between C_2_A and C_2_B) (Kiyonaka, et al. 2007; Uriu, et al. 2010; Kaeser, et al. 2012; Südhof 2012), allowing the protein to functionally link exocytosis-ready vesicles with the excitation-dependent Ca^2+^ signaling. Indeed, the broad conservation of this functionality across bilateria (Kaeser, et al. 2011; Graf, et al. 2012; Kushibiki, et al. 2019), points to an early evolutionary adaptation of I-RIM for regulating fast, synchronous synaptic exocytosis that requires nanometer proximity between Ca_V_2 channels and exocytotic vesicles (Eggermann, et al. 2012; Wang and Augustine 2015; Stanley 2016). That the N-terminal and C-terminal interactions/functions of I-RIM might be considered separate is suggested by a recent study on the mechanisms for exocytosis of large dense-core vesicles in isolated neurons from conditional RIM1/2 knockout mice. Here, genetic reintroduction of I-RIM1 variants bearing disrupted N-terminal sequences failed to rescue dense-core vesicle exocytosis, while variants lacking the PDZ domain were successful at rescuing exocytosis (Persoon, et al. 2019). Hence, in these neurons, the N-terminus-associated functions of RIM (priming and docking of vesicles) appears essential, while its role in Ca_V_ channel localization does not.

The recent characterization of RIM as one of only 25 genes that are unique to animals and that have broadly resisted genetic loss (Paps and Holland 2018), including in animals that lack nervous systems and synapses, points to a general functionality for this gene that is perhaps distinct from its role in synchronous neuronal Ca^2+^/excitation-dependent exocytosis. For example, I-RIM is present in the gene data for both placozoans (*Trichoplax adhaerens*) and poriferans (*Oscarella carmela*) (Figure 2), both of which lack synapses. Homologues from both of these early-diverging animals lack PDZ domains (Figure 2), likely lost from a common ancestor, and as a result, their putative capacity to interact with Ca_V_2 channels (although *Oscarella* has a second I-RIM homologue with a PDZ domain). Poriferans also lost the majority of genes required for fast electrical neural signaling, including voltage-gated sodium and potassium channels (Moran, et al. 2015), and thus lack the capacity for canonical electrical signaling in the form of action potentials, and by extension, excitation-secretion coupling. Perhaps, the bimodal functionality of I-RIM is phylogenetically conserved, where its regulation of vesicle-cell membrane interactions is widely conserved, while its roles in nanodomain coupling of Ca_V_2 channels is restricted to select neurons in animals that utilize fast, synchronous synaptic transmission. Interestingly, we show here through *in situ* hybridization that only a subset of cultured neurons from the CNS of the mollusc snail *Lymnaea stagnalis* express I-RIM (Figure 4B), implying that this gene is not essential for synaptic exocytosis in all neuron types, or instead, that phylogenetically distinct proteins can carry out redundant functions in neurons that do not express I-RIM. Nonetheless, the importance of I-RIM is underscored by genetic disruption studies in vertebrates and invertebrates, where for example, double knockout of RIM1 and RIM2 in mouse is postembryonic lethal, attributed to disrupted neurotransmitter release (Schoch, et al. 2006).

Notable is that *Trichoplax* is the most early-diverging animal to possess I-RIM plus all three types of Ca_V_ channels found in cnidarians and bilaterians (Ca_V_1-Ca_V_3 channels). However, *Trichoplax* Ca_V_2 lacks a C-terminal D/E-D/E/H-WC-_COOH_ like ligand motif, and as noted above, its I-RIM gene lacks a PDZ domain. All three cnidarian Ca_V_2 channel subtypes possess D/E-D/E-WC-like motifs (Ca_V_2c bears an atypical ETWC motif), and I-RIM is broadly conserved in these animals (Figure 2). Thus, based on current models, cnidarians might have the capacity for a I-RIM/Ca_V_2 pre-synaptic interaction, akin to bilaterians. Indeed, Ca_V_ channels are known to drive synaptic transmission in cnidarians (Bullock 1943; Kerfoot, et al. 1985). However, whether they similarly exhibit nanodomain and microdomain synapses, distinguished by the proximity between Ca_V_ channels and synaptic vesicle Ca^2+^ sensors, is not known (Senatore, et al. 2016).

### Identification of a novel clade of RIM homologues

Here, we report the identification of a previously unknown clade of metazoan RIM genes (II-RIMs), with similar domain architectures as I-RIMs, but generally shorter in length (Figure 2), and bearing sequence differences at key loci including the PDZ domain ligand interface (Figure 4). We acknowledge that our analysis is only as good as the available sequence data and, as we obtained sequences from both genomic and transcriptomic databases, we cannot say whether II-RIM is expressed in all of the organisms that harbor it within their genomes. While we demonstrate that I-RIM is ubiquitously present in all animals with the exception of ctenophores, II-RIM appears to have undergone independent losses in multiple animal lineages, including at the subphylum level (Figures 2 and 7). In the context of leading animal phylogenies placing ctenophores as the most early-diverging animals (Ryan, et al. 2013; Moroz, et al. 2014; Whelan, et al. 2017), II-RIM likely emerged at the stem lineage of the Metazoa, while I-RIM emerged in the common ancestor of poriferans, placozoans, cnidarians and bilaterians (Figure 7). That II-RIM has been repeatedly lost, but no animal lineage has lost both I-RIM and II-RIM (Paps and Holland 2018), alludes to the importance of RIM genes in animals. This also suggests that these two genes exhibit some functional redundancy, where I-RIM might be more essential given its ubiquity in cnidarians and bilaterians (Figure 2). Molluscs, and perhaps other Lophotrochozoans, represent an interesting case in that they have broadly retained both genes (Figure 2). Based on our qPCR expression analysis of I and II-RIM in various tissues of the freshwater snail *Lymnaea*, it is evident that the two genes differ in their tissue expression levels (Figure 3A). Nevertheless, despite the observation that I-RIM is enriched in the CNS compared to II-RIM, the two RIM genes overlapped in their neuronal expression (Figure 3). Hence, in the CNS, it is possible that the two genes are functionally complementary. Instead, in different tissues, there might be differential requirements in terms of abundance for one gene over the other. *Trichoplax* also retains both homologues, but our mRNA expression and localization studies of RIMs and Ca_V_s point to low level expression (Figures 1 and S1), making it difficult to interpret their cell-type expression profiles and possible roles. We note that in ongoing studies being carried out in our lab, all three *Trichoplax* Ca_V_ channels express *in vitro* to conduct voltage-sensitive Ca^2+^ that resemble those of Ca_V_1-Ca_V_3 homologues from other animals ((Smith, et al. 2017); unpublished data). However, the role of Ca_V_ channels and transient membrane Ca^2+^ signaling in *Trichoplax* biology is unknown. Speculation of the roles RIM proteins might play in ctenophores is equally, if not more, intriguing. Ctenophores exclusively possess II-RIM, making them the only animals with synapses that lack I-RIM. The ctenophore II-RIM is also atypical in that it is predicted to lack an N-terminal Zn^2+^-finger domain, conserved in I-RIMs, II-RIMs and rabphilins (Figure 2). As noted previously, this particular domain is crucial for direct interaction with Munc-13 (Betz, et al. 2001; Dulubova, et al. 2005; Lu, et al. 2006; Quade, et al. 2019), and thus, II-RIM might not play a role in synaptic vesicle priming in ctenophore synapses. Nevertheless, we note broad conservation of predicted N-terminal alpha-helical structures associated with Rab3 binding (Figure S2, Supplementary file 1), suggesting that II-RIMs can interact with vesicles. To our knowledge, ctenophores are the only animals in which the requirement for pre-synaptic Ca^2+^ influx for exocytosis and synaptic transmission has not yet been confirmed, and little is known about the mechanisms for synaptic transmission in these animals (Senatore, et al. 2016). However, microscopy studies have revealed structures with hallmark features of synapse active zones (Hernandez-Nicaise 1973), and ctenophores possess Ca_V_2 channels. Given the proposal that ctenophores independently evolved the nervous system and synapses (Moroz, et al. 2014; Moroz and Kohn 2016), it will be particularly interesting to decipher the roles that Ca_V_2 and II-RIM play in this group of animals.

**Figure 7.**
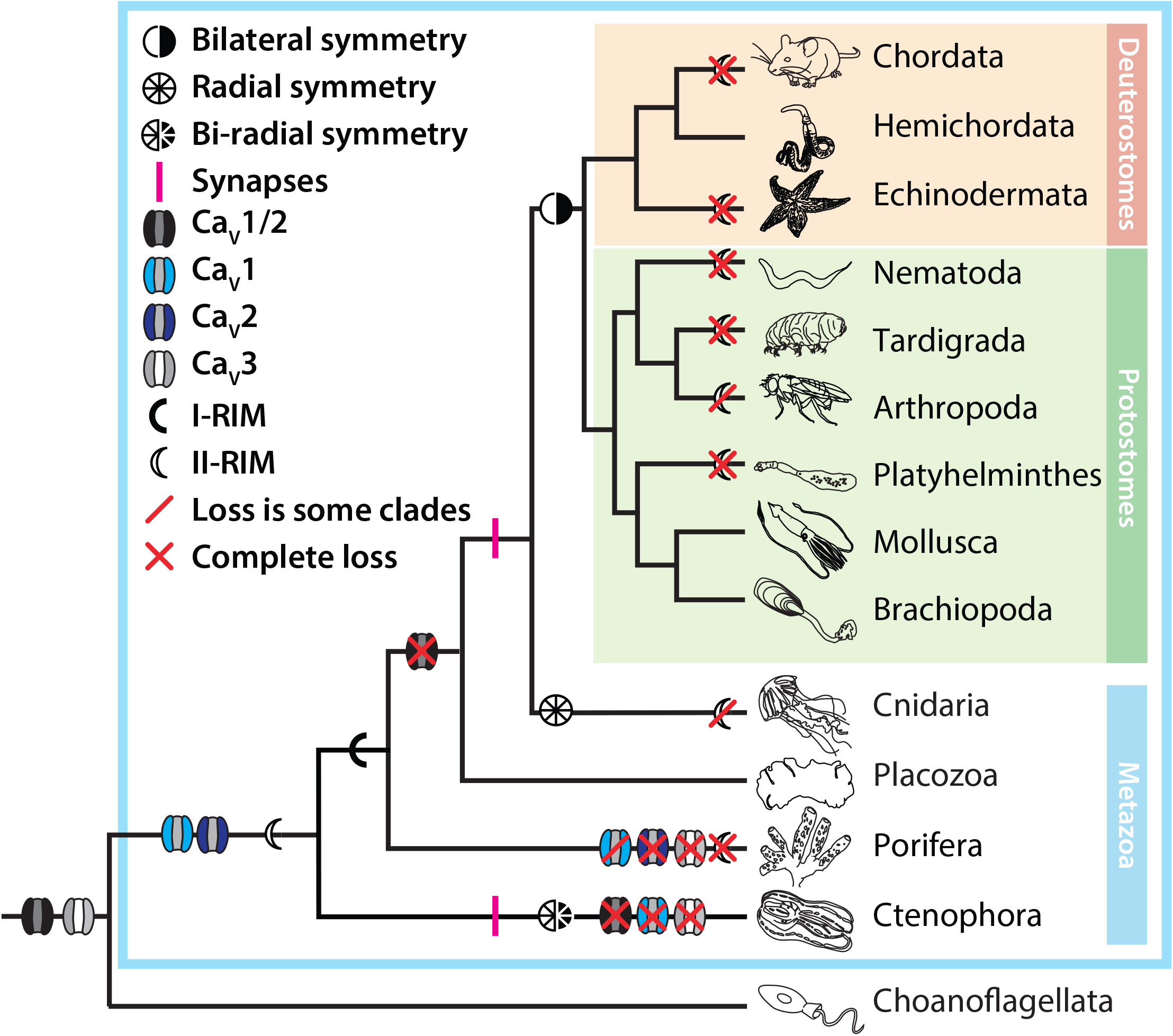
Summary figure of Ca_V_ channel and I-RIM/II-RIM evolution highlighting the putative emergence of Ca_V_1 and Ca_V_2 channels at the stem of Metazoa, and the frequent independent complete or partial loss of II-RIM in various animal phyla. Evident is that Ctenophora is the only animal phylum with synapses but lacking I-RIM, possessing only II-RIM without N-terminal Zn^2+^-finger motifs. The phylogenetic relationships depicted in the tree correspond to contemporary phylogenomic observations and the corresponding hypothesis that ctenophores, and not sponges, are the most early-diverging animals.

The PDZ domain of I-RIM is the physical link for selectively recruiting pre-synaptic Ca_V_2 channels to the active zone (Kaeser et al., 2011). We detailed the primary and secondary structures of I and II-RIM PDZ domains to gain insight into whether they might differ in their ligand binding. Generally, the PDZ domains of both RIM homologues shared a common secondary structure but II-RIM featured lower sequence conservation (Figure 4A). However, the two PDZ domains differed in residue positions involved in ligand selectivity. Specifically, of the three amino acids in the RIM PDZ domain that both interface with bound ligands (Lu, et al. 2005), and are sites for evolutionary divergence in PDZ domain ligand specificity (Sakarya, et al. 2009), two are different between I-RIM and II-RIM, with respective consensus sequences of T**<u>K</u>**V**<u>K</u>** and T**<u>W</u>**I**<u>V</u>** (Figure 4A). That they differ in this key locus, with I-RIM having positively charged lysines and II-RIM neutral tryptophan and valine, provides additional support for II-RIM being an independent clade. Given that these motifs interface with bound ligands, it is also tempting to speculate that the positively charged lysines in the I-RIM PDZ domain provide charge attraction for the conserved negatively charged glutamate and aspartate residues of Ca_V_2 channel D/E-D/E-WC-_COOH_ like motifs (Figure 5A), and hence by extension, that II-RIMs select ligands with different chemical profiles.

One note that is important to mention with respect to the I-RIM/Ca_V_2 interaction, is that although the ligand specificity of the I-RIM PDZ domains appears conserved between rodent and fruit fly (Kaeser, et al. 2011; Graf, et al. 2012), Ca_V_2 channels from nematodes of the rhabditomorpha, including *C. elegans*, lost the D/E-D/E-WC-_COOH_ like motif (Figure 5A). Furthermore, phage display screening of the *C. elegans* I-RIM PDZ domain identified a consensus ligand sequence of F-S/C/I/D-F-W-L/I-_COOH_ (Tonikian, et al. 2008), which is quite different from the C-terminal sequence of its corresponding Ca_V_2 channel, and the acidic motif of other Ca_V_2 channels. Despite these differences, genetic experiments have established that RIM and RIM-BP are redundantly required for active zone localization of Ca_V_2 channels in *C. elegans* (Kushibiki, et al. 2019), suggesting they both directly interact with the channel. A possible explanation for this inconsistency might be that, as has been shown for the human Ca_V_2.1-RIM interaction, RIM can bind at secondary internal sites independently of the distal PDZ ligand motif (Hirano, et al. 2017). Alternatively, the interaction between Ca_V_2 and RIM might be indirect, mediated by shared interactions with RIM-BP (Figure 1A). Interestingly like *C. elegans*, cnidarian I-RIMs exhibit sequence divergence in the PDZ domain <u>TK</u>V<u>K</u> motif, but nevertheless conserved negatively charged D/E-D/E-WC-_COOH_ like motifs in their Ca_V_2 channel C-termini (Figure 5A). Clearly, future wet lab experiments aimed at characterizing these interactions in cnidarians, ctenophores and other animal lineages will be essential towards our understanding of RIM evolution and function in animals.

### Insights into Ca_V_ channel evolution

Our phylogenetic analysis of metazoan Ca_V_ channels revealed deep conservation of C-terminal PDZ ligand motifs for both Ca_V_1 and Ca_V_2 (Figure 5A). Ca_V_1 channels, including the homologue from *Trichoplax*, bear hydrophobic C-termini that fall within the class I PDZ ligands with motifs of X-S/T-X-φ-_COOH_. Little is known about the conservation of PDZ-mediated interactions for Ca_V_1 type channels, which in contrast to Ca_V_2 tend to localize to post-synaptic sites in neurons and muscle. In vertebrates, the scaffolding protein Shank interacts with Ca_V_1.3 channels through both PDZ and SH3 domains, localizing the channels at appropriate post-synaptic locations (Zhang, et al. 2005). Shank is also known to be important in invertebrate post-synaptic functions (Harris, et al. 2016), but, to our knowledge, a direct interaction between Ca_V_1 and Shank has not been reported for any invertebrate. Interestingly, the identified Ca_V_1 homologue from *Oscarella*, and the Ca_V_1/2 homologues from fellow sponges *Amphimedon* and *Haliclona* sp., bear E-T-S/T-V-_COOH_ motifs, corresponding to the consensus sequence for PDZ domains of DLG synaptic scaffolding proteins from human and nematode worm (Tonikian, et al. 2008). This is in contrast to Ca_V_ homologues from pre-metazoan organisms, which have more variable (and positively charged) residues in their extreme C-termini. Based on these observations, it may be that animal-specific adaptations in Ca_V_ channel function occurred early and involved incorporation into specific PDZ domain-mediated interaction networks, a process that is proposed to have given rise to expansion and complexification of PDZ interactions networks in metazoan proteomes (Kim, et al. 2012). The significance of the presence of these motifs in channel homologues from early-diverging animals is unclear, especially given how little is known about Ca_V_ channel function in these animals (Senatore, et al. 2016). Nevertheless, Ca_V_ channel signaling functions are highly dependent on cellular localization and proximity to Ca^2+^-sensitive cytoplasmic proteins. This is because Ca^2+^ can be cytotoxic and tends to be quickly extruded and chelated once inside the cytoplasm (Clapham 2007), restricting high concertation zones to just micrometers from the channel pore (Rizzuto and Pozzan 2006).

Also interesting is that we identified a Ca_V_1 channel in the gene data for the sponge *Oscarella carmela*, significant because Ca_V_1 channels were thought to be absent in sponges (Moran and Zakon 2014; Moran, et al. 2015). Furthermore, we identified a structural feature that distinguishes Ca_V_1 and Ca_V_2 channels in the alpha helical structure predicted in C-termini of Ca_V_1 channels including the *Oscarella* homologue (Figure 6A and B). We also identified an additional structural feature that phylogenetically distinguishes Ca_V_1/Ca_V_2 channels and Ca_V_3 channels, the EVH1 binding motifs in the C-termini of Ca_V_1/Ca_V_2 channels upstream of the IQ motif (Figure 6C). Here, the potential for interactions with Homer and other EVH1 domain-bearing proteins further alludes to differential integration into membrane-localizing protein complexes as a mechanism for Ca_V_ channel adaptation for distinct cellular functions.

Our analysis of proline-rich SH3 ligands in Ca_V_ channel C-termini was less clear than for PDZ ligands. Our impetus for performing this analysis was the consideration of the tripartite interaction between Ca_V_2, I-RIM and RIM-BP conserved between protostome and deuterostome bilaterians, though we were aware that SH3 domain ligands exhibit a considerable degree of sequence entropy and are difficult to predict with confidence (Teyra, et al. 2017). Indeed, although it is likely that the various linkers and N-/C-termini of Ca_V_ channels bear conserved binding sites for RIM-BP SH3 domains, we point out a flaw in this analysis where SH3 ligand motifs were not predicted for the *C. elegans* Ca_V_2, despite its expected interactions with RIM-BP *in vivo* (Gracheva, et al. 2008). Nevertheless, it is notable that SH3 ligands appear enriched in Ca_V_1 and Ca_V_2 channels relative to Ca_V_3 and pre-metazoan Ca_V_s, outside of Ca_V_3.2 and Ca_v_3.3 in chordates (Figure 5A and B).

Based on our presented analyses, Ca_V_1/2 channels appear to have emerged just prior to the divergence of animals from closely-related eukaryotes (Figure 7), upon which they took on the capacity to interact with PDZ domain bearing proteins, constraining their Ca^2+^ signaling functions to discrete subcellular locations. The identification of a Ca_V_1 channel in sponges, Ca_V_2 channels in ctenophores, and a Ca_V_3 channel in choanoflagellates, suggests that the last common ancestor to all animals possessed a full complement of Ca_V_ channels: Ca_V_1-Ca_V_3 plus Ca_V_1/2 channels. Under this model, and consistent with reports of substantial loss of ion channel genes in early-diverging groups (Liebeskind, et al. 2015), ctenophores lost Ca_V_1, Ca_V_3 and Ca_V_1/2 channels, sponges lost Ca_V_3, Ca_V_2 and either Ca_V_1 or Ca_V_1/2, while placozoans and the remaining animal groups lost Ca_V_1/2 channels but retained Ca_V_1-Ca_V_3 (Figure 7). This model supports the notion that Ca_V_1 and Ca_V_2 channels evolved from an ancestral Ca_V_1/2-like channel (Moran and Zakon 2014; Moran, et al. 2015), but suggests that all three channel types co-existed in an ancestral species. If so, early in the divergence between Ca_V_1 and Ca_V_2 channels, they took on differential functional attributes, such as the pronounced Ca^2+^-dependent inactivation of Ca_V_1 compared to Ca_V_2 channels mediated by interactions with calmodulin at the C-terminal IQ motif (Catterall 2011; Taiakina, et al. 2013). Included in the divergence between Ca_V_1 and Ca_V_2 channels, which are respectively specialized for post- and pre-synaptic functions (Senatore, et al. 2016), might have been differential incorporation into distinct membrane complexes including those mediated by scaffolding proteins bearing PDZ domains.

## Materials and Methods

### mRNA quantification and localization

*Trichoplax adhaerens* animals were prepared for fluorescence *in situ* hybridization (FISH) by freezing in tetrahydrafuran (THF) overnight on dry ice followed by fixation in 3% acetic acid in methanol (MeOH) for 30 minutes at −20°C and then 4% paraformaldehyde in methanol for 30 minutes at room temperature, as described (Mayorova, et al. 2019). *In situ* hybridization was performed with RNAscope probes for I-RIM (#72781-C3), II-RIM (#572791-C2), Ca_V_1 (#442461), Ca_V_2 (#442471) and Ca_V_3 (#488711) and Multiplex Fluorescent Assay reagents (#320850) from Advanced Cell Diagnostics (Hayward, CA, USA). For dual labeling with probes for Ca_V_1, Ca_V_2 or Ca_V_3 and CF-405 conjugated wheatgerm agglutinin (#29027-1, Biotium, Freemont, CA), animals were frozen in THF as described above and then fixed in 4% formaldehyde in MEOH for 30 minutes at −20°C and 30 minutes at room temperature. Following *in situ* hybridization, the samples were incubated in CF-405 WGA diluted 1:200 in PBS for 1 hr at room temperature. Fluorescence images were collected with a 63X NA 1.4 objective on a LSM880 laser scanning confocal microscope (Carl Zeiss Microscopy LLC, Thornwood, NY, USA). Images in Figure 1C-E, left panels, were collected with a 32-channel spectral detector in the lambda mode and processed by linear unmixing. Enlarged views in Figure 1C-E insets were collected with an AiryScan detector. Projected images were generated with Zen software (Carl Zeiss Microscopy LLC).

For the qPCR experiments, young adult *Lymnaea stagnalis* albumen gland, buccal mass, brain, heart, and prostate gland were micro-dissected from anesthetized animals and pooled into triplicate tubes (5-10 individual tissues per tube), and total RNA extracted as previously reported (Senatore, et al. 2014). Complimentary DNA (cDNA) libraries were prepared from each RNA isolate with SuperScript III reverse transcriptase (ThermoFisher Scientific, Canada) and an anchored oligo-dT_18_ primer (Table 1). Gene-specific primers for *Lymnaea* I-RIM (NCBI accession FX186940.1), II-RIM (NCBI accession FX181400.1) and elongation factor-1α (EF-1α; NCBI accession DQ278441.1) (Table 1) were used for quantitative PCR using the iQ SYBR Green Supermix (BioRad, Canada) and the following cycling conditions: denaturation at 95°C for 3 minutes, followed by 40 cycles of 95°C for 15 seconds (denaturation) and 57–60°C for 30 seconds (extension). To ensure that single amplicons were produced in each PCR reaction, melting curve protocols were performed after each run. Real-time PCR fluorescence measurement and melt curve analyses were done using a Bio-Rad C1000™ Thermal Cycler equipped with a CFX96™ System (Bio-Rad). Transcript expression levels were quantified and normalized relative to EF-1α using the ΔΔ cycle threshold (ΔΔ CT) method (Andersen, et al. 2004): ratio = (E_target gene_) ΔCT_Target gene_/(E_EF-1α_) ΔCT_EF-1α_, where E denotes PCR efficiency for respective PCR primer pairs. Normalized transcript abundance of all tissues was standardized to the transcript abundance of the I-RIM and II-RIM in the albumen, which was set to 100%. One-way ANOVA was also performed to confirm that transcript abundance of EF-1α did not significantly differ between tissues (P = 0.217).

**Table 1.**
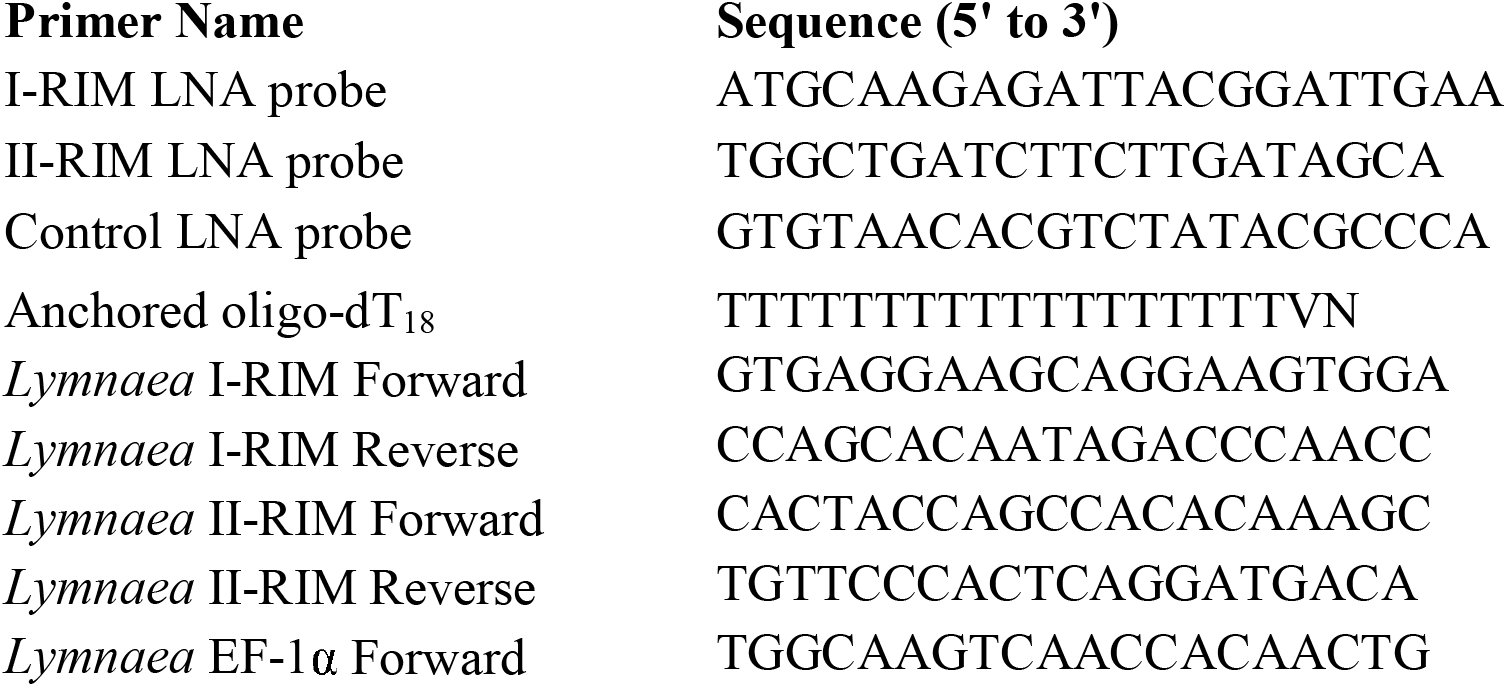
Sequence of oligonucleotides used for *Lymnaea* qPCR and *in situ* hybridization.

For the FISH experiments on isolated neurons from the CNS of *Lymnaea stagnalis*, central ring ganglia (CNS) from young adult *Lymnaea stagnalis* (16-18 mm in length) were isolated and exposed to antibiotic washes prior to cell culture. Individual ganglia were then desheathed, enabling removal of specific, identified neurons using suction applied via a fire polished pipette. Individual cells were plated on poly-L-lysine (Millipore Sigma) coated Falcon dishes (VWR) following isolation. Cells were given 10-30 minutes to attach to the cell culture plates and were then fixed using 4% paraformaldehyde. Prior to staining, cells were treated with 3% H_2_O_2_ (v/v in 1xPBS; Millipore Sigma, Canada) to eliminate endogenous peroxidase activity and dehydrated overnight in 70% ethanol. The next day, cells were incubated in hybridization buffer (25% formamide, 0.05M EDTA, 4x saline-sodium citrate buffer (SSC), 10% dextran sulphate, 1x Denharts solution, 0.5mg/mL *E.Coli* tRNA, 20 mM ribonucleoside vanadyl complexes and 9.2 mM citric acid; Millipore Sigma) at 55°C for 2 hours. Following pre-hybridization, cells were incubated in either 10 nM of LNA enhanced detection probes (Qiagen) targeting mRNAs for I-RIM, II-RIM or a negative control probe (Table 1) at 55°C for 1 hour. Next, cells were washed in a series of stringency washes, including 4x SSC, 2x SSC, 1x SSC, and 0.1x SSC at 37°C for 30 minutes, incubated in blocking buffer (3% bovine serum, 4x SSC, 0.1x Tween-20) for 30 minutes, then horseradish peroxidase-conjugated streptavidin (Thermo-Fisher) for 30 minutes. Cells were then washed in TNT buffer (0.1 M Tris HCl, 0.15 M NaCl, 0.05% Tween-20), then incubated in tyramide (Perkin-Elmer) according to the manufacturer’s instructions for 1 hour at room temperature. Following additional washes in TNT buffer, cells were mounted in Fluoroshield™ containing DAPI (Millipore Sigma) as a counterstain to label nuclei. Cells were imaged using a Carl Zeiss Axio Observer.Z1 inverted light/epifluorsecence microscope, with Apotome.2 optical sectioning (Zeiss). For all images, Z-stack slices were taken at 0.29 µm intervals and were rendered into 2D maximum intensity projections using the Zeiss Zen 2 microscopy software.

### Phylogenetic inference

Both maximum likelihood and Bayesian inference strategies were used to infer the phylogenetic relationships of I-RIMs, II-RIMs and rabphilins. Briefly, candidate protein sequences were identified in various databases using reciprocal BLAST search, smart BLAST, and InterPro domain analysis. Protein sequences were then aligned with MUSCLE (Edgar 2004) and trimmed with the trimAl (Capella-Gutiérrez, et al. 2009) using a gap threshold of 0.6 (accession numbers listed below; raw sequences and trimmed alignment are provided in supplementary file 1). The trimmed alignment was then used to infer a maximum likelihood phylogenetic tree using IQ-TREE (Nguyen, et al. 2014) with default parameters and 1,000 ultrafast bootstrap replicates for estimating node support, and a best-fit substitution model of VT+F+G4. Using the same alignment and model, bayesian inference was done using MrBayes version 3.2.6 (Ronquist, et al. 2012), with four Markov chains, 10,000,000 generations, a tree sampling frequency of 100, and a burn-in fraction 0.25. Phylogenetic trees were visualized with MEGA X (Kumar, et al. 2018) and FigTree 1.4.3 (Rambaut 2007), and shared nodes with respectively strong bootstrap support and high posterior probability values were annotated on the maximum likelihood tree shown in Figure 1. NCBI protein sequence accession numbers (unless otherwise indicated) are as follows: *H.sapiens* I-RIM1α: NP_055804.2; *M.musculus* I-RIM1α: NP_444500.1; *R.norvegicus* I-RIM1α: Q9JIR4.1; *G.gallus* I-RIM1α: XP_025004772.1; *H.sapiens* I-RIM2α: Q9UQ26.2; *M.musculus* I-RIM2α: NP_444501.1; *R.norvegicus* I-RIM2α: NP_446397.1; *G.gallus* I-RIM2α: XP_015138471.1; *S. kowalevskii* I-RIM: sakowv30037565m (OIST Marine Genomics Unit); *P.flava* I-RIM: pfl_40v0_9_20150316_1g5731.t1 (OIST Marine Genomics Unit); *S.purpuratus* I-RIM: XP_011664650.1; *A.planci* I-RIM: XP_022091188.1; *A.californica* I-RIM: XP_005106594.2; *C.gigas* I-RIM: XP_019921964.1; *M.yessoensis* I-RIM: XP_021377513.1; *O.bimaculoides* I-RIM: XP_014789722.1; *E.granulosus* I-RIM: XP_024347539.1; *S.haematobium* I-RIM: XP_012796083.1; *L.anatina* I-RIM: XP_013421607.1; *L.polyphemus* I-RIM (1065aa): XP_022257304.1; *L.polyphemus* I-RIM (1630aa): XP_022243248.1; *C.sculpturatus* I-RIM: XP_023224599.1; *H.azteca* I-RIM: XP_018007323.1; *D.melanogaster* I-RIM: NP_001247161.2; *H.dujardini* I-RIM: OQV23161.1; *R.varieornatus* I-RIM: GAV01719.1; *C.elegans* I-RIM: NP_741831.1; *N.vectensis* I-RIM: evg1261940 (from in house transcriptome assembly (Wong, et al. 2019)); *E.pallida* I-RIM: XP_020898842.1; *H.vulgaris* I-RIM: XP_012564401.1; *T.adhaerens* I-RIM: evg1642237 (from in house transcriptome assembly (Wong, et al. 2019)); *O.carmela* I-RIM (739aa): m.21147 (Compangen); *O.carmela* I-RIM (1164aa): m.26069 (Compangen); *S. kowalevskii* II-RIM: sakowv30000298m (OIST Marine Genomics Unit); *A.californica* II-RIM: XP_012944640.1; *C.gigas* II-RIM: XP_011422085.1; *M.yessoensis* II-RIM: XP_021345230.1; *O.bimaculoides* II-RIM: XP_014774175.1; *L.anatina* II-RIM: XP_013397099.2; *C.sculpturatus* II-RIM: XP_023227959.1; *L.polyphemus* II-RIM (1313aa): XP_022258855.1; *L.polyphemus* II-RIM (1323aa): XP_022240908.1; *M.leidyi* II-RIM: evg198193 (Wong, et al. 2019); *B.ovata* II-RIM: combined overlapping fragmented transcripts TR51711|c1_g1_i2 and TR51711|c1_g3_i7 from unpublished transcriptome generated by Mark Martindale and Joseph Ryan, Whitney Lab, Florida; *H.californiensis* II-RIM: evg158061 (Wong, et al. 2019); *E.pallida* II-RIM: KXJ21887.1; *T.adhaerens* II-RIM: evg1176111 (Wong, et al. 2019); *H.sapiens* rabphilin: NP_001137326.1; *M.musculus* rabphilin: NP_001289273.1; *R.norvegicus* rabphilin: NP_598202.1; *G.gallus* rabphilin: XP_015131024.1; *S. kowalevskii* rabphilin: XP_006817123.1; *S.purpuratus* rabphilin: XP_030848070.1; *A.planci* rabphilin: XP_022085716.1; *A.californica* rabphilin: XP_012945559.1; *C.gigas* rabphilin: XP_011424844.1; *M.yessoensis* rabphilin: XP_021345406.1; *O.bimaculoides* rabphilin: XP_014779575.1; *L.anatina* rabphilin: XP_013406958.1; *L.polyphemus* rabphilin: XP_013782034.1; *R.varieornatus* rabphilin: GAV03979.1; *H.dujardini* rabphilin: OQV25295.1; *D.melanogaster* rabphilin: NP_572651.1; *H.azteca* rabphilin: XP_018018219.1; *C.elegans* rabphilin: NP_001022566.1; *S.haematobium* rabphilin: XP_012797599.1; *E.granulosus* rabphilin: CDS16155.1; *A.digitifera* rabphilin: XP_015773295.1; *E.pallida* rabphilin: XP_020895735.1; *H.vulgaris* rabphilin: XP_012561655.1; *M.leidyi* rabphilin: evg19024 (Wong, et al. 2019); *B.ovata* rabphilin: TR42475|c0_g1_i1 (Mark Martindale and Joseph Ryan); *H.californiensis* rabphilin: evg140073 (Wong, et al. 2019); *T.adhaerens* rabphilin: evg1107189.2 (Wong, et al. 2019); *O.carmela* rabphilin: m.15463 (Compangen).

To infer the phylogenetic relationships of Ca_V_ channels, a similar approach was used as described for the RIM/rabphillin maximum likelihood tree, with manual trimming of the protein alignment, and a substitution model of LG+I+G4 selected by IQ-TREE (sequences and trimmed alignment provided in respective supplementary files 2 and 3). NCBI protein sequence accession numbers (unless otherwise indicated) are as follows: *C.porosus* Ca_V_1.1: XP_019401893.1; *G.gallus* Ca_V_1.1: NP_001292076.1; *P.vitticeps* Ca_V_1.1: XP_020660962.1; *G.japonicus* Ca_V_1.1: XP_015267320.1; *H.sapiens* Ca_V_1.1: NP_000060.2; *D.rerio* Ca_V_1.1: NP_001139622.1; *C.porosus* Ca_V_1.2: XP_019400097.1; *G.gallus* Ca_V_1.2: XP_015142138.1; *H.sapiens* Ca_V_1.2: Q13936.4; *P.vitticeps* Ca_V_1.2: XP_020644644.1; *G.japonicus* Ca_V_1.2: XP_015278355.1; *C.picta bellii* Ca_V_1.2: XP_008162493.1; *D.rerio* Ca_V_1.2: XP_009298610.1; *P.vitticeps* Ca_V_1.3: XP_020659553.1; *G.japonicus* Ca_V_1.3: XP_015274327.1; *C.picta bellii* Ca_V_1.3: XP_023956012.1; *G.gallus* Ca_V_1.3: XP_015148473.1; *H.sapiens* Ca_V_1.3: NP_001122312.1; *C.porosus* Ca_V_1.3: XP_019392560.1; *D.rerio* Ca_V_1.3: XP_021335256.1; *P.vitticeps* Ca_V_1.4: XP_020648615.1; *A.carolinensis* Ca_V_1.4: XP_016846283.1; *P.humilis* Ca_V_1.4: XP_005533405.1; *H.sapiens* Ca_V_1.4: NP_005174.2; *C.intestinalis* Ca_V_1: XP_018667169.1; *H.roretzi* Ca_V_1: BAA34927.2; *A.melifera* Ca_V_1: XP_016766333.1; *A.aegypti* Ca_V_1: XP_021699870.1; *D.melanogaster* Ca_V_1: AAF53504.1; *D.pulex* Ca_V_1: EFX89598.1; *D.magna* Ca_V_1: KZS09199.1; *C.elegans* Ca_V_1: NP_001023079.1; *C.nigoni* Ca_V_1: PIC34019.1; *D.pachys* Ca_V_1: PAV74346.1; *P.pacificus* Ca_V_1: PDM72512.1; *S.ratti* Ca_V_1: XP_024499702.1; *T.britovi* Ca_V_1: KRY55670.1; *T.pseudospiralis* Ca_V_1: KRY90155.1; *L.stagnalis* Ca_V_1: AAO83839.1; *C.gigas* Ca_V_1: XP_011452714.1; *O.bimaculoides* Ca_V_1: XP_014774811.1; *L.anatina* Ca_V_1: XP_023932094.1; *A.planci* Ca_V_1: XP_022081221.1; *S.purpuratus* Ca_V_1: XP_011670134.1; *N.vectensis* Ca_V_1: combined overlapping fragmented transcripts evg1131374 and TRINITY_DN40077_c0_g1_i4 (Wong, et al. 2019); *E.pallida* Ca_V_1: XP_020903719.1; *A.digitifera* Ca_V_1: XP_015778662.1; *C.capillata* Ca_V_1: AAC63050.1; *T.adhaerens* Ca_V_1: evg1032956 (Wong, et al. 2019); *O.carmela* Ca_V_1: combined overlapping fragmented transcripts m.84361 and m.186460 and g4118.t1 (Compagen); *A.mississippiensis* Ca_V_2.1: XP_019354952.1; *S.vulgaris* Ca_V_2.1: XP_014748708.1; *G.gallus* Ca_V_2.1: ATE62979.1; *C.picta bellii* Ca_V_2.1: XP_008161515.1; *H.sapiens* Ca_V_2.1: O00555.3; *P.vitticeps* Ca_V_2.1: XP_020656028.1; *G.japonicus* Ca_V_2.1: XP_015274358.1; *D.rerio* Ca_V_2.1: XP_021330116.1; *C.picta bellii* Ca_V_2.2: XP_008172570.1; *C.porosus* Ca_V_2.2: XP_019396155.1; *G.gallus* Ca_V_2.2: XP_015134766.1; *P.vitticeps* Ca_V_2.2: XP_020668759.1; *G.japonicus* Ca_V_2.2: XP_015276634.1; *H.sapiens* Ca_V_2.2: NP_000709.1; *D.rerio* Ca_V_2.2: XP_021331856.1; *C.porosus* Ca_V_2.3: XP_019386495.1; *G.gallus* Ca_V_2.3: XP_015145962.1; *C.picta bellii* Ca_V_2.3: XP_008164802.1; *P.vitticeps* Ca_V_2.3: XP_020645405.1; *A.carolinensis* Ca_V_2.3: XP_008107103.1; *H.sapiens* Ca_V_2.3: NP_001192222.1; *D.rerio* Ca_V_2.3: XP_017206695.2; *C.intestinalis* Ca_V_2: XP_018670105.1; *A.melifera* Ca_V_2: XP_016766516.1; *A.aegypti* Ca_V_2: XP_021710362.1; *D.melanogaster* Ca_V_2: AFH07350.1; *H.azteca* Ca_V_2: XP_018019172.1; *C.elegans* Ca_V_2: NP_001123176.1; *C.nigoni* Ca_V_2: PIC16379.1; *H.contortus* Ca_V_2: CDJ96819.1; *S.ratti* Ca_V_2: XP_024503488.1; *T.britovi* Ca_V_2: KRY49212.1; *T.pseudospiralis* Ca_V_2: KRY73966.1; *L.stagnalis* Ca_V_2: AAO83841.1; *A.californica* Ca_V_2: AVD53847.1; *C.gigas* Ca_V_2: XP_019920407.1; *H.bleekeri* Ca_V_2: BAA13136.2; *A.planci* Ca_V_2: XP_022085274.1; *S.purpuratus* Ca_V_2: XP_011662956.1; *N.vectensis* Ca_V_2a: combined overlapping fragmented transcripts evg194605 (Wong, et al. 2019) and NVE4667 (David, et al. 2013); *E.pallida* Ca_V_2a: XP_020910418.1; *O.faveolata* Ca_V_2a: XP_020626975.1; *A.digitifera* Ca_V_2a: XP_015773841.1; *N.vectensis* Ca_V_2b: combined overlapping fragmented transcripts TRINITY_DN6634_c0_g1_i1 and evg1115545 (Wong, et al. 2019); *E.pallida* Ca_V_2b: XP_020906771.1; *O.faveolata* Ca_V_2b: XP_020612369.1; *S.pistillata* Ca_V_2b: XP_022780503.1; *N.vectensis* Ca_V_2c: combined overlapping fragmented transcripts TRINITY_DN35757_c0_g1 and evg1243599 (Wong, et al. 2019); *O.faveolata* Ca_V_2c: XP_020613420.1; *S.pistillata* Ca_V_2c: PFX33508.1; *T.adhaerens* Ca_V_2: evg1041627 (Wong, et al. 2019); *S.rosetta* Ca_V_1/2: XP_004989719.1; *A. queesnlandica* Ca_V_1/2: Aqu2.38198_001 (Fernandez-Valverde, et al. 2015); *H.ambioensis* Ca_V_1/2: combined overlapping fragmented transcripts m.28368 and m.28207 (Compagen); *H.tubifera* Ca_V_1/2: m.43115 (Compagen); *C.porosus* Ca_V_3.1: XP_019398025.1; *G.gallus* Ca_V_3.1: XP_015150965.1; *P.vitticeps* Ca_V_3.1: XP_020670763.1; *G.japonicus* Ca_V_3.1: XP_015266211.1; *H.sapiens* Ca_V_3.1: NP_061496.2; *D.rerio* Ca_V_3.1: XP_021336020.1; *A.mississippiensis* Ca_V_3.3: XP_019344978.1; *G.gallus* Ca_V_3.3: XP_015144216.1; *G.japonicus* Ca_V_3.3: XP_015274652.1; *P.vitticeps* Ca_V_3.3: XP_020665783.1; *H.sapiens* Ca_V_3.3: NP_066919.2; *D.rerio* Ca_V_3.3: XP_021329632.1; *C.porosus* Ca_V_3.2: XP_019397684.1; *G.gallus* Ca_V_3.2: XP_015149910.1; *P.vitticeps* Ca_V_3.2: XP_020633697.1; *H.sapiens* Ca_V_3.2: NP_066921.2; *D.rerio* Ca_V_3.2: XP_009297960.1; *C.intestinalis* Ca_V_3: XP_018667817.1; *A.melifera* Ca_V_3: NP_001314887.1; *A.aegypti* Ca_V_3: XP_021706157.1; *H.azteca* Ca_V_3: XP_018025902.1; *L.stagnalis* Ca_V_3: AAO83843.2; *B.glabrata* Ca_V_3: XP_013096433.1; *C.gigas* Ca_V_3: XP_011439125.1; *C.elegans* Ca_V_3: CCD68017.1; *C.nigoni* Ca_V_3: PIC19332.1; *D.pachys* Ca_V_3: PAV61762.1; *P.pacificus* Ca_V_3: PDM63609.1; *S.ratti* Ca_V_3: XP_024510580.1; *T.britovi* Ca_V_3: KRY57796.1; *T.pseudospiralis* Ca_V_3: KRZ43346.1; *A.planci* Ca_V_3: XP_022091428.1; *N.vectensis* Ca_V_3a: NVE5017 (David, et al. 2013); *E.pallida* Ca_V_3a: XP_020900273.1; *A.digitifera* Ca_V_3a: XP_015765864.1; *O.faveolata* Ca_V_3a: XP_020622701.1; *S.pistillata* Ca_V_3a: XP_022794522.1; *N.vectensis* Ca_V_3b: NVE7616 (David, et al. 2013); *E.pallida* Ca_V_3b: XP_020912298.1; *A.digitifera* Ca_V_3b: XP_015766817.1; *O.faveolata* Ca_V_3b: XP_020626608.1; *S.pistillata* Ca_V_3b: XP_022784214.1; *T.adhaerens* Ca_V_3: evg954676 (Wong, et al. 2019); *S.rosetta* Ca_V_3: XP_004995501.1; *C.reinhardtii* CAV: XP_001701475.1; *G.pectorale* CAV: KXZ52368.1; *C.eustigma* CAV: GAX86028.1; *P.tetraurelia* Ca_V_1a: GSPATP00010323001 (Lodh, et al. 2016); *P.tetraurelia* Ca_V_1b: combined overlapping fragmented transcripts GSPATG00033414001 and GSPATG00033415001 (Lodh, et al. 2016); *P.tetraurelia* Ca_V_1c: GSPATP00010323001 (Lodh, et al. 2016); *S.cerevisae* CCH1: NP_011733.3; and *S.pombe* CCH1: NP_593894.1.

### Protein alignments and structural predictions

Protein alignments were generated using MUSCLE (Edgar 2004) within the MEGA X software (Kumar, et al. 2018), and visualized with JalView (Waterhouse, et al. 2009). Jalview was also used to generate consensus and conservation plots (Figures 4A and S2). PROMALS3D was used to predict all secondary structures (Pei and Grishin 2014) (Figures 4A, S2 and 6B), with the exception of alpha helices predicted at the N-termini of RIM and rabphilin homologues (Supplementary file 1). EMBOSS plotcon (Rice, et al. 2000) was used to generate conservation vs. position in alignment plots with a running amino acid alignment window of 15 (Figure 4B) or 6 (Figure 6). Protein domains, including PDZ, SH3, Zn^2+^ and C_2_A/C_2_B (Figures 1A and 2), were predicted with InterProScan (Jones, et al. 2014), and secondarily with hmmscan (Finn, et al. 2011). PDZ ligand motifs (Figure 5A) were predicted using PDZPepInt (Kundu, et al. 2014), and SH3 ligand domains (Figures 5A and S3) were predicted using three separate algorithms: 1) Find Individual Occurrences (FIMO) (Grant, et al. 2011); 2) LMDIPred (Sarkar, et al. 2018) and 3) SH3PepInt (Kundu, et al. 2014). FIMO, part of the MEME Suite of sequence analysis tools, identified all cases of the consensus SH3 binding motif PXXP in Ca_V_ C-terminal sequences, providing a liberal estimate of the actual number of motifs. SH3PepInt used a graph-kernal algorithm to predict SH3 motifs based on peptide-array data for 69 human SH3 domains and 31 regular expressions for canonical SH3 motifs (run using the default 15mer peptide window and a step size of 5 amino acids). Linear Motif Domain Interaction Prediction (LMDIPred) used four independent methods (support vector machine (SVM) prediction, position specific scoring matrix (PSSM), motif instance matching, and regular expression scanning) to predict 6-mer SH3 binding motifs. We counted a SH3 motif only if it was identified using three or more of the independent LMDIPred algorithms. Statistical analysis of SH3PepInt-predicted SH3 ligand motifs was performed first conducting normality assessment with Shapiro-Wilk tests, and homogeneity of variance with Levene’s test (ANOVA on residuals). Post-hoc analysis was done using Kruskal-Wallis and Dunn’s tests with Benjamini-Hochberg p-value adjustment. This same approach was used for comparing the three different algorithms used to predict SH3 ligand domains (Figure S3). For comparing the lengths of intracellular regions of various Ca_V_ channels (Figure S4), PSIPRED (McGuffin, et al. 2000), TMHMM (Krogh, et al. 2001) and ExPASy ProtScale Kyte-Doolittle plots (Gasteiger, et al. 2005) were used to identify interfaces between transmembrane and cytoplasmic/extracellular regions. *De novo* motif identification (particularly SLiMs; Figure 6) was performed using Swiss Institute of Bioinformatics (SIB) MyHits Motif Scan (Pagni, et al. 2007), using HAMAP, PROSITE, and Pfam HMM databases; and hits were cross-referenced with existing entries in the Eukaryotic Linear Motifs (ELM) database (Dinkel, et al. 2011); and manually inspected to identify tandem amino acid repeats (AARs)

## Acknowledgements

We would like to thank Drs. Mark Martindale and Joseph Ryan (Whitney Lab, Florida) for providing us with access to gene sequences from unpublished genome and transcriptome data of the ctenophore *Beroe ovata*.

## Figure Legends

**Figure S1.** Fluorescence *in situ* hybridization (FISH) with probes for RIM and voltage gated calcium channel (Ca_V_1, Ca_V_2 or Ca_V_3) genes in wholemounts of *Trichoplax*. **A)** Merged and color-separated images of a narrow region extending halfway across the animal. Each image is a projection of a complete series of horizontal optical sections through the animal. Ca_V_1 is more highly expressed in a narrow zone near the edge and in the zone corresponding to the area occupied by lipophil cells (lipophil zone) than in the intervening area. Ca_V_2, Ca_V_3, I-RIM and II-RIM have more uniform radial expression patterns. Additional images from the same samples are illustrated in Figure 1C-E. **B)** FISH with probes for Ca_V_1, Ca_V_2 or Ca_V_3 in wholemounts labeled with fluorescent wheatgerm agglutinin (WGA), a marker for mucocytes. Each image is a single 0.8 µm (theoretical estimate) optical section. Arrowheads mark mucocytes that contain Ca_V_1 or Ca_V_2 label. Examination of three-dimensional reconstructions of entire mucocytes (not shown) confirmed that that some had Ca_V_1 or Ca_V_2 label in their interiors. Few mucocytes contained Ca_V_3 label. Scale, 20 µm (A); 5 µm (B).

**Figure S2. A)** Alignment of the Zn^2+^-finger structural motif of I-RIM, II-RIM and rabphilin sequences from representative species of the Metazoa. Conserved cysteine residues putatively involved in Zn^2+^ binding are delineated by yellow boxes. The SGAWFF structural element mediating binding to Rab3 is outlined in green. **B)** Alignment of the C_2_A and C_2_B domains of I-RIM, II-RIM and rabphilin sequences from representative species. Conserved aspartate (D) residues, putatively involved in Ca^2+^ binding, are delineated by red boxes.

**Figure S3.** Average number of predicted SH3 biding motifs in the C-termini of metazoan and pre-metazoan Ca_V_s using the FIMO, SH3PepInt and LMDIPred algorithms. Differences in the number of SH3 motifs predicted by each algorithm were statistically significant for Ca_V_1, Ca_V_2, Ca_V_3 and pre-metazoan Ca_V_s (Kruskal-Wallis and Dunn’s post-hoc test with the Benjamini-Hochberg p-value adjustment). Statistical analysis could not be performed for Ca_V_1/2 (ancestral HVA) due to lack of sufficient number of sequences. Error bars denote standard deviation.

**Figure S4.** Variability in amino acid sequence length of the intrinsically disordered termini and linkers of metazoan and pre-metazoan Ca_V_ channels. The top schematic illustrates the linear transmembrane structure of Ca_V_ channel α subunits, defined by four homologous domains comprised of six transmembrane segments (S1-S6) connected by intrinsically disordered cytosolic linker regions, and the similarly disordered N- and C-termini. The HVA channel alpha interaction domain (AID) and IQ motifs, located in the I-II linker and C-terminus respectively, are also depicted (both are absent in LVA Ca_V_3 channels). Diamonds indicate lengths of individual channel linkers/termini. Vertical lines denote average linker/termini lengths per clade. Black Xs denote truncated or missing regions for select entries. Uppercase letters denote statistical significance for comparisons between Ca_V_ channel types, and lowercase letters denote significance for comparisons between different animal clades for any particular Ca_V_ channel type.

