## Supplementary figures and images for "Early metazoan origin and multiple losses of a novel clade of RIM pre-synaptic calcium channel scaffolding protein homologues"

### Figure S1

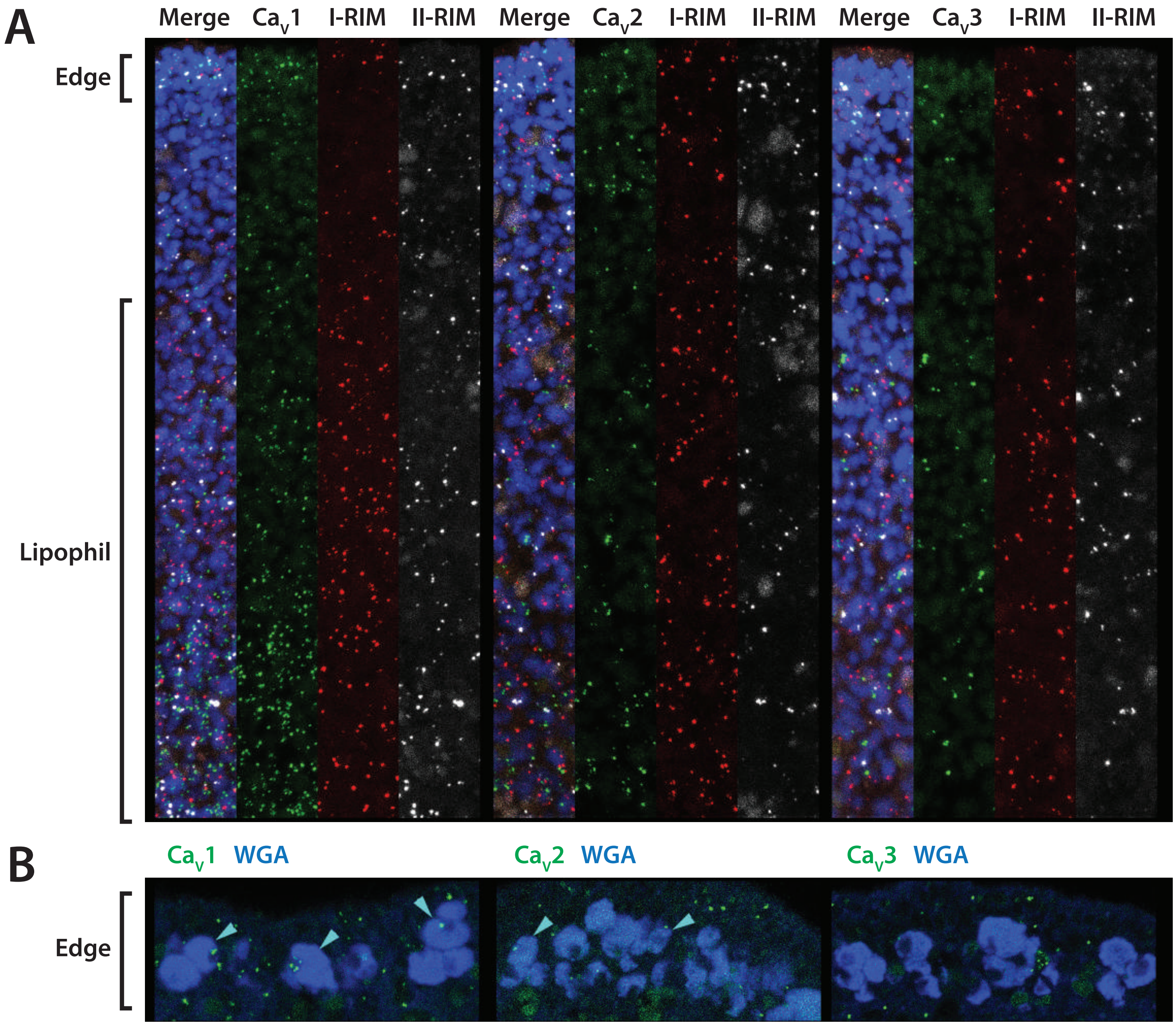


Supplementary Figure 1.

### Figure S2

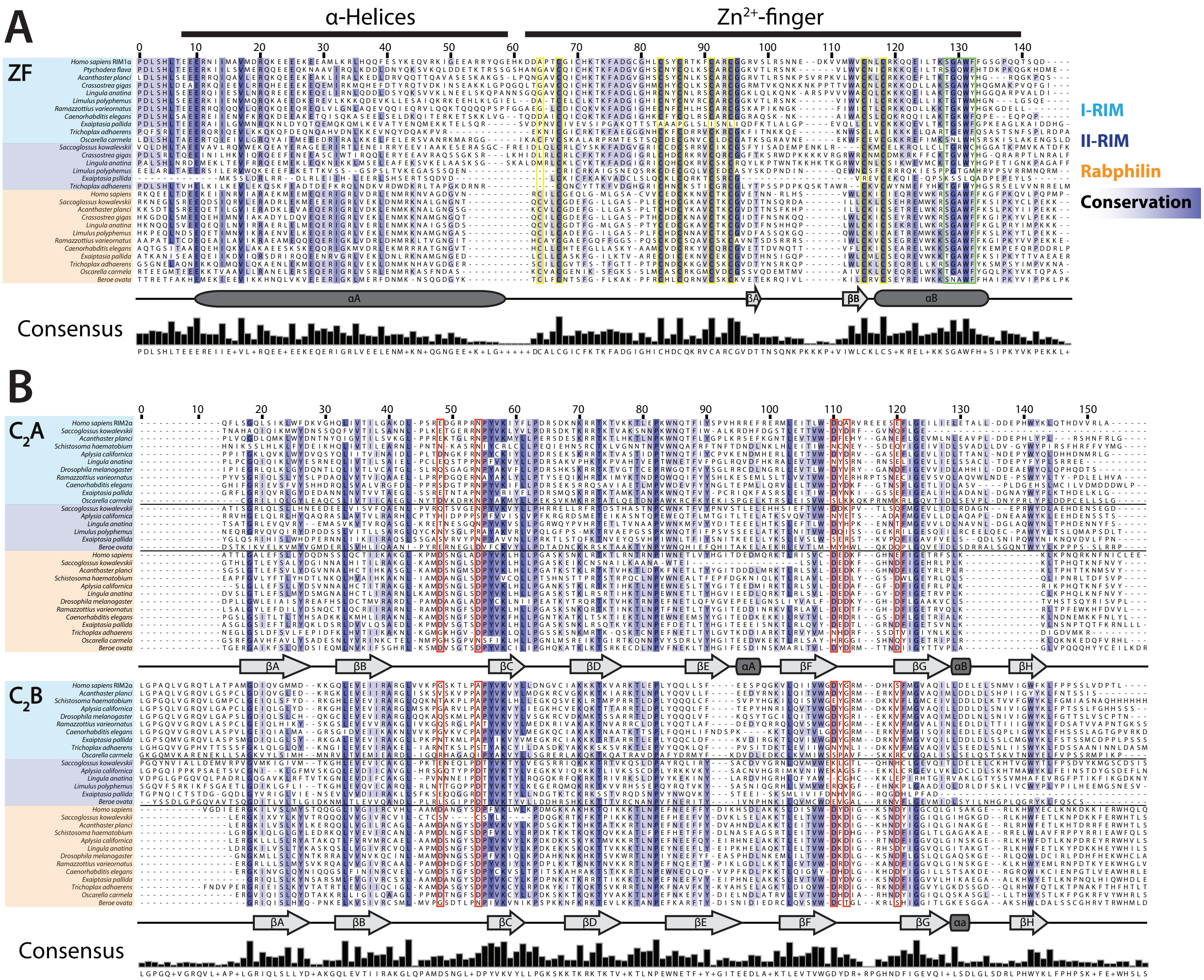


Supplementary Figure 2.

### Figure S3

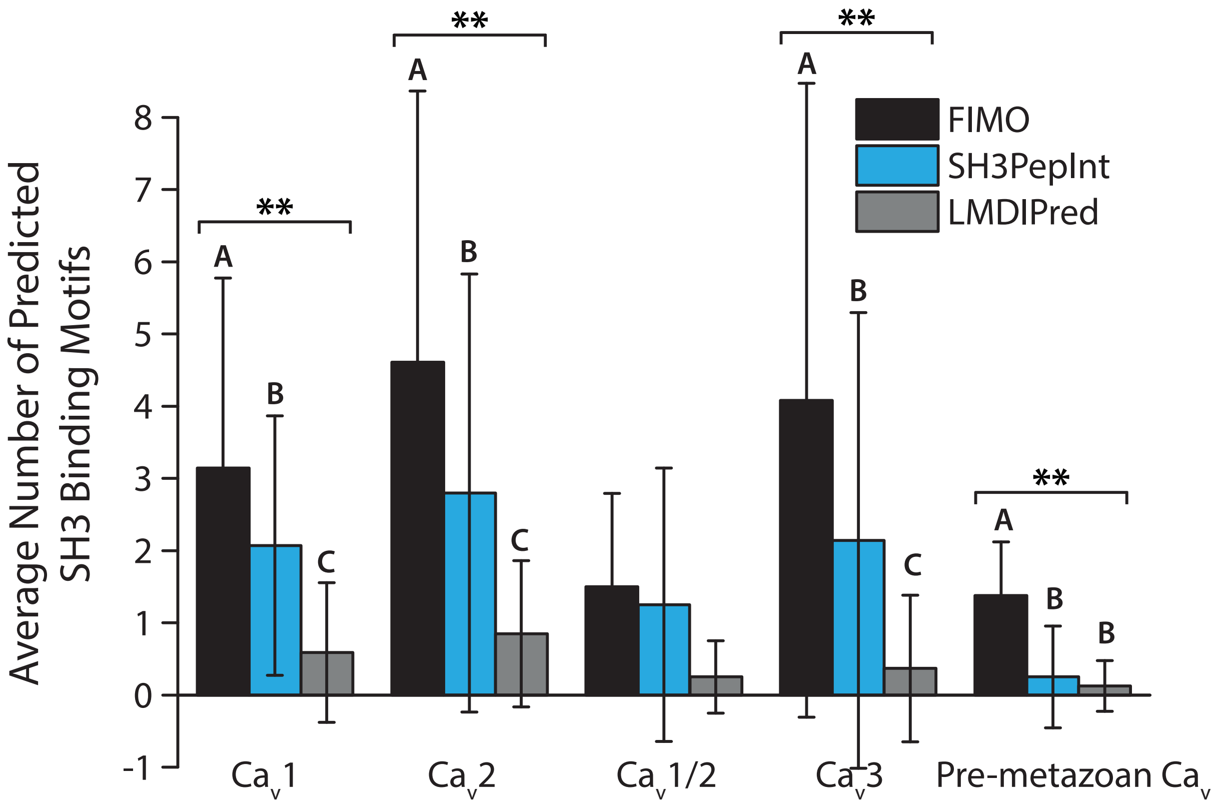


Supplementary Figure 3.

### Figure S4

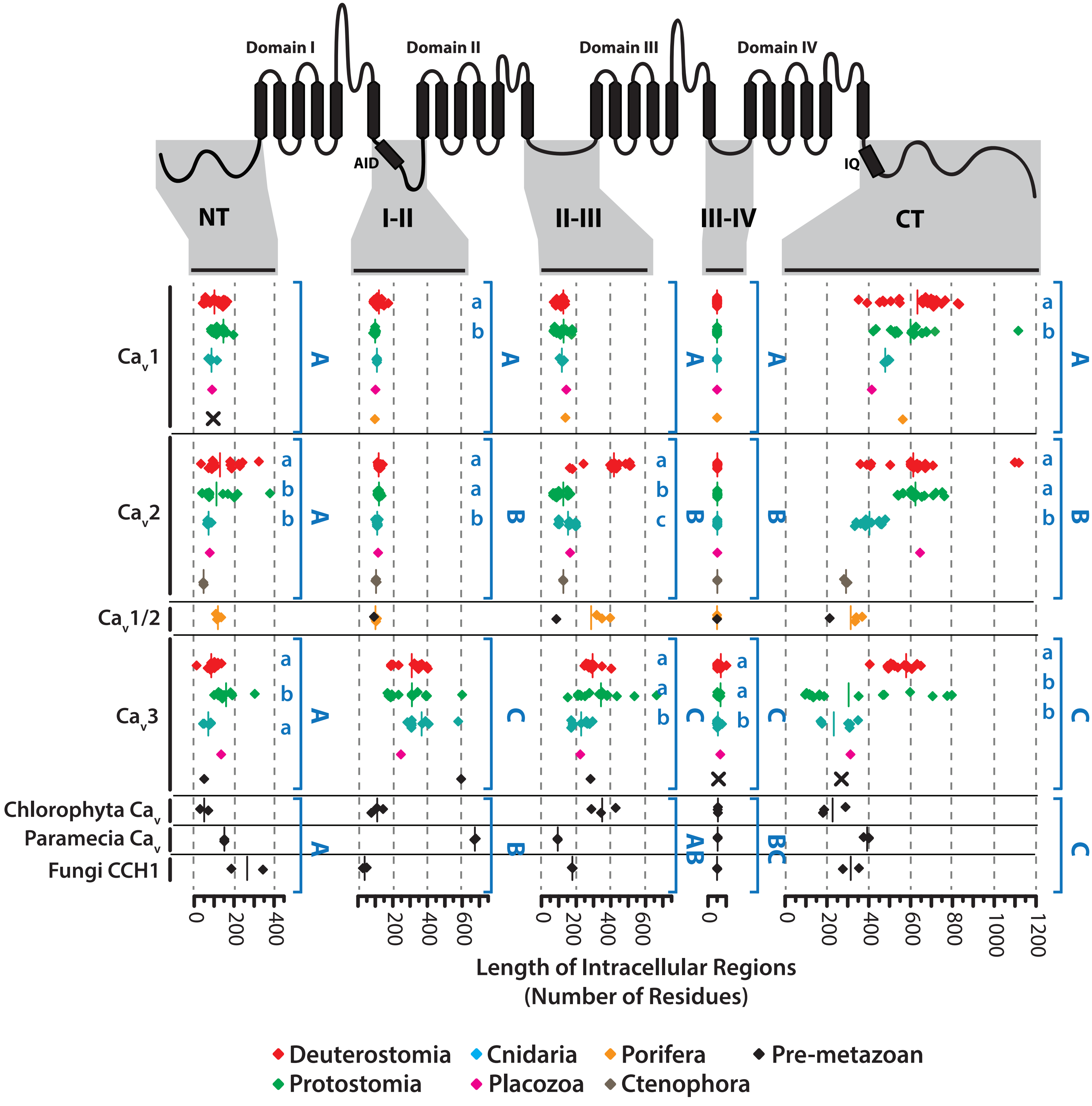


Supplementary Figure 4.
