## Supplementary material for "Early metazoan origin and multiple losses of a novel clade of RIM pre-synaptic calcium channel scaffolding protein homologues": File S1

Zinc-Finger Motif

PDZ Domain

C2a Domain

C2b Domain

Alpha Helices preceding the PDZ domain (Underlined)

>Q86UR5|Homo_sapiens_I-RIM1α

MSSAVGPRGPRPPTVPPPMQELPDLSHLTEEERNIIMAVMDRQKEEEEKEEAMLKCVVRDMAKPAACKTPRNAENQPHQPSPRLHQQFESYKEQVRKIGEEARRYQGEHKDDAPTCGICHKTKFADGCGHLCSYCRTKFCARCGGRVSLRSNNEDKVVMWVCNLCRKQQEILTKSGAWFFGSGPQQTSQDGTLSDTATGAGSEVPREKKARLQERSRSQTPLSTAAASSQDAAPPSAPPDRSKGAEPSQQALGPEQKQASSRSRSEPPRERKKTPGLSEQNGKGALKSERKRVPKTSAQPVEGAVEERERKERRESRRLEKGRSQDYPDTPEKRDEGKAADEEKQRKEEDYQTRYRSDPNLARYPVKPPPEEQQMRMHARVSRARHERRHSDVALPRTEAGAALPEGKAGKRAPAAARASPPDSPRAYSAERTAETRAPGAKQLTNHSPPAPRHGPVPAEAPELKAQEPLRKQSRLDPSSAVLMRKAKREKVETMLRNDSLSSDQSESVRPSPPKPHRSKRGGKKRQMSVSSSEEEGVSTPEYTSCEDVELESESVSEKGDLDYYWLDPATWHSRETSPISSHPVTWQPSKEGDRLIGRVILNKRTTMPKDSGALLGLKVVGGKMTDLGRLGAFITKVKKGSLADVVGHLRAGDEVLEWNGKPLPGATNEEVYNIILESKSEPQVEIIVSRPIGDIPRIPESSHPPLESSSSSFESQKMERPSISVISPTSPGALKDAPQVLPGQLSVKLWYDKVGHQLIVNVLQATDLPARVDGRPRNPYVKMYFLPDRSDKSKRRTKTVKKILEPKWNQTFVYSHVHRRDFRERMLEITVWDQPRVQEEESEFLGEILIELETALLDDEPHWYKLQTHDESSLPLPQPSPFMPRRHIHGESSSKKLQRSQRISDSDISDYEVDDGIGVVPPVGYRSSARESKSTTLTVPEQQRTTHHRSRSVSPHRGNDQGKPRSRLPNVPLQRSLDEIHPTRRSRSPTRHHDASRSPVDHRTRDVDSQYLSEQDSELLMLPRAKRGRSAECLHTTRHLVRHYKTLPPKMPLLQSSSHWNIYSSILPAHTKTKSVTRQDISLHHECFNSTVLRFTDEILVSELQPFLDRARSASTNCLRPDTSLHSPERERGRWSPSLDRRRPPSPRIQIQHASPENDRHSRKSERSSIQKQTRKGTASDAERVLPTCLSRRGHAAPRATDQPVIRGKHPARSRSSEHSSIRTLCSMHHLVPGGSAPPSPLLTRMHRQRSPTQSPPADTSFSSRRGRQLPQVPVRSGSIEQASLVVEERTRQMKMKVHRFKQTTGSGSSQELDREQYSKYNIHKDQYRSCDNVSAKSSDSDVSDVSAISRTSSASRLSSTSFMSEQSERPRGRISSFTPKMQGRRMGTSGRSIMKSTSVSGEMYTLEHNDGSQSDTAVGTVGAGGKKRRSSLSAKVVAIVSRRSRSTSQLSQTESGHKKLKSTIQRSTETGMAAEMRKMVRQPSRESTDGSINSYSSEGNLIFPGVRLGADSQFSDFLDGLGPAQLVGRQTLATPAMGDIQIGMEDKKGQLEVEVIRARSLTQKPGSKSTPAPYVKVYLLENGACIAKKKTRIARKTLDPLYQQSLVFDESPQGKVLQVIVWGDYGRMDHKCFMGVAQILLEELDLSSMVIGWYKLFPPSSLVDPTLTPLTRRASQSSLESSTGPPCIRS

>NP_444500.1|Mus_Musculus_I-RIM1α

MSSAVGPRGPRPPTVPPPMQELPDLSHLTEEERNIIMAVMDRQKEEEEKEEAMLKCVVRDMAKPAACKTPRNAESQPHQPPLNIFRCVCVPRKPSSEEGGPDRNWRLHQQFESYKEQVRKIGEEARRYQGEHKDDAPTCGICHKTKFADGCGHLCSYCRTKFCARCGGRVSLRSNNEDKVVMWVCNLCRKQQEILTKSGAWFFGSGPQQPSQDGTLSDTATGAGSEVPREKKARLQERSRSQTPLSTAAVSSQDTASHGAPLDRNKGAEPSQQALGPEQKQASRSRSEPPRERKKAPGLSEQNGKGGQKSERKRVPKSVVQPGEGTADERERKERRETRRLEKGRSQDYPDRLEKREDGRVAEDEKQRKEEEGVSTPEYTSCEDVELESESVSEKGDLDYWLDPATWHSRETSPISSHPVTWQPSKEGDRLIGRVILNKRTTMPKESGALLGLKVVGGKMTDLGRLGAFITKVKKGSLADVVGHLRAGDEVLEWNGKPLPGATNEEVYNIILESKSEPQVEIIVSRPIGDIPRIPESSHPPLESSSSSFESQKMERPSISVISPTSPGALKDAPQVLPGQLSVKLWYDKVGHQLIVNVLQATDLPPRVDGRPRNPYVKMYFLPDRSDKSKRRTKTVKKLLEPKWNQTFVYSHVHRRDFRERMLEITVWDQPRVQDEESEFLGEILIELETALLDDEPHWYKLQTHDESSLPLPQPSPFMPRRHIHGESSSKKLQRSQRISDSDISDYEVDDGIGVVPPVGYRASARESKATTLTVPEQQRTTHHRSRSVSPHRGDDQGRPRSRLPNVPLQRSLDEIHPTRRSRSPTRHHDASRSLADHRSRHAESQYSSEPDSELLMLPRAKRGRSAECLHMTSELQPSLDRARSASTNCLRPDTSLHSPERERGRWSPSLARRRPASPRIQIQHASPENDRHSRKSERSSIQKQSRKGTASDADRVLPPCLSRRGYAIPRATDQPVIRGKHTTRSRSSEHSSIRTLCSMHHLAPGGSAPPSPLLTRTHRQGSPTQSPPADTSFGSRRGRQLPQVPVRSGSIEQASLVVEERTRQMKMKVHRFKQTTGSGSSQELDHEQYSKYNIHKDQYRSCDNASAKSSDSDVSDVSAISRASSTSRLSSTSFMSEQSERPRGRISSFTPKMQGRRMGTSGRAIIKSTSVSGEIYTLEHNDGSQSDTAVGTVGAGGKKRRSSLSAKVVAIVSRRSRSTSQLSQTESGHKKLKSTIQRSTETGMAAEMRKMVRQPSRESTDGSINSYSSEGNLIFPGVRVGPDSQFSDFLDGLGPAQLVGRQTLATPAMGDIQIGMEDKKGQLEVEVIRARSLTQKPGSKSTPAPYVKVYLLENGACIAKKKTRIARKTLDPLYQQSLVFDESPQGKVLQVIVWGDYGRMDHKCFMGVAQILLEELDLSSMVIGWYKLFPPSSLVDPTLTPLTRRASQSSLESSSGPPCIRS

>Q9JIR4.1|Rattus_norvorvegicus_I-RIM1α

MSSAVGPRGPRPPTVPPPMQELPDLSHLTEEERNIIMAVMDRQKEEEEKEEAMLKCVVRDMAKPAACKTPRNAESQPHQPPLNIFRCVCVPRKPSSEEGGPERDWRLHQQFESYKEQVRKIGEEARRYQGEHKDDAPTCGICHKTKFADGCGHLCSYCRTKFCARCGGRVSLRSNNEDKVVMWVCNLCRKQQEILTKSGAWFFGSGPQQPSQDGTLSDTATGAGSEVPREKKARLQERSRSQTPLSTAAVSSQDTATPGAPLHRNKGAEPSQQALGPEQKQASRSRSEPPRERKKAPGLSEQNGKGGQKSERKRVPKSVVQPGEGIADERERKERRETRRLEKGRSQDYSDRPEKRDNGRVAEDQKQRKEEEYQTRYRSDPNLARYPVKAPPEEQQMRMHARVSRARHERRHSDVALPHTEAAAAAPAEATAGKRAPATARVSPPESPRARAAAAQPPTEHGPPPPRPAPGPAEPPEPRVPEPLRKQGRLDPGSAVLLRKAKREKAESMLRNDSLSSDQSESVRPSPPKPHRPKRGGKRRQMSVSSSEEEGVSTPEYTSCEDVELESESVSEKGDLDYYWLDPATWHSRETSPISSHPVTWQPSKEGDRLIGRVILNKRTTMPKESGALLGLKVVGGKMTDLGRLGAFITKVKKGSLADVVGHLRAGDEVLEWNGKPLPGATNEEVYNIILESKSEPQVEIIVSRPIGDIPRIPESSHPPLESSSSSFESQKMERPSISVISPTSPGALKDAPQVLPGQLSVKLWYDKVGHQLIVNVLQATDLPPRVDGRPRNPYVKMYFLPDRSDKSKRRTKTVKKLLEPKWNQTFVYSHVHRRDFRERMLEITVWDQPRVQDEESEFLGEILIELETALLDDEPHWYKLQTHDESSLPLPQPSPFMPRRHIHGESSSKKLQRSQRISDSDISDYEVDDGIGVVPPVGYRASARESKATTLTVPEQQRTTHHRSRSVSPHRGDDQGRPRSRLPNVPLQRSLDEIHPTRRSRSPTRHHDASRSPADHRSRHVESQYSSEPDSELLMLPRAKRGRSAESLHMTSELQPSLDRARSASTNCLRPDTSLHSPERERHSRKSERCSIQKQSRKGTASDADRVLPPCLSRRGYATPRATDQPVVRGKYPTRSRSSEHSSVRTLCSMHHLAPGGSAPPSPLLLTRTHRQGSPTQSPPADTSFGSRRGRQLPQVPVRSGSIEQASLVVEERTRQMKVKVHRFKQTTGSGSSQELDHEQYSKYNIHKDQYRSCDNASAKSSDSDVSDVSAISRASSTSRLSSTSFMSEQSERPRGRISSFTPKMQGRRMGTSGRAIIKSTSVSGEIYTLERNDGSQSDTAVGTVGAGGKKRRSSLSAKVVAIVSRRSRSTSQLSQTESGHKKLKSTIQRSTETGMAAEMRKMVRQPSRESTDGSINSYSSEGNLIFPGVRVGPDSQFSDFLDGLGPAQLVGRQTLATPAMGDIQIGMEDKKGQLEVEVIRARSLTQKPGSKSTPAPYVKVYLLENGACIAKKKTRIARKTLDPLYQQSLVFDESPQGKVLQVIVWGDYGRMDHKCFMGVAQILLEELDLSSMVIGWYKLFPPSSLVDPTLAPLTRRASQSSLESSSGPPCIRS

>XP_025004772.1|Gallus_gallus_I-RIM1α

MSSSAGPRGPRPPTVPPPMQELPDLSHLTEEERNIIMAVMDRQKEEEEKEEAMLKRLHQQFESYKEQVRKIGEEARRYQGEHKDDAPTCGICHKTKFADGCGHLCSYCRTKFCARCGGRVSLRSNNEDKVYRNTVPTHCGSPVPVRVMWVCNLCRKQQEILTKSGAWFFGSGPQPSPSQDGTLSDTATGASSDAPREKKARLQERSRSQTPLSTAAASSQEISPSSVQSDRRKGAEVSQPAMGLDQKQVSSRSRSEPPRERKKTLVSDQNGKVVKSERKRVPKTSLQKEGPADDRERKERHENRRLEKGKSQDYPDLPEKLEEGKVPDDEKQKKEDEYHTRYRSDPNLARYPVKPHPEEQQMRMHAKVSKARHERRHSDVALPHTEMEEAEVPENNLGKRSQLQGTQDRKSLVETQRSYSIDRTGDVRISVSKQLTNHSPPTPRHSPVPIEHVEYKNHDTFKKQSRLDPSSAILMRKAKREKMETMLRNDSLSSDQSESVRPSPPKPHRAKRGGKKRQMSVSSSEEEGASTPEYTSCEDVEIESESVSEKGDLDYYWLDPATWHSRETSPISSHPVTWQPSKEGDRLIGRVILNKRTTMPKESGALLGLKVVGGKMTELGRLGAFITKVKKGSLADVVGHLRAGDEVLEWNGKPLPGATNEEVYNIILESKSEPQVEIIVSRPIGDIPRIPESSHPPLESSSSSFESQKMERPSISVISPTSPGALRDAPQVLPGQLSVKLWYDKVGHQLIVNVLQATDLPPRVDGRPRNPYVKMYFLPDRSDKSKRRTKTVKKSLEPKWNQTFLYSHVHRRDFRERMLEITVWDQPRVQEEESEFLGEILIELETALLDDEPHWYKLQTHDESSLPLPQPSPFMPRRHVHGGESSSKKLQRSRPISDSDISDYDVDDGIGVVPPVGYRSSTRESRSTTLTVPEQQRTTHHRSRSVSPHRGDDQGRTRSRLPNVPSQRSLDEIHQMRRSRSPTRHHEASRSPADYRSRDMDSQYLSDQESELLMLPRAKRGRSAECLHTISELQPSLDRARSASTNCLRPDTSLHSPERERQSRKVERYSSQKQTRKGSAAETERGLLPCLSRRGLPAPRTTEQPVIRGKHHARSRSSEHSSVRALCSVHHLAPGGSAPPSPLLTRIHQQGSPTQSPPADTSFSSRRGRQLPQVPVRSGSIEQASLVVEERTRQMKMKVHRYNQTSGSGSSQEHEREQYTKYNIQTDQYRSCDNVSAKSSDSDVSDVSAISRTSSASRLSSTSFMSEQSERPRGRISTFTPKMQGRRMGTSGRITKSTSVSGEMYKLEHNDGSQSDTAVGMVGTGGKKRRSSLSAKVVAIVSRRSRSTSQLSQTEAGNKKLKSTIQRSTETGMAAEMRSRMVRQPSRESTDGSINSYSSEGNLIFPGVRLGADSQFSDFLDGLGPAQLVGRQTLATPAMGDIQIGMVDKKGQLEVEVIRARGLTQKPGSKSTPAPYVKVYLLENGACIAKKKTRIARKTLDPLYQQTLVFDESPQGKVLQVIVWGDYGRMDHKCFMGVAQILLEELDLSSVVIGWYKLFPPSSLVDPTLTPLTRRASQSSLESSTGPPCIRS

>Q9UQ26.2|Homo_sapiens_I-RIM2α

MSAPVGPRGRLAPIPAASQPPLQPEMPDLSHLTEEERKIILAVMDRQKKKVKEEHKPQLTQWFPFSGITELVNNVLQPQQKQQNEKEPQTKLHQQFEMYKEQVKKMGEESQQQQEQKGDAPTCGICHKTKFADGCGHNCSYCQTKFCARCGGRVSLRSNKVMWVCNLCRKQQEILTKSGAWFYNSGSNTPQQPDQKVLRGLRNEEAPQEKKPKLHEQTQFQGPSGDLSVPAVEKSRSHGLTRQHSIKNGSGVKHHIASDIASDRKRSPSVSRDQNRRYDQREEREEYSQYATSDTAMPRSPSDYADRRSQHEPQFYEDSDHLSYRDSNRRSHRHSKEYIVDDEDVESRDEYERQRREEEYQSRYRSDPNLARYPVKPQPYEEQMRIHAEVSRARHERRHSDVSLANADLEDSRISMLRMDRPSRQRSISERRAAMENQRSYSMERTREAQGPSSYAQRTTNHSPPTPRRSPLPIDRPDLRRTDSLRKQHHLDPSSAVRKTKREKMETMLRNDSLSSDQSESVRPPPPKPHKSKKGGKMRQISLSSSEEELASTPEYTSCDDVEIESESVSEKGDSQKGKRKTSEQAVLSDSNTRSERQKEMMYFGGHSLEEDLEWSEPQIKDSGVDTCSSTTLNEEHSHSDKHPVTWQPSKDGDRLIGRILLNKRLKDGSVPRDSGAMLGLKVVGGKMTESGRLCAFITKVKKGSLADTVGHLRPGDEVLEWNGRLLQGATFEEVYNIILESKPEPQVELVVSRPIGDIPRIPDSTHAQLESSSSSFESQKMDRPSISVTSPMSPGMLRDVPQFLSGQLSIKLWFDKVGHQLIVTILGAKDLPSREDGRPRNPYVKIYFLPDRSDKNKRRTKTVKKTLEPKWNQTFIYSPVHRREFRERMLEITLWDQARVREEESEFLGEILIELETALLDDEPHWYKLQTHDVSSLPLPHPSPYMPRRQLHGESPTRRLQRSKRISDSEVSDYDCDDGIGVVSDYRHDGRDLQSSTLSVPEQVMSSNHCSPSGSPHRVDVIGRTRSWSPSVPPPQSRNVEQGLRGTRTMTGHYNTISRMDRHRVMDDHYSPDRDRDCEAADRQPYHRSRSTEQRPLLERTTTRSRSTERPDTNLMRSMPSLMTGRSAPPSPALSRSHPRTGSVQTSPSSTPVAGRRGRQLPQLPPKGTLDRKAGGKKLRSTVQRSTETGLAVEMRNWMTRQASRESTDGSMNSYSSEGNLIFPGVRLASDSQFSDFLDGLGPAQLVGRQTLATPAMGDIQVGMMDKKGQLEVEIIRARGLVVKPGSKTLPAPYVKVYLLDNGVCIAKKKTKVARKTLEPLYQQLLSFEESPQGKVLQIIVWGDYGRMDHKSFMGVAQILLDELELSNMVIGWFKLFPPSSLVDPTLAPLTRRASQSSLESSTGPSYSRS

>NP_444501.1| Mus_Musculus_I-RIM2α

MSAPLGPRGRPAPTPAASQPPPQPEMPDLSHLTEEERKIILAVMDRQKKEEEKEQSVLKIKEEHKAQPTQWFPFSGITELVNNVLQPQQKQPNEKEPQTKLHQQFEMYKEQVKKMGEESQQQQEQKGDAPTCGICHKTKFADGCGHNCSYCQTKFCARCGGRVSLRSNKVMWVCNLCRKQQEILTKSGAWFYNSGSNTLQQPDQKVPRGLRNEEAPQEKKAKLHEQPQFQGAPGDLSVPAVEKGRAHGLTRQDTIKNGSGVKHQIASDMPSDRKRSPSVSRDQNRRYEQSEEREDYSQYVPSDGTMPRSPSDYADRRSQREPQFYEEPGHLNYRDSNRRGHRHSKEYIVDDEDVESRDEYERQRREEEYQARYRSDPNLARYPVKPQPYEEQMRIHAEVSRARHERRHSDVSLANAELEDSRISLLRMDRPSRQRSVSERRAAMENQRSYSMERTREAQGQSSYPQRTSNHSPPTPRRSPIPLDRPDMRRADSLRKQHHLDPSSAVRKTKREKMETMLRNDSLSSDQSESVRPPPPRPHKSKKGGKMRQVSLSSSEEELASTPEYTSCDDVELESESVSEKGDSQKGKRKTSEQGVLSDSNTRSERQKKRMYYGGHSLEEDLEWSEPQIKDSGVDTCSSTTLNEEHSHSDKHPVTWQPSKDGDRLIGRILLNKRLKDGSVPRDSGAMLGLKVVGGKMTESGRLCAFITKVKKGSLADTVGHLRPGDEVLEWNGRLLQGATFEEVYNIILESKPEPQVELVVSRPIGDIPRIPDSTHAQLESSSSSFESQKMDRPSISVTSPMSPGMLRDVPQFLSGQLSIKLWFDKVGHQLIVTILGAKDLPSREDGRPRNPYVKIYFLPDRSDKNKRRTKTVKKTLEPKWNQTFIYSPVHRREFRERMLEITLWDQARVREEESEFLGEILIELETALLDDEPHWYKLQTHDVSSLPLPRPSPYLPRRQLHGESPTRRLQRSKRISDSEVSDYDCEDGVGVVSDYRHNGRDLQSSTLSVPEQVMSSNHCSPSGSPHRVDVIGRTRSWSPSAPPPQRNVEQGHRGTRATGHYNTISRMDRHRVMDDHYSSDRDRSHPRTGSVQTSPSSTPGTGRRGRQLPQLPPKGTLERSAMDIEERNRQMKLNKYKQVAGSDPRLEQDYHSKYRSGWDPHRGADTVSTKSSDSDVSDVSAVSRTSSASRFSSTSYMSVQSERPRGNRKISVFTSKMQNRQMGVSGKNLTKSTSISGDMCSLEKNDGSQSDTAVGALGTSGKKRRSSIGAKMVAIVGLSRKSRSASQLSQTEGGGKKLRSTVQRSTETGLAVEMRNWMTRQASRESTDGSMNSYSSEGNLIFPGVRLASDSQFSDFLDGLGPAQLVGRQTLATPAMGDIQVGMMDKKGQLEVEIIRARGLVVKPGSKTLPAPYVKVYLLDNGVCIAKKKTKVARKTLEPLYQQLLSFEESPQGRVLQIIVWGDYGRMDHKSFMGVAQILLDELELSNMVIGWFKLFPPSSLVDPTLAPLTRRASQSSLESSTGPSYSRS

>NP_446397.1| Rattus_norvorvegicus_I-RIM2α

MSAPLGPRGRPAPTPAASQPPPQPEMPDLSHLTEEERKIIQAVMDRQKKEEEKEQSVLKKLHQQFEMYKEQVKKMGEESQQQQEQKGDAPTCGICHKTKFADGCGHNCSYCQTKFCARCGGRVSLRSNKVMWVCNLCRKQQEILTKSGAWFYNSGSNTPQQPDQKALRGLRSEEAPQEKKAKLHEQTQFQGPPGDSSVPAVERGRAHGLTRQDSIKNGSGMKHQIASDMPSDRKRSPSVSRDQNRRYDQSEEREEYSQYVPSDSTMPRSPSDYADRRSQREPQFYEEPDHLNYRDSNRRGHRHSKEYIVDDEDVESRDEYERQRREEEYQARYRSDPNLARYPVKPQPYEEQMRIHAEVSRARHERRHSDVSLANAELEDSRISLLRMDRPSRQRSVSERRAAMENQRSYSMERTREAQGQSSYPQRTTNHSPPTPRRSPIPLDRPELRRADSLRKQHHLDPSSAVRKTKREKMETMLRNDSLSSDQSESVRPPPPRPHKSKKGGKMRQVSLSSSEEELASTPEYTSCDDVEIESESVGEKGDMEYSWLEHASWHSSEASPMSLHPVTWQPSKDGDRLIGRILLNKRLKDGSVPRDSGAMLGLKVVGGKMTESGRLCAFITKVKKGSLADTVGHLRPGDEVLEWNGRLLQGATFEEVYNIILESKPEPQVELVVSRPIGDMPRIPDSTHAQLESSSSSFESQKMDRPSISVTSPMSPGMLRDVPQFLSGQLSSQSLSRRTTPFVPRVQIKLWFDKVGHQLIVTILGAKDLPSREDGRPRNPYVKIYFLPDRSDKNKRRTKTVKKTLEPKWNQTFIYSPVHRREFRERMLEITLWDQARVREEESEFLGEILIELETALLDDEPHWYKLQTHDVSSLPLPHPSPYMPRRQLHGESPTRRLQRSKRISDSEVSDYDCEDGVGVVSDYRHDGRDLQSSTLSVPEQVMSSNHCSPSGSPHRVDVIGRTRSWSPSVPPPQRNVEQGLRGTRATGHYNTISRMDRHRVMDDHYSSERDSHFLTLPRSRHRQTSEHHHRDGRDCEAADRQPYHRSRSTEQRPLLERTTTRSRSSERADTNLMRSMPSLMTGRSAPPSPALSRSHPRTGSVQTSPSSTPVTGRRGRQLPQLPPKGTLERMITEDMDSTRKRNSGAMDIEERNRQMKLNKYKQVAGSDPRLEQDYHSKYRSGWDPHRGADTVSTKSSDSDVSDVSAVSRTSSASRFSSTSYMSVQSERPRGNRKISVFTSKMQSRQMGVSGKSMAKSTSISGDMCSLEKNDGSQSDTAVGALGTSGKKRRSSIGAKMVAIVGLSRKSRSASQLSQTEGGGKKLRSTVQRSTETGLAVEMRNWMTRQASRESTDGSMNSYSSEGNLIFPGVRLASDSQFSDFLDGLGPAQLVGRQTLATPAMGDIQVGMMDKKGQLEVEIIRARGLVVKPGSKTLPAPYVKVYLLDNGVCIAKKKTKVARKTLEPLYQQLLSFEESPQGKVLQIIVWGDYGRMDHKSFMGVAQILLDELELSNMVIGWFKLFPPSSLVDPTLAPLTRRASQSSLESSTGPSYSRS

>XP_015138471.1|Gallus_gallus_I-RIM2α

MSAPAGPRGGPAPPQPLPATQPEMPDLSHLTEEERKIILAVMDRQKKEEEKEQSVLKVKEEQKPQLTQWFPFSGITELVNNVLQPQQKQQNEKEPQTKLHQQFEMYKEQVKKMGEESQQQQEQKGDAPTCGICHKTKFADGCGHNCSYCQTKFCARCGGRVSLRSNKVMWVCNLCRKQQEILTKSGAWFYNSGSNAPQKPDQEGIRGLRNEEAPQEKKAKLQEHLQYQGPPGDISTQVLDKNRSQGLTRQDSIKNGSGVKHQITSDTTSDRKRSPSISREQNRRYDQREERDEYSQYATSDSAMPRSPSDYSDRRSQRGPQLYEEPELGDYRDSNRRSRRRSKEYPVEEEDAQNREEYERQRREEEYQARYRSDPNLARYPVKPQPYEEQMRIHAEVSRARHERRHSDVSLANTELEDSRISMLRMERPSRQRSVSERRAAMENQRSYSMERTREAQGPSPNRQRTTNHSPPTPRRSPIPLERPDMRRSDSLRKQHHLDPNSAVRKTKREKMETMLRNDSLSSDQSESVRPPPPKPHKTKKGGKMRQVSLSSSEEELASTPEYTSCDDVEIESESVSEKGDSQRGKRKTSEQAVVLDSNTLSERQKRMVCFGDQSFEEDLEWSEPQIKDSGVDTCSSTTLNEEHSHSEKHPVTWQPSKDGDRLIGRILLNKRLKDGSVPRDSGAMLGLKVVGGKMTESGRLCAFITKVKKGSLADTVGHLRPGDEVLEWNGRLLQGATFEEVYNIILESKPEPQVELVVSRPIGDIPRIPDSTHAQLESSSSSFESQKMDRPSISVTSPMSPGMLRDVPQFLSGQLSIKLWYDKVGHQLIVTILGAKDLPSREDGRPRNPYVKIYFLPDRSDKNKRRTKTVKKTLEPKWNQTFIYSPVHRREFRERMLEITLWDQARVREEESEFLGEILIELETALLDDEPHWYKLQTHDVSSLPLPHPSPYLPRRQLHGESPTRRLQRSKRISDSEVSDYDCDDGIGVVSDYRHNGRDLQSSTLSVPEQVMSSNHCSRSGSPHRGDSIGRTRSWSPSVPPPQSRNVDQGPRGTRSTAAHYNTLNRMERHRVIDDHYSPDRDRNCEAADRQPYQRSRSTEQRPMLERTNSRSRSTERPDSNLIRSMPSLMTGRSAPPSPALPRSHPRTGSVQTSPSSTPVVGRRGRQLPQLPPKGTLERKAGGKKLRSTVQRSTETGLAVEMRNWMTRQASRESTDGSMNSYSSEGNLIFPGVRLAADSQFSDFLDGLGPAQLVGRQTLATPSMGDIQVGMMDKKGQLEVEIIRARGLVVKPGSKTLPAPYVKVYLLENGVCIAKKKTKVARKTLEPLYQQLLSFEESPQGKVLQIIVWGDYGRMDHKSFMGVAQILLDELDLSNMVIGWFKLFPPSSLVDPTLAPLTRRASQSSLESSTGPSYARS

>Sakowv30037565m|Saccoglossus_kowalevskii_I-RIM

MFFFRQLKDDLENYQKSVKQLDEETKQRGPQQSGAVCQICHKTKFADGVGHSCNYCQLKSCARCGGKVTLRSNKVLWVCNICRKKQEILIKSGKWLGPEKGKSPDSGTETASQVSTKEQEAQKRVNADGSRGSDKENIPKGAHPPPGPRGRELKRQYSMNKDETRRERTLDNQISRDMRGVPRRPDHWRGSHGDINTQREYDERTRSHRRSSSRHPPDDVHYYSDHSDRQYPINSKERIRDRYRHPDRYPDDRGRRLDKDRFPPERGRSYTPDRDRLRYPDQDRYSDSERGYADERVDDRIYTDNERSYAEQHSRRYDRHSSDLEKGRRPVEGMSERPPSAHSDTPNSRYLSERRLSDSVTRSDPDFRSSRDRASMRMQHEQREREQRQREPQEKEPSSTERERHVRPHQSDYSKHSEHSSSMGRPGREFAPSTGHTPRPSHRHSITIIEDVENQKRSPTRQRELESRPLDISPYKQHLDPSSAAVNKTRRTPSRKADTMIRNDSLSSDQSECVRPPPPKPHKHKKGSRKARQLSLSSSDEEIRSTPECTSCEDVEIESESVSEKESEMSAAKKKIVRFGRGEGRSMDEDLEWSEPLIKDSGVDTSSSTTLNEEHSLTAKHPVTWQPDLDNNRLIGHMILKKVTHDGTQPRDSSAILGLKVVGGKMTEIGKKGAFITKVKKGSIADTVGHLRAGDEVLEWNGRDLQGATFDEVYNIVLDSKAEPLVELVVARVMGDIGPGIPTKQTRDIISKRASLRSHSVTITSPGSPTPPNNRTNAHAQIQIKMWYDNSSQQFVVTILSANNLPLKETGERRNPYVKLYVLPDRSDKTKRRTKTVAKSNNPKWNQTFIWALKRDHFDGSTLETTVWDYDRFGVNQFLGE

>pfl_40v0_9_20150316_1g5731.t1|Ptychodera_flava_I-RIM

MAAPAPMPDLSHLTEEERKIILSVMERQQEEEQKNAAVIRQLKDDLDNYQKSVKQLDDETKTRSSGSHANGAVCQICHKTKFADGVGHSCNYCQLKSCARCGGKVTLRSNKVLWVCNICRKKQEILTRSGQWFHTDKPKQGKHDMESPSGSETASLVSTKDSEQKTNEGSRGSDKENIPKGAPTQGPRGRELKRQYSMSKDETRRERTPDSRDPRDIRGDPRHEHWRGSHGDVRTHREYDERTRGHPRSSSRHPDDVHYYSDHSERQYPVSARDREHSRDRYRYPDDRGRHSERDRFPPERGRSYTPDRDRHRYERDRFSDPERGYADAYADHRERGYADPSRRYDRHSAESDKRRHNDGLPPERPPSAHSDLQSPNSRYLAERRLSNDSVASRTDSEYSRGGRERSARMQHEREQREREQRQREKELEMERQIERERQRELEHQRQLEREHERQRQNELEQQRLLEVERQQEIERQRELERKELERQRELEIQRNLEIERQRQLEHQRQVDIEHKRQLELERKREAERERQRANDIPRPSSADRERHNRPHDLDYSKHSEHSQMGRAGRDFDPTQGQSPRPAPRHSITIEEVDNKKRSPTRQRELESRTLEISPYRQHLDPSSAAMRKSRRDTSRKADVMMIRNDSLSSDQSECVRPPPPKPHKHKKGSRKARQFSLSSSEEEIRSTPECTSCEDVEIESESVSEKGEVESWDDRWHSDEMPSQPKLYSSHPVTWQPDLDNNRLIGHMILKKVSRDGTPPRDSSAILGLKVVGGKMTDHGKLSAFITKVKKGSIADTVGHLRAGDEVLNWNGRNLQGATFDEVYDIVLDSKSEPLVELVVARAMGDIGPGIPEKQNRDIISKRASLTSSGYDSGKMKEDEEPSKPRRPSVTITSPGSPTPPGNRPTSLHSMGQIQIKMWYDNTLQQLVVTILSAEHLSHKELGERRNPYVKLYILPDRSDKTKRRTKTIAKSNNPKWNQTFMWDLKRAEFRGSTLEVTMWDFDRFGSNEFLGEVLVDLDRAPLDDEPHWYILSNNHDKNMPIPGGSPKVSRRSKKSHRDGEMKNTPRGQISGNHISDSEQSDLDYDDGIGVVTDCESDVVEMPNTTENATLTDVIESSTVVGVGDGASVSSYGSSCSPPPRNEGNSEHIRQQRNAREQERRQHTAPQQPVSPSPRRRTGTVLAPPREELNDRPQPHTLTVPDQPPRPRTRSPSPGRRGRPPSGEYDLQRSRSPTRRSDIERSRSPTRRSDIERARSPTRRSDIPRADLHRRSQSEIRGNRDLEREYHNAMVGTMSPPDMRSPVTSLPSSPIRHQATSPSRSSPSSTPSTPRKHRQLPQLPVHQTRGDRGK

>XP_011664650.1|Strongylocentrotus_purpuratus_I-RIM

MASPQSMSKPAAALTPGPSPPGPSLPAAKPTPELPDLDLSHLTEEERQQILAVLDRQKNEEEKEAKLIRSLQDDISKAENTVKVISEENKVKNVQQNGAVCQICHKTKFADGVGHSCNYCNLKSCARCGGRVTLRSNKPSPDKSAGRSEVLWVCKLCRKKQEYLTKTGQWYHGGTKTPPSTPGDKLKKLAMSTESLSAVLSSAPSTPGAGGPPSTPSSSTTARAGTRTTTTSPATTTTVTTTSKTVSPASSTSSSLNLRDRERLSDSKAPSLRGQLGKDGTPPSGGQSKGLTRGGSQHAPLTRQLSKDDDSGPGTTRPVHRPHGRDSLRHVDDRDIRGNRGARREPLYGSSGDLNHRDQDRHRDRYPRRDRPGDRDRSISGDRYGPRQAGDRANYAGRGKLPGPEQEPDSRKYPEPDQYRDRRERYPENDRYPPRYHEGDRPGRGFPPEERFYSDDPERDRHNYHKEPVSRTASGRSLDRQRRMYDDPPQDFLPDGYPDSSSVVGKRDSLNRIEGMEPPSGPSEIEPHIERRPRERDRRDHITHTVPSNVREKERIPSDFGPPPVQYGMQRSSNYGSSAGKRGNHGHRHANAGEDIPDALSQDLYGVRTDVTPKQHLDPSSAAGKASAKDSRKTDSMIRNDSLSSDQSECVRPPPPKPHKGKGMRKRRDYSLSSSEEEIRSTPDYTSCDEGEIESGSISERGSGELDITSSPATAGSNNHPGIPPGHGAKSSPTGAVAPPSSSHHIRHHSESEINLVSSPKKIVHFGRGEAGHSLEEDDRQIKDSGIDTCSSTTLNEEHNLGAKYPVNWQPDMDNNVLIGQMVLRKNQSEACTHQDSSAILGLKVIGGKMKEAGRLGAFISKVKRGSIADTVGHLRAGDEVLSWNGHNLQGKDLNEVYNIIFESKSEPQVELVVARPMGSEIPPSSRDKIAQFQSPRPDSIMKRSSQSELQCSSGYESNKPKEEEEPMPIVSRVKRPSVTITSPGSPGLPRRNTSPTALGEIQIKLWYESNNYLINMTVLTMHGAQPRDTGDLRNPYVKVFLLPDRTEETRRRTKTIQNTLSPKWNQTFTFGPIKRADFRGRDLDVTVWDFEQYGAKEFLGQMMLPLEGAPFDGQSHWYSLYPHGSSRHQPSSPPQQPPPPPQPKKQLPLTTNNNGNPRMHDMIDKREMIGKHEMIGKHDMIGKHDMIGKRMAQENGNMREKLSPSSVGRITDSDVSDFDDGLTGLSGGTTTGAGDVASISSFGSSCSPPPVSDQDDRPSSQDTDFPRSQAPHRRAGIVTPPTQEDLTPNEEPMPNKQRRNSQSTLAVPERASRRPRSPSPIRKERGRTRHSEPAIEYDRQDGAQDLQQDPDREQDRDRNRDRDRDRDRDRDRPLDRDRDRDRNHTRDRGRERERDRDRHRDPDRDLDHDLDRDRDRDMYSYRGTEVDNIRTSPNSISSRGNDQRESPHRLDRSSPRSSPPRRGEREHRSRTLSPPNVYEYSEPRRSRSPSRRSVEGSQSDSNQRRSRSEVRGEMEYDRQYRGRNMLPPPTSGLDSSSLPNSPIHGGRNSPSSTPSTPRKQQRRLPQIPPSSKADKGTADIEERARQIKLKMTQYKQAASANTLSPHGGSGGIGGSGGGGGSLERGSHMGSPGHDMTNPFQHSSHRRKQSPDNISIKSSDSNLSSTSDVSAVTAASTASAFSTQSERPRPTRKFSAFTAKMQEHHPAPTRKPLNRSTSSTDMYMYEKNEGSISDSAAEQGSVEGKKRRGSIAKVASLVGLSKKSNSTSNLAGKKPRASIQRSEEVLPMEMRDRLQKQASRESNDGSICSFGSDSSSSFMLPANFRFGGESQFNDFLDGLGPSQLVGRQALGSPWMGDIQLSLEDKRGNLEVEVIRARSLVAKPGVKVPPAPYVKVYLMDGKHCVAKLKTKIARKTLDPLYQQTLKFIEDDHSNKVLQITVWGDYGRLERKIFMGVAQILLSELDLSHPVMGWYKLFNTSSVADIHQRGSISSLEGSMVSLASTK

>XP_022091188.1|Acanthaster_planci_I-RIM

MMATPQSKAPATKTVQPLATKPGAAAPAPPGPAAQPPAGPVVPELDLSHLSEEERRQIQAVLERQKQEEEKEAKLIRALKEDLDRVQQTTQAVASESKAKGSLPQNGAVCQICHKTKFADGVGHSCSYCNLKSCARCGGRVTLRSNKPPEKAPSSVRPKVLWVCKLCRKKQEFLTKTGQWYHGRQGKPQSLDLSAVKAIDGGSKPQTPQPAKDGPQQQQQQIIKPQMEQILHSAKPSNNNNKENLGAGQPPSRPRSTTPHSQLTRQLSKNEDETSSPLPQTNSKPVHRTHSRELAGKRPGAHDERTHNARADPSYDPSSDVHYQDRRRESRYSMRDRTPDRDRSTSSERTTMTERHGRDPAKYSGREREHFMEQDPRRFKNQDHIRDHGRYPDHERERDRERFNHGYESPDRHGYPEERYYAGDHDRGRHYTKEPLVRQVSGRTLDRQRRMYADDPPSPHYPHDRYPEHTSSDRERRDMDQREIDRRETEHRELSDRREMDRRDMPERREGVGAQMDMREFERHPSERRTRARRERDRSPGESKRSSPRKDEERFGEEYRRGAGPPYPQEGVSQKRPNSSPRHSITVEDVDAQRERLLLAESYGAPGRAPELGPKQHLDPSSAATKTSKQESLSRNSKAADSMIRADSLSSDQSECVRPPPPKPHKTKGSRKRRQFSLSSSEEEIRSTPEYTSCEEGEIESESVSERGSGELEIGTGGGNNNSSSKDKGMHGGRGEPVAADDHHHSESELALSAAKKKIVRFGQGEGRSMEEDPEWSEPQIKDSGVDTGSSTTLNEEHSLASKQPVTWTPDLDNNRLIGHMILKKSQGDSSTPQDSSAILGLKVKGGKVKEAGKLGAFISRVKRGSIADTVGHLRAGDEVLSWNGHNLQGKDVNEVYTVIFESKSEPQVELKVARPMGSEMDQTTREKLAQQFRSQRPDSITKRSSQSVCSCACAGSSGYESTTKHKEEEEIPPQVRHKRPSVTITSPGSPGMTRRNGSPLVQGDLQMKLWYDNTNYQIGVTILSVKGLPPREKTGDLRNPYVKMYLLPERSDESKRRTKTLHKTLNPKWNQTFLYGPLKRSEFKGNTLEVTVWDFEHYGANEFLGEVMLNLEAVPLDDEPHLYPLRSHNKQVPLPLVSPVPARHQMKHHSHRNSDTGSREQLSSPASGGRITDSEFSDWDDGIGVVTGASVVGVGDGASSLGERRATPPMNEGSSVVGIGDGASVSSLGSSCSPPPASEDDRPPNEQEYPRSPTAQRRSGIVIPPQQDELSSHTRERRGSASTLAVPERAPRSRPRSPSPTRRERMSTEPVDVYRKHQEATSRFAHYQDDTDGQSRQSPTAMRSQRERDRGGWEYERSRDLRDRDLRDQDLREHELRELDLRERDLRSRESRERELRDLPPHELDLPSRGLSPREPHDQELLDREGRDPRDHRTHRALSPPTRYDYERMSGRRSRSPSRRADLQHGDPHRRTQSENRGEMDYERQYRSHARTITSNIDASSLPNSPVHGGRGSPVSTPSTPRKHRQLPQVPAHLKSDKGTAEIEERARQMKLKMKINQYKQAAATANTLSPHGAVASTSAGGTPILDAPPHPRRKQSPDNISIKSSDSNVSSTSDVSAITQASTASAFSTQSERRPTRKLSAFTAKMQESAPVKKPLNRSNSSADMYTYDRNEGSISDSAADTNIQEGKKRRASIGYKMASLVGLSKKSNSTSNLAGKKPRSSIVRSEEVGLAAEMRNRLQKQASRESTDGSVCSFSSDSSSQLWLPGNFRLGPEGQFDGFLEGLGPAQLVGRQVLGSPCLGDIQVGLEDKRGKLEVEIIRARGLISKSVSKVPPAPYVKVYLMDGKRCVGKKKTKIARKTLNPLYQQVLLFEEDYRNKILQITVWADYGRMERKVFMGVAQIMLDDLDLSHPIIGYYKLFNTSSISELHGHRASISSLEGSMTSLASAK

>XP_005106594.2|Aplysia_californica_I-RIM

MLDEGAVCEICHKTKFADGVGHSCHFCHKKSCARCGGRVQGKGLNKEKPSMIWACNLCKKKQDLLAKTGAWYHGGMARPVALDVGDTASGSDAASTKTDVSPSTEKRAKMMEKGQHDSSQGSEKENFDRQRSSFSRAGSLQGKELKRQFSMSDAVNRGHDSGLGGSTTSSSSQQGSLNASQTSGGTSVGSLGEQEKLRDHGQILDRGRGKDRGQAKHRFHSESRISDTDRRYANETQHHGERDRHNKVDRWDGGGGGGGGGGGREGGGGREGGGGRHEERGSGSSSGHKGSHSEDRKGDHGSFRGDPDPRDGDRKSERHRDGSQHDRRHDRSGGQLDERDQMDRRREGKDRRKDGRVEAADRERGDRYHRQSPVNGRRDYGSRERLNDMQQQQQQQQQQQQQQQQQQPQVVDVSLQEGEGSRLERRGGSSRHHRERGERSVKDQRRSPSRERERDRGGGSGSGGGGGAGSGSGGGGGGGGGVVVGVGGGKGEPPDVHEGQFSRHRILIDHASDTQAEDPRDVVDNDPGDKKQHLDPSSAGGRSRNNRKKLESMLRNDSLSSDPSDCVRPPPPKPHKHRRGKKQRQQSISSSEDEIQSTPECSSCEEVDLESESVSEKGELESVTGSLRSRGSHDGYGWRGSKPDCLAKPTLYDTGRGTSVETESGSSEPPPLKENGNDTGLSSSSNLTLNTSSVDDFVSKRQLQSIKHPVTWQSSADGTKWIGHMILKKTVLEGSGEKRDSSAILGLKVIGGKVSDSGKLGAFITKVKKGSIADTVGHLRPGDEVVEWNGRSLQGATFDEVYDIILESKQEPQVELIVHRSAKHGETPPGVQSSFTDHARDYGLKEAIPDVQRSSRHPVPTPQPRSRLRSRPLSPPITGKLQVKLWYDVQSYQLIITVINAIDLPLTDNGKFRNPYCKIYLLPDRSEKSKRRTKTVASTIDPTWNQTFIYCPVKENDMHERLLETTVWDYDRVGASEFLGEVLIDLTTANLNDEPYWYQLDHHDNASIPLPASSPRSKTSQDPYDLRKDHLSPPISTRGLSDSDMSELDFDDSIGVVPVVARDDLGSRQRHPSAADKLEVPEVGGSGQRSRSPSREGSAKNSRPRSRSPGNIRVTDASRSLSPPELRAVPSGTSQRSFPPASSRSLGGTPTSTPSPKKRQLPAIPLDAQKASRDRVTQDLEERARIMKMKMKLAQQGQGGGIPHPESDSRLTRRRDRHDGSDRMHDRSISHDRVERMRERAMERNYERGRDHDRGHRTRRNKEFSPDMSDDVASDVSETSDMSEVSKVSTISVRSTQSERPRRKLSEFASKMESRTTMPQRRQQQQQQAARSSSNESQGFDQNDGSVSDSAVTASVTEGRKRRPSIGHKMAALVGLNRRSSSASQLAGAGLKKSGRGGGKKRSSFQRSEEIGPVDLRGGMSKQASKDSTDGSIGSISSDSSSVLWLPTGMRLGPEGQFGEFVDGLGPSQLVGRQVLGAPCLGEIQLGLYDRKGHLEVEVIRARGLMSKPGARVLPAPYVKVYLIEGKHCVEKQKTTTARRTLDPLYQQQLFFTEPYHNRILQVTVWGDYGRMDKKVFMGVAQILLDDLDLSNIVIGWYKLFPTSSLFGHHSSSTGLSQSGLGSSTSLETLMSSRT

> XP_019921964.1|Crassostrea_gigas_I-RIM

MMSTTSRRPSLPPMPDLSHLDEAERKQIEEVLRRQHEEEEKEQEMIRKMQHEFDTYRQTVDKINSEAKKLGPQGQLDTGAVCQICHKTKFADGVGHMCHYCQLKSCARCGGRMTVKQNKKMEDTPPVPVNKPPTTVWACNLCRKKQDLLAKSGAWYHQGMAKPVQFGIDAQSGSDTGSIKADISPQNEKRPKLYEKKLDGTPSGSDKENMNSGPRGPPHPKEGIRRQSSLDTGSTQHSRGRHNEQYALAPEDGDKLREHGPERGRNRDRSPASRRHYSETRLSETDRRFAQEFRHGERLGRGEGPSRSRDPSRERGGPRDDMKRGRDSHIDKRVRDGPRDPRDERHRDLHRSEHDMVYKRSKDPEHYRQPPVDSVKGHDYADKDTIRGSRERVDMLDGDKRRTRDSSLSKVNNMQPPSAGKMDRGSSGRLSSRQERSPRDASSSRRSPTTMDRDKSKEGLDVYHGETSRSPSVSSRHKIRNASPTVKDESGRPIDSPYGYSHSVSSHLDPSTAAGKNLSKNNNNNNRKKMENLRADSLSSDPSDCARPPPPKPHKHRKGNKKQRQHSLSSSDDEIRSTPENSSYEGEEESESIVSEKGLSIVGGRETPLGQHGAYITKVKKGSIADTVGHLRPGDEVLEFNGRCLQRATFEEVRDIIMESKQEPQVELIVNRNASSGEVPPGLQQTDHNTRRSPLDHDKRGKRATDITNGHTRRVNLSETAPNFTSPRSRSLTPKSMGKIQVKLWYNMKEQELIVTLISATNLPPTTKGQFRHPYCKVYLLPDRSEKSRRRTKTIMNSNEPTWNQTFMYYGVTSEDLQNLLLEITVWDYDRMGPNEFMGEVLLDLHSANLEDDPFWYPLSQHDENSIPLPQSSPRSKTQQEAYYGRKDHLSPPVSSRGHSDSDISEFDVDERFARSVPAEVRRGLDKTSSCSSYGSSAGSDDLSDPTDYKYSRTRTRPIPILKSQFRPEISPLAREDPLARLREAIVTSREEHEHSHRHPKTAVEKLSVPGEQSSLPRTASPKTPDMHPRNRSRTPSSSYHRVPENASRSLSPPDLSSLMANSPSNTPSPKKRQLPAIPTDAQRESRDRVVKDLEERARQIKSRMRPGSVTPNLSDSEAMRGRHRSSYDRSISRERGYPLYDRDPSRGRRKDDYDIPSDADSDISEISKVSTISVRSTQSEKPHRKFSQFKEKINTKSTIPRSQQPSVRTPTGDSQGFQKQSSSGSMSDSEVGSCVTEGRRRRPSLGHKIGSLIGLQRRSSSASQLTDGKKKSSIQRSEEVLSGRGLVKQPSKDSTDGSIGSISSDSGSVLWLNPAGMRLGPEGQFGDFIEGLGPAQLVGRQVLGSACLGEIQLSLYDRKHHLEVEVIRARGLIPKAGAKILPAPYVKVYLIDGKHCVEKQKTTVARRTLDPLYQQQLAFTEDYRGKILQVTVWGDYGRMDRKVFMGVAQILLDHLDLSNIVIGWYKLFTSSSLAGHHSSSSTIGHSRKGSTTSLDSGYNNNSPRT

>XP_021377513.1|Mizuhopecten_yessoensis_I-RIM

MSTNRPKPSVRPSLPPMPDLTHLTEDERKIIEDVLKRQHDEEEKEQDMIRQMQTDLDTFKVSVEKVNEEAKKIGPPNQDTGAICQICHKTKFADGVGHRCNYCHLRSCARCGGRMALKPNKKPPPGAKTETPVMVWACNMCKKKQDILAKTGQWYHGGMAKPVQLDNLPQGEGPAGQNISPPNEKRQKIHGEGPDKENSEDKKGHKIVRTGSLQGRELKRQFSLDAPKNHSDRSGSQNLTPDMADKEKGSRDRGHLPSERGRTRDRSPAANRHHSESRLSETDRRFNAQEWRNIEQESRYRRGDTGREPRSRDQSRDPDHMEKGKYTDDKRRDPRDAHVDRLMDDRDRKREISRERSRGSREELGRSTRDGREDRSNRDPRDWEEGSRRRDGPGDKRKDRQTAPVDNLNIYDFADKDNKGSREKLDSLENDKRRVRDPHSKPGGSPHSRDNTQNSRHLNNRHPRDRLGSGSKDRRSPTTGDTRERRKEPEIYNSHVRSPSLNRHNIRIERVGDSPTVMEEQVDHDKYHDKKQHLDPSTAATGRNSRSRKLDISMSRNDSLSSDPSDCARPPPPKPHKHRRGKKLQQHSISSSDDEIRSTPECSSCEEPDLEIESVISEKGEFEMGEERWRKDEILAAKIKKFLSHPVTWQTSADGVKRIGHMILKKTVLEGTGKDDSGAILGLKIVGGQGTEQGQFGAFITKVKKGSIADTVGHLRPGDEVIEWNGRGLQRATFEEVKDIILESKQEPQVELIVHRSVSSGEIPPGVQPTVDHSLGRDGHSNRDYTHNREHRDHRDRDHRDRDYRHSKDTLSTGHHSSRPSVMVTSPGSSVEPRSRASSPPISGKIQVKIWYNSKGNELIITIISALDLPLTDKSKFRNPYCKLYLLPDRRYLRNFAERHREVAFEPCDKSKRRTKTLSNTNDPTWNQTFMFYPITESDFRTRVLEITVWDYDRIGASEFLGEVLIDIHSTNLNDEPLWYQLGHHDETSIPLPQSSPRTKSSQDPYDLRKALSPPVSLRGLSDSDMSELDFDDSVGVIPAGIFPDCVRDDRTERRRLSGKNNEMLSVPRDQAPQTHQSSRSPHPPEKGHSSRTRSRSPSSYHRVPESASRSLSPPELSKSSPKRRKLPAIPLEAQMAGRDKATQDLEKRAQKLTMKMKMSSSGANMSDSESYRSRRDYHRGDFRERSRDRGMSGDRGEMMYERSGHGSNDFDRPHHHRRSRRNPEFNPDIGDDMGSDASETSDMSEVSKVSTLSVRSTQSDHPRRTFSEFSQRMGIRTTVPRRQITPSSSSDSRDSYEKTDGSMSDSALSASVTEGRKRRPSLGHKMATFVGLPWKSNSASQLAEAGKKKNSFQRSVEVGHGLEAKYGRMVKQASKDSTDGSIGSISSDSGSVVWQPPGGNPALGPEGQFGEFIEGLGPAQLVGRQVLGSPCLGEIQLGLYDRKGHLEVEVIRARGLMAKSGAKILPAPYVKVYLIDGKTCVEKQKTVVARRTLDPLYQQQLVFMEQYGGKILQVTVWGDYGRMDRKVFMGVAQILLDHLDLTNIVIGWYKLFTSNSLVGHHSSTSTLGHSRKGSTTSLDSGYNNSSPRT

>XP_014789722.1|Octopus_ bimaculoides_I-RIM

MSTNRRPSLPPMPDISHLTEDERRIIEDVLKRQREEEEREQQMIQQMQDEFDSYQQAVKKLSEETKQVTPQDEGAVCQICHKTKFADGVGHSCHYCHLKSCARCGGRITMRGNKPVPVSGKEVKDLSGQNVTVIWSCNLCRKKQEILAKTGAWYHGGMARPVALDVGDTASGSETASTKTDVSPPNEKKAKILAKQQTDNGASSEKENIDKKSVSRSGSLQNRGLKRQYSLSEVQGNKQNDLKDRPTDNTSGTIVSEKEKLREKGQLQDRGRDVERGPSARGSNVEGSKHHDGKVHSREESARRYDKRSDRDKSHEKDERNREKHQGRFRDGSTDRFNDEKRVDHSGRETGDHYHERDQRKPRDSSREPSRDRLKGSKESLSSRSHRTDRADRSERVADRNERVDRTAEPHEDWMADRHREGDRRRGRINLESEHYQDPVLNNQDYHESDTRPTREPRQKDGYEEEQWKNVDSSSGHHGRRNATRHQRDHQRADQNKDHPQEAVRGHRDSLNKRDHIREHPAHSRDHRGELQRELPREQHPRERKDHITDSKDSRSPSRDRERTRGSKGDKYSSYITDQSSIQVVSTGHHDREASEIEHRGETDRYFDRKQHLDPNSAANKSSRTNRRKVESMLRNDSLSSDPSDCVRPPPPKPHKHKKGKKQRQQSISSSDDEIRSTPECSSCEEPDMESESVSEKAAFYWSQKGDFEALTSDDHWKKDEILAAKIKKFLSHPVTWQRSANGRYYIGHMILKKTVLEGTGSRDSSAILVCLERPRQKRRSKEWEGSFKASLKVVGGRMTEKGQLGAFITKVKKGSIADTVGHLRAGDEVVEWNGRSLREATFEEVYDIIFETKHEPQVELIVHRNAKTGEIPPGLQQIEPSGRDYSSKDGLMTRPSVMVTSPGSPSASGKRVHSPQITGKIQIRMWYDDSRCELSICIVSALDLLPSADGKLRNPYCKLYLLPDKSEKTKRRTKTIPNTNDPTWNQTFVYVAGKESHFKQRLLEITVWDYDRLGASEFLGEVIIDLSRANFKDDPLWYPLRQQDESNIQLPTSSSRTKSSHDPGDIQKVHLSPPVGSRGFSDSDMSEFDYDDSFQTGAALLERMGGSSISSLSSAISAAHLNEDYSEIMDGRCKSRRDRGVGSPTTRRRSGTVIARDELDDHRKHNHPEGGSLVVPEKGTRQRSRSPVEDMNHGSSRTTRSPARHQVPETLSRSLSPHSSRQNRGMSPIRVLPTTSRSAGGTPSSTPSPKKRQLPSIPIEAQIASRDRVTQDLEERARQMKLKMKINAQHRHGTANHGAPSSDSEIAHHHHRSRLRDGDYRHDRYDIHGYHSRGRDLERGHRSRRRKEFSPDVSDDNLPSDASETSDMSEVSKVSTISMRSTQSERPHRKFSEFTSKMESRTTIPKRQIIRSPSSESSSFERNDGSISDSAVSSSVIEGRKRRPSLGHKVAAFVGLSRRSSSAVQLAETNKKKGSVIRSEEVGGGLEMRSRLAKQSSRDSTDGSMGSISSIESSVMWLPPSHYGDFVEGLGPSQLVGRQVLGAPCLGEIQLGLYDRKGHLEVEVIRARGLMAKPGAKVLPAPYVKVYLIQAKHTVEKQKTTIARRTLDPLYQQQLVFNESYQGRVLQVTVWGDYSRTDRKVFMGVAQIVLDELDLSNIVIGWYKLFPTSSLVGHHSGPSSASSPHTRKSSASSLETLSTATSWTQ

>XP_024347539.1|Echinococcus_ granulosus_I-RIM

MPSNCALLLAAATVAAGAGSLSASLSLPIALTLALARLCVRATLSLIYFPRLRHVSPLGSCSLAFADAAASYLALYISQALLSIPICQVVHRGSKVFQRLPALTDASPSSSSTSTTNTASASNAQLANSIGGPVARAKDEFFGGVTRRGLCAERPLPSVHHNRTSGDLFGAVSNFFTRTGGGSGNFGDVCEICHVTKFTGNAGHNCVQCNLKTCVRCGGWFGTQSSWMCKNCFSNLSSPSAMTHVAKDAKTPLPSGLTSYRRESFSSPLNTTESTLNPRMIPKIPPTSTPATASTSSSLSASNVAPVERSLSSYAGAPHLYTTPLLKSSSWLTEVRNRSLSEEDRVWSHYQPTPSSHTYLLHSHQSPKNYYQHPPVPPHSHPHKQQHPQRQQSYSSIPPRYLQQHSSHYPAAATATAQHPYNAPHFRHSTGPPVDRAYRRPFGEAAWIRPVMPPPLRSERHTFRSMEGGGVEGGGVSTTSCGSTEDSPHGGFRAGYLRRSGGGQMDPGSVSEPCKSEAYSRGDEKLKPSDSDWLMCQNFTLLESRISLSCGRSAFWSPSQDGESLIGQVYLRKCRASGEIGLLSSFGLKLVIGKRVPSDMVGTFVSSVKEGSVADVMGELQTGDEILEWNGHNLRGLHQEEVSEILALSRNADEVWLTVQRVIDDYCEESDGANSTSLSRPGEDARFQEEEEEEEYGDNFGPGDADPHLQASKILKLLYDEEKEALIVSVLAARDLPPQLNTQTWLCSSFCQITILPETNKYEPRCTRVARNTNDPVWIHNVTFENFVINSKELEVAVFDYLRGRTAFIGEVLINLQVADLSGRAYWYPIPPGWDNTNQDFSSQVQSESGTFEADEWSGEASERVTAQVSHRPVSGGSGGRRALRNISPRSVPSQLSASASAVADRQKRQQPGGQRTSRRESRGPAQPGPPKTHAADRGPRFSLSEDDQSFLSDNSEASGFSSFSKLSLQSNRLRQRQHQRLNASKLGSAARRISRSRADCYSPSQDLLAETGEEEEAELEEAAAMEDEGKQSRGREERPPDDQPKSSSTEEKSVSSSTAAAASTVAVSSGGFTETTTMTTAGATKATAAAATPSTARKRRSSIGHRFSNVLGISKKNTATSGTDKKTKASFQRSEEVLPAYIQTNEKGTVSGGYKAPGVVAEGAGFGGIAASSVQSSVGGGFTGQAGHHGSGLQHTLSSVLHSSSSCSSSAAATAARKDQLALEMGEVHVGEFVEGLGPGQLVGRQVLGMPCLGEIQLSFFDRKGHLEVEVIRARGLQQKNTSKPLPTPYVKLHLLEGKQSVEKMRTTAPARRTLDPLFQQQFSFSNSYKDKILQAMLHSGLAINDSPHTLSQVSDPYLGGDGTAPLSDRSRETVSVWGEYGRMDKKVFLGMCEIVLDDLNLRSIVFGWYKLFGMIAATAQHHKQVRRHTSFGGTSTATTTTSGGTSQTSSSGKRHASKNRQPRGKS

>XP_012796083.1|Schistosoma_haematobium_I-RIM

IPVGNQSIRDPIITSRSTHITNPSEKSPINIATVNGKNVTKLSSSSNTNTSLPFGNTVVNLMAEAAHAVTGGSTGTTGGFSAGHSVNVCESCHKTKFADVSGNPCIYCKRRTCTRCGGHMEIKPNIVHWVCKQCAQNLTSDRQIVTTSGGNLPHHMESSINSSITGRLGESIKDFISKPKSSDVKRQLPQLNVKNQNVPYISDGSRGQTIQHRNSGRTLAMNRSNSLDADKTAYISPDHSVYHVDQQRPFGDKFKKYDQIPGEYGHSARSRHTSGAYYPSDEITSYREVNNRCPPNDPRLHGNTYLDSTYPGDSFPSDPSDSPCQSGGGDSSTKYGLSYGKGPAHEFGRHQEIRYRSDQNFPSQSPYYKQNYSSSPFSAQRRNKPNWETYRYGAVPPRYRDQLRHHQLHHLSFSSSDGEDFSVVDDGSVLYQEPRRRLLPTNFERKNSPYSQGTVTDDDDPASQFYISKRSTTPYYDRQLKPTVWDQDKALSLCDEPPVRYAYSDEAQPVYHRQLPENRSMGYQKDVQYNHKDGTVVTKRLPPNISSKTDPHFGEPGLGLKLAGGRLDKSGHLSAFVTQIKPGSPADIIGHLKSGDEILEWNGQPLRGLSADDVSQIIYQSRDELQVELLAQRELNEPANTPTDDFVLMVPNRSDCLNPNEMDSVEHNIKSSLHLKLFYDEIKHQLIVTILEARNLPNTNKASRMPCNIYCRLCLLPEQIEHNSRCTKLVTCSGNNTNPVWNQAFRFTNFTDNDLNKLHLEITIWNCNEYSDEQYKIGEIVIDLSVADLLDVGYWYPIRSFHRFFSATELSNMNPDVNSAPYDLQPCTTTQQQWSPKSEYRHSDIPDDNTRKLKSIRDNYSDIDEQRFSASIRQNRKFTDHEYYRDGTEVHHRGSRLHPSAQNFEYGGLEENAIQSDASEFSDLSEISRMSLHTSQSDRQRMESIQTSRNGSQQFETPTQSYNERHSRNVQKPDRLDENFLPSVEQSTRARRDSNRIEKDIIVPTEDEEQHELIAKDNRNFESGSSPRITSADDYGGGQVSRSLIENKVSNFEIAQNVHRDQVDGRKRRPNIGQKFSNVLGRSNKSSSTSTLDKKSRTSFQRSEEVLPFGGYGAIFEGQEQEGSFSHGRLSMMHCQSSISSGKSDGRNSRYGTNSDTTDQESVYMGRNESHVGEFVEGLGPGQLVGRQVLGMPCLGEIQLSFYDRKGHLEVEVIRARGLQQKNTAKPLPAPYVKLHLLEGKQSVEKLRTMYPPRRTLDPLYQQQLCFSVPYHGKILQISVWGEYGRMDKKVFLGMCEIVLDDLDLKTVVFGWYKLFGMIASNAQHHHHHHHNNHNQKQQQQQQQQERRKSLSSSSTPLDNAQNINNNNNKLIGSKLDKLTNRSSSISSSSNHNNKNNERNNRTKSKSSQEINKSIQQSGFNKTNNDRIQSNLNMISSNLIKR

>XP_013421607.1|Lingula_anatina_I-RIM

MSRRPSLPPMPDLSHLSEEERQIIESVLNRQREEEEKEQELIRNLQDDLDSYQKSVVKLNEELSKQPANNSGGGGNQGAVCQICHKTKFADGLGHRCNYCNLRSCARCGGKLQLKQNKARSPVVVWACQLCKKKQEILAKTGQWYHGGQARPVALDIEVGSGSETASTKTDISPTSDRRARVSGGPRGQGQGQSDSSGQGSEKENIVRRPDDPRMQMSRSGSLRGSRELKRQYSMSEVGKIQDGSKEQGHYDRSPGGHEDRMSERGRARGRPLEQLPPDQARTRSLPPDDRHLEGMHIHGRRRGANGQDPMHDGHGTLHPPGSRSRDPSVERRSSDGRRPSPAELDRMRPDHDSHLRPDQDPRMDRGRMHPDGDPRQKDLPPPGRSRSPRGRRGELSPHDVIPPSDQELRRRSPSPARVREQEMREAHMQQQQQQQRDLNADHFHGDRRRESPPDRYADKEMVQKRHRREGVVHPPDGTKDMRDYQDGGRETIAMDRKRDMGEDRRRDGTMDRHREIHEPHDHEQLIRQDSSTSRRSDHSSRSVRERGMPGDQMQKPLGSERERELARTAQQPLQPDVHITAKPPQSPRHERMVVGTSPVSPREEVMDSMRDRRGGASGREEISDLDARSGDSRRMSQTHLDPSSAANRNSKNNRRKMETMLRNDSLSSDPSDCVRPPPPRPHKHKKGKSKQPVQQPSFSSSDDEIRSTPEYTSCDDQDLESESISEKGVGELETHPNPDSHAMQKKGVTLSSRGRSQGRLGQGEGQLGQGHMRNDRNDHRWSEVPIKDSGIDTGLSSSNQTLYEDSTKHPVTWRTADDGKKVIGHMILKKGVADGKGKDSGGILGLKVVGGKVTEGGQLGAFVTKVKKGSIADVVGKLRPGDEVIEWNGRSLQAATFEDVYDIIMESKQEPQVELIVHRNIESDMVHGNHDREHRDREHREHREHREREKVDRQDYPDQPSSASLRHKRPSISITSPGSPGTSPRRVSPLPCGQIQIKLWYESRNEQLIVTILSAIELPCLEQSTPPNPFVKVYLLPDRSAKVPHKTKPVAHTVEPTWNETFVFPGLRQVDFHKRALEVTVWNYSRNRNSELIGEVLIELSSACLTDEPYFYPLSRREEEASTPSTPRGQQKSRRGDPYDLPGSGGLPYTDSDISDYDDSVGVVSGSSLGACSQILVMCPLLCLLDNIDIIWSM

>XP_022257304.1|Limulus_polyphemus_I-RIM(1065aa)

MSDKSGSAIVPPLPDLSHLTEEERNIIENVMLRQKEEEKKDVEVLKRKQEEVNILESTIKKRAEDQKKMGLTLEATCELCLKTKFADGVGHVCYYCSVRCCARCGGKVALRSNKVIWVCILCRKKQELFTKTGQWVHGNLSDPLYRPILEKSHSTESENIPLDRSNSLHPGGEQSLPGSRRGSLERSGSSKSRDLRRQNSVQRQSREGALRTADLTLRDRGRSHSERRSPPNSDRQRRRSCSEPRVDAADEKHFPHDIQRGRHMQRDGARSRDYSPSPPPRDDTELHETGLERTREASKRDRRRGTREIKREDVQFHNPITRDDYRARGATSRDGSVERRSQRINESEVERYTDRGRATYRTPTYDRKTSRRDTQYRSRGDSENSRHIKGKYIREVPSDLCLPYQDHQNDLLVAYSGDADRQYHSPGDESEKGISGFSSPLMSNTDTRGHAGCYGRRKDVKSDLIVSENSEHVRTRLHKHRRSKKQRHRSPSSSDEDVTAISDWSSCDEEDLGGPNLGGLEKGVRSHSRYKTRHSEPQSTSRKNHTKQRGPSYYYNDQDICEEASLKKTVRFNRDNATSRQNSEELWDEQQTKDSGIDTSSSATLNEDNNNKHPVGWKLSLDGTRMSGHMILKKSVKEGESGASTAAILGLKVVGGRFLESGRLGALVEKVKMGSVADTVGHLQPGDEVLEWNGHSLQDRTYEAVFELIAESREDHQVEMTVSRPVRDIGWTEKISCRDKENKDFITLEDVSKHNMQQDWWPSVMVTSPGSPETVRLCPRSPVIGGRLQLKLWYNLQALQLIVTIICATELATRRGKLMPNPYAKVFLLPDKSEKSKRRTKTLSNTNEPRWNQTFVYSPLRQLDLKTRALQIRVWDYDCYGANDFLGEVLIDLSLITLDNKGHWYSLVRHEDSVNAQLRQQATFLDTDTGSSTLADHLSPLSNVSSHLSDGDTSDIDDGTVVGGNRNSVGGLDRISLSSLGSSSSPPVAGEEFSDVVERWVTRDLSPLGRKRSETGEETRYNHRGDKFCKVEEFINSTSRERPSQRGVEYTSRSLSPPVRRLQNL

>XP_022243248.1|Limulus_polyphemus_I-RIM(1630aa)

MESQSPAVVPPMPDLSHLSEEERKIIENVMQRQKAEEDKEREVLKKKQDEVKLLESAIQKRKEEHLKLGIELDATCELCLKTKFADGIGHICNYCNVRCCARCGGKVTLRSNKVIWVCILCRKKQELLIKTGTWMHGNLGSQEGSLDQGSGGESTPIKSELTPSSDRRMRLERGHSSEKENILQPTLTHPGGGQSLPSSRRGSLQRTGSSQSRDLKRQFSQETRPSSEYTYKVADKPPSDRGRGGSDKSPSQGESRKRRSIHEDDSRHHREGSDRGRHLHKDGARSREHSPEPPTRQERERGEQDFECERTRERSRERKREVSSERKREERKGSRELRRDELITRDDIRLREISRDQNVERGREDDRIRVREPSVERSRSYDERRPSRERRSFEERRPSRENVNQNYEREPQERLRDHEHRSRRDTSGVRHVHPDDKRDSSYRQKLPNSSEREHYVVKPVVSSIDLESRARPEYIIRRNHLDPSSASVVTIDSRGRSNNNNNRRKIESVVRNDSLSSDQSECVRPPPPKPHKHKRGKKQRQRSLSSSDDEIRSTPEYSSCEEQDIESESVSEKGEHAHFRFRSHCCDLNSALRKSCKLQRDPECYYDSQDIRSPFDSRTHKKTVRFNRESAPCRQHSDEFWDEQQTKDSGIDTSSSATLNEENNRKHPVSWQPSTDGTKMIGHMILKKTLKEGAGSVSSASILGLKVVGGKFFESGRIGAMIEKVKRGSIADTVGHLRPGDEVLEWNGRSLQGKTYEEVYDIIAESRQEPQVELIVCRPLSDVGRVDLRERAIRRLVGAPSETTLDPRMHSSIKDRRPSVTITSPGSPETVRVRPQSLVVGGRIQVKLWYDVTALQLVVTIVSAAGLTPRANQQARNPYVKMFLLPDRSEKSKRRTKTIANASEPKWNQTFVYSPLRRSDLKTRALEITVWDYDRYGANDFLGEVIIDLSSVPLNKEVQWFFLTSHEDSLNSLLRRQNMYLDTEAASTITSTDHLSPPSTLSRLSDSDISEFDLDEGSSLLRDRRLAGVDGASISSLGSSSSPPLVREEFGNIMERRSRRDMSPSGRRQSGTIATRNELVYDQNGRGPVIMTENIPPTLSAIRGRSRSAVHQNESRISRSRSPSRRGSEGTTRSLSPPEIRPHSPVIGPPNLAWRYPFTREGFSSSVNSPKKRQLPAIPPSLRHTSRNQMTLDLEERARQLKLRMQMQRRGGAITPVAMYSDSEVTSRNAVEKQLHVHPHRGRLQPGAMKGSSRHMSPGVGMSPEKETVEMITGIESDGSETSSVSKFSINSAFSIQSERPRGSRTLSEFTTRMQGYGPVHPPRPVRRTLNRSLSSEGGSDEKADGSLSDTAVGTITEKAGMLEQHGAPGKSWGGGKMAQLMGLSKKSSSTSQLSVTGHKKRLGFRRKRSSTINVHRSEEIAPQECRHLVKQSSSVSSDGEGSLSSDSTAWIPSLRLTPDGEYSHFVEGLGPGQLVGRQVLASPSLGDIQLSLCDRKGNLEIEVIRARGLQPRLGAKLLPAPYVKVYLVNGRKCIAKAKTSIARRTLDPLYQQQLVFHEDYRGCVLQVTVWGDYGRMEKKVFMGVSQIMLDDLDLSNIVIGWYKLFHPSSLVNLPASGQRNNLVSMDSFG

>XP_023224599.1|Centruroides_sculpturatus_I-RIM

MTTEKKGPGIGPVASMAPPVPDLSHLTEEERRIIESVMARQKEEEDKEMEILRKKQDEVRLLETSIQKRSQEKQKIGPEQLDATCQICLKTKFADGIGHICNYCQVRCCARCGGKVTLRSNKVIWVCIVCRKKQELLIKTGQWVHSGMAGKPSLVQQMELDVGLGNESISMSPTVDKRPKLEHRHSAEKENIHQQHSGLVPGMPFTARRGSLQKTSSSAAKELKRQFSQERDRELRPLTSELPLRGERTEHIMFEKSNERQIHSGSRTHPSSERGSPDIRFRRAASDEGPREEIRHYRENERGRTIPREGARSREPSPDLHHRRDYDHEVERSRGHISERELIRDGRERSREPRESSSERHRSQQRSNREYLRDDYSRDDRRRDDVKHRESMRDLSRPVDRLRNDTIQRRESSVDRRHGRESGHHEMIERHLDYSHDMESYPYREPAHVSGRLSRQEEQRDLIHRHKVPSGIDRDIYETHVEIIGHDDPHYRITPLREVSRRNHLDPNSAVSVIVDRGHNSRRKVESVVRNDSLSSDQSECVRPPPPKPHKNKRGKKLRQRSLSSSDDEIRSTPECSSCEEPEIESESVSEKGEEVLGQTWKKEEILDAKIKKFLAHPVSWQPSADGTKMIGHMILKKTVKEGIRGGSSAAILGLKVVGGKMMETGRLSAVIEKVKKGSIADTVGHLRPGDEVVEWNGRSLQNKTREDVSEIILESRQEPQVELIVSRPLRTCYFLRFVLVKLWYDIQALQLVVTIVCATELPFDRNRPQRNPYVKMFLLPDRSEKSKRRTKTIANTNEPKWNQTFVYSPLRRSDLKTRALEITVWDYDRYGANDFLGEVSRCDIFMFIFLKWTASLDKNCNKSSSPCGHPLLQRRQNMFLDTEAASSITSTDHLSPPSTISRLSDSDISEFDFDSRLPGTDAASLSSVASSPTLVAEEVGKIIERRSRRDMWPEDHSHSDLCTDSRLPGTDAASLSSVASSPTLVAEEVGKIIERRSRRDMWPEDHSHSVSREDMIYGHTTRRESTIPVDEIMSRGRSRSTVAGEMRMSRTRSPRRMPESTSRSLSPPELRPRSPITTISNRRSAHPHSVGGTPSPKKRQLPQIPLRRASRDQMTLDLEERARQLKMRMQMQKRHDAMTPVGIYSDSEITSRHGFEKQFHVHMHRGNRALPVNSVNSRNGGTHNVPPSIISPEKEVNVTVGIESDGSETSSVSKFSITSAFSTQSERPTGSRTLSEFTSRMQGYGPVHPPRPSHGPMDHSLSSSDGKVDGSLSDTAIGSVTEKDNGEKHHRSTRNVGGGRITQFGGLSKRSSSTSQLSVTGHKKRLGFRRKRSSTFSVHRSEEVAPQECRHLVKQASSISSDGEGSLSGDSST

>XP_018007323.1|Hyalella_azteca_I-RIM

MAGEGGEDLMPDLSHLDDHERRIIEQVMQRHRLEEAKESELVRRKREEVRVLEDTIRLRSESQRRRGAELEATCQICVKTKFADGIGHLCHYCNVRCCARCGGKVTLRSNKVIWVCILCRKKQELLTKTGSWMSGGPGDDPEFDYGAGTHRSPSTDRRAEHRGSHDVSIKPEALDKLTSSGNKGILREIFTHSGTELNRQVISTQHPQSQRSWNALSGGISEQYVGRMSLEASGTLARIGTEGSDVQAESSPRLPEPALASSGSDFTGGTDDRADKCFQNVGSSSVFGYSGQRQSESGTGLASPMDSSGVSGRGSTEGSSHRSDESRNEPVASTTRDRKRMDQVVRNDSLSSDQSESVRPRPPRPHRTRQNRDRRARQRSLSSSDEELWSTPEYTSGDEPTENIPEKGCSELVSNHNWLREQMVEAQIKEFLAKNSGLFPSDSYLGSRREKHYHKDYRKPKDEIRDPISEAELSRNSDAFSRPIFSRYRHRQNLYGTRLGKAYPVQVREECCDSRATDNRSYTERRKKTVRFNSEGWDTIEDDLDPSAVEDTRWSSAYNVYYDEERCPPGRPGPTSLHPNYAPMRDHGSQLLPLPRRVGGWGGPNSGPNRELPIPAVRRQSWDVERQESQDSQTKDSGIDSGTSSNFNSSEDSSKGDLPRTCRHPVSWQPSQDGSKMIGHMILNKAMRPEHGGQSSSAAILGLKVGGGKILDSGKIGAVVEKVKKGSIADTVGHLRPGDEVLEWNGRVLSGLTFEEVYDVISESCHEAQVELIVARQLTDIDRHPARRHTHAGLISRGGSHDGGFDPRKLSEPGRDRRPSVTITSPGSPDNYRTRLHSPSIGGKLQLKLFLDSQVQQLIVTVISASELPLRATGVPRNPYAKLFLLPDRSEKSKRRTKTIAGTCQPRWNQTFVYAPVRVAELRQRSLQVTVWDYDRFGTNDFLGEVLVDLNAAALDDEPEWYSLAPPDDMLTSHSRRTMCLDTESASTLPSIDHLSPPSTASRLSESDMSDADLEDSLHQRKMEQASLSSVGSSSSPPPDDHRGFGSYEHRSRREDPRFRSPGHMMGFGGSLGASVGPPYKDYIPAPIPMSARGRSQSAAPTDSPSLHVSRSRSKSPRRIPDVASRSLSPPEARPEYGGMGGGRPLMTSRSETATPTSSPKKRQLPQIPAALQHANRDKITQRNRDSQDLEERARHMKLRLRQMQPQGHSRSALNSGYSDSEIGARRYDRYTRGASRGALASPERDPLERDLGDSASDIESVVSGASTFSTQSERPRGSVKHRNSDSGVERRWRDEARQYVDRRPRRRDAPPRHALPARQRPCRDAGHDAGTYYGYDSEYSGAETEVFLSDRSFDRSFSQENDSDPYNSGSQCDCMYAMDECPQIVNQTPSSLPIIERISPDWPPRKPLLQPRKLVRAKTEGSLIDYARKDQRLRQETPGKVDYSNTLPRTESSSKKKKQVNFPFPSPTPKVKGNQHIGQNSELDSTKMRSDYNRRNHPHSQQYLINATSKGAHADQSSFCSDCTTPFPDEIIHSRFDSCYDKSGIDNDRYGPSFHSYDSSDATCSSCEERRSKERYLLKHDNCPIHCGELSSYRERDTDSIASPMHSLNLSYNTSACPSVKRQTIYTADCFSHSTDRAYREPIPPKPDSSHECFSVPFQLHVNTRVRPSSSVSINEQPQYFEFDANSPPTKEYRVIADVAPSSDPASELSDRMSGYGAPGGAFGGSGKRGTFARSLSTSEVPETEKAAKKGVRSEAKPTSPLAGAALGIPTALPLSPSSSTTKVVSSSSAAVSSPSNNGMKKLEFHVNLGDTPSTEGAKPDPDGQLASTPSNSSSPRLPLFDSADTQILELPESSKSRGQERGKSHTSGSTTLRTGVAGVPATEACSSLPEQEEESEKGTSSTSSERELGATKQSELHDNVDGSLSDSAAGSFTLDPKERTRLATGDSFAASTPGGGSSPGKSDSSIAGLAKKSSSTSKLSDTGRKRKLGFRKKSRSTITVHRSEEILPTASRHLLRQSSSQSSEEVDAELWESRGTRRRAPTGELTEFIEGLGPGQLVGRQVLAAPTLGDIQLSMCERKNKLEVEVIRARGLQCKSGCKTLPAPYVKVYLVDGKKCVAKAKTATARRTLDPLYQQQLIFHERYHGCVLQVTVWGDYGRMEGRKIFMGVAQILLDDLDLSNIVIGWYKLFGTSSLVSLPTFTRRGSVVSLESFEERP

>NP_001247161.2|Drosophila_melanogaster_I-RIM

MDEMPDLSHLTPHERMQIENVLMRQKQEEEKQNEIMRRKQDEVVTLEMQIRQRSEQQKKAGVELDATCHICLKTKFADGVGHICHYCNIRCCARCGGKVTLRSNKVIWVCILCRKKQELLSKTGQWINKTAAQQDGFIRRIEPDGSSDISQHPIVDPHDTTDKRPKLERTRSAAEKENLPMQRAGSMLRRQYSQQEQPTNRRLSASDSGMDPVMSPGQQMQHHQQQQRARMQPMNPQQAQQGYGMQQQQQPRSGGAYPDDDPRYYQGELDGLMRQHPHLAHPSQVQPQHTPQSHSQQQQSQHQQQHLMPQQHRPSPQQHSQQQQQQQQHSQQFAARQQQHQQQQHQQQVQQSYHAQIHQQPQQHPHPHSQQQQHHQQMQHQHGQSVPQMGGYQKPPPLTRNLTISGGHAMDSSAYSQQQQQQQRRALAGSQYASQQQRSFSSSEEEFQHQLLAAQGAVTVTAGSDYDGGRLQVRTSLDPWRQQRSLNERDLLSSGGGGGGGVGGADFRHLSHATQRLYASADQRDRYELQLQHRERLEQYPLTRTARLDVQQRNSLNRSNHERQFSGGGAGVGPGGAGGGGAAGAYRRAPPLAAAGGGGSSSSSTTSSSYYPQSKPQIVIGPGSGYDRGSGSMASMASAAAATAADAQGQAARNSQRGGLLGATRGGGGSSSVSSTEAAYWDEPATSSSSSAAQDSRRFTERRIKKTVRFDAHDEEASILASSSLPPVTAILPSTAVDVGAATILSSGKSLDAVGVGGADWSRWEAERQGSQDSATKDSGIDTSSTFTSSEDSNRGDGPKNPVNWQTSADNTRLIGHMILRKYYDGEDILGLKVNGGQPLATGGSVSGAGTGAAIGAGGPCGAIVEKVKRGSVADLEGRIRPGDEILEWNGRGLHNKSADEVYDIIDESRLDAQVELIVSRPIGSGGGSGGSSANVSPISGASGGSANSVPARRSSANFPHSGLASSMAGSAVVSSGGRYLQRKAPAVEAIEIHRDKPSVLITSPGSPDIHTSVSGPGSVSGSGLRQRGGVGQTQRLGPSHSTHSHSSSGSGSGSGSSTSSVHAPASAHGPVHASASASGSAPGGLHGTTTAGHHHHPHPHHYHPHQHPLPHPHQHQGPPAAHLQGHGVGGGIGMGSGSGTTTQPIPIEGRLQLKLGYDQNTLQLIVTLVCATGLSLRQSGAGRNPYAKVFLLPDRSHKSKRRTKTVGTTCEPRWGQTFVYSGLRRCDLNGRLLEVTLWDYVRYGANDFIGEVVIDLAHHILDDEAEWYQLQPHQDTSYLLRDEGSDVDGLILTPTDHLSPPSTMSRLSDSDTTSDCDIDGMTPGASISSMGSSASPPPLLELDLNERRSRRDMSPQGRKRVAGMVARDYRTVSGIGQSYHNQASATGYYRRGGGNGVGSAIGPGGMSLSQRSHSAAPSDGYHSSGVGGAASGGPAYGYRSTSPRRGSLSPPDDRYIDYPVLPVHGSSPNAPSVYQGPGGVSSAAASASQQQRFQSRSATATPTGSPKKRQLPQVPQTSRSAMLRDRLGQDFDERLASGGRFGRHRTRQPHHQATYRSTGMGGWERHYTGLSDSDLHSMDARMRPRHSLSPDKDFMGEFGDSDMESVVSVTSSAFSTQSERPRTSRGLSEYDKDKNGAEEAGGSKSAEGAGAEDKVDGSLSDTANDRKKGGGVDQERSPKGGSGMGKKSNSTSQLSATEQEQYQAQFAIAGSSTSSAGHHHITLPPPHPYHQRQSRGSAGSIQRSVEVPPVPPVTRYSAGSIHSITSSEGSSFSPSLRNDGALNEFVDGLGPGQLVGRQVLGAPSLGDIQLSLCHQKGCLEVEVIRARGLQQKAQSKMLPAPYVKVYLVSGKRCVDKMKTSSARRTLDPLYQQQLVFKQSYTGCILQITVWGDYGRIEKKVFMGVAQIMLDDLNLSNIVIGWYKLFGTTSLVSCPTNIGLGSRRSSIASLDSLKL

>OQV23161.1|Hypsibius_dijardini_I-RIM

ADMPMPDLSNLTEEERRQIEEVINRQKHEEAKDKEVVQNLANEVKQIEQRVIQQKTQQPFSGLLPDPNKEGLCDICQKVKFSDGIGHQCNYCRLHCCVRCGGRVPIKNGKFTWVCSLCKKKQDLLKKTGQWYHGNPMDLLGGDSAVSPNASMVGAPGMMSNHHAGMGNNVPQNGGNNRQQQQQQQHHPPSLQRQKSHDPGGGGPHQQQRPGGMSHSHSQLPPQSLHGSSSTQMQRQQSLRRHQTVDQGGVMSRSDEERLLSQGRGTGGRGQQQGGSQGLARQGSVIEGGGSSRRDRDASRTRQPQSGRGMAETGGPSRRTQPGGSADPFDILHAPTAARRPDSQGGNTRERRYDESGHLVANHSDYSGSRAPDYTGSRGPLRRAQSHDRYGSQDYADDYDPRRGASYDAWPRTSGDVAVGGRGAGGGGGGPGAYGGPSRASSALPPPLLPSQQQQTPRRYPSVEDRRRTYQQSSFSSEEDLQSNISEFTMADDGSIRSYSSEQSTRTHPDLRHMNSGGGGAGGRPGNRRMWRQSSQQEQGYYSQEGYSISGDTRPSEVSWRPTDDGRRMIGHIILRKDNFNEMNMGRQMNCAAALGLRIEGGKMTHSGRLAAFIVRVKRGSVAELMGHLRAGDEVLEWNAVPLRGLTFQEVYDIIEDSRNDSLVELMVSRRSAADYRLQLQQQPGHYFRSQLQQEDRRFSPQPSLSLNHPDGGNIASGGAAAGLLPTANKAPVPYVSGRIQLSFYYSHTDAQLAVTVIQAAALLPRPDGSARNAYAKLYLLPNRSEQTKRRTKIISKTNDPQWQQIFYYAPLKESELEDLTLEIMVWDYERYKQTSNEFVGEVLIGLGSDWRNDCPAWYYLTYPDTGLENRRAPPRRGGYEDPDFNTFSSMDGISPANSSASRLSDIPMDSYISDADMLEAVDRRKQRGAMASHQLSRYGGNGSPRHQGGGGYGFASPVDSPRQMMDPRDHRRGGGRESPHGLHYDTNRMHDRGRGRREAYGDSPQDDVRHTSRRHTSAPPVGDDYYYTDEQGGYEQGSPPGSRTSSARKRQLPIRPNDNALHGGTSPSHSHQLSPRRTSLSTSNKNSSSSQQLLASDDQLLLNLPSHVPSSPASNASRHSQSKAPIMQQLHNQTLKVPGSEDDTTPIMVPPSISVHGSSIRHAPALNQLLNNTSQEENIDPQTMMMMQEAAETRHSRSSSRRSVMRKPSASLSRSSEPTESDLMLLVTNSDGRSRKLSESLPRLQSLNIDDGASQLSALAVQKSVSNTDVYSKPKFDFDDYANKFLEQQSTAKDRKTKSSAKAAAILGLQKKDSKKPTFSRSEEVGHQSPASVQRALLHKQMSKDSTDGSVSSSGYGGNKAHRMDHTHLASFVDGLGPGQVVGRQVLALPCLGEIQLHIKYTKGNLEVEVVRAKGLVNKSGTRQLPACYVKVYIMEDKRIVAKQKTVVSRRSLDPLWQQTLGFEDHQYIDRASLQVSVWGEVGRMEKKLFMGVALINLDELDLSTIIIGWYKLFDSATVLAPATKSSLYAASVSGVGSNNSSMDEKSSPHGVNPRSSAVKT

> GAV01719.1|Ramazzottius_varieornatus_I-RIM

MADAFPEPDLSHLTEEERRQIQQVLQRQKQEEQREQLVLGNLAQEVQQIEQRVLQQKTQQQQPSPFGGASDKEGLCDICQKVKFADGIGHECNFCHLHSCARCGSKAPIKNGKFIWICTLCRKKQDLLKKTGKWYHGKGDELLAEIDEAALAGAGVGLSRDGTSLVVHGHAAASAVPPPSRILQRQQSHDPNAVPRGALSSQQQRQQSLRRHDTVSHNGPPAAEALRAQGRRPSLLAGPLPPPSSAFGPAANSARPQATSAAPPAAASGHRFGPPALGPPHAPPQARQAHPQATASGSLPKRGPAHPVHAAPSGGPSGNQGPAPLLRRAHSQDRYGSEYPSDHSEYEARRGHWQTTTAPADPHPSLHRPGPLHRSVAGQQARSASAAALYPQFSFSSSEEEMQSNISDFTMADDGSLRSYDSQHSIRTHPDLRHVSSPRGLQRRMWRQASQQEQGYYSQEGYTSSDSRAGSRDVVWRESEDGRRLIGHLMLRCDQFNNGEGGMGRGYSLPSQIGLRVEGGKRTASGRLAAFIVRVKRGSVADVVGQLRPADEVLEWNGVGLRGLAFHEVYSILEDSRQDGQVQLLVSRKLQRSAGPSAFYPAQAEGAYGGPPTMASLPAPGPSPALPYVSGRIQLSLYYSLPDAQLVVTVIQAAELLPRPDGQERNAYAKLYLLPTYSEQIKHRTKIMYKTNDPQWQQIFYYSHLKESEMLSLTLVVTVWDYERHRPTSNECIGEVSVQLAQVWLNDEPSWYYLTYPDLELQQQQRQPASLRSHRYRSMEEDYGPSQGFSPTNSPSASRLSGDSILLDDMEVVDHRRRRQQPSSISASRHNLAQAAAAGYYDRTDSPRSRHRSSTLQGQQYYRQNNHDFQALAYPNERRGRREHHEYNDDLAEELGPLHGQRSQRRSAHHSAPPVERDQAYLNHYDEDAMGPYSDDAASHRRPASSSSRKRHLPMAPQSTAALRDRRTSLHNHQREEGANGSAPRLGAEHNVPLSLSVPPPQGPSSPSAHSAHSASSRHSQATNNPPSLKTANGKAPITQQFHHQTLKVPAGEEEDSAPKAPPSPNPSHRSRLEQLSKNASQEENLDPLQLLSAEASNAAVATATHSRSSSLRRSFRGSSSNVAAGPLQDSNNRKPSVSLSRSSEPTDSNEAGDLSLLLPSSSASQASGSQSRSRKHNDLNRMRSLDLEETASQASGLTAINLGSIGTGNMSDGGGSQAGKKTFDYDEFQARFLNQQTANKNQQKSKSSAKAAAILGLKKDSKRPTFTRSEEVGHQSPASLQRQALSKQMSKDSTDGSVSSGGYTGRNHQRADHSHVASFVEGLGPGQVVGRQVLASPCLGEIQLHIKYSKGSLEVEVVRAKGLIVKGQSRGLPATYVKVYLMEDKRIVAKAKTNVARRCLDPLYQQTLAFDEDYQRRQLQVSVWGDFGRMEKKVFMGVALINLEDLDLTTIIIGWYKLFDSATVVAPNTGKSSSSSLYAASVSGVASNNSSLDDSTPIAVSSTSRPSPMQTQRSSNALQKVS

>NP_741831.1|Caenorhabditis_elegans_I-RIM

MDDPSMMPDLSHLSAEEREIIENVFKRQKDEEAKETQISQKASEELSELDKQITERKETSKKLVGTQDDAICQICQKTKFADGIGHKCFYCQLRSCARCGGRAQSKNKAIWACSLCQKRQQILAKTGKWFQPEEQPQPKISGSEPSPSPQPLTDDNVPEPQQRRAEPPDKMNTPNYQNNQQPRGMGMQPNHNQTQQNFMNQNQNSNQHPNQNHNQNQMQNPHQNQNHVQNNHQGANNHQQNNRRAMQQQPMSQNQANQINQMNQNQNQQQSHNQNMTQNQRNQTGPQNQQRTNDSRTMKQTPQQQPSQYQNNVGAAHQHHNQHGQQEQHHQQMNEQRTDNNRMRENTNGQGGMFNRQPSLEQTTPMNKYNHVEDDGMNQRPTFYTGNSENDQRQFDGQMQQGSQQNNQNQNQNNRNLRKNTVSRVTEEDYASSSNFESKKQRNNSSQSQSNTQGVRACPSTDDHLNRVKNRLHRQLRSMSSSEEDIIAGGGGNTLKMSTSAVVASGGKTAFHDDMGASNVQRLSEECNSEKDLLRYIYGDHKNSDSSSLGAGVGGSGGGGVLSGNSVLKSKSHALLKGSYQGGLDMQANRRRDKSLSLSPSRNDHFGTGSISGGDLLASRIRTFLSHPVTWQPSADQKKLIGHMILHRTENSAANGDLGLKIVGGRRTDTGKLGAFITQVKPGSVADTIGRLRPGDEVVEWNGQSLQNATYEQVYDSIAASRYDTSVELIVSRSAIIPGGDDFLNLTPSQMSSSAYSRVPSAYPSQFQRQLPNPDLFLDIHPALQQLSLPHSQSAVFPHNNTLTSRNRSTSSYYYSDVPDLGVPSNREMQESQAFGTGHIFGRIEVSFVYSHHDRQLSVALVRGFDLPPRSDGTPRNPYVKIFLLPDRSEKSRRQSAVIAETLMPVWDEVFYYNGLTEPMLLQRVLELTVWDYDKFGTNSFLGETLIDLASVPLDGEHSLMCILVDMDDDNPLRTVISIFKFHVIILTFLQRLKLRKASYNAPTRRPQSELNYYDHSSNYYDHISQNIDKQPHHHHLAPNDEENDEYIDDDELENDIDLATGGGARKSRTYRREKGMHGGHGYADWTQNHQRQSGYTSDHGYGRQNMIGRAYNRRQQRRPRSATALSQMEREDMYDPTRKHRDDNEYSMRESVRHGSQYYLGDQPLYEDGRYKISQGQMTPKQHNQQHQPHPLSQAHQQQQTAGVQPQHHQGFQQQQHPQQPNQQMQQMQPPMPNQGYYSDGSETLSVHSTNSMPTTMTTVNRRNMNANNTSNDNTSFAETPTANTNRVPIKETKQNSLASSSSVAGGGSAANNVMKERKKSLMTRFIPGRGAEGKRTGFARSEEVGIPGNLSSDRLTEPTPPFLKQASKESTDSAHSDKFARQCWLPVLADGPLGTFVDNLGPGQVVGRQVLASPVLGEIQIALMAGRSGIDVEIIKAKNLVVKPGVKVCPAPYVKVYLMEGKQCIAKAKTNAATKTTSPLFQQHLIFNDSPKKKTLQVTVLGDYGRMERKVFMGISQIRLEDLELGSQPLIGWYKLFHSSSLAGTGPVRKDSDVSVGGAQQ

>evg1261940|Nematsotella_vectensis_I-RIM

MFILFQPAATGPAQGFSLFTNLINEVKTLALGPEVQWLCLICKKKQEVLTKTGSWFGPKDIKALDDPSSQFGNDFKAIPSEEDKRAMHLQSALHRSEPDEEVESIQDSAYNFNHYYSDDEASRYGDNRGRRKVTFHDEQRLGMSRQPYGPGRGRGHPNGRPMQGRGMQPGIRGGMMPQQGPGVRPGFPPGPQAPGQPPGMRGGYPQQQGARQMAPGMRPGVPHQQMMMQQQAPGMGRGGPQMMQLGPRPGVPPQQQQMPQQPPVSTAAGTGRGGTVPPTPATAPPPNPDGTPRPEINGKKIPPRGMRPPAAKKAAAEAAAAAAQAERGAGQQQQKTSVPQQQVQQQQQQQPQQHLDTRMGGQPSPRPGMPGSVQQQQMIGQAPTHMGGHPGQRPMGPHAHQQGHPGQQMGGHPHQMGVHPHQQMGQHPVRQMGPHPGQMGSHPQQMGVHPQQMGGHPQQMGGHPQQMGGHPQQMGGHPQQMGGPQQQMGGPQQQMGGPQQQMGGPQQQMGGPPQQMGGPQQQMGGPQQQMGGPQQQIGGPPQQMGGPQQQMGGPPQQMGGPPQQMGGPPHMAGQMPGPQPMGTPTSGRRSQQGFSERRTGMVGSPPKIHNLGPSPQHLDGGMPPGQFTGHVDGRPGLNQGMQRPDQRMVPFQNQGMNADGQFSDGRSGFGQGLPGGQGFSPGSQGTPSQGYTPNRLASPSGYGPSDSRPGSSQPGMDRFGQGQKPSRFGSQGQDFGPESKRSDGRDYGSSFNGRDGRDGYSSPRTSMDSRVSDDVMGKRVDDSPGFAGEQRDGFRDGIGDDMVHKERRPTPEDNNNLESSIKWSPPDDDNKCIGEVLLTKDSRVHPSETDPSAMYGLKVVGGKMSDDGKLAAFVTDVVRGGPADLQAKLQKGDEVLEWNEQSFVDCTFEEVLEIINQPIDTPELHLVLCRDKSKFSPQDLERRDSPSRRFPTSSSGSPVKRDRTSPLTTMASPQDTGRLNMGPGPDTGRLNMGPGPNTGRLNMGPGPDTGRYNMGPDRTNDAIGSNMNDQSIFGSQVSNDDFFFGGGPDKSEASRKPSQGRVQAKLFYDEDSSNLNVTVVGAEGLLSRDAHNPPNPYVRIYFLPDRSMQSKRRTKTAMKTANPKWNQTFVYPCRLQKFQGRSLEITVWDYNKVGSSEFIGEVVINMTEANLDGNPYWYNMKSHDENGDPLNPPTPTQSPQGSFRSKGQRGPGDLGPMGGYGSDYEDDTMRGHGAPNGMIPGGHGYSPDNERLKQQQMQKQQQQHLQQQLQQHQQQQMQHRDSPPDSYGQNQMIQAYTPQYNTSVTTTAHGTTYPYTAADHNRTQPRSLQHPGIGRIQDGRMSPAGIPGYHTMPAHAQYAMGRTSPSHGPHGQKKTRALPQLPPGREPMDEEERIRMAKQKLREMEIDRSKLVIGGSRGRMGYGQAPDHVYGEMPYDPTMSPRPGRHSPRMSGSPVHQPRPGQGAPHLSPQARGPPRQRRNSTSMIPYTERDVMQGQMVAQPAPSPVPLGQSGKSASITYLPVTEGGERISNGHIPSNGVTRQGPQLTQQSRSGSVNDIAVFSTGGYLKTSRQAMRNEINYGGSGRSSRSSSIADSDRSGSLTSIPGSVASSESGSPWVPPSLKLGNEGHLGDFIEGLGPSQMVGRQVLASPSMGDIQLGLCYRKGALEVEVIRAKGLLPKPGTKILPAPYVKVYVMEGKKCIAKKKTRTTRRTLEPLYQQMLDFRVDINGKTLQIIVWGDYGRMDRKVFMGVVQILLDDLDLSNLCMGWYKLFSTSSMCDPPLPSPMGGSPKKSPAPSPRSSHGLSPRMSPIPTIMPPPGHMGGHGGHMGGHVRGHMDGGYDEQDGDYV

>XP_020898842.1|Exaiptasia_pallida_I-RIM

MSGMRGPSSPPPPDLSHLTEEERAIILGVMNRQKNLDYQTQEMQKQMLKEVATYENQMEKKTELSQRSVDPNVCEVCHKTKFTEGTGKECKYCKLKCCSRCIVEVSIPGAKQTTSTAAAPGLSLISNLIQDFKTLALGPEVQLLCLVCKKKQEVLTKTGSWFGPKEAGLKAIDDHGSPFGTDFRAIPSEEDKRAMRRQGSARGSFRRGANMENESDRARRLEHERQNYLRRPSLSDEEVESIQSRPYGQYYSDDEGTRYGERGRRKVTFHDEKGLGRGGMPSHGRGTKGPYQGKPGMGPQGQMGRGGQPGVRPGFGTQQGPGTRPGFPSQQGQPPGGRGFGPQQQGPRAMAPGSQQGIRPQMQQQQGQGIGRGGPTGSQQGIRQGYPQNQGPGRGGPGRGSPSQQQPGMGPKQQMSAGRGMPPGSSNVGRGGMPQSSPQMAGGRPGTQPSPRQGYPQQQMGGPQQQVRQQHQGRPMGGPQQMGGLHQMGGPHQIGGPNQMGGPNQMGGPNQMGGPNQMGGPNQMGGPNQMGGPNQMGGPNXMGGPNQMGGPNHMGGPQQMGSPHGQPLGAPKQMGXPQHMGGSQQFSGHPQMGSSPHMGGHQPIGGSPHMGGPQQLGSPQQMGSPQQMGSPQQMGSSPHMGGPPQQVGTPRSARRQMQQQGYQQQQQQPGSVAERRTGMVGSPPKISNLHGHNDPNQQGIYPGQDRPTQNGTMPHMDGRGSSAPVKIPDGRGGYRPDSQNYARSPDNRPGMGPQQGLSAQNQPMYGSSPXSYGMNGKSMDGRGQFGSPTNMYGRDSPQSRLSNLSGRGGSGLDDSYGGRQSGYSDGIGGDMMDDRSTTGSQPTSSESAIKWSPPDDDNKCIGEVLLTRDPRILPSXMSEPSSLFGLRVVGGKMSDDGKMAAYVVEVKRGTPADLQAKLQEGDEVMEWNDQSLVDCTFEEVQEIINSTDDSPELHLVLCRDKNKFAPKDNQPTRRGPTSTGSPVKRDRTSPMSGPQDSRSNFGADRANGPVGSRMNDQPGFGSQIADDDYFFGGGPPTDNRMADSAGHKGRFLGRIQVRLGYDEDANNLNVTVVTAEGLSSRETNTAPNPYVRIYFLPDRSMQSKRRTKTAMKTANPKWNQTFVYPCRIQKFASRSLEITIWDYNKIGSSEFIGEALIHLADANLDGNAYWYPLKTHDENGEPLAPPTPNQSPQSSFRSKGQRVPGDTGYGSDYEDESSRYHAPNGMIPGTQGFPTETDRLHRDSPPNSYNQQIVPGKLPPDDNTSGSLNNLTVLPTGGYMKPSRPGLRNDNYGGSGRSSRSSSIADSDRSGSINSIPGSIASSESGRHGLYAIMNAENPIPWVPPSLKLGNEGHLGDFIDGLGPSQMVGRQVLASPSMGDIQLGLSYRKGALEVEVIRAKGLLPKPNTKMLPAPYIKVYVMEGKRCIAKKKTRTARRTLEPLYQQCLDFKVDIRGKTLQVIVWGDYGRMDRKVFMGVVQILLDDLDLSSLCMGWYKLFSTSSMCDPPLPSSSPLGGSPKKSPAPSPRSSMQHGGSQRMSPLPSIMPPDHRRGVHNYEDPDGDFV

>XP_012564401.1|Hydra_vulgaris_I-RIM

MIMSQLSTGNDYYPASPPPIDLSHLTEAEKRKIKNVLDRQKELENETAFIQSGIVRELATYQVRYELKSNQAPLSDANMCELCHKNKEFCVNNYKRYCKFCRSKLCDECSIQSGSGKMVTWSCLICKKKQDLLLKTGRWFHGKEHKFPNPAAEIEQLLTVPTQQKKPLSPKIIKTGYPASSVTKAQLADKNCTKVLDQTNNESTELSNTKKVTFGNKSKSNMLEEQQIKEMFQSLVSHHVHWSRPDDENKILGDILLSIGSNIIKKFPDYCTAFGIKLSMGVSDEHGLLKTLVKSVVPGSIGDIIGKIKPGDELLEWNELSLVGCSKEEVEDVLANEDSLDLHLLLSRKCHKEKILANIQSVPPSHHSPPSQPMGILKTIDGSFDKTDCQNGAVECDSSKSNVVQANQSRGTPILKANNISGQLQIKLKYEEDTSCLYVSIIGAQGLNHRQNMQLRNPYVKLYFLPDRSLQSKRRTKTVLRSLSPTWNQTFMYTLKSQKRDTSILSQTSVLDILPYLSPEDINEVKLCHNSKSSDSKNYNVFKGRYLEITLWDYERSESNEFLGEILISMLEAPLDNLCHWYPLQLHDQQNMLPQPTPTPTLSPQCSKAQFILQDNGEREHPVENQTVSSFSPADINEKINEVGTVKTSFSYDSTGGTSTDENTPVPSPSPSQVFEMNKELSQITSSKNSDNEKRQYLKPALVQESSVIRPSLLQRAQLFLQKQLDVLTSSSSSETQLTKNSTSLWTSNPTNYESKLTTVTQTENQSPLSSLSVDTSSISSDMSTVNNIRYGKEVYGRFSSEKYPQIKRVDSVDSSRKIEPRLTDYNLSLSQQGDRAGDKSAEEWRQSGNDNNYGVKFNNKPTLLRMDSMSGSSPKNFEKKAENYYREQHNYNEDKRFQPRSNHEISFMRSTDFDHNGYSAMRNVNRYRQNSYGSGHSSPVSSIGSAKSESSSSIASSMMTLTSVGSSMSWIPPSLRLSGDVQLIDFVEGLGPAQVVGRQVLGSPLMGDVQLSLLDRRGVLDVEIIRAKSLIIKPDSKTLPAPYVKVYLLCGKKCVEKRKTKTSSKRTLDPHYAQHLTFVQNYQGKVLQIMVWGDYGRLDRKVFMGVCQILLDDLDLSQLVVGWYKLFTTTSLVHPPSNPSPNPSNPTTSSTNSDRF

>evg1642237|Trichoplax_adhaerens_I-RIM

MWPSKIQNVFFGNTNRSTSHNGHDGDDFEVVEHLPQFSRLTEEERQRIQEVLKKQKQFDEQNQAHVDNLKKEVNQYDQAKPVREKNICGICRKTKFAEGGGNHCKFCDRRVCKRCGKFITNKKQEKNWSCLACIKKKELQARTGEWFQSASTSNFSPLRDPAAVDNMIDPDKQYEHQNTGENVNQKPTLLSTSVSSLSKAFVTLKQNANSVKQALSNANANNNDENQTSQLQQSQSLNSSTVSLRTNSTSSRREHTGSLLLTPHDKMDGEGIPVSRSSSFSAMPKRKSSGPRPPRPPPPRSNSTSSSRSPTRSPLSQSPTPDKQEQSIPNGEMNGHDVELAKSSQEEPKLATRNDHHVEDGVTPDDSKTKKLTPSRPPLPSRASLKRRSSRSQSTSQPSDSATPSNDDNRIIPAHDNNVNATPPTDFSDNPRQPEKEPTEDSSDSLPNGEEHQPTSPKASTVQPPKVENDSNTTTAEKPTKTREAAPSNMTSVNQDTPQNNDQETAEVKAKPSESKKVPPSRPPRPHPNALKRTLSNKMKTQQQPRKDSKTDQEVPKTMHVSDDIPQAQQPTSVQLHPVPKEVTKAAKEEPITAEENDATLNEMPAQTNEHSATSKDSPTRGHSVNDLPEDAAIHDSHDNHAEISPKDQDKSITPHGDAITQKQEHQNHAEVIPEDSGKVGDLHLKADRQPVSKPVVMPEVQKEAPQTVIKVNNDEHIKEANHTGSNNIAPPDTTSTHSRSSSVGNFSDDDVFEDALSDGETEEHTKAAAVGTSESAAPTDGIKEPINTPVTVNEAKKTSLESPVQKRIRIPSVEDTVDAALASGIATAAVLSSPKNLQNANTKNQAFKLPPDHPQDGIKSSEPATVSKPQQADVKPKIENEGPTSDNKQTNLKHNQSGSKQHQDSKIPNSVSSSNKMKNTITNRNEQPQVTVQTKSALPVKIIENQTPASQLKQSQQLTKVEKEPSPAMASNHLQNNTNNGHNGAMNETKSVPTSKSKSKSPKSTPSKTKSAIAAGIVLADTTAKNKSPRSTPSKTKAAAVANAMNADSKTGKGSTKVKNKSANSTPSKTKAAATAAAVNSSNSTPVDNSSSNKKSSVGSRSKAAEIKSSKPQTSTRRPLSALFGGKSKDDKPSKSPEVTKKSAGQNIQPSNSTSKDMNRIQNNVKPTVSDSVATDGTLQASQIQSQAKLEKAETQSKTTESQTSKENPPNAKPSLHQQSVSTPSSHEVGPIPNNQEPKVNDDHHEEGEPPNIVINRRSSKIETKILKPSELDTAIQRKSQLDSNDNPNTLNDINKLPATKQDDTKPMTSSNQQENNITSPESKPTNNNSSKFDNNPVATEEPLTNNYHTNNKSQIDLSPVSKSKEVEENTKVTPQNYANDTLKAEPTSNRNVSLTSDATTVNHNSQITNVSSKESINVSPKQHLQDEPKTITNSSQEIEITPVPKTKGAVNAFLRSGMDENDSKSDNHSHEPPTDPIVEEEEIKPTVHRVTASMPLHFLLEPEEDTELDHDGKSSEPSVTETAQSNPKVQVPTDMKVNHIPSADEEAKERVTVHPINEHDSNKVVKDLKHGSQATPYTKSNSPKVAIKSKQEEVDNKPHTNLHDTPSYTKAYPQKEEESIHHDERNNVNAATEPKVNTTETANLSPKMNKIEESKTIHTTDQSDHKTANLSPKMSANESKNISQPDHSDHDHETSGVKAPMTSTAAVNSSITTTSHDVPAAASIEKTSAHDEIANARTSEVAPPSSTAEKAPVKAETAAAVAGTTEAVATTSATISTASTIPANTAHEITPAASTSIKTVRNATVEDATVVKLNSNTTNESMSATPHVEAGNDAKKEPNHVVAQDDHDHAKVVAAAVGATAAAAATAAAVSVTADATAHPSPEINTKEATKAAANLHADMHHEDVKSVAAAVIASNRARKRLQNRIASQTSIQTVDKVTGRMKIKLWHEKVTENLVITVFGAEDLYPCDHKVYRNAYAKVFMYPEASQFGKRRTKTCDRQLNPTWNQTFMYFSISDKRLESSSLQVTIWDNDKDHGIKFLGEVIIKMATARLDNQPHWYLLRNHDHSVGQLPPPTPIKKTASDKQSSHGLRRSNVSLHATNTNDDSVGAISEDSVVGHPEEVYHSSVNANKYRVADSSNYAEHAMSNTGHEDAPSQSQDHPSEADDNTKNQNHAATSDASNDDKKTPPRKPPRIDRPPSEIENPAVQENQTPLKTNHNDAHQTNHVSTTPTSNHTTPESQISAHQTTSSNQELTPRSNSHGRRQQNSGSVSSSGSNDSKKTNLPDIHNGNNINPPTSHTSNSPGGSSSRMPQNWTNGATKTPEKIRKMRDALGHGQVVGPHVTTSSSFGKLQLGLQYKNGILEVEVIRAKELIARNTKSLPSTYAKCYILDASDKYIAKKKSKSVRKTLEPVYQQKLQFPVSITDKTLEVIIWGNYNLIDKKVFMGVAQIVLSEEDLSNLIISWYKLFDSSAANINNLDASMNSN

>m.21147|Oscarella_carmela_I-RIM(739aa)

MDMAAYLPVERPLITSSNPRLDVHGLTPTEIAQIQQVLARAERANEIVAKKTRQRKDEYEHLKSQVEAAERIMKSYHGGTAKEKKNLCHICFRVNVGLTVSHMCCLCQRKFCSSCGCRVKTGTKPGTPMEWTCALCYKEHRYMSLSGDWLSGRNQCNLSVSPGSAFSATSKTLSRSCGDLLDCIGSELNPTGDLDSMGTCGRILIQLGYLEAQCSLIVTIICAEGLNYTDVKDRKCSINPYAKMYLLPEKSVKMKRRTATLQETDNPAWKRCFKYEKISPGELKTRSLEISVWSLKKQKPRNMKRQLGQVTIDLSEVPLDNYPRLYPLDDPCELLMPRSCHSLPRQNAVDGGGVTHRHRRSTSDISSRNSMTSDESSCSDEVLKRGGLCHPSARLPHVFSDRDSGMAVSLSSVDTCTSGRKSPLSMIGQLFRKPCDDVDAHVREYRMSQAKQQAVKSKWSKRSSIEEVNATLKSTGDCSSLAIPNDDTYKSSSLPRKVSLDIPEGTSDDKRRHSMPVVTLRGGRSTSPALSNERRSTTVGLDRSHLSPRRYSRVNETYCLSPTPSEDGSLGSTVSALHMSDMQGLGIGQLLRSAQIGVENSGRIHLGFSLANHRLQISVYAANDLVFNSEDLANGQANAPYVKLYLLHDTKPVAGLKEQRTKCGDSLNNPQFGQTLTFRTTAIHNKILKVKVKYDTVKKTFKRKSHNHEIGQALIALDDLRVNNEEVKDGWYQLFPSRS

>m.26069|Oscarella_carmela_I-RIM(1164aa)

MGNAAMAKRAEHSSDAADSSAAFGASDALRAAFPSLHELDLSHLTAEERTQIEIVLGRQYAIEMEEEDRIRALKKELFELERSVANRKMARGGIKACFVCSKALARPRMTEKGVLPESTSQCDECGNVCCLDCGATKSGRAVNETEKWFCEVCGKKREFIMKSNLWHSRSKISLAKENGSLDLPSDRKRRSASPNPVIMLTSPDFDANSILFNAKGDDASTGSATGSSEVGGFTPPSFAVDALGSSGGSSYGGVGANRKAESRRKREKMAEEVASSDDGNGDPTSRARRNQKRRRLPEVPADAKPWTSPASAAVYRARMRENEGSEEDGAELRNHTTSDGDGVLTEARVSRLLEQNEALYAVSGVSSQPPPLPLSPPPPLTPSPEKGERGRADSGSSVDDPAHPYARIEYKVRGEAAEEIPDPGYAKLRRPGDTAVKAEEESEDSFGNDGGYSTVGKKVGTVIVDQDDSQYTSVPEITKKTKAEESLPSQHKVDNGKLQDISDLYAKVDKSEKTQHKVGNEKLQDISDLYAKVDRSKKTKAKAKAKDKDKDKDPSAIYAKVDKSKKKKKKEKEAREDPVGEKKQSDEEPEKPKDVISLFLGDSEKEKLAQEQPGKTKTARHFLVKKPEWRKGREMSRSQSEEVRSPKGLVKARKAMFEGMSSKASTLPKTTTEENQDERTGKPEKATEETWNHPSKADSSPQSFSQSLSSSQASISSATTHTRDSPKPPVIHISNEEIVDIVLSVHDSAAADADPGHPFGLHLGPASNGIRVCVQGIDSGSPADLCSKIDIGDEIVSVKNISYLAKRLPQLEELLVKEKTVKMKIRHKLKKGSAFKEKIKEKQRELNSEFSLAEAVRAGDENSGGEVPTLPPVTEIEEEQRNESVSCLDSGLEAELSSPIATSSPSDGVVLREKKRPGRVKAFFSRLSRPFGNRESMLELRSTSMDFLNVDRGKKTVSLPRKFGGSMNVLNMKEDDMASSLSDSLSLNDDLTRHRETTPESECDLKCQNPEWHLGKGQMVKARENEKLLSAGKVYLSIMHMNRVLTVEMIEARDLIGQRHAGAQIPYAKVYLLSSGRERLAKQKTKQATRAVDPAFYDRMYFKDVDLEDKCLKVMIWRRSSPAVFSKKRCVGQALISLSELQSTSKVTNWYELFVPSSRMPIKP

>sakowv30000298m|Saccoglossus_II-RIM

MFKSTSAHEVKTPLSPIPDAREGPMPDVQLDHLTAEEVAVLRQVWEKQEAYERAGEERIRTLENEIRRYEEVIAAKESERASGCFREIDLQLCRLCYSKKFADGVGRICCDCSRRVCIKCGSFYISADEMPENKLRCKMCELKRQLLCLTGRWCHGGATKPMVKATDFKDKVGTLEAGPYVNRKTTHVNKQDQRVPRRRISHQEFTYHTPDRIQLSPSIHKPRSLSCRRKADGEVPAQRETAMSERRRRMPRRSRQQSWKTLSRGSTSDSEAELLKENQHLLVVGSYPSHGYGEQMPLSHSAHEMDYCLSSHSQNIATVYSTSANEDERYRERVVHQQSVSKRRPRRERVLPKIRRMDTIDLTEGDARATAALQQLERQRSTAETTVRRVLLCQDKHDKSTRTNGFGMRVVGGKVGQNGVLGAYVTMVVDGGPARSDAGILEGDQILEWNGLSFINKTFEEVKMAINESDEEVNLLVAYKPSATHEPLDRKSSDNCENSNQTMQKAIHQTRASLASQVSEDQYKEMLHSPTTPRRKLPKSPVDLQAARATISGRLQLSLLHNEEDEELVISVFQAENLPVRQTSVGEQQMPNPYVKVYLLPHRRELLRFRTDSTHASTNPCWNKTFVYPNVSTLELEHHSIEFTVWDDKPVTLSQFMGEVLIDLRDAGNEPRWYDLAEHDENSPPLPKPSPTSVRKVSPCQRMEKTMQAVMYHARMRERMQRSMSISEPPPEEQSEEEEFPGAKNIKPLLRKLDELNVDENGASCNRRSSRDSTKEGDSTSERSCSSNPSSAGSLTLRNRSLAAPEHPLTGFRGSLKDILKKKLSTSKNPSLKPDSKKRSVSSHDLERYFDETGKDKRMMSSRRSLNDIGFPKSVSENTISFNRRRYSWMSPEERAVYRGRGLSRASSLDDGSDTNDSVASFDSDPMRCPSYTRPAPEGDDVASVRSSNELGSLGPGQYNVIALLDEMVRPVGVMKIGIVMTKGHLEVEVICAKGLPKTENEQLPDTYVKTYLVEGSRRIQKKKTRTVKQSLDPAYRQLIRYSACDVYGRNLQVMVWEKLGTLQHNHCLGEVQITLDELELCKHTVGWYTLFPSDVYKMGSCDSISSW

>XP_012944640.1|Aplysia_californica_II-RIM

MTFLLKKVIKPWGSSKDSRLSVPEKPVCPPSPDLSHLTPDEISVINDVIRRQEEFDRQEAYRIKKLKDELDGLQQQLVQRSESALQDRKQVDLRLCRLCFKTKFADGVGRVCHDCQQRVCSNCGAFSKPRWNAKKNKNVRGRWRCKLCSVRREVLCRTGGWYHATPDSSEGIRNKLSLALDPEAETDAGSSTCQGNGTNMEDSEHYPSGNRTDSELLRRIPTHATNGGMRRVPPGTKRFNRRRSIPPDSAGGKDEKLGKDKKSDERLFLVDDDDDNDSLDSMILERRMSRKRRQEKKQRRKWRAGQDASLESLPVSDRPPSCSSGSGEMVQSPQSDVSCPVKGRSGQRRDSVTRPLWGHSSSLNTSIADRRPSPIQEAKGGYSDTDSETSSVFSSFRDSSFSDNGTCSPYPGPLLLRIIRQDAVSLNSSCFSLASANTDSRCGGLGARSQMGSSDPLSARSRDSLRATSPVVHERKVISSKSKESPKGGSEKSLTRWKNEGFDNCFVFTLHHTPGKEPPYGIKLTGGIVDSGHVGRTVITWVSPDLKSMLGPGHEILEWNGERLRGQTFDQVVTTVSNVQPHVQIVLDRPECRPDEGKTAPEEVMKPSQSLDRTATWSPTIDQAGKGGKRRMLPKTPVEIKRRTRRVHGELQLRLHYQAQRASLAVTVLRARHLCPTYHIDPAPPSPFVVISLVPFKRGRDSMETEIKASNTQPEWEQTFLMTGITLTELATKSVQVTVWNYETSADAFMGEVLLDLAQAPLDNEAAWYKLEEHDENSARLPPRRRSHSASMSASLTSGSFRDSLLTSPTASPEVPSLRQHRQSPCSIRSASISPVPFGSASRVGSREFHLNSAFQPDSFTTKVRRKMRSTVNKMSSLSLIEKKDNAGSDESYHSDAHRGPNDRSDRSGAHSRSPSLMKRGQNSSHQSMDLLAPPQAPSRTNSSSSFFISDDEESQGRFSTSPVAPDRPAPDGDDITSTLGPGQIPPKPSAETSVCGNIKLGFMVSK**GQLEVD**IICAVGLHRSGQTYPPDTYVKTYLVEGKKVIQRKKTLIVKSSCDPLYRKKVKYSACNVHGRIMKVNIWEKAGSFEKKLCRGEVLVKLDCLDLSKHTMAWYKLFEINSTDYGSDEFLNSW

>XP_011422085.1|Crassostrea_gigas_II-RIM

MAHMFRKTKSVRSGHPSTSGRVSPVEPPSPDLSRLTQEEINILHKVIQRQEEFENEEASCIWTIRQQLERYEEAVRAQSSGKSRLKHIDLRLCRLCFKTKFADGIGRLCHDCRKRVCQRCGGFTKSRWDPKKKKMVRGRWRCNMCDMKRKFFCKTGVWYHGQRARPTLNRALFSQKTRSLEADLTEDERCWRALSPGSSDVVVTDTDLPRTHASYQAMMQARGRRKSMPIFRTESVDISDNDSDEDPHSHSARRRRSKRHSRRHSRRNRPVSPSQAVLTSPGHQRADDTYEDSCPHTGDLAFVCSDMPNQGELFFYMSGDSDNESVPETSGRLLNHRKPDGQSHDSSEHLVNVLSPDHMGSISYIRNRESQFSYPSNRSPDKIVIDSVGHDVERRLSNESDIRYETAYQHNTPRSERRRKNQSVDPKYLTPENPERNRSYSSDSNSSMRSFVSSMTDHDHAEIHQIPFTQDYFTSLEPREILLYRDVKDQSIRSSGLGMRVMFGKKGHGGKLGAFVSKVERNGPADSYGIRKGDQILLWNGKSLMGTTLEESMEVIGQSSDVVQLLVVHHTNSFSDNEDTGLSVHEKTLPHNTSVPCDDRSGCATPPSSLNLTRKRRMLPKTPVEIKKEDRSVSGCVRIALFHEEPCLSVTLVSAENLVSADPMTSLNPVVMIHLLPNRGSFPKHQTQPRSNTANPIYNETFTFQGLKRDELSRLSLELTLWSVVGEEHLFLGEVLLDFCDLQLDNQVRTYNLQDHDENSSPLPLRKRKESSDVTASPLTSVNDGSSWSTSPGVQSSPRRELNHHDDRAGHHVTTGYANGRDRRFKLSVTQSVPNSPQLSPISQSEESVYLGGDHRGSISTLVKKRMSNAMSKVSVLSAESRRHSMYDRRKDKFTSRSSSVSEATLESAWSSRNYDNPSPVSRRSAASNISGIEELGKPPQPLNTGDSAGHFREDGDDSSGENEAGLCTTSGRLDPDGNDVTSLLGPGQVPPKPSSETAICGSIRLAFMVSKGQLELDIINVVDLNRSQHITPPDTYVKAYLVGGSKVIQKKKTHVVKGSFCPFFRRTIKYSACNIHGRAIKIILWARHGAFDKKQSLGEAVVKLDGLDLMHQSTNWYKLFPPGATDFGSNESLHFW

>XP_021345230.1|Mizuhopecten_yessoensis_II-RIM

MHKSSTDNQHGSYNSSASLKLLDCYNNSSASQHQNGSFNRKTSVSVDETFKKPFSSKHRAYNNVSAPHLGRSYSPFTNNSASYNLKQHDGSQDSDSLCDIMPPVLPSPDISHLTSEELDILKRVFKREEMFERDEEERLREMERRLETYEHTVRQQATGKGKVKHMDLRLCRLCHETKFADGVGRVCHDCRRRVCLRCGSCETPKGSPRKQKNVRGKWRCNICQLKLKFVCQSGKWFHGNKSASMVNHPIRNKIQFHDADTSDTDNQTSVDSSSLFSASEICTDSEQRKFNRAVSMSGSKNMQNRNRRRSGSLIPRSSPSDNDSETDDSVERARAHVKQKRTQRRNSSRRKSLKQRQLLTDVPDQDNKYNGDLTSGYVVNGGNISPCAHLHSMQRLNVSSDLENRVRRHSDESVRSGSVSVYFKPDDSQTETTSSKRSSITRPNPSNCGFSHMQAGLTRDSWGYGLQPSQQSPPLISSGHRRSMPGLIPRSTPNTLTVDDGNSPSVRRRHSHNISNSGVLGFKLDGKRFSVDQHTSGHKIKQTYLMPPESPAARCRRWSSDSGSSRRSSGNNETAESTDWNEDIKRQEYLASLKSRDVLLYRDKNDQNCRCNGLGMRVVGGQCRFGNRLGAFVTYVDTDGPADSYGILEGDQVLMWNGKSLVDTTFEETKRIVNQSSEIVQLVVVHHEENYGMVDITETAEYTEQESIPSIVQPIPLPPSVLSLNKPKRRMLPKTPIEMKKDERLVSGKLQMSVSYSEEAECLSVTLVQADDLRPPGQLELSVINPTALVHLLPGRGFYPPYETIPQFNTSSPRWKETFNILEISASEVNDKSIEITLWNKQHAENMFLGEVLLDLVDANTTDEIMTYDLEDHDENSSPLPRRRRKESDVTTTSQISPLTPTIDTSSNWSTSPSHVSNQNLNTEGRQSADERNSRYSPLQTRHSLPRSGSFSAKMKRRMAVAVSRMSSTFSSGERRMSLQEDKIATMGTPSPDAMLDYLVPPIPISPRHSDELSKGQLSDISNSARSCSETGASMYDNNSDIDERRKRFGSTSPDRPAPDGDDITSMLGPAQVPPKPSSEQSVCGDVRLGLMVTQGKLEIDIICVTGLLRDSENAPPDTYVKTYLVEGSKVIQKKKTSTVKASYEPTFHRKIKYSACNIHGRHIKVKIWARQGAFDKKLCLGETVIKLDGLDLSQHTHTLSWYKLFPQGATDFSSNESLSF

>XP_014774175.1|Octopus_bimaculoides_II-RIM

MAHYVFAVPVLILRSGLLDWMLDEIRAIDVKTCKILIGTHNFHINSNNFNAAIQAEVSNSSLTPPDQFTFHSQYGSTLQVVPRWNFKEIVVLKKIDLSGKFARVFGFGRGKDNSQSDEYGNKSTTQAESKVLVDNSCEASLAASAQCPPMPDLTGLTNEEIQIITNVFRRQEQFEVETKDRIRNIKMALEKYETLSTEFATQKTSGYVDLRLCQLCFRVKFPDGIGRVCCDCEKRVCHKCGSFARPRWDPKRSKSIRGKWRCAHCQLRREMMCKTGTWYHGDDTVDTIDSSESTIKPKLENEDAITTLVDLMYNKQDADKYLSNVNKDTLTDVSSDVFRSAVETEGHPTIAQNPVNRRSYRARSLPTGYTCEYRDEEASGDTEQFETERKMRRKKRLDRKFRRRSVLRNRSEFDREIPEIIVNPPEEPYRRSARSRTLSASSVSSRKQQENNVKRKPIRKGDIVCDEKKNEKLTEKRSDSHNKIQRQDAVCYSSSSNSLTSESKVDIQGNDKTKTLEANDYTIKKSSADEKKNRFRSVSLTPTRVRTLDVTPVFRSSTYDVPMVRGNDLLAPDTSPNKCRRITLPLCQDLGLAAQKAKGDQVNVPPSNPHEVILHRNKDDSSCRTRGLGMRVIGGKQGKWGRMGAYVTNVEEGGPADLQGNIREGDQIITWDGQSLVDVTFEEAQEIMDHSGNIVQIVIWHRPVTEEETVAPTAASRREKFQRADTTDMELTQSTNKEFESFGVRRKKRILPKTPLEIKKDTTRLITGRLQMKLNYKPEQMKLFVTLIKVTNLVPPEKPERKIPNPVAMLHLLPKRNESQVFETTTILECSSPEWNETFTFVEIPPSR

>XP_013397099.2|Lingula_anatina_II-RIM

MFQGLLRKVVKKGSTVPEQPEEVARIPGAPPSPALSHLNRDEMEKLTEVFRRQEEMEKLEQKNIKKLMSELEAFEKSVKELAASKSGSKALDMRLCKLCYKTKFADGIGRICQHCQKRVCSKCRQYLKPTWNTKKHKFTKGRWVCNLCSIKWEVMCKTGLWYHGPETIGNKPAGAFFGKLQREWCLSVTDGDSQMDSDSSVCRRGRNNESAVTDSEMHTCNQTPNEQHRRRSTPNDGESRDRRRRLQRQSNRTAVVAPTESYSDSDTGQDRDRLRTRRRMTLDSYGRLNNDARSKIRTHFEPSDNSDSDTLEQNCRFRKKEGSHANGEFTINVLDNVRKKLEANVRPSSDSETGHTQSHVLPRKRVSSIANGELIFNEWNKARHNLRIGEKHNTEAGVKHRRRLLGVASDEDEEFRGIESNHRDSVRNDVGEVRNNGGQRQALLLHLQSPITAEVDGDVFSKMSSFHVDVRNSIRHSSGRQESRSTESTTQASLSHPRTMAKVAPFSREVPHGAKEGERGRRQLPTLNIIPNIDVSLMGEVLQVTLQRSSEEAICSNFGFGVRVVGGKISENGILYAYIARIVAGSPADVEGNVSVGDAVLEWNNQSLINLTFEATQQIIHSSYVQESVHLLISRVQKRASLLHSRKTLQLSDHGKGDRFSDINDYDSNRRHTDHFQFEVRATESRKHRLLPPTPVDSVRPSTSATGRLEVQVRYEMASVKVTVRQASGLKRETNESGQLDLPNPYVKVSLLPGRWQYPVHRTETLTVNAAPVWNKMFVYYDITEEELHTKSLEFTLWDYHPKENNTFMGEVVLDLNAVNLDGLAQWYNLTPHDENNGQLMQPTPKKSDQKGESEPAEDIQTPVPPMPAHERAKSLLKTHHRGSGTLKRFLQRHLTRSSLTSPTSSLRSSVQGRDLHLSDSELSDHTQTTSQDYTALRDNSDISQLPCNSESISSTPVCNTVPNTPDCNNITESRKNSEKSDLSESDVDNYFSDFKERHAPKSQVTVDPGLGPGQVQLPADRLGQGAVKLRFTVTKGNLEVDIFCAKGLPVNNNEIPDTYVKTYLRQGQKRFQKRKTRIVRKSLDPVYKEKINYLARDVHGRHLQFYAPWCGHCKKLEPIFTEVGLALRHTGIRVAKLDGTAYSSVMHAFEVRGFPTIKFIKGDKNYTFTGDRTKEDIVQFAKKADGPGVRFLSSLGKFREAKSEHKDEDTVFFVYVGNQDPQVSALLSAYKQIAQERKVQTYFYAGEIRILPEEIKASLKAESTVIVFKDDVYEEYE

>XP_023227959.1|Centruroides_sculpturatus_II-RIM

MISTNVMSFMKKIVKSGNSGGEEIREKPSAINKLKQTVTLGMTLGRDKPKPIGEDLAYLTAEERSILNRVWQKEKEFEKESQKFRIQSPQSTPQAVQIKEKAVEKPQTPVNVPSGPNCRICRKIIEPGEQRNNCDECGQVVCDDCASYSKTSANQSQGWKCSFCRRRQGQDRLGIEPPPGSGMHRVPSVRRMAEKLARTNKEVGIYGSSEKLFTKEGAILLEKRQQSTHDVINKEYASCLTPFNPDSKEMSNKNIEDESPRSLRKHLSLEKSQDGRTRRETKSSVKRQKSIEDRRRSGRTRNSSPVGRDQSKERYYSSETPSDRRSSSESQKGPIHPLPVHNPARVSSCGVCSLTSSHQHSHGTEQSSPSESSLDEMSQSPEYDKRRKYRVKRKSKIQRQKNYIEEPDSDPCGYRDERARSNATSSLESSSALESIDNGLDYWPKEYKNSSEKPLRQIASDAALYVRRFSSESSDISDVSGEYSTSDRSGKYISTSTKRQRSSQKHPRRRRDIVSPVVMDSSPSADEQIIEHRTPAKSSSLDDTDKSPKGATTQLRKSIPCVYVDSVADLQTSRRNHVQDHTEWDYQRRSSTGRALPAIPDGHSFLLPHTRTQSTHSLDLPREEVGRPERRASAPERENIKIVIDDVDSEQKGRYGRQTYLRNILIRRDPDDEGARTKGFGMRVMGGKNGDDGRLYAQISWIVPGSAVEKHGLKPGDKIIKWNTECLTDKTYEEVSSIMERSSDVVELLVEVSGRTKSFDSISSIKTKIAPKLIISTEGYKESDVSVVSPTRRKLPKTPDQLGQQHSTVCGEVGVQIWYNQDGNDLVVTVLAARGLRRRKSSSAALPRAFAKIRLVPHSAYNTMKTAVAEPTCNPEWSKTFIFPEVTSDDILESVFELSIWDHCPLGSNVFLGELQLPLRKADLQDCPEWYPLSAHQQTPVALSPTHKITPPSLVPSTPAHDIARRMLQRGTEEYSQSLTYDEIKDESSLPPENKNMQRSSSMDLTLLHPDDAWDRHPTQPSSGNQERKPSAPSVVTRVKSTPPSRRSSNNLVSQDSLPELYGTGPKVKERVRSASFRVYRSADSDSEKGDHKKLGQLVRRKLSRTLSLKSDKDKRSVALGSLIVPPEQRVSNLSSPELTPDSDTGDTQNRNPGQIIPKRFSGNALAPKLGEIKLGFVMTKGQLEVDVVCAKGLANNSNGNPPDTYVKTYLKDEDRQMQKRKTKVARHSTDPQYRQTLKYNASEVFGRTLLVMIWERQKGFEHNQALGATEIQLDKLELSKLTVGWYQLFPYSHIKAESTEST

>XP_022258855.1|Limulus_polyphemus_II-RIM(1313aa)

MISTNVMSFMKKIVKPSRDETGEASGPMNKLKQTVTTGLQEMSIAREQTTSSKTRKTATSRLTPQERSILRMIWQKNDDIEEEPRRASLARPAEPVKQTAPTRESCRICLKNISDGEQNRECDECGQNVCEDCASYSGDPTDSLQEWKCSFCRRRQGQDRLGIELPPGSGMNRVPSVRRMEQRAREAGKEGYFSSLDPGSDFDILKTSEEATDFKAFNSIDESVPSFNIVATNIKGLTKDQDREIRNKRRSKTLATKANHRRQVSLKRKDSRPTVEKKKSTDVSKRIKEKRRRSILTSTGEYLSRSSSPEKHARERHFSFENLSDRCSSSESTKGMITSKMLALKAVYRKDDSHLGSYLSHADFVQLNKHSSASGSSLEESSHKEEKERKRRSRTKRRSKIQRQNYYVREPDSDPISYRDENFPKANTKTSFESSSAIDSLEKDLNYESQRRHSMRVPTVLEKPLRQIASETALDKRRVSSESSDTSDISEDISTHSDRSGFDGSHEGYQKGEKVAVQNPMIQRDDSKNIPAHLQFVEGGYFSPTGTADSSLSLDEHSHLSSDKSFSLDDRDIFRTSTPHIRRSIPSVLVDSVVDHQRSASVSLLTSGHYIMDNPFPRRNSTGRMLPMLPVTDNQLGLYPASQSRSWSTHSLDLPREESGWAERRASAPEGENIKIVVDDVDSHPNGVRGKSSRQPLLRLVRLHRDRKVPVRGFGLKVKGGKFSDDGHLHAYIAWTVAGGSAEKKGLAQGDRVLEWNGVSLVDKTYEEVSTIIGRSPDTVDLLVEKVAWRRLSDQLGSVSSPQLKHASLKTDHDTKTDVYSTSPTRRKLPKTPNDNEQRGRVQVQIRYDPDDSSLVITLLSARGLQYCKNYSGKLPRAYAKVRLVPQLGFPSMKTRVAEPGSSPKWNQTFIFPSVSSEELATKTLNITVWDQISSGEKRILGESQIILRCEELQECPAWYYLTSLQMAPMKRSRTCSMSSDDLAQRQLQHNGTNQIICFRSYSDGKSYLDRKEESLVKNTSMDSSLLHPDDACCEERAVASDGATNMEIIPLKQNRDSKQAIDTKNKFCFHSQDNLSDLNSTRTKTTAKVKSSSFRIHRPSNLDKEQGETVEPMTLGKLVKNKLCRTLSMKSDKNQRSVGLELEKRSTRSRSDLTSENEIPIMTIGPNGIRRGSGQVFSRKIKFSGAETLGDIKLGFLITKGHLEVQVICARQLRLNTTGQPPDTYVKTYLKEGERQMQKRKTKVVRHSSEPQYRQTVKYSASNIQGRHLLVMVWERQKGFEHNKPLGAADIQLDRLDFSKLIVCWYPLYPILHEEMGSNESV

>XP_022240908.1|Limulus_polyphemus_II-RIM(1323aa)

MRWGVLSDPELRNVRVRLALNPSQQQQTPRDSDRCFISYSQASQKRSGMIVKTGTEEESERPSAITKLRQITTGLQEGRDRKRSCRAPQEELARLTAEERSILERVWQKEEEFEKETTKVSSGSPSLTVKEGASSSQENCRICRKAIGSNEQSRKCDECGQLVCEDCASYSKDPNDINQEWNCSFCRRRQKQDRLGIESPPGTGMHRVPSVRRMNQRMELLGREGSILVPELSVDLEEKIGLEGSPPFRLEKKESKRLAEESDRSFSLESRDLNMIQQSDQEHEIKQVFETKTSQVEKQKPIECIKESKIIRGSSRSTSRASSPEMKGVSRSKDGHLSLDFSLERRPSSEYRKGTRRCQTEHGLYTSSTSSHHQHHQKDSSQQRHSSASESSIDEHRRDEVDERKQRNRIRKRSRIQRQKYYVEEPDSDPTIYRVNILGRSDAKSSLASREIISGSAEALNRRLNYVTDHLTKKPLSLAGVDIANRHSSDSSELSDVSGDVSPFSDRSGHKRLHGPYYSGERLEVRTSKKHQKSRKPSVEFAKERTISYAGAMDSSLSSDEQELIRLSTDKSYSLEDPEMFLYPNVSLGKSIPSVFVETVPDIDSPPSPYPQNEEIIRRHSTGRVLPPVPVPENQLGLQLLPSGDRHSTYSLNLPYDEKGLHERRSSAPESEDIKIAINEVDSKLKTKSKPRLKKVRLHRDKTDTGARTRGFGMRVLGGKSAEDGKLYATVAWTLPGGQADQKGIHQGDIILEWAGIILIDRTFEEVSSIIESTGDTVEIVFQKESIKAGEDKSSRYSPGEPAQMSHHKTALGTSSVQRKPRQLLMLTPEPIMDTDVQPVSPTRRKLPRTPDQPLPPFFVGDVGVQMWFDRNRSDLIITLLSARGLRLKKTTADLPRAFAKLRLVPHSGFGPKETQLADPSTSPKWNKKFVFPSVTSDELLEKSVEVTVWNQSPTGNKSILGGTEIPLRKTDLQDCPEWYPLTIIQAPTQKPPHSGVSCHDVARQFMKQSGNYQGSTQSLSEEKCDKQSDNQNESVVKSSSMDFTLLHPEDAWDRLQQISPEDDTKLCPHTDTKSFRSTSPSGRNSGKLTSQDSLPAEIVREQNSQNKTRRTKFSRTLSLRTDKEDRSVVLDSKLASSSLSSPDLTPENEIPIMTIGPGDIRRGAGQVFSTKMKLDRVETLGEIKLGFLLTKSQLEIEVFCARQLPASSLGQPPDTYIKTFLKEGQRQMQKRKTRVVRCTSNPQYRQTLKYSASDILARHLLVMVFQRQKGFDHNQPIGAVEIELGRLDLTKSVCWYPLYPLPSEDFDSNEST

>evg198193|Mnemiopsis_leidyii_II-RIM

MNPNEKALSNKRKANRVNSEPDALSALRTKERMSASVSEGSNQEPDSQNPGFMTRIKGLKSDIMKRVSRVMSDEDMRNLRTIRTKLTNIEEPQRAQMSTAVQEVEDPNLKQKKSEQSLHDVYKQLYIRRITSVDPPSSHALKSSDELRKPHVDVKRRTFELPKAKVTENVTRPTSTKTEESDHEISYVLQALKEGKPAHEIACSRNIVQRTVKLERLKDFPVQREEQGGQFGLRIVGGKKMAGGTLCAFIASITPGSPASRSPDLHEGDLVLSWNTKMLIGLTYEECQACLDNCDFADLVVSNFLRGSYRRYRPRSSNTSRTGSVESILRGTTNKWITLTRRGILARQSGQSRGTDESNTIEESEKEEENVNEPAEEAKSSKLKLEIRLVHVAMENRLSVHLIQATNIPYREGMEGLDVFCKLYLLPDINQSADGKKRSKTAVRTYNPYWNQHFEYQPISESELSDKKLEVTLMMHHWLQKDQSLGQVEINLGDKKVLSGQADWFYLQPAPSTDLLNTSPGSREQSRDHSESDVTSPGLGSKSLSEMMREKVTIQHRTTFSSFDNKKEIVSDIRRTASDSTKLAKPLKQKSKQQQTSDEDSSDNVFNPAAIHTHLSNIKESDEQRPPIQVRGSPRRRPRVIRNKSSAKDHRKSDSITYSRQCSNSSNEIITPETQPNITKLRRAGRSNSEDLSVTKHRVLPKTPERSKRVSAMTLPTKRMTSPLLTAKTSQEFSNTRVILTAPSSEECGSSSTDSVSSHLTAIHRQQPEGRPLIGTGEPSSQLGPGQISVSYSVTVQGSLVLTLGIQKNMLTLEVVQANNLPLRPNGPAPDTYVKTYIMLDHNRSHKEKTTTVRCNCNPVFNERITYIVNPNNQIIQVMVWDENGPLRRNILLGEILIDLDYIASHGPLQGRYKLFPATFSKPP

>TR51711|c1_g1_i2-TR51711|c1_g3_i7|Beroe_ovata_II-RIM

MSDNEHTTSVNKREPTSVSDGVSELKIAGGGPPKTGQTRPKPSRSISDPCHGGRGGGKDSSRVRIPSSTTSRDQDPANSGIMTRIRGLKSDIMKRVSRVMSDEEMRNLKSVKTRGQENDEEGQGREREDRPGGAETALIGSQGDNMVDDPNIRHKKCEHTLHDVYKQLRYRRITGTIDSSPATTSPTLSRGSRSPVELVRRSPILRRKSPHNVKSTKEEEVPTKKAEETDLEVFNVLQALKDGKSTHEIECSRNIVERSATLSRSPDFKPHREDQGGQFGVRIVGGKKTSGGLLCAFVASIVPGSPASYSPDIHEGDLVLRWNDKILIGLTYEECQDCLDDCQSVDLVVSNFLRGSYRRYKPRTSASASRSGSVESILRGTTQKWLTITRRGMLARLPETREDASVNSGSIEEQPGIEDQLTDSTKIKVELKVMYVGMDERLSVHLIQAANIPYRERNEGLDVFCKVYLLPDTGADNCKKRSKTAAKTYNPYWNQHIEFQHITAKELAEKRLEVTLMMYHWLQKDQPLGQVEIDLSDRRALSGQNTWYYLKPPPSSSPSSSLECPSSPYGESRKSSLAGLADSRKSSQGSASMNLCDRKSSQNEGKKSYAIENRKISSDTSESGQTVSRYLSDLVKEKVSIQHRITFSTFDSKKDIVPDIRRSASDSTKPQKPPKQKTLAQQQPGGAGDDDSSDNVFSTAATIQSHLSNIKEYESDVERRPIDVRGSPRRKAKVIRHRSQTRDDTARFSDVVTRSSDPSDDLLRSLIETSTISPEPVAPSKLRSPPVRSNSEDLMTTKHRVLPKVPQPRDIRYPARIRCDPPAGLILPSKKPTSPSSSGSVAMTTTRPTAFSETKVILTAPSSEEWTGNSSSTDSELSSQKLNLRRPSPEGRVLIGTGDYSSDLGPGQVAVTSQGDIPGTLVLTLGIDKNMLTLEVVQANDLPLRLSGIPPDTYVKTYVLLDDHRSHKEKTTTVRCNANPVFNERITYIVNPHNKILQVMCWDEVGALRRNVFLGEVMIDLSYILNHGPLQGRYKLFQSCSAGVHSKPP

>evg158061|Hormiphora_californiensis_II-RIM

MSASAAADHKPAKRKEKRVNSEPDALAAVRRKENSEENDQSSTSGLFTRIKGFKTDFMKRVSRVVLDDDQVRQLRREHEERNAAAAAQPPSAPVDPNLKAQKTEQSLHDVYKELQFRKFMQLSDSTGQNTESPSSGGERPKSPDSERPEEETEVSRVLQELKDGKEEISCSRHIVQRAVTLRSSADFKPQRPTQGGQFGIRIVGGKKVSGGVLCAFIASIVPGSPASKSLQLQEGDLVLSWNGRMLIGMTYEEVLGTLDDENAATLVVSNYLRGSYRRYRPRSSNSQRTGSIESILRGTTNKWIAITRRGMAARSQELRVPECPEEERSAAGDQAAPAAAAVPRPELQDRSDTSITPAPHPPTAARCKLEMSVVYVTMEKRLSFHLIQAGNIPYREGVEGLDLFCKLQLTPEIGSCESKKRSRTAMKTYSPYWNQHFEFQPISEPEVRERKLSIVLLAHHWLHKDEILGEVDIDLSQKEALSGHASWFYLTKPIDQSADERNVSEASKEKSSSLQHRITFSTFDAKKVSADIRRSASDTCKPAGLKPIVSKRSGEQLSGSDSSDNVFSPYSVHSHLSNIKETEDGQRPPIQVRGSPRRRPRVAKERPATRLEKGRDSTSSAELITPAIPSQQSPPSAGDRPSDCSSSKPAHPCSASTCPAWSRTCRSNSEDFSITKHRILPTPPTHKGSIVMTLPSKKMTSPLLISSSSHNFGTTRVVVTRPEEQQCSSSESISMKTLKRPSPDGKSWVGTGDDPGNLGPGQLSVVAVPSGVSGSIVLTLVIEKNMLILEVVQANDLPVRPSTGLPPDAYVKTYVLMEQHRSHKEKTRLVRGNCNPVFNERITYIVNPSNQILQVMVWDEVGALRRNTLLGEILVDLNYVYQHGPLQGRYKLYPALCPAEKS

>KXJ21887.1|Exaiptasia_pallida_II-RIM

MKSSLDRLRRDLRLEIIHLEERLSHSERTSQDVDLCKICFKAKVADCLSLQCKLCQRKTCSRCGVREVCEKQIEQPSKSSLQADPEPEYLSLPVDRQSIPRDCDSVILSVEDSDCCTSAYFSPSPTPSPTPSEEFYPGIHALNMAGAIEYDSPDSSYSTGHPSDMTTHQGRPIIERLTLTKYNASTTTEGLPFGTRIVGGKMSDGVVIGAYVTQINPSSKASTSLNIGDQIIEWNNISLVDSTFEETQEAINSSPNSVTLVVGHIKRKKQTLLRPSSSQIINDAIEKTRANFAKEGTLCLSQEQLLSRLEKNRSTDSIIKEQALCNRYGLQSRIHISLWHDPERNNLIIKIIRAQSLSSGASVRVEYPNPYAVVNLLPRKTLSTQFKTNVEYQSVHPIWNLTFIYSNITKEELLYKSLEVSLWSERSTKVPQFIGAVFVELSQDVLSNIPQWYNIKNQVDVLHNITIPTSRRFTVSSSHDVINSIISTSKEKEHFVEISRTSRNVRSCIVRQRSPRDVVDKDNTFVYNKRRKTDSGATASYLVRSRTPSKQSIKDDAFDGEITMEDTFRGRGNSILSCKGLDTQGFPRTHSAPATPCGKTDFHPMFAYSPRFNCPFWQKSRDDVASCTNSSQGSLRGSPLWDKTTSPRSEKTTVSEERENNNTKMLPAVYLTAAIDNESNTSEASLQEDRKKKRLFFRFPNPDFSSESSYSFDDGPQDNETGPNQICTSTDGGQDLGYVRIGLVIENSEVLVVDIIRAKLLSFKSEQQGGRIPHLDTYVKTYLLNDGTKTCKRKSSTVRQDSNPVYNWQVKYSTEEISKSMLYVRIVIFDNHVRGRIQGDHGLPFAD

>evg1176111|Trichoplax_adhaerens_II-RIM

MCLECLRGIFSTGSKRVPSQQKSDQCCGMMVSSLAKLFKKKARPIISRRNTEQENVLEKEIPYITPDLSHLTVHELKILKEVLEKQSKFEADTDEFKRQLNDKVRDWDKRLTAPGKDRNRCQNCYTTKFVDGHGRICHNCKSTICGRCGLYTPSSPDPKQSKTAWRCKVCWYNMEFYHKTGFWYHGSRSELVVNRRELMGSVMENLKSGLSSKDDINHESKLPGLDDESKWILLRHNDNHRPYPWCMGILILGGVLVGKDTTYAVIDNMERETELEYESKMTIGDIVISWDGHTLINKSFEEVRSIVSTTADAVCIRVCRATESAFVQDLRMIPDTMTGENNFIKIVLSSSWETFNMEDTRKVSTENTKKTLLSVQDTSADDMESSQEITQEAFNDNPRNGRIKLSVKYHDSGRLEVYIAEAADINPDEITSDSIFYVQLCFITQDGEISEVWTSLSMDDSKSTKLEWNQLYSCTINHNELLKTSLAVILWQLSSNITKVREVIVNPERAALEVEPQWYQLYPHDDNFFELPSPPCSEKVHEPEMKNIQHAIPEAIVQDNDLTDHPEQVQKVDVKASAKLREVIAVKTLRKASNLASSSQGMTRNPGEDPEYKKDLFQRRRSSLDGVACKDVSETLALSYNDAQRADRTDSANISESGIMENYCFSGIRSTSHVVPRSDSVSSFCSYQDMTSTTDYSDTEVKKQESTSGTRFRKKRGRTNSKSSLLERLLLKGRLTENNDKIFSCPPSPNSERYRRTSKNSVDNSPIPGDATPVRTRWQLNSEVIEDRLEAISQQNLGPGQIAANSGQFGVNISRNVGALQLTIREVENSLQLHIIRAVNLMATVVLPDTFVKIYLMEGGIHNIKNKKRNIVHKAKSRSVKNTKSPIYDFMLSFSNYKKSQVLYLTVWSMESSNRSKKRLIGDITIALDLVDLQYCVNRWYCLMSQQTTEKPIPHTRPRLSIASTNSMSSIDAPVHSDIFPTRDQRKQKLKQQRDRYRTKASSLSEPRTSQTIDNQGKDSAETSEADKISAMESDNSYLERDLYMLQSTGATADTTEVSVHGQEDKISAVSETEFVLPPRSSSETLPANATTDDEQWF

>NP_001137326.1|Homo_sapiens_rabphilin

MTDTVFSNSSNRWMYPSDRPLQSNDKEQLQAGWSVHPGGQPDRQRKQEELTDEEKEIINRVIARAEKMEEMEQERIGRLVDRLENMRKNVAGDGVNRCILCGEQLGMLGSACVVCEDCKKNVCTKCGVETNNRLHSVWLCKICIEQREVWKRSGAWFFKGFPKQVLPQPMPIKKTKPQQPVSEPAAPEQPAPEPKHPARAPARGDSEDRRGPGQKTGPDPASAPGRGNYGPPVRRASEARMSSSSRDSESWDHSGGAGDSSRSPAGLRRANSVQASRPAPGSVQSPAPPQPGQPGTPGGSRPGPGPAGRFPDQKPEVAPSDPGTTAPPREERTGGVGGYPAVGAREDRMSHPSGPYSQASAAAPQPAAARQPPPPEEEEEEANSYDSDEATTLGALEFSLLYDQDNSSLQCTIIKAKGLKPMDSNGLADPYVKLHLLPGASKSNKLRTKTLRNTRNPIWNETLVYHGITDEDMQRKTLRISVCDEDKFGHNEFIGETRFSLKKLKPNQRKNFNICLERVIPMKRAGTTGSARGMALYEEEQVERVGDIEERGKILVSLMYSTQQGGLIVGIIRCVHLAAMDANGYSDPFVKLWLKPDMGKKAKHKTQIKKKTLNPEFNEEFFYDIKHSDLAKKSLDISVWDYDIGKSNDYIGGCQLGISAKGERLKHWYECLKNKDKKIERWHQLQNENHVSSD

>NP_001289273.1|Mus_musculus_rabphilin

MTDTVVNRWMYPGDGPLQSNDKEQLQAGWSVHPGAQTDRQRKQEELTDEEKEIINRVIARAEKMEAMEQERIGRLVDRLETMRKNVAGDGVNRCILCGEQLGMLGSACVVCEDCKKNVCTKCGVETSNNRPHPVWLCKICLEQREVWKRSGAWFFKGFPKQVLPQPMPIKKTKPQQPAGEPATQEQPTPESRHPARAPARGDMEDRRPPGQKPGPDLTSAPGRGSHGPPTRRASEARMSTAARDSEGWDHAHGGGTGDTSRSPAGLRRANSVQAARPAPAPVPSPAPPQPVQPGPPGGSRATPGPGRFPEQSTEAPPSDPGYPGAVAPAREERTGPAGGFQAAPHTAAPYSQAAPARQPPPAEEEEEEANSYDSDEATTLGALEFSLLYDQDNSNLQCTIIRAKGLKPMDSNGLADPYVKLHLLPGASKSNKLRTKTLRNTRNPVWNETLQYHGITEEDMQRKTLRISVCDEDKFGHNEFIGETRFSLKKLKANQRKNFNICLERVIPMKRAGTTGSARGMALYEEEQVERIGDIEERGKILVSLMYSTQQGGLIVGIIRCVHLAAMDANGYSDPFVKLWLKPDMGKKAKHKTQIKKKTLNPEFNEEFFYDIKHSDLAKKSLDISVWDYDIGKSNDYIGGCQLGISAKGERLKHWYECLKNKDKKIERWHQLQNENHVSSD

>NP_598202.1|Rattus_norvegicus_rabphilin

MTDTVVNRWMYPGDGPLQSNDKEQLQAGWSVHPGAQTDRQRKQEELTDEEKEIINRVIARAEKMETMEQERIGRLVDRLETMRKNVAGDGVNRCILCGEQLGMLGSACVVCEDCKKNVCTKCGVETSNNRPHPVWLCKICLEQREVWKRSGAWFFKGFPKQVLPQPMPIKKTKPQQPAGEPATQEQPTPESRHPARAPARGDMEDRRAPGQKPGPDLTSAPGRGSHGPPTRRASEARMSTTTRDSEGWDHGHGGGAGDTSRSPGGEQGLRRANSVQASRPAPASMPSPAPPQPVQPGPPGGSRAAPGPGRFPEQSTEAPPSDPGYPGAVAPAREERTGPTGGFQAAPHTAGPYSQAAPARQPPPAEEEEEEANSYDSDQATTLGALEFSLLYDQDNSNLQCTIIRAKGLKPMDSNGLADPYVKLHLLPGASKSNKLRTKTLRNTRNPVWNETLQYHGITEEDMQRKTLRISVCDEDKFGHNEFIGETRFSLKKLKANQRKNFNICLERVIPMKRAGTTGSARGMALYEEEQVERIGDIEERGKILVSLMYSTQQGGLIVGIIRCVHLAAMDANGYSDPFVKLWLKPDMGKKAKHKTQIKKKTLNPEFNEEFFYDIKHSDLAKKSLDISVWDYDIGKSNDYIGGCQLGISAKGERLKHWYECLKNKDKKIERWHQLQNENHVSSD

>XP_015131024.1|Gallus_gallus_rabphilin

MTDAVVGGSADRWMCPGDRTMSLRASEKEQIPAGWAARGGQPERQRKGEELTDEEKEIINRVIARAEKMEEMEQERIGRLMTRLEDMRRSVLGDGVNRCILCGEQLGPRGSACVVCEDCKKNVCTKCGVETTNSRPHPIWLCKICSEQREVWKRSGAWFFKGLPKQMLPQPMPVSKSKVPPAPSEPSPAEPPAPDPKVPSRTPGRGQADEQDHCGSEVTMAARVRKPAEGRTGPCGSEDTESREPTTESGVSRSPGVKRANSMQSNSTPPPRPPAVTAGPAAPAAARPGPGAAGRMLETQASPPPGPPEPARAAPKEERAGGYAAPPARDERPARPPGAPQPPAPLRQPPPEEEEEDANSYDSDEGTTLGALEFSLLYDQENSALHCTLIRAKGLKPMDSNGLADPYVKLHLLPGASKSNKLRTKTLRNTRNPVWNETLVYHGITDEDMTRKTLRISVCDEDKFGHNEFIGETRVSLKKLKANQKKNFNICLERVIPMKRAGTTGSSRGMALYEEEVDRGGDVEERGKILVSLMYSTQQGGLIVGIVRCVHLAAMDANGYSDPFVKLWLKPDMGKKAKHKTQIKKKTLNPEFNEEFFYDIKHSDLAKKSLDISVWDYDIGKSNDYIGGCQLGITAKGERLKHWYECLKNKDKKIERWHTLQNENHVASD

>XP_006817123.1|Saccoglossus_kowalevskii_rabphilin

MAVRSGGIKKWTCPDDRQLALRAKLGTGWSFHTNEASRYRKNEGLSREEQDSIQRVLERADRLEKMEEERIGRLVEKLDNMRKNALGNGNTQCVLCGDEFGLLGASPLECHDCGKAVCTKCGVDTTNSQRDAILLCKLCSEHREMWKRSGAWFFKSIPKYCLPEKKMDTQKYQTVGGRRIEAKTRNRKPGGGRNYNTWSRAGYASESESESSSSSDDEVSIGRKRKPKRHDSDLSRTSSVRSATSASATIEEDDVDRAFNHYGTNDIRGSGRGYHSQHPSYSQHQTPYSQQQQQQQHQPAYAQQQSPYAQESSAPHSPVKEREITPPPQEVLLESSPDDDFGTHLGTLEYSALYDGINNALHITILRAKGLKAMDSNGLSDPYVKLHLLPGASKEIKCNSNFIAILKAANAWTEILSVLDEDKFGHNDFIGEHRLPLKKLTPHQTKNFNVYLEKPLPLEKDDELASVIRGKIXVYLKYVTRQQQLVVGIVRCVILCTCSVCSYLKPDQGKRTKFKTPIKKKTLNPEFHEEFVYDVKLSELAKKTLELTVWDKDIGKSNDYIGGIQLGIHGKGDRLKHWFETLKNPDKKFERWHTLSEEHFDDA

>XP_030848070.1|Strongylocentrotus_purpuratus_rabphilin

MGEFAGGNAVDRWTCPNDRQLALRAKLGTGWSFHTNSAKKFQKSEGLNRDEQDTILKVIEKAEKLEVSEQERIGRLVDKLDNMKKNSLGNGTTQCILCGDEFGLLGASPMTCYDCYKAVCSKCGVDTTNSIKQPIMLCKLCSETRELWKRSGAWFFKALPKYTVPEKKSEMHHNRYPGSFRRTEAKPIRRRRADSESSDASSSDDDVSLGRKRRAKRDGDQDDTSISSSGLSSANLLGGGSKNDYPGTLYSSNHMAGSRTSVGSGGWYGSANDSSVGDAGHDDSDSERSTPGATLGKDSSLGTLEFSVLYDGINNALHCTVTKARGLKAMDSNGLSDPYVKLHLLPGATKSTKLRTKTVAKTLNPDFNETLTYYGVTEDDLSRKILRLSVLDEDRFGHNDFIGEYRLPLRKLTPYQTKSLSVYLEKPLPLEKDDELAGERGKLMVGLKYVSTRQCLVVSIIRGAGLAAMDSNGYSDPYVKVYLKPDAGKRTKHKTAVKKRTLNPEFNEEFYYEVKHPELAKKTLEITVWDKDIAKANDYIGGVQLGITSKGERLRHWFETLKGIDKKYERWHTLSDESFGDE

>XP_022085716.1|Acanthaster_planci_rabphilin

MPINGAATNMGEFVGGSAVDRWTCPNDRQLALRAKLGSGWSFHTSASKYKKSDTLSREEQETILGVIERADKLEVAEQERIGRLVDKLENMKKNALGNGSGKCVLCGDEFGLLGASPLTCHDCYKAVCTKCGVDTTNSFKHPILLCKICSETRELWKRSGAWFFKSLPKYIIPEKKVDQSRLQGWRFETRKSGRKSTTGRVNNTWSKADVSESESEYSSSEDEVSIGRKRKPKRETNTDDNISIGSKNGMASSIYGNNYMAGSRTSIGSAGWYGGASTTESSHDAARDESDSERSTLGDNEHSRTSSVRSLTSTNISVSGSLGTLEFSVLYDGVNNALHCTIHKASGLKAMDSNGMSDPYVKLHLLPGATKSTKLRTKTVHKTLDPKFNETLTYYGITDDDLMRKTLRLSVLDEDKFGHNDFIGEYRLPLRKLTPHTTKNMSVYLEKQMPLEKDDELMGERGKILVSLKYLKERQQLLVGILRCAGLAAMDANGYSDPYVKVYLKPDSGKKTKHKTATKKRTLNPEFNEEFFYDVAHNDLAKKTLEITVWDKDIGKHNDYIGGIQLGINAKGERLRHWFETLKHTDRRFERWHTLSDEIFAGDD

>XP_012945559.1|Aplysia_californica_rabphilin

MGEFVSSGGVDRWVCPNDRQLSLRAKLGTGWSVHTNRGGSFGRPDQLSLEEQEQILGVIGRAEYLEGVEQERIGRLVDKLDNMKKNAIGNGNSQCVLCGDEFKLLGASPTYCDDCAKAVCSKCGVDTFNCNRQPLWLCKICAETREVWKRSGAWFFKGIPKYILPEKKAESGKYPVRMRPSPRGSIKGRPAPGAGGVSRDSYTWHKARGSASTASYGESSDQESSESSEEDFPMGKVKAKKSVDSGNPASSPKSDNISIGSSASRNAPHGNHQTYTRAESKTSITSATYMGGQLSATESSRGDDLHDETDSERSSTYDFGRPTALSIANPNFAFSREGSTRSVGKASLGTEEELDNIDMAINRYVKASHAVDEEQLSSPDSEGTSLGLLEFSLLYDSVNNALHCTITRAKGLKAMDSNGLSDPYVKLHLLPGANKSTKLRTKTIHKTLNPEWNESLTYYGITEEDMIRKTLRLSVLDEDAFGFDFIGEFRLPLKRIKPHQTKNFSVYLEKQLPMEKDDDLIQIRGKLLLSLRYATAKQTLFVRVMRCAELAAMDSNGYSDPYVKVYLKPDKDKKSKYKTMVKKRTLNPEFNEEFQYEIRHNELAKKTLEITVWDKDIGRTNDFIGGVQLGINSKGERLKHWFDTLKNPDREYRRWHVLSAELTPEPDD

>XP_011424844.1|Crassostrea_gigas_rabphilin

MGEITGSGAMDRWVCPNDRHLALRAKLGTGWSVHTNKLSTFHKGDQLSMEEQEHIMNVIAKADYLDQVEQERIGRLVEKLENMKKNAMGNGQTQCILCGDEFGLLGASPTHCDDCKNAVCTKCGIDTFNSNKQQLWLCKICSEYREVWKRSGAWFFKGIPKYVLPEKKLETSKLAGQKGHKEMRGSMRSRPGSVRGYNTWSRGRASHGQSSYGESSEQESSSSSDDEVSIGKRKTVKHSGDSGESDNISVTSSASGQYYGNHVNKEPRSSITSSNYYGGQLSATESSRGDDVHEDTDNESLGGGTIHHATNGSHDPCPPSKLADEADIDDAFTKYGKNHPHSEHEDNLSSPESPTNGSLGSLEMSLLYDAANNALHCSIVRARGLKAMDSNGLSDPYVKLHLLPGASKSNKLRTKTIHKTLNPDWNKTLTYYGITEDDMYKKTLRLAVLDEDAFGFDFIGETRVPLKTLQPHQTKHFNVCLEKAIPTDKDDDLLSHDRGKILLGLKYSTAKQCLVVNVVRCVELAACDSNGYSDPYVKLYLKPDPEKRSKFKTAIKKKTLNPEYNEEFIYDIKHNELAKKTLEITVWDRDIGKANDFIGGVQLGINSKGQRLKHWYDTLKNPDHKFERWHILAAELIPEV

>XP_021345406.1|Mizuhopecten_yessoensis_rabphilin

MDRWTCPNDRQLALRAKLGTGWSVHTNRLGMFHKGDNLSIDEQEHIMNVICKADFLERNEQERIGRLVEKLDNMKKNAIGNGTTQCILCGDEFGLLGASPTFCDDCKNAVCTKCGIDTYNCNKQPLWLCKICSENREVWKRSGAWFFKGLPKYTIPEKKAMEATKVVTTKGRSESRGVQRGRPGSTRNYNTWSRGRGNHAYGESSEQETESSSEDEISIGKRKTGRNRGDSESDNVSVTSSASGQYYSGHNPKETRGSVTSSNYHGGQLSATESSRGDDLHEDTDSESVGEFSRQGSTQSHDLPVIQARGRQEDDHNIDNAFSKYAHNPHQEEYEDGPPSSPDSDGVTLGALEFSLLYDSAANALHCTIVRARGLKAMDSNGLSDPYVKLHTLPGASKSNKLRTKTIHKTLNPEWNETLTYYGITEDDMLKKTLRLAVLDEDAFGYDFIGETRVPLKRLRAGQTKHFNVYLEKQLPLEKDDDLMSNDRGKILLSLRYISAKQALTVGVVRCVELASMDSNGYSDPYVKL

>XP_014779575.1|Octopus_bimaculoides_rabphilin

MDDPNSQSNTERWVCPNDRQLALRAKLGTGWSYHTNRYTQFRKDDQLSVDEQEQILNVISRAEFMDNVEQERIGRLVEKFDNMKKNAMGNGDNQCILCGDEFGLLGASPTYCDDCKKAVCIKCGVDTFNSQKTPLWLCKLCSESREVWKRSGAWFFKGIPKYTVPKHHHEGGKFSSNARNQLETSAHQKSRSGSFRSYNTWSRGRHNGNKEGFGGESSGQDASSSSEDEVSIGKRTKESDNISIGSSQSSRDHQGGNGKVISSNTNVHGGHLKSTDNNQDFHNASDNEKTGTTGPYQKGSQLPEDRKGSTADGDIDVAFNHYSQQITDLENGNNSCEMLGILEFSLLYDADNNALNCTVVRGKNLKAMDMNGFSDPYVKLHLLPGASKANKLRTKTIHKTLNPEWNETLTYYGITEEDVAKKAVRLSVLDEDPIGYDFIGEYRISLSQLKRNQTKRFSVLLEDHQPLEISDSQSVERGKILLSLCYKPSEQLLLIGVVRCANLAAMDSNGYSDPFVKIYMKPDPGKKSKFKTETKRKTLNPEYNEVFNYNIRYNELTHKTLEITVWDKDIGKPNDFIGGLEFGSQSKGDTLQHWYEMLRNPDQKVEYWHTLTPDFCP

>XP_013406958.1|Lingula_anatina_rabphilin

MGEFTSGGGIDHWVVPNDRQLALRARLGTGWSVHTHKLQTFHKNQQLSVEEQEQILNVIRRAERLELMEQERIGRLVERLENMKKNAIGSGDSQCVLCGDTFGMLGASPTYCEDCGKAVCTKCGVDTFNSHHQPLWLCKICSEKRELWKRSGAWFFKGLPRYIMPKKKTELDKVPGRGGHPGQGQRSRTGSPAPQYNTWTRGRVQYGESTDEQESSESSDDEINISKRKVRRQQDSESESEMSTTSDFRPWASRGSMGTVESDSVSIASSGRPNPLMASRPLSDSKTSLDSGQLPSSGQGTGQGSLPSEGPRVQESQDSTVSDSERSATSGVGSFSRGNSLQSADTGSIKEKHEEEADIDDAFTKYGKHEGEDEDQLSSPESDVSLGTLEFSLMYDSMGNALHCTLIRARNLKALKAMDSNGLSDPYVKLHLLPGASKSTKLRSRTIHKTLNPEWNETLTYHGVIEEDLQRKTLRLSVLDEDAFGYDFIGETRVPLKCLKPHQLKNFNVYLDRQMPLEKDDDLVAHERGKILVSLTYKASKQSLLVGIVRCVQLAPMDSNGFSDPYVKVYLKPDKEKKSKQKTTVKKKTLNPEYNEEFLYEIPKHELAQKTLELTVWDKDIGRHSDYIGGVQLGITAKGTRLKHWFETLKYPDKKHEYWHILSADVVPDSP

>XP_013782034.1|Limulus_polyphemus_rabphilin

MGDFKSSRDKWVCPNDRELALRAKLKSGWSVKTNSANFFHKPEQLNDSEQETIMNVIKRAENVELKEQERIGRLVEKLENMKKNAMGNGASQCILCADEFGILGASPLFCHDCNNAVCTKCGVDFASSQKGQQWLCKICAETREMWKKSGAWFFKSLPKYTLPEKKTDHNKYGGGKGVVGAGSGGRGLISWSHGKAGLESSERESTDSSDDEVKVSRAIRRPIIHNVVHESTDGSESGQNGKTSATVAPPDTPRSTIVIARPLGIERYGEESHIEGSDNGHSMSSEGATANSSRVLLDHDTYRSVSHDSTDHVPYSPSHWRHARDWSRYGMEEEGNADDVHTQFESCNRQDSQVSVVSTGSVGSVGPYIRYSPTPSGYSRDRDSVTEEVSLDERLSPEDTSNLGSLEFTLMYDPSDQALHCTIHRAKGLKAMDYNGFSDPYVKLHLLPGASKANKLRTRTVHKTLNPEFNETVSYYGITDYEISKKTIRLSVLDDDIFGNDFIGEVRFPLKRLKPHQSKHLNVYLERPLSIDRDEEEEMERGRILLSLMYSNDRLGLTVGIIKCAHLTARDANGYSDPFVRIQLKPDPLRRKYKTSTKKKTLNPEYNEEFIFDIKPQDLCKKTLEVTIWDKDYGKPDDYIGGLQLSIHSKGDRLQHWVETLKNPDRRIEQWHKLSGVLLLH

>GAV03979.1|Ramazzottius_varieornatus_rabphilin

MSFRAASPGTMAGSRQDPWVCPDDRQLALRGRLKAGWSVHTGKASAVAKPAAPATLTCDEQEAILAVIKRAEAMEQQEQERIGKLVDRLDHMRKLTVGNGITHCAYCGAEFGQFFGGSPQSCKDCSQAVCSKCAVNTSDSRRRSIWLCKVCSEGREVWKKSGAWFFKGIPKYVVPEKKENGTIGNKRITALDAEPAPSALTTAHTGSSPPRSPMLAPPSTSPTDRRPSLTKVWSFITKSREELTPPSPPIANTASSSAKQADGRKSAPSVNPDEAIVIRRSFSRKAQDTDSESGLSTRGSIDSTASEAVSEIRIGRSSFRRRRPSTVMTDSPKLSTSSRLSESPSPPPLLPETSSAAASVPTRVITTSEPDNPSTNPSSTTVLPVPTSPRPFSLSRKSLRSRKSKMNGASMEAEESVMDSPEDDLSALGYLEFDILYDSNQCTLQCRIIRARNLRAMDRNGFSDPYVKLHLLPGASKSNKQRTKTINKTLNPTFNETLTYYGITEDDINRKVLRLAVLDEDTFGHDFIGETRVQLKRLTPFEWKHFDVVLEKRLPTGDKPDDIIDERGRLLLSLMYSVKRQALVVGIVRCAALPAMDHDGFSDPYVKIYLKPDPLKKTKNKTNVKKRTLNPEFNQEFTYPIKLGDLLKKTLEVTVWDRDYGKANDFIGGVQLGINSKGEKLKHWYEMLRNPDTKYEKWHGLVNEEFHDSS

>OQV25295.1|Hypsibius_dujardini_rabphilin

MASLKPPTTPTGPSSQDPWVCPDDRQLALRGRLRTGWSVHTGKGTSAPKKSAPTTLTDEEQEAILAVIKRAETMEQQEQERIGKLVDRLDNMKKLTVGNGVTHCAFCGREFGRVLGGSPQSCKDCGQAVCSKCGVDTSDSKKRTVWLCKVCSEAREVWKKSGAWFFKGIPKHVMPEKKEGNGNIGNKRPANVEPPPIPMTSTTSTSSPAHSSPPKSPLFATPNTPSHRRPSMSMVWSFISKSKDESSAPPSPPINSNSSGTATLSLGQKQSASPEDSEDGPVVIHRSFSRRRHHESDSESGMSGRNSVDSTASDASKTVSEIRVGRSSFRRRKNTTPAARAAAAAAAAAQQVADPTAPTLMPPPMPDLPPFVEPPSSSSTSIELSPARPNLSAPTPSAAVITRAQSVTSSKNKHNPPSRLNGNGIEAEESTVMDSPEDDPSSLGYLEFEIQYNSAQCTLQCNIIRARNLRAMDRNGFSDPYVKLHLLPGASRSNKQRTKTINKSLNPVFNEILTYYGITEDDINRKVLRLAVLDEDVFGHDFIGETRVILKYLPVMEWKKFDVVLEKRIPTADKTDDMLDERGRILLSLLYSAKRQALVVGIVRCAALAAMDRDGFSDPYVKIYLKPDPLKKTKNKTTVKKRTLNPEFNEEFTYPIKLGDLLKKTLEITVWDRDYGKSNDYIGGVQMSINSKGEKLKHWYEVLRNPDTAYVKWHGLVDEEFHDSS

>NP_572651.1|Drosophila_melanogaster_rabphilin

MDFQNRNNTANKFVCPSDRQLALRAKLKAGWSSSKTSEPLRPEEQEAIISVIRRNEEIEVAERQRVGRLVERVEKIKQHAVERGPNCCRLCGDTFGILRPQRILCEDCRQSVCTKCSVDINIRYHTSERSREIWLCRICSETREMWKKSGAWFFKGLPKYDMPRSASATPIPNPGPGSMAGDTRAVQSCHATPTRPARVKKLTIRVNDSSSSSSGHSEPEDEVDTGVGIGVGVVGGGMAKGIASTRLQREDSFRLRAYGSIRSFIDGGERKLSNSFFFNRQQSQRPSECPEYDSVSMSKLRRESSFLRRGSVSSSWSISDSSGSGSNNSGNSQTQSQYQQHQQQLQQHCRDPLLGWLEIAISYREAFHSLDCTMVRARDLPAMDAAGLADPYCKLNIITPEAHTKYTRWQRTKTVHKTRNPEFNETLQFVGVEPEELGNSLIYVALFDDDKYGHDFLGAAKVCLSTVHSTSQYRISVPLGVEDQYSNAAEMAQNWPNGKMLLSLCYNTKRRALVVNVKQCINLMAMDNNGSSDPFVKIQLKPDAHKNKKHKTSVKWRTLNPIYNEEFYFEASPHDLNKEMLILTVWDKDLGKSNDFLGSLQLGAQSKGERLQQWLDCIRLPDHFHEKWHCLAPDNPAH

>XP_018018219.1|Hyalella_azteca_rabphilin

MQRRNSAGSLTASCSERWVCPSDRSLALRAKLRMGWCTGQASSPARPEGLSAAETAAILEVIQRAEQLDMAEHQRIGRLVGRVTAIREKAQGDGKASCILCGERLPLLPPASICHHCRQGVCNKCAHEQLSGTEKVLLCKICSETREMWKRTGAWFYRGLPQKLLPSGGEGNGRLANAGSQSLGGSSISIALLGRHSKAPPRRAHSLLSTDLTSLLRIGDKESDSESGSRQLQHFPGSPSLSSSLSPAASTSSVASLAASHCSPRLPRNSPHHHQQNHEVNQLSQQTHLQQKKTAGFSDIFGGNVVPEEKLENGENGTADSALFEKLVSDLNAKKNTAGCAHGCNGAVAFDYDSDSSDAPEEDHARHGRENGCVTSALRNDVTAAAPSDWNEAQDLQFCASETPRTILSAKSSKESAAPSDWNEAQDLQFCASETPRNILSAKSSKESKDSFDSPVFTTSVEDETSSVSLDWCPETRETLQRHGSITSSVCSSSSNSTLPLGHNKGNSNNVGVPATAGDRSKPSKDNLGSLEFTLFYDSHHQSLHCTIHHAQNLKSRSSTGSADPYVKLHLLPGASKSNKLRTRTVPKNLNPEFQETLTYHGISTDDISNKTLRLLVMDEGRLHRYFLGEARVALKRVKAHESRRLSVQLSDKINGSDDQTSTGQELGRLLLSLRYTSARGALIVGIVRCAHLRARDKNGYSDPYVKVQLKPDPHKRKLKTAVKWKNLNPEFNSEFTFEVRRNELPKRQLDVRVYDKDVGRSDDFIGGLVLGHDSRGPELRHWYDVLQYPDRRHDRWHNLRPMHSC

>NP_001022566.1|Caenorhabditis_elegans_rabphilin

MFSRRTTPSPSITTASSSTFSISNLTNNNATSTSDLPASAISNIVPQIPPTPRRVPPKIGLLRHLSGFLESKKDLMNDWEIGGTQNKWVCPSDRHLHLRAQLKSGWSVRTATARSPTNSKAQTGSITAAEQEHIQKVLAKAEESKSKEQQRIGKMVDRLEKMRRRATGNGVTHCLLCHTEFGLLASKSYAAMCVDCRKYVCQRNCGVETTDVNQTTGKVETVFLCKICSEAREVLWKKSGAWFYKEMPEFQRPDDRLPYYVPVTTNGTLPNASSAATPLSGTPGGAGPQPMTMPSTSSCQMTTPKWASPGVCNSPGLQMNGGPTSPLPNGTRRNTGHGGIEFPSSSRPSICSVLQAIEPLDRSKSPRPRIQPRWVNEKVMSSMSVDDEEKAASSSDGESFVQSGVPRRALNNKTPVGSTSATTSPAPPPTSTTPTSRREANMERFSRHTHAHANRLYSTDDDDDSSPESRPSTRSTSPRHSLATPSSYAHDTCHDTSLPDADTRSIDSGVVQSDHSNPQQSGLTCSSSSLTPLQQQASHDHHSGGGTPRRISNPDRTTSRVAQSASGTSLVTPPPPISSRTSPDNCNSSPLNVMEHKSSSASTASSGGNRRVGSAEPVLNNHHAMHNNQNHNDINKKLISQTSRAESPLAASSSFLSSPDDDTKQKNRRRDGVGRVNSLQLRTSLDDVAPPVAPISKMNGHIVSSEPTSSTTSNQNHTSVPIPTVPVVPEEEEEKAITASTESASEPGSLGSITLTLTYHSADKKLKMHLIRAKNLKAMDSNGFSDPYVKFHLLPGNTKATKLTSKTIEKTLNPEWNEEMSYYGITEDDKEKKILRVTVLDRDRIGSDFLGETRIALKKLNDNEMKKFNLYLESALPVPQQTKEEENEDRGKINVGLQYNIQQGSLFININRCVELVGMDSTGFSDPYCKVSLTPITSKAHRAKTSTKKRTLNPEWNEQLQFVVPFKDLPKKTLQIGVYDHDLGKHDDYIGGILLSTSAKDERGRQWIKCIENPGTLVEAWHRLELDS

>XP_012797599.1|Schistosoma_haematobium_rabphilin

RLGTGWSSRAQSAQRRRAQPLTTDEEEQILRVLRRNELLAENEKARVEAMLQKLDRLKQTADSRKNEECALCHKEFGHLSNLPIVCVECGRYVCTTCCVELVIPPLYRQPITKRNSLLNFQWMSNTYGSGNLLNISKHQTYFTMDQTRQNGNTTLNNSNNLKRLTNDTSKPRFLNRRSSLMGTLNSAALHSQQLFEKVKVKANSRQRDNHSFIWKRSGAWFYKSLPRSRMTSCPSSPSSTPANNNCQIFNDITDLVKPLQINTNNESHIEHTDNESEQWIQLKPNRRPSKPEGSEADSSSLPSTSKDTPQNEVTSSTKTRGISKGFPKSPEHSSYTPPDQSTVDVPKSNSPTYTNVPATTNSQDITHISPASCLLSDSLNTSSLWRPQSPVSAQSFHPSFVSNKSVNHSTASALASSFTNANSTQKVASLTGEKEPSNLLSPSSGATSTVRRSTVSSSEEAPFGVLYFTLYHDTLNKQLHVAIHKAKNLIAMDANGLSDPYVVCQLLPTSHNSTTPRTSTRPQCLNPVWNEALTFEPFDGKNIQLKTLRLAVLDEDLYGSDWLGEYRLQLSQLIPNRLTDFSVPLGPHKPIQRGEFDLACPTRGKIQLGLGYLEDRKQLYVEVIRCANLAPMDLNGFSDPFVKLYLRPDKTKKTKQKTQVKKATLFPEFHETFFYDLNASEAGSRTLEITVWDFDHGPSNDFIGGLTLGAGAKAERRELWLAVFRPPYRRIEAWFQLSNRTENGNQFTTPGVCD

>CDS16155.1|Echinococcus_granulosus_rabphilin

MEGYVCPSDRQLALRARLGTGWSTVAQTNQRRTNTRLSDEELDHIYRVLQRNESINERESRRVESMLKRLENMQVATTPQNKTACSICGHEFGLLLKSSTHCADCGRLVCNRCSMEFTEELPEKPRLASDVSNVSPSKYLPPGSLGVSGGQRMGSRVLRRLWRSRQLSERRDDPKQGLGPQDRSAYNGAVGAHSKYKSFSLERRQSMSRLSNFFSIPLSLTPSASLTPTSNRFHLCKLCCEAREVWKKSGAWFHRGLPKCGLHSSCPASPLISKPMTTVDGWTSLLARGPAELIPSTGGAQASDDDWTRIEKRPPEPVPTSEGVGSHSSFTIPSYSLEDVKSSLEEPSNSSSAVITNSGNNGGIGAVTHPGSPHNLFESPTWKPSMEQQQRQHQTLLLSPTSKTDQKTPSQIHESLSGNSLNSTERSPQRGRESKVQSLFPETRLGLGLSDPKIAPSTFTLSPELKFNKGSPSRRKSAIPFSEAPLGILYFSVTHDVTNCELHVRIHNAKHLIAMDANGLSDPFVVCQLLPTVGKRFRTRTIPQSLNPVWDETFTFVDFDNRKLEQKILRLAVLDADTFGADWLGEYRLGLGELLPDRLSEFAVPLNPRKPLSKEIDDLTNPTRGKIQIALRFIEETKQLCVEILRCADLAPMDRNGLSDPFVKLCLRPDKLRKTKQKTKIKKSTLFPEFYEEFFFAMAALEVNKRTLEITVWDFDRGMRNDFIGGLTLGAKAKAERREVWQAVFRPPYRRFEAWFQLASRSDTEYPGSEPQTNVCAQGRVRACISSSRYAVPESVINFRIILILPSPLLSSLLPSFPILWGDTVPLRITFPPPLFICDSC

>XP_015773295.1|Acropora_digitifera_rabphilin

MSEQIDEEQRWVCPNDRQLCLRAKLNSGWSFHTSGPARKPASKNNISVSEQEIIRDVLERSERLRQIEEERVGRLVDKLENMHNRAIGDGRETCLLCNSKFGTLKVVAKRCDICEKNVCQKCGLDTHDSYGDAIWLCLLCSEHRELWKRTGAWFFKAIPNYRHPRQDQESTGKRKETKSSNSSDYLSLSWTHKYDQTWPNYSNAGDSVDGVRVESESEEEESSSSSDEMIFDPRQKSALYLDMDQSAADDERPLYSSRLSDLSMPGSIILENSDTPPPFRKRTLPLTEQDKHRRPGGVSPRLALKEDVNGQTEPPEKSYTKQKTSPFLGRASEEEVKVDVHRTLTIKQETEEEDIDEVVRTYKESEAVGQSDTGNAGLGTIEFSVHYDKQYSQLQITIECAKGLKAMDHRGTSDPYVKLHLLPGASKSNKLRTHTKYKTLNPRYGETLVYHGITDDDLSKKSLRLQVLDEDKLGRNNFIGETFVPLKVFHSKPSQSLKRSLVSRSSVEDGGEPLVSNLGRIQISLRYKSQKNQLVVGIIRCAGLAAMDSNGYSDPFVKV

>XP_020895735.1|Exaiptasia_pallida_rabphilin

MSGIANDVGNRWTCPNDRQLGLRAKLNSGWSFHSANPAKKATKANISEAEQEIIKDVIQRSDRIRQQEENRVGRLVEKLDNLRKNAMGNGEDLCLLCASKFGILKTVARECDICEKNVCEKCGVDTHNSIGQPIWLCMLCSEQREVWKKTGAWFFKSIPKYTTVAEAERNRANSLSRSDTAVSGHYSDYYSDGRSGWTPVMPRRSRDPVVNGDALSSIANHSDDDGSSSEDEMILDRQKRPRASTGSASTASDDFSLYRESGNLLDAPSDGFRGRTGRVSPADRMILKKRASRGGSIAMDTSPLDKSDGAKLGGLGDEEDIDELVRNYKENAESQKKPDNASGLGSIEFTLRYQKLDSRLDVVLQSAEDLKPMDVSGTSDPYVKLHLLPGASKSNKLRSQTKYKTLNPEFDETLTYHGITEEEISNKTLRLQVYDEDTLGRNDFIGETSVNLKVLNSNPTQTFKRNLLAKAFIDTGKDSPSPNSSSNLGRIQLSLKYNSARSMLFVGIVRCSXLAAMDANGYSDPYVKCYLKPDPNKKTKRRTTIKKKTLNPEFNEEFVYEIPHNELAKKSLEVTVWDYDVGKSNDFIGGVVLHIKAEGAALKHWYETLKNPNQLHIQWHDLETMTVQDD

>XP_012561655.1|Hydra_vulgaris_rabphilin

MRMVELSIDGDDRWTIPDDRQLALRGRLDTGWSSYKGYSRKGEVLTEEEFLILSDVMSRRDAKQKQEELRIGTLVKKLENMRTNVLGDGIQTCLICGTKAGLLKDSLNYCLHCGHLVCQKCGVDSKITTETSKKLFYCVICNEERQLWKSSGAWFYNSIPKYVISDVGSELNSNMRVLGRNNSFNTPRPVWIENRSMSHESDATDDDLETTSFNAFEISDERSQSWHGQIEVNSPQLKFSNVKRSCSDLPNIAPKSPTLSYSSNQVDGVFSDRMQTNGLTSRGHLNNNSNSDLAYERQRSYGSTLSLEMKPDKKNLLGIKRLVRNQSRDSDLDKVSKGSKQNDFGSSDNMSNCSEQDIDSLFKEEAQNQAVKEKKEYYGKIGMIEFTLKYDASTEELHVIIQNCKGLCGDVEKGKLPFPYVKTYLLPEANKATKKRTSTLKKTADPVFNETLVYHGISETDIRSKGLKLSVMDEKSRHSTVIGETSIPLKNLSMQPVQQFRRILDVRNNSNPLYLESHISSNNPGRIELALHYLSKQEKLVVGIIRCAGLKAHDSNGYSDPFVKCYLLPDPQKHTKQKTAIKKKTLNPEFNEEFTYKIAHHELAKRTLHITVWDHDVGRTNDFIGGITLGIQSTPEALKHWFETLKTADKKVIKWHTLSEDCPAID

>evg19024|Mnemiopsis_leidyii_rabphilin

MAKAGLHKWVCPTDRELCLRSKLNNGMGWSYKSVQSGLKPTCTRETFAKHEIDKIEEVIKKHNELVQLEEERIGKLMERFENMKCSQGDGYKTCLFCNASFGFLKAKPFRCHMCQRNVCSKCKVETEKRQIVLPKILCRVCSEYREIWKKSNAWFFHAIPKYVIPPKLPKQQPKIKNSPVQKKPIERPNKVNRERFWDPDSDYLSTTDDSDESDDSDSDDSDSDDAVDKSNKDDELDGDDSTSGGISCDEDQVLDAIASDEINSHEKKDDLETGELGAIRFSLQYDPIEQMLNIQVFSGHKLKPMDVGGTSDPYVKCTLIPGHPKATKLKTSRKECDLNPVFNEKLTYHGIFDSDIKTKMVRLTVLDYDRMTKNDMIGITTVPLDGLIPQKQHLYGEILKEPEDETHEASAIVDRGRIQISLHYQANKELKVTIVRCSMLMVPGSDTLPNPMVKVNIRPSPKRNTFKRKTEILKQTSNPEFGHKAKFTYDITSFDIESNDRCLEVTVWDCTGLKRHSYIGGVVLGKESGGSTRSHWEDALSSCGHRVTRWHMLEGESTSTIGECSSIELTDQSQADVANPAAAP

>TR42475|c0_g1_i1|Beroe_ovata_rabphilin

MAQEGLHKWVCPTDRELCLRAKLNNGMGWSYKSVQSGVVPSTTRETFAKHEMEKIEEVIKKHNQLVKVEEERIGKLMERFDNMKNSQGDGYKTCLFCNTSFGFLKAKPFRCHLCQRNVCSKCKVETEKRQIVLPKILCRVCSEYREIWKKSNAWFFHAIPKYVIPPKLPKQPMKIHNSPIQKKPQERPVKGHSDSYWDNDTDYASTTDDDTTSDETDSDDTDSESGENDSNPDNSKDDGGESTSVEIQCEEDQVLDIIGSDDTNFIENDAEMETGERGAIKFSLQYDSIEQMLNVKIFSGHRLKSMDVSGTSDPYVKCILIPGHQKATKLKTSRKECDLNPVFNEKLTYHGIFESDINSKMARLTVLDYDRMTTNDIIGYTTVPLAGLVPQQQHYYCEILKEPEDETHEASALGDRGRIQISLHYQPNKELKVSIVRCSMLMVPGSDTLPNPIVKVNIKPSPKKNFFKRKTEVLKKTANPEFGPKARFTYDISNIDIESSDRCLQVSVWDCTGLKRLSHIGGITLGKESGGAAKSHWLDALSSCGHRVTRWHMLDGDSFGMIGETSSIDMTDQLSVQPSTP

>DN21272_c0_g2_i1|Hormiphora_californiensis_rabphilin(partial)

QAFLSSTGMTKDGLHKWVCPTDRELCLRSKLNNGMGWSYKSVQAGFKQPSSKDTFAKHELEKIQEVIERHNGLVKLEEERIGRLMERFENIKCSQGDGTKTCLFCATSFGFLKAKPFRCHLCQRNVCGKCKVETEKRQAVVPQILCRVCSEYREIWKKSNAWFFHAIPKYVVPPKQPKQPLTKLKNSPVPMRSLERRKEGAGANSHGYHDHSEPGYDSSSTSEEETEDDSESEDSDADDSESEGEEQEEGDGTSLSLPCCEEDQVLDILAEEVKVDYSEAELETGELGAINFSLQYDPIEQMLNVKIFSGHKLKPMDVGGTSDPYVKCTLIPGNPKATKLKTSRKECDLNPVFNEKLTYHGIFEFDINSKLLRLTVLDYDRMTKNDMIGVTTVPLAGLVPQQQHMYREVLREPEEEREVEGSAVAAGLGAASGDRGRIKISLHHLGGKELKVSIIRCSMLMV

>evg140073|Hormiphora_californiensis_rabphilin(complete)

MPPLSDSWGSDRRRLLERPCDESIGSPIPGIIPQAFLSSTGMTKDGLHKWVCPTDRELCLRSKLNNGMGWSYKSVQAGFKQPSSKDTFAKHELEKIQEVIERHNGLVKLEEERIGRLMERFENIKCSQGDGTKTCLFCATSFGFLKAKPFRCHLCQRNVCGKCKVETEKRQAVVPQILCRVCSEYREIWKKSNAWFFHAIPKYVVPPKQPKQPLTKLKNSPVPMRSLERRKEGTGANSHGYHDHSEPGYDSSSTSEEETEDDSESEDSDADDSESEGEEQEEGDGTSLSLPCCEEDQVLDILAEEVKVDYSEAELETGELGAINFSLQYDPIEQMLNVKIFSGHKLKPMDVGGTSDPYVKCTLIPGNPKATKLKTSRKECDLNPVFNEKLTYHGIFEFDINSKLLRLTVLDYDRMTKNDMIGVTTVPLAGLVPQQQHMYREVLREPEEEREVEGSAVAAGLGAASGDRGRIKISLHHLGGKELKVSIIRCSMLMV

>evg1107189.2|Trichoplax_adhaerens_rabphilin

MTDRNRKASTNAWICPNDRQLCLRAKLNIGWSYHSDSPRGSGNELAQNEKKQIMQVLQKAENLEKMEQERIGRLVEKLDNMKNHAVGDGERSCILCGVTFSTFGTTPTPCNDCKKGVCSKCGVDTYNSRKQPLWLCKLCSEQRELLKRSGAWFYKSMPSYIMPVKNADNLGSKNIRSTASATSSTSKDNYGTSLTKRPTMQADGSSEDETDTSLSTDSDTDDGSSNDDSIGINDSVKPRPAWASLAHKDHSSNSEISLKKSMDDERNEESSDYSSEVSFSASKVNPKEKAENENEINLAFQEFKPSQADFRSPITENESIDSTGNEGLGSLDFSVLFEPIDFKLHVTIIKAKNLKGMGKHGVDPYVKLQLLPGSSKSNKMRTKQSKTCNPEFNETLTYLGVTKDDIARKTLRVYVMNHDRFSSDTVIGIYNLKLKDIGDVMKRFNDVPLHKISQEDLEEDRVNSIERGRIEISLKYVTATRTLEVGIIQCIGLKAMDASGYSDPYVKCYLRPDKYKITKQKTSVKRRTLNPKFNAVFKYKYAHSELAGKTLDIQVWDRDIGRKNDFIGGVYLGKDSSGDQLRHWFQTLKTPNAKFTHFHTLTDELRHPAE

>m.15463|Oscarella_carmela_rabphilin

GHHFRSLPPMDFDDLPLKWLCPNDRQLEVRSRLGSGWSTRTIDLDRRTEEGMTEEEKKTVAAVLLRANELERSEQERIGRLVSRLENMRKASFNDASKCVACGENIKSFGKSSLMCASCRKGMCVDCGSSVQDEMTMVEAIVCKLCVEQRELWKKTGAWFYGQLPKYVKTQPASRSKFRSSPSPSYDASREPSDNAPSSSESSSDNDLDYHLLQVPSKNRKQSSDTKKESLFCELQKSNQLSVSSTNVSQLSATSGNSATSVNSAQSFHSLNFDSKPFKDGSEMKRPAAPPKRLSAPLPTANGSSNGHGHSLAIPEGRPRRLSGWKEQLENEKLLIFGSKGTRIEKTADKEEEEENENEDDEDDIDALFEQAGKSVKLDDDESVDIGAGSRFGAVHFAVLYSADESNLYVRINKCTELNMPGHSGPVNSFIKLHLLPGNLMSREIKTGIRTKQNNPVYSDRLVYHGITEEDWKEKTLRLSAYHRGGISHHNQYIGETRIPLRKLQKNKEDQFVRHLEKQPLQPTDETPLANDRGVIQISLQYDTAKKRLLIGILQAAGLPPMDTNGFSDPYVKCYLLPDKSKKTKQKTYVAKKTLNPEFNKEFVYDIELNDLATRVLEVTVWDHDIARHNDYIGGIQLQSKASGLLLTHWYSTLKFPGKRFVYWHRLSPIQPTKSTNDKQVIPMNV
