## Supplementary material for "Early metazoan origin and multiple losses of a novel clade of RIM pre-synaptic calcium channel scaffolding protein homologues": File S2

>XP_019401893.1|Crocodylus_porosus_Cav1.1

MELASAQEDQKKKQQKEKSKKPVPPAAPRPARALFCLTLQNPLRKACISIVEWKPFEIIILLTIFANCVALAVYLPMPEDDTNATNSRLEKIEYVFLIIFTIEAMLKIIAYGFLFHTDAYLRSGWNVLDFAIVSLGLFTVTLEQISVMQGAPPSGKGGFDVKALRAFRVLRPLRLVSGVPSLQVVLNSIIKAMVPLLHIALLVLFMIIIYAIVGQELFKGKMHKTCYYTGTDIIATVGSEKPAPCTSTGHGRHCTLNGTDCRGGWPGPNNGITHFDNFGFAMLTVYQCITMEGWTEVLYWVNDAIGNEWPWIYFVSLILLGSFFILNLVLGVLSGEFTKEREKAKSRGTFQKLREKQQLEEDLKGYMDWITHAEVMDSVRTRGEGMLPLDEGSSETESLYEIEGMNKWILFFRHWRRWNRLFRRKCREVVKSKFFYWLVILLVALNTLSIASEHHMQPDWLTHVQDNANRVLLSLFAAEMLLKMYALGLRQYFMSLFNRFDCFVVCAGILETILVEIRLMSPLGISVLRCIRLLRIFKITKYWTSLSNLVASLLNSIRSIASLLLLLFLFIVIFSLLGMQLFGGKYDFEDMEVRRSTFDNFPQALISVFQVLTGEDWNSIMYNGIMAYGGPSFPGMLVCIYFIILFVCGNYILLNVFLAIAVDNLAEAESLTSAQKAKAEERKRRKMSRGYPGKSEEEKQLLAKKLEQKAKGEGIPTTAKLKVDEFESNVNEIKDPYPSADFPGDDEEDEPEIPLSPRPRPLAELQLKEKAVPMPEASAFFIFSPTNKFRILCHRIVNATWFTNFILLFILLSSISLAAEDPIRAESFRNQILGYFDIGFTSVFTVEIVLKMTAYGAFLHKGSFCRNSFNILDLLVVAVSLISMGIQSSTISVVKILRVLRVLRPLRAINRAKGLKHVVQCVFVAIKTIGNIVIVTTLLQFMFACIGVQLFKGKFNSCTDPSKITERECRGYFITYVDGDPTQIQLQERVWQHNDFHFDNVLSAMMSLFTVSTFEGWPQLLYKAIDTHTEDMGPIYNYRVEMAIFFIIYIILIAFFMMNIFVGFVIVTFQEQGESEYKNCELDKNQRQCVQYALKARPLRRYIPKNPYQYQIWYVVTSSYFEYLMFFLILLNTICLGMQHYNQSDEMNHASDILNVTFTILFTVEMFVKLMAFKAKGYFGDPWNVFDFLIVIGSIIDVILSEIDDSEDNSRVSITFFRLFRVLRLVKLLSRGEGVRTLLWTFIKSFQALPYVALLIVMLFFIYAVIGMQMFGKVAMVDGTQINRNNNFQTFPQAVLLLFRCATGEAWQEILLAASYGKLCDPESDYAPGEEYTCGTGFAYFYFISFYMLCAFLIINLFVAVIMDNFDYLTRDWSILGPHHLDEFKRIWAEYDPEAKGRIKHLDVVTLLRRIQPPLGFGKFCPHRVACKRLVCMNMPLNSDGTVTFNATLFALVRTALKIKTEGNFEQANEELRAIIKKIWKRTSMKLLDQVIPPIGDDEVTVGKFYATFLIQEHFRKFMKHQEEYYGYRPKKNPIEIQAGLRTIEEEAAPEIHRAISGDLTAEEELERAMVEAAMEEGIYRRQGGLFGQVDNFIEHHSPLQPHVASQRPLQFTEQGSEDIDSPVFLDDFTPERNTNASNANNNNAASRVEYEDELQGKKISCQTRQLSALAPKCASHHEKLQQELSRRRAASVGVCPVSQL

>NP_001292076.1|Gallus_gallus_Cav1.1

MEPASHQDDVKKRQQKEKSKKPVPPVAPRPPRALFCLTLQNPLRKACISIVEWKPFEIIILLTIFANCVALAIYQPMPEDDTNVANSSLEKLEYVFLIFFAIEAMLKIIAYGFLFHTDAYLRNGWNVLDFSIVSLGLVTMTLEQINAKEGGSLGGKGGFDVKALRAFRVLRPLRLVSGVPSLQVVLNSIIKAMVPLLHIALLVLFMIIIYAIVGQELFKGKMHKTCYYLGTDVIATVGSEKPAPCTTSGHGRHCSINGTECRGGWPGPNNGITHFDNFGFAMLTVYQCITMEGWTEVLYWVNDAIGNEWPWIYFVSLILLGSFFVLNLVLGVLSGEFTKEREKAKSRGTFQKLREKQQLEEDMKGYMDWITHAEVMDSDRARGEGMMPLDEGGSETESLYEIEGMNKWILYFRQWRRWNRMFRRKCRDVVKSKFFYWLVILLVALNTLSIASEHHFQPEWLTIVQDNANRVLLALFVAEMLLKMYALGLRQYFMSLFNRFDCFVVCAGVLEIILVELSTLSPLGISVLRCIRLLRIFKITRYWTSLSNLVASLLNSVRSIASLLLLLFLFIIVFALLGMQLFGGMYDFEDMEVRRSTFDNFPQALISVFQILTGEDWNSIMYNGIMAYGGPSFPGMLVCIYFIILFVCGNYILLNVFLAIAVDNLAEAESLTSAQKAKAEERKRRKMSRGYPEKSEDEKQMLAKKLEQKAKGEGIPTTAKLKVDEFESNVNEIKDPYPSADFPGDDEEDEPEIPLSPRPRPLAELQLKEKAVPMPEASSFFIFSPTNKFRMLCHRIVNATWFTNFILLFILLSSISLAAEDPIRAESFRNQILGYFDIGFTSVFTVEIVLKMTAYGAFLHKGSFCRNSFNILDLLVVAVSLISMGFESSTISVVKILRVLRVLRPLRAINRAKGLKHVVQCVFVAIKTIGNIVVVTTLLQFMFACIGVQLFKGKFYSCTDPSKLTEKECRGHFINYVDGDPTQIELKERVWFHNAFHFNNVLSAMMSLFTVSTFEGWPELLYRAIDTNDENKGPIYNYRVEIAMFFIIYIILIAFFMMNIFVGFVIVTFQEQGESEYKNCELDKNQRQCVQYALKARPLRRYIPKNPYQYQIWYVVTSSYFEYLMFFLIMLNTICLGMQHYNQSAEMNHVSDILNVAFTVLFTLEMILKLMAFKAKGYFGDPWNVFDFLIVIGSIIDVILSEIDTVLASSGGLYCLGGGCDSIDPDDNSRVSITFFRLFRVMRLVKLLSRGEGVRTLLWTFIKSFQALPYVALLIVMLFFIYAVIGMQMFGKIAMVDGTQINRNNNFQTFPQAVLLLFRCATGEAWQEILLDCSYGKRCDPESDYAEGEEYTCGTGFAYFYFISFYMLCAFLIINLFVAVIMDNFDYLTRDWSILGPHHLDEFKRIWAEYDPEAKGRIKHLDVVTLLRRIQPPLGFGKFCPHRVACKRLVCMNMPLNSDGTVTFNATLFALVRTALKIKTEGNFEQANEELRTIIKKIWKRTSMKLLDQVIPPIGDDEVTVGKFYATFLIQEHFRKFMKRQEEYYGYRPKKNPVEIQAGLRSIEEEAAPEIHRAISGDLTAEEELERAMVEAAMEEGIYRRTGGLFGQVDSFLEPSSPLQPHVASQRPLQFTEVGSEDLDSPVFLDDFPQEGNTNVNNANWQQGLEYEDEVLRRAVPRPPVREPDLPIPLERLRRRRGAASEHHPASGPSLHPQQGSTVTAAASSSEHRWHESIDTNTSSTPAITFLIQEALISGGLAALARDPSFVAVTRDEMAASSQMEMDEVEKAAVELLQGREALQDTEISSAPRDTLAASAVSTPGLSVAASHQTSSAARL

>XP_020660962.1|Pogona_vitticeps_Cav1.1

MEPASALDDSLKKKQQKEKAKKAALELPPRPPRSLLCLTLQNPVRKACIAIVEWKPFETIVLLTIFANCVALALYLPMPEDDTNKMNSRLEKLEYFFLIVFAIEATLKIIAYGFLFHADAYLRNGWNVLDFTIVFLGVFTVILEKISVIEGALLSGQGGFDVKALRAFRVLRPLRLVSGIPSLQVVLNSIVKAMLPLFHIAVLVVFMLTIYAIMGQELFKGKMHKTCYYNGTDIIATVENEKPSPCTSAGHGHQCTIPGSECRGKWPGPNNGITHFDNFGFAMLTVYQCISMEGWTQVLYWVNDAIGNEWPWIYFVSLILLGSFFILNLILGVLSGEFTKEREKAKSRGTFQKLREKQQLDEDMKGYMDWIVHAEVMESERKRGEGMMSEDEGGSETESLYELEGMNKFILFFRHWRRWNRLFRRKCREVVKSRFFYWLVILIIALNTFSIASEHHNQPDWLTQAQDVANRVLLALFTVEMILKMYALGLRQYFMSLFNRFDCLVVCTGILEIILVESASMSPLGISVLRCIRLLRLFKITKYWRSLNNLVASLLNSVRSIASLLTLLFLFMVIFALLGMQLFGGKFDFDDVEIRRSTFDNFPQALITVFQVLTGEDWTSVMYNGIMSYGGPSYPGMLVCIYFIVLFVCGNYILLNVFLAIAVDNLAEAETLTSAQKAKAEEKKRKKLARGYPEKSEEEKQLLAKKLEQKAKGEGMPTTAKLKVDEFESNVNEIKDPYPSADFPGDDEEDEPEIPLSPRPRPLAELQLKEKAVPMPESSAFFIFSPTNKIRVLCHRIVNATWFTNFILLFILLSSISLAAEDPIRAESFRNKILGHFDTGFTTVFTVEIVLKMTAYGAFLHKGSFCRNYFNILDLLVVAVSLISMGLESSAISVVKILRVLRVLRPLRAINRAKGLKHVVQCVFVAIKTIGNIVLVTFLLQFMFACIGVQLFKGKFFYCTDTTKITESECWGYFITYVDADPTQIVLNQRMWLQNEFHFDNVFSAMMSLFTVSTFEGWPKLLYRAIDTHTENMGPIYNYRMGIAIFFIIYLILIAFFMMNIFVGFVIVTFQEQGETEYKDCELDKNQRQCVQYALKARPLRCYIPKNPYQYQIWYLVTSSYFEYLMFFLIMLNTVCLGMQHYNQSETMNQVSDVLNVVFTILFTVEMIVKLIAFKAKGYFGDPWNVFDFLIVIGSIIDVILSQIDTSPPALQGYSSAGEATPQVPAIDPDETGRISITFFRLFRVLRLVKLLSRGEGIRNLLWTFIKSFQALPHVALLIVMLFFVYAVIGMQMFGKIALVDGTQINRNNNFQTFPQAVLLLFRCATGEAWQEIMLAASYGKKCDSESDFAPGEEYSCGSGFAYFYFISFYMICAFLIINLFVAVIMDNFDYLTRDWSILGPHHLDEFKKIWAEYDPEATGRIKHLDVVTLLRRIQPPLGFGKFCPHRVACKRLVGMNMPLNSDGTVTFNATLFALVRTALKIKTEGNFEKANEELRMIIKKIWKRTSMKLLDQVIPPIGDDEVTVGKFYATFLIQEHFRKFIKRQEEYYGYRPKKNAVEIQAGLRTIEEEAAPEIKRAISGDLTAEEELERAMVEAAMEEGIYRRTGGLFGQVDTFMEPSSPLPPHVASQRPLQFTEMGSEDLDSPVFLDEIPQRGNTNTNNSNSNANNSVTARMTYEDELRGEMRLPSSHQLSHRANKCSIQLEKEQRKRSVLPASGVKTNPILQFSSQQTPFALHEAVDDEAPQLTSSSEGNGLRDMEASTPVTIQFIQEALTSGGLGSLAKDYSFLKLARDEMAAAFQVEVPEMEEVAATLLKRGKISTTSL

>XP_015267320.1|Gekko_japonicus_Cav1.1

MEPTSVQDDIQKKKQQKEKAKKAELEAPPRPLRSLFCLTLQNPVRKACIAIVEWKPFETIILLTIFANCVALAIYLPMPEDDSNKANSRLEKLEYFFLMVFAIEAVLKIIAYGFLFHADAYLRNGWNVLDFTIVFLGVFTVILERISVMESALLSGQGGFDVKALRAFRVLRPLRLVSGIPSLQVVLNSIGKAILPLFHIAVLVVFMLIIYAIVGEELFKGKMHKTCYYIGTDIIATVENEEPSPCTNSGHGRHCTINGSECRGWWPGPNNGITHFDNFGFAMLTVYQCISMEGWTKVLYWVNDAIGNEWPWIYFVSLILLGSFFILNLILGVLSGEFTKEREKAKSRGTFQKLREKQQLEEDMKGYMDWIVHAEVIDSDRTRGEGMMPSDEGGSETESLYEIEGMNKWILFFRHWRRWNRLFRRKCREVVKSRFFYWFVILIVSLNTLSIASEHHRQPGWLTHVQDIANRVLLALFTVEMILKMYALGLHQYFMSIFNRFDCLVVCTGILEIILVEIVSMSPLGISVLRCIRLLRIFKITKYWTSLNNLVASLLNSVRSIISLLTLLFLYIVIFALLGMQLFGGRFDFDDAEIRRSTFDNFPQALISVFQVLTGEDWTSIMYDGIMAYGGPSYPGILVCIYFIILFVCGNYILLNVFLAIAVDNLAEAETLTSAQKAKEEEKKRKKMARGYPEKSDEEKLLLAKKLEQKAKGEGIPTTAKLKVDEFESNVNEIKDPYPSADFPGDDEEDEPEIPLSPRPRPLAELQLKEKAVPMPEASAFFIFSPTNKIRVLCHRIVNATWFTNFILLFILLSSISLAAEDPIRAESFRNQILEYFDYVFTSVFTVEIVLKMTAYGAFLHKGSFCRNSFNILDLLVVAVSLISMGIQSSAISVVKILRVLRVLRPLRAINRAKGLKHVVQCVFVAIKTIGNIVLVSALLQFMFACIGVQLFKGKFYSCTDPLKITEEECRGQFIDYVDADPTQIILRERIWFHNEFHFDNVLSAMMSLFTVSTFEGWPKLLYQAIDTHTEDMGPIYNYRMGIAIYFIVYLILIAFFMMNIFVGFVIVTFQEQGETEYKSCELDKNQRQCVQYALKARPLKCYIPKNPYQYQIWYVVTSSYFEYLMFFLITLNTICLGMQHYNQSETMNQVSDILNVVFTLLFTVEMILKLIAFKAKGYFGDPWNVFDFLIVIGSIIDVILSQIDTVLASRGGLYCLSGGCDGIPPSTGDDDNGRISITFFRLFRVLRLVKLLSRAEGIRNLLWTFIKSFQALPHVALLIVMLFFIYAVIGMQMFGKVGLVDGTQINRNNNFQTFPQAVLLLFRCATGEAWQEVLLAASYGKLCDPESDYTPGEEYSCGTGFAYFYFISFYMICAFLIINLFVAVIMDNFDYLTRDWSILGPHHLDEFKRIWAEYDPEAKGRIKHLDVVTLLRRIQPPLGFGKFCPHRVACKRLVGMNMPLNSDGTVTFNATLFALVRTALKIKTEGNFEQANEELRMIIKKIWKRTSMKLLDQVIPPIGDDEVTVGKFYATFLIQEHFRKFMKRQEEYYGYRPKKNAVEIQAGLRTIEEEAAPEIHRAISGDLTAEEELERAMVEAAIEEGIYRRTGGLFGQVDTFIEPSSPLPPHMASQRPLQFTEVGSEDLDSPVFLDDFPPEQHTNVNNSNSNANNNKTTWMAYENELRGETVLPPSHRPLVLANKCFLQPEKQPRKISQVQRHGVTRCSELPLSTCQLPQKMPIDSQQEAENGKVLQSRNSTEANTSRDKETSTPAILQLIQEALISGGLHSLVKDSSFLKLVRDEMAAAFQTEVTEMEEIAATLLNGKESLARSLCRSRSFTSSLEATPDTRL

>NP_000060.2|Homo_sapiens_Cav1.1

MEPSSPQDEGLRKKQPKKPVPEILPRPPRALFCLTLENPLRKACISIVEWKPFETIILLTIFANCVALAVYLPMPEDDNNSLNLGLEKLEYFFLIVFSIEAAMKIIAYGFLFHQDAYLRSGWNVLDFTIVFLGVFTVILEQVNVIQSHTAPMSSKGAGLDVKALRAFRVLRPLRLVSGVPSLQVVLNSIFKAMLPLFHIALLVLFMVIIYAIIGLELFKGKMHKTCYFIGTDIVATVENEEPSPCARTGSGRRCTINGSECRGGWPGPNHGITHFDNFGFSMLTVYQCITMEGWTDVLYWVNDAIGNEWPWIYFVTLILLGSFFILNLVLGVLSGEFTKEREKAKSRGTFQKLREKQQLDEDLRGYMSWITQGEVMDVEDFREGKLSLDEGGSDTESLYEIAGLNKIIQFIRHWRQWNRIFRWKCHDIVKSKVFYWLVILIVALNTLSIASEHHNQPLWLTRLQDIANRVLLSLFTTEMLMKMYGLGLRQYFMSIFNRFDCFVVCSGILEILLVESGAMTPLGISVLRCIRLLRIFKITKYWTSLSNLVASLLNSIRSIASLLLLLFLFIVIFALLGMQLFGGRYDFEDTEVRRSNFDNFPQALISVFQVLTGEDWTSMMYNGIMAYGGPSYPGMLVCIYFIILFVCGNYILLNVFLAIAVDNLAEAESLTSAQKAKAEEKKRRKMSKGLPDKSEEEKSTMAKKLEQKPKGEGIPTTAKLKIDEFESNVNEVKDPYPSADFPGDDEEDEPEIPLSPRPRPLAELQLKEKAVPIPEASSFFIFSPTNKIRVLCHRIVNATWFTNFILLFILLSSAALAAEDPIRADSMRNQILKHFDIGFTSVFTVEIVLKMTTYGAFLHKGSFCRNYFNMLDLLVVAVSLISMGLESSAISVVKILRVLRVLRPLRAINRAKGLKHVVQCMFVAISTIGNIVLVTTLLQFMFACIGVQLFKGKFFRCTDLSKMTEEECRGYYYVYKDGDPMQIELRHREWVHSDFHFDNVLSAMMSLFTVSTFEGWPQLLYKAIDSNAEDVGPIYNNRVEMAIFFIIYIILIAFFMMNIFVGFVIVTFQEQGETEYKNCELDKNQRQCVQYALKARPLRCYIPKNPYQYQVWYIVTSSYFEYLMFALIMLNTICLGMQHYNQSEQMNHISDILNVAFTIIFTLEMILKLMAFKARGYFGDPWNVFDFLIVIGSIIDVILSEIDTFLASSGGLYCLGGGCGNVDPDESARISSAFFRLFRVMRLIKLLSRAEGVRTLLWTFIKSFQALPYVALLIVMLFFIYAVIGMQMFGKIALVDGTQINRNNNFQTFPQAVLLLFRCATGEAWQEILLACSYGKLCDPESDYAPGEEYTCGTNFAYYYFISFYMLCAFLVINLFVAVIMDNFDYLTRDWSILGPHHLDEFKAIWAEYDPEAKGRIKHLDVVTLLRRIQPPLGFGKFCPHRVACKRLVGMNMPLNSDGTVTFNATLFALVRTALKIKTEGNFEQANEELRAIIKKIWKRTSMKLLDQVIPPIGDDEVTVGKFYATFLIQEHFRKFMKRQEEYYGYRPKKDIVQIQAGLRTIEEEAAPEICRTVSGDLAAEEELERAMVEAAMEEGIFRRTGGLFGQVDNFLERTNSLPPVMANQRPLQFAEIEMEEMESPVFLEDFPQDPRTNPLARANTNNANANVAYGNSNHSNSHVFSSVHYEREFPEETETPATRGRALGQPCRVLGPHSKPCVEMLKGLLTQRAMPRGQAPPAPCQCPRVESSMPEDRKSSTPGSLHEETPHSRSTRENTSRCSAPATALLIQKALVRGGLGTLAADANFIMATGQALADACQMEPEEVEIMATELLKGREAPEGMASSLGCLNLGSSLGSLDQHQGSQETLIPPRL

>NP_001139622.1|Danio_rerio_Cav1.1

MEGNGEAPMSSFIMDEETLKRKQREKLKKLQATGGNPRPARSLLFLTLKNPFRKACINIVEWKPFEIIILLTIFANCVALAVFMPMPEEDTNNTNSNLESLEYIFLIIFTMECFLKIVAYGFLFHADAYLRNCWNILDFVIVTMGLFTVVVDFINSISGVEAPVEQKGGFDMKALRAFRVLRPLRLVSGVPSLQVVMSSILKSMLPLFHISLLVFFMVTIYAIIGLELFKCKMHKTCYHTGTDIIATGDDAQAAPCAQAGNGRRCTLNGTECRGDWPGPNNGITHFDNLGFSMLTVYQCITTQGWTDVLYWVNDAIGMEWPWLYFVTLILLGSFFILNLVLGVLCGEFTKEREKSSRSGEYQILRERQQFDEDLKGYMEWITQAEVMDNDQEGQGLLPLQDGSETETLYELDILNKLMFYVRHARRWNRFFRRKCRVWVKSKLFYWLVILLVFFNTLAIATEHHQQPDSLTNFQDNTNKALLSLFAVEMFLKMYAMGLPSYFMSLFNRFDCFVVSVGILELILVRMDVMSVMGISVLRCIRLLRIIKITRHWTTLSNLVASLLNSVRSIASLLLLLFLFIVIFALLGMQVFGGKFNFPDDRVRRSNFDNFPQALITVFQILTGEGWNYVMYDGIMAHGGPAIPGILVSIYFIILFICGNYILLNVFLAIAVDNLAEAESLTSAQKEKAEEKKRKRLLRENLPDKGEEEKALLAKKLAEQRAKIDGIPTTAKLKVDEFESNVNEIKDPFPPADFPGDDEEEEPEIPLSPRPRPMADLQLKETEVPMPEASAFFLFGPQNKFRKLCHRIINATTFTNIILLFILLSSISLAAEDPIDPMSFRNQVLAYADYVFTSVFTAEIVLKMTTYGAFLHKGSFCRNSFNILDLIVVGVSLLSMGMESSAISVVKILRVLRVLRPLRAINRAKGLKHVVQCVFVAIKTIGNIVLVTMLLDFMFACIGVQLFKGKFLYCTDPLKMTAEECQGTFIQHQENALHDMVVSQRLWMNSDLNFDNVLNGMLALFTVSTFEGWPDLLYKAIDSNLENMGPVYNNHIEISIFFIVYLILIAFFMMNIFVGFVIVTFQEQGEQEYKNCELDKNQRQCVQYALKARPLRCYIPKNPYQYQVWYIVTSCYFEYLMFLLIMLNTMCLGMQHCKQSDHITDLADTLNVIFTVLFTVEMILKLGAFKAKGYFGDPWNVFDFVIVVGSIVDVILSEIDAALAAQGGLYCLTGCSEVNPMQAIADSENMSVSITLFRLFRVMRLVKLLNRFEGIRNLLWTFIKSFQALPYVALLIVMLFFIYAVIGMQVFGKIALLDGTIINRNNNFQTFPQAVLLLFRCATGEGWHEIMLGCLYGQRCDPKSEYLPGEEYTCGSGFAILYFMSFYMLCAFLIINLFVAVIMDNFDYLTRDWSILGPQHLDEFKKIWAEYDPEATGRIKHLDVVTLLRRIQPPLGFGKFCPHRVACKRLISMNMPLNSDGTVTFNATLFALVRTGLKIKTEGNFEQANEELRAIIKKIWKRTSMKLLDQVIPALGDDEVTVGKFYATFLIQDHFRKFMKRQEEYYGYRPSKKNKNTTEIQAGLRSIEEEAAPELQRAISGDLLNDEEMDRAMEESGEESIYRRAGGLFGNHVDPFSMERGTGAAQVTSQRPLQLADSRADRTETTYASYHANNNNTNNNNRRRHRCEPHFPACISASRLHSPASAAEFLIQQVLTSGGLESLASDWRFVSVTKSEMAAALHLDLQELERKSAAILNQHKGQCVLRKKKMETSHSDITHCHV

>XP_019400097.1|Crocodylus_porosus_Cav1.2

MVNENKRMYIPEENHQGSNYGSPRPAHANMNANAAAGLAPEHIPTPGAALSWQAAIDAARQAKLMGSAGNATISTASSTQRKRQQYGKQKKQGTTTATRPPRALLCLTLKNPIRRACISIVEWKPFEIIILLTIFANCVALAIYIPFPEDDSNATNSNLERVEYLFLIIFTVEAFLKVIAYGLLFHPNAYLRNGWNLLDFIIVVVGLFSAILEQATKADGGNSIGGKGAGFDVKALRAFRVLRPLRLVSGVPSLQVVLNSIIKAMVPLLHIALLVLFVIIIYAIIGLELFMGKMHKTCYHAHGALADTPAEEDPSPCAPQSAHGRQCQNGTECKAGWEGPKHGITNFDNFAFAMLTVFQCITMEGWTDVLYWVNDAIGRDWPWIYFVTLIIIGSFFVLNLVLGVLSGEFSKEREKAKARGDFQKLREKQQLEEDLKGYLDWITQAEDIDPENEDEGMDEEKPRNMSMPTSETESVNTDNVAGADIEGENCGARLAHRISKSKFSRYWRRWNRFCRRKCRAAVKSTVFYWLVIFLVFLNTLTIASEHYHQSNWLTEVQDTANKVLLALFTAEMLLKMYSLGLQAYFVSLFNRFDCFIVCGGILETILVETKIMPPLGISVLRCVRLLRIFKITRYWNSLSNLVASLLNSVRSIASLLLLLFLFIIIFSLLGMQLFGGKFNFDEMQTRRSTFDNFPQSLLTVFQILTGEDWNSVMYDGIMAYGGPSFPGMLVCIYFIILFICGNYILLNVFLAIAVDNLADAESLTSAQKEEEEEKERKKLARTASPEKKQEIEKPAVEEETKEEKIELKSITADGESSPATKINVEDYQPNENEEKNPYPTTEAPGEEDEEEPEMPVGPRPRPMSELHLKEKAVPMPDASAFFIFSPSNRFRVHCHRIVNDNIFTNLILFFILLSSISLAAEDPVRHYSFRNQILFYFDIVFTVIFTIEIALKMTAYGAFLHKGSFCRNYFNILDLLVVSVSLISFGIQSSAINVVKILRVLRVLRPLRAINRAKGLKHVVQCVFVAIRTIGNIVIVTTLLQFMFACIGVQLFKGKLKSCSDSSKQTPAECKGYFITYKDGEVTQPMIQPRSWENSKFDFDNVLNAMMALFTVSTFEGWPELLYRSIDSHMEDVGPIYNHRVEISIFFIIYIIIIAFFMMNIFVGFVIVTFQEQGEQEYKNCELDKNQRQCVEYALKARPLRRYIPKNQYQYKVWYVVNSTYFEYLMFILILLNTICLAMQHYGQSCMFKEAMNILNMLFTGLFTVEMVLKLIAFKPKGYFSDPWNVFDFLIVIGSIIDVILSETNPAEHTQCSSSMNAEENSRISITFFRLFRVMRLVKLLSRGEGIRTLLWTFIKSFQALPYVALLIVMLFFIYAVIGMQVFGKIALNDTTGINRNNNFQTFPQAVLLLFRCATGEAWQEIMLACLSNKKCDPESEQANSTEEDQSCGSSFAIFYFISFYMLCAFLIINLFVAVIMDNFDYLTRDWSILGPHHLDEFKRIWAEYDPEAKGRIKHLDVVTLLRRIQPPLGFGKLCPHRVACKRLVSMNMPLNSDGTVMFNATLFALVRTALRIKTEGNLEQANEELRAIIKKIWKRTSMKLLDQVVPPAGDDEVTVGKFYATFLIQEYFRKFKKRKEQGLVGKPSQRNALSLQAGLRTLHDIGPEIRRAISGDLTAEEELDKAMKEAVSAASEDDIFRRAGGLFGNHVSYYQSDGRSTFPQTFTTQRPLHINKSGNNQGDTESPSHEKLVDSTFTPSSYSSSGSNANINNANNTALCRFPSPPNFPSTVSTVEGHGTPLSPTIRVQEAPWKLGSKSSGSRDSQLAIVCQEEVSQDETYDENLNEDVEYCSEPSLISTEMLSYQDDENRQLTPPENNKGEDNRQSPRRGFLCSSSLGRRASFHLECLKRQKNQGVDVSQKTVLPLHLVHHQALAVAGLSPLLQRSHSPTMFSRLCATPPATPCSRGWPEQTIPTLRLDGAESSEKLNSSFPSIHCSSWYSDRASCSSARRARPVSLTVPSQTGGSGRQFHGSAGSLVEAVLISEGLMQFAQDPKFIEVTTQELADACDMTIEEMENAADNILNGNSKQSPNGNLLPFVNCRDPGQDSAGEEEEEVQNPDCRKSQEELKDSRIYISSL

>XP_015142138.1|Gallus_gallus_Cav1.2

MFQAFVRQAYQPLSSHQQGESETKYRGKLVHETQLNCFYIKPGGSNYGSPRPAHANMNANAAAGLAPEHIPTPGAALSWQAAIDAARQAKLMGSAGNATISTASSTQRKRQQYGKQKKQGTTTATRPPRALLCLTLKNPIRRACISIVEWKPFEIIILLTIFANCVALAIYIPFPEDDSNATNSNLERVEYLFLIIFTVEAFLKVIAYGLLFHPNAYLRNGWNLLDFIIVVVGLFSAILEQATKADGGNSIGGKGAGFDVKALRAFRVLRPLRLVSGVPSLQVVLNSIIKAMVPLLHIALLVLFVIIIYAIIGLELFMGKMHKTCYHVQGGLIDTPAEDDPSPCAPQSAHGRQCQNGTECKAGWEGPKHGITNFDNFAFAMLTVFQCITMEGWTDVLYWVNDAIGRDWPWIYFVTLIIIGSFFVLNLVLGVLSGEFSKEREKAKARGDFQKLREKQQLEEDLKGYLDWITQAEDIDPENEDEGMDEEKPRNMSMPTSETESVNTDNVPGTDIEGENCGARLAHRISKSKFSRYWRRWNRFCRRKCRAAVKSNVFYWLVIFLVFLNTLTIASEHYNQPDWLTEVQDTANKVLLALFTAEMLLKMYSLGLQAYFVSLFNRFDCFIVCGGILETILVETKIMSPLGISVLRCVRLLRIFKITRYWNSLSNLVASLLNSVRSIASLLLLLFLFIIIFSLLGMQLFGGKFNFDEMQTRRSTFDNFPQSLLTVFQILTGEDWNSVMYDGIMAYGGPSFPGMLVCIYFIILFICGNYILLNVFLAIAVDNLADAESLTSAQKEEEEEKERKKLARTASPEKKQEIEKTAVEEETKEEKIELKSITADGESPPATKINMDDYQPNENEEKSPYPTTEAPAEEDEEEPEMPVGPRPRPMSELHLKEKAVPMPDASAFFIFSPNNRFRVHCHRIVNDNIFTNLILFFILLSSISLAAEDPVRHLSFRNQVLFYFDIVFTVIFTIEIALKILGNADYVFTSIFTLEIILKMTAYGAFLHKGSFCRNYFNILDLLVVSVSLISFGIQSSAINVVKILRVLRVLRPLRAINRAKGLKHVVQCVFVAIRTIGNIVIVTTLLQFMFACIGVQLFKGKLYSCTDSSKQTEAECRGYYITYKDGEVSQPMIQPRSWENSKFDFDNVLTAMMALFTVSTFEGWPELLYRSIDSHMEDVGPIYNHRVEISIFFIIYIIIIAFFMMNIFVGFVIVTFQEQGEQEYKNCELDKNQRQCVEYALKARPLRRYIPKNQYQYKVWYVVNSTYFEYLMFVLILLNTICLAMQHYGQSCMFKEAMNILNMLFTGLFTVEMVLKLIAFKPKGYFSDPWNVFDFLIVIGSIIDVILSETNHYFCDAWNTFDALIVVGSIVDIAITEVNPAENTQCSSSMNAEENSRISITFFRLFRVMRLVKLLSRGEGIRTLLWTFIKSFQALPYVALLIVMLFFIYAVIGMQVFGKIALNDTTEINRNNNFQTFPQAVLLLFRCATGEAWQEIMLACLPDKKCDPDSEPANSTEADHSCGSSFAVFYFISFYMLCAFLIINLFVAVIMDNFDYLTRDWSILGPHHLDEFKRIWAEYDPEAKGRIKHLDVVTLLRRIQPPLGFGKLCPHRVACKRLVSMNMPLNSDGTVMFNATLFALVRTALRIKTEGNLEQANEELRAIIKKIWKRTSMKLLDQVVPPAGDDEVTVGKFYATFLIQEYFRKFKKRKEQGLVGKPSQRNALSLQAGLRTLHDIGPEIRRAISGDLTAEEELDKAMKEAVSAASEDDIFRRAGGLFGNHVSYYQSDGRSAFPQTFTTQRPLHINKSGNNQGDTESPSHEKLVDSTFTPSSYSSSGSNANINNANNTALCRFPSPPSYPSTVSTVEGHGTPLSPTIRVQEAPWKLPSKRTHYYETLGRSSSRDSQLAIVCQEEVSQDETYDENLNEDIEYCSEPSLLSTEMLAYQDDENRQLTPPENNKGEDTRHSPKKGFLCSSALGRRASFHLECLKRQKNQGVDVSQKTVLPLQMVHHQALAVAGLSPLLQRSHSPTTFSRLCATPPATPCSRGWPQQPIPTLRLEGAESSEKLNSSFPSVHCSSRFPDSSDCGSPRRARPVSLTVPSPTAGSSRQFHGSASSLVEAVLISEGLMQFAQDPKFIEVTTQELADACDMTIEEMENAADNILNGNSKQSPNGNLLPFVNCRDAGQDSAGEEEEEVQNPDCRKSQEELKDSRIYISSL

>Q13936.4|Homo_sapiens_Cav1.2

MVNENTRMYIPEENHQGSNYGSPRPAHANMNANAAAGLAPEHIPTPGAALSWQAAIDAARQAKLMGSAGNATISTVSSTQRKRQQYGKPKKQGSTTATRPPRALLCLTLKNPIRRACISIVEWKPFEIIILLTIFANCVALAIYIPFPEDDSNATNSNLERVEYLFLIIFTVEAFLKVIAYGLLFHPNAYLRNGWNLLDFIIVVVGLFSAILEQATKADGANALGGKGAGFDVKALRAFRVLRPLRLVSGVPSLQVVLNSIIKAMVPLLHIALLVLFVIIIYAIIGLELFMGKMHKTCYNQEGIADVPAEDDPSPCALETGHGRQCQNGTVCKPGWDGPKHGITNFDNFAFAMLTVFQCITMEGWTDVLYWVNDAVGRDWPWIYFVTLIIIGSFFVLNLVLGVLSGEFSKEREKAKARGDFQKLREKQQLEEDLKGYLDWITQAEDIDPENEDEGMDEEKPRNMSMPTSETESVNTENVAGGDIEGENCGARLAHRISKSKFSRYWRRWNRFCRRKCRAAVKSNVFYWLVIFLVFLNTLTIASEHYNQPNWLTEVQDTANKALLALFTAEMLLKMYSLGLQAYFVSLFNRFDCFVVCGGILETILVETKIMSPLGISVLRCVRLLRIFKITRYWNSLSNLVASLLNSVRSIASLLLLLFLFIIIFSLLGMQLFGGKFNFDEMQTRRSTFDNFPQSLLTVFQILTGEDWNSVMYDGIMAYGGPSFPGMLVCIYFIILFICGNYILLNVFLAIAVDNLADAESLTSAQKEEEEEKERKKLARTASPEKKQELVEKPAVGESKEEKIELKSITADGESPPATKINMDDLQPNENEDKSPYPNPETTGEEDEEEPEMPVGPRPRPLSELHLKEKAVPMPEASAFFIFSSNNRFRLQCHRIVNDTIFTNLILFFILLSSISLAAEDPVQHTSFRNHILFYFDIVFTTIFTIEIALKILGNADYVFTSIFTLEIILKMTAYGAFLHKGSFCRNYFNILDLLVVSVSLISFGIQSSAINVVKILRVLRVLRPLRAINRAKGLKHVVQCVFVAIRTIGNIVIVTTLLQFMFACIGVQLFKGKLYTCSDSSKQTEAECKGNYITYKDGEVDHPIIQPRSWENSKFDFDNVLAAMMALFTVSTFEGWPELLYRSIDSHTEDKGPIYNYRVEISIFFIIYIIIIAFFMMNIFVGFVIVTFQEQGEQEYKNCELDKNQRQCVEYALKARPLRRYIPKNQHQYKVWYVVNSTYFEYLMFVLILLNTICLAMQHYGQSCLFKIAMNILNMLFTGLFTVEMILKLIAFKPKGYFSDPWNVFDFLIVIGSIIDVILSETNHYFCDAWNTFDALIVVGSIVDIAITEVNPAEHTQCSPSMNAEENSRISITFFRLFRVMRLVKLLSRGEGIRTLLWTFIKSFQALPYVALLIVMLFFIYAVIGMQVFGKIALNDTTEINRNNNFQTFPQAVLLLFRCATGEAWQDIMLACMPGKKCAPESEPSNSTEGETPCGSSFAVFYFISFYMLCAFLIINLFVAVIMDNFDYLTRDWSILGPHHLDEFKRIWAEYDPEAKGRIKHLDVVTLLRRIQPPLGFGKLCPHRVACKRLVSMNMPLNSDGTVMFNATLFALVRTALRIKTEGNLEQANEELRAIIKKIWKRTSMKLLDQVVPPAGDDEVTVGKFYATFLIQEYFRKFKKRKEQGLVGKPSQRNALSLQAGLRTLHDIGPEIRRAISGDLTAEEELDKAMKEAVSAASEDDIFRRAGGLFGNHVSYYQSDGRSAFPQTFTTQRPLHINKAGSSQGDTESPSHEKLVDSTFTPSSYSSTGSNANINNANNTALGRLPRPAGYPSTVSTVEGHGPPLSPAIRVQEVAWKLSSNRERHVPMCEDLELRRDSGSAGTQAHCLLLRKANPSRCHSRESQAAMAGQEETSQDETYEVKMNHDTEACSEPSLLSTEMLSYQDDENRQLTLPEEDKRDIRQSPKRGFLRSASLGRRASFHLECLKRQKDRGGDISQKTVLPLHLVHHQALAVAGLSPLLQRSHSPASFPRPFATPPATPGSRGWPPQPVPTLRLEGVESSEKLNSSFPSIHCGSWAETTPGGGGSSAARRVRPVSLMVPSQAGAPGRQFHGSASSLVEAVLISEGLGQFAQDPKFIEVTTQELADACDMTIEEMESAADNILSGGAPQSPNGALLPFVNCRDAGQDRAGGEEDAGCVRARGRPSEEELQDSRVYVSSL

>XP_020644644.1|Pogona_vitticeps_Cav1.2

MFQSIVRQATSTYRPLPAHPPEEPGVKYTGRMVHETQLSCFYIPPGGSNYVSPRPAHANMNANAAAGLAPEHIPTPGAALSWQAAIDAARQAKLMGAAGNATISTASSTQRKRQQYGKQKKQGTTTATRPPRALLCLTLKNPIRRACISIVEWKPFEIIILLTIFANCVALAIYIPFPEDDSNATNSNLERVEYLFLIIFTVEAFLKVIAYGLLFHPNAYLRNGWNLLDFIIVVVGLFSAILEQATKADGVNSIGGKGAGFDVKALRAFRVLRPLRLVSGVPSLQVVLNSIIKAMVPLLHIALLVLFVIIIYAIIGLELFMGKMHKTCYVTGILSDTPAEEEPSPCAPAYAHGRQCQNGSDCRPGWEGPKHGITNFDNFAFAMLTVFQCITMEGWTDVLYWVNDAIGRKWPWIYFVTLIIIGSFFVLNLVLGVLSGEFSKEREKAKARGDFQKLREKQQLEEDLKGYLDWITQAEDIDPENEDEGMDEEKPRNMSMPTSETESVNTDNVTGGDIEGENCGARLAHRISKSKFSRYWRRWNRFCRRKCRAAVKSNVFYWLVIFLVFLNTLTIASEHYNQPDWLTEVQDTANKVLLALFTAEMLLKMYSLGLQAYFVSLFNRFDCFIVCGGILETILVETKIMSPLGISVLRCVRLLRIFKITRYWNSLSNLVASLLNSVRSIASLLLLLFLFIIIFSLLGMQLFGGKFNFDEMQTRRSTFDNFPQSLLTVFQILTGEDWNSVMYDGIMAYGGPSFPGMLVCIYFIILFICGNYILLNVFLAIAVDNLADAESLTSAQKEEEEEKERKKLARTASPEKKQEPEKPAVEEEMKEEKIELKSITADGESPPSNKGNTDEYQPNENEEKNAYPTTETPGEDEEDEPEMPVGPRPRPMSELHLKEKAVPMPEASAFFIFSPSNRFRVHCHRIVNDNIFTNLILFFILLSSISLAAEDPVQHYSVRNQILFYFDIFFTVIFTIEIALKILGNADYVFTSIFTLEIILKMTAYGAFLHKGSFCRNYFNILDLLVVSVSLISFGIQSSAINVVKILRVLRVLRPLRAINRAKGLKHVVQCVFVAIRTIGNIVIVTTLLQFMFACIGVQLFKGKLYSCSDSSKQTEAECKGSFITYKDGEVSQPMIQPRRWENSKFDFDNVLTAMMALFTVSTFEGWPELLYRSIDSHMEDVGPIYNHRVEISIFFIIYIIIIAFFMMNIFVGFVIVTFQEQGEQEYKNCELDKNQRQCVEYALKARPLRRYIPKNQYQYKVWYVVNSTYFEYLMFVLILLNTICLAMQHYGQSCLFKEAMNILNMLFTGLFTVEMVLKLIAFKPKGYFSDPWNVFDFLIVIGSIIDVILSETNHYFCDAWNTFDALIVVGSIVDIAITEVNPAEHTQCSSSMNAEENSRISITFFRLFRVMRLVKLLSRGEGIRTLLWTFIKSFQALPYVALLIVMLFFIYAVIGMQVFGKIALDDTTDINRNNNFQTFPQAVLLLFRCATGEAWQEIMLACLPDKRCDPESFEPNNSIEGEHSCGSSFAVFYFISFYMLCAFLIINLFVAVIMDNFDYLTRDWSILGPHHLDEFKRIWAEYDPEAKGRIKHLDVVTLLRRIQPPLGFGKLCPHRVACKRLVSMNMPLNSDGTVMFNATLFALVRTALRIKTEGNLEQANEELRAIIKKIWKRTSMKLLDQVVPPAGDDEVTVGKFYATFLIQEYFRKFKKRKEQGLVGKPSQRNALSLQAGLRTLHDIGPEIRRAISGDLTAEEELDKAMKEAVSAASEDDIFRRAGGLFGNHVNYYQSDGRSSFPQTFTTQRPLHINKSGNSQGDTESPSHEKLVDSTFTPSSYSSSGSNANINNANNTALSRFSSPPNYPSTVSTVEGHNTPLSPPARIREVSWKLGCRRSCSRDSQTAIVCEEQEVSQEEETYDKQRKEETAYCIEMLSYQDDEHRQLTPHEDNKGEETQYSPKSGFLHSLSLSRRASFHLECQRKQKKQNTEVNQKTVLPVHLVHHQALAVAGLSPHLQRSHSPLRYSRSCATPPTTLCTQDWPQQPIKTLQLDGEESNEKLNNSFPSVYCSSLYSDSTSARRTQKARPVSLTVPSQTRVCGRQFHGSANSLVEAVLISEGLIQFAQDPKFIETTTQELADACDLTIEEMENAADNILNGNSKPSPNGNLLPFVNCRDPGQDCMGDQAEAAQNLDCRKSQEEPVDSRIYISSL

>XP_015278355.1|Gekko_japonicus_Cav1.2

MNANAAAGLAPEHIPTPGAALSWQAAIDAARQAKLMGAAGNATISTASSTQRKRQQYGKQKKQGTTTATRPPRALLCLTLKNPIRRACISIVEWKYPFXPFEIIILLTIFANCVALAIYIPFPEDDSNATNSNLERVEYLFLIIFTVEAFLKIITYGLLFHPNAYLRNGWNLLDFIIVVVGXVLAILEQATKADGVNSIGGKGAGFDVKALRAFRVLRPLRLVSGVPSLQVVLNSIIKAMVPLLHIALLVLFVIIIYAIIGLELFMGKMHKTCYLLGVTDTPAEEDPSPCAPHLAHGRQCQNGTECRAGWEGPKHGITNFDNFAFAMLTVFQCITMEGWTDVLYWMQDAMGYELPWVYFVSLVIFGSFFVLNLVLGVLSGEFSKEREKAKARGDFQKLREKQQLEEDLKGYLDWITQAEDIDPENEDEGMDEEKPRNMSMPTSETESVNTDNVAGGDIEGENCGARLAHRISKSKFSRYWRRWNRFCRRKCRAAVKSNVFYWLVIFLVFLNTLTIASEHYNQSDWLTEVQDTANKVLLALFTAEMLLKMYSLGLQAYFVSLFNRFDCFIVCGGILETILVETKIMSPLGISVLRCVRLLRIFKITRYWNSLSNLVASLLNSVRSIASLLLLLFLFIIIFSLLGMQLFGGKFNFDEMQTRRSTFDNFPQSLLTVFQILTGEDWNSVMYDGIMAYGGPSFPGMLVCIYFIILFICGNYILLNVFLASTVDNVADAESLTSAQKEEEEEKERKKLARTASPEKKQEPEKPAVEGETKEEKIELKSITADGESPPSNKSNVDEYQPNENEEKNPYPTTETPGEEEEEEPEMPVGPRPRPMSELHLKEKAVPMPDASAFFIFSPSNRFRVHCHRIVNNNIFTNLILFFILLSSISLAAEDPVRHSSVRNQILFYFDIFFTVIFTIEIALKMTAYGAFLHKGSFCRNYFNILDLLVVSVSLISFGIQSSAINVVKILRVLRVLRPLRAINRAKGLKHVVQCVFVAIRTIGNIVIVTTLLQFMFACIGVQLFKGKLYSCSDSSKQTEAECKGKFITYKDGEVSHPILNIRNWENSKFDFDNVLTAMMALFTVSTFEGWPELLYRSIDSHLEDVGPIYNHRVEISIFFIIYIIIIAFFMMNIFVGFVIVTFQEQGEQEYKNCELDKNQRQCVEYALKARPLRRYIPKNKYQYKVWYVVNSTYFEYLIFILIMLNTICLAMQHYGQSCLFKEAMNILNMLFTGLFTVEMVLKLIAFKPKGYFSDPWNVFDFLIVIGSIIDVILSETNPAEHTQCSSSMNAEENSRISITFFRLFRVMRLVKLLSRGEGIRTLLWTFIKSFQALPYVALLIVMLFFIYAVIGMQVFGKIALNDTTEINRNNNFQTFPQAVLLLFRCATGEAWQEIMLACLPDKKCDSDSLEQNNSTEDEYSCGSSFAIFYFISFYMLCAFLIINLFVAVIMDNFDYLTRDWSILGPHHLDEFKRIWAEYDPEAKGRIKHLDVVTLLRRIQPPLGFGKLCPHRVACKRLVSMNMPLNSDGTVMFNATLFALVRTALRIKTEGNLEQANEELRAIIKKIWKRTSMKLLDQVVPPAGDDEVTVGKFYATFLIQEYFRKFKKRKEQGLVGKPSQRNALSLQAGLRTLHDIGPEIRRAISGDLTAEEELDKAMKEAVSAASEDDIFRRAGGLFGNHVSYYQSDGRSSFPQTFTTQRPLHINKSSNNQGDTESPSHEKLVDSTFTPSSYSSSGSNANINNANNTALSRFPSPPRYPSTVSTVEGHSTPLSPSARIREPPWKLGCRRSCSRDSQVAIVCEEEPSQEEETYDERLNEEVEYCSEPSLLSTEMLSYQDDEHRQLTPPENCKGEDTRHSPKSGFLRSASLNRRASFHLECLKRQKNPSAVGPQKTVLPLHLVHQQALAVAGMSPLLQRCHSPVRASRSCTTPFTTPCTQSWPQQPIKTLQLDGEESNEKLNSSFPSVQCSSLYSDGTNSGSTRKVRPVSLTVPSQTTECGRQFHGSANSLVEAVLISEGLVQFAQDPKFIEVTTQELADACELTIEEMENAADNILNGNSKPSPNGKLLPFVNCRDSGQDCMGEEEAAAQNPDCKKSQEESKDSRIYISSL

>XP_008162493.1|Chrysemys_picta_bellii_Cav1.2

MFQAFVRQATSAYRPLPSLLPGEPEMKYKGKLVHETQLNCFYIQPGGSNYGSPRPAHANMNANAAAGLAPEHIPTPGAALSWQAAIDAARQAKLMGSAGNTTISTASSTQRKRQQYGKQKKQGTTTATRPPRALLCLTLKNPIRRACISIVEWKPFEIIILLTIFANCVALAIYIPFPEDDSNATNSNLERVEYLFLIIFTVEAFLKVIAYGLLFHPNAYLRNGWNLLDFIIVVVGLFSAILEQATKADGVNSIGGKGAGFDVKALRAFRVLRPLRLVSGVPSLQVVLNSIIKAMVPLLHIALLVLFVIIIYAIIGLELFMGKMHKTCYFLQGGLPAEEEASPCAPLSAHGRQCQNGTECKPGWEGPKHGITNFDNFAFAMLTVFQCITMEGWTDVLYWVNDAIGRDWPWIYFVTLIIIGSFFVLNLVLGVLSGEFSKEREKAKARGDFQKLREKQQLEEDLKGYLDWITQAEDIDPENEDEGMDEEKPRNMSMPTSETESVNTDNVAGADIEGENCGARLAHRISKSKFSRYWRRWNRFCRRKCRAAVKSNIFYWLVIFLVFLNTLTIASEHYNQPYWLTEVQDTANKVLLALFTAEMLLKMYSLGLQAYFVSLFNRFDCFIVCGGILETILVETKIMSPLGISVLRCVRLLRIFKITRYWNSLSNLVASLLNSVRSIASLLLLLFLFIIIFSLLGMQLFGGKFNFDEMQTKRSTFDNFPQSLLTVFQILTGEDWNSVMYDGIMAYGGPSFPGMLVCIYFIILFICGNYILLNVFLAIAVDNLADAESLTSAQKEEEEEKERKKLARTASPEKKQEIEKPAVEEETKEEKIELKSITADGESPPTTKINVDDYQPNENEEKNPYPTTEAPAEEEEEEPEMPVGPRPRPMSELHLKEKAVPMPDASAFFIFSPTNRYLRSCYAAEDPVKHYSFRNQILFYFDIVFTVIFTIEIALKILGNADYVFTSIFTLEIILKMTAYGAFLHKGSFCRNYFNILDLLVVSVSLISFGIQSSAINVVKILRVLRVLRPLRAINRAKGLKHVVQCVFVAIRTIGNIVIVTTLLQFMFACIGVQLFKGKLYSCSDSSKQTEAECRGHYISYKDGEVTQPIIQSRNWENSKFNFDNVLTAMMALFTVSTFEGWPELLYKSIDSHMEDVGPIYNHRVEISIFFIIYIIIIAFFMMNIFVGFVIVTFQEQGEQEYKNCELDKNQRQCVEYALKARPLRRYIPKNQYQYKVWYVVNSTYFEYLMFVLILLNTICLAMQHYGQSCPFKQAMNILNMLFTGLFTVEMILKLIAFKPKGYFSDPWNVFDFLIVIGSIIDVILSETNHYFCDAWNTFDALIVVGSIVDIAITEVNPAEHTQCSSSMNAEENSRISITFFRLFRVMRLVKLLSRGEGIRTLLWTFIKSFQALPYVALLIVMLFFIYAVIGMQVFGKIALNDTSYINRNNNFQTFPQAVLLLFRCATGEGWQDIMLDCLPDKKCDPESEPTNSTEGDHSCGSSFSIFYFISFYMLCAFLIINLFVAVIMDNFDYLTRDWSILGPHHLDEFKRIWAEYDPEAKGRIKHLDVVTLLRRIQPPLGFGKLCPHRVACKRLVSMNMPLNSDGTVMFNATLFALVRTALRIKTEGNLEQANEELRAIIKKIWKRTSMKLLDQVVPPAGDDEVTVGKFYATFLIQEYFRKFKKRKEQGLVGKLSQRNALSLQAGLRTLHDIGPEIRRAISGDLTAEEELDKAMKEAVSAASEDDIFRRAGGLFGNHVSYYQSDGRSSFPQTFTTQRPLHINKSGNNQGDTESPSHEKLVDSTFTPSSYSSSGSNANINNANNTALSRFHSPPSYPSTVSTVEGHGTPLSPTIRVQEAPWKLTSKRTHYYETLGRSSSRDSQLAIVCQEEVSQDETYDENLNEDVEYCSEPSLISTEMLSYQDDENRQLTPPENNKGEGSRQSPKRGFLRSSSLGRRASFHLECLKRQKNQGVDVSQKTVLPLHLVHHQALAVAGLSPLLQRSHSPTTFSPRLCATPPAVPCSRGWPQKTIPTLRLDGVESSEKLNSSFPSIHCSSWYSDSTSCSSPRRARPVSLTVPSQTGGSSRQFHGSAGSLVEAVLISEGLMQFAQDPKFIEVTTQELADACDMTIEEMENAADNILNGNSKQSPNGNLLPFVNCRDPGQDSAGEEEEEVQNPDCRKSQEELQDSRIYVSSL

>XP_009298610.1|Danio_rerio_Cav1.2

MVNESKNMYIPEDTLENHQGSNYSSPSLAPVPSLNEDEHVGGGGGVLGLAPEHIPTPGAPLSWQAAIDAARQAKLMGTTGAPISTASSTQRKRQHYTKPKKQASTASTRPPRALLCLTLKNPIRRACINIVEWKPFEIIILMTIFANCVALAVYIPFPEDDSNATNSNLERVEYLFLIIFTVEAFLKVIAYGLLCHPNAYLRNGWNLLDFIIVVVGLFSAILEQATKGDGGTSMGGKAAGFDVKALRAFRVLRPLRLVSGVPSLQVVLNSIIKAMVPLLHIALLVLFVIIIYAIIGLELFMGKMHRTCFFYKDGHKGHIAEEKPAPCAPSSAHGRHCSPPNITQCMMGWEGPNDGITNFDNFAFAMLTVFQCITMEGWTDVLYWMQDAMGYELPWVYFVSLVIFGSFFVLNLVLGVLSGEFSKEREKAKARGDFQKLREKQQLEEDLKGYLDWITQAEDIDPENDDEGLDDDKPRNLSMPASENESVNTDNAPAGDMEGETCCTRMANRISKSKFSRYSRRWNRLCRRKCRAAVKSNVFYWLVIFLVFLNTLTIASEHHQQPEWLTNVQDIANKVLLALFTGEMLLKMYSLGLQAYFVSLFNRFDSFVVCGGILETILVETKIMSPLGISVLRCVRLLRIFKITRYWNSLSNLVASLLNSVRSIASLLLLLFLFIIIFSLLGMQLFGGKFNFDETRRSTFDNFPQSLLTVFQILTGEDWNSVMYDGIMAYGGPSFPGMLVCIYFIILFICGNYILLNVFLAIAVDNLADAESLTSAQKEEEEEKERKKLARTASPEKRQNSEKPPLEDEKKEEKIELKSITSDGETPTATKINIDEYTGEDNEEKNPYPVNDFPAGEDDEEEPEMPVGPRPRPLSDIQLKEKAVPMPQAKAFFIFSPSNKFRVLCHKIVNHNIFTNLILFFILLSSISLAAEDPVKNDSFRNQILGYADYVFTGIFTIEIILKMTAYGAFLHKGSFCRNYFNILDLVVVSVSLISSGIQSSAINVVKILRVLRVLRPLRAINRAKGLKHVVQCVFVAIRTIGNIVIVTSLLQFMFACIGVQLFKGKFFYCTDTSKQTQAECRGAYILYKDGNVGKPEKAQRSWENSDFNFDDVLQGMMALFAVSTFEGWPGLLYRAIDSHAEDVGPIYNYRVVISIFFIIYIIIIAFFMMNIFVGFVIVTFQEQGEQEYKNCELDKNQRQCVEYALKARPLRRYIPKNPYQYKVWYVVNSTYFEYLMFTLILLNTICLAMQHHGQSQSFNKAMNILNMLFTGLFTVEMILKLIAFKPRGYFSDPWNVFDFLIVIGSIIDVILSEINPADPSSSPPSSVVRPMGLQNTEDNARISITFFRLFRVMRLVKLLSRGEGIRTLLWTFIKSFQALPYVALLIVMLFFIYAVIGMQMFGKIALRDNSQINRNNNFQTFPQAVLLLFRCATGEAWQEIMLACSPNRPCEKGSEINHSSEDCGSHFAIFYFVSFYMLCAFLIINLFVAVIMDNFDYLTRDWSILGPHHLDEFKRIWAEYDPEAKGRIKHLDVVTLLRRIQPPLGFGKLCPHRVACKRLVSMNMPLNSDGTVMFNATLFALVRTALRIKTEGNLEQANEELRAIVKKIWKRTSMKLLDQVVPPAGDDEVTVGKFYATFLIQEYFRKFKKRKEQGLVAKIPPKTALSLQAGLRTLHDMGPEIRRAISGDLTVEEELERAMKETVCAASEDDIFRVKRSGGLFGNHVNYYHQSDGHVSFPQSFTTQRPLHISKSGSPGEAESPSHQKLVDSTFTPSSYSSSGSNANINNANNTAIGHRYPKPTVSTVDGQTGPPLTTIPLPRPTWCFPNKRSCFYDTFMRSDSSDSRLPIIRREEASTDETYDETFLDERDQAMLSMDMLEFQDEESKQLAPMVEAEVGEERRPWQSPRRRAFLCPTALGRRSSFHLECLRKHNRPDVSQKTALPLHLVHHQALAVAGLSPLLRRSHSPTLFTRLCSTPPASPSGRSGGGPCYQPVPSLRLEGSGSYEKLNSSMPSVNCSSWYSDSNGNHSGRAQRPVSLTVPPVTRRDSISLAHGSAGSLVEAVLISEGLGRYAHDPSFIQVAKQEIAEACDMTMEEMENAADNILNANAPPNANGNLLPFIQCRDTGSQESRCSLSLGLSPATGSDGALEAELEESEGAGQRNSPLMEDEDMECVTSL

>XP_020659553.1|Pogona_vitticeps_Cav1.3

MSTTPAQPVGSLSQRKRQQYAKSKKQGNTSNSRPARALFCLSLNNPIRRACISIVEWKPFDIFILLAIFANCVALAVYIPFPEDDSNSTNHNLEKVEYAFLIIFTIETFLKIIAYGLLLHPNAYVRNGWNLLDFVIVIVGLFSVILEQLTKEAEGGSHSGGKPGGFDVKALRAFRVLRPLRLVSGVPSLQVVLNSIIKAMVPLLHIALLVLFVIIIYAIIGLELFIGKMHKSCYFDDTDILAEEDPAPCAFSGSGRQCPINGTECKGGWPGPNGGITNFDNFGYAMLTVFQCITMEGWTDVLYWVNDAIGSEWPWIYFVSLIILGSFFVLNLVLGVLSGEFSKEREKAKARGDFQKLREKQQLEEDLKGYLDWITQAEDIDPENEEEGDGEGKRNTSMHASETESVNTENVGGDGENPPCCGSLCQVISKSKFSRRWRRWNRFNRRRCRAAVKSVSFYWLVIVLVFLNTLTISSEHYDQPEWLTQIQDIANKVLLALFTCEMLVKMYSLGLQSYFVSLFNRFDCFVVCGGIVETILVELEIMSPLGISVFRCVRLLRIFKVTRHWTSLSNLVASLLNSMKSIASLLLLLFLFIIIFSLLGMQLFGGKFNFDETQTKRSTFDNFPQALLTVFQILTGEDWNAVMYDGIMAYGGPSSSGMIVCVYFIILFICGNYILLNVFLAIAVDNLADAESLNTAQKEEAEEKEKKKAARKESLDIKKGDKLEGEQKKSKDSKVTINDYGEGEDEDKDPYPPCDVPGEEEEEEDEEEPEVPAGPRPRRISELNMKEKMTPIPEGSSFFLFSNTNPIRVGCHRLINHHIFTNLILVFIMLSSASLAAEDPIRSHSFRNNILGYFDYAFTAIFTVEILLKLTVFGAFLHKGSFCRNYFNLLDLLVVGVSLVSFGIQSSAISVVKILRVLRVLRPLRAINRAKGLKHVVQCVFVAIRTIGNIMIVTTLLQFMFACIGVQLFKGKFYRCTDEAKQNPVECRGIFIVYKDGDVDSPMVRERIWQNSDFNFDNVLTAMMALFTVSTFEGWPALLYKAIDSNGENIGPIYNYRVEISIFFIIYIIIIAFFMMNIFVGFVIVTFQEQGEQEYKNCELDKNQRQCVEYALKARPLRRYIPKNPYQYKFWYMVNSTVFEYIMFVLIMLNTLCLAMQHYGQSKLFNDAMDILNMVFTAVFTVEMVLKLIAFKPKGYFSDAWNTFDSLVVLGSIVDIVLSEADHYFTDAWNTFDALIVVGSVVDIAITEVNNSEDSARISITFFRLFRVMRLVKLLSRGEGIRTLLWTFIKSFQALPYVALLIAMLFFIYAVIGMQVFGKVALKDGSQINRNNNFQTFPQAVLLLFRCATGEAWQEIMLACMPGKRCDPESDFNPGEEYTCGSNFAIIYFISFYMLCAFLIINLFVAVIMDNFDYLTRDWSILGPHHLDEFKRIWSEYDPEAKGRIKHLDVVTLLRRIQPPLGFGKLCPHRVACKRLVAMNMPLNSDGTVMFNATLFALVRTALKIKTEGNLEQANEELRAVIKKIWKKTSMKLLDQVVPPAGDDEVTVGKFYATFLIQDYFRKFKKRKEQGLVGKYPAKNSTIALQAGLRTLHDIGPEIRRAISCDLQEDEPEQNHHEEEEVYKRNGALFGNHVNHVSNETRDAFQQINTTHRPLHVQRPSIPSSGETTEKNVHPPAGNSVFHNHHSHHSHHSHHNHNSVGKHVPNSTNANLNNANMSKVANGRHIRSGSHEHISETGHRSHSRRDHHEKRRRSHKRRSRYYEAYIRSESGDRHLPTICREDHEIRDYYNDDCYLDDQEYYSGEEYYEEDSMLSRGRHTHDFHSRYHCNDLDFERPKGYHHPHGFYEEDESQMCYDSKRSPRRRLLPPTPPPHRRSSFNFECLRRQSSHDEIPLSPTFHHRTALPLHLMQQQVMAVAGLDATKAHRHSPSRSTRSWATPPATPPNPDRSPYYTPLIQVDQADSTEYMNGSLPSLHRSSWYTDDPDIAYRTFTPANLTVPSDFRHKHSDKQRSADSLVEAVLISEGLGRYAKDPKFVSATKHEIADACDMTIDEMESAASNLLNGSISNGANGDMFPILNRQDYELQDYGPGYSDEEPETGKYEEDLADEMICITTL

>XP_015274327.1|Gekko_japonicus_Cav1.3

MLVEQTVGGKTMSYSSEKVEYAFLIIFTIETFLKIIAYGLLLHPNAYVRNGWNLLDFVIVIVGLFSVILEQLTKEAEGGTHSGGKSGGFDVKALRAFRVLRPLRLVSGVPSLQVVLNSIIKAMVPLLHIALLVLFVIIIYAIIGLELFIGKMHKSCYFDDADILAEDDPAPCAFSGSGRQCSTNGTECKSGWVGPNGGITNFDNFAFAMLTVFQCITMEGWTDVLYWVNDAIGCEWPWIYFVSLIILGSFFVLNLVLGVLSGEFSKEREKAKARGDFQKLREKQQLEEDLKGYLDWITQAEDIDPENEEDGDEEGKRNRVTLEDLLNDKKKSRFNCFRGATNKHASMQASETESVNTENVGGEGEGHVCCGALCQVISKSRFSRRWRRWNRFNRRRCRAAVKSVSFYWLVIVLVFLNTLTISSEHYNQPDWLTQIQDIANKVLLALFTCEMLVKMYSLGLQSYFVSLFNRFDCFVVCGGIVETILVELEIMSPLGISVFRCVRLLRIFKVTRHWTSLSNLVASLLNSMKSIASLLLLLFLFIIIFSLLGMQLFGGKFNFDETQTKRSTFDNFPQALLTVFQILTGEDWNAVMYDGIMAYGGPSSSGMVVCIYFIILFICGNYILLNVFLAIAVDNLADAESLNTAQKEEAEEKERKKNARKESLDTKKGDKTESEQKKAKDTKVTINEYGEGEEEDKDPYPPCDVPVDEEEEEEEEEEEPEVPAGPRPRRISELNMKEKMTPIPEGSSFFIFSNTNPIRVGCHRLINHHIFTNLILVFIMLSSASLAAEDPIRSHSFRNNILGYADYVFTSMFTFEIILKLTVFGAFLHKGSFCRNYFNLLDLLVVGVSLVSFGIQSSAISVVKILRVLRVLRPLRAINRAKGLKHVVQCVFVAIRTIGNIMIVTTLLQFMFACIGVQLFKGKFYRCTDEAKQNPEECRGIFIVYKDGDVDSPMVRERIWQNSDFNFDNVLAAMMALFTVSTFEGWPALLYKAIDSNGENIGPVYNYRVEISIFFIIYIIIIAFFMMNIFVGFVIVTFQEQGEQEYKNCELDKNQRQCVEYALKARPLRRYIPKNPYQYKFWYVVNSTGFEYIMFVLIMLNTLCLAMQHYGQSKLFNDAMDVLNMVFTAVFTIEMVLKLIAFKPKICVPKKKRMLGYFSDAWNSFDSLVVLGSIVDIVLSEADPKPTETVTTDESGNNEDSARISITFFRLFRVMRLVKLLSRGEGIRTLLWTFIKSFQALPYVALLIAMLFFIYAVIGMQVFGKVALRDGTQINRNNNFQTFLQAVLLLFRCATGEAWQEIMLACLPGKRCDEDSDYNPGEENTCGSNFAIIYFITFYMLCAFLIINLFVAVIMDNFDYLTRDWSILGPHHLDEFKRIWSEYDPEAKGRIKHLDVVTLLRRIQPPLGFGKLCPHRVACKRLVAMNMPLNSDGTVMFNATLFALVRTALKIKTEGNLEQANEELRAVIKKIWKKTSMKLLDQVVPPAGDDEVTVGKFYATFLIQDYFRKFKKRKEQGLVGKHPAKNTTVALQAGLRTLHDIGPEIRRAISCDLQEDEPEQNHHEEEEVYKRNGALFGNHVNHVPNDTRESFQQINTTHRPLHVQRPSIPSANETTEKNVHPSTGNSVFHNHHSHHNHHHHNSIGKHVPNSTNANLNNANMSKVVNGRHINTGSHENMTETGHRLHSRSDHHEKRRRPSNRRARYYEAYIKSESGDRHLPTICREDHEVRDYYNDDHYLDDQEYFSGEEYYEEDSMLSGSRHAHEYHSRYYYNDSDFERPKGYHHPHGFYEDDESQMCYDSKRPPRRRLLPPTPTPNRRSSFNFECLRRQGSQDEIPLSPSFYHRTALPLHLMQQQVMAVAGLDATKAHRHSPCRSTRSWATPPATPPNRDRTPYYTPLIQVDRAESTEHMNGSLPSLHRSSWYTDDPDIAYRTFTPANLTVPSDFRHKHSDKQRSADSLVEAVLISEGLGRYAKDPKFVSATKHEIADACDMTIDEMESAASNLLNGNISNGTNGDMFPILNRQDYELQDFGPGYSDEEPDTGKYEEDLADEMICITTL

>XP_023956012.1|Chrysemys_picta_bellii_Cav1.3

MHHHQQQEQYPEEANYASSTRIPLPGDGPTIQSNSSAPSKQTVLSWQAAIDAARQAKAAQTMSTTTAQPVGSLSQRKRQQYAKSKKQGNTSNSRPPRALFCLSLNNPIRRACISLVEWKPFDIFILLAIFANCVALAVYIPFPEDDSNSTNHNLEKVEYAFLIIFTIETFLKIIAYGLLLHPNAYVRNGWNLLDFVIVVVGLFSVILEQLTKETEGGSHSGGKPGGFDVKALRAFRVLRPLRLVSGVPSLQVVLNSIIKAMVPLLHIALLVLFVIIIYAIIGLELFIGKMHKSCFLVDSDILVEDDPAPCAFSGNGRQCAINGTECRGGWVGPNGGITNFDNFAFAMLTVFQCITMEGWTDVLYWVNDAIGCEWPWIYFVSLIILGSFFVLNLVLGVLSGEFSKEREKAKARGDFQKLREKQQLEEDLKGYLDWITQAEDIDPENEEEGDEEGKRNTSMPTSETESVNTENVSGEGESPACCGSLCQTISKSKFSRRWRRWNRFNRRKCRAAVKSVSFYWLVIVLVFLNTLTISSEHYNQPDWLTQVQDTANKVLLALFTCEMLIKMYSLGLQAYFVSLFNRFDCFVVCGGIVETILVELEIMSPLGISVFRCVRLLRIFKVTRHWTSLSNLVASLLNSMKSIASLLLLLFLFIIIFSLLGMQLFGGKFNFDETQTKRSTFDNFPQALLTVFQILTGEDWNAVMYDGIMAYGGPSSSGMVVCIYFIILFICGNYILLNVFLAIAVDNLADAESLNTAQKEEAEEKQRKKNARKESLENKKGDKSEGDQKKAKDNKVTIAEYREGEDEDKDPYPPCDVPVGEDEEDEEDEPEVPVGPRPRRISELNMKEKITPIPEGSAFFIFSSTNPIRVGCHRLINHHIFTNLILVFIMLSSVSLAAEDPIRSHSFRNNILGYFDYAFTAIFTVEILLKILGYADYVFTSMFTFEIILKMTAFGAFLHKGSFCRNYFNLLDLLVVGVSLVSFGIQSSAISVVKILRVLRVLRPLRAINRAKGLKHVVQCVFVAIRTIGNIMIVTTLLQFMFACIGVQLFKGKFYRCTDEAKQNPEDCRGIFIVYKDGDVDNPMVRERVWQNSDFNFDNVLSAMMALFTVSTFEGWPALLYKAIDSNAENIGPVYNYRVEISIFFIIYIIIIAFFMMNIFVGFVIVTFQEQGEQEYKNCELDKNQRQCVEYALKARPLRRYIPKNPYQYKFWYMVNSTGFEYIMFVLIMLNTLCLAMQHYGQSKLFNDAMDILNMVFTGVFTVEMVLKLIAFKPKIFVRKKERWLGYFSDAWNAFDSLIVIGSIVDVVLSEADHYFTDAWNTFDALIVVGSVVDIAITEVNPKPTETVTTDESGNSEDSARISITFFRLFRVMRLVKLLSRGEGIRTLLWTFIKSFQALPYVALLIAMLFFIYAVIGMQVFGKVAMRDNNQINRNNNFQTFPQAVLLLFRCATGEAWQEIMLACLPGKRCDPDSDYSPGEEFTCGSNFSIIYFISFYMLCAFLIINLFVAVIMDNFDYLTRDWSILGPHHLDEFKRIWSEYDPEAKGRIKHLDVVTLLRRIQPPLGFGKLCPHRVACKRLVAMNMPLNSDGTVMFNATLFALVRTALKIKTEGNLEQANEELRAVIKKIWKKTSMKLLDQVVPPAGDDEVTVGKFYATFLIQDYFRKFKKRKEQGLVGKYPAKNTTIALQAGLRTLHDIGPEIRRAISCDLQDDEPEENNHEEEEDIYKRNGALFGNHINHVSSDRRDSFQQINTTHRPLHVQRPSIPSASDTEKNMYHQASNSVFHNHHNHNSIGKHVPNSTNANLNNANMSKVANGRHPNIGNHEHRSENGYHSYSRADHDRHRRSNSKRTRYYETYIRSDSGDGHLPTICREDHEVRDYCNDDHYMGEQEYFSGEEYYEEDYMLSGSRHTYDYHNRYHCNDLDFERPKGYHHPHGFFEEDDSQICYDSKRSPRRRLLPPTPTSNRRSSFNFECLRRQSSQDEIPLSPTFHHRTALPLHLMQQQVMAVAGLDSSKAHKHSPSRSTRSWATPPATPPNRDRTPYYTPLIQVDRAESTEQMNGSLPSLNRSSWYTDDPDISYRTFTPANLTVPNDFRHKHSDKQRSADSLVEAVLISEGLGRYAKDPKFVSATKHEIADACDMTIDEMESAASNLLNGNISNGTNGDMFPILSRQDYELQDFGPGYSDEEPDTGRYEEDLADEMICITTL

>XP_015148473.1|Gallus_gallus_Cav1.3

MQHHQQQQPEQHPEEANYASSTRIPLPGDGPTTQSNSSAPSKQTVLSWQAAIDAARQAKAAQNMNTTTAQPVGSLSQRKRQQYAKSKKQGNTSNSRPPRALFCLSLNNPIRRACISLVEWKPFDIFILLSIFANCVALAVYIPFPEDDSNSTNHNLEKVEYAFLIIFTVETFLKIIAYGLLLHPNAYVRNGWNLLDFVIVVVGLFSVILEQLTKETEGGSHSGGKPGGFDVKALRAFRVLRPLRLVSGVPSLQVVLNSIIKAMVPLLHIALLVLFVIIIYAIIGLELFIGKMHKSCFLIDSDILVEEDPAPCAFSGNGRQCVMNGTECKGGWVGPNGGITNFDNFAFAMLTVFQCITMEGWTDVLYWMNDAMGFELPWVYFVSLVIFGSFFVLNLVLGVLSGEFSKEREKAKARGDFQKLREKQQLEEDLKGYLDWITQAEDIDPENDEEADEEGKRNTSMPTSETESVNTENVSGEGENPACCGSLCQTISKSKFSRRWRRWNRFNRRKCRAAVKSVTFYWLVIVLVFLNTLTISSEHYNQPDWLTQIQDIANKVLLALFTCEMLVKMYSLGLQAYFVSLFNRFDCFVVCGGIVETILVELEIMSPLGISVFRCVRLLRIFKVTRHWASLSNLVASLLNSMKSIASLLLLLFLFIIIFSLLGMQLFGGKFNFDETQTKRSTFDNFPQALLTVFQILTGEDWNAVMYDGIMAYGGPSSSGMIVCIYFIILFICGNYILLNVFLAIAVDNLADAESLNTAQKEEAEEKERKKNARKESLENKKSEKSEGDQKKPKDSKVTIAEYGEGEDEDKDPYPPCDVPVGEDEEDEEDEPEVPAGPRPRRISELNMKEKITPIPEGSAFFIFSSTNPIRVGCHRLINHHIFTNLILVFIMLSSVSLAAEDPIRSHSFRNNILGYFDYAFTAIFTVEILLKMTAFGAFLHKGSFCRNYFNLLDLLVVGVSLVSFGIQSSAISVVKILRVLRVLRPLRAINRAKGLKHVVQCVFVAIRTIGNIMIVTTLLQFMFACIGVQLFKGKFYKCTDEAKQNPEECRGIYIVYKDGDVDNPMVKERVWQNSDFNFDNVLSAMMALFTVSTFEGWPALLYKAIDSNGENVGPVYNYRVEISIFFIIYIIIIAFFMMNIFVGFVIVTFQEQGEQEYKNCELDKNQRQCVEYALKARPLRRYIPKNPYQYKFWYVVNSTGFEYIMFVLIMLNTLCLAMQHYGQSKLFNDAMDIMNMVFTGVFTVEMVLKLIAFKPKGYFSDAWNTFDSLIVIGSIVDVVLSEADHYFTDAWNTFDALIVVGSVVDIAITEVNPKPTETVTTDESGNSEDSARISITFFRLFRVMRLVKLLSRGEGIRTLLWTFIKSFQALPYVALLIAMLFFIYAVIGMQVFGKVAMRDNNQINRNNNFQTFPQAVLLLFRCATGEAWQEIMLACLPGKRCDPESDYNPGEEYTCGSNFAIIYFISFYMLCAFLIINLFVAVIMDNFDYLTRDWSILGPHHLDEFKRIWSEYDPEAKGRIKHLDVVTLLRRIQPPLGFGKLCPHRVACKRLVAMNMPLNSDGTVMFNATLFALVRTALKIKTEGNLEQANEELRAVIKKIWKKTSMKLLDQVVPPAGDDEVTVGKFYATFLIQDYFRKFKKRKEQGLVGKYPAKNTTIALQAGLRTLHDIGPEIRRAISCDLQDDEPEENNPDEEEEVYKRNGALFGNHINHISSDRRDSFQQINTTHRPLHVQRPSIPSASDTEKNIYPHTGNSVYHNHHNHNSVGKQVPNSTNANLNNANVSKVVHGKHANFGSHEHRSENGYHSYSRADHEKRRRPSSRRTRYYETYIRSDSGDGRRPTICREERDIRDYCNDDHYLGEQEYYSGEEYYEEDSMLSGNRHVYDYHCRHHCHDSDFERPKGYHHPHGFFEEDDSQTCYDTKRSPRRRLLPPTPASNRRSSFNFECLRRQSSQDDIPLSPNFHHRTALPLHLMQQQVMAVAGLDSSKAHKHSPSRSTRSWATPPATPPNRDHTPYYTPLIQVDRAESTEHMNGSLPSLHRSSWYTDDPDISYRTFTPANLTVPNDFRHKHSDKQRSADSLVEAVLISEGLGRYAKDPKFVSATKHEIADACDMTIDEMESAASNLLNGNISNGTNGDMFPILSRQDYELQDFGPGYSDEEPEPGRYEEDLADEMICITSL

>NP_001122312.1|Homo_sapiens_Cav1.3

MMMMMMMKKMQHQRQQQADHANEANYARGTRLPLSGEGPTSQPNSSKQTVLSWQAAIDAARQAKAAQTMSTSAPPPVGSLSQRKRQQYAKSKKQGNSSNSRPARALFCLSLNNPIRRACISIVEWKPFDIFILLAIFANCVALAIYIPFPEDDSNSTNHNLEKVEYAFLIIFTVETFLKIIAYGLLLHPNAYVRNGWNLLDFVIVIVGLFSVILEQLTKETEGGNHSSGKSGGFDVKALRAFRVLRPLRLVSGVPSLQVVLNSIIKAMVPLLHIALLVLFVIIIYAIIGLELFIGKMHKTCFFADSDIVAEEDPAPCAFSGNGRQCTANGTECRSGWVGPNGGITNFDNFAFAMLTVFQCITMEGWTDVLYWMNDAMGFELPWVYFVSLVIFGSFFVLNLVLGVLSGEFSKEREKAKARGDFQKLREKQQLEEDLKGYLDWITQAEDIDPENEEEGGEEGKRNTSMPTSETESVNTENVSGEGENRGCCGSLCQAISKSKLSRRWRRWNRFNRRRCRAAVKSVTFYWLVIVLVFLNTLTISSEHYNQPDWLTQIQDIANKVLLALFTCEMLVKMYSLGLQAYFVSLFNRFDCFVVCGGITETILVELEIMSPLGISVFRCVRLLRIFKVTRHWTSLSNLVASLLNSMKSIASLLLLLFLFIIIFSLLGMQLFGGKFNFDETQTKRSTFDNFPQALLTVFQILTGEDWNAVMYDGIMAYGGPSSSGMIVCIYFIILFICGNYILLNVFLAIAVDNLADAESLNTAQKEEAEEKERKKIARKESLENKKNNKPEVNQIANSDNKVTIDDYREEDEDKDPYPPCDVPVGEEEEEEEEDEPEVPAGPRPRRISELNMKEKIAPIPEGSAFFILSKTNPIRVGCHKLINHHIFTNLILVFIMLSSAALAAEDPIRSHSFRNTILGYFDYAFTAIFTVEILLKMTTFGAFLHKGAFCRNYFNLLDMLVVGVSLVSFGIQSSAISVVKILRVLRVLRPLRAINRAKGLKHVVQCVFVAIRTIGNIMIVTTLLQFMFACIGVQLFKGKFYRCTDEAKSNPEECRGLFILYKDGDVDSPVVRERIWQNSDFNFDNVLSAMMALFTVSTFEGWPALLYKAIDSNGENIGPIYNHRVEISIFFIIYIIIVAFFMMNIFVGFVIVTFQEQGEKEYKNCELDKNQRQCVEYALKARPLRRYIPKNPYQYKFWYVVNSSPFEYMMFVLIMLNTLCLAMQHYEQSKMFNDAMDILNMVFTGVFTVEMVLKVIAFKPKGYFSDAWNTFDSLIVIGSIIDVALSEADPTESENVPVPTATPGNSEESNRISITFFRLFRVMRLVKLLSRGEGIRTLLWTFIKSFQALPYVALLIAMLFFIYAVIGMQMFGKVAMRDNNQINRNNNFQTFPQAVLLLFRCATGEAWQEIMLACLPGKLCDPESDYNPGEEYTCGSNFAIVYFISFYMLCAFLIINLFVAVIMDNFDYLTRDWSILGPHHLDEFKRIWSEYDPEAKGRIKHLDVVTLLRRIQPPLGFGKLCPHRVACKRLVAMNMPLNSDGTVMFNATLFALVRTALKIKTEGNLEQANEELRAVIKKIWKKTSMKLLDQVVPPAGDDEVTVGKFYATFLIQDYFRKFKKRKEQGLVGKYPAKNTTIALQAGLRTLHDIGPEIRRAISCDLQDDEPEETKREEEDDVFKRNGALLGNHVNHVNSDRRDSLQQTNTTHRPLHVQRPSIPPASDTEKPLFPPAGNSVCHNHHNHNSIGKQVPTSTNANLNNANMSKAAHGKRPSIGNLEHVSENGHHSSHKHDREPQRRSSVKRTRYYETYIRSDSGDEQLPTICREDPEIHGYFRDPHCLGEQEYFSSEECYEDDSSPTWSRQNYGYYSRYPGRNIDSERPRGYHHPQGFLEDDDSPVCYDSRRSPRRRLLPPTPASHRRSSFNFECLRRQSSQEEVPSSPIFPHRTALPLHLMQQQIMAVAGLDSSKAQKYSPSHSTRSWATPPATPPYRDWTPCYTPLIQVEQSEALDQVNGSLPSLHRSSWYTDEPDISYRTFTPASLTVPSSFRNKNSDKQRSADSLVEAVLISEGLGRYARDPKFVSATKHEIADACDLTIDEMESAASTLLNGNVRPRANGDVGPLSHRQDYELQDFGPGYSDEEPDPGRDEEDLADEMICITTL

>XP_019392560.1|Crocodylus_porosus_Cav1.3

MLKNALGHRILSWYNEKVEYAFLIIFTIETFLKIIAYGLLLHPNAYVRNGWNLLDFVIVIVGLFSVILEQLTKEAEGGSHSGGKPGGFDVKALRAFRVLRPLRLVSGVPSLQVVLNSIIKAMVPLLHIALLVLFVIIIYAIIGLELFIGKMHKSCFLIGTDILVEEDPAPCAFSGNGRQCSTNGTECKGSWVGPNGGITNFDNFAFAMLTVFQCITMEGWTDVLYWMNDAMGFELPWVYFVSLVIFGSFFVLNLVLGVLSGEFSKEREKAKARGDFQKLREKQQLEEDLKGYLDWITQAEDIDPENEEGDEEGKRNTSMPTSETESVNTENVSGEGENPICCGKLCQTISKSKFSRRWRRWNRFNRRRCRAAVKSVSFYWLVIVLVFLNTLTISSEHYNQPNWLTQIQDIANKVLLALFTCEMLVKMYSLGLQAYFVSLFNRFDCFVVCGGIVETILVELEIMSPLGISVFRCVRLLRIFKVTRHWTSLSNLVASLLNSMKSIASLLLLLFLFIIIFSLLGMQLFGGKFNFDETQTKRSTFDNFPQALLTVFQILTGEDWNAVMYDGIMAYGGPSSSGMIVCIYFIILFICGNYILLNVFLAIAVDNLADAESLNTAQKEEAEEKERKKNARKESLENKKGDKTEGDQKKSKDSKVTIAEYGEGEDEDKDPYPPCDVPVGEEEEEEEEEPEVPAGPRPRRISELNMKEKITPIPEGSAFFIFSSTNPIRVGCHRLINHHIFTNLILVFIMLSSISLAAEDPIRSHSFRNNILGYFDYAFTAIFTVEILLKMTTFGAFLHKGSFCRNYFNLLDLLVVGVSLVSFGIQSSAISVVKILRVLRVLRPLRAINRAKGLKHVVQCVFVAIRTIGNIMIVTTLLQFMFACIGVQLFKGKFYRCSDEAKQNPEECRGIYIVYKDGDVDNPVVRERVWQNSDFNFDNVLSAMMALFTVSTFEGWPALLYKAIDSNGENVGPIYNYRVEISIFFIIYIIIIAFFMMNIFVGFVIVTFQEQGEQEYKNCELDKNQRQCVEYALKARPLRRYIPKNPYQYKFWYVVNSTGFEYIMFVLIMLNTLCLAMQHYEQSKLFNDAMDILNMVFTGVFTVEMVLKVIAFKPKHYFTDAWNTFDALIVVGSVVDIAITEVNNSEDSARISITFFRLFRVMRLVKLLSRGEGIRTLLWTFIKSFQALPYVALLIAMLFFIYAVIGMQVFGKVAMRDNNQINRNNNFQTFPQAVLLLFRCATGEAWQEIMLACLPGKRCDPESDYNPGEEYTCGSNFAIIYFISFYMLCAFLIINLFVAVIMDNFDYLTRDWSILGPHHLDEFKRIWSEYDPEAKGRIKHLDVVTLLRRIQPPLGFGKLCPHRVACKRLVAMNMPLNSDGTVMFNATLFALVRTALKIKTEGNLEQANEELRAVIKKIWKKTSMKLLDQVVPPAGDDEVTVGKFYATFLIQDYFRKFKKRKEQGLVGKYPAKNTTIALQAGLRTLHDIGPEIRRAISCDLQDDEPEENNQEEEEDVYKRNGALFGNHINHVSSDRRDSFQQINTTHRPLHVQRPSIPSASDAEKNMYAHAGNSVFHNHHNHNSVGKQVPNSTNANLNNANMSKVANGKHPNTGNHEHRSENRYHSYSRVDHDKHRRSSSKRTRYYETYIRSESGDGHLPTICREDHDVRDYCNDDHYFGDPEYFSGEEYYEEDSMLSGNRHTSDHNYRYHCHDSDFERPKGYHHPHGFFEEDDSQICYDAKRSPRRRLLPPTPTPNRRSSFNFECLRRQSSQDDITLSPTFHHRTALPLHLMQQQVMAVAGLDSSKAHKHSPSRSTRSWATPPATPPNRDRTPYYTPLIQVDRAESTEHMNGSLPSLHRSSWYTDDPDISYRTFTPVSLTIPNDFRHKQSDKQRSADSLVEAVLISEGLGRYAKDPKFVSATKHEIADACDMTIDEMESAASNLLNGNISNGTNGDMFPILSRQDYELQDFGPGYSDEEPDTGRYEEDLADEMICITTL

>XP_021335256.1|Danio_rerio_Cav1.3

MSANGPAPPAATPAAPPAAAVPVPSVVPVGSLAQKKRAQYAKSKKQGSSANTRPQRALFCLNLNNPIRRACISLVEWKPFDIFILIAIFANCMALAVYVPFPEDDSNSTNHDLETVEYAFLIIFTIETFLKIIAYGLVMHQNAYVRNGWNMLDFVIVVIGLFSVVLEVLTKEGGEKEEVGENLSAHGHGGKPGGFDVKALRAFRVLRPLRLVSGVPSLQVVLNSIIKAMVPLLHIALLVLFVIIIYAIIGLELFIGKMHASCYFQGTDILEDEPAPCAVNGHGRTCPINGTLCKEGWQGPNGGITNFDNFMFAMLTVFQCITMEGWTDVLYWMNDAMGLELPWVYFVSLVIFGSFFVLNLVLGVLSGEFSKEREKAKARGDFQKLREKQQLEEDLKGYLDWITQAEDIDPENEEEEEESKRNPSMPASETESMNTENEKGEDEKATCCGPTCQKISKSKFSRRWRRWNRLCRRNCRLAVKSVPFYWLVIILVFLNTLTISSEHYNQPMWLTQVQDVANKVLLAMFTCEMLVKMYSLGLQAYFVSLFNRFDCFVVCGGITETILVEFEIMSPLGISVFRCVRLLRIFKVTRHWASLSNLVASLLNSMKSIASLLLLLFLFIIIFSLLGMQVFGGKFNFDETQTKRSTFDNFPQALLTVFQILTGEDWNAVMYDGIMAYGGPSSSGMIVCIYFIILFICGNYILLNVFLAIAVDNLADAESLNTDDTKKPDEIDEIEDEAKAGEEDEKDNAEEDEEEPDVPAGPRPKISELVKKEKITPIPEGSAFFIFSNTNPVRVACHKLINHHIFTNLILVFIMLSSASLAAEDPIRNFSARNIILGYFDYAFTAIFTVEIVLKMTTYGAFLHKGAFCRNYFNLLDLLVVGVSLVSFGIQSSAISVVKILRVLRVLRPLRAINRAKGLKHVVQCVFVAIRTIGNIMIVTTLLQFMFACIGVQLFKGKFYRCNDEAKSSPEECKGTYIMYKEGDVNQPIIQKRHWHNSDFNFDNVLMAMMALFTVSTFEGWPALLYKAIDSNRENMGPIYNYRVEISIFFIIYIIIIAFFMMNIFVGFVIVTFQEQGEKEYKNCELDKNQRQCVEYALKARPLRRYIPKNPYQYKFWYVVNSTGFEYIMFVLILLNTICLAVQHYGQSELFNYVMDILNMVFTAVFTVEMVLKLIAFKPRHYFTDAWNTFDALIVVGSVVDIAITEVNPTEAPQVDESGNTEDSARISITFFRLFRVMRLVKLLSRGEGIRTLLWTFIKSFQALPYVALLIAMLFFIYAVIGMQVFGKIAMVDHTQINRNNNFQTFPQAVLLLFRCATGEAWQEIMLACMPGKLCDPESDYNPGEEMTCGSSFAIIYFITFYMLCAFLIINLFVAVIMDNFDYLTRDWSILGPHHLDEFKRIWSEYDPEAKGRIKHLDVVTLLRRIQPPLGFGKLCPHRVACKRLVAMNMPLNSDGTVMFNATLFALVRTALKIKTEGNLEQANEELRAVIKKIWKRTSMKLLDQVVPPAGDDEVTVGKFYATFLIQDYFRKFKKRKEEGLVGVHPAQNNTAIALQAGLRTLHDIGPEIRRAISCDLQDDELVDFIPEEDEEIYRRNGGLFGNHINHINGDPRRSSGHQTNATQRPLQVQPPPHYVHMEQPVGRLGRANAMAQQNHHRHHHHHHHHHHHNNSYNKSPKSTNINLNNANVSSLPNGGHNRYYEHAPANGYPGSYYGEYDKPRTPHGQRRRYYETYIRSQGSDRRRPTIRREEEYEEDRYSGEYYSGEEFYEDDSMLSGDRYPNSDQEYETPRGYHHPDSYYEDDEQPLYHDSHRSPKRRLLPPTPQGNRRPSFNFECLRRQSSQDDLPHQRTALPLHLMQHQVMAVAGLDSSRAHRLSPTRSTRSWASPPPTPASKDRTPYYTPLIRVDRPLRDSASSSHSSIRKSSWYTDDPEYQQRNFSPVHLQVPPEYRNQYLQKRGSATSLVEAVLISEGLGRYAKDPKFVAATKHEIADACEMTIDEMESAASHLLNGGITPVVNGVNVFPILGHRDYELQDVSASYSDEEPEPEPRPRYEEDLADEMICITTL

>XP_020648615.1|Pogona_vitticeps_Cav1.4

MSECNGIPTGPPEVPRSAAEAFVEATGQALAWTLTSGCTDTLGSTSSNAQKKRLQHNKHKTQQGANVVHRSPRALFCLRLNNPIRRAAISIVEWKPFDILILMTIFANCVALGVYIPFPEDDSNVSNHNLEQVEYVFLIIFTVETFLKILAYGLVMHPSAYIRNGWNLLDFVIVVVGLFSVILEQVSHKPGEAHHMSGKPGGFDVKALRAFRVLRPLRLVSGVPSLHIVLNSIMKAMVPLLHIALLVLFVIIIYAIIGLELFIGRMHKTCFIIGSDLEAEEDPSPCAFSGHGRECTVNNSECRGKWEGPNGGITNFDNFFFAMLTVFQCITMEGWTDVLYWMQDAMGHELPWLYFVSLVIFGSFFVLNLVLGVLSGEFSKEREKAKARGDFQKLREKQQMEEDLKGYLDWIMQAEDIEPDEEGDETNEKHARVTVEDLTSKRKRKWFRQGSSHSTDTHTSLQASETTSVNTENVGDEEHHQDNCCDVCLGKLSKTKFCRRIRRVNRLFRKKCRLAVKSVTFYWIVLILVFLNTLTIASEHYMQPDWLTQIQAYANKVLLSLFTLEMLVKMYSLGLQAYFVSFFNRFDCFVVCGGILETVLVEFEIMEPLGISVLRCVRLLRIFKVTRHWASLSNLVASLLNSMKSIASLLLLLFLFIIIFSLLGMQLFGGKFNFDETQTKRSTFDTFPQALLTVFQILTGEDWNAVMYDGIMAYGGPVFPGMLVCVYFIILFICGNYILLNVFLAIAVDNLADGDNINTSKEEEQKEKKRKKKKVVKNKWPSRKSQSKSSFYSPAQTGEEDEEGDQKSLAESHLDKVDEAPKQKVVPIPDGSAFFCLSKTNPLRVGCHKLIHHHIFTNLILVFIILSSISLAAEDPIRAHSFRNIILGYFDYAFTSIFTVEILLKMTAYGAFLHQGSFCRNWFNLLDLLVVSVSLISFGIHSSAISVVKILRVLRVLRPLRAINRAKGLKHVVQCVFVAIRTIGNIMIVTTLLQFMFACIGVQLFKGKFYSCTDEARHTPKECKGTFIVYKDGDVAHPMVRDRLWLNSDFNFDNVLSGMMALFTVSTFEGWPALLYKAIDANAENHGPIYNYRVEISIFFIVYIIIIAFFMMNIFVGFVIITFRAQGEQEYKNCELDKNQRQCVEYALKAQPLRRYIPKNKYQYKFWYIVNSTGFEYIMFVLILLNTIALAVQHYEQSQPFNYVMDLLNMVFTGLFTIEMVLKIIAFKPKHYFVDAWNTFDALIVVGSVVDIAVTEVNSSEDSSRISITFFRLFRVMRLVKLLSKGEGIRTLLWTFIKSFQALPYVALLIAMIFFIYAVIGMQTFGKVAMQDGTPINRNNNFQTFPQAVLLLFRCATGEAWQEIMLASLPGKRCDPESDYEPGEEFTCGSNFAIVYFISFFMLCAFLIINLFVAVIMDNFDYLTRDWSILGPHHLDEFKRIWSEYDPAAKGRIKHLDVVTLLRRIQPPLGFGKLCPHRVACKRLVAMNMPLNSDGTVTFNATLFALVRTSLKIKTEGDLDVANKELRAVIKKIWKRTKPKLLDEVIPPPEEEEVTVGKFYATFLIQDYFRKFRKRKEKGMLGPDGSPSNSTALQAGLKSLQDLGPEIRRAMSCDLEEEEEEEEENGETLCEEETVTYKSTEALYGSAPPDHHVSTAGASPDQDGKAVANGLVSSRQPSSASLSHMTNGPTPYEENGGDQDEPAELPNIATRRRMSRRSSDGSCIPPTVKEEATHPDDYGEDTSVEQGYYSREEDSESLASRDRQPSLHHPQPRRRRWDTGYGGSLPLPSRRVNNGLINGPLEGRQTKRRRLLPPTPTGRKPAFSIQCLQRQGSCDDIPIPGTYHQNAPPCRARGPGYGSYDSWHSGSRSSTASSHSWATPPKRGRLLYAPLILVEEAGLPGHSWEKNSSSLPPMSRARWYLNEPEVPYHTYGHLHVPGPLKHSYNDKRGSADSLVEAVLISEGLGLYARDPKFVAFTKREIADACHMTIDEMESAATDLLSRRFVGSEHGGQGGSGGGSHVLYSSDGEHSGVGSLGPVYSDEEPLRTREEEDLADEMACVTSY

>XP_016846283.1|Anolis_carolinensis_Cav1.4

MDAESEDTEDRLEKGQKEWKVLLYEVEKHLDQSASIFWAVTHPPIQPFQPATSSNKPAKCQSRSATEDNLSLLSFLPTVTEPPELPRSAAEAFVEATGQALAWTLTNGCTDTLGSTGSNAQKKKLQHNKHKTQGTNVVHRSPRALFCLRLNNPIRRAAISIVEWKPFDILILMTIFANCVALGVYIPFPEDDSNVANHNLEQVEYIFLIIFTVETFLKILAYGLVMHPSAYIRNGWNLLDFVIVVVGLFSVILEQFSHKPGEAHHMSGKPGGFDVKALRAFRVLRPLRLVSGVPSLHIVLNSIMKAMVPLLHIALLVLFVIIIYAIIGLELFIGRMHKTCFIIGSDLEAEEDPSPCAFSGHGRECTVNNSECRGKWEGPNGGITNFDNFFFAMLTVFQCITMEGWTDVLYWMQDAMGHELPWLYFVSLVIFGSFFVLNLVLGVLSGEFSKEREKAKARGDFQKLREKQQMEEDLQGYLDWIMQAEDIEPDDEGDEADEKHTRVTVEDLTGKKKKKWFRHSSSTDTHSNFLIFLHMSPYYVLFMVFYKLSQSHPRRSRKFLERASFMRCVKVELQLLVLLVQGHIVLLSLSSVQPSLLQGKLAKTKFGRRMRRINRLLRKRCRLAVKSVSFYWMVLILVFLNTLTIASEHYNQPDWLTQIQAYANKVLLSLFTLEMLLKMYSLGLQAYFVSFFNRFDCFVVCGGILETVLVEFEIMQPLGISVLRCVRLLRIFKVTRHWASLSNLVASLLNSMKSIASLLLLLFLFIIIFSLLGMQLFGGKFNFDETQTKRSTFDTFPQALLTVFQILTGEDWNTVMYDGIMAYGGPYFPGMLVCVYFIILFICGNYILLNVFLAIAVDNLADGDNINTSKNKEAPAEGEQSNETEEKDVKLECEEEEEEEEAEEGSEEAGEEEEEEGDQQSLAESHLDKVEDTPTEKVMPIPDGSSFFCLSKTNPLRVGCHKLIHHHIFTNLILVFIILSSISLAAEDPIRAHSFRNNMTAFGGFLHQGSFCRNWFNLLDLLVVSVSLISFGIHSSAISVVKILRVLRVLRPLRAINRAKGLKHVVQCVFVAIRTIGNIMIVTTLLQFMFACIGVQLFKGKFYSCTDEAKHTPNECKGTFIVYKDGDVAHPMVRDRLWLNSDFNFDNVLSGMMALFTVSTFEGWPALLYKAIDANAENEGPIYNYRVEISIFFIIYIIIIAFFMMNIFVGFVIITFRAQGEQEYKNCELDKNQRQCVEYALKAQPLRRYIPKNKYQYKFWYMVNSTGFEYIMFVLILLNTIALAVQHYEQSQPFNYVMDLLNMVFTGLFTVEMVLKIIAFKPKHYFCDAWNTFDALIVVGSVVDIAVTEVNSSEDSSRISITFFRLFRVMRLVKLLSKGEGIRTLLWTFIKSFQALPYVALLIAMIFFIYAVIGMQTFGKVAMQDGTPINRNNNFQTFPQAVLLLFRCATGEAWQEIMLASLPGKRCDPESDYEPGEEFTCGSNFAIVYFISFFMLCAFLIINLFVAVIMDNFDYLTRDWSILGPHHLDEFKRIWSEYDPAAKGRIKHLDVVALLRRIQPPLGFGKLCPHRVACKRLVAMNMPLNADGTVTFNATLFALVRTSLKIKTEGDLDVANKELRAVIKKIWKRTKPKILDEVIPPPEEEEVTVGKFYATFLIQDYFRKFRKRKEKGMLGADGSPSNSTALQAGLKSLQDLGPEIRRAMSCDLEEEEEEEENGETVCEEETVTYKSTEALYGSAPDQSAAPASPEQEGKTLMNGLVGSRHPSSTSLSHVANGPTHYEDNVGSHDEPTELPSMPTRRRLSRRSSDGSCIPPTVKEETNHNEDYGEDTSVEQGYYSREEDSESLASRERQPSLHNPRPHRHHWDNGYGGSLPLPSRRMNNGLVNGMLDSRQPKRRRLLPPTPTGRKPTFNIQCLQRQGSCDDIPIPGTYHQNAPPCRARAQGYGSYDSWQSGGRNSTASSQSWATPPKRSRLLYAPLILVEGDELPGHSWEKNSSLPPMSRARWYMNEPEMPYRTYSHLHVPGPLKHSYDKRGSADSLVEAVLISEGLGLYARDPKFVAFTKREIADACHMTIDEMESAATDLLSRKYVGSENGSRSSSGGGHALYSSDGEHSGVGSVGPVYSDEEPFRSPREEEDLADEMACVTSY

>XP_005533405.1|Pseudopodoces_humilis_Cav1.4

MTGVVGGKTWCPPLPPRPFDILILATIFANCVALGVYIPFPEDDSNTSNHNLEQVEYVFLIIFTVETFLKIIAYGLVLHPSAYIRNGWNLLDFVIVIVGLFSVILEQVSHKPGDAHHMSGKPGGFDVKALRAFRVLRPLRLVSGVPSLHIVLNSIMKAMVPLLHIALLVLFVIIIYAIIGLELFIGRMHKTCFFIGSDLESEDDPSPCAFSGHGRACLQNNTECRGRWEGPNGGITNFDNFFFAMLTVFQCITMEGWTDVLYWMQDAMGHELPWIYFVSLVIFGSFFVLNLVLGVLSGEFSKEREKAKARGDFQKLREKQQLEEDLRGYMDWITQAELEGDEEEGEHEKHRLTAEDLMGKRKPRLKWLRHASHSTDTHASLPGSETTSVNTETVGEEEAPPTACDRCLGKITKTKFCRRLRRLNRLWRRRCRAAVKSVSFYWTVLLLVFLNTLTIASEHHGQPPWLTETQAYANKALLSLFAAEMVLKLYALGPSCYFASFFNRFDCFVVCGGVLETALVERGAMEPLGISVLRCVRLLRVFKVTRHWASLSNLVGSLLNSMKSIASLLLLLFLFIIIFALLGMQLFGGRFSFDETQTKRSTFDTFPQALLTVFQILTGEDWNAVMYDGIMAYGGPVFPGMLVCVYFVILFICGNYILLNVFLAIAVDNLADGDNINSGGDKKXAQSPGALSGVMRGEPQPPGPQGGVEGDQPQEEEEEEEEEVEEGEARLESLEEAPKPKVTPIPEGSAFFVLSSTNPLRVKCHALINHHIFTNLILVFIILSSISLAAEDPVRAHSPRNHILGYFDYAFTSIFTVEILLKMTVFGGFLHKGSFLRNWFNLLDVLVVGVSLISFGMHSSAISVVKILRVLRVLRPLRAINRAKGLKHVVQCVFVAIRTIGNIMIVTTLLQFMFACIGVQLFKGKFYSCTDEAKHTPGECKGTFLVYKDGDVSHPSVRERLWLNSDFNFDNVLAGMMALFTVSTFEGWPALLYKAIDANAENQGPIYNYRVEISIFFIVYIIVIAFFMMNIFVGFVIITFRAQGESEYRNCELDKNQRQCVEYALKAQPLRRYIPKNRTQYRVWAMVNSTAFEYIMFVLILLNTIALAVQHYEQSKPFNYVMDLLNMVFTGLFTVEMVLKIIAFKPRHYFCDAWNTFDALIVVGSVVDIAVTEVNNGGHVGESSEDSSRISITFFRLFRVMRLVKLLSKGEGIRTLLWTFVKSFQALPYVALLIAMIFFIYAVIGMQTFGKVALQDGTQINRNNNFQTFPQAVLLLFRCATGEAWQEIMLASLPGKRCDPESDVGPGEEFTCGSNFAIAYFISFFMLCAFLIINLFVAVIMDNFDYLTRDWSILGPHHLDEFKRVWSEYDPAAKGRIKHLDVVTLLRRIQPPLGFGKLCPHRVACKRLVAMNMPLNSDGTVTFNAPLFALVRTSLKIKTEGNLDVANKELRAVVKKIWKRTKPKLLDEVIPPPEEEEVTVGKFYATFLIQDYFRKFRRRKERGMLGASAGPSNECALQAGLQTLQALGPEMRRALSDLEGDEGGDAPAEEPLTYAAPETLYGSAPGSPLLSEPPPVPSPAPTEGEDPAPPAHGGAPRPGGRRRSEPGDQDEEAPAEDVEEPGADPGDVSHDEDLESSAPPRHRWAGHRAPGPPRWAAELPRGGSLPLPPRHFALGSGAREGQQLKRRRLLPPTPAGRKPSFTIQCLRRQGSCEDEPIPGTYNPSGPPGSARPQAWAAPARGHVLYAPLILVEGAPAAGGSLPPLNRWFPPAPLRLRPCGPHDALARGSADSLVEAVLISEGLGLFARDPKFVAVAKREIADACDMTMDEMESAAADLLTRRRAPPAPAVYSDEEPLRPPAEEELADEMGWAGGV

>NP_005174.2|Homo_sapiens_Cav1.4

MSESEGGKDTTPEPSPANGAGPGPEWGLCPGPPAVEGESSGASGLGTPKRRNQHSKHKTVAVASAQRSPRALFCLTLANPLRRSCISIVEWKPFDILILLTIFANCVALGVYIPFPEDDSNTANHNLEQVEYVFLVIFTVETVLKIVAYGLVLHPSAYIRNGWNLLDFIIVVVGLFSVLLEQGPGRPGDAPHTGGKPGGFDVKALRAFRVLRPLRLVSGVPSLHIVLNSIMKALVPLLHIALLVLFVIIIYAIIGLELFLGRMHKTCYFLGSDMEAEEDPSPCASSGSGRACTLNQTECRGRWPGPNGGITNFDNFFFAMLTVFQCVTMEGWTDVLYWMQDAMGYELPWVYFVSLVIFGSFFVLNLVLGVLSGEFSKEREKAKARGDFQKQREKQQMEEDLRGYLDWITQAEELDMEDPSADDNLGSMAEEGRAGHRPQLAELTNRRRGRLRWFSHSTRSTHSTSSHASLPASDTGSMTETQGDEDEEEGALASCTRCLNKIMKTRVCRRLRRANRVLRARCRRAVKSNACYWAVLLLVFLNTLTIASEHHGQPVWLTQIQEYANKVLLCLFTVEMLLKLYGLGPSAYVSSFFNRFDCFVVCGGILETTLVEVGAMQPLGISVLRCVRLLRIFKVTRHWASLSNLVASLLNSMKSIASLLLLLFLFIIIFSLLGMQLFGGKFNFDQTHTKRSTFDTFPQALLTVFQILTGEDWNVVMYDGIMAYGGPFFPGMLVCIYFIILFICGNYILLNVFLAIAVDNLASGDAGTAKDKGGEKSNEKDLPQENEGLVPGVEKEEEEGARREGADMEEEEEEEEEEEEEEEEEGAGGVELLQEVVPKEKVVPIPEGSAFFCLSQTNPLRKGCHTLIHHHVFTNLILVFIILSSVSLAAEDPIRAHSFRNHILGYFDYAFTSIFTVEILLKMTVFGAFLHRGSFCRSWFNMLDLLVVSVSLISFGIHSSAISVVKILRVLRVLRPLRAINRAKGLKHVVQCVFVAIRTIGNIMIVTTLLQFMFACIGVQLFKGKFYTCTDEAKHTPQECKGSFLVYPDGDVSRPLVRERLWVNSDFNFDNVLSAMMALFTVSTFEGWPALLYKAIDAYAEDHGPIYNYRVEISVFFIVYIIIIAFFMMNIFVGFVIITFRAQGEQEYQNCELDKNQRQCVEYALKAQPLRRYIPKNPHQYRVWATVNSAAFEYLMFLLILLNTVALAMQHYEQTAPFNYAMDILNMVFTGLFTIEMVLKIIAFKPKHYFTDAWNTFDALIVVGSIVDIAVTEVNNGGHLGESSEDSSRISITFFRLFRVMRLVKLLSKGEGIRTLLWTFIKSFQALPYVALLIAMIFFIYAVIGMQMFGKVALQDGTQINRNNNFQTFPQAVLLLFRCATGEAWQEIMLASLPGNRCDPESDFGPGEEFTCGSNFAIAYFISFFMLCAFLIINLFVAVIMDNFDYLTRDWSILGPHHLDEFKRIWSEYDPGAKGRIKHLDVVALLRRIQPPLGFGKLCPHRVACKRLVAMNMPLNSDGTVTFNATLFALVRTSLKIKTEGNLEQANQELRIVIKKIWKRMKQKLLDEVIPPPDEEEVTVGKFYATFLIQDYFRKFRRRKEKGLLGNDAAPSTSSALQAGLRSLQDLGPEMRQALTCDTEEEEEEGQEGVEEEDEKDLETNKATMVSQPSARRGSGISVSLPVGDRLPDSLSFGPSDDDRGTPTSSQPSVPQAGSNTHRRGSGALIFTIPEEGNSQPKGTKGQNKQDEDEEVPDRLSYLDEQAGTPPCSVLLPPHRAQRYMDGHLVPRRRLLPPTPAGRKPSFTIQCLQRQGSCEDLPIPGTYHRGRNSGPNRAQGSWATPPQRGRLLYAPLLLVEEGAAGEGYLGRSSGPLRTFTCLHVPGTHSDPSHGKRGSADSLVEAVLISEGLGLFARDPRFVALAKQEIADACRLTLDEMDNAASDLLAQGTSSLYSDEESILSRFDEEDLGDEMACVHAL

>XP_018667169.1|Ciona_intestinalis_Cav1

MHGPRKRKQAQQDTAKAETSLLCLSLKNPFRKACLKIVEWRPFDVLILLTIFANCCALAIYVPFPGEDSNATNEILEKVEYVFLAIFTVESFMKIIAFGFAFHPNAYLRNGWNILDFIIVIVGLISIVFEMADVGSTDKVRALRAFRVLRPLRLVSGVPSLQVVLNAIIRAMLPLLHIALLVMFVIIIYAVVGLELFKGKLHKTCYFNETGMTDVIANEDPQPCAGPNEWGRHCPDDTVCKEGWDGPANGIINFDTFYFSFITVFQCITMEGWTEVLYYTNDAMGSHLPWMYFVSLIIVGSFFVMNLILGVLSGEFSKEREKANARGEFQKLREKQQLDEDVRGYMEWITQAEDIDPVNEDDDMDEKRQGDNEDGSSDVTAAQADDSWWQKQRKKLCKTCYSRRWKRWNRKTRRKCRLMVKSQTFYWLVIVLVFFNTLSLATEHYQQPDWLTSVQEISNKVLLGIFTLEMLLKMYALGMQVYFVSLFNRFDCFVVCGGIVEMVLTSAKVMEPLGISVLRCVRLLRIFKVTRYWSSLSNLVASLLNSIRSIAGLLLLLFLFIVIFSLLGMQLFGGRFNSIAEGDQKIRSNFDTFLQALLTVFQILTGEDWNVVMYYGIRAYGGASSIGLITSIYFIILFVCGNYILLNVFLAIAVDNLADAESLNVAQKEKEEEQKRKKTMRLKKLRNLFKKKETTSVETAEGADEYGDRQKYDKNEDGIPLQNIAESSLQTDEIDHEIRIEVTEASETNSDRHLPEDGGSDSEPEVPIGPRPRRMSELNLKETKSPMPQATSFFIFTPTNPFRKWCHFIANNNIFNNGIFVCIMLSSVALACEDPIDSKSELNEVLKYFDYVFTGIFTVEIILKMVAYGVILHKGSFCRNSFNLLDLLVVGVSLISIFGNSDGFSVVKILRVLRVLRPLRAINRAKGLKHVVQCVIVAISTIGNIFIITTLLQFMFACIGVQLFKGRLYGCTDESKSTREECKGDFYAIPQDGFGQPHIKKREWVNNDFNYDNVLNAMLTLFVVATFEGWPALLYKSIDSWKEGVGPKYDARPAVALFYFIYIIVIAFFMMNIFVGFVIVTFQEQGEQEYRNCELDKNQRQCVEYALKAKPTRRYIPKNPWQYKAWFVVNSTYFEYFMLVLILLNTVCLAIQHYQQDAGLTRILNHMNLVFTTLFTIEMIFKLIAFKPRGYISDPWNIFDFLVVIGSIVDILLSKIDTGGDKSFSINFFRLFRVMRLVKLLSRGEGIRTLLWTFIKSFQALPYVALLIVLLFFIYAVIGMQVFGKVKPIDGEQINRNNNFQTFIQSVLLLFRCATGESWQEVMLAAASGKECDDRSDWNSTGLASPEDKFACGSDFSYTYFLTFYMLCAFLIINLFVAVIMDNFDYLTRDWSILGPHHLDEFKTVWSEYDPEAKGRIKHLNVVKLLRRIQPPLGFGKLCPQRMACRKLVTMNMPLNSDGTVMFNATLFALIRTSLNIKTEGNIDQANEELRAVIKKIWKRTSIKLLDQIAPPAGNDDITVGKFYATYLIQDYFRKFRERKAAAKLANDKSKGLSGANMQNHKDVFSLQAGLRALQDAGPEIKRAISGGIAHDEPYASNDENEPEHRRRHSLFGMLRRNSATTPTASKRPLQVESDDHKKRYSGRFLTSQSPMLNRLTPKSRSIHDIVMAARAQHSGQSGDETTSSIASPTDESLGRNDYHSRSAVTHGENSNVNNSSSNYSFSSYATDDELNSVGGYYDEHAPLHPAARSERKYNDHQRYPSYAKRWDRSVYRCDDDSRTVLYPLLPVHSRDEESEYETDHTTYAAAPYRCNDETCSVDFEKRSNYQAQNGAVSNRTNRSLPNVPLRTSIDRPVALNPHDHHHHRESVNNHTNSNDSGPMHSRYTITTPKLRIMDSDYVPTIDRFVPRSRQSKSKPHRRLLPIPPLQHARSGTIPSLHLSNQEAWRTPPSSPRQALSVRSNLNSPIGSSDEGGWATPAQRWTSESNMDTFGRGNHINYHDSRRQSGNDLIEKVLLDGGLPVLAQDKNFVQTISKELADACELSLPELHSAATNIINSDHSLYGSDESLCGDVSPDNHIDDDVTSLEEGRVCFRRRSKTRKDEHGLEDEIIVVPNITS

>BAA34927.2|Halocynthia_roretzi_Cav1

MNGTTNPTTRKRKIKPDPNAGRAPQALLCLSLKNPIRKACMKIVDWRPFDVLILLTILANCVALAVYVPFPGDDSNRTNEILEKVEYIFLGIFTIEAILKIIAYGLFFHPNAYLRNGWNVIDFVIVVIGLVSIVLETANVGSTDKVRSLRAFRVLRPLRLVSGVPSLEVVLNAIIRAMVPLLHIALLVIFVIIIYAVVGLELFKGKLHKTCYHNEVAVLIMEDEAKPCADSDSWGRHCSGGMICESDWAGPSKGIINFDTFYFAVITVFQCITMEGWTDVLYYMNDAVGNLWPWIYFVSLIIIGSFFVMNLILGVLSGEFSKEREKANARGEFQKLREKQQTDEDMKGYMDWITQAEDLDPMNDEDREDRRSASNEQLNDADSEVSGLQIDETWWQMQRRALFKVCYSRRWRRWNRKTRRRCRTMVKSKSFYWLVIVLVFCNTLSLATEHYRQPPWLTLAQDLANKILLTLFTIEMLVKMYSLGMQQYFVSLFNRFDCFVVCGGIVELVLTSSKIMEPLGISVLRCVRLLRIFKMTSSWNSLSNLVASLLNSIRSIASLLVLLFLFIIIFALLGMQMFGGRFSEIEQEDKIRSNFDTFLQALLTVFQILTGEDWNVVMYNGIEAYGGASTIGLLTSVYFIVLFIGGNYILLNVFLAIAVDNLADAESLGAAQKEKEEEKKMKKTLRLKKLRKLFKKKDQTSVENQEVNQEDTLNRVEEMPNYYTPDHDIRIEVTEASDTNSDKHLPEVSDGEMEPEVPVGPRPRRMSEMHLSEKKVPLPEGSSFFILSNTNRLRVFCYDIVNYNWFNNAILACIILSSIALACEDPVSAHSARNKVLEYFDYVFTGVFAVEIVLKMTAFGVFLHKGSFCRSYFNLLDLLVVAVSLVSMLSNSDKFSVVKILRVLPSVLRPLRAINRAKGLKHVVQCVFVAISTIGNIMVITGLLQFMFACIGVQLFKGRLYYCTDQSKETKEECHGKFFVYSKDGNGEPRVEERLWENSEFNYDNVMNAMLTLFVVATFEGWPGLLYKSIDSWSENHGPRYDARQAVALFYFVFIIVIAFFMMNIFVGFVIVTFQEQGEQEYKNCELDKNQRQCLEYALKAKPVKRYIPKNPWQYKVWFIVNSTYFEYFMLVLILLNTVCLAVQHHQQSKELTVILNHMNYVFTALFALEMIVKLVAYKPRGYLSDPWNVFDSLIVIGSIVDIVFSELDHGNEKSFSINFFRLFRVLRLVKLLSRGEGIRTLLWTFIKSFPALPYVALLIIMLFFIYAVIGMQIFGKIKPNDGSQINRNNNFQTFLQAVLLLFRCATGESWQEVMLACASGNECDDESDWNYYGDKDISAKFTCGNDFAYTYFLTFYMLCAFLIINLFVAVIMDNFDYLTRDWSILGLHHLDEFVRVWSEYDHEASGRIKHLHVVKLLRHIQPPLGFGKLCPQRMACRKLVSMNMPLNSDGTVMFNATLFALVRTSLKIKSEGNIDQANEELRAVIKKIWKRTSMKLLDQVVPPAGNDDVTVGKFYATYLIQDYFRKFKESKERRLRERDGRHHPNNANTLTLQAGIREVQDVSPELKRSISGNLVPNEYEESELVSMDDEPEHRRRHSIFGQLRSLASPGTPVMTSRRPLTVSDNTPAVGHRKHEQNSDQNPLVNRLTPQQPFPNATPSHAQATEKSVNRSLDGIHTMPRDANIGRHHGSIDDFSRYHKSPTQPDGFVVEPTFTSPSHNTNRNNSANPTECMVHRPNGKCQKMQRAYSEEDKGERGDRNFIVPYLSMDGGMNTVQHWGRDAREQRPCYCGKPRRLLRPCSKRSLNLDYTPLQRNETPVKSPTTENLSKRTLQSCDRSFSFPGKNRIDSFTSIEDECLLSRDSRDADSIFFNEIGGSGISSSDSDMFESDSGSCDDECGSVRTYRTANTISPQCAFKTFSEVNSQPGSKSDVGVTRKLPPTPEAPSRIFRLPMMSKNWRKSRENMPLCDIKTSETAPLINSRNETSFHRNSIPRHKLTRQFSTNYDNRHHADDMDYRRDNEVIIYNKGDDGRGCSKTRSFVQIFQSLTNGDSDNDSTNVCQPKSYCRSDSVWDEEEETPIKRSQVEISPSGSRTRSPVPRNGHEGCIERDEEADLASEMIVPNVK

>XP_016766333.1|Apis_melifera_Cav1

MSAGGDGGSGLGPPELAGAQPTAATPPILGQHRNPETGQDGQQAATAQSQTGQTLGTSTGAAAAAAAAAAAAKSATKRPARRGGKAPPDRPNRTLFCLPLKNPLRKMCIDVVEWKPFEWLILMTIFANCIALAVYTPYPYGDSNLTNQYLEKIEYIFLVIFTVECVMKIIAYGFVAHPGAYLRNGWNILDFSIVVIGMVSTVLSVLMKEGFDVKALRAFRVLRPLRLVSGVPSLQVVLNSILRAMIPLLHIALLVLFVIIIYAIIGLELFSGKMHKTCRHNMTDAIMDDPVPCGPGGYQCDNVGSDYYCSKQFWEGPNWGITNFDNFGLAMLTVFQCVTLEGWTEVLYNIEDAMGSSWQWIYFISMVILGAFFVMNLILGVLSGEFSKEREKAKARGDFHKLREKQQIEDDLRGYLDWITQAEDIEPETDEPKMQDGKTKQQSEMESTDQLEGDEEGVQQESLWRRKKLDFDRVNRRMRRACRKAVKSQVFYWLIIVLVFLNTGVLATEHYNQPHWLDDFQEITNMFFIALFTMEMMLKMYSLGFQGYFVSLFNRFDCFVVIGSITEMILTNTHVMPPLGVSVLRCVRLLRVFKVTKYWRSLSNLVASLLNSIQSIASLLLLLFLFIVIFALLGMQVFGGKFNFNVLENKPRHNFDSFWQSLLTVFQILTGEDWNAVMYDGIRAYGGVSSFGMLACFYFIILFICGNYILLNVFLAIAVDNLADAESLTAIEKEAEEEAEKNKSHSASPTRDKDSGEQGDDGGEGTGGEDEGGGTDLEHDPNETMEDYEAAVDTETSEKSDDMNTHAKVRLNIESDEEVEEEEEVEHNEMHGERFIYDGTEQGVSARPRRMSEFNMATKKQPIPAGSAFFIFSQTNRIRIFCHWLCNHSTFGNVILVCIMISSAMLAAEDPLRASSSRNLVLQKFDYFFTTVFTIEICLKMISYGFIIHEGAFCRSAFNLLDLLVVCSSLISMSFSSGAFSVVKVLRVLRVLRPLRAINRAKGLKHVVQCVIVAVKTIGNIVLVTSLLQFVFAVVGVQLFKGKFFYCTDASKMTKEECQGTYLEFENGNINKPIMKERNWCQQRFHFDDVAKAMLTLFTVSTFEGWPSLLDYSIDSNKEDHGPIHNFRPIVAAYYIIYIIIIAFFMVNIFVGFVIVTFQNEGEQEYKNCELDKNQRNCIEFALKAKPVRRYIPKHRIQYKVWWFVTSQPFEYTIFTLIMINTVTLAMKFYRQPEIYTQALDVLNMIFTAVFALEFIFKLAAFRFKNYFGDAWNVFDFIIVLGSFIDIVYSEVNPGSTIISINFFRLFRVMRLVKLLSRGEGIRTLLWTFIKSFQALPYVALLIIMLFFIYAVIGMQVFGKIAIDDETSINRNNNFQSFPQAVLVLFRSATGESWQEIMMDCSVQPGKVKCDPNSDEALNTNGCGSDIAFPYFISFYVLCSFLIINLFVAVIMDNFDYLTRDWSILGPHHLDEFIRLWSEYDPDAKGRIKHLDVVTLLRKISPPLGFGKLCPHRVACKRLVSMNMPLNSDGTVLFNATLFAVVRTSLRIKTEGNIDDANAELRAVIKKIWKRTSPKLLDQVVPPPGGDDEVTVGKFYATFLIQDYFRRFKKRKEQEMKDGDKECHNTVTLQAGLRTLHEAGPELKRAISGNLEELLDDNPEPMHRRNHSLFGSVWSSMRKGHHSFNRARSLKVNSTSKASPTNSIDFVPYSSFHRGGGDPSNQITARSHQVVPNVAGGLSDSAMNQMGIDPKLTGIEESIPLRPLAVFGNPVQQQSYHHTSYKVLDGPGSGNYLHPNNEYVSWAGESNGSIGAERLSHSLPGSPADRKPNFEVIGSAESLVGRVLVEQGLGKYCDPDFVRYTSREMQEALDMTREEMDRAAHQLLLQERRGQPLSYQLQQGVDQQWTSSYQPSQSTGIGYQPLQEQQSSGQPRQYRSYYRGGGQATTTSDPSSIQQQQQQQQQQQQQQQQQQQQQQSPPS

>XP_021699870.1|Aedes_aegypti_Cav1

MDADPALRSSGDHKPPSASVSSSKHATEADKRIPEVKTDLIVGPKADSKSRASDTKKRSKNKSFIVGLLRKPSEPEATAKKKNNKNKKRSSRFFSRRRHSLSDVWQATLKSTTAMSQAAAPINGQNAAGIGLDPGTGPPKGPGETGIANTTIPVAPKKPVRRAGVKPQPDRPVRALFCLTLKNPLRKLCIDIVEWKPFEYLILLTIFANCVALAVYTPFPNSDSNTTNAALEKIEYIFLVIFTAECVMKLIAYGFIMHPGSYLRNGWNILDFTIVVIGMISTALSNLMKEGFDVKALRAFRVLRPLRLVSGVPSLQVVLNSILRAMVPLLHIALLVLFVIIIYAIIGLELFSGKLHKTCFHNVTGEKEVFSQEYVLILFLFLDEIMDDPHPCGDDGFQCASISEDMVCRYYWAGPNFGITNFDNFGLSMLTVFQCVTLEGWTDMLYYIQDAMGSTWQWVYFISMVILGAFFVMNLILGVLSGEFSKERTKAKNRGDFQKLREKQQIEEDLRGYLDWITQAEDIDPENEANVVQEGKTMTANEIDSSDHMGEEGEIQQESWLARKRKNIDRVNRRLRRACRKAVKSQAFYWLIIVLVFLNTGVLATEHYQQPPWLDDFQEYTNMFFVALFTMEMLLKMYSLGFQGYFVSLFNRFDCFVVIGSIGEMVLTSTQIMPPLGVSVLRCVRLLRVFKVTKYWQSLSNLVASLLNSIQSIASLLLLLFLFIVIFALLGMQVFGGKFNFNSETDKPRSNFDSFVQSLLTVFQILTGEDWNAVMYDGIQAYGGVASIGILASIYFIILFICGNYILLNVFLAIAVDNLADADSLTTVEKEEGEGEEGADVEKSGSHEGTPLGITDDGFMEHDKNNLDSENEMNISEDYEHNGSETKMTLPDDDEGYEDQDQDYQDKEYNLARPRRMSELNVANKIIPIPPGSSFFIMSQTNRFRVFCHWLCNHSTFGNIILVCIMFSSAMLAAEDPLNANSERNQILNYFDYFFTTVFTIELLLKVISYGFLFHDGAFCRSAFNLLDLLVVCVSLISMFFSSGAISVIKILRVLRVLRPLRAINRAKGLKHVVQCVIVAVKTIGNIVLVTCLLQFMFAVIGVQLYKGKFFSCSDGSKMQESECHGTYLVFEGGNVDKPVSKEREWSRNRFHFDDVSKAMLTLFTVSTFEGWPGLLYVSIDSHEEDSGPIHNFRPIVAAYYIIYIIIIAFFMVNIFVGFVIVTFQNEGEQEYKNCDLDKNQRNCIEFALKAKPIRRYIPKHRIQYKVWWFVTSQPFEYMIFILIMINTITLSMKFYRQPEIYTEVLDLLNLIFTAVFALEFVFKLAAFRFKNYFGDAWNVFDFIIVLGSFIDIVYSEVNVSKGMKGGSSIISINFFRLFRVMRLVKLLARGEGIRTLLWTFIKSFQALPYVALLIVMLFFIYAVIGMQVFGKIAMDDETSIHRNNNFQTFPQAVLVLFRSATGEAWQDIMLDCSSRPGEVMCDPRSDDANSPEGCGSSIAFPYFISFYVLCSFLIINLFVAVIMDNFDYLTRDWSILGPHHLDEFVRLWSEYDPDAKGRIKHLDVVTLLRKISPPLGFGKLCPHRVACKRLVSMNMPLNSDGTVLFNATLFAVVRTSLKIKTEGNIDDANAELRATIKQIWKRTAPKLLDQVVPPPGVDDEVTVGKFYATFLIQDYFRRFKKRKENDNKHVDYDNRRTMTLQAGLRTLHEAGPELKRAISGNLEDIGDDNPEPMHRRNHTLFGNMWSSIRRPGQFPKKGNNVAPKTSATSAAATAAMHAVLSGAEAYKAPYCQINNGFNRMAKNEPAHDNIPLHPLLLDGSNTAQTGRVYNANELSKENKLTTSTSVTSTKSTTTVLNDDVESNSSSSIRRRNRLNFNNMLMLMVDDKTKEKYCYDSVDEGSEGSVMTSKTSSTNDTCDGFKNMFSARNKNDLQKIPLSYMRSPNQHNIGKYQFFVRQNAFQSSPQNLRKKITPSKVSGEAQMSPTIYSQSPRQQMTSKSTDTEESALLCDSKSDSIVIGHSSSSQEPNSLSLELTDPSLIEGSCWKKPSQAEDWSIRPNRKNQRYSHCGRVTNGGNIGRGKFIPARVIHSTPSSPQDKRTKSIEVIGSAESLVGRVLLEQGLGKYCDPEFVHCAQMEMQEALEMTQEEMDLAAHELMLQERFNRNNNQPKNVQPKQKRKHDNQIL

>AAF53504.1|Drosophila_melanogaster_Cav1

MGGGELVNCIAYDDNTLVIERKPSPSSPSTSRRYLKAETPTRGSRKYNRKSSAKSDLEVVVVKPEHHHQHRSPTITLPVPANPLTTSASAGSSPTGAGLAAGLGTASGTVLQQSCSALDPPEDSNQPSGTRRRPTSTELALSNVTSQIVNNATYKLDFKQRRHKSNNGGSESGSLTGIATGPATSPAGPTGPTSSSGKRRKSSCTSCGGGGISAPPPRLTPEEAWQLQPQNSVTSAGSTNSSFSSGGGRDDNSSYSAVGGDSSSSNSCNCDITGDNSTLHGFGVGDVCSFIADCDDNSEDDDGDPNNQDLSSQTLRTAAIVAAVAAAAKEQAQEQSLADCESFSDRRQDADEDVRIIQDCCGGNNDSLEDVGEVDDNADVVVRKNSRNRPSIRRTCRITEEDDDEDENADYGDFDREDQELDDEEPEGTTIDIDEQEQQHDQGDSAEEEDDDEDVDEYFEEEEDDTQAFSPFYSSSAELIDNFGGGAGKFFNIMDFERGASGEGGFSPNGNGGPGSGDVSRTARYDSGEGDLGGGNNIMGIDSMGIANIPETMNGTTIGPSGAGGQKGGAAAGAAGQKRQQRRGKPQPDRPQRALFCLSVKNPLRALCIRIVEWKPFEFLILLTIFANCIALAVYTPYPGSDSNVTNQTLEKVEYVFLVIFTAECVMKILAYGFVLHNGAYLRNGWNLLDFTIVVIGAISTALSQLMKDAFDVKALRAFRVLRPLRLVSGVPSLQVVLNSILKAMVPLFHIALLVLFVIIIYAIIGLELFSGKLHKACRDEITGEYEENIRPCGVGYQCPPGYKCYGGWDGPNDGITNFDNFGLAMLTVFQCVTLEGWTDVLYSIQDAMGSDWQWMYFISMVILGAFFVMNLILGVLSGEFSKERNKAKNRGDFQKLREKQQIEEDLRGYLDWITQAEDIEPDAVGGLISDGKGKQPNEMDSTENLGEEMPEVQMTESRWRKMKKDFDRVNRRMRRACRKAVKSQAFYWLIIVLVFLNTGVLATEHYGQLDWLDNFQEYTNVFFIGLFTCEMLLKMYSLGFQGYFVSLFNRFDCFVVIGSITETLLTNTGMMPPLGVSVLRCVRLLRVFKVTKYWRSLSNLVASLLNSIQSIASLLLLLFLFIVIFALLGMQVFGGKFNFDGKEEKYRMNFDCFWQALLTVFQIMTGEDWNAVMYVGINAYGGVSSYGALACIYFIILFICGNYILLNVFLAIAVDNLADADSLSEVEKEEEPHDESAQKKSHSPTPTIDGMDDHLSIDIDMEQQELDDEDKMDHETLSDEEVREMCEEEEEVDEEGMITARPRRMSEVNTATKILPIPPGTSFFLFSQTNRFRVFCHWLCNHSNFGNIILCCIMFSSAMLAAENPLRANDDLNKVLNKFDYFFTAVFTIELILKLISYGFVLHDGAFCRSAFNLLDLLVVCVSLISLVSSSNAISVVKILRVLRVLRPLRAINRAKGLKHVVQCVIVAVKTIGNIVLVTCLLQFMFAVIGVQLFKGKFFKCTDGSKMTQDECYGTYLVYDDGDVHKPRLREREWSNNRFHFDDVAKGMLTLFTVSTFEGWPGLLYVSIDSNKENGGPIHNFRPIVAAYYIIYIIIIAFFMVNIFVGFVIVTFQNEGEQEYKNCDLDKNQRNCIEFALKAKPVRRYIPKHGIQYKVWWFVTSSSFEYTIFILIMINTVTLAMKFYNQPLWYTELLDALNMIFTAVFALEFVFKLAAFRFKNYFGDAWNVFDFIIVLGSFIDIVYSEIKSKDTSQIAECDIVEGCKSTKKSAGSNLISINFFRLFRVMRLVKLLSKGEGIRTLLWTFIKSFQALPYVALLIVLLFFIYAVVGMQVFGKIALDGGNAITANNNFQTFQQAVLVLFRSATGEAWQEIMMSCSAQPDVKCDMNSDTPGEPCGSSIAYPYFISFYVLCSFLIINLFVAVIMDNFDYLTRDWSILGPHHLDEFIRLWSEYDPDAKGRIKHLDVVTLLRKISPPLGFGKLCPHRMACKRLVSMNMPLNSDGTVLFNATLFAVVRTSLSIKTDGNIDDANSELRATIKQIWKRTNPKLLDQVVPPPGNDDEVTVGKFYATYLIQDYFRRFKKRKEQEGKEGHPDSNTVTLQAGLRTLHEVSPALKRAISGNLDELDQEPEPMHRRHHTLFGSVWSSIRRHGNGTFRRSAKATASQSNGALAIGGSASAALGVGGSSLVLGSSDPAGGDYLYDTLNRSVADGVNNITRNIMQARLAAAGKLQDELQGAGSGGELRTFGESISMRPLAKNGGGAATVAGTLPPEANAINYDNRNRGILLHPYNNVYAPNGALPGHERMIQSTPASPYDQRRLPTSSDMNGLAESLIGGVLAAEGLGKYCDSEFVGTAAREMREALDMTPEEMNLAAHQILSNEHSLSLIGSSNGSIFGGSAGGLGGAGSGGVGGLGGSSSIRNAFGGSGSGPSSLSPQHQPYSGTLNSPPIPDNRLRRVATVTTTNNNNKSQVSQNNSNSLNVRANANSQMNMSPTGQPVQQQSPLRGQGNQTYSS

>EFX89598.1|Daphnia_pulex_Cav1

MESQNGLAMSPETPVGVPGTPAAAGVAAQATAPSATVGPAKTLAEAEAAAAAAAAAAKRKAMSRRGKPPPERPQRALLCLSLTNPLRKLCISVVEWKPFEYLILLTIFANCVALAVYTPYPNGDSNITNAYLEKVEYVFLVIFTIECVMKIIAYGFVAHSGAYLRNTWNLLDFTIVVIGAVSTALSTMMKDGFDVKALRAFRVLRPLRLVSGVPSLQVVLNSILKAMVPLLHIALLVIFVIIIYAIIGLELFSGKLHTTCYDPETGDMMKDPHPCSNSSEVGFDCRTIGMVCLPDWEGPNDGITNFDNFGLAMLTVFQCVTLEGWTDVLYQIEDAMGNSWQWIYFISMVIIGAFFVMNLILGVLSGEFSKEREKAKARGDFHKLREKQQIEEDLRGYLDWITQAEDIEPEGEDRHHDEPKHEQDNSVDKTEESSNADTSQDSWWAQKKRNWDRSNRRMRRACRKAVKSQAFYWLIIILVFLNTGVLATEHYQQPEWLDFFQDVTNIFFIVLFAFEMLLKMYSLGFQGYFVSLFNRFDCFVVISSIVEVVLTKTDLMPPLGVSVLRCVRLLRVFKVTKYWRSLSNLVASLLNSIQSIASLLLLLFLFMVIFALLGMQVFGGKFNFENQELPRSNFDSFWQSLLTVFQILTGEDWNAVMYDGIKAYGGVASLGILACIYFIILFICGNYILLNVFLAIAVDNLADAESLTAIEKEEADQAEAKAEAEADLDKSAASDSATKDGDDRSQEELDEEEEDNGLGDDQGSYDDDNNGEEEEAEVGEEEEEEEDEDGQPVTSRPRRASAVARSNNKIIPIPPYTSFFILSHSNRFRVFCHWFCNHNYFGNIILACILISSAMLAAEDPLSADTDRNKILNHFDYFFTTVFTIEIALKVIAYGLVLHKGAFCRSAFNLLDLLVVCVSLISFGFSSGAISVVKILRVLRVLRPLRAINRAKGLKHVVQCVIVAVKTIGNIMLVTCLLEFMFAVIGVQLFKGKFFKCNDPSKMTRDDCIGTFIQYEDGDIERPTVSNRTWENNDFHFDDVGKAMLTLFTVSTFEGWPGLLYVSIDSHTEDMGPMHNYRPMIACFYFIYIIIIAFFMVNIFVGFVIVTFQNEGEQEYRNCELDKNQRNCIEFALKAKPFRRYIPKQRFQYKVWWMVTSQPFEYVLFTLISMNTITLGMKFYGQPEVYTQALDILNMIFTSVFALEFLLKLAAFRFKNYFSDPWNVFDFVIVLGSFIDIVYSQLNVSKEMLVLLYGVPDSNLISINFFRLFRVMRLVKLLSRGEGIRTLLWTFMKSFQALPYVALLIVMLFFIYAVIGMQVFGKIALDDDTAIHRNNNFQTFPQAVLVLFRSATGEAWQDVMLGCSSQKAPACDPLSDEIRNNSTDHCGTEFAIPYFISFYVLCSFLIINLFVAVIMDNFDYLTRDWSILGPHHLDEFVRLWSEYDPDARGRIKHLDVVTLLRKISPPLGFGKLCPHRVACKRLVSMNMPLNSDGTVMFNATLFAVVRTSLKIYTEGNIDSANETLRAVIKKIWKRTNGKLLDQVVPPPGIDDEVTVGKFYATFLIQDYFRRFKKRKEMLNKDSALDGEGAGVDTVTLQAGLRTLHDAGPELKRAISGNLDELMERDPEPMHRRNHSLFGNVWSNMRRGGGPSSRQRPTESSPLKLSPQMLLKATVAVPNGQSVPINSSSDDLSEAMPLRPLRLRQQQNQTNNNGSLPKYDENFLHPSYNAQEEAVAGSAESLVGKVLVEQGLGKFIDDDFIRTTSREMQEAMDMTQEEMDRAAHHLLMLTQDGDGTAVRTPALSGAAIRRPGGQPLTLALNSAFNEVDHVPQSRHKS

>KZS09199.1|Daphnia_magna_Cav1

MMESQNGLATSPETAVVSATPAAAASGIAAPVTAPPTTGGPAKSLAEAEAAAAAAAAAAKRKAMSRRGKPPPERPQRALFCLSLTNPVRKLCISVVDYCIGYNILNPLSLCTPFEYLILLTIFANCVALAVYTPYPNGDSNITNAYLEKVEYVFLVIFTIECVMKIIAYGFVAHSGAYLRNTWNFLDFTIVVIGAVSTALSTMMKDGFDVKALRAFRVLRPLRLVSGVPSLQVVLNSILKAMVPLLHIALLVIFVIIIYAIIGLELFSGKLHTTCYDPETGDMMKDPHPCSNSSEVGFDCATIGMVCLPDWEGPNDGITNFDNFGLAMLTVFQCVTLEGWTDVLYQIEDAMGNSWQWIYFISMVIIGAFFVMNLILGVLSGEFSKEREKAKARGDFHKLREKQQIEEDLRGYLDWITQAEDIEPEGEDRQHDESKHAKNKREQDNSVDKTEESSNADTSQDSWWTQKKRNWERSNRRMRRACRKAVKSQAFYWLIIILVFLNTGVLATEHYQQPEWLDFFQDVTNIFFIVLFAFEMLLKMYSLGFQGYFVSLFNRFDCFVVISSIVEVVLTKTDLMPPLGVSVLRCVRLLRVFKVTKYWRSLSNLVASLLNSIQSIASLLLLLFLFMVIFALLGMQVFGGKFNFENQELPRSNFDSFWQSLLTVFQILTGEDWNVVMYDGIKAYGGVSSLGILACIYFIILFICGNYILLNVFLAIAVDNLADAESLTAIEKEEADQAGAKAEAEADLDKSATSDSATKDGDDRSQEELDEEEEDNMDDNGKEGDGDDNAQADGNHGDISERKRSSKKKKSDGGSTRIEIEQDYDDEGTEDASKDDEGLGDDQGSYDDDNNGEEEEAEVGEEEEEEEDEDGQPGTLRPRRASAAARSNKIIPIPPYTSFFILSHSNRFRVFCHWFCNHNYFGNIILACILISSAMLAAEDPLSADTDRNKILNHFDYFFTTVFTIEIALKVIAYGLVFHKGAFCRSAFNLLDLLVVCVSLISFGFSSGAISVVKILRVLRVLRPLRAINRAKGLKHVVQCVIVAVKTIGNIMLVTCLLEFMFAVIGVQLFKGKFFKCNDPSKMTRDDCIGTFIQYEDDDIERPVVSNRTWENNDFHFDDVGKAMLTLFTVSTFEGWPGLLYVSIDSHTEDMGPMHNYRPMIACFYFIYIIIIAFFMVNIFVGFVIVTFQNEGEQEYRNCELDKNQRNCIEFALKAKPFRRYIPKQRFQYKVWWMVTSQPFEYVIFTLIITNTITLGMKFYGQPEVYTQALDVLNMIFTSVFALEFVLKLAAFRFKNYFSDPWNVFDFVIVLGSFIDIVYSQLNPDSNLISINFFRLFRVMRLVKLLSRGEGIRTLLWTFMKSFQALPYVALLIVMLFFIYAVIGMQVFGKIALDNDTAIHRNNNFQTFPQAVLVLFRSATGEAWQDVMLGCSSQKAPACDPKSDEFRNNATEHCGTEFAIPYFISFYVLCSFLIINLFVAVIMDNFDYLTRDWSILGPHHLDEFVRLWSEYDPDARGRIKHLDVVTLLRKISPPLGFGKLCPHRVACKRLVSMNMPLNSDGTVMFNATLFAVVRTSLKIYTDGNIDTANETLRAVIKKIWKRTNSKLLDQVVPPPGIDDEVTVGKFYATFLIQDYFRRFKKRKEMLNKDVTLDGEVTGVDTVTLQAGLRTLHDAGPELKRAISGNLDELMERDPEPMHRRNHTLFGSVWSNMRRGGGPSSRQRPTESSPLKLSPQMLMKTANASNGQSLPVSSSSDELSEAMPLRPLRLRQNQTNNNGSLPYNAHEEAVAGSAESLVGKVLVEQGLGKFIDDDFIRTTSREMQEAMDMTKEEMDRAAHHLLMLTQDEIGTSRSPQALSGAAVRRPGGQPLTLALNSAFSEVDHVPQSRHNS

>NP_001023079.1|Caenorhabditis_elegans_Cav1

MSVLASMMSSGEDEEQAAADEQERTDLWQQTLQAAVAASSSQDATKKRPAQRKPLRQTNVVERSERSLLCLSLNNPIRKLCISIVEWKPFEFLILFMICANCIALAIYQPYPAQDSDYKNTLLETIEYVFIVVFTIECVLKIVAMGFMFHPSAYLRNAWNILDFIIVVIGLVSTILSKMSIQGFDVKALRAFRVLRPLRLVSGVPSLQVVLNAILRAMIPLLHIALLVLFVILIYAIIGLELFCGKLHSTCIDPATGQLAQKDPTPCGTDTEGSAFKCQPSDSLTNMGVRWECSSNTTWPGPNNGITNFDNFGLAMLTVFQCVSLEGWTDVMYWVNDAVGREWPWIYFVTLVILGSFFVLNLVLGVLSGEFSKEREKARARGLFQKFREKQQLEEDLKGYLDWITQAEDIEPVNEDEQEDEPVAQTVVGEEADEEGEERVEDVRPSKWAARMKRLEKLNRRCRRACRRLVKSQTFYWLVILLVLLNTLVLTSEHYGQSEWLDHFQTMANLFFVILFSMEMLLKMYSLGFTTYTTSQFNRFDCFVVISSILEFVLVYFDLMKPLGVSVLRSARLLRIFKVTKYWTSLRNLVSSLLNSLRSIISLLLLLFLFIVIFALLGMQVFGGKFNFNPQQPKPRANFDTFVQALLTVFQILTGEDWNTVMYHGIESFGGVGTLGVIVCIYYIVLFICGNYILLNVFLAIAVDNLADADSLTNAEKEEEQQEIEGEDEEFEEGEDEGEEHGMDEPEGDEEMTSARPRRMSEVPAASTVKPIPKASSLFILSHTNSFRVFCNMVVNHSYFTNAVLFCILVSSAMLAAEDPLQANSTRNMILNYFDYFFTSVFTVEITLKVIVFGLVFHKGSFCRNAFNLLDILVVAVSLTSFVLRTDAMSVVKILRVLRVLRPLRAINRAKGLKHVVQCVIVAVKTIGNIMLVTFMLQFMFAIIGVQLFKGTFFLCNDLSKMTEAECRGEYIHYEDGDPTKPVSKKRVWSNNDFNFDNVGDAMISLFVVSTFEGWPQLLYVAIDSNEEDKGPIHNSRQAVALFFIAFIIVIAFFMMNIFVGFVIVTFQNEGEREYENCELDKNQRKCIEFALKAKPHRRYIPRNRLQYRVWWFVTSRAFEYVIFLIIVMNTVSLACKHYPSSRGFEDFLDVFNLIFTGVFAFEAVLKIVALNPKNYISDRWNVFDLLVVVGSFIDITYGKLNPGGTNLISINFFRLFRVMRLVKLLSRGEGIRTLLWTFMKSFQALPYVALLIVLLFFIYAVIGMQFFGKVALDDSTSIHRNNNFHSFPAAILVLFRSATGEAWQDIMLSCSDREDVRCDPMSDDYHKGGLNESRCGNNFAYPYFISFFMLCSFLVINLFVAVIMDNFDYLTRDWSILGPHHLEEFVRLWSEYDPDAKGRIKHLDVVTLLRKISPPLGFGKLCPHRLACKRLVSMNMPLNSDGTVCFNATLFALVRTNLKIYTEGNIDEANEQLRSAIKRIWKRTHKDLLDEVVPPAGKEDDVTVGKFYATFLIQDYFRRFKKRKEMEAKGVLPAQTPQAMALQAGLRTLHEIGPELKRAISGNLETDFNFDEPEPQHRRPHSLFNNLVHRLSGAGSKSPTEHERIEKGSTLLPFQPRSFSPTHSLAGAEGSPVPSQMHRGAPINQSINLPPVNGSARRLPALPPYANHIHDETDDGPRYRDTGDRAGYDQSSRMVVANRNLPVDPDEEEQWMRSGGPSNRSDRRNPHLREPMLVARGAALALAGMSSEAYEGTYRPVGEGKSVRLPFSSRPVLRPAEDSRPVDRLIGQSLGLGRYADARIVGAARREIEEAYSLGEQEIDLAADSLAPLMQHVGMHDIRDINENSRSALLRPAENSSRQHDSRGGSQEDLLLVTTL

>PIC34019.1|Caenorhabditis_nigoni_Cav1

MSVLASMMSSGEDEEQAAADEQERSDLWQQTLQAAVAASSSQDATKKRPAQRKPLRQTNVVERSERSLLCLSLNNPIRKLCISIVEWKPFEFLILFMICANCIALAIYQPYPAQDSDYKNTLLETIEYVFIVVFTIECVLKIVAMGFLFHPSAYLRNAWNILDFIIVVIGLVSTILSKMSIQGFDVKALRAFRVLRPLRLVSGVPSLQVVLNAILRAMIPLLHIALLVLFVILIYAIIGLELFCGKLHSTCIDPATGQLAQKDPTPCGNDGSAFKCRPSDSLTNMGVRWECSSNTTWPGPNNGITNFDNFGLAMLTVFQCVSLEGWTDVMYWVNDAVGREWPWIYFVTLVILGSFFVLNLVLGVLSGEFSKEREKARARGLFQKFREKQQLEEDLKGYLDWITQAEDIEPVNEDEQEDEPAAAAVTGEEVDEEGEERVEDVRPSKWSARVKRLEKFNRRCRRACRRLVKSQTFYWLVILLVLLNTLVLTSEHYGQSEWLDHFQTMANLFFVILFSMEMLLKMYSLGFTTYTTSQFNRFDCFVVISSILEFVLVYFDLMKPLGVSVLRSARLLRIFKVTKYWTSLRNLVSSLLNSLRSIISLLLLLFLFIVIFALLGMQVFGGKFNFNPQQPKPRANFDTFVQALLTVFQILTGEDWNTVMYHGIESFGGVGTLGVIVCIYYIVLFICGNYILLNVFLAIAVDNLADADSLTNAEKEEEQQEIEGEDEEFDEGEEEGDEHGAEEPEGEEDMTSARPRRMSEVPAASTVKPIPKASSLFILSHTNSFRVFCNMVVNHSYFTNAVLFCILVSSAMLAAEDPLQANSTRNMVLNYFDYFFTSVFTVEITLKVIVFGLVFHKGSFCRNAFNLLDILVVAVSLTSFVLRTDAMSVVKILRVLRVLRPLRAINRAKGLKHVVQCVIVAVKTIGNIMLVTFMLQFMFAIIGVQLFKGTFFLCNDLSKMTEAECRGEYIHYEDGDPTKPVSKKRVWSNNDFNFDNVGDAMVSLFVVSTFEGWPQLLYVAIDSNEEDKGPVHNSRQAVALFFIAFIIVIAFFMMNIFVGFVIVTFQNEGEREYENCELDKNQRKCIEFALKAKPHRRYIPRNRLQYRVWWFVTSRAFEYVIFLIIVMNTVSLACKHYPSSRGFEDFLDVFNLIFTGVFAFEAVLKIVALNPKNYISDRWNVFDLLVVVGSFIDITYGKLNPGGTNLISINFFRLFRVMRLVKLLSRGEGIRTLLWTFMKSFQALPYVALLIVLLFFIYAVIGMQFFGKVALDDSTSIHRNNNFHSFPAAILVLFRSATGEAWQDIMLSCSDREDVRCDPMSDDYSKGGFNESRCGNNFAYPYFISFFMLCSFLVINLFVAVIMDNFDYLTRDWSILGPHHLEEFVRLWSEYDPDAKGRIKHLDVVTLLRKISPPLGFGKLCPHRLACKRLVSMNMPLNSDGTVCFNATLFALVRTNLKIYTEGNIDEANEQLRSAIKRIWKRTHKDLLDEVVPPAGKEDDVTVGKFYATFLIQDYFRRFKKRKEMEAKGVLPAQTPQAMALQAGLRTLHEIGPELKRAISGNLETDFNFDEPEPQHRRPHSLFNNLVHRLSGAGAKSPTEHERLERGATLLPYEARSFSPTHSLAGAEGSPVPSQMHRGAPINQSINLPPVNGSARRLPALPPYANHIHDETDDGPRYRDTGDRAGYDPSQNRMVVANRNLPVDPDEEERWMRGGPSNRSDRRNLPIREPMLVARGAALALAGMSSEAYEGTYRPVGEGKSVRLPFSSRPVLRPAEDSRPADLLIGQSLGLGRYADARVVGAARREIEEAYQLGEQEIDMAADSLAPLMQHVGMHDIRDINENSRSALLRPAGESSRRQHDSHGGSQEDLLLVTTL

>PAV74346.1|Diploscapter_pachys_Cav1

MMSSGEDEEPPTGIGGGDSQGQGPTGSQAGGSIGGGVSGEHEQERTDLWQQTLQAAVAASAGGDPNKKRPAQRKPLRQANVVERPFEFLILLMICLNCIALAVTQPYPAQDSDSINAKLERVENVFIVVFTIECILKVIAYGFMFHPSAYLRNAWNVLDFIIVVIGIISTVLAGMHLQGFDVKALRAFRVLRPLRLVSGVPSLQVVLNAILRAMIPLLHIALLVLFVILIYAIIGLELFCGKLKQTCIDPATGQLAMKDPTPCGNDGTSFQCIPTDAMIAMGVSWECSNNTNWTGPNFGITNFDNFGLAMLTVFQCVSLEGWTEVMYWVNDAVGREWPWIYFVSLVILGSFFVLNLVLGVLSGEFSKEREKARARGLFQKFREKQQLEEDLKGYLDWITQAEDIEPVNEDEQDQEPQPQTQQQQGEEQDEEGEERPDEARPSKWSGRIKRLEKLNRRCRRSCRRLVKSQTFYWLVILLVFLNTLVLTSEHYGQSDWLDEFQNWANLFFVILFTMEMFLKMYSLGLTTYTTSQFNRFDCFVVISSILEFILVNCKLMKPLGVSVLRSARLLRIFKVTKYWTSLRNLVSSLLNSLRSIMSLLLLLFLFIVIFALLGMQVFGGKFNFNPQAPKPRANFDTFIQALLTILTGEDWNTVMYNGINSFGGVGTVGVVVSIYYIVLFICGNYILLNVFLAIAVDNLADADSLTNAEKEEEQQDLEAEDEYDENEEGAELDEHEGDDDLVTARPRRMSELPQTTNIKPIPKASSLFILSHTNPFRVFCNKVVNHSYFTNSVLVCILVSSAMLAAEDPLKADSPRNLILNKFDYFFTTVFTIEITLKVVFFGLILHKGSFCRNAFNLLDIIVVAVSIISPILKSDAFSVVKILRVLRVLRPLRAINRAKGLKHVVQCVIVAVKTIGNIMLVTFMLQFMFAIIGVQLFKGTFFYCNDISKMTEAECRGEYIHYEDGDPTKPVAKKRAWQNNDFNFDHVGNAMISLFVVSTFEGWPDLLYIAIDSNEENKGPIHNNRQAVALFFIAFIIVIAFFMMNIFVGFVIVTFQNEGEREYENCELDKNQRKCIEFALKAKPHRRYIPRNRFQYRVWWFVTSRAFEYLIFLIIVLNTVALACRHDPSSQSFNDILDKLNLCFTSVFAFEAFFKIIALNPKNYFGDRWNAFDFIIVLGSFIDITWSNLNPDNKSLISINFFRLFRVMRLVKLLSRGEGIRTLLWTFMKSFQALPYVALLIVLLFFIYAVIGMQVFGKIALDDSTEIHRNNNFQTFFSAVLVLFRSATGEAWQQIMLSCSERDEVRCDRKSDDPGQKCGTNFAYPYFISFFMLCSFLVINLFVAVIMDNFDYLTRDWSILGPHHLEEFVRLWSEYDPDAKGRIKHLDVVTLLRKISPPLGFGKLCPHRLACKRLVSMNMPLNSDGTVCFNATLFALVRTNLKIYTEGNIDEANEQLRSAIKRIWKRTPAKMLDEVVPPAGKEDDVTVGKFYATFLIQDYFRRFKKRKELEAKGIVPTHTPQAMALQVEHLHLFFISAKNFHVLVWNTRNTLFKNSMHQIS

>PDM72512.1|Pristionchus_pacificus_Cav1

MSVFAAMGSSPTEEEQVDEERGDLWQATLQAAVQSAQEGGKKRPVARKPLRATNTVERPDRSLFCLNLANPLRKACIAITEWRPFEWLILFMICANCIALAVYQPYPAQDSDLKNNILEQIEYLFIVVFTIECVLKVIAQGFLMHPGAYLRNAWNSLDFVIVVIGLVSTILARMNIQGFDVKALRAFRVLRPLRLVSGVPSLQVVLNAILRAMIPLFHIALLVLFVIVIYAIIGLELFCGKLHSTCVDQNTQQFAMKEQTPCGTSATAFNCNPDTAFLNTGESANWVCIQNSSWTGPNNGITNFDNFGLAMLTVFQCVSLEGWTDVMYWVNDAVGVEWPWIYFVSLVILGSFFVLNLVLGVLSGEFSKEREKARARGLFQKFREKQQLEEDLKGYLDWITQAEDIEPINEEEQEEEPPPPPKNGGLAVSVGDEENEEGVETVDETRPSAFRLRMRRFEKMNRRFRRACRRLVKSQTFYWLVIFLVFLNTLVLTTEHHNQPPWLDHFQTAGNLFFVILFSLEMLLKMYSLGFTSYTRSQFNRFDCFVVISSIIEFVLVYLELMKPLGVSVLRSARLLRIFKVTKYWASLRNLVASLLNSLRSIMSLLLLLFLFIVIFALLGMQVFGGKFNFNPQAPKPRANFDTFIQSLLTVFQILTGEDWNAVMYNGIESFGGVGSMGMLVCIYYIVLFICGNYILLNVFLAIAVDNLADADSLTNAEKEEEAHEMDEEEEMDEGLYDEDGMEKEERELDEELADDMESARPRRMSEVPAVNAAKPIPKASSLFVLSHTNPFRVFCNKIINHAYFTNAVLVCILVSSAMLAAEDPLNSESDRNQVLNYFDYFFTSVFTVEISLKVVVYGLILHKGAFCRNAFNLLDILVVAVSLVSFLLKSNAISVVKILRVLRVLRPLRAINRAKGLKHVVQCVIVAVKTIGNIMLVTFMLQFMFAIIGVQLFKGTFYACNDQSKVTERDCRGQYLQYEDGDPTKPVSKLRVWSNNDFNFDNVANAMVSLFVVSTFEGWPDLLYVAINSNEEDHGPVYNSRQSVAVFFIAFIIVIAFFMMNIFVGFVIVTFQNEGEREYENCELDKNQRKCIEFALKAKPHRRYIPRNRFQYRVWWFVTSRAFEYVIFLIIVLNTLSLACKHYPSGESFDHVLDMLNLIFTGVFAFEAFFKIIALNPKNYFGDRWNAFDFIIVLGSFIDIIYGRVSPGQNFISINFFRLFRVMRLVKLLSRGEGIRTLLWTFMKSFQALPYVALLIVLLFFIYAVIGMQMFGRVALDDTTEIHRNNNFHTFPMAVLVLFRSATGEAWQLIMLSCSSREDVKCAKGSDDRRQEELTWKDASEIPMCGNDFAYPYFISFFMLCSFLVINLFVAVIMDNFDYLTRDWSILGPHHLEEFVRLWSEYDPDAKGRIKHLDVVTLLRKISPPLGFGKLCPHRLACKRLVSMNMPLNSDGTVCFNATLFALVRTNLKIYTEGNIDEANEQLRSAIRRIWKRTPIGILDEVVPPAGKEDDVTVGKFYATFLIQDYFRRFKKRKELEAKGIVGGNHSSTPHAMALQAGLRTLHEIGPELKRAISGNLETDFNFEAEEEPTHRRPHVLFNHIVSALSNRSPLGHPERQAVLLPYNPSGPRQFSPTNSMMEMDGIDPPGRMRPVTHSSLNLPSNGGTIPHRRLPNAPPIETAAGAAMIMVNRTHQFDSDDDDNWSGDRDRHRPNRIMADYPYDHSRTSMASPVRLARNQAIAMVGMGPDVPEATYRPAPDGKSVRLPFAPRPVLRPADRLVSEALGLGRYGDERVVGAARREIEEAYSLDDSQIDQAAAGLAPNLMEHAGMNEMRDYNEYSRSSLLRPANSEDDDRSSHDDLLLVTTL

>XP_024499702.1|Strongyloides_ratti_Cav1

MSVLASMIQSSGEDEENVHEEVHNRGDLWQQTLQAAVAASGQAGDVNRKRAQQRKPLRQTNVVERNERSLMCLSVRNPIRRACIGIVEWRPFEWLILLMICANCIALAVYQPYPAQDSDTKNTILEQIEYLFIIVFTIECILKVIALGFLCHSGAYLRNAWNMLDFLIVVIGLVSTVLSRMNIQGFDVKALRAFRVLRPLRLVSGVPSLQVVLNAILRAMVPLLHIALLVMFVIIIFAIIGLELFCGKLHSTCVDVNTGELAMKNPTPCGFAASAYHCEPNFNQINTTISKWICSSNTTWQGPNNGITNFDNFGLAMLTVFQCVSLEGWTDVMYWVNDAVGVEWPWIYFVTLVILGSFFVLNLVLGVLSGEFSKEREKARARGLFQKFREKQQLEEDLKGYLDWITQAEDIEPVNDDDNVEDEQQFNEDGMDEEGEERTEDSRPTWFIIKMRLFQKWNRRCRRACRRLVKSQSFYWLVIILVLLNTLVLTSEHYGQSEWLDNFQTAANLFFVILFSMEMLLKMYSLGLTTYTTSQFNRFDCFVVISSILEFILVYFNLMKPLGVSVLRSARLLRIFKVTKYWTSLRNLVSSLLNSLRSIMSLLLLLFLFIVIFALLGMQVFGGKFNFDPQQPKPRANFDTFIQSLLTVFQILTGEDWNTVMYNGIESFGGVGSIGVLVSIYFIVLFICGNYILLNVFLAIAVDNLADADSLTNAEKEEEQGEVEYEEGEEDYEDEKYLENRGDDDQEEDEHAVIPEGDDDGETKEEVGARPRRMSHLPQQTKEKQIPKASSLFILSHTNPFRVFCNKIVNHQYFTNSVLVCILVSSAMLAAEDPLQAQSPRNLILNYFDYFFTTVFTIEISLKVIVFGLVIHKGSFCRNAFNLLDILVVAVSLISFVLKSDAISVVKILRVLRVLRPLRAINRAKGLKHVVQCVIVAVKTIGNIMLVTFMLQFMFAIIGVQLFKGTFFSCNDPSKMTEAECRGEFIQYEDNNPTKPMRLRRAWTKNDFNFDNVLDAMISLFVVSTFEGWPDLLYVAINSNEEDRGPVHNARQAVAFFFITFIVVIAFFMMNIFVGFVIVTFQNEGEREYENCELDKNQRKCIEFALKAKPHRRYIPRNRFQYRVWWFVTSQFFEYAIFIIILLNTTTLAMKHYPPDTQMDHILDVLNLIFTAVFAFEALFKIIALNPKNYFGDRWNAFDFVIVLGSFIDIIYGKLSPGSNIISINFFRLFRVMRLVKLLSRGEGIRTLLWTFMKSFQALPYVALLIVLLFFIYAVIGMQVFGKVALDSETQIHRNNNFHTFPAAVLVLFRSATGEAWQEIMLACADREEVKCDPASDEYKKDPNALCGVNFAYPYFISFFMLCSFLVINLFVAVIMDNFDYLTRDWSILGPHHLEEFVRLWSEYDPDAKGRIKHLDVVTLLRKISPPLGFGKLCPHRLACKRLVSMNMPLNSDGTVCFNATLFALVRTNLKIYTEGNIDEANEQLRSAIRRIWKRTPLKMLDEVVPPAGRDDDVTVGKFYATFLIQDYFRRFKKRKELESKGVNSHQHSQTMALQAGLRTLHEIGPELKRAISGSLDTDWAPETEEPQHRRTHSLFNNIVHALSGGSIKLSKEERNRRLLLVGDNNSSLSPSHSIPGDELMSLAIHSENNQGTDIPNSHSIPHNLFNEMDNNTMYPGPQQRQPRSRILPSIHLPNYKTPDLHRSNFVPYQRISKGSSTDGEYNIPQAGQKYIFINRSAPSDSEEEDWYERRRRIDAEEKMMCHRASSTLMRDPLLLPASSGYNDAYDCHVYRPAPDGKGVRLPYTSRPMIGGPIDVPPPPPPSSFIPRQELTDNLVAKALSLGRYMDHNVVEIARREIAEAYQLDEPSMVRAAYSLSENPPGPAPINIPNSSNISHHQHPLHHSHHIDMNDGYHEPEYLEHVGGIRPEEVSDINQYSQQGLFRPVDPHTLRNNTNQNRGNDGNSSSQEDLLLVTTL

>KRY55670.1|Trichinella_britovi_Cav1

MLAGGETLLQIEFSWLLWSRRRSRASYLCWHRESRPEQVVPSPIAEPRVASIQTMIVRQVRHHARSEANFNDDVNIGKELSGNEAAALLHGDEARASRADLWQQTLQAAVAAQTESSTTTKKRQQQRKMQRNVQQERPERSLLCLGLKNPIRKLFISIVEWKPFEWLILCMICANCIALAVYQPFPAHDSDRKNAVLLCAIGKTFFDFVPAELRPAIDRQPGTAFQNRKHKLNPQPGQRTATVAAVGHTGWGVGLFSEEQVEYIFIVVFTIECVMKVIAYGFLFHPGAYLRNGWNLLDFLIVVIGLISTALSTLNIHGFDVKALRAFRVLRPLRLVSGVPSLQVVLNSILRAMVPLFHIALLVLFVIIIYAIIGLELFCGKLHKTCVDQWTGEHVPDPGPCGESHTSFHCDRSKNLVCTENHTWPGPNDGITNFDNFGLAMLTVFQCISLEGWTDVMYWVNDSVGREWPWIYFITLVILGSFFVLNLVLGVLSGEFSKEREKARARGLFQKFREKQQLEDDLKGYLDWITQAEDIDLVNEEDEEQEAMDREEFGADGEGGEEGGSKEEFQRQSWFSMKIKRLKKLNRRCRRSCRRIVKSQAFYWLVIVLVFLNTMVLTSEHYGQPEWLDHFQEIANLFFVVLFTLEMFLKMYSLGFVNYFVALFNRFDCFVVIGSILEFALTFAGLMKPLGVSVLRSARLLRIFKVTKYWNSLRNLVASLLNSLRSIASLLLLLFLFIVIFALLGMQVFGGKFNTIDPNMNKPRANFDTFVQALLTVFQILTGEDWNAVMYNGIAAFGGVHSIGVIVCIYFIVLFICGNYILLNVFLAIAVDNLADAESLTAAEKEEENKRANEADPEDDMVVKATADEDAVSVGQFHESGEVKLPFNEPTDAEDEHLASGDDDQEREMENQSQFKPTARPHRQSELNLPKKTKPIPDASSLFLFSSTNKVRIFCNKVINHSYFTNSVLVCILVSSAMLAAEDPLQASSFRNEVPKFVLNYFDYFFTTVFTIEISLKVLVYGLILHKGSFCRNAFNLLDMLVVGVSLTSFGLKSGAISVVKILRVLRVLRPLRAINRAKGLKHVVQCVIVAVKTIGNIMLVTFMLEFMFAIIGVQIFKGSFFRCTDRARLTAEECKGTFIEFEGGDVTRPHVRNREWTNYDFNFDNVQNAMVALFVVSTFEGWPDLLHVAMDSSDEGIGPQYNARVSVAIFFITFIVVIAFFMMNIFVGFVIVTFQSEGEREYENCELDKNQRKCIEFALTAKPQRRYIPKNRFQYKIWWFVTSQPFEYAIFIIIILNTLILGMKHYKSSAAFDEALDVLNLFFTTVFALEFICKLFALTFKHHKMTRTFDDVLDTMNLIFTGIFAMEFILKVMAFRCKNYFGDAWNVFDFIIVLGSFIDIIYGKVSPGSNIISINFFRLFRVMRLVKLLSRGEGIRTLLWTFMKSFQALPYVALLIVLLFFIYAVIGMQIFGKIALNSNTEIHRNNNFQTFPSAVLVLFRSATGEAWQLIMLSCANTPAAMCDPESDDRGQPCGNDFAYPFFISFFMLCSFLIINLFVAVIMDNFDYLTRDWSILGPHHLDEFVRLWSEYDPDAKGRIKHLDVVTLLRKISPPLGFGKLCPHRLACKRLVSMNMPLNSDGTVCFNSTLFALVRTNLKIYTEERPFLFLKKSEKVESQSLVSSNIEEANEQLRAVIKRIWKRTPQRLLDEIVPPSGRDDEITVGKFYATYLIQDYFRRFKKRKEVEQKETNMQGNITMSLQFGEEFLSDLEAFDVLIVFLHVLLLQAGLRTLHEIGPEIKRAISGNLETDWSKEFEEPQHRRDHSLFGTLVHALQAHYKPFIEGNLTNPAYSNVNGDLKSQEGDDEAEQQHQQHQQEQQEQQQQLMFNKMNDLKSPPVKSDRDKISEFESDESVCNKPERRCNSFFSNIRRRMDSIVSPSIVLFDSDHDSSEHEMQERVRFVPRGSKPTGLSFEGSRWLPRKLSFRRTPIGGAAAVSNLQQGNNKRLQFSNAIILDDSDPDSVLHSENVVDLTESGDDSKLSVSPPLELRDDAYYLDGIDYNTWRPAPMGQGVRLRGRGNGRRKSRSMPLPHHEHQRNYADVLVEKVLADQGLGRYADPNLIRTTQLEIAEAYNMTEAQMHSAARSLMQRSPKYFEHMGGQRPADIKDFNQYSKTALLKPREINSSEEVDISDDMNMFMSVV

>KRY90155.1|Trichinella_pseudospiralis_Cav1

MIKLLILLSHHARSEANFNDDVNIGKELSGNEAAALLHGDEARASRADLWQQTLQAAVAAQTESSTTTKKRQQQRKMQRNVQQERPERSLLCLGLKNPVRKLFISIVEWKPFEWLILCMICANCIALAVYQPFPAHDSDRKNAVLEQVEYIFIVVFTIECVMKVIAYGFLFHPGAYLRNGWNLLDFLIVVIGLISTALSTLNIHGFDVKALRAFRVLRPLRLVSGVPSLQVVLNSILRAMVPLFHIALLVLFVIIIYAIIGLELFCGKLHKTCVDQWTGEHVPDPGPCGESHTSFHCDRSKNLVCTENHTWPGPNDGITNFDNFGLAMLTVFQCISLEGWTDVMYWVNDSVGREWPWIYFITLVILGSFFVLNLVLGVLSGEFSKEREKARARGLFQKFREKQQLEDDLKGYLDWITQAEDIDLVNEEDEEQEAMDREEFGADGEGGEEGASKEEYQRQSWFSMKIRRLKKLNRRCRRSCRRIVKSQAFYWLVIVLVFLNTMVLTSEHYGQPEWLDHFQEIANLFFVVLFTLEMFLKMYSLGFVNYFVALFNRFDCFVVIGSILEFALTFAGLMKPLGVSVLRSARLLRIFKVTKYWNSLRNLVASLLNSLRSIASLLLLLFLFIVIFALLGMQVFGGKFNTIDPNMNKPRANFDTFVQALLTVFQILTGEDWNAVMYNGIAAFGGVHSIGVIVCIYFIVLFICGNYILLNVFLAIAVDNLADAESLTAAEKEEEGKRANEADPDDTVTKAPADEDAVSVEQFHESGEVKLPFNEPTDAEDEHLASGDEDQEREMENQSQFKPTARPHRQSELNLPKKTKPIPDASSLFLFSSTNKVRIFCNKVINHSYFTNSVLVCILVSSAMLAAEDPLQASSFRNEVLNYFDYFFTTVFTIEISLKVLVYGLILHKGSFCRNAFNLLDMLVVGVSLTSFGLKSGAISVVKILRVLRVLRPLRAINRAKGLKHVVQCVIVAVKTIGNIMLVTFMLEFMFAIIGVQIFKGSFFRCTDRARLTAEECKGTFIEFEGGDVTRPHVRNREWTNYDFNFDNVQNAMVALFVVSTFEGWPDLLHVAMDSSDEGIGPQYNARVSVAIFFITFIVVIAFFMMNIFVGFVIVTFQSEGEREYENCELDKNQRKCIEFALTAKPQRRYIPKNRFQYKIWWFVTSQPFEYAIFIIIILNTLILGMKHYKSSAAFDEALDVLNLFFTTVFALEFICKLFALTFKHHKMTRTFDDVLDTMNLIFTGIFAMEFILKVMAFRCKNYFGDAWNVFDFIIVLGSFIDIIYGKPGSNIISINFFRLFRVMRLVKLLSRGEGIRTLLWTFMKSFQALPYVALLIVLLFFIYAVIGMQIFGKIALNSNTEIHRNNNFQTFPSAVLVLFRSATGEAWQLIMLSCANTPTAMCDPESDDRGQPCGNDFAYPFFISFFMLCSFLIINLFVAVIMDNFDYLTRDWSILGPHHLDEFVRLWSEYDPDAKGRIKHLDVVTLLRKISPPLGFGKLCPHRLACKRLVSMNMPLNSDGTVCFNSTLFALVRTNLKIYTEVSSNIEEANEQLRAVIKRIWKRTPQRLLDEIVPPSGRDDEITVGKFYATYLIQDYFRRFKKRKEVEQKETNLQGNITMSLQAGLRTLHEIGPEIKRAISGNLETDWSKEFEEPQHRRDHSLFGTLVHALQAHYKPFIEGNLANPAYSNVNGDLKSQEGDDEIEHHHYHQQQQEPEHQQEPQQMMMFKMNDLKSPPVKSDRDKISEFESDDSVCNKPERRCNSFFSNIRRRVDSIVSPSIVLFDSDHDSSEHEMQERLRFVPRGSKPTTGLSFEGSRWLPRKLSFRRAPIGAAAVSNSQQGNSNNNNNNKRLQFSNAIILDDSDPDSVMHPENVVDLTESGDDSKSSVSPPLELRDDAYYLDGIDYNTWRPAPMGQGVRLRRRGNGRRKSRSMPLPHDQRNYADVLVEKVLADQGLGRYADPNLIRTTQLEIAEAYNMTEAQMHSAARSLMQRSPKYFEHMGGQRPADIKDFNQYSKTALLKPRQMNSSEDVDISDDMNMFMSVV

>AAO83839.1|Lymnaea_stagnalis_Cav1

MASSPTGVHNDQPTGQHREIGGGGGHNPGGAVNSPSGAGGGGGRGGGGGEDPGGTGGGSGYKPLVPQNTTALATVVGGLSLNTTDSGRALSTGWSTALAAAQGAATVRKRANMRKQQNQNVRPARALFCLTLKNPIRKFCIQVAEYKAFEFLVLITIFANCVALAIYTPYPMSDSNEVNSALDRIEYVFLVIFLLEGILKIIAYGFVMHQGAYLRNGWNALDFTIVVIGIISSILSFVQEKGFDVKALRAFRVLRPLRLVSRAPSLQVVLNSIVRAMVPLLHIALLVIFVIIIYAIIGLELFYGKLHNACYKINSTEFSGDPRICGQGYSCDDIASGGEKGECREGWEGPNYGITNFDNFGLAMLTVFQCITMEGWTTVLYDVNNALGNEWPWIYFISLIIIGSFFVLNLVLGVLSGEFSKEREKAKARGDFQKLREKQQLEEDLRGYLDWITQAEDIDPEVEDENEEGATPRHSKITSDIDSEDKVEEGENVEIQQSWLQQKRLHLQKLNRRYRRFCRRIVKSQAFYWGVIVLVFLNTVVLTSEHYKQPVWLDDFQAIANLFFVVLFTMEMLVKMYSLGFQGYFVSLFNRFDSFVVVCSILEVILQYTNVFPPLGISVLRCARLLRVFKATRYWSSLRNLVASLLNSMRSIASLLLLLFLFIVIFALLGMQLFGGKFNFPQGEKPRSNFDTFWPSLLTVFQILTGEDWNAVMYDGIRAYDGVKFPGILVCLYFVVLFIVGNYILLNVFLAIAVDNLADAQSLTEIEEEKEEEKERTRSLRRSKSRSPNKEQQEVGEEEGLDVNGNDSGEAGEKESNHLSRQPSSRSRRSYRKESTQLNHHVRLDISPDHNMPGEYRNSHKVPLRDDEDEDENNEESDEEEGTEGTETEEDEEGEEEDEEEDEDETPSSARPRRMSELHISSRVKPIPPFSSLFIFSATNKFRIICHKICNHSYFGNVVLACILISSAMLAAEDPLRNESPRNQILNKFDYFFTSVFTIEIIIKIITYGLMLHKGSFCRSLFSILDLVVVAVSLISFPLDNQAISVVKILRVLRVLRPLRAINRAKGLKHVVQCVIVALRTIYNIMLVTFLLNFMFSVMGVQLFKGKFSMCTDESKLTEDECQGNFITYDDGKFNSPNIEERKWKKNDFNFDDVSNGMLTLFTVSTFEGWPGLLYKSIDSHAEGKGPIQNSKPAVAVFYFIFIIVIAFFMMNIFVGFVIVTFQNEGEQEYKNCELDKNQRKCIEFALKVKPIRRYIPKARWQYKIWWFVTSQAFEYGIFVLIMINTVALAMKYHGQSASYSDALDYLNIIFTGVFTVEFVLKLAAFRFKNYFGDAWNVFDFIIVLGSFIDIIYAEVNPGKAFISINFFRLFRVMRLIKLLSRGEGIRTLLWTFIKSFQALPYVALLIVMLFFIYAVIGMQMFGRIKLDYNTQIHPNNNFQTFPHAVLVLFRSATGESWQEIMLACSDKTAVKCDPAVIEHGVNVTDAEANRCGNDFAYVYFISFYILCSFLIINLFVAVIMDNFDYLTRDWSILGPHHLDEFVRLWSEYDPEAKGRIKHLDVVTLLRKISPPLGFGKLCPHRVACKRLVSMNMPLNSDGTVMFNATLFALVRTSLKIKTEGNIDTANEELRTVIKKIWKRTSPKLLDQVVPPAGRDDDVTVGKFYATFLIQDYFRRFKKRKEQMVKISKGQEHTNALQAGLRAVHDLGPEIRRAISGNLDEEEMMDKEVEEPMHRRNHSLFGSVVSAMAGVRPILPFLNRTQSLKGTKGHNTSNLSPIIAVSTAATKMIGMNGNIGNAPSCNHLNLDLSNINHSIQQTHITVPNEDSRRPSKTSETKITIEPPEESFHDADDNIDEDEKFEDVEDGGNKEQEGEKVELDSVMTETSSHPQSVGDGPLPHTTIPNNPNASPSTKKHGIYVYRDLPQEDSDFEREQTPPTPPPRRLSRKGASFKLGCIGKQGSDENPLMRRPGRPLRLTGHHNNQGLTPPTLPDPDGSSQQRPGSQGSSGGRLFSFFRKLGRRPDREPEESPVQHRHMDSSPAPDRSMHLHRSIGRGPLIIPRAVQQQQSLAPRQLENRRSSAETLVAKVLLDEGLNRYVDPQCLQREIAEATNMTQEEMNLAAREIIRRSRHLEPGYYENDPAGILSPDAGPRPRRSPSGQHQQHELLERSRYSTPSNNIDKHPSVN

>XP_011452714.1|Crassostrea_gigas_Cav1

MASPGPGRNNENYNSPSGNTHEGPHIYSKPQPGGGITIRTDSNHSKPLSSAWTTALGAATANVGNTNTMTRKRAPPRKQQNQNVRPPRALFCLTLENPIRKLCIRIVEWKAFEYLILLTIFANCVALAVFQPFPNLDSNEVNLALERVEYVFLVIFTLEAIMKIIAYGFMLHSGAYLRNGWNILDFIIVVIGIITPVFSLFNIHGFDVKALRAFRVLRPLRLVSRAPSLQVVLNAIVRAMVPLLHIALLVIFVIFIYAIIGLELFSGSMHETCFDKQTNSVMTLSDVHPCGKGFSCPENSTCRRYWAGPNDGITNFDNFGLAMLTVFQCITLEGWTDVLYNINDSLGNSWPWTYFISLIIIGSFFVLNLVLGVLSGEFSKEREKAKARGDFQKLREKKQLEEDLRGYLDWITQAEDIDPENEEENEEGATPRHKNQEIPSVKTEDVESGEIQQTWWHRKSRRLRKWNRRCRRMCRKLVKSQAFYWTVIVMVFLNTLVLTSEHHKQPQWLDSFQAIANLFFVILFTLEMLLKMYSLGLQGYFVSLFNRFDSLVVLFSIIEVILIYAKVLPPLGVSVLRCARLLRVFKATRYWSSLRNLVASLLNSMRSIASLLLLLFLFIVICALLGMQLFGGKFNVISNTEDKPRSNFDTFWQSLLTVFQILTGEDWNMVMYDGINSYGGVNSIGLITCLYFVILFICGNYILLNVFLAIAVDNLADAQSLTEDEAEKEDEKERIRSLRRSKTPEGDEKSNEAELEPDEGVGMVENGGEEENVYHSRQPSQRSHRSRQDVGIESEEHRVRINLGHTEDDYKESNHIPIEEEDDTCASDTSSYDLLEGETAVEDEVDEEDMEDEEDEEESQTPSTARPRRMSELHIPEKIKPIPKASSLFILQPSNKFRIICHKICNHPYFGNIVLACILISSGMLAAEDPLQSQSKRNEILNYFDIFFTSVFTVEIIIKVITYGLIVHKGSFCRSFFNILDFTVVGVSIISFVLDNQAISVVKILRVLRVLRPLRAINRAKGLKHVVQCVIVAIRTIYNIMLVTFLLQFMFAVIGVQLFKGRFFSCSDKSKLTESECRGQYIDYPTGDIDEAEIKVREWTNNPLNYDNVPEAMLTLFTVSTFEGWPTLLYKSIDANEENNGPIHNNQPIVAVFYFIFIIVIAFFMMNIFVGFVIVTFQNEGEQEYKNCELDKNQRKCIEFALKTRPTRRYIPKARWQYKIWWFVTSRAFEYGIFTLIILNTVILAMKYDGQSAAYSDALDYLNMIFTGVFTIEFILKLMAFRFRNYFGDPWNVFDFIIVLGSFIDIIYTEVNPGQGIISINFFRLFRVMRLVKLLSRGEGIRTLLWTFIKSFQALPYVALLIVMLFFIYAVIGMQLFGKISTQQDDSQIHRNNNFQTFPQAVLVLFRSATGEAWQDIMLSCVGNEIPCDEMSDADPSQTCGNDVAYFYFISFYMLCSFLIINLFVAVIMDNFDYLTRDWSILGPHHLDEFVRLWSEYDPEAKGRIKHLDVVQLLRKISPPLGFGKLCPHRVACKRLVSMNMPLNSDGTVMFNATLFALVRTSLKIKTEGNIDEANEDLRRVIKTIWKRTSPKLLDQVVPPAGREDDVTVGKFYATFLIQDYFRRFKKRKEQMKKIEKGQEHTNALQAGLRAVHDLGPEIRRAISGNLDEEDFADKDVEEPMHRRNHKLFGFGSVMTALAGVHKTAIPFASRTQSLLINQTPKISSHANLSPQNSLNGKVTPAESCNHLSVEQRNAINRSPSPLAPVRVSPMMGNDSRRHSHTSGISSSESHRDMLPPRSPTSQQDSEPEAPHWDKYKPQDITKKQGLYVYRDLPQEDSDYERDHDHAPPTPPPRKLSRRGASLRLACIGKQQSDENPLMKKIAEPLKLAQTQAMAVAGLTSDGRQRPNEYPRTGLPLSRASYQNTAHLTFSRSNSVIEGHSMSPRQNIHRGSDTMAFDDQTGMYSRTKEAGPLIIPNYVLANQRYRQGSHTKSSAESLVEKVLTEEGLDRYVDAHTLRQEIAEANDMTADDLDKAARMLLHPPDENHTSTTHPGQRSPYYEHLGGFLVQEMKDYNCYSTKDDLNHKGYSQSSSEPSDDEMKFRLTRHAHSFRY

>XP_014774811.1|Octopus_bimaculoides_Cav1

MATIQDAGAAHNEKDTAACSSSSSFATSNTLSRPPRGILSQDRTAVGEQSLSQSLQNFSQLQSQEQGATSSTTGPEAGTDPAKPLSSAWATALAATTVTNMTRKRVNIRKQAQQNIRPARSMFCLTLKNPLRKVCIRIVEWKAFEALILMTIFANCVALAIYTPYPESDTNEINMALENVEYVFLVIFTLECAMKIIAYGFVLHPGAYLRNGWNILDFIIVVIGVISTVLSFLNNDGFDVKALRAFRVLRPLRLVSRAPSLQVVLNSILRAMVPLLHIALLVIFVIIIYAIVGLELFSGKMHKTCFIKNTEHLALEKPHPCGKGYSCHADTEDCRTYWIGPSDGITNFDNFGLAMLTVFQCITMEGWTGVLYNINDAMGNSWPWIYFISLIIIGSFFVLNLVLGVLSGEFSKEREKAKARGDFQKLREKQQIEEDLRGYLDWITQAEDIDPENEEGEEEGTTPRHNKPSDTESSEKNEDLDSGEIQQTWWSVKSRRFQKWNRRCRRLCRRLVKSQAFYWVVIVMVFLNTGVLTSEHYLQPSWLDQFQEVANLFFVVVFTCEMLLKMYSLGFESYFVSLFNRFDSFVVICSIVEVILIHTNLIKPLGVSVLRCARLLRVFKATRYWTSLRNLVASLLNSMRSIASLLLLLFLFIVIFALLGMQLFGGKFNFDETQDKPRSNFDTFWQSLLTVFQILTGEDWNEVMYNGIKAYGGVRNVGVLVCLYFVILFICGNYILLNVFLAIAVDNLADAQSLTEIEQEKEEEKERSRSIRRSKSKSPEAKGGEENNEGEKSNELAEKLDSREPHDDQHIRIDINSDEVAGEYRENTQIPSQEEEIPDEEHATEDEETTGEDDESDAAATARPQRMSEVKMNDKVKPIPKASALFVFSHDNRFRVFCHFVCNHNYFGNFVLACILISSAMLAAEDPLDQSAERSKILNYFDYIFTSVFTIEIIIKLISYGLVLHKGSFCRSYFNLLDLLVVGVSLISINPSNEAISVVKILRVLRVLRPLRAINRAKGLKHVVQCVIVAVRTIGNIMLVTFLLQFMFAVIGVQLFKGTFHMCSDSSKRTMEECQGNYIVYKNGDVNYPELREREWTNNDFNFDDVGKAMLTLFTVSTFEGWPNLLYISIDSHQEGMGPVYDNRPVVAIFYFVYIIVIAFFMVNIFVGFVIVTFQNEGEQEYKNCELDKNQRKCIEFALKVKPARRYIPKARWQYKVWWFVTSQPFEYAVFALIMINTITLAMKYDGENKGYSDVLDYLNMIFTGLFTVEFMLKLAAFRFKNYFGDAWNVFDFIIVLGSFIDIIYTEVNRIDNDDPSQPTLSSGAPGSPIISINFFRLFRVMRLVKLLSRGEGIRTLLWTFIKSFQVLIYFFFLFFFFFFAFSISIKXTFGRIALNESTEIHRNNNFQTFPQAVLVLFRSATGEAWQEVMLSCVDSHDVFCDKLVKTNSTCGTNFAYPYFISFYILCSFLIINLFVAVIMDNFDYLTRDWSILGPHHLDEFVRLWSEYDPEAKGRIKHLDVVTLLRKISPPLGFGKLCPHRVACKRLVSMNMPLNSDGTVMFNATLFALVRTSLKIKTEGNIDQANEELRAVIKKIWKRTSPKLLDQVVPPAGRDDEVTVGKFYATFLIQDYFRRFKKRKEQIQKIQKGQEHTNALQAGLRAVHDLGPEIRRAISGNLEEEDFPERDVEEPMHRRNHSLFGTVVSALTGHKPIPFSNRTQSLHLNHTQPHPKVSPTNSLNAPQSKLSPQNSINGKVSPALSCNHINIDLINSINRSATPLGPVKVAPANEIMPLRKDSHTSKPSSEHSSVEVDSVLDNEQGLGSRPSEVQDSDTESIPMRTVVSDLSQNVQPQSDHAAGVSDPDHGSDQISDKFSDKVLFDHMPDRVPVTHPGPPDKKQDIYVYRDLPQEDSDYEREQTPPSPPPASRKVSKRGASIKLSCIGKQGSDENPLVKKIAQPLKLAQRQAMAVAGVPPGTKYMSTDPRDQSIPAANQKYPYLSPGHRFMFFKKRWKRPRDDEHLRRHPRSGDYPEGTDSAIPYRAYERGPLVIPTHVLSRSKYGFERGSAESLVEKVLEEEGLRKYVDVKYLQQEIKEAGGGMSQEELDRAAHDLIRSSVHGGIHPSYIDQRLGGYKGHEMKDLNQFSRSATPTHRLRRQPHLPHPEFIDDYHDSMDDRMEPCLQIQHALPKK

>XP_023932094.1|Lingula_anatina_Cav1

MSGQDSGLRYDPGESPEPTAARVSTGSPGAQKPHSSPAGKNLLGADAGGVGGHLSSSTPGNGGPPGLGQQQQGGGAQGQQQQQQRTLSSAWATALGAAATAASMNRAKRSNQRGRGGQTNQNPRPPRALFCLTLKNPIRRTCISIIEWKPFEALILLTIFANCVALAIYLPYPKGDSNEVNDALDNVEIVFLIIFAVEAILKIIAYGFLFHQGAYLRNGWNILDFTIVVIGTLSTVLSYMRIKGFDVKALRAFRVLRPLRLVSRAPSLQVVLNSIIMAMVPLLHIAMLVCFVIIIYAIVGLELFSGKMHKTCYHSITDEIMDDPTPCGSSYMCDDGYNCTEKPQPQRWEGPNSGITNFDNIGLAVLTVFQCVTMEGWTTVLYWINDAVGNEWPWMYFISLIILGSFFVLNLVLGVLSGEFSKEREKAKARGDFQKLREKQQLEEDLKGYLDWITQAEDIEEGDEDEEEEEVEPTPTHKANESQSEKTEELGSGEMTQNLSWWHRKSKRLRTWNKRCRRTCRKIVKSQGFYWIVIVMVFFNTCVLTTEHYKQPEWLDEFQFIGNLFFVILFALEMLLKMYSLGFQGYFVSLFNRFDCFVVICSIVEVVLTSTKVMPPLGVSVLRCARLLRVFKVTRYWSSLSNLVASLLNSMRSIASLLLLLFLFIVIFALLGMQLFGGKFNFTDKSKPRSNFDDFFQSLLTVFQILTGEDWNEVMYDGINSYKGVASPGVLVCLYFVILFIVGNYILLNVFLAIAVDNLADAEQLTEMEKEKEEEKERQRSIIRSKSGSPEREKEVQGPCSCCTCCWSRKGDNEEDEGVGSGDNGENEPGDDNQNTANYHDRSSFKHEKSPSSIASEHRVHIELDDDPEKYRGGKPLQGNGGHGRGKCRSXTHKANESQSEKTEELGSGEMTQNLSWWHRKSKRLRTWNKRCRRTCRKIVKSQGFYWIVIVMVFFNTCVLTTEHYKQPEWLDEFQFIGNLFFVILFALEMLLKMYSLGFQGYFVSLFNRFDCFVVICSIVEVVLTSTKVMPPLGVSVLRCARLLRVFKVTRYWSSLSNLVASLLNSMRSIASLLLLLFLFIVIFALLGMQLFGGKFNFTDKSKPRSNFDDFFQSLLTVFQILTGEDWNEVMYDGINSYKGVASPGVLVCLYFVILFIVGNYILLNVFLAIAVDNLADAEQLTEMEKEKEEEKERQRSIIRSKSGSPEREKEVQGPCSCCTCCWSRKGDNEEDEGVGSGDNGENEPGDDNQNTANYHDRSSFKHEKSPSSIASEHRVHIELDDDPEKYRGGKPLQAGDRNPDESGDEVDEDEEDEHHSSNATARPRRMSELHIPDKVKPIPNASSLFIFSPTNKIRIFCHQVCNHSYFTNIVLACILISSAMLAAEDPLNADSERNQILNYFDYFFTSVFTVEIIIKVIAYGFFVHKGSYCRSIFNLLDLLVVSVSLISIFLEKGAFSVVKILRVLRVLRPLRAINRAKGLKHVVQCVIVAIKTIGNIMLVTFLLNFMFAVIGVQLFKGRFYHCTDESKMTEEECQGWFIEYAGDDLSNPSEQKREWVNNPLNYDDVSQGLLTLFTVATFEGWPGLLYTSIDSNEESQGPIYNYRQAVAVFYIIFIIIIAFFMVNIFVGFVIVTFQNEGEQEYKGCELNKNQRKCIEFALKAKPTRRYIPKARIQYKIWWFVTSRAFEYSIFGLILFNVMALAMKYDGMSDGYSQFLSVMNIIFTAAFTAECVLKLIAFKFKNYFGDAWNVFDFIIVLGSFIDIIYTKVNEGEKMISVNFFRLFRVMRLVKLLSRGEGIRTLLWTFIKSFQALPYVALLIVMLFFIYAVIGMQVFGKIALDDDTPMTRNNNFQTFPQAVLVLFRSATGEAWQDVMLGCISTEDVACDPASDSQDKESCGSDFAYVYFISFYILCSFLIINLFVAVIMDNFDYLTRDWSILGPHHLDEFVRLWSEYDPDAKGRIKHLDVVTLLRKISPPLGFGKLCPHRVACKRLVSMNMPLNSDGTVMFNATLFALVRTSLGIYTTGNIDSANEELRSVIKKIWKRTSPKLLDQVVPPAGRDDDVTVGKFYATFLIQDYFRRFKKRKEQLAKMQNLGHEHTSALQAGLRAVHDLGPEIRRAISGNLEEDEFPTQEELPMHRRNHSLFGTVMSAFGSKPRSNSLQVNSTQPHPKVSPTNSLNVTPVPHLTLSPQHSYSNGGPSPQPSPVPSTNHLNVDHHNNVNHSPQPAHDRRSPSSPINLSSTGSLYHVETDPMLGQRDPPLSNLSLEVPHYSNTFIPPVPSRLEAHRGSGSGSAEHIPLQAFGNEERTSSSSPRILPLTRKQGPFVYRDLPQEDSDIEREQHSPPSPPPPRKLSRRGASFKLACIGKQDSDENPLRLVSMNMPLNSDGTVMFNATLFALVRTSLGIYTTGNIDSANEELRSVIKKIWKRTSPKLLDQVVPPAGRDDDVTVGKFYATFLIQDYFRRFKKRKEQLAKMQNLGHEHTSALQAGLRAVHDLGPEIRRAISGNLEEDEFPTQEELPMHRRNHSLFGTVMSAFGSKPRSNSLQVNSTQPHPKVSPTNSLNVTPVPHLTLSPQHSYSNGGPSPQPSPVPSTNHLNVDHHNNVNHSPQPAHDRRSPSSPINLSSTGSLYHVETDPMLGQRDPPLSNLSLEVPHYSNTFIPPVPSRLEAHRGSGSGSAEHIPLQAFGNEERTSSSSPRILPLTRKQGPFVYRDLPQEDSDIEREQHSPPSPPPPRKLSRRGASFKLACIGKQDSDENPLMHRKIAEPLKLTQSQAMAVAGISPEGRSGPLEIPRQSSTPRMQPQPPPPSPGYASHQSSLPTAPSLDPSSSSPSSNREATAAYFQALEEGRLSIPSTQSPQWSPQHKMYGVGGKDIFNKLFVAPSLDPSSSSPSSNREATAAYFQALEEGRLSIPSTQSPQWSPQHKMYGVGGKDIFNKTLEEEGLKHYIDAQCLQREIAEANDMTQAEIDQAARDLLHRSPYYDHLGGYSAQEMRDFNQYSTPEDRGDMADTAEKKKDQTNYITTV

>XP_022081221.1|Acanthaster_planci_Cav1

MDKFRTVVHNNVAAATASSGTYVSGSASTPTPTTASPTGLPATTNAPLSSAWRTTLAATTTVSTMNRRRNTYNKKKQHSGTSLRPPRALFCLTLDNPVRRMCISIVEWKPFEYLILLTIFANCFALAIYTPFPHEDTNTTNKNLENVEYIFLFIFTLEAMLKIVAMGFLFHSGAYLRNAWNFLDFIIVIIGVVSTILSHTAQQIAGFDVKALRAFRVLRPLRLVSGVPSLQVVLNSIVRAMVPLLHIALLVIFVILIYAVIGLELFIGKLHRTCWIVEDGVRRHVEEEPHPCGDRGFNCSELREDAFCDEYWEGPNEGITNFDNIGQAMLTVFQCITMEGWTDVLYNVNYAIESWWPWFYFVTLILLGSFFVLNLILGVLSGEFSKEREKAKARGAFQKFREKKQIEEDLKGYLDWIMQAEDIDPENEIEQHGEPPKHVLWKRLGTLNTKVPKPMSESDSSEKSEELGSTDIPQQSWMQRKKRQMRRWNRRCRRLCRQAVKSQAFYWVVIIMVFLNTIILASEHYRQPKWLMDFQDIGNLLFVVIFTVEMLIKMYSLGLQGYFVSLFNRFDCFVVCSSMVEVVLMYAGVIQPIGISVLRCVRLLRVFKVTRYWASLRNLVASLLNSMRSIASLLLLLFLFILIFALLGMQVFGGRFNFNKTQDKPRSNFDNFWQSLFTVFQILTGEDWNEVMYDGIAAYGGVHSIGIIASSYFIILYIWGNYILLNVFLAIAVDNLADAESLTALEKEREEEKRNKSIRRANEVDLSKQLPPGVDPKGVIRGPKRLQEKRKALNKEAPSSNNHKAIEDGEKQVHIEEDEAKKALKEPGDEDEDEEEEETLTTQSARPRRLSELDLPNKKAPMPKESSLFVLGPENRFRKGCYFVCTHNYFSNVVLLLILISSIMLAAEDPIDKNKTLNFILNCFDYGFTVAFTIEILLKVISFGLVIHKGAFCRNFFNLLDLLVVTVSYISIALQGQGAISAVKILRVLRVLRPLRAINRAKGLKHVVQCVFVAIKTIGNIMMVMLLLVFMFACIGVQLFSGKFYSCTDLSKMTEEHCHGNFIEYKDGNYLEPVVKPRVWKLNEFSFNNVGSGMLALFTICTFEGWPQLLYVAIDAASEPDHGPIRNNQLGVAIFFFIYIIVVAFFMVNIFVGFVIVTFQNEGEQEFKNCELDKNQRQCLEFALKARPKKKYIPKNSKQLKVWKIVTSRPFEYLIFVLIMVNTIVLAMKYYDQSDEYSEVLDRVNIVFTAIFLLECILKIIAFKIKNYVRDLWNLFDFVIVVGSIIDIILSEGKASQMDSESRFSINFFRLFRVMRLVKLLSKGEGIRTLLWTFIKSFQALPYVALLIVMLFFVYAVIGMQLFGKIALTTDGPINRNNNFQSFVAALLVLFRSATGEAWQQIMLACASSPEAKCDPFVLEHQPEVGETCGNDFAYIYFLTFYSFCSFLVINLFVAVIMDNFDYLTRDWSILGPHHLDEFVRNWAEYDPEATGKIKHLDVVALLRQISPPLGFGKLCPYRIACKRLVSMNMPLNSDGTVMFNATLFALIRTSLKIKTEGNIDKANDELRQVIRKIWKRTSTKMLDQVVPPAGADDDVTVGKFYATFLIQDYFRRFKKRKHDQLKMQGHEQGTVALQAGLRTLQEIGPQIKRAISGNLEDMDDEVSEAVELDEPSHRRSHSLFGNMLNMYHNRRQSAPMANTHPHHSIPLTNNLTVSPSSQHRKLSPANSLNSYSYTPRSRSPYSMSSGGGSFRSSRPPAGNALHVSSPNSAGFSPVGTQRYSPNNVAGRNHRHSSGQSHEATQPLLSEEDHHLDPPYRNGSVHADDQNYGTLGQGRQEMEHNAGDSREPFQDPYREPYRDYSRIPKSILRITRRLREPPPTLTTECYDDEDDDTSSENSHPATSPLLSHEHTPSPSRSCSSGSRSVSASPQTSSLTDFPTATGTSSSRYPGDESSLPGGECGGDGGNSYPAMAGRKVPPLSSARTSGSSLLSCVGKAEPRPASERKTAVPLKLAQAQVMAVAGMTPEGKPRDDGRPSLWLTPPSSPRRVRSSYHTPTPPRGRASGRGGGDSLLPTDNNNASAKEPLALRHSTGEIGTSRDPSTAFFKALTKPRVRQPLIPKGPKAAASQMPENVEGSAESLVAQVLEEEGISPIHDRDLISTVERELAEACNMSQSEMDAAAHRLIQANQQGDTSLPYFDHLGGYELQDYTQRSRSRKRGHSREGGFRTNSKGTSQEEDDLESENTDSENDGEDDHDDDDDDAKPKDSSGDMVYVTTL

>XP_011670134.1|Strongylocentrotus_purpuratus_Cav1

MQRQRTGPVNNAASAGTGSSGTYVTGTVPLSRPGPTTRPPAAAGSTTTPAVAAATTTTTAPPSNNAAAAPDPPGPDQDGAVAKQPLSNAWVQALAAANTANSMKYGKKKQHIGAPTRPPRSLFCLTLDNPMRRMCISIVEWKPFEYLILITIFANCIALATYTPFPKQDSNDVNRNLEYVEYAFLIIFLIEALLKICAQGFLFHPGAYLRNGWNILDFLIVAVGVISTILSLRNVENTNFDVKALRAFRVFRPLRLVSGVPSLQVVLNSIFRAMVPLLHIALLVIFVIIIYAVIGLELFMEYMHKTCYFKDTSIIAMDDPHPCGNGFRCTDLLEVGINNTDCLEKWEGPNDGITTFDNIGLAMLTVFQCITMEGWTDIMYDINDGAGPWWPFLYFVSLIIIGSFFVLNLVLGVLSGEFSKEREKAKARGAFQKLREKQQIEEDLRGYLDWITQAEDIDPDNDSEQNPGPPKHVPKPISESDSDDKSEEMGSSEIPQQQWMQKERRQLRRWNRRCRRLCRQAVKSQAFYWIVIIMVFFNTVILASEHYSQPAWLTDFQDFGNLCFVVIFTIEMIIKMYSLGLQGYFVSLFNRFDCFVVCSSIIEVVFIYAHIIPPIGISVLRCVRLLRVFKATRYWTALRNLVASLLNSMRSIASLLLLLFLFILIFALLGMQVFGGHFNFDSTKLKPRSNFDSFFQSLFTVFQILTGEDWNEVMYDGIQAYGGVKSIGFLASTYFIILYICGNYILLNVFLAIAVDNLADAESLTALEKEKEEEKRNKSIRRANEFDLSKQLPPGVDPKGVIRGPKRLQEKRKAAFGKESDGSLNHKSKSQDSEENVNVIDEDDVKKSLKEPDEDDKEEEEAESLTTASARPRRLSNINIPTKTRPIPKANSLFVFSNTNRFRRLCYNLVNHPYFTNVVLVLILISSTMLAAEDPLDEDKRRNYILSLFDYGFTSIFTIEILLKVVAYGLVFHQGAFCRNSFNLLDLLVVTVAYISIIFDDTKISAVKTLRVLRVLRPLRAINRAKGLKHVVQCVFVAIKTIGNIMLVTLLLVFMFACIGVQLFRGRFFSCNDSSKLYEEDCQGHYFVYKNGDLDQVSVEQRVWSKNEFHFNDVGNAMLALFTVATFEGWPKLLYVAVDSTEDDKGPVHSSRMPVAVFFFAYIIVIAFFMVNIFVGFVIVTFQNEGEQEYKNCELDKNQRNCLAFALKAKPVRKYIPKNPKQHLVWKLVTSRAFEYFIFVLIMVNTIILAMKYRTQTEAYKNVLDYMNIVFTAVFTVEFLLKIIAYKPKNYFRDYWNAFDFIIVLGSIIDIMIDMFSRVVTAANEKKQFSINFFRLFRVMRLIKLLSRGEGIRTLLWTFIKSFQALPYVALLIVMLFFVYAVIGMQMFGKIKLSLDGALNRNNNFRTFPTAVMVLFRSATGEAWQQIMMACSQSPKAPCQRDDTEICGNNFAYVYFISFYSICSFLIINLFVAVIMDNFDYLTRDWSILGPHHLDEFVRQWSEFDPDATGRIKHLDVVTLLRSISPPLGFGKLCPHRIACKRLVTMNMPLNSDGTVMFNATLFALIRTSLKIKTEGNIDQCNEELRAVIKKIWKRTSTKLLDQVAPPAGADDDVTVGKFYATFLIQDYFRRFKKRKQEGLKIPGQPDSTFALQAGLRTLHEIGPQIKRAISGNLEGMDENEQFEHDEPNHRRSHSLFGTVLNMYHNRRQSTPMANTHYQGRDPPPAYYPNNATNNLVSSPPSNQHRKLSPANSLNNYNHNPQNKSPFNMSSSKSPSAIRGRGGPYSPNANSLQVSSPHTRGFSPVGNRNPGGRGMPNNRGPGNRYSTKEGRQPLLQNEDRHHHHHHNNNHHHSPDSPYDDDDVEEIEPKAPSSPQVVDEEYQPEPYQQPYREPYRDYSSLYTVYEEPPTLRTECYQEEDEDNLSETSSHPPTSPLLSPQRSPSSSRGHSPTPPPHQNSSNHPSSPSPTKLTLDFSTTVPSPTETSLPKLEGESIGSRSVTHSPRKSPAGSPSRRLLLPSGKSDGLVPAGDRKTAVPLKLAQAQVMAVAGMTPEGKTRNSTGRMWLTPPSSPRRIRSSLHTPAPRPTTQSSHLLPTTGSTSNSTSTHMMVKEPLAQRHSTGEIGTSRDPSAAFFKSLTKPKTSVPRNNPSPEEVAGSAESLVAQVLAGEGISPIRDRDLINMAERELAEACNMTQAQLDSAAHRILFPKDKLANVPVVTTTVTQDTQLPYFDHLGGYELRDYTRRSGGHDRTNDDDDEEGGRRAGREADSDTESGTIEADEHGVLYVTTL

>evg1131374+ TRINITY_DN40077_c0_g1_i4|Nematostella_vectensis_Cav1

MDSNVYSREQPSAPMKSSWPTAAELAQNRAKLNGQPKYGKPPPQPKRQKKPTNAVRPKRALFCLTLGNPIRSTAISLVEWRPFDVMILITIFANCAALAAFQPLPEQDSSLINEELEVAEFVFLGIFTMESVLKIIAYGFVMHPGAYLRNGWNILDFVIVVVGLATIIVKLYTPDSFDVKALRAFRVLRPLRLVSGVPSLQVVLNSIIKALIPLFHIALLVVFVVIIYAIIGVELFMGKLHSTCYDNVTGQPTFDESHPCSTESEGYSCSNAGPGQVCLKKWEGPNYGITNFDNIGLACLTVFQCITLEGWTDVMYSINDAIGNSWPWLYFVTLIIWGSFFVLNLVLGVLSGEFAKEKARAQKSGEFQKLREKQLVDDAYHGYLDWISQAEDIEGDSSAGEDEEGKADRKPSFRRRKENDDISKNKENQEDSAASDQGWIDRKKKILKRFHHRLRRSCRKAVKTQWFYWTVIVFVFLNSLTLALEHYNQPEFLTQFLDKANKLFLALFTLEMVVKMYCLGFHGYFASLFNRFDCLVVISSLLELGLTEAMDQRPIGISMLRCVRLLRIFKVTRYWSSLSNLVASLLNSMRSIMGLLLLLSLFMVIFSLLGMQIFGGKFNLGDEDVPRSNFDSFWRALVTVFQILTGEDWNAVMYTGIQSWGGITESLSAIPILYFIFLVVVGNYILLNVFLAIAVDNLADAESLTEMEEEKKKEREEELEMKLRMEEKESSSNKDLENKRASTSSGAGKTTSQDLHSNGNSIPRALSSDAESPSLAEDSKATLNREGDHESVRSATSTEVMDHEPMPPESSLFIFSNTNCFRVVCHRIATNSYFVNFILLLIIVSSCMLAAEDPLNSNSKRNQVLNYFDYFFTAVFTIEITIKIIAYGVILHKGSFCRSAFNLLDFLVVAVSIVSIALRDSSSQISVVRILRVLRVLRPLRAINRAKGLKHVVQCVFVAVKTIGNIMLVTVLFNFLFAVIGVQLFKGTFFSCTDAEKITKRECQGQYIEFKGPGLTNPVVKDREWQPQTFNFNDVPQAMLTLFTVMTFEGWPGILESSMDSTDVDEGPFLNNRPWVAIYYVIYIIIIAFFMINIFVGFVIVTFQNEGEEEFKDCELDKNQRKCVEFALKARPTRRYIPTNRLQFHVWRVVTSQPFEYLIFAFITGNTILLMMQYYNEPKLYTRVLDGFNIGFTSVFLLECILKLFAFKPKNYFLDRWNLFDFVVVVGSVVDITMNEVSSEQMFAFGFFRLFRALRLVKLLNQGSGIKTLLWTFIKSFQALPYVGLLIIMTFFIYAVVGMQMFGRIAIDPETQINRNNNFQTFPQSLMVLFRSATGENWQLIMLACTDTPNAKCDPNAYPQDTDGLCGTDFAYAYFCSFYAICSFLIINLFVAVIMDNFDYLTRDWSILGPHHLDEFVRVWSEYDPEASGCVKHVDIVTVLKRIAPPLGFGKFCPHREACKRLVSMNMPLNRDGTVNFNATLFGLVRTSLSIKRPEGKGSLEKANEEMRAIIMKVWPKTSQEFLDEIIQPPGGDDEVTVGKFYATYLIQEYFRRFKARQRAENAQYQDAHSTQALQAGLRTLHGLGPQLRRAISGHLDDDDEELFLKEDIQQKSQLDFLSEQHKRFWESLKNAVTPSPRHSRPSSLRRPGSIRLSAFIKKGNSTPEAQRKSSLPATLTVPSFSHGPLLTSDRQLLVPPQQSDVNANDKESKQGAHVPSKPPLKGARSFPLPIMANGDLKEPALDTRERSKSEILERPASGEAVLEPTELDKEKSSSAQNIVEQALQDEELDEDDTIVRVVEQEIAEAFDLSTDQLNDAAERLLSELEETLKNEEEVEKEEARLEAEAQADKSEEGKDNFWKKGLQPSPDSCIVITDL

>XP_020903719.1|Exaiptasia_pallida_Cav1

MATSKPTWPVPDTSQTRTKLNGQPKYGKPAPQTKRPKKPPIAVRPKRAVFCLTLGNPIRSAAISLVEWKP

FDVMILITIFANCAALAAYQPLPEQDSSSVNEELEVAEYVFLAVFTLEALLKIIAYGFVMHPGAYLRNGW

NILDFVIVVVGLATILVKALNLESFDVKALRAFRVLRPLRLVSGVPSLQVVLNSIIKALIPLFHIALLVV

FVVIIYAIIGVELFMGKLHKTCYDNVTGQMAFDEAHPCSTDGEGYACSAADGQVCLAKWDGPNFGITNFD

NIGLACLTVFQCITLEGWTDVMYSINDAVGNSWPWLYFVTLIIWGSFFVLNLILGVLSGEFAKEKSRAQK

SGEFQKLREKQLIDDAYHGYLEWIAQAEDIEDEEGADQDQEGVPGRKLSTKRRKEDGELAENEENVESST

ALDQGGGWLEHKKKVIKRLHHRLRRFLRKAVKTQAFYWTVIVVVFLNSLTLALEHYNQPEFLTQFLDKAN

KLFLALFTLEMLVKMYCLGFHVYFASLFNRFDCLVVVSSLLELALTEALDQRPIGISVLRCVRLLRIFKV

TRYWSSLSNLVASLLNSMRSIAGLLLLLSLFMLIFSLLGMQIFGGKFNIDDTEVPRSNFDSFWRAIVTVF

QILTGEDWNAIMYIGILSWGGITDSSSVIPILYFIFLVIVGNYILLNVFLAIAVDNLADAESLTEMENEK

KKKEEEKKELESSRSSLSESPTKGALSKASQELHSNGNGIPRAISEGDVESQSHEKGSKITLDRSTHSVR

STTSTDENLDREPMPLESSMFIFSSTNCFRILCHKFVTNIYFVNFILILIIVSSALLAAEDPLNANSKRN

QILNYFDYFFTTAFTIEITVKHRFNIDDTEVPRSNFDSFWRAIVTVFQILTGEDWNAIMYIGILSWGGIT

DSSSVIPILYFIFLVIVGNYILLNVFLAIAVDNLADAESLTEMENEKKKKEEEKKELESSRSSLSESPTK

GALSKASQELHSNGNGIPRAISEGDVESQSHEKGSKITLDRSTHSVRSTTSTDENLDREPMPLESSMFIF

SSTNCFRILCHKFVTNIYFVNFILILIIVSSALLAAEDPLNANSKRNQILNYFDYFFTTAFTIEITVKII

AYGVFLHKGSFCRSAFNLLDALVVAVSIISIALRGSKSSQISTVRILRVLRVLRPLRAINRAKGLKHVVQ

CVFVAVKTIGNIMIVTVLFNFLFAVIGVQLFKGTFFHCTDGGKITQEECQGQYLEFKGPGLSNPVTQQRE

WKHRDFNFDNVLNAMLTLFTVMTFEGWPGILENSMDSTDVDQGPFLNNRPWVAVYYVIYIIIIAFFMVNI

FVGFVIVTFQNEGEAEYEDCELDKNQGTFFHCTDGGKITQEECQGQYLEFKGPGLSNPVTQQREWKHRDF

NFDNVLNAMLTLFTVMTFEGWPGILENSMDSTDVDQGPFLNNRPWVAVYYVIYIIIIAFFMVNIFVGFVI

VTFQNEGEAEYEDCELDKNQHKRFWESLKNAVTPSPRHSRPNSFRRPGSVRLSAFIKKSNNNTPETTRKI

SLPATLSVQKAPRERQVLVPLHSDDSDSEDKRLFVPEHQPLKGARSFPLTVTANGNVNTDTSLETRERSK

SEILSPRTPSQELQQDNLNREKSSSAYEAVGRALVEEGLDDDDTMVRVVEQEIAEAFDVSTERLNEAAER

LLNELEATLSSDEEGEKADSETSVLKSDDDSDSDXPSHFWREGLKSMPDNCIVITDL

>XP_015778662.1|Acropora_digitifera_Cav1

MEQNGYPRANFTSATKSLWPNGTDLSQYKSRLNGHATKYTKPGASAKRQKKSGNAVRPKRALLCLSLGNP

IRSAAINLVEWKPFDVMILITIFANCAALAAYEPLPGRDSSEVNEGLEIAEYVFLAIFTLEAILKIIAYG

FFFHSGAYLRNGWNILDFVIVVVGXVNLVTIWELSVLYILGTEVGVYCSVLYGFMWAEFHTVLCLFQGAE

AMENPHPCSSGGSGFHCNASEAQVCEAGWKGPNYGITNFDNIALACMTVFQCITLEGWTDVLYMINDAVG

NSWPWIYFVTLIIWGSFFVLNLVLGVLSGEFAKEKARAQKSGEFQKFREKQQVEDAYNGYLDWITQAEDI

EGDSESETGDESKSSRRALLANCILKLNLNEWFGDFEEKTRHSRIDDIEMIDKNERQEIAVQEAHHGWCH

NEKYWSSLSNLVASLLNSMRSIAGLLLLLSLFMLICSLLGMQIFGGSHCIPLPCKDILLNVFLAIAVDNL

ADAENLTEMEEEKKKKKEKAKEKLRASTESQTKIGQDGAIVPHHSSATHSNMTLDKSNQELHSAGNLNGN

AVAQTASHSDIEAQSVEHLEPEDSKSAVNNNEESAAVGSTEDIDYTPMPPESALFIFSSTNITVLCLFQG

AEAMENPHPCSSGGSGFHCNASEAQVCEAGWKGPNYGITNFDNIALACMTVFQCITLEGWTDVLYMINDA

VGNSWPWIYFVTLIIWGSFFVLNLVLGVLSGEFAKEKARAQKSGEFQKFREKQQVEDAYNGYLDWITQAE

DIEGDSESETGDESKSSRRASRHSRIDDIEMIDKNERQEITVQEAHHGWCHNEKKVLKRWHHRTRRELRK

AVKTQAFYWIVIVVVFLNSLTLALEHYGQPHFLTIFLGEASCQASVFREVTKHNLLGKHSLPVSTCPLRD

LFLVLFLVTCCILCFQVVVSSLLELAIVEAMSQRPIGISVLRCIRLLRIFKVTRYWSSLSNLVASLLNSM

RSIAGLLLLLSLFMLICSLLGMQIFGGRFSMDGEDVPRSNFDSFWKALITVFQILTGEDWNTVMYDGIRS

WGGIGEGGAILAILYFIFLVVVGNYILLNVFLAIAVDNLADAENLTEMEEEKKRRKXKAKEKLRASTESQ

TKIGQDGAIVPHHSSATHSNMTLDKSNQELHSAGNLNGNAVAQTASHSDIEAQSVEHLEPEDSKSAVNNN

EESAAVGSTEDIDYTPMPPESALFIFSSTNIQSNSPVLNYFDYFFTSVFTLEILIKFVAYGLILHKGSFC

RSAFNLLDLLVVSVSVISISLKNSQFSVVRILRVLRVLRPLRAINRAKGLKHVVQCVFVAVKTIWNIMLV

TMLFNFLFAVIGVQLWKGTFFYCTDQKKRFEDECKGNYFEYNGAGLSNPVAKKREWKRRDFNFDNVGNAM

LTLFTVMTFEGWPGILYNSIDSTEVDEGPLQNNRPWVAVYYIIYIIIIAFFMVNIFVGFVIVTFQSEGEE

EFKDCELDKNQVSKRSFNYFIDRWNLFDFIIVVGSIIDITMNEVSSEQMFAFGFFRLFRALRLVKLLNQG

SGIKTLLWTFIKSFQALPYVALLIVMMFFIYAVIGMQMFGRIALDPETAINRNNNFQTFPHSLMVLFRSA

TGENWQEIMLSCTNREDVKCDPNADPKDPSGLCGSDFAYFYFVSFYSICSFLIINLFVAVIMDNFDYLTR

DWSILGPHHLDEYVRVWSEYDPEARGCIKHVLLKRIAPPLGFGKFCPHREACKRLVTMNMPLTKDG

MVDFNATLFGLVRSSLNIKKPEGKGSIDKANGEVRNIILRIWPKTSMQLLDKVVQPPGVRKYRRWNQSGY

ALTPALCVKSSLSKTFGRLVAEWLADPVHLTYTHRFKFPSDYYMSVLRVRTCVKRLVCVCCFFLQHKGFW

ESLKNAVSVSPRHSFRRPNSFRLSTFLGKNESATEKKKKSSSMSNLGENRNANLVNNERFLAPSTSADEN

GNENESEAETSLMEPPPKGPEKEASELFKRDHATIKAASSLPLTGDNRKIFGPITYLRQRSRSETLPNRS

SSEDEIHRLSHEDRSESARSLIEEAMLDEGISITLDDPLLRIAEQEIAEAFDVSEDDLHSAAERMLADED

DRDSADVDSARHSPSVPFRLSGGPDSELVITDL

>AAC63050.1|Cyanea_capillata_Cav1

MFDVQVKQSDGETERQGKWESRPSEDDDDSIPFENSSPQYDWDDNDLTSGKDQDDEKNLALLATQKMATS

TYPTKKTKKQPGAGQNLRPKRALFCLTLDNPVRSAAITIVDWKPFDFFILASIFANCAALAAYEPLPAGD

MTSTNQDLEVAEYFFLAVFAIEGLLKIIAYGFILHPGAYLRNGWNILDFSIVVIGFASMIFEEYLKSGFD

VKALRAFRVLRPLRLVSGVPSLQVVMNSIVKAMLPLFHIALLVVFVIIIYAIIGVELFTGKLHQTCYDNI

TNLPASSEPKPCSTTTSYGRQCPSGSICKNNWEGPNYGITNFDNVALAALTVFQCTTLEGWTDVLYDINN

VSGGGWPWIYFVTLITFGSFFVLNLILGVLSGEFAKEKARQTKSGEFHKIREKHMLDDAVKGYLDWINQA

SDIENVTVTQEGAVEPGDSTSERKLSSRRASGISHASSGIYNIAAPNPLTTKERIEKNLTKFHHRLRRQC

RGIVKSQTFYWMVIIAVFLNSLVLAVEHYDQPDYITMFLDRANYFFLGLFTFEMLLKIYCLGIYGYLNSL

FNRFDCLVVLSSLLEVAITVPTGWPPIGISVLRCVRLLRIFKVTRYWESLSNLVQSLVNSIKSIGSLLLL

LSLFILIFSLLGMQIFGGRFNLDEQAPPRTNFDSFWRSLITVFQILTGEDWNAVMYVGIQSWGGIKNPSS

IIAIIYFVALVIVGNYILLNVFLAIAVDNLADAENMTKVNEEEKRKKKDAKLMKKLARKKMDSLDSAEGK

KETTESITDGSKDPLSGSEGPEDVELGNPKSKNGTLRHMGETTSTEMSEGKEARIRPLRLSELNLLKDIP

DPMPPESSFFIFSANNKLRYLCYRLAVNKIFINSILVLIIMSSVALAAEDPIGRDVLRNKILGYFDIFFT

AMFTFEVTVKMIAFGVILHKRSFCRSFFNQLDLVIVAVSWAAIMLSRGSATSVVRILRVLRVLRPLRAIN

RAKGLKHVVQCVFLAIKSIGNIMIVTLLFQFLFAVIGIQLFKGTFFYCTDRSKMTAEECKGTFNSYPEVN

LANPVVAARTWEKHTFRFDNVFQAYLSLFVVMTFEGWPSILEHSIDSTTVNQGPKFNNRPFVAIYYVIYI

IIIAFFMINIFVGFVIVTFQNEGEEEFADCELDKNQRKCVEYVLTVKPTYRFVPRKRFQYHIWRVVTSRL

FEYMIFGFILGNTIVLAAQYHNASKLYERVLDGFNIGFTAVFLLECVLKLMAFNAKNYFRDPWNIFDFVI

VVGSIADIIIGEISKDGGIKVNFFRLFRALRLVKLLSQGDGIRTLLWTFMKSFQALPFVGLLILLLFFIY

AVIGMQVFGTIRLDSGTVINSNNNFQTFPQALIVLFRSATGENWQQIMMACVNSESVKCEVDPSKTCGTD

FAYLYFMSFYMICSFLIINLFVAVIMDNFDYLTRDWSILGAHHLEEYVRIWAEYDPEASGRMKHVDIVSM

LKRIEPPLGFGKCCPHREACKRLVSMNMMMNNDGTVDFHATLFALVRTSLNIKKPDATETILHANNELRG

ILKHLWPRTNENLFDKLIPPPDYAEGITVGKFYATFLIQEYFRKFKKKRQEERKKKNADSTVALKAGLRT

LQELGPKIKRAISGDLMEEGTKEEEEKEKEPNKRHSFMAGLRKGIISSFAGDKKRNSYYNALQDTPKQTP

KQTPTSSVSHLAAPNTQGKRISLPSLQTKTNSSQDNIPRGALHEGPSIMQRLSPRLKRKGQRPKSTPFEI

IHHDSDDLALPQIVLSSQSDVSNPSSKDRDNSRSSTPQHASSIPGADLNSNLLDSNLGPRRSRSSSTSSL

VGQALAHEGVPQDEAILRTVQKELSDSLNISESQLEGAARNIIYDFEKESGTSIPPPEAAPPQEQPNAEQ

LSPYNDNNDNRSSRPDRTSYL

>evg1032956|Trichoplax_adhaerens_Cav1

MADDKVGTDDGKSTLDLSDIRDRLPPSVAGTLLETAAIARAKRRGAIQGRIQHYKAKQGALTPRAARVLFFLKTDNPIRQFATTVVEKKAFEYLILFTIFANCVALALYQPLPNNDNTLLNENMEKVEYVFLAIFTIESFLKIITYGFAIPSGAYLRNGWNILDFIIVIVGIINIIFTATSSSNQNIDVLRALRAFRVLRPLRLVSGVPSLQVVMNAIMKAMVPLFHVAALVVFVIIIYATIGLELFNGVLHRACYHNITKKLINDPRPCAASESFESAHHCKSGYVCLRNWTGPNAGITSFDNIGLSMLTVFQCITMEGWTNIMYSINDAVGSEWPWIYFVTLIILGSFFVLNLVLGVLSGEFSKEREKARVSGDFKKLREKRQIEEDYRGYLEWIGKAEDLEVDVDDLQVDMEQQVAAEGVRFDDVEHGSQGSAVEEKKVYCFRFRHQLKRWHKRSRRYCRLVVKSQTFYWLVILAVLLNTICLAVEHYEQQRVVTQFLSITNSVFVGLFTIEMSIKMYALGIEGYFMSLFNRFDFLVVLVSIIELIFIAIIGGAAALGLSVLRCVRLLRIFKITRYWNTLRNLVASLLNSMRSIASLLLLLFLFVLIFALLGMQIFGGRFNFNKSIPRSNFDSFWQSLLTVFQILTGEDWNEIMYNGIKALDGIDRFGILAVFYFIILVVVGNYILLNVFLAIAVDNLANAESLTEINERRNAKRKMAKEQRLQKLQTPVQSLRYGNISIGSAVENVSSEENPDDIQKLSKVMYEGDLENDFSNPVYKMAVFSETESELSRVEELDSNSDCTEEPARPRLDASFRTNTTQINWKKIPDRVIVPIPKATSMFLFKSTNRFRCKIFDFVTNLYFSNIILIIIILSSITLAAEDPLGKDKIRNQVLSYCDKTFTAIFCVEAAMKMIAFGVIMHEGSYFRNIFNILDMIVIAVSIADYTIAEENLKQLKVLRVLRVLRPLRALNRARGLKHVVQCVFVAIKTIWSIMLVTLLLVFMFAVIGVQLFKGRFYFCTDASKMDNSTCKGSYYNYPTGDLNYLQVEPRIWKESSFNFDNVPNAMMTLFTITTFEGWPSILYRAIDATEAGRGPSRNHQPLVAIYFVIYIIIVAFFMVNIFVGFVIVTFQTEGEQEYRNCDLDKNQRNCIEFALKTKPQKRYIPKNKLQLAVWKLATSLGFEYTIFGLITLNTMTLMMQVYRPSRVYNDVLEYLNIGFTVLFGLEAILKIVAFKPQNYFRDKWNVFDFVIVVGSIIDIVISEIYKDSSNVTVDFSVNFFRLFRAMRLVKLLSRGGGMRTLLWTFMKSFQALPYVGLLIVFVFFIYAVIGMQLFGTVLTSPGSAITEYNNFHSFFSSVLVLFRCATGENWQLITLSCTWRTGNGCSENSCGSNISYLYFSSFYVLSSFLVINLFVAVIMDNFDYLTRDISILGPHHLDEFVQAWSFFDPEATGRIKFDDMVKLLRRISPPLGLGKLCPDSTAYRNLMGMHMPLGTDNNVKFNATIFALVRKQLKIKLDGNLDLRDRELRNIIKRIWPRTPNKLLDEIVPLSLKDNDITIAKIYALLIIRNFIRRRRERQAKQRREQTSNEIQVGIRVLQDLGQTNLRRASGELRSDDDDEEEDIKTHAKSQFTLRTFNQSLYRFGSSLKINSSSDLASQKSSSIDSIKNKYCKAPIVYSSQNTHFSEAETPNSVDNRYEVSTSKDMRHETSQQNKRLELQTFTDSKEGTSRLNEGLRNADIPDSGYTKDKDGGIVNKLYGIDITNTGSSGNGSGVIDSGSMPYGPYGDTLKTKVSDRINRSTLTGKIHYRKSSFERVSVV

>m.84361+m.186460+g4118.t1(Compagen)|Oscarella_carmela_Cav1

MGLAMLTVFQCMTTEGWSQILYWTQDAVGSKWPWVYFITLLSIGSFFVLNLVLGVLSCEFSRERDRAQARGDFKKLRDRQKVANDVKQFLDWIAEGERGERPQDVVTSTKGEEIQASDDTTVDCKMTSRWKCNFSHNRFYKYLQRLNARFRPLVRRLVKSRIFYWLIIALVSANTISLAAHHYGEGSFLRSFDEKSYVVFLAIFVLEMMLKLYGLGVQGYFMSNFNRFDFCIVVTCCMEFFATVFGGVRKIGISVLRCIRLLRVFKFVPSYWVSLKNLVASLLNSIRSIISLLFLLFLFILIFALLGMQIFGGRFNFDYGVPRPNFDSFVSSLLTVFQILTGEDWNEVMYLGVEAYGGIGGAGAVAVLYFVVLVVFGNYILLNVFLAIAVDNLADAQLLTKEEELHWMERKRSREETRKIIEEYSEQNKASKCLFTDSINQSCSINRTSSMDAKSADSMSASKASGLDGAQPDEIQSIGTTPTKVDEEPNTESSSYIYMRSFLRMSQAEAGVKHVAKANGRRSMPDTRSCICISPRNRLRRLCHSVVCHRYFDTVSLTLILICCVAMAVEDPLLDAENSTKNQVLQYFDYFFTFLFSLEMLAKILVFGFVCGKESYLRSFYNILDFCVVLLSVSSFILKTFFGGVDLEVIKVLRVFRVLRPLRAINRTSRLKTAVRCMIASVRAMGKILLVTGLLQFAFAVIGMQLFKGTFFYCSDPSKMIQEECQGRFFTYPDGDLQRPMVANREWRKRDFNFDNVLEATKTLFPVATFEGWPTFLYWGIDSNKEDHGPIRDSRPAVALFYVIYIVVISFFMVNIFVGFVIVSFQRVGEEEFKDSELNKNQRACLEVALTAKPSRIYRPSHRTLFALWVVATSKKFELFTLVIIGINVLVMTLKFHGQSEFYGITLEYINIVFTSLYTVECILKLVAFTPKHYFRDRWNVFDFLIVLGSLIDTALLYSLSGDSYEGVNLNFFRVFRAFRLVKLLRKQKGIKTLFWTFFKSLKTLPYIGIIIALIFFIYAIVGMQVFGRIKLDSSTEIFRHNNFQTFFQALLVLFRAATGENWQKIMTACATNSKCDPLSDSPGSCGNGFSYFYFVSFIVLCSFLIINLFLAVILDNFDYLTRDRSILGPQDLDNFIDVWKTFDPEGSGRIHHEQVVPLLQKLEPPLGLGTCCPNRLAYRRMISMNMPMNEDETVNFNGTLFALVRTALKIKSGVNRAESDAELRETLKHFWPKICSKNVVNEVIPLDDDNNMKATVGKFYAVMLIQSCFQQFKARKRRQSQTKEIEDYNSNYLSIKTEEADDSQKLCNLVAWRRRMSRDSSIFSDDGQNCLGGAEQLSSAWCAGGTAEDRRQRWQGFVQEKRKVNSLILVAECRDDESDDDSNSNSNTEFSSEETVVQQEFSHQRTHPITISITSAGDIEREINPMSSVRQSEQSASFQLKAEEVGRHRCDSWSSEDTFYTARCESACPSLVTASSRFGETEKSKTASELQLPLFLKPRSNASTSDVSSTGTTSSSRSGVERVLEEEGCGGILNQGMMEEVEKQLFEVCNVCSEDLEELLLDMGSEGGGEIEEKAEKEMGNEESGQKEEEEGEPSLVLGLRRAESQHCYPGLVSSISLDTMSNSSSSTCQAVEVDSDDFSGQVSWHNKIDQSVQRKRFVRETSV

>XP_019354952.1|Alligator_mississippiensis_Cav2.1

MAMHAVARSAMPSPPSYYDAVHDGCGLPDAPSHDGRYHQMSPEQEPPALKRRGGVVDHRDVIMAHQAHKIHSTPQAQRKEWEMARFGDEMPARYGAGGVGGAGSGGAAGGGGSRGAGGGRQGGPPGAQRMYKQSMAQRARTMALYNPIPVRQNCLTVNRSLFLFSEDNVVRKYAKKITEWPPFEYMILATIIANCIVLALEQHLPDEDKTPMSERLDDTEPYFIGIFCFEAGIKIIALGFAFHKGSYLRNGWNVMDFVVVLTGILAKVGSEFDLRTLRAVRVLRPLKLVSGIPSLQVVLKSIMKAMIPLLQIGLLLFFAILIFAIIGLEFYMGKFHTTCFDSVTGEIKDRVPCGMDEPARTCPNGTRCSKYWEGPNYGITQFDNILFAVLTVFQCITMEGWTDLLYNSNDASGNTWNWLYFIPLIIIGSFFMLNLVLGVLSGEFAKERERVENRRAFLKLRRQQQIERELNGYMEWISKAEEVILAEDETEGEQRHPFDALQRAAIKKSKTDLLNPEEADDQLADIASVGSPFARASIKSAKLENSTFFHKRERRMRFYIRRMVKTQAFYWTVLSLVALNTLCVAVVHYSQPDWLSDFLYYAEFIFLGLFMSEMFIKMYGLGTRPYFHSSFNCFDCAVIIGSIFEVVWAVMKPGTSFGISVLRALRLLRIFKVTKYWASLRNLVVSLLNSMKSIISLLFLLFLFIVVFALLGMQLFGGQFNFDTGTPATNFDTFPAAIMTVFQILTGEDWNMVMYDGIKSQGGVKRGMVFSVYFIVLTLFGNYTLLNVFLAIAVDNLANAQELTKDEQEEEEAATQKLALQKAKEVAEVSPLSAASMSIAVKEQQKNQKSSKSVWEQRTSEMRKHNLLASREALYNELDPEERWKVSYARPSRPDIKTHLDRPLVVDPQENRNNNTNKTRPSDPPLEPRFGPPQAEDFLRKPPRYHDRPRDPGSGRTYPPPESGVPELRRPHSGSREMEPPVEERSYHEADPERLKAAEPPRRHPHHQAGGKESRGGSPRDGDREHRRHRGHRRAGEDGGGTDEGPKSERRPRHRPGEGEMPDGERRRRHRHAAQSTYDGDGKREDKERRHRRRKENQGPVPVHGPHLSTTRPIQQDMGRPEPPVAEDIDNMKNNKLATTETSNPHPQSPTKLGNHANCTPSRAPDALGQLPPNSQNVANRWAPDNPSQPSNPRPPKTPENSLIVTNPGPQNNPTKTAKKPEYTAVEIPTTFPPPIHNTVVQVNKNANPEPLPKKDEEKKEEEADDQEENRPKPMVPYSSMFILSTTNPFRRLCHYIVNLRYFEMCILMVIAMSSIALAAEDPVQPNAPRNNVTPFLDLVFTGVFTFEMVIKMVDLGLVLHQGAYFRDLWNILDFIVVSGALVAFAFTGSKGKDINTIKSLRVLRVLRPLKTIKRLPKLKAVFDCVVNSLKNVLNILIVYMLFMFIFAVVAVQLFKGKFFYCTDESKEFENDCRGEYLVYEKDEVKAEKREWKKYEFHYDNVLWALLTLFTVSTGEGWPQVLKHSVDATYENQGPSPGYRMEMSIFYVVYFVVFPFFFVNIFVALIIITFQEQGDKMMEEYSLEKNERACIDFAISAKPLTRHMPQNKQSFQYRMWQFVVSPPFEYTIMAMIALNTVVLMMKFYGASDAYENVLKMFNNVFTSLFSLECLLKIMAFGVLNYFRDAWNIFDFVTVLGSITDILVTEFGNNFINLSFLRLFRAARLIKLLRQGYTIRILLWTFVQSFKALPYVCLLIAMLFFIYAIIGMQVFGNIGIDDKDDESAITEHNNFRTFFQALMLLFRSATGEGWQEIMLSCLSGKPCDENSGIMEHECGNEFAYFYFVSFIFLCSFLMLNLFVAVIMDNFEYLTRDSSILGPHHLDEYVRVWAEYDPAAWGRLTFTDMYEMLRHMSPPLGLGKKCPPRVAYKRLLRMDLPVADDNTVHFNSTLMALIRTALDIKIAKGGADKQQMDAELRKEMMAIWPNLSQKTLDLLVTPHKSTDLTVGKIYAAMMIMEYYRQSKAKKLQAMREEQNRTPLMFQRMEPPSPGQEGGPGQDALPPPPLDQGGGLGHEGGMKESQSWVTQRAQEMFQKTGTWSPERARPDDLPNSRPSSQSVEMREIGKDGYSDSDHYPPMQGHGRAASMPRLPAENQRRKVRPRGNNLSTIPDVSPMKRSASVLGHPRARGIRLDDYSLERVPPDDAQRHHLRRRDRDRDRAHRASDRSLGRYADADAGLGTDLSMTPPSGELPLKERDGERGRPKDRRHHHHHHHHHHRHEDGRPRDRDRDRRWSRSPSEARDHPPPRQGSSSVSGSPVLSTSGTSTPRRGRRQLPQTPATPRPHVSYSPVVRKAAPPVPRRHPDPAPERPTAASPRSYRHASSRWQPPDAPPGAHHSYYRGPEFNETPHGTPRTCRATASPSRHGRRLPNGYYHSQGHGHGPPKARPAGSRHGLHEPYSETDEDDCC

>XP_014748708.1|Sturnus_vulgaris_Cav2.1

MARFGEEVAARYGPGGGGSGPGAAAGGRGAGGGPRQGPPGAQRLYKQSMAQRARTMALYNPIPVRQNCLTVNRSLFLFSEDNVVRKYAKKITEWPPFEYMILATIIANCIVLALEQHLPDEDKTPMSERLDDTEPYFIGIFCFEAGIKIIALGFAFHKGSYLRNGWNVMDFVVVLTGILASVGSQFDLRTLRAVRVLRPLKLVSGIPSLQVVLKSIMKAMIPLLQIGLLLFFAILIFAIIGLEFYMGKFHTTCFDLVTDEIKVEVPCGTDEPARICPNGTKCKKYWEGPNYGITQFDNILFAVLTVFQCITMEGWTDLLYYSNDASGNTWNWLYFIPLIIIGSFFMLNLVLGVLSGEFAKERERVENRRAFLKLRRQQQIERELNGYMEWISKAEEVILAEDEPEAEPRRPFDALRRATSKRSKTDLLSPEEGEEQLGDIAAMGSPFARASLKSAKLENATFFHKRERRMRFHIRRMVKTQAFYWTVLSLVALNTLCVAIVHYDQPDWLSDFLYYAEFIFLGLFMSEMFIKMYGLGTRPYFHSSFNCFDCAVIIGSIFEVIWAVVKPGTSFGISVLRALRLLRIFKVTKYWASLRNLVVSLLNSMKSIISLLFLLFLFIVVFALLGMQLFGGQFNFDDGTPPTNFDTFPAAIMTVFQILTGEDWNAVMYDGIKSQGGVKGGMVFSVYFIVLTLFGNYTLLNVFLAIAVDNLANAQELTKDEQEEEEAANQKLALQKAKEVAEVSPLSAANMSVTMKEQQKNQKSSKSVWEQRTSELRKQNLLASREALYSELDPEERWKVPYARHLRPDMKTHLDRPLVVDPQENRNNNTNKTRPGEPPPDPRFGPLRAEELRRQQPRYPEHGVGPRPADPAGPPGSRRPRSSSREPEREPSQERGYVEGERPKGRHRSRDGRAGSPPGEREHRRHRGHRRGPEEGEEGARPERRPRHREGGRPPRGEGDPEHGDGERRRRHRHGPPPGYDGEGRREDKERRHRRRRETQPPPAPSGPTLSTTRPIQQDLGRREPPVAEDIDNMKNNKLATAEPPEPPAMGNHTPCPPSGPPRRPPPTTNPPPPENSLVVTNPGPPNNPAKAGGKPEHTAVDIPPPFPPPPSNALVQVNRNANPEPLPRKEEEEKKEEEGDGQDENGPKPMVPYSSMFILSTTNPFRRLCHYIVNLRYFEMCILMVIAMSSIALAAEDPVQPNAPRNNVLRYFDYVFTGVFTFEMVIKMVDLGLVLHQGSYFRDLWNILDFIVVSGALVAFAFTGSSKGKDINTIKSLRVLRVLRPLKTIKRLPKLKAVFDCVVNSLKNVLNILIVYMLFMFIFAVVAVQLFKGKFFYCTDESKEFEKDCRGEYLVYEKDNEVKAQKREWKKYEFHYDNVLWALLTLFTVSTGEGWPQVLKHSVDATYENQGPSPGYRMEMSIFYVVYFVVFPFFFVNIFVALIIITFQEQGDKMMEEYSLEKNERACIDFAISAKPLTRHMPQNRQSFQYRMWQFVVSPPFEYTIMAMIALNTIVLMMKFYDASDTYENVLKMFNNVFTSLFSLECLLKIMAFGVLNYFRDAWNIFDFVTVLGSITDILVTEFGNNFINLSFLRLFRAARLIKLLRQGYTIRILLWTFVQSFKALPYVCLLIAMLFFIYAIIGMQVFGNIGIEEEDDESAITQHNNFRTFFQALMLLFRSATGEAWHEIMLSCLSGKPCDENSGIKEDECGNEFAYFYFVSFIFLCSFLMLNLFVAVIMDNFEYLTRDSSILGPHHLDEYVRVWAEYDPAAWFRPFIGSMSLIGFHRLIIHDRPIIYRATRVSPACHPRVPAGGADKQQMDAELRKEMIAIWPNLSPKNLDLLVTPHKSTDLTVGKIYAAMMIMEYYRQSKAKKLQAMREEQNRTPLMFQRMEPPSPTQEGPPGPDSAPAAEPXVPPGVLGVLGVAGVPAGPPXAPLSRRAARDGDMKESQSWVTQRAQEMFQRTGTWSPERGHPDDVPNSRPNSQLVEMREMPKDGHSDSDYLPMEGHGRAASMPRLPADTQVRGQPRVPPPGTPGRLGTDLSVTTQSGDAPTKERDPERGRPKDRRHRHHHHHHHHHHHGGGAGGDRERCVPPPERHDYGRPRSRDRRWSRSPSEGREHAAPRQVGAPLRSPPLRL

>ATE62979.1|Gallus_gallus_Cav2.1

MARFGDDHPSRYGAGGGGALGSSMGRVSGGSRAAGGGPGGGGGGGAPPGGQRVYKQSMAQRARTMALYNPIPVRQNCLTVNRSLFLFSEDNAVRKYAKRITEWPPFEYMILATIIANCIVLALEQHLPDDDKTPMSERLDDTEPYFIGIFCFEAGIKIIALGFAFHKGSYLRNGWNVMDFVVVLTGILATVGSQFDLRTLRAVRVLRPLKLVSGIPSLQVVLKSIMKAMIPLLQIGLLLFFAILIFAIIGLEFYMGKFHTTCFDLVTNEIKVEVPCGTDEPARICPNGTKCRKYWEGPNYGITQFDNILFAVLTVFQCITMEGWTDLLYYSNDASGNTWNWLYFIPLIIIGSFFMLNLVLGVLSGEFAKERERVENRRAFLKLRRQQQIERELNGYMEWISKAEEVILAEDEGEGEPRHPFDALRRATIKKSKTDLLSPEDAEEQLADIASVGSPFARASLKSAKLENATFFHKRERRMRFYIRRVVKTQAFYWTVLSLVALNTLCVAIVHYDQPEWLSDFLYYAEFIFLGLFMSEMFIKMYGLGTRPYFHSSFNCFDCAVIIGSIFEVIWAVVKPGTSFGISVLRALRLLRIFKVTKYWASLRNLVVSLLNSMKSIISLLFLLFLFIVVFALLGMQLFGGQFNFDTGTPPTNFDTFPAAIMTVFQILTGEDWNAVMYDGIKSQGGVKGGMVFSVYFIVLTLFGNYTLLNVFLAIAVDNLANAQELTKDEQEEEEAANQKLALQKAKEVAEVSPLSAANLSIAVKEQQKNQKGSRSVWEQRTSELRKQNLLASREALYGELEPEERWKPPYGRHLRPDAKTHRDRPLVVDPRENRNNNTNKTRPAVAADGAALRSALRRRAAPAAPPRRPRRPPPRRGARGRPAAPEPGGGARRRGRGAGEGAEAPRPGGARRVAANGAPEAPRAPPGGGGRRGAALAAPRRGARGRGDGRRRALRLREAEAAPARSGGRRGLRRRRPPGRQGEEAPQEAGEPRPRGAARAVHHAPHPAGPGPAAAARGGGHRQHEIPACPPRPPAPLKTIKRLPKLNGQPEGPPPPDDGLVVTNPTAHNDPTAALRRAEPKAEPKAEPKAEHTAVEIPPLLPPPPSSALVQMNRNANPEPLPRKEEEEKKEEEGNGDEENGPKPMVPYSSMFILSPTNPFRRLCHYIVNLRYFEMCILMVIAMSSIALAAEDPVQPNAPRNNVLRYFDYVFTGVFTFEMVIKMVDLGLVLHQGAYFRDLWNILDFIVVSGALVAFAFTGSSKGKDINTIKSLRVLRVLRPLKTIKRLPKLKAVFDCVVNSLKNVLNILIVYMLFMFIFAVVAVQLFKGKFFYCTDESKEFEKDCRGEYLVYEKNEVKAQRREWKKYDFHYDNVLWALLTLFTVSTGEGWPQVLKHSVDATYENQGPSPGYRMEMSIFYVVYFVVFPFFFVNIFVALIIITFQEQGDKMMEEYSLEKNERACIDFAISAKPLTRHMPQNRQSFQYRMWQFVVSPPFEYTIMAMIALNTIVLMMKFYDASDAYENVLKMFNNVFTSLFSLECLLKIMAFGVLNYFRDAWNVFDFVTVLGSITDILVTEFGNNFINLSFLRLFRAARLIKLLRQGYTIRILLWTFVQSFKALPYVCLLIAMLFFIYAIIGMQVFGNIGIEEEDDESAITQHNNFRTFFQALMLLFRSATGEAWHEIMLSCLSGKPCDENSGIKEDECGNEFAYFYFVSFIFLCSFLMLNLFVAVIRDSSILGPHHLDEYVRVWAEYDPAAWGRLTLMDMYAMLRNMSPPLGLGEKCPPRVAYKRLLRMDLPVADDNTVHFNSTLMALIRTALDIKIAKGGADKQQMDAELRKEMVAIWPNLSPKNLDLLVTPHKSTDLTVGKIYAAMMIMEYYRQSKAKKLQAMREEQNRTPLMFQRMEPPSPTQEGPPGTDAAPTAEPAVRDGGIKESPSWVTQRAQEMFQRTGTWSPERGHEDVPNSRPNSQSVELREMPRDGSDGEYLPVEGHGRAASMPRLPADNQRRKVRPRGNNLSTIADSPIRRSASTLGSGRGRAVRLDEFSLERIAPDGGQRHHPRRGHRGHRSSERSGGRYTDGDTGLGTDLSITTQSGELPPPPPKDRDPERGRPKDRRHRHCMNRNN

>XP_008161515.1|Chrysemys_picta_bellii_Cav2.1

MARFGDEVPGRYGAGGSGGAGSGGAAGGGGGRGAGGSRQGGPPGAQRMYKQSMAQRARTMALYNPIPVRQNCLTVNRSLFLFSEDNVVRKYAKKITEWPPFEYMILATIIANCIVLALEQHLPDEDKTPMSERLDDTEPYFIGIFCFEAGIKIIALGFAFHKGSYLRNGWNVMDFVVVLTGILAKVGSDFDLRTLRAVRVLRPLKLVSGIPSLQVVLKSIMKAMIPLLQIGLLLFFAILIFAIIGLEFYMGKFHTTCFDSVTGEIKVDVPCGTEEPARMCPNGTECREYWEGPNYGITQFDNILFAVLTVFQCITMEGWTELLYYSNDASGNTWNWLYFIPLIIIGSFFMLNLVLGVLSGEFAKERERVENRRAFLKLRRQQQIERELNGYMEWISKAEEVILAEDETEEEQRLPFDVGALRRATIKKSKTDLLNPEEADDQLADISSVGSPFARASIKSAKLENSTFFHKRERRMRFYIRRMVKTQAFYWTVLSLVALNTLCVAIVHYSQPDWLSDFLYYAEFIFLGLFMSEMFIKMYGLGTRPYFHSSFNCFDCAVIIGSIFEVIWAVMKPGTSFGISVLRALRLLRIFKITKYWASLRNLVVSLLNSMKSIISLLFLLFLFIVVFALLGMQLFGGQFNFDDGTPSTNFDTFPAAIMTVFQILTGEDWNMVMYDGIKSQGGVHQGMVYSVYFIVLTLFGNYTLLNVFLAIAVDNLANAQELTKDEQEEEEVANQKLALQKAKEVAEVSPLSAASMSIAVKEQQKNQKSSKSVWEQRTSEMRKHNLLASREALYNEMDPEDPWKVTYARAMRPDIKTHLDRPLVVDPQENRNNNTNKTRPNEPSLEQRFSQQRAEDFLRKQARYHERSREFSSSRSYEQPDAEGPEARRQRSGSKETNLNQDGTYREQGYESSREQGYRDSASERIQAVDPHRRHQHRQGGSKESRSGSPRMAEGERDHRRQRAHRRMGEEGGGGRDEGSKVERRLRHREGSRPSRGDVEGEQPDGERRRRHRHAAQSTYDTDSKRDDRERRHRRRKENQGPSPVPGPNLSTTRPIQQDMGRHEPQVAEDIDNMKNNKLATTEPSTPHNASHLQSPAKVGHHANCTASRASHNLPGTLGQPPPNSQNAANRQAPSNPPQPGPPKTPENSLIVTNPSTQNNPAKAAKKPEHTTVDIPATFTPPINNIVVQVNKNANPDPLPKKEEEKKEEEDGEQEENGPKPMVPYSSMFILSTTNPFRRLCHYIVNLRYFEMCILMVIAMSSIALAAEDPVQPNATRNNVLRYFDYVFTGVFTFEMVIKMVDLGLVLHQGAYFRDLWNILDFIVVSGALVAFAFTGSSKGKDINTIKSLRVLRVLRPLKTIKRLPKLKAVFDCVVNSLKNVLNILIVYMLFMFIFAVVAVQLFKGKFFYCTDESKEFEKDCRGEYLVYEKDNEVKAQSREWKKYEFHYDNVLWALLTLFTVSTGEGWPQVLKHSVDATYENQGPSPGYRMEMSIFYVVYFVVFPFFFVNIFVALIIITFQEQGDKMMEEYSLEKNERACIDFAISAKPLTRHMPQNKQSFQYRMWQFVVSPPFEYTIMAMIALNTIVLMMKFDKASTAYEDVLKMFNHVFTSLFSLECLLKIMAFGVLNYFRDAWNIFDFVTVLGSITDILVTEFGNNFINLSFLRLFRAARLIKLLRQGYTIRILLWTFVQSFKALPYVCLLIAMLFFIYAIIGMQVFGNIGIEDEEEDSAITEHNNFRTFFQALMLLFRSATGEAWHEIMLACLSGKPCDENSGIKEHECGNEFAYFYFVSFIFLCSFLMLNLFVAVIMDNFEYLTRDSSILGPHHLDEYVRVWAEYDPAAWGRMTFTDMYEMLRHMSPPLGLGKKCPPRVAYKRLLRMDLPVADDNTVHFNSTLMALIRTALDIKIAKGGADKQQMDAELRKEMMAIWPNLSHKNLDLLVTPHKSTDLTVGKIYAAMMIMEYYRQSKAKKLQAMREEQNRTPLMFQRMEPPSPTQEGVPGQNALPSSQLDQGGGLSAPEGGIKESQSWVTERAQAMFQKTGTWSPERGRPEDMPNSRPNSQSVEMREIGKDGYSDSDHYLPMEGHGRAASMPRLPAENQRRKVRSRGNNLSTITDASPMKRSASMLGHSRAKGIRLDDYSLERVVPEENQRHHQRRRERSHRASERSLSRYADVDTGLGTDLSMTTPSGDLPPKERDQERGRPKDRKHHHHHHHHRHHHPSSDKERYSQERHEHSRPRSRDRRWSRSPSEGREHMPHRQGSSSVSGSPVLSTSGTSTPRRGRRQLPQTPSTPRPHVSYSPVVRKVISTGPQQQPHHPSRISTPAPRRYPSLPADQQAAERPGRSPAMERHTVPPRNDSPRAYRHASSRWSAPHVSEPHPGPRRGGYYRSPDCTAPPTEDAFSYETTHISSSGRSPRTSRAGGGSSPSRPGRRLPNGYYHSHGAAKPRSRQGLHEPSSETDEDDCC

>O00555.3|Homo_sapiens_Cav2.1

MARFGDEMPARYGGGGSGAAAGVVVGSGGGRGAGGSRQGGQPGAQRMYKQSMAQRARTMALYNPIPVRQNCLTVNRSLFLFSEDNVVRKYAKKITEWPPFEYMILATIIANCIVLALEQHLPDDDKTPMSERLDDTEPYFIGIFCFEAGIKIIALGFAFHKGSYLRNGWNVMDFVVVLTGILATVGTEFDLRTLRAVRVLRPLKLVSGIPSLQVVLKSIMKAMIPLLQIGLLLFFAILIFAIIGLEFYMGKFHTTCFEEGTDDIQGESPAPCGTEEPARTCPNGTKCQPYWEGPNNGITQFDNILFAVLTVFQCITMEGWTDLLYNSNDASGNTWNWLYFIPLIIIGSFFMLNLVLGVLSGEFAKERERVENRRAFLKLRRQQQIERELNGYMEWISKAEEVILAEDETDGEQRHPFDALRRTTIKKSKTDLLNPEEAEDQLADIASVGSPFARASIKSAKLENSTFFHKKERRMRFYIRRMVKTQAFYWTVLSLVALNTLCVAIVHYNQPEWLSDFLYYAEFIFLGLFMSEMFIKMYGLGTRPYFHSSFNCFDCGVIIGSIFEVIWAVIKPGTSFGISVLRALRLLRIFKVTKYWASLRNLVVSLLNSMKSIISLLFLLFLFIVVFALLGMQLFGGQFNFDEGTPPTNFDTFPAAIMTVFQILTGEDWNEVMYDGIKSQGGVQGGMVFSIYFIVLTLFGNYTLLNVFLAIAVDNLANAQELTKDEQEEEEAANQKLALQKAKEVAEVSPLSAANMSIAVKEQQKNQKPAKSVWEQRTSEMRKQNLLASREALYNEMDPDERWKAAYTRHLRPDMKTHLDRPLVVDPQENRNNNTNKSRAAEPTVDQRLGQQRAEDFLRKQARYHDRARDPSGSAGLDARRPWAGSQEAELSREGPYGRESDHHAREGSLEQPGFWEGEAERGKAGDPHRRHVHRQGGSRESRSGSPRTGADGEHRRHRAHRRPGEEGPEDKAERRARHREGSRPARGGEGEGEGPDGGERRRRHRHGAPATYEGDARREDKERRHRRRKENQGSGVPVSGPNLSTTRPIQQDLGRQDPPLAEDIDNMKNNKLATAESAAPHGSLGHAGLPQSPAKMGNSTDPGPMLAIPAMATNPQNAASRRTPNNPGNPSNPGPPKTPENSLIVTNPSGTQTNSAKTARKPDHTTVDIPPACPPPLNHTVVQVNKNANPDPLPKKEEEKKEEEEDDRGEDGPKPMPPYSSMFILSTTNPLRRLCHYILNLRYFEMCILMVIAMSSIALAAEDPVQPNAPRNNVLRYFDYVFTGVFTFEMVIKMIDLGLVLHQGAYFRDLWNILDFIVVSGALVAFAFTGNSKGKDINTIKSLRVLRVLRPLKTIKRLPKLKAVFDCVVNSLKNVFNILIVYMLFMFIFAVVAVQLFKGKFFHCTDESKEFEKDCRGKYLLYEKNEVKARDREWKKYEFHYDNVLWALLTLFTVSTGEGWPQVLKHSVDATFENQGPSPGYRMEMSIFYVVYFVVFPFFFVNIFVALIIITFQEQGDKMMEEYSLEKNERACIDFAISAKPLTRHMPQNKQSFQYRMWQFVVSPPFEYTIMAMIALNTIVLMMKFYGASVAYENALRVFNIVFTSLFSLECVLKVMAFGILNYFRDAWNIFDFVTVLGSITDILVTEFGNNFINLSFLRLFRAARLIKLLRQGYTIRILLWTFVQSFKALPYVCLLIAMLFFIYAIIGMQVFGNIGIDVEDEDSDEDEFQITEHNNFRTFFQALMLLFRSATGEAWHNIMLSCLSGKPCDKNSGILTRECGNEFAYFYFVSFIFLCSFLMLNLFVAVIMDNFEYLTRDSSILGPHHLDEYVRVWAEYDPAAWGRMPYLDMYQMLRHMSPPLGLGKKCPARVAYKRLLRMDLPVADDNTVHFNSTLMALIRTALDIKIAKGGADKQQMDAELRKEMMAIWPNLSQKTLDLLVTPHKSTDLTVGKIYAAMMIMEYYRQSKAKKLQAMREEQDRTPLMFQRMEPPSPTQEGGPGQNALPSTQLDPGGALMAHESGLKESPSWVTQRAQEMFQKTGTWSPEQGPPTDMPNSQPNSQSVEMREMGRDGYSDSEHYLPMEGQGRAASMPRLPAENQRRRGRPRGNNLSTISDTSPMKRSASVLGPKARRLDDYSLERVPPEENQRHHQRRRDRSHRASERSLGRYTDVDTGLGTDLSMTTQSGDLPSKERDQERGRPKDRKHRQHHHHHHHHHHPPPPDKDRYAQERPDHGRARARDQRWSRSPSEGREHMAHRQGSSSVSGSPAPSTSGTSTPRRGRRQLPQTPSTPRPHVSYSPVIRKAGGSGPPQQQQQQQQQQQQQAVARPGRAATSGPRRYPGPTAEPLAGDRPPTGGHSSGRSPRMERRVPGPARSESPRACRHGGARWPASGPHVSEGPPGPRHHGYYRGSDYDEADGPGSGGGEEAMAGAYDAPPPVRHASSGATGRSPRTPRASGPACASPSRHGRRLPNGYYPAHGLARPRGPGSRKGLHEPYSESDDDWC

>XP_020656028.1|Pogona_vitticeps_Cav2.1

MARRAVVRSSLLSPPSYYEAVHDGRGPTSPEYPFLGSWYHYMGQEPLSYKRRGGVVDHRDIIMAHQAHKIHNTPQAQRKEWEMARFGDEVPARYGAGGAGGTGSGAAGSGGGRGAGGVRQGGPPGAQRIYKQSMAQRARTMALYNPIPVRQNCLTLNRSLFLFSEDNVVRKYAKKITEWPPFEYMILATIIANCIVLALEQHLPDNDKTPMSERLDDTEPYFIGIFCFEAGIKIIALGFAFHKGSYLRNGWNVMDFVVVLTGILATVGTEFDLRTLRAVRVLRPLKLVSGIPSLQVVLKSIMKAMIPLLQIGLLLFFAILIFAIIGLEFYMGKFHTACKDIHTDEITQEAPCGTEAPSRLCPNGTRCAEYWQGPNYGITQFDNILFAVLTVFQCITMEGWTDLLYDSNDASGSAWNWLYFIPLIIIGSFFMLNLVLGVLSGEFAKERERVENRRAFLKLRRQQQIERELNGYMEWISKAEEVILAEDDIEGEPRHPFDGALRRATIKKSKTDLLNPDEADDPLADISSVGSPFARASIKSAKLENATFFHKKERRLRFYIRRIVKTQAFYWTVLSLVALNTLCVAIVHYNQPDWLSNFLYYAEFIFLGLFMSEMFIKMYGLGTRPYFHSSFNCFDCAVIIGSIFEVIWAVVKPGTSFGISVLRALRLLRIFKVTKYWASLRNLVVSLLNSMKSIISLLFLLFLFIVVFALLGMQLFGGQFNFDEGTPPTNFDTFPAAIMTVFQILTGEDWNEVMYHGIQSQGGVDKGMMFSIYFIVLTLFGNYTLLNVFLAIAVDNLANAQELTKDEQEEEEAANQKLALQKAKEVAEVSPLSAANMSIAVKEQQKNQKGAKSIWEQRTNEMRKQNLLASREALYNELDPDDRWKVSYARQMRPDMKTHLDRPLVVDPQENRNNNTNKTRPNEPPLDQHFRQQRPEEFLRKPPRYHEYPRDPHMYERPDSGSLEPRRPRSCSKEIDFNQESAYHEIDYASKEHNAILEHGYHGDCDSERIKAFDPHRRHRGSKESRSGSPRTADGDREHRRHRAHRRTPDEGGGEDSSKAERRLRHREGSRPARSDVDGEQPDGERRRRHRHAAQSTYDADGKERRHRRRKENQGSGLPSGPNLSTTRPIQQDMGRLEPPVAEDIDNMKNNKLATNETSNSHPSGHPQNPTKGGSHLNSTPSRASQNPPVPLDQFLPNSQNSVSRQAPNNPANPASPGPHKNPENSLIVTNPTTQNNPPQTAKKPEHTVVEVPPTFSPPINNAIMQVNKNANPEPLPKKEEEKKEEEDDEGGEDGPKPMPPYSSMFILSTTNPFRRLCHYIVNLHHFEMCILTVIVMSSIALAAEDPVQPRAPRNNVLRYFDYVFTGVFTFEMVVKMIDLGLILHRGSYFRDLWNILDFIVVSGALVAFAFTGSKGKDINTIKSLRVLRVLRPLKTIKRLPKLKAVFDCVVNSLKNVLNILIVYMLFMFIFAVVAVQLFKGKFFYCTDESKEFEKDCRGEYLVYEKNNEVKAERREWRKYEFHYDNVLWALLTLFTVSTGEGWPDVLKHSVDATYENQGPSPGYRMEMSIFYVVYFVVFPFFFVNIFVALIIITFQEQGDKMMEEYSLEKNERACIDFAISAKPLTRHMPQNKQSFQYRMWQFVVSPPFEYTIMAMIALNTIVLMMKFYNATPVYDNLLKMFNIVFTSLFSLECILKIIAFGLLNYFRDAWNIFDFVTVLGSITDILVTEFGDPNNFINLSFLRLFRAARLIKLLRQGYTIRILLWTFVQSFKALPXVCLLIAMLFFIYAIIGMQVFGNISIDYDEDMSEDDKPAITEHNNFRTFFQALMLLFRSATGEAWHEIMLSCLSGKPCDEKADVEGNECGNEFAYFYFVSFIFLCSFLMLNLFVAVIMDNFEYLTRDSSILGPHHLDEYVRVWAEYDPAAWGRMTFTDMYEMLRHMSPPLGLGKKCPARVAYKRLLRMDLPVADDNTVHFNSTLMALIRTALDIKIAKGGADKQQMDAELRKEMMAIWPNLSQKTLDLLVTPHKSTDLTVGKIYAAMMIMEYYRQSKAKKLQAIREEQNRTPLMFQRMEPPSPTQEGVQGQNALPSSQLDQGVEMSMEEGRMKQSESWVTQRAQEMFQKTGTWSPERGRPEEIPNSRTNSQSVEMREMSKDGYSDSDQYLPMEGHGRAASMPRLPAENQRRKARPRGNNLSTITDASPMKRSASMLGHSRSRGMRLDDYSLERVIPEENQRHHRRRERSHRTSERSLSRYTDVDTGIGTDLSMTTQSGDLPAKERDQERGRPKDRRHHHHHHHHHHHHTSSDKERYSQERHEYSRPRSRDRRWSRSPSEGRDQMPHRQGSSSVSGSPVLSTSGTSTPRRGRRQLPQTPSTPRPHISYSPVVRKGAANTGAQPPPPPPSRPRLAPAPRGEAAANSHRPLLRPGHISPTPPWSAAAAPPPPHGSSSSSAGRSPRTSRVGGSGSSSLSPSRHGGRRLPNGYYPSHGPAKSRGPGSRKGLHEPYSETDEDDCC

>XP_015274358.1|Gekko_japonicus_Cav2.1

MARFGDEVPARYGAGGSGGTGSGTASVGGSRGTGGVRQGGPPGAQRIYKQSMAQRARTMALYNPIPVRQSCLTVNRSLFLFSEDNVVRKYAKKITEWPPFEYMILATIIANCIVLALEQHLPDDDKTPMSERLDDTEPYFIGIFCFEAGIKIIALGFAFHKGSYLRNGWNVMDFVVVLTGILATVGTDFDLRTLRAVRVLRPLKLVSGIPSLQVVLKSIMKAMIPLLQIGLLLFFAILIFAIIGLEFYMGKFHTTCLDIHTEEIKLEVPCGTAAPARLCPNGTKCDKYWDGPNYGITQFDNILFAVLTVFQCITMEGWTDLLYYSNDASGNAWNWLYFIPLIIIGSFFMLNLVLGVLSGEFAKERERVENRRAFLKLRRQQQIERELNGYMEWISKAEEVILAEDDIEGEPRHPFDALRRATIKKSKTDLLTEEEEDPLADISSVGSPFARASIRSAKLENATFFHKKERRLRFYIRRMVKTQAFYWTVLSLVALNTLCVAIVHYKQPDWLSDFLYYAEFIFLGLFMSEMFIKMYGLGTRPYFHSSFNCFDCAVIIGSIFEVIWAVVKPGTSFGISVLRALRLLRIFKVTKYWASLRNLVVSLLNSMKSIISLLFLLFLFIVVFALLGMQLFGGQFNFDQGTPPTNFDTFPAAIMTVFQILTGEDWNAVMYDGIQSQGGVNKGMVFSIYFIVLTLFGNYTLLNVFLAIAVDNLANAQELTKDEQEEEEAANQKLALQKAKEVAEVSPLSAANMSIAVKEQQKNQKVSKSVWEQRTTEMRKQNLLASREALYNELDPEDRWKVNYARQMRPDMKTHLDRPLVVDPQENRNNNTNKTRPSEPPLDQHFGQQRPEEFLRKPPRYHEHARDPRAYERPDSGSFEPRRPRSSSKETDFGQEGRYHEIDYPSKEHNAIIERSYHGDCDSERIKVLDPHRHHRGSKESRSGSPHAGDGDREHRRHRAHRRTPDEGGSEETGKGERRSRHREGSRPPRGGDEGEHLDGERRRRHRHTAQSTYDTDGKEHRHRRRKENQGSSLPPGPNLSTTRPIQQDMGRIEPPVAEDIDNMKNNKLATNETSDPHPISHPQNLTKGGSHPNSTPSHNPQNLPVPLDQPLPNSQNSGSRRVPNNPSNPSNPGPPKNPENSLIVTNPTTQNNPPQTAKKPEHTTVEIPPSFPPPINNATMQVNKNANPEPLPKKEEEKKEEEDDDGGDNGPKPMPPYSSMFILSTTNPFRRLCHYIVNLRYFEITILMVIAMSSIALAAEDPVQPSAPRNNVLRYFDYVFTGVFTFEMVVKMIDLGLVLHQGAYFRDLWNILDFIVVSGALVAFAFTGSKGKDINTIKSLRVLRVLRPLKTIKRLPKLKAVFDCVVNSLKNVLNILIVYMLFMFIFAVVAVQLFKGKFFFCTDESKEFEKDCKGEYLVYEKNNEVKAEKREWKKYDFHYDNVLWALLTLFTVSTGEGWPQVLKNSVDATYENQGPSPGYRMEMSIFYVVYFVVFPFFFVNIFVALIIITFQEQGDKMMEEYSLEKNERACIDFAISAKPLTRHMPQNKQSFQYRMWQFVVSPPFEYTVMAMIALNTIVLMMKYYNASNLYEQVLKMFNIVFTALFSLECILKIIAFGLVNYFRDAWNIFDFVTVLGSITDILVTEFGDPNNFISLSFLRLFRAARLIKLLRQGYTIRILLWTFVQSFKALPYVCLLIAMLFFIYAIIGMQVFGNIGIKDEDSAINEHNNFRTFFQALMLLFRSATGEAWHEIMLSCLGGKACDEKADAKDNECGNEFAYFYFVSFIFLCSFLMLNLFVAVIMDNFEYLTRDSSILGPHHLDEYVRVWAEYDPSAWGRMTFMDMYEMLRHMSPPLGLGKKCPARVAYKRLLRMDLPVADDNTVHFNSTLMALIRTALDIKIAKGGADKQQMDNELRKEMMAIWPNLSQKTLDLLVTPHKSTDLTVGKIYAAMMIMEYYRQSKAKKLQVMREEQNRTPLMFQRMEPPSPSQEGGQGQNALPSSQLDQGGGMSMQEGGMKESESWVTQRAQEMFQKTGTWSPERGHPEDTPNSRTNSQSVEMREMAKDGYSDSDPYLPMEGHGRAASMPRLPAENQVRVFLSISFVNLRTVVISHPCSLSCPRRQQSSSLSLVFVFPVLMQIERTLEPSNSL

>XP_021330116.1|Danio_rerio_Cav2.1

MARFGDEVPSRYGGGPGHAQGGPGRGGSRQGGPPGAQQRMYKQSMAQRARTMALYNPIPVRQNCFTVNRSLFIFSEDNFVRKYAKKITEWPPFEYMILATIIANCIVLALEQHLPDGDKTPMSERLEDTEPYFIGIFCFESGIKILALGFAFHKGSYLRNGWNVMDFVVVLTGILSTVGSDFDLRTLRAVRVLRPLKLVSGIPSLQVVLKSIMKAMIPLLQIGLLLFFAILMFAIIGLEFYMGKFHTTCFDKITDEIREEFPCGEEVPARICPNGTVCKKYWLGPNYGITQFDNILFAVLTVFQCITMEGWTDLLYYSNDAAGSAWNWMYFIPLIIIGSFFMLNLVLGVLSGEFAKERERVENRSEFLKLKRQQQIERELNGYLEWICKAEEVILADEDNDPDDRMPFDGSRRRPTIKKSKTDLLDAEDGDGDIGSPFARGSLKSSKLEGSSFHKKERRLRFFIRRIVKTQAFYWTVLCLVGLNTLCVAVVHYDQPETLSDFLYFAEFIFLGIFMSEMCIKMYGLGTRPYFHSSFNCFDCIVICGSIFEVLWAMIQPGTSFGISVLRALRLLRIFKVTKYWASLRNLVVSLLNSMKSIISLLFLLFLFIVVFALLGMQLFGGQFNFEAGTPPTNFDTFPAAIMTVFQILTGEDWNMVMYDGIESQGGVKKGMVFSVFFIVLTLFGNYTLLNVFLAIAVDNLANAQELTKDEQEEEQAANKKMALQTAKEVAEVSPLSAANLSIAAKEQQKNHKGCKSVWEQRTSELRRQTLVNSREALYNELDPEDRWKVSYSRHIRPDMKTHLDRPLVVDPQENRNNNTNKTRPGDGQPQHSHQLLHKQPNFNEAETGGGGGSESWPALERGDSTGSRRSLLGSEHYEEGRESHRRHQHSQHCSQARQHRKAAEGGGEPGGRRHRSKAKERGPECEHSDGGERKHRRHRHEGEGRKERGVRHRNRRDGHASGPTLSTTRPIQKTLSRQDSQYSEDLDNAMNNKLATQPHDSLLNLANTAHSAGLAHTGTESSLILTNPSSTIANVTSIGHLGMKPEYTAVDIPPMFPSSNAILQVNKNANTEPLPKKEDTKGDDDDDEKDDGGPKPMPPYTSMFILTTTNPFRRLCHYIVTLRYFEMCILLVIAMSSIALAAEDPVWPESPRNNVLRYFDYVFTGVFTFEMLIKMVDLGLVLHQGSYFRDLWNILDFIVVSGALVAFAFTGSSKGKDISTIKSLRVLRVLRPLKTIKRLPKLKAVFDCVVNSLKNVLNILIVYMLFMFIFAVVAVQLFKGRFFYCTDESKEFERDCRGEYLVYERDNEVRSQKREWKKYDFHYDNVLWALLTLFTVSTGEGWPQVLKHSVDATYENQGPSPGYRMEMSIFYVVYFVVFPFFFVNIFVALIIITFQEQGDKMMEDYSLEKNERACIDFAINAKPLTRHMPQNKQTFQYRMWEFVVSPPFEYTIMALIALNTIVLMMKYDGASLTYEDVLKYLNIVFTSLFSMECILKIIAFGALNYFKDAWNIFDCVTVLGSITDILVTELGNNFINLSFLRLFRAARLIKLLRQGETIRILLWTFVQSFKALPYVCLLIAMLFFIYAIIGMQLFGNIKIEENSDSAITQHNNFRTFFQALMLLFRSATGEAWHDIMLSCLGKKPCDILSDNPKPECGSEFAYLYFVSFIFLCSFLMLNLFVAVIMDNFEYLTRDSSILGPHHLDEYVRIWAEYDPAACGRISYRDMYEMLRHMSPPLGLGKKCPARVAYKRLLRMDLPVADDNTVHFNSTLMALIRTALDIKIAKGGVDKHQMDAELRKEMMAIWPNLSQKNLDLLVTPHKSTDLTVGKIYAAMMIMEYYRQSKAKRTQALHDEQNRTPLMFQRLEPPSPSQDVGPGLTGLPDTQMHPANHLPVDERIPESQSWVTARAQEISQKAGNWSPEGQHPDDTTDNRRQSQTVEMREMGRDGYSDTEQYLPMEGHGRAASMPRLPADNQQPRRKGRHRGENLTPITDNSPMKRSASSLGHGRAGRSMRGEDYAMERVIPEEGHRHGHRHRDRSHRASERSLSRYTDADTGLGTDLSTTTQSGDLPSKERDRGRAKDRKHHHHHHHHHGSLDKEHYGHERDRGEYGHRQSRERDRRWSRSPSEGRECLTHRQGSSSVSGSPVPSTSGTSTPRRGRRQLPQTPATPRPHVTYSPVVRKPISSTPPPGQQSRLPTPTSRRFSPTGPEPPLPPHHHPSPPHHGSPRSGRHAHWGPEPAESVEGDGFYDDQDYEFNHHEPPSYEQSPVQGNPHPHSPRTSRHNTPPQGQAHPRRMPNGYRSSSPSPHHHAPAHPGTHKPPHPRGPRKGLHEPYSETDEDDWC

>XP_008172570.1|Chrysemys_picta_bellii_Cav2.2

MARFGDDLPTRYGGGGPAGAGRGGSRQGGPPPGQRMYKQSMAQRARTMALYNPIPVKQNCFTVNRSLFIFSEDNVIRKYAKRITEWPPFEYMILATIIANCIVLALEQHLPEDDKTPMSERLDDTEPYFIGIFCFEAGIKIIALGFVFHKGSYLRNGWNVMDFVVVLTGILATAGTDFDLRTLRAVRVLRPLKLVSGIPSLQVVLKSIMKAMVPLLQIGLLLFFAIVMFAIIGLEFYMGKFHKTCFSNETGEEVGDFPCGEDLPARQCENGTTCRKYWPGPNYGITNFDNILFAVLTVFQCITMEGWTDILYNTNDAAGNTWNWLYFIPLIIIGSFFMLNLVLGVLSGEFAKERERVENRRAFLKLRRQQQIERELNGYLEWIFKAEEVMLAEEDKNAEEKSPLDVLKRATIKKSKNDLIHAEEGEDHFTDICSVGSPFARASLKSGKNESSSYFRRKEKMFRFFIRRMVKAQSFYWIVLCVVTLNTLCVAMVHYDQPEGLTTALYFAEFVFLGLFLTEMSLKMYGLGPRNYFHSSFNCFDFGVIVGSIFEVIWAAVKPGTSFGISVLRALRLLRIFKVTKYWNSLRNLVVSLLNSMKSIISLLFLLFLFIVVFALLGMQLFGGQFNFQDETPTTNFDTFPAAILTVFQILTGEDWNAVMYHGIVSQGGVHSGMFSSIYFIVLTLFGNYTLLNVFLAIAVDNLANAQELTKDEEEMEEATNQKLALQKAKEVAEVSPMSAANISIAAKQQNSSKSKSVWEQRTSQIRMHNFRASCEALYNELDPEERVRYATTLHIRPDMKTHLDRPLVVEPRTEGRNDVGKLSPVDVQETEQAKATSADNAEVPRKHHRHRDKDKMGEQEKSDVAKDENGESGTNNKEERHRQHRSRSKEAEGGSKEGKSERNRSQEGGKRHHRRGSMEEGAEKEYRRHRPHRHSAERQAKEGNGTANGARSERRSRHRGGSRSGNREGDPGLKGENGEEPHRRYKIRHRALSTYESVEKENGEKEGEAGEKDLKNHQPRETQCEIEAGGSVSVVPVHTLPSTYLQKVPEQPEDADNEKNVTRMTQPPLDKTTTVNIPVTITAPPGETTVIPMNNVEFESKTEEKKDMEIEDLTKNGPKPILPYSSMFILSPTNPIRRLFHYIVNMRYFEMVILIVIALSSIALAAEDPVQAESPRNDALKYLDYIFTGVFTFEMVIKMIDLGLLLHPGSYFRDLWNILDFIVVSGALVAFAFSGSKGKDINTIKSLRVLRVLRPLKTIKRLPKLKAVFDCVVNSLKNVLNILIVYMLFMFIFAVIAVQLFKGKFFYCTDESKELEKDCRGQYLDYEKSEVEAQPRQWKKYEFHYDNVLWALLTLFTVSTGEGWPTVLKHSVDATYEEQGPSPGYRMEMSIFYVVYFVVFPFFFVNIFVALIIITFQEQGDKVMSECSLEKNERACIDFAISARPLXRYMPQNKQSFQYKMWKFVVSPPFEYFIMVMIALNTIVLMMKFYGAPDAYEEMLKCLNIVFTSMFSMECVLKIIAFGVLNYFRDAWNVFDFVTVLGSITDILVTEIAETDNFINLSFLRLFRAARLIKLLRQGYTIRILLWTFVQSFKALPYVCLLIAMLFFIYAIIGMQVFGNIALDDDTSINRHNNFRTFLQALMLLFRSATGEAWHEIMLSCLSNRACDKLSGLTKNECGSDFAYFYFVSFIFLCSFLMLNLFVAVIMDNFEYLTRDSSILGPHHLDEFIRVWAEYDPAACGRISYTDMYEMLRHMSPPLGLGKKCPARVAYKRLVRMNMPISDPDLTVHFTSTLMALIRTALEIKLASAGVKQHQCDAELRKEISLVWPNLSQKTLDLLVPPHKPDEMTVGKVYAALMIFDFYKQNKNSREQVHQPPGGLCQTGPVSLFHPLKATLEQTQPLVFNHAKAFLRQKSCTSLNNGGTLPAPESGIKESVSWGTQRTQDVFYETRTPAFERGHSEEIPIERTSKQVVEMREISPTLANGEHQPGLESQGRAASMPRLAAETQPIPDTSPMKRSISTLTPQRPHAMHLYEYSLERMPPDQAHHHHHHRCHRRRDKKQKSLDRSPNQLADGDAGEASSKDKKQERGRSQERKLHSSSSSEKQRFYSCDRYGSRDRSQPKSADHSRPTSPNGGLEPGPHRQGSGSVNGSPLLSTSGASTPCRGRRQLPQTPLTPRPSITYKTANSSPVHFTSFQTSLPAFSPGRLSRGLSEHNALLHGDSQSHSRSPVARIGSDPYLGHRDDSDSPYRVVPEDTLTFEEAVATNSGRSSRTSYVSSLTSQSHQIRRVPNGYHYTLGLSTGPGTCARTRSYYHEADEDDWC

>XP_019396155.1|Crocodylus_porosus_Cav2.2

MARFGEELPSRYGGGGPAGAGRGSSRQGGPQPGQRVYKQSMAQRARTMALYNPIPVKQNCFTVNRSLFIFSEDNVIRKYAKRITEWPPFEYMILATIIANCIVLALEQHLPDGDKTPMSERLDDTEPYFIGIFCFEAGIKIIALGFVFHKGSYLRNGWNVMDFVVVLTGILATAGTDFDLRTLRAVRVLRPLKLVSGIPSLQVVLKSIMKAMVPLLQIGLLLFFAIVMFAIIGLEFYMGKFHKTCFSNETGEEVGDFPCGEELPARQCESGSTCRKYWQGPNYGITNFDNILFSVLTVFQCITMEGWTDILYNTNDAAGNTWNWLYFIPLIIIGSFFMLNLVLGVLSGEFAKERERVENRRAFLKLRRQQQIERELNGYLEWIFKAEEVMLAEEDKNAEEKSPLDVLKRATIKKSKNDLIHAEEGEDHFTDICSVGSPFARASLKSGKNESSSYFRRKEKMFRFFIRRMVKAQSFYWIVLCVVTLNTLCVAMVHYAQPEKLTTALYFAEFVFLGLFLTEMSLKMYGLGPRNYFHSSFNCFDFGVIVGSIFEVIWAAVKPGTSFGISVLRALRLLRIFKVTKYWNSLRNLVVSLLNSMKSIISLLFLLFLFIVVFALLGMQLFGGQFNFQDETPTTNFDTFPAAILTVFQILTGEDWNAVMYHGIESQGGVHSGMFSCVYFIVLTLFGNYTLLNVFLAIAVDNLANAQELTKDEEEMEEATNQKLALQKAKEVAEVSPISAANISIAAKQQNSSKSKSVWEQRTSQIRMHNFRASCEALYNELDPEERVRYATTLHIRPDMKTHLDRPLVVEPRTEGRNNVNKLSPGDVQEIEQTKTISAECAEAPRKHHRHREKDKTGEQEKSDMTKDENGESGTNNKEERQRQHRSRSKEAEGSSKEGKIERSRGQEGGKRHHRRGSVEEGAEKEHRRHRTHRHSAERQGKEGNGTINGARTERRSRHRGGSRSGNREGDPGSKGENGEEPHRRHRIRNRALSTYDPAEKENGEKEGDVGEKDHRNHQPKENQCEIEAGASVSVVPVQTLPSTYLQKVPEQPEDADNQKNVTRMTQPPLDKTTTVNIPVTITAPPGETTVIPMNSVEFESKTEEKKDVDGDDLTKNGPKPILPYSSMFILSPTNPIRRLFHYIVNLRYFEMVILIVIALSSIALAAEDPVQAESPRNDALKYLDYIFTGVFTFEMVIKMIDLGLLLHPGSYFRDLWNILDFIVVSGALVAFAFSGTKGKDINTIKSLRVLRVLRPLKTIKRLPKLKAVFDCVVNSLKNVLNILIVYMLFMFIFAVIAVQLFKGRFFYCTDESKELEKDCRGQYLDYEKSEVEAKPRQWKKYEFHYDNVLWALLTLFTVSTGEGWPTVLKHSVDATYEEQGPSPGYRMEMSIFYVVYFVVFPFFFVNIFVALIIITFQEQGDKVMSECSLEKNERACIDFAISAKPLTRYMPQNKQSFQYKMWKFVVSPPFEYFIMVMIALNTIVLMMKFYDAPEAYEEMLKCLNIVFTSMFSMECVLKIIAFGVLNYFRDAWNVFDFVTVLGSITDILVTEIADTDNFINLSFLRLFRAARLIKLLRQGYTIRILLWTFVQSFKALPYVCLLIAMLFFIYAIIGMQVFGNIALDDDSSINRHNNFRTFLQALMLLFRSATGEGWHEIMLSCLSNRACDPRSGLTKDECGSDFAYFYFVSFIFLCSFLMLNLFVAVIMDNFEYLTRDSSILGPHHLDEFVRVWAEYDPAACGRISYTDMYEMLRHMSPPLGLGKKCPARVAYKRLVRMNMPISDQDLTVHFTSTLMALIRTALEIKLASGGVKQHQCDAELRKEISLVWPNLSQKTLDLLVPPHKPDEMTVGKVYAALMIFDFYKQNKNSREQVHQPPGGLCQTGPVSLFHPLKATLEQTQPPAFNNAKAFLRQKSSASLNNGGTLPAPEGGIKESSSWGTQRTQDIFYETRMPAFERGHSEEIPIERTSKQVVEMREISPTLANGEHQPGLESQGRAASMPRLAAETQPIPDTSPMKRSISTLTPQRPHPMHLYNYSLERMPPDQVHHHHHHRCHRRKEKKQKSMDRPSHHLADGDAVAQPGETSSRDRKQERGRSQERKQHSSSSSEKQRFYSCDRYGSRDHSQPKSADHSRPTSPNGGPEQGPHRQGSGSVNGSPLMSTSGASTPCRGRRQLPQTPLTPRPSITYKTANSSPVHFTSFQTGLPTFSPGRLSRGLSEHNALLRGDSQNHSHTMVARIGSDPYLGHRDDSDSPYRVVPEDTLTFEEAVATNSGRSSRTSYVSSLTSQSHQIRRVPNGYHYTLGLNTGPGTGSRGRSYYHEADEDDWC

>XP_015134766.1|Gallus_gallus_Cav2.2

MARFGDDLPTRYGGGGPAGAGRGSSRQGGPQAGQRMYKQSMAQRARTMALYNPIPVKQNCFTVNRSLFIFSEDNVIRKYAKRITEWPPFEYMILATIIANCIVLALEQHLPDGDKTPMSERLDDTEPYFIGIFCFEAGIKIIALGFVFHKGSYLRNGWNVMDFVVVLTGILATAGTDFDLRTLRAVRVLRPLKLVSGIPSLQVVLKSIMKAMVPLLQIGLLLFFAIVMFAIIGLEFYMGKFHKTCFSNETGEEVGDFPCGEEPPARQCESGTTCREYWQGPNYGITNFDNILFAVLTVFQCITMEGWTDILYNTNDAAGNTWNWLYFIPLIIIGSFFMLNLVLGVLSGEFAKERERVENRRAFLKLRRQQQIERELNGYLEWIFKAEEVMLAEEDKNAEEKSPLDVLKRAAIKKSKNDLIHAEEGEDHFTDICSVGSPFARASLKSGKNESSSYFRRKEKMFRFFIRRMVKAQSFYWIVLCVVALNTLCVAMVHYDQPEKLTTALYFAEFVFLGLFLTEMSLKMYGLGPRNYFHSSFNCFDFGVIVGSIFEVIWAAVKPGTSFGISVLRALRLLRIFKVTKYWNSLRNLVVSLLNSMKSIISLLFLLFLFIVVFALLGMQLFGGQFNFRDETPTTNFDTFPAAILTVFQILTGEDWNAVMYHGIESQGGVHSGMFSSIYFIVLTLFGNYTLLNVFLAIAVDNLANAQELTKDEEEMEEATNQKLALQKAKEVAEVSPMSAANISIAAKQQNSSKSKSVWEQRTSQIRMHNFRASCEALYNELDPEERVRYATTLHIRPDMKTHLDRPLVVEPRGEGRNNISKLSPVDVQEVEQTKVSSTDGAEAPRKHHRHRDKEKLGEQEKGDVTKDENGESGINNKEERHRQHRSRSKEVEGGSKEGKSDRSRGQEGGKRHHRRGSVEEGVEKEHRRHRTHRHSAERQGKEGNGTINGARSERRTRHRGGSRSGNREGEPGSKGENGEEPHRRHRFRSRALSTYDSVEKENREKEGETAEKEHQNHQPKENQCEIEASGSVSIPVHTLPSTYLQKVPEQPEDADNQKNVTRMIQPPLDKTTTVNIPVTITAPPGETTVIPMNNVEFESKTEEKKDVDDLTKNGPKPILPYSSMFILSPTNPIRRLFHYIVNLRYFEMVILIVIALSSIALAAEDPVQAESPRNDALKYLDYIFTGVFTFEMVIKMIDLGLLLHPGSYFRDLWNILDFIVVSGALVAFAFSGTKGKDINTIKSLRVLRVLRPLKTIKRLPKLKAVFDCVVNSLKNVLNILIVYMLFMFIFAVIAVQLFKGRFFYCTDESKELEKDCRGQYLDYEKNEVEAQPREWKKYEFHYDNVLWALLTLFTVSTGEGWPTVLKHSVDATYEEQGPSPGYRMEMSIFYVVYFVVFPFFFVNIFVALIIITFQEQGDKVMSECSLEKNERACIDFAISAKPLTRYMPQNKQSFQYKMWKFVVSPPFEYFIMVMIALNTIVLMMKFYDAPEAYEEMLKCLNIVFTSMFSMECVLKIIAFGVLNYFRDAWNVFDFVTVLGSITDILVTEIADTDNFINLSFLRLFRAARLIKLLRQGYTIRILLWTFVQSFKALPYVCLLIAMLFFIYAIIGMQVFGNIALNDETSINRHNNFRTFLQALMLLFRSATGEAWHEIMLSCLSNRACDPLSGLTKNECGSEFAYFYFVSFIFLCSFLMLNLFVAVIMDNFEYLTRDSSILGPHHLDEFVRVWAEYDPAACGRISYTDMYEMLRHMSPPLGLGKKCPARVAYKRLVRMNMPISPEDLTVHFTSTLMALIRTALEIKLASGGVKQHQCDAELRKEISLVWPNLSQKTLDLLVPPHKPDEMTVGKVYAALMIFDFYKQNKNSREQVHQPPGGLCQPGPVSLFHPLKATLEQTQPSAFSSAKAFLRQKSSASLNNGGTLPAPEGGIKESSSWGTQRTQDVFYETRTPAFERGHSEEIPIERVVEMREISPTLANGEHQPGLESQGRAASMPRLAAETQPIPDTSPMKRSISTLTPQRPHPMHLYEYSLERVPTDQVHHHHHHRCHRRKEKKQKSLDRTTHQLADGEAVAQSGESSSKDKKQERGRSQERKQHSSSSSEKQRFYSCDRYGSRDRSQPKSADQSRPTSPNGGPEQGPHRQGSGSVNGSPLLSTSGASTPCRGRRQLPQTPLTPRPSITYKTANSSPVHFSTSPGGLPPFSPGRLSRGLSEHNALLRGDQQPPPAAVARIGSDPYLGHRDAADSPIGAAPEDTLTFEEAVATNSGRSSRTSYVSSLTSQSHQARRVPNGYHYTLGLNTGPGTGTRGRSYYHEADEDDWC

>XP_020668759.1|Pogona_vitticeps_Cav2.2

MILATIIANCIVLALEQHLPDGDKTPMSERLDDTEPYFIGIFCFEAGIKIMALGFVFHKGSYLRNGWNVMDFVVVLTGILATAGTQFDLRTLRAVRVLRPLKLVSGIPSLQVVLKSIMKAMVPLLQIGLLLFFAIVMFAIIGLEFYMGKFHKACFSNETGERVGDFPCGEEKPARVCEEGTCKKYWEGPNFGITNFDNILFAVLTVFQCITMEGWTDILYNTNDAAGNMWNWLYFIPLIIIGSFFMLNLVLGVLSGEFAKERERVENRRAFLKLRRQQQIERELNGYLEWIFKAEEVMLAEEDKNAEEKSPLDVLKRATIKKSKNDLIHAEEGEDHFTDVCSVGSPFARASLKSGKNDSSTYFRRKEKMFRFFIRRMVKAQSFYWMVLCVVALNTMCVAIVHYDQPEGLTTALYFAEFVFLGLFLTEMSLKMYGLGPRNYFHSSFNCFDFGVIVGSIFEVIWAAVKPGTSFGISVLRALRLLRIFKVTKYWNSLRNLVVSLLNSMKSIISLLFLLFLFIVVFALLGMQLFGGQFHFNNETPTTNFDTFPTAILTVFQILTGEDWNAVMYQGIQSQGGVHSGMFSSAYFIVLTLFGNYTLLNVFLAIAVDNLANAQELTKDEEEMEEATNQKLALQKAKEVADVSPLSAANISIAAKQQNSSKSKSVWEQRTSQIRIKNRASCEALYNELDPEERVRYATNLHIRPDMKTHLDRPLVVEPRSDGRNNHVSKLSPVDVQEAPEQGKAPAPEGAEGARRHHRHREKDKGGEHEKISPPKEENGESGPNHKEECHRPHRSRSKEGEGGGREGRGEHNRGQEGGRRHHRRGSMEEGAEREHRRHRGHRHSAERQGREGNGTVNGAKTERRSRHRGGSRPGNREGEPGSKGEGGEEPHRRHRMRQKALSTYESVEKENGEKEGELGDKELRNHQPRAGPGETEPGGSVPVAPLHTLPSTCLQRVPEQPEDADNQRNVTRMGQPPLDKTATVNIPVTVTAPPGDTAVIPINNVDFETKTEEKKDMDADNDLTENGPKPILPYSSMFILSPTNPIRRLCHYIVNMRHFEMVILFVIVLSSIALATEDPVQAESPRNEALKYLDYIFTGVFTFEMVIKMIDLGLLLHPGSYFRDLWNILDFIVVSGALVAFAFSGNKGKDINTIKSLRVLRVLRPLKTIKRLPKLKAVFDCVVNSLKNVLNILIVYMLFMFIFAVIAVQLFKGRFFYCTDESKDLEKDCRGQYLSYENDEVEAQPRQWKKYEFHYDNVLWALLTLFTVSTGEGWPTVLKHSVDATDENQGPSPGYRMEMSIFYVVYFVVFPFFFVNIFVALIIITFQEQGDKVMSECSLEKNERACIDFAISAKPLTRYMPQNKQSFQYKMWKFVVSPPFEYFIMVMIALNTIVLMMKFYDAPEPYENMLKCLNIVFTSMFSLECVLKIIAFGALNYFRDAWNIFDFVTVLGSITDILVTEIADTDNFINLSFLRLFRAARLIKLLRQGYTIRILLWTFVQSFKALPYVCLLIAMLFFIYAIIGMQVFGNIALNDDTAINRHNNFQTFLQALMLLFRSATGEAWHEIMLACLSHQACDELSNLSKNECGSDFAYFYFVSFIFLCSFLMLNLFVAVIMDNFEYLTRDSSILGPHHLDEFVRVWAEYDPAACGRISYTDMYEMLRHMSPPLGLGKKCPARVAYKRLVRMNMPISNDDLSVHFTSTLMALIRTALDIKLASGALQQQCDAELRKEISLVWPNLSPKTLDLLVPPHKSEAMTVGKVYAALMIFDFYKQNKNTRDPSHPAPGGLGQTGTVSLFHPLKAALEQSQPAFPNAFLRQKSSASLNNGGALPTPEVRRIKESVSWGTQRTQDVFYETRTPAFERGRSEEIPVAQASKQVVEMREMSPTLANGEHQPGLESQGRAASMPRLAAETQPIPDTSPMKRSISTLNPQRPHTAHLYEHSRPAEHTHHHHHHRCHRRKERKQKSMDKPPSHLAESGAPAGEAASKSKKQERGRSQERKLHSSSSSEKQRFYSCDRFGSREPSQAKPSERSQPTSPNGEQDHGAQKQGSGSVNGSPLLMTSGASTPCRGRRQLPQTPLTPRPSVTYKTANSSPVHFPGLQTSLPTFSPGRLSRGLSEHNALLHGEAQGVPSQPLVSRIGSDPYLGHREDGDSLYHAVPEDTLTFEEALATNSGRSSRTSYVSSLTSQSHQIRRVPNGYHYTLGLSGGPGTSTRARSYYHERSEDDWC

>XP_015276634.1|Gekko_japonicus_Cav2.2

MVLKHFPAIWASAWSWAGIHNVVQFFPLTIASTPFEYMILATIIANCIVLALEQHLPDGDKTPMSERLDDTEPYFIGIFCFEAGIKIIALGFVFHKGSYLRNGWNVMDFVVVLTGILATAGTQFDLRTLRAVRVLRPLKLVSGIPSLQVVLKSIMKAMVPLLQIGLLLFFAIVMFAIIGLEFYMGKFHKACFSNETEERVEDFPCGEEPPARQCDKGTVCQKYWEGPNFGITNFDNILFAVLTVFQCITMEGWTDILYNTNDAAGNTWNWLYFIPLIIIGSFFMLNLVLGVLSGEFAKERERVENRRAFLKLRRQQQIERELNGYLEWIFKAEEVMLAEEDKNAEEKSPLDGRATSEGPILRGTTTLEISSGGSCNMLKRATIKKSKNDLIHAEEGEDHFTDVCSVGSPFARASLKSAKNESSSYFRRKEKMFRFFIRRMVKAQSFYWIVLCVVALNTLCVASVHYDQPEGLTTALYFAEFVFLGLFLTEMSLKMYGLGPRNYFHSSFNCFDFGVIVGSIFEVIWAAVKPGTSFGISVLRALRLLRIFKVTKYWNSLRNLVVSLLNSMKSIISLLFLLFLFIVVFALLGMQLFGGQFHFNDETPTTNFDTFPTAILTVFQILTGEDWNAVMYQGIESQGGVHSGMFSSIYFIVLTLFGNYTLLNVFLAIAVDNLANAQELTKDEEEMEEATNQKLALQKAKEVADVSPLSAANISIAAKQQNASKSKSVWEQRTSQIRMHNFRTSCEALYNELDPEDRVRYATTLHIRPDMKTHLDRPLVVEPRSSEGQNSNGSKLSPTDVQESAEQANSAPAECTEGPWKHHRHRDKDKAGEQEKSNVPKEENGESGTNNKEECHRPHRSRSKEGEGGGKEGKSERNRTQEGGRRHHRRGSMEEGAEREHRRHRSHRHSAERQAKEGNGTVNGTKTERRSRHRGGSRPGNREGDPGSKGEGCEEPHRRHKTRHKALSTYESVEKENGEKEGELGEKELRNHQPREGQCDTEAGGSVIVAPLHTLPSTCLQRVPEQPEDADNQKNVTRMGQPSLDKTATVNIPVTVTAPPGDATIIPMNNVEFETKTEEKKDMDADDLTKNGPKPILPYSSMFVLSPTNPIRRLCHYIVNMRYFEMVILIVIALSSIALAAEDPVQAESPRNEALKYLDYIFTGVFTFEMVIKMIDLGLILHPGSYFRDLWNILDFIVVSGALVAFAFSQNKGKDINTIKSLRVLRVLRPLKTIKRLPKLKAVFDCVVNSLKNVLNILIVYMLFMFIFAVIAVQLFKGRFFYCTDESKDLEKDCRGQYLDYEKDEVEAQPRKWNKYEFHYDNVLWALLTLFTVSTGEGWPTVLKHSVDATYENQGPSPGFRMEMSIFYVVYFVVFPFFFVNIFVALIIITFQEQGDKVMSECSLEKNERACIDFAISAKPLTRYMPQNKQSFQYKMWKFVVSPPFEYFIMAMIALNTIVLMMKFYDAPAPYEDMLKCLNIVFTSMFSLECVLKIIAFGALNYFRDAWNIFDFVTVLGSITDILVTEIADTDNFINLSFLRLFRAARLIKLLRQGYTIRILLWTFVQSFKALPYVCLLIAMLFFIYAIIGMQVFGNIALNDETSINRHNNFQTFLQALMLLFRSATGEAWHEIMLSCLSNQACDKLSKLSKNECGSDFAYFYFVSFIFLCSFLMLNLFVAVIMDNFEYLTRDSSILGPHHLDEFIRVWAEYDPAASCRIHYKDMYNLLRAIAPPLGLGKKCPHRVAYKRLVRMNMPISNDDLTVHFTSTLMALIRTALEIKLASGVLQHQCDAELRKEISLVWPNLSQKTLDLLVPPHKPEAMTVGKVYAALMIFDFYKQNKNTRDPAHPATQTGTVSLFHPLKAALEQSQPVFPNAFLRQKSSASLNNGGALPPPEGRGIKESISWGTQRTQDVFYETRTPAFVRGHSEEISVEQTIEMQEISPMLANGEHQPGLESQGRAASMPRLAAETQRSKARSPGRYLAPIPDTSPMKRSISTLNPQRPHATHLYEHSRPTEHTHHHHHHRCHRRREKKQKSIDKPPNHLADGEAAAQTGEAASKAKRQERGRSQERKLHSSSSSEKQRFYSCDRYGSRDHSQPKSTERSRPTSPSGGQEHGMQKQGSGSVNGSPLLSTSGASTPCRGRRQLPQTPLTPRPSITYKTANSSPVHFSSFQTGLPTFSPGRLSRGLSEHNALLHGDSQGHSQTLVARIGSDPYLGHREDSDSPYHVVPEDTLTFEEAVATNSGRSSRTSYVSSLTSQSHQIRRVPNGYHYTLGLSSGPGTSTRARSYYHERDEDDWC

>NP_000709.1|Homo_sapiens_Cav2.2

MVRFGDELGGRYGGPGGGERARGGGAGGAGGPGPGGLQPGQRVLYKQSIAQRARTMALYNPIPVKQNCFTVNRSLFVFSEDNVVRKYAKRITEWPPFEYMILATIIANCIVLALEQHLPDGDKTPMSERLDDTEPYFIGIFCFEAGIKIIALGFVFHKGSYLRNGWNVMDFVVVLTGILATAGTDFDLRTLRAVRVLRPLKLVSGIPSLQVVLKSIMKAMVPLLQIGLLLFFAILMFAIIGLEFYMGKFHKACFPNSTDAEPVGDFPCGKEAPARLCEGDTECREYWPGPNFGITNFDNILFAILTVFQCITMEGWTDILYNTNDAAGNTWNWLYFIPLIIIGSFFMLNLVLGVLSGEFAKERERVENRRAFLKLRRQQQIERELNGYLEWIFKAEEVMLAEEDRNAEEKSPLDVLKRAATKKSRNDLIHAEEGEDRFADLCAVGSPFARASLKSGKTESSSYFRRKEKMFRFFIRRMVKAQSFYWVVLCVVALNTLCVAMVHYNQPRRLTTTLYFAEFVFLGLFLTEMSLKMYGLGPRSYFRSSFNCFDFGVIVGSVFEVVWAAIKPGSSFGISVLRALRLLRIFKVTKYWSSLRNLVVSLLNSMKSIISLLFLLFLFIVVFALLGMQLFGGQFNFQDETPTTNFDTFPAAILTVFQILTGEDWNAVMYHGIESQGGVSKGMFSSFYFIVLTLFGNYTLLNVFLAIAVDNLANAQELTKDEEEMEEAANQKLALQKAKEVAEVSPMSAANISIAARQQNSAKARSVWEQRASQLRLQNLRASCEALYSEMDPEERLRFATTRHLRPDMKTHLDRPLVVELGRDGARGPVGGKARPEAAEAPEGVDPPRRHHRHRDKDKTPAAGDQDRAEAPKAESGEPGAREERPRPHRSHSKEAAGPPEARSERGRGPGPEGGRRHHRRGSPEEAAEREPRRHRAHRHQDPSKECAGAKGERRARHRGGPRAGPREAESGEEPARRHRARHKAQPAHEAVEKETTEKEATEKEAEIVEADKEKELRNHQPREPHCDLETSGTVTVGPMHTLPSTCLQKVEEQPEDADNQRNVTRMGSQPPDPNTIVHIPVMLTGPLGEATVVPSGNVDLESQAEGKKEVEADDVMRSGPRPIVPYSSMFCLSPTNLLRRFCHYIVTMRYFEVVILVVIALSSIALAAEDPVRTDSPRNNALKYLDYIFTGVFTFEMVIKMIDLGLLLHPGAYFRDLWNILDFIVVSGALVAFAFSGSKGKDINTIKSLRVLRVLRPLKTIKRLPKLKAVFDCVVNSLKNVLNILIVYMLFMFIFAVIAVQLFKGKFFYCTDESKELERDCRGQYLDYEKEEVEAQPRQWKKYDFHYDNVLWALLTLFTVSTGEGWPMVLKHSVDATYEEQGPSPGYRMELSIFYVVYFVVFPFFFVNIFVALIIITFQEQGDKVMSECSLEKNERACIDFAISAKPLTRYMPQNRQSFQYKTWTFVVSPPFEYFIMAMIALNTVVLMMKFYDAPYEYELMLKCLNIVFTSMFSMECVLKIIAFGVLNYFRDAWNVFDFVTVLGSITDILVTEIAETNNFINLSFLRLFRAARLIKLLRQGYTIRILLWTFVQSFKALPYVCLLIAMLFFIYAIIGMQVFGNIALDDDTSINRHNNFRTFLQALMLLFRSATGEAWHEIMLSCLSNQACDEQANATECGSDFAYFYFVSFIFLCSFLMLNLFVAVIMDNFEYLTRDSSILGPHHLDEFIRVWAEYDPAACGRISYNDMFEMLKHMSPPLGLGKKCPARVAYKRLVRMNMPISNEDMTVHFTSTLMALIRTALEIKLAPAGTKQHQCDAELRKEISVVWANLPQKTLDLLVPPHKPDEMTVGKVYAALMIFDFYKQNKTTRDQMQQAPGGLSQMGPVSLFHPLKATLEQTQPAVLRGARVFLRQKSSTSLSNGGAIQNQESGIKESVSWGTQRTQDAPHEARPPLERGHSTEIPVGRSGALAVDVQMQSITRRGPDGEPQPGLESQGRAASMPRLAAETQPVTDASPMKRSISTLAQRPRGTHLCSTTPDRPPPSQASSHHHHHRCHRRRDRKQRSLEKGPSLSADMDGAPSSAVGPGLPPGEGPTGCRRERERRQERGRSQERRQPSSSSSEKQRFYSCDRFGGREPPKPKPSLSSHPTSPTAGQEPGPHPQGSGSVNGSPLLSTSGASTPGRGGRRQLPQTPLTPRPSITYKTANSSPIHFAGAQTSLPAFSPGRLSRGLSEHNALLQRDPLSQPLAPGSRIGSDPYLGQRLDSEASVHALPEDTLTFEEAVATNSGRSSRTSYVSSLTSQSHPLRRVPNGYHCTLGLSSGGRARHSYHHPDQDHWC

>XP_021331856.1|Danio_rerio_Cav2.2

MARFEDDLPTRYGGGGGGSPAGPGRGATRQPGPPGGPRVYKQTMAQRARTMAIYNPIPVKQNCLTVNRSLFIFSEDNIIRKYAKKITEWPPFEYMILATIIANCIVLGLEQHLPALDKTPMSKRLDDTEPYFIGIFCFEAGIKIIALGFAFHKGSYLRNGWNVMDFVVVLTGILTIVGPDFDLRTLRAVRVLRPLKLVSGIPSLQVVLKSIMKAMVPLLQIGLLLFFAILMFAIIGLDFYMGKFHRTCFRTDTGEQVDEFPCGLESPAWTCENGTECREYWIGPNFGITNFDNILFAVLTVFQCITMEGWVDILYNANDASGNTWNWLYFIPLIIIGSFFMLNLVLGVLSGEFAKERERVEKRQEFLKLRRQQQIERELTGYLEWICKAEEVMLAEEDKNAEDKDDVAWYKRKCNNPGGFTRSLGADFGLPVLKRAKKSKNDLINAEEGEDHFTDISSVAPQGSPFTRTSVKSSKIESLSYFRRKEKRFRFFIRRMVKAQSFYWTVLCIVGLNTLCVAIVHYDQPEWLTYALYLAEFVFLGLFLIEMSLKMYGLGPRTYFHSSFNCFDFGVIVGSIFEVIWAAVKPGASFGISVLRALRLLRIFKVTKYWNSLRNLVVSLLNSMKSIISLLFLLFLFIVVFALLGMQLFGGQFNFEDETPTTNFDTFPAAILTVFQILTGEDWNAVMYHGIESQGGVHRGMFCSVYFIVLTLFGNYTLLNVFLAIAVDNLANAQELTKDEEEQEEAISKKLALQKAKEVKEVSPMSAANISITAKEQQRSLKMMSVWEQRTTQIRKHMLASSEALYQEELNPRLMSNLPLRTDIKTHLDRPLVVEPRCDGPISPHSGWDKPQDLQGEGSTPEERVPPVFEPPQSHPPRKHHRHRERLSNETNENGDTGHAKEGRHHVHHSRSKEHDGTRCKEGKGDRSRSREGGRRHHHQSSVDEGVGGGEREHRHHHSHRHNREGNGVVSAGGKSERRSRQKEGSRSGTREGERCSRGENGGNGGRRRHRPRTSKAQSTLEGEEQHVNGEEEGKSQSHRHMDMGGDSMAVNPHQPSVGSRWCLEKPEDSDNKRNIGQTGGMSTHIPVTVTSPPGETTLIPMNSIDSETVPMTEKNLEDLNQSSIRPILPFSSMFIFNPTNPVRRLCHYIVSLRYFEMCILVVIAMSSIALAAEDPVQANAPRNNVLKYLDYVFTGVFTFEMVIKMVDLGLILHPGSYFRDLWNILDFIVVSGALVAFAFSGTKGKDISTIKSLRVLRVLRPLKTIKRLPKLKAVFDCVVNSLKNVLNILIVYILFMFIFAVIAVQLFKGKFFYCTDESKGLEKDCRGLFLDYDKDDVTAQPREWKKYEFHYDNVLWAFLTLFTVSTGEGWPTVLKHSVDATFEDQGPSPGYRIEMSIFYVVYFVVFPFFFVNIFVALIIITFQEQGDKVLSECSLEKNERACIDFAINAKPLTRYMPQNQQSFQYRLWKFVVSPPFEYSIMIMIALNTVVLMMKFHGAPDFYEAMLKYLNIVFTVLFSLECILKIIAFGPLNYLKDAWNVFDFVTVLGSITDILVTEINTTERQLNFSFLRLFRAARLIKLLRQGYTIRILLWTFVQSFKALPYVCLLIAMLFFIYAIIGMQVFGNIDLNDDTAINRHNNFRTFLQALMLLFRSATGEAWHDIMLSCLSERTCDPSSGTLGKECGSDFAYFYFVSFIFLCSFLMLNLFVAVIMDNFEYLTRDSSILGPHHLDEFIRVWAEYDPAACGRISYKDMYNLLRIISPPLGLGKNCPNRVAYKRLVKMNMPIADDNSVHFTSTLMALIRTALEIKLASGMVAQRLSDAELKKELSTVWPNLSQKTMDLLVTPHKPNELTVGKVYAALMIFDYYKQNRAKRLQMQQQQLKEQAGPGSQNKVSALLEPILPLTHMQDQPVNGVDSASECQAQTRPSSTTLNNGRTTLDNTIKGSSSWASEKSKEVQKSKRRPMSRGQSEDASNANTAQESVEMRKMENSANTVTSGGLEGQGRAASMPKLNAEMQRSHSRQSPGTLLEPIPDSPMRRSASTFAPQRPHEVNLNDYILEKPVQERHHHHHHHHRCHHRRDRDKKQRSLDRSPTGHHKTTGTATDSVEERAHDRGRSHERKHHSSSGEKQRYYSCDRYASREHCSSKSAVASCATSPSETQEANLNKQGSGWAKGSPVLLTSGASTPSRGRRQLPQTPLTPRPSVAYRTANSSPVQLLATTTSLLPPSPGRLSRGLSEHNALLLSGSISSSPAPVSRINSEPFLGPPDHPAGLQPEAFAVFQENPSPQMGRSPQTVGMPFVAIPPPPPQQFRRVPNGYHFSTGPSASRGRAQQYYPEAEVEDWC

>XP_019386495.1|Crocodylus_porosus_Cav2.3

MARFGEAVAGRLGSADGGSEQSRSRPGAAAPAGGTPGAFKQTKAQRARTMALYNPIPVRQNCFTVNRSLFLFGEDNVVRKYAKKLIDWPPFEYMILATIIANCIVLALEQHLPGDDKTPMSRRLEKTEPYFIGIFCFEAGIKIVALGFVFHKGSYLRNGWNVMDFIVVLSGILATAGTHFNTHVDLRTLRAVRVLRPLKLVSGIPSLQIVLKSIMKAMVPLLQIGLLLFFAILMFAIIGLEFYSGKLHRACYANNSGELEELDPPHPCGVQGCPPGYECKEWIGPNDGITQFDNILFAVLTVFQCITMEGWTTVLYNTNDALGATWNWLYFIPLIIIGSFFVLNLVLGVLSGEFAKERERVENRRAFMKLRRQQQIERELNGYRAWIDKAEEVMLAEENKNAGTSALEVLRRATIKRNRTEAMNRDSSDEHCVDISSVGTPLARSSIKSAKVDGASYFRHKERLLRISVRHMVKSQVFYWIVLSLVALNTACVAIVHHNQPLWLTHLLYYAEFLFLGLFLLEMSLKMYGMGPRLYFHSSFNCFDCGVTVGSIFEVVWAIFRPGTSFGISVLRALRLLRIFKITKYWASLRNLVVSLMSSMKSIISLLFLLFLFIVVFALLGMQLFGGRFNFMDGTPSANFDTFPAAIMTVFQILTGEDWNEVMYNGIRSQGGVRSGMWSSIYFIVLTLFGNYTLLNVFLAIAVDNLANAQELTKDEQEEEEAFNQKHALQKAKEVSPMSAPNMPTVERDRRRRHHMSMWEPRSSQLRERRRRHHMSVWEQRTSQLRRHMQMSSQEGINKDEPPLLNPHASIFRRKKPGENVASDKPEEEQGGKAEQPQVEGLEQQTTGTNLSASEDKRSPSPRAKRERDQWHQKSCHGNCDPTEQDGSGGGGSGIEDRARLRQSQRRSRHRRVRTEGKEPAGALESQSTSQDAGLEDANPAEGQEKRNEGGKEENLIQGEQADEELKRTNGIPASEAEVPGATKDRSPPKVPHELNQGNNVSLTEQDCSSLDTSDQALLGSLQMEMSRAISRSEPDLSSITANTEKATESTTIMIDVQDSTVVQISNKTDGEASPLKEAETKEDEEEAEKKKRKKEKSSETGKAMVPHSSMFIFSTTNPVRRACHYIVNLRYFEMCILLVIAASSIALAAEDPVLTNSDRNKVLRYFDYVFTGVFTFEMVIKMIDQGLILQDGSYFRDLWNILDFIVVVGALVAFALANALGTNKGRDIKTIKSLRVLRVLRPLKTIKRLPKLKAVFDCVVTSLKNVFNILIVYKLFMFIFAVIAVQLFKGKFFYCTDSSKDTEKDCIGNYVDHEKNKMEVKCREWKRHEFHYDNIIWALLTLFTVSTGEGWPQVLQHSVDVTEEDRGPSRSNRMEMSIFYVVYFVVFPFFFVNIFVALIIITFQEQGDKMMEECSLEKNERACIDFAISAKPLTRYMPQNRHTFQYRVWHFVVSPSFEYTIMAMIALNTVVLMMKYYSAPYTYELALKYLNIAFTMVFSLECVLKIIAFGFLNYFRDTWNIFDFITVIGSITEIILTDTKLVNTSGFNMSFLKLFRAARLIKLLRQGYTIRILLWTFVQSFKALPYVCLLIAMLFFIYAIIGMQVFGNIKLDEGSHINRHNNFRSFLGSLMLLFRSATGEAWQEIMLSCLEGKGCEPDTTATSGQNENERCGTDLAYVYFVSFIFFCSFLMLNLFVAVIMDNFEYLTRDSSILGPHHLDEFVRIWAEYDRAACGRIHYTEMYEMLTLMSPPLGLGKRCPSKVAYKRLVLMNMPVAEDMTVHFTSTLMALIRTALDIKIAKGGADWQQLDSELQKEILTIWPHLSQKMLDLLVPMPKTSDLTVGKIYAAMMIMDYYKQSKAKKQKQQLEEQKNAPMFQRMEPSSLPQEIISNAKALPYLQQDTLSGLSSRSGFPSLSPLSPQEIFQLACMDPTHGQYQEHQSLEPEVREFKRVQPSNRGNYLPVDTQERAVSGRASSMPRLTVDPQVVTDTSSMRRSFSTIRDKRTNSSWLDEFSMERSSENTYKSRRRSYHSSLQLTARRLNADSGHRSDTHRSGGRERGRSKERKHLLSPDISRCNSEERSPQADCESPERRRQSRSPSEGRSQTPNRQGTGSLSESSIPSISDTSTPRRGRRQLPPVPPKPRPFISYTSMMQHAGDTFPQPDESEGGSPLLSKALETNDTCLTESSSSLQGKQSRHSTPQRYISEPYLALHDDSHASDCGEEETLTFEAAVATSLGRSNTIGSAPPMRHSWQMPNGHYRRRRRGAAQGVMCGASIDVLSDTEEDDKC

>XP_015145962.1|Gallus_gallus_Cav2.3

MLGLGTCSVSAPGRAAEPAASPHYALALPEEEMWREHRLALMHSTQSCNLLEPQSLAEVLVSRATSFDALYQPRDADGDSGSALDLGLGTYVPISPDVIKRRRGGLIEQRDIIKAHEAHKMQSTPQARRKEWEMARFGEAVAGRLGSGDGGSEQNRSRQGAAAPAGGSPGAFKQTKAQRARTMALYNPIPVRQNCFTVNRSLFLFGEDNIVRKYAKKLIDWPPFEYMILATIIANCIVLALEQHLPEDDKTPMSRRLEKTEPYFIGIFCFEAGIKIVALGFVFHKGSYLRNGWNVMDFIVVLSGILATAGTHFNTHVDLRTLRAVRVLRPLKLVSGIPSLQIVLKSIMKAMVPLLQIGLLLFFAILMFAIIGLEFYSGKLHRACYTNNSGELEELDPPHPCGVQGCPPGYECREWIGPNDGITQFDNILFAVLTVFQCITMEGWTTVLYNTNDALGATWNWLYFIPLIIIGSFFVLNLVLGVLSGEFAKERERVENRRAFMKLRRQQQIERELNGYRAWIDKAEEVMLAEENKNAGTSALEVLRRATIKRNRTDAMNRDSSDEHCVDISSVGERRLAQRRPLHPTLGNPLSRSGLKGVDGASYLRHKERLLRISVRHMVKSQVFYWIVLSLVALNTACVAIVHHNQPAWLTHFLYYAEFLFLGLFLLEMSLKMYGMGPRLYFHSSFNCFDCGVTVGSIFEVVWAIFRPGTSFGISVLRALRLLRIFKITKYWASLRNLVVSLMSSMKSIISLLFLLFLFIVVFALLGMQLFGGGFNFIDGTPSANFDTFPAAIMTVFQILTGEDWNEVMYNGIRSQGGVRSGMWSSIYFIVLTLFGNYTLLNVFLAIAVDNLANAQELTKDEQEEEEAFNQKHALQKAKEVSPMSAPNMPAIERDRRRRHHMSMWEPRSSHLRERRRRHHMSVWEQRTSQLRRHMQMSSQEGINKDEPPLINPHASIFRRKKPGDGVALEKCDEEQGGKGERPPAEGPEQPAPGTNPGGGEDRKSPSPRARRDKEMWQHKACHGNCEPGEEGAGGGIEERARMRQSQRRSRHRKARMEGKETTGALESRAGSQEMGLEEPCPAEGMQDGERRGDSAAALDLIQGELEASGDPTRTNGVPAEEGSLPASAPEPSRGMEGSLGEQDCSSPDTSEQALLGGAALGASRTVSHSEPDLSSVTANTEKATESTTIMIDVQDSTVVQISNKTDGEASPLKEAESKEDEEEMEKKKRKKEKSETGKAMVPHSSMFIFSTTNPVRRACHYIVNLRYFEMCILLVIAASSIALAAEDPVLTNSDRNKVLRYFDYVFTGVFTFEMVIKMIDQGLILQDGSYFRDLWNILDFIVVVGALVAFALANALGTNKGRDIKTIKSLRVLRVLRPLKTIKRLPKLKAVFDCVVTSLKNVFNILIVYKLFMFIFAVIAVQLFKGKFFYCTDSSKDTEKDCIGNYVDHEKNKMEVKCREWKRHEFHYDNIIWALLTLFTVSTGEGWPQVLQHSVDVTEEDRGPSRSNRMEMSIFYVVYFVVFPFFFVNIFVAFIIITFQEQGDKMMEECSLEKNERACIDFAISAKPLTRYMPQNRHTFQYRVWHFVVSPSFEYTIMAMIALNTVVLMMKYYSAPYTYELALKYLNIAFTMVFSLECVLKIIAFGFLNYFRDTWNIFDFITVIGSITEIILTDTKLVNTSSFNMSFLKLFRAARLIKLLRQGYTIRILLWTFVQSFKALPYVCLLIAMLFFIYAIIGMQVFGNIKLDEESHINRHNNFRSFLGSLMLLFRSATGEAWQEIMLSCLEGKGCEPDTTATSGQNENERCGTDLAYVYFVSFIFFCSFLMLNLFVAVIMDNFEYLTRDSSILGPHHLDEFVRIWAEYDRAACGRIHYTEMYEMLTLMSPPLGLGKRCPSKVAYKRLVLMNMPVAEDMTVHFTSTLMALIRTALDIKIAKGGADWQQLDSELQKEILTIWPHLSQKMLDLLVPMPKTSDLTVGKIYAAMMIMDYYKQSKAKKQRQQLEEQKNAPMFQRMEPSSLPQEIISNAKALPYLQQETLTGLSSRSGFPSLSPLSPQEIFQLACMDPAHGQFQEHQSLEPEVREFQRVQPSNRGSYLPVDTQDHAVSGRASSMPRLTVDPQVVTDTSSMRRSFSTIRDKRTNTSWLDEFSMERSSDNTYKSRRRSYHSSLQLSARRLNADSGHRSEGHRSGGRERGRSKERKHLLSPDISRCNSEERSPQAHDESPERRRESRSPSEGRSQTPNRQGTGSLSESSIPSISDTSTPRRGRRQLPPVPPKPRPLLSYASMLRHAAGDASPPPDETEGGSPLLSKEPNATGLTESSSSPAGKPSRPSTPQRYISEPYLALHDDSHASDCGEEETLTFEAAVATSLGRSNTIGSAPPLRHSWQMPNGHYRRRRRGAGQGVMCGASIDVLSDTEEDDKC

>XP_008164802.1|Chrysemys_picta_bellii_Cav2.3

MQATQSCSLLEPEALGDALVSRTASFDILYEPGSREGEASLGFEGSSTFDLGLSQYIPVSPDVIKRRRGGLIEQRDIIKAHEAHKMQSTPQARRKEWEMARFGEAVAGRLGSGDAGSEQNRIRQGPPAPSGATPGAFKQTKAQRARTMALYNPIPVRQNCFTVNRSLFLFGEDNMVRKYAKKLIDWPPFEYMILATIIANCIVLALEQHLPEDDKTPMSRRLEKTEPYFIGIFCFEAGIKIVALGFVFHKGSYLRNGWNVMDFIVVLSGILATAGTHFNTHVDLRTLRAVRVLRPLKLVSGIPSLQIVLKSIMKAMVPLLQIGLLLFFAILMFAIIGLEFYSGKLHRACYTNNSGELEELDPPHPCGVQGCPSGYECKEWIGPNDGITQFDNILFAVLTVFQCITMEGWTTVLYNTNDALGATWNWLYFIPLIIIGSFFVLNLVLGVLSGEFAKERERVENRRAFMKLRRQQQIERELNGYRAWIDKAEEVMLAEENKNAGTSALEVLRRATIKRNRTEAVNRDSSDEHCVDISSVGTPLARGSIRSAKVDGASYFRHKERLLRISVRHMVKSQVFYWTVLSLVALNTACVAIVHHNQPPWLTHLLYYAEFLFLGLFLLEMSLKMYGMGPRLYFHSSFNCFDCGVTVGSIFEVVWAIFRPGTSFGISVLRALRLLRIFKITKYWASLRNLVVSLMSSMKSIISLLFLLFLFIVVFALLGMQLFGGRFNFMDGTPSANFDTFPAAIMTVFQILTGEDWNEVMYNGIRSQGGVRSGMWSSIYFIILTLFGNYTLLNVFLAIAVDNLANAQELTKDEQEEEEAFNQKHALQKAKEVSPMSAPNMPSIERDRRRRHHMSMWEPRSSQLRDRRRRHHMSVWEQRTSQLRRHMQMSSQEGLNKDAPPLINPHASIFRRKMPGENVTLEKSEEDHGGSAEHPQAECLDQPIAGTNSCPSTKEDQKSPSPRAKRDREQWHQKSCHGNCDLTEQEGGGRGGGGGGAGIEDRARLRQSQRRSRHRRVRTEGKEPAGALESRSASQEAGLEEANPTEGEQARNEGGKEDTNQEEPLAGEELNRTNEIPAGDAAFAGAMQEMSPLKVPHELNEEKNTSLTEQDCSGLDTSDQALLGSLQMEMSRAVSRSEPELSSVTANTEKATESTTIMIDVQDSTVVQISNKTDGEASPLKEAETKEDEEKEKKKRKKEKSSETGKAMVPHSSMFIFSTTNPVRRACHYIVNLRYFEMCILLVIAASSIALAAEDPVLTNSDRNKVLRYFDYVFTGVFTFEMIIKMIDQGLILQDGSYFRDLWNILDFIVVVGALVAFALANTLGTNKGRDIKTIKSLRVLRVLRPLKTIKRLPKLKAVFDCVVTSLKNVFNILIVYKLFMFIFAVIAVQLFKGKFFYCTDSSKDTEKDCIGNYVDHEKNKMEVKCREWKRHEFHYDNIIWALLTLFTVSTGEGWPQVLQHSVDVTEEDRGPSRSNRMEMSIFYVVYFVVFPFFFVNIFVALIIITFQEQGDKMMEECSLEKNERACIDFAISAKPLTRYMPQNRHTFQYRVWHFVVSPSFEYTIMAMIALNTVVLMMKYYSAPYTYELALKYLNIAFTMVFSLECVLKIIAFGFLNYFRDTWNIFDFITVIGSITEIILTDTKLVNTSSFNMSFLKLFRAARLIKLLRQGYTIRILLWTFVQSFKALPYVCLLIAMLFFIYAIIGMQVFGNIKLDEESHINRHNNFRSFLASLMLLFRSATGEAWQEIMLSCLQGKGCEPDTTATSGQHENEHCGTDLAYVYFVSFIFFCSFLMLNLFVAVIMDNFEYLTRDSSILGPHHLDEFVRIWAEYDRAACGRIHYTEMYEMLTLMSPPLGLGKRCPSKVAYKRLVLMNMPVAEDMTVHFTSTLMALIRTALDIKIAKGGADWQQLDSELQKEILAIWPHLSQKMLDVLVPLPKTSDLTVGKIYAAMMIMDYYKQSKAKKQKQQLEEQKNAPMFQRMEPSSLPPEILSNAKALPYLQQDTLSGLSSRSGYPSLSPLSPQEIFQLACMDSTHGQFQEQQSLEPEVREFQSVQPSNRGNYLPVDTQERAVSGRASSMPRLTVDPQVVTDTSSMRRSFSTIRDKRPNSSWLDEFSMERSSENTYKSRRRSYHSSLQLSARRLNADSGHRSETHRSGGRERGRSKERKHLLSPDISRCNSEERSPQADGESPVRRQSRSPSEGRSQTPNRQGTGSLSESSIPSISDTSTPRRGRRQLPPVPPKPRPLLSYASMMRHPGDTSPQPDESEGGSPLFSQALETNDTCLTESSSSPQGKQSRHSTPQRYISEPYLALHDDSHASDCGEEETLTFEAAVATSLGRSNTIGSAPPLRHGWQMPNGHYRRRRRGAAQGVMCGASIDVLSDTEEDDKC

>XP_020645405.1|Pogona_vitticeps_Cav2.3

MMLSLGPCSISAHILSEGGSFPAGESTELDSNPLRCALALPEEETWRQHRLGLMHSTQSCNLLEPDTLTDPLVSRTASFDALYELRSQERDSSLGLDGSSTFDVGQSQLIPVSPEVIKRRRGGLIEQRDIIKAHEAHKMQSTPQARRKEWEMARFGEAMAGRLGSGDGAAEQNRNRQGAPASSTGGTPVAYKQTKAQRARTMALYNPIPVRQNCFTVNRSLFIFGEDNIVRKYAKKLIDWPPFEYMILATIIANCIVLALEQHLPEDDKTPMSRRLEKTEPYFIGIFCFEAGIKIVALGFVFHKGSYLRNGWNVMDFIVVLSGILATAGTHFNTHVDLRTLRAVRVLRPLKLVSGIPSLQIVLKSIMKAMVPLLQIGLLLFFAILMFAIIGLEFYSGKLHRACYMNNSGKLEEMDPPHPCGVQGCPAGYECRDWIGPNDGITQFDNILFAVLTVFQCITMEGWTTVLYNTNDALGATWNWLYFIPLIIIGSFFVLNLVLGVLSGEFAKERERVENRRAFMKLRRQQQIERELNGYRAWIDKAEEVMLAEENKNSGTSALEVLRRATIKRSRTEAMNRDSSDERVDISAVGTPLARASIKSAKLDGASYFRHKERLLRISVRHMVKSQVFYWLVLSIVALNTACVAIVHHDQPPWLTHLLYYAEFIFLGLFLLEMSLKMYGMGPRLYFHSSFNCFDCGVTVGSIFEVVWAIFRPGTSFGISVLRALRLLRIFKVTKYWASLRNLVVSLMSSMKSIISLLFLLFLFIVVFALLGMQLFGGRFNFADGTPSANFDTFPAAIMTVFQILTGEDWNEVMYNGIRSQGGVSSGMWSSIYFIILTLFGNYTLLNVFLAIAVDNLANAQELTKDEQEEEEAFNQKHALQKAKEVSPMSAPNMPAIERDRRRRHHMSMWEPRSSNLRERRRRHHMSVWEQRTSQLRRHMQMSSQEGINSEDPPLLNPHASIFRRRKVLENSALEKNEESQGGRLERTGGEGQEQPISGSNPCPTIREGQRSPSPQGKREWDEWHHKSFHGNCDLNELEGGGSVGFEERARLRQSQRRSRHRRVRTEAKEHRPTAQESAPEGSGPTEGEADGEPKKNEEKEALIEKKEEASEELKTTNEAPANDADFPVPPQETNSSDSHLDLKKGCSSKDVSPADQDSSLDTSEQVLLSDLPMETGKTISRSEPDLSSVTTNMEKPTESTSIMIDVQDSAVVQISTKTDGEASPLKEAETKEEVEVEAEKKQKQKKKRSQKGKPMVPHSSMFIFSTTNPIRRACHYIVNLRYFEMCILLVIAASSIALAAEDPVLTNSDRNKVLRYFDYVFTGVFTFEMVIKMIDQGLILQDGSYFRDLWNILDFIVVVGALMAFALANALGTNKGRDIKTIKSLRVLRVLRPLKTIKRLPKLKAVFDCVVTSLKNVFNILIVYKLFMFIFAVIAVQLFKGKFFYCTDSSKDTEKDCIGNYVDHEKGKMEVKCREWKRHEFHYDNIIWALLTLFTVSTGEGWPQVLQHSVDVTEEDRGPSRSNRMEMSIFYVVYFVVFPFFFVNIFVALIIITFQEQGDKMMEECSLEKNERACIDFAISAKPLTRYMPQNRHTFQYRVWHFVVSPSFEYTIMAMIALNTVVLMMKYYSAPYTYELALKYLNIAFTMVFSLECVLKIIAFGFLNYFRDTWNIFDFITVIGSITEIILTDSKLVNTSSFNMSFLKLFRAARLIKLLRQGYTIRILLWTFVQSFKALPYVCLLIAMLFFIYAIIGMQVFGNIKLDEESHINRHNNFRSFLGSLMLLFRSATGEAWQEIMLSCLEGKGCETDTTATSGQNENEQCGTDLAYVYFVSFIFFCSFLMLNLFVAVIMDNFEYLTRDSSILGPHHLDEFVRIWAEYDRAACGRIHYTEMYEMLTLMSPPLGLGKRCPSKVAYKRLVLMNMPVAEDNTVHFTSTLMALIRTALDIKIAKGGADWQQLDSELQKEILTIWPHLSQKQLDLLVPMPKTTDLTVGKIYAAMMIMDYYKQSKAKKQRQQLEEQKNAPMFQRMEPSSLPQEIISNAKALPYLQQDTLSRLSSRSGFPSLSPLSPQEIFQLACMDPNHGEYQEQQSLEPEVREFKRVQPSNCGSYLPVDTQERAVSGRASSMPRLTVDPQVVTDPGSMRRSFSTIRDKRTNTSWLDEFSMERSSENTYKSRRRSYHSALHLSSRRLNTDSGHRSDAHRSGGRERGRSKERKHLLSPDISRCNSEERSLQADGDSPERHQSRSPSEGRSQTPNRQGTGSLSESSIPSISDTSTPRRSRRQLPPVPPKPRPFISYTAMMQHCGNTSPQPEDSEGGSPLLSGVLETNNTCLTESSSSPQGKQNLLSTPQRYISEPYLMLQDDSHASDCGEEETLTFEAAVATSLGRSNTIASGPRSRHSWQMPNGHYRRRRRGASQGVMCGASIDVLSDTEEDDKC

>XP_008107103.1|Anolis_carolinensis_Cav2.3

MARFGEAMAGRVGSGDGATEQNRNRQGAPAPPGGTPVQAYKQTKAQRARTMALYNPIPVRQNCFTVNRSLFIFGEDNIVRKYAKKLIDWPPFEYMILATIIANCIVLALEQHLPEDDKTPMSRRLEKTEPYFIGIFCFEAGIKIVALGFVFHKGSYLRNGWNVMDFIVVLSGILATAGTHFNTHVDLRTLRAVRVLRPLKLVSGIPSLQIVLKSIMKAMVPLLQIGLLLFFAILMFAIIGLEFYSGKLHRACYVNNSGELQELDPPHPCGVQGCPAGYECRDWIGPNDGITQFDNILFAVLTVFQCITMEGWTTVLYNTNDALGATWNWLYFIPLIIIGSFFVLNLVLGVLSGEFAKERERVENRRAFMKLRRQQQIERELNGYRAWIDKAEEVMLAEENKNSGTSALEVLRRATIKRNRTEVMNRDSSDERVDISSVGTPLARASIKSAKLDGANYFRHKERLLRISVRHMVKSQVFYWLVLSIVALNTACVAIVHHNQPPWLTHLLYYAEFIFLGLFLLEMSLKMYGMGPRLYFHSSFNCFDCGVTVGSIFEVVWAIFRPGTSFGISVMRALRLLRIFKVTKYWASLRNLVVSLMSSMKSIISLLFLLFLFIVVFALLGMQLFGGRFNFMDGTPSANFDTFPAAIMTVFQILTGEDWNEVMYNGIRSQGGVSSGMWSSIYFIVLTLFGNYTLLNVFLAIAVDNLANAQELTKDEQEEEEAFNQKHALQKAKEVSPMSAPNMPAIERDRRRRHHMSMWEPRSSNLRERRRRHHMSVWEQRTSQLRRHMQMSSQEGINNEDPPLLNPHASIFRRRKLLENSGPEKNEEGQGDRSERANGEGQDQQIPGSNPCPSIQENQKSPSSRPKKEWEEWHHKSFHGNCDLIEPEGGGSGGFEERARLRQSQRRSRHRRVRTEAKEHRAASQEMAPEEAVPAEGEQNVEQKKNEEEKEALIEKDQGTGEEQKITNEVSVNDAAFHVATHETSPSEGHLDLNQSKDVSPVEQDGNSPDTSEQALLGDLPMESGRTISRSEPDLSSITTNVEKATESATIMIDVQDSAVVQISNKTDGEASPLKEPETKEEEEVEVEKKKPKKKKSATGKPMVPHSSMFIFSTTNPIRRACHYIVNLRYFEMCILLVIAASSIALAAEDPVLTNSERNKVLRYFDYVFTGVFTFEMVIKMIDQGLILQDGSYFRDLWNILDFIVVVGALMAFALANALGTNKGRDIKTIKSLRVLRVLRPLKTIKRLPKLKAVFDCVVTSLKNVFNILIVYKLFMFIFAVIAVQLFKGKFFYCTDSSKDTQKDCIGNYVDHEKNLMEVKCRQWKRHEFHYDNIIWALLTLFTVSTGEGWPQVLQHSVDVTEEDRGPSRSNRMEMSIFYVVYFVVFPFFFVNIFVALIIITFQEQGDKMMEECSLEKNERACIDFAISAKPLTRYMPQNRHTFQYRVWHFVVSPSFEYTIMAMIALNTIVLMMKYYSAPYTYELALKYLNIAFTMVFSLECVLKIIAFGFLNYFRDTWNIFDFITVIGSITEIILTDSKLVNTSSFNMSFLKLFRAARLIKLLRQGYTIRILLWTFVQSFKALPYVCLLIAMLFFIYAIIGMQVFGNIKLDEESHINRHNNFRSFLGSLMLLFRSATGEAWQEIMLSCLGGKGCESDTTATSGQNGSEQCGTDLAYVYFVSFIFFCSFLMLNLFVAVIMDNFEYLTRDSSILGPHHLDEFVRIWAEYDRAACGRIHYTEMYEMLTLMSPPLGLGKRCPSKVAYKRLVLMNMPVAEDNTVHFTSTLMALIRTALDIKIAKGGAEWQQLDSELQKEILTIWPHLSQKQLDVLVPMPKTTDLTVGKIYAAMMIMDYYKQSKAKKQRQQLEEQKNAPMFQRMEPSSLPQEIISNAKALPYLQQETFSGLSSRSGFPSLSPLSPQEIFQLACMDPTHGEYQEQQSLEPEVREFKRVQPSNCGSYLPVDTQERAVSGRASSMPRLTVDPQVVTDPGSMRRSFSTIRDKRTNTSWLDEFSMEKSSENTYKSRRRSYHSALYLSSRRLNTDSGHRSDTHRSGGRERGRSKERKHLLSPDISRCNSEERSPQADGDSPERRQSRSPSEGRSQTPNRQGTGSLSESSIPSISDTSTPRRSRRQLPPVPPKPRPLISYAAMMQHCGDTSPNPDDSEGGSPLLSGTLETNNPCLTESSSSPTGKQSLLSTPQHYISEPYLMLQDDSHASDCGEEETLTFEAAVATSLGRSNTIASGPRSRHSWQMPNGHYRRRRRGASSQGVMFGAGIDVLSDTEEDDKC

>NP_001192222.1|Homo_sapiens_Cav2.3

MARFGEAVVARPGSGDGDSDQSRNRQGTPVPASGQAAAYKQTKAQRARTMALYNPIPVRQNCFTVNRSLFIFGEDNIVRKYAKKLIDWPPFEYMILATIIANCIVLALEQHLPEDDKTPMSRRLEKTEPYFIGIFCFEAGIKIVALGFIFHKGSYLRNGWNVMDFIVVLSGILATAGTHFNTHVDLRTLRAVRVLRPLKLVSGIPSLQIVLKSIMKAMVPLLQIGLLLFFAILMFAIIGLEFYSGKLHRACFMNNSGILEGFDPPHPCGVQGCPAGYECKDWIGPNDGITQFDNILFAVLTVFQCITMEGWTTVLYNTNDALGATWNWLYFIPLIIIGSFFVLNLVLGVLSGEFAKERERVENRRAFMKLRRQQQIERELNGYRAWIDKAEEVMLAEENKNAGTSALEVLRRATIKRSRTEAMTRDSSDEHCVDISSVGTPLARASIKSAKVDGVSYFRHKERLLRISIRHMVKSQVFYWIVLSLVALNTACVAIVHHNQPQWLTHLLYYAEFLFLGLFLLEMSLKMYGMGPRLYFHSSFNCFDFGVTVGSIFEVVWAIFRPGTSFGISVLRALRLLRIFKITKYWASLRNLVVSLMSSMKSIISLLFLLFLFIVVFALLGMQLFGGRFNFNDGTPSANFDTFPAAIMTVFQILTGEDWNEVMYNGIRSQGGVSSGMWSAIYFIVLTLFGNYTLLNVFLAIAVDNLANAQELTKDEQEEEEAFNQKHALQKAKEVSPMSAPNMPSIERDRRRRHHMSMWEPRSSHLRERRRRHHMSVWEQRTSQLRKHMQMSSQEALNREEAPTMNPLNPLNPLSSLNPLNAHPSLYRRPRAIEGLALGLALEKFEEERISRGGSLKGDGGDRSSALDNQRTPLSLGQREPPWLARPCHGNCDPTQQEAGGGEAVVTFEDRARHRQSQRRSRHRRVRTEGKESSSASRSRSASQERSLDEAMPTEGEKDHELRGNHGAKEPTIQEERAQDLRRTNSLMVSRGSGLAGGLDEADTPLVLPHPELEVGKHVVLTEQEPEGSSEQALLGNVQLDMGRVISQSEPDLSCITANTDKATTESTSVTVAIPDVDPLVDSTVVHISNKTDGEASPLKEAEIREDEEEVEKKKQKKEKRETGKAMVPHSSMFIFSTTNPIRRACHYIVNLRYFEMCILLVIAASSIALAAEDPVLTNSERNKVLRYFDYVFTGVFTFEMVIKMIDQGLILQDGSYFRDLWNILDFVVVVGALVAFALANALGTNKGRDIKTIKSLRVLRVLRPLKTIKRLPKLKAVFDCVVTSLKNVFNILIVYKLFMFIFAVIAVQLFKGKFFYCTDSSKDTEKECIGNYVDHEKNKMEVKGREWKRHEFHYDNIIWALLTLFTVSTGEGWPQVLQHSVDVTEEDRGPSRSNRMEMSIFYVVYFVVFPFFFVNIFVALIIITFQEQGDKMMEECSLEKNERACIDFAISAKPLTRYMPQNRHTFQYRVWHFVVSPSFEYTIMAMIALNTVVLMMKYYSAPCTYELALKYLNIAFTMVFSLECVLKVIAFGFLNYFRDTWNIFDFITVIGSITEIILTDSKLVNTSGFNMSFLKLFRAARLIKLLRQGYTIRILLWTFVQSFKALPYVCLLIAMLFFIYAIIGMQVFGNIKLDEESHINRHNNFRSFFGSLMLLFRSATGEAWQEIMLSCLGEKGCEPDTTAPSGQNENERCGTDLAYVYFVSFIFFCSFLMLNLFVAVIMDNFEYLTRDSSILGPHHLDEFVRVWAEYDRAACGRIHYTEMYEMLTLMSPPLGLGKRCPSKVAYKRLVLMNMPVAEDMTVHFTSTLMALIRTALDIKIAKGGADRQQLDSELQKETLAIWPHLSQKMLDLLVPMPKASDLTVGKIYAAMMIMDYYKQSKVKKQRQQLEEQKNAPMFQRMEPSSLPQEIIANAKALPYLQQDPVSGLSGRSGYPSMSPLSPQDIFQLACMDPADDGQFQERQSLEPEVSELKSVQPSNHGIYLPSDTQEHAGSGRASSMPRLTVDPQVVTDPSSMRRSFSTIRDKRSNSSWLEEFSMERSSENTYKSRRRSYHSSLRLSAHRLNSDSGHKSDTHRSGGRERGRSKERKHLLSPDVSRCNSEERGTQADWESPERRQSRSPSEGRSQTPNRQGTGSLSESSIPSVSDTSTPRRSRRQLPPVPPKPRPLLSYSSLIRHAGSISPPADGSEEGSPLTSQALESNNACLTESSNSPHPQQSQHASPQRYISEPYLALHEDSHASDCGEEETLTFEAAVATSLGRSNTIGSAPPLRHSWQMPNGHYRRRRRGGPGPGMMCGAVNNLLSDTEEDDKC

>XP_017206695.2|Danio_rerio_Cav2.3

MAQPEDAPKEMLSAEGKAESESAGEIKRARTMALYNPIPAKHNCLTVNRSLFIFAENNMIRKYAKRIIEWPPFEYMILATIIANCIVLSLEQHLPGEDKTPMSKRLEKTEPYFIGIFCFEAGIKLVALGFVFHKGSYLRNGWNVMDFIVVLSGILATAGSHMNIPVDLRTLRAVRVLRPLKLVSGIPSLQIVLKSIIKAMVPLLQIGLLLFFAILMFAIIGLEFYSGKLHHTCLADLDILDNETVDSSEVDFACGVRKCPDKYTCSGSWIGPNDGITQFDNILFAVLTVFQCITMEGWTAVLYNTNDALGPTWNWIYFIPLIIIGSFFVLNLVLGVLSGEFAKERERVENRRAFMKLRRQQQVERELNGYRAWIDRAEEVMLQEENKNSGRSALDVLKRATSKKNARRRGPGEEKYSEISTVAGGSRPRVGMRTPRRGPAAYLRRKERMLRFSIRRMVKTDSFYWIVLSLVALNTISVSIVHHNQPEWLTVIQYYTEFVFLGLFLAEMFLKMYGLGFRLYFHSSFNCFDCGVIVGSIFEVVWGFFRPGVSFGISVLRALRLLRIFKITKYWSSLRNLVVSLMSSMKSIISLLFLLFLFIVVFALLGMQLFGGRFIFEDYTPTNFDTFPASIMTVFQILTGEDWNEVMYNGIRSQGGVHYGMWSSIYFIVLTLFGNYTLLNVFLAIAVDNLANAQELTKEEEEEEEEFFNQRYKSREFGLSERRRRPYLYRKRAIHRGRPSPAEAEEEAAGKQAAEEQPVNAFAGRRERRKKINMSVWEQRANQLRKRRQMASREVLFGGPTEDQDTPGSDHHQQHSLSTETSPLHLPESPMSVGMPLPEPPMSITIPLPEPPESEPLSGSGLEERPSANNHHANTERRHRVARKFRAAAAAGVAEGGRHKRHRHRESRTEARNAENHQQFERQAEDGEGKTEEAQSKHMMNMCDPERQIENQEECQREFVCDLTGLSQASGHQEEDQAESQEAQLLRHKEDFSTESKYMTEVSDIENNGSYLNQDECMTETSLKRESSSQPLLEMAPMTVSFKDVEASLSDKADDDDDDDMDDDDDYEDDENGGEKSTASKPERPDSMFIFKSKNPIRRICHYVVTLRYFEMTILLVIVASSIALAAEDPVCTSSERNKVLRYFDYVFTGVFTFEMIIKMIDQGLILHDGSYFRDMWNILDFIVVVGALIAFALTNLLGNNKGRDIKTIKSLRVLRVLRPLKTIKRLPKLKAVFDCVVTSLKNVFNILIVYKLFMFIFAVIAVQLFKGKFFYCTDGSMGTQKECQGYYIDYGRDRKEVKKREWRRHEFHYDNVLWALLTLFTVSTGEGWPQVLQHSVDVTEEDHGPSRGNRMEMSIFYVIYFVVFPFFFVNIFVALIIITFQEQGDKMMEECNLEKNERACIDFAISAKPLTRYMPQNRQTLQYRLWHFVVSPSFEYTVLVMIALNTVVLMMKYYSAPTAYDIVLKHLNTAFTVLFSLECILKIMAFGFMNYFRDTWNIFDFITVLGSITEIVVDLQSVNTINMSFLKLFRAARLIKLLRQGYTIRILLWTFVQSFKALPYVCLLIAMLFFIYAIIGMQVFGNIKLNDESHINQHNNFKTFFGALMLLFRSATGESWQEIMLSCLEGKECEPDPSITPPFTSPDHEGGCGTDFAYFYFVSFIFFSSFLMLNLFVAVIMDNFEYLTRDSSILGPHHLDEFVRIWGEYDRAACGRIHYNAMYEMLTHMSPPLGLGKKCPAKIAYKRLVLMNMPVDEDMTVHFTSTLMSLIRTALEIKIARGGEDRLQLDMELQKEISFIWPHLSQKTLDLLVPINKDTDMTVGKIYASRMIMDFFKQSKAKKLRLQVEAQRAAALPLISHSAGSALTSGGFVALSPISPQELLLQPITQRTDSSEDYDASQVDESGLNGVWGEDRPPEHAFRARHKSYKAAVSHSEQTQYMEKVRGRPKEQRLLLSPKNSNSKSRSRSPSEERVHGVTRQGNSSESPVPSTSESSTPSGKRRLSQTSTCSRPHISYSPLVSHTQSSSQPTNNNEEDEEDDPAVQQMLGCCSTALWDDSAEDQDARARQQEKSHFEVDEFLGSSVRRYPSESFLALQEEEHSPDSATAMETLTFEAAVATSLGRANTVSASMRTRSRIGWQVPNGHFRKRLAQSVSLAGGETLSDVEDDKY

>XP_018670105.1|Ciona_intestinalis_Cav2

MKSSAKNNWYGVVRAVNSAKTSFLGTHEKDNEKQRAQEENETRKQAARWRKRHLSQTGFEPSSFGRVNPAFNSSSELTRFDFQNDKLGYDEAKCWEYYYSNTSMDNLKDRGESTKDLNPGSSWSIDRMTEDRNHNFAVDETYRGNNGTSNDYERHTNERSNSIYDRFGIQPQHIVDEPADISDASSWHSDLYRISPNTQRRSMKDTSPARCRRGAVIGGQKDIAVAHEKHKMARFAEEAYPNKRQRGFPGGGGGFGGPGGPKGRKGRGQGGPYKMTLAQQARTMSYYNPIPKQQNCITANRSLFVFGVDNVVRKLAKRIIEWPPFEYLILATIVANCIVLALEEHLAAGDKTPRTIRLEGTEPYFLGIFIVEAAVKILALGFVLHKDSYLRYGWNIMDFTVVVTGCVTYFDTSLGSGFQTLRAVRVLRPLKLVSGIPSLQVVLKSIMKAMVPLLQIAVLLLFFIVVCSIVGLELYMGRFHRTCYSNSTNKIIRDNQICSEDNSASYGKYVCENGSYCGDWPLGPNHGITTFDNIIFSMMTVFQCITMEGWTDILYFADDATGSIYNWAYFIMLIIVGSFFMLNLVLGVLSGEFAKERERVENRMAFLKVRKKQQMDRELDGYLEWMQKAEEVILAEEDRTLEKSAQEAHRRKAAYRKSKVDLLDSAESSFTDISATVIGNSNRFTRSNPNTKPTNTCLAWLSHKEKRFRVKCRHLVKSPVFYWIVLFLVLLNTAFLSSVHYKQPKWWEDFLYYAEFVFLGLFSGEILLKVYGLGPRTYFRSSFNIFDFVVIIGSIFEIIWSVIRPDASFGISVLRALRLLRVFKFTSAWSGLRNLVVSLMSSLRSIVSLIFLLFLFLVVFALLGMQIFGGRFSFKDGKPNSNFDSFPSAILTVFQILTGEDWNMVMYNGVEAKGGVKNGGLWWSLYFIFLVMFGNYTLLNVFLAIAVDNLANAQELTKDEEEEKDNKEAQKALRVAREIESVSPGSMENVRIKLSDSFNKNNLQRTAWERRQHEFRKQRRTLRMKRDQNGLVTDDDSDDDLYVPGERRSGKYFDISSFNKDKEGIQGAATANGISSQHGSPTKQVGNDDHIIGQSIIGGESRFLSRQNSLRVRNTPMRELGILPSSGEVAEINQTENTRENSIMADTPVAMEDGKGTVRSSGEVDEEEDNGPKPVLPYSSMFIFSPTNPIRLACHYIVNLKYFETTILVIIILSSLTLATEDPVTKDSQRNNVLKYFDYIFTAVFTFEMVMKMIDLGLVLHPGSYFHSLWNILDFVVVCSALVGFALTASNNPQLDLGVIKSLRVLRVLRPLKTIKRLPKLKAVFMCVVNAFRNVATILIVYMLFMFIFAVIAVELFKGKFFYCTDPMINVESECKGKFFYQSPESDNKEVSERQWLQHDFHYDNVLYSFLTLFVISTGEGWPEVLWHSIGSTYEDKGPERGFRMEVSIFYIVFFVVFPFFFVNIFVAFIIITFQEEGDKAMSNCSLEKNERACVDFAISSKPLTRFMPEDKHSLQYHVWKVVVSPVFEWIIMALIVLNTVVLMLKDYWRIKQSPNYENILQYCNMAFTAIFTIECIIKMAAFKPINYFRDSWNIFDFITVVGSIADVTITLVAMIQKLYLPLDQQHDMSGFINLSFLRLFRAARLIKLLRQGETIRILLWTFVQSIKALPYVCLLIAMLFFIYSIIGMQLFGNIQLDPNSAINHHNNFTHIFQALMLLFRCATGEAWQSVMLACVSANCDPRAGIDKENGCGSVISYIYFTSFIFFCSFLMLNLFVAVIMDNFEYLTRDSSILGPHHLDEYIRVWAEYDPQASGYIKYTDMFTMLRHMEPPLGFGKNCPYRVAYRRLIQMNMPINEEKQVHFTTTLLALIRTSLKIKMLPQDAAGIDKTYRDQWQLDQELRQEIRLAWPNLNEKKMNLLLPPEGETQFTVGKVYGAMLVYEHWRAYKSRRDGKNPTLGQRPNIFPRMAAPATSSNNTDPQLDNPPIGDPLSSQALSMVSIVSAASANAGVHLPKGCNSLSPYGSRDTTPYSSPTTERRSKLQVPHPYKQLEPLRNNAATSQVEPVYAQCVDHDITPTIDDVTVAYQSGGFETNDYMQYTTPEDCMANNTYRTTTLADDVMTRDVISRLTTNDKATTFTLLSSFLQTSPTSPRGKTKLE

>XP_016766516.1|Apis_melifera_Cav2

MFRVFESVFFLRRESNHLNSFIISCYTMIIFLITSGLRHNEGVHRIIWMLGGVGGRHMSTRRRGSSPLVRGGAGLTGYAGPGASGNSNDVAAIPPDMQRYAGRRRRAVTTSDHKSCALVQTRIKLGDIMLAAQEAAQRDPGYASQYRRRPRLAGLSFGDWTSFGGEVPGLVDMAPGRDQGGGAGGGGGGGKGTTSLFILSEDNCIRKHTRFIIEWPPFEYAVLLTIIANCVVLALEEHLPKQDKTILAQKLEATEIYFLGIFCVEASLKILALGFVLHRGSYLRNIWNIMDFFVVVTGFITAFSQGIELDMDLRTLRAIRVLRPLKLVSGIPSLQVVLKSIIKAMAPLLQIGLLVLFAIVIFAIIGLEFYSGTLHKTCYSIRDINVIVKEGEQASPCNTDNKSEAPFGAHVCDANISTCMDHWEGPNFGITSFDNIGFAMLTVFQCITMEGWTAILYWTNDALGSTYNWIYFIPLIVLGSFFMLNLVLGVLSGEFAKEREKVENRQSFLKLRRQQQLEHELYCYLNWICKAEEVILAEERTTEEEKKHILEGRKRAEAKKKKLGKSKSTDTEEEEGDDDQDDGFSRSSSTKEKGPCKQFWLAEKRFRYWIRKSVKSQKFYWFVIVLVFFNTVCVAVEHYGQPQWLTDFLYFAEFVFLALFMLEMFIKVYALGPRTYFDSSFNRFDCVVISGSIFEVIWSEVKSGSFGLSVLRALRLLRIFKVTKYWKSLRNLVISLLSSMRSIISLLFLLFLFILIFALLGMQLFGGQFNFDSGTPPTNFNTFPIALLTVFQILTGEDWNEVMYQGIESQGGHKKGMIYSLYFIVLVLFGNYTLLNVFLAIAVDNLANAQELSAAENEEEEEDKQKQAQEIEKEIQSLQNPKDGGAPKVEICPPSPNQNFKDGKGGKQSSEEEKKQDEDDDTGPKPMLPYSSMFILSPTNPVRRAAHWVVNLRYFDFFIMVVISLSSIALAAEDPVWEDSPRNEVLNYFDYAFTGVFTVEMILKIIDLGIILHPGSYLREFWNIMDAVVVICAAVSFAFDMTGSSAGQNLSTIKSLRVLRVLRPLKTIKRVPKLKAVFDCVVNSLKNVINILIVYILFQFIFAVIAVQLFNGKFFYCSDESKYTQQDCQGQYFVFEDGALLPEPKKREWQSQFFHYDNVMAAMLTLFAVQTGEGWPQILQNSMAATYEDKGPIQNFRIEMSIFYIVYFIVFPFFFVNIFVALIIITFQEQGEAELQDGEIDKNQKSCIDFTIQARPLERYMPKERNSVKYKIWRIVVSTPFEYFIMGLIVLNTVLLMMKFHRQSDAYKNTLKYMNMCFTGMFTVECILKIAAFGVRNFFKDAWNTFDFITVIGSIVDALVIEFGERFSSMPSGGQLGEKKENFINVGFLRLFRAARLIKLLRQGYTIRILLWTFVQSFKALPYVCLLIAMLFFIYAIIGMQVFGNIALDADTSITKHNNFQSFIQGLMLLFRCATGEAWPNIMLSCVKGRPCDAKAGKQEGGCGSNIAYAYFVSFIFFCSFLMLNLFVAVIMDNFDYLTRDSSILGAHHLDEFVRIWAEYDPNATGKIHYTEMYDMLKNMDPPLGFGNKCPNRLAYKKLIRMNMPVDVDLKVNFTTTLFALIRENLNIKVRRASERNQANEELRDTIRSIWPLQAKKMLDLLIPRNEEIGRGKMTVGKIYVCLLILESWRTTRFGQIESAGQNDNDLIDNDNAGQSPAAQAQAFLNCILDVTAVNHTGNNHGAGSRANSLEHVVRPTPETKLDQHKEHREEDDEHRRQRSIRNKKVNWKMSHQYDILGSSGESSQGGGGSGVVPVVNDEDRAPELEPLLAEVPSQQDQNYYYHHHHHDQYYDTDNDLYYDHHHHHHHHHHHHHHHAHRPPSYYQHQNTYAPRSSIRYHKMELQDVVVSDSRAGSLESLTHTGKRLHPPVQPVRHPSRSPSLRRHSPGRPGYDHHGHYYHEGPGFSDTVSNVVEIQRHTHHPHPSQYNHRHRMRDYYDHCDYYDDGPWSASTSPARTPSPIHHIDRGRHYGTTSLEQRSRSPSPIGGRQPPHTHQHYHRHHPHQHSYPVLVTRRGRGRRLPPTPNKPSTLQLKPANINFPKLNASPTHGSHIHVPIPAGMQHPPPGQHLPPMQPSHCPLSFEQAVAMGRGGRLLPSPVPNGYKPQPQAKQRTPRSRHSDSDEDDWC

>XP_021710362.1|Aedes_aegypti_Cav2

MIGSVGGRHMSTRRRGSSPMVRLGGGSLMAHLEPPDGLNTPRVSPRRRRAVTTSDHKSCALIQTRLKLGDIMLAAAQEAALAGQGSNGGQFGRKGGQYLADKMGGPSSQGGQPPSGGGPTSLFILSEDNIIRKYTRFIIEWPPFEYAVLLTIIANCVVLALEEHLPHGDKTLLAQKLEKTEAYFLGIFCVEASLKILALGFVMHKHSYLRNIWNIMDFFVVVTGFITLFPQEGPEVDLRTLRAIRVLRPLKLVSGIPSLQVVLKSIIKAMAPLLQIGLLVLFAIVIFAIIGLEFYSGALHRSCYSLEDISQIVKEGEFPTPCNADNDTIAPTGAYVCNSSDSMCVEQWEGPNFGITSFDNIGFAMLTVFQCITMEGWTAILYWTNDALGSTFNWIYFVPLIVLGSFFMLNLVLGVLSGEFAKEREKVENRQEFLKLRRQQQLEKELNGYVEWICKAEEVILAEERTTEEEKMHIMEARRRAAAKRKKLRNLGKSKSTDTEEDDPDDDCGDDGYLKSKVKTQGKCIGFWRAEKRFRFWIRHTVKTQWFYWFVIVLVFFNTVCVAVEHYGQPNWLTQFLYYAEYVFLGLFMMEMWIKMYALGPRIYFESSFNRFDCVVISGSIFEVVWSEVKGGSFGLSVLRALRLLRIFKVTKYWSSLRNLVISLLNSMRSIISLLFLLFLFILIFALLGMQLFGGQFNLPDGTPPTNFNTFPIALLTVFQILTGEDWNEVMYQGIESQGGHKKGMIYSLYFIILVLFGNYTLLNVFLAIAVDNLANAQELTAAEEEQMEENKEKQQMELDKEMGALHLQTDGSPPRVEVSSPSPTRGNGSKANKKEEDKEEDDDDIPDGPKPMLPYSSMFVLSPTNPIRCAAHWVVNLRYFDFFIMVVISLSSIALAAEDPVEEDSPRNKILNFFDYAFTGVFTIEMLLKIVDLGVILHPGSYLREFWNIMDAVVVICAAVSFGFDMTGSSAGQNLSTIKSLRVLRVLRPLKTIKRVPKLKAVFDCVVNSLKNVINILIVYILFQFIFAVIAVQLFNGKFFYCTDDSKHTSEECKGSFFVYDGTDQLPRREVREWKTQSFHYDNVATAMLTLFAVQTGEGWPQVLQNSMAATYEDKGPIQNFRIEMSIFYIVYFIVFPFFFVNIFVALIIITFQEQGEAELQDGEIDKNQKSCIDFTIGARPLERYMPNKRNSFKYKVWRIVVSTPFEYFIMMLIVFNTLLLMMKYHNQGKEFEKSLKYLNMGFTGMFSVETILKIIGFGVKNFFKDPWNIFDFITVIGSIIDALVLELGENSFNVGFLRLFRAARLIKLLRQGYTIRILLWTFVQSFKALPYVCLLIAMLFFIYAIIGMQVFGNIELEPESAITRHNNFRSFVQGLMLLFRCATGESWPNIMLACLKGRPCDPRAGKSNETCGSTLAYAYFVSFIFFCSFLMLNLFVAVIMDNFDYLTRDSSILGAHHLDEFVRIWAEYDPNATGKIHYTEMYDMLKNMDPPLGFGNKCPNRLAYKKLIRMNMPLDAEGKVGFTTTLFALIRENLNIKMRTAEEMDQADQELRQTISHIWPLQAKKLLDLLVPRKDELNTGKLTVGKIYAGLLILESWRNTKFGQIESDLPELQDVSRQPSLESLTGGDGHLHPGHAYHNGHRRSPSLRRESPLVRTPSSRRRHHDVGFSDTVSNVVEIQKEEHKRGRHHFGYAHRHNRGSWSASTSPARSPSPNRFDVHSSAQHRSRKDRSKNHPAHQTYDTTILCERSRSPSPASLLQELRDRDPTRRKYRNGMGPPGHVQHSYPVLVARRQGQGRRLPPTPCKPSTLQLKQTNINFPKLNASPTHTLHSTHTTPHSIHSLPPTREFLRERDRERDRERERDLYYRERDRERDRERYRSSREREARDYGPLHYEFRDRERELYEREIEREFEREYERMEHQMPLSYEQALAMGRTGGRVLPSPILNGYKPKGGLHSSRHSDSDDEDWC

>AFH07350.1|Drosophila_melanogaster_Cav2

MGGPKKEENPPGGGPTSLFILTEDNPIRKYTRFIIEWPPFEYAVLLTIIANCVVLALEEHLPGGDKTVLAQKLEKTEAYFLCIFCVEASLKILALGLVLHKHSYLRNIWNIMDFFVVVTGFMTQYPQIGPEVDLRTLRAIRVLRPLKLVSGIPSLQVVLKSIIKAMAPLLQIGLLVLFAIVIFAIIGLEFYSGALHKTCYSLEDPNKLVKEGESETPCNTDNILEKATGSFVCNNTTSMCLEKWEGPNSGITSFDNIGFAMLTVFQCITMEGWTAILYWTNDALGSAFNWIYFVPLIVIGSFFMLNLVLGVLSGEFSNERNRVERRMEFQKCRFRAMFQTAMVSYLDWITQAEEVILAEERTTEEEKMHIMEARRRNAAKRKKLKSLGKSKSTDTEEEEAEEDYGDDGYLKTRSKPQGSCTGFWRAEKRFRFWIRHTVKTQWFYWFVIVLVFLNTVCVAVEHYGQPSFLTEFLYYAEFIFLGLFMSEMFIKMYALGPRIYFESSFNRFDCVVISGSIFEVIWSEVKGGSFGLSVLRALRLLRIFKVTKYWSSLRNLVISLLNSMRSIISLLFLLFLFILIFALLGMQLFGGQFNLPGGTPETNFNTFPIALLTVFQILTGEDWNEVMYQGIISQGGAQKGMIYSIYFIVLVLFGNYTLLNVFLAIAVDNLANAQELTAAEEEQVEEDKEKQLQELEKEMEALQADGVHVENGDGAVAPSKSKGKKKEEEKKEEEEVTEGPKPMLPYSSMFILSPTNPIRRGAHWVVNLPYFDFFIMVVISMSSIALAAEDPVRENSRRNKILNYFDYAFTGVFTIEMLLKIVDLGVILHPGSYLREFWNIMDAVVVICAAVSFGFDMSGSSAGQNLSTIKSLRVLRVLRPLKTIKRVPKLKAVFDCVVNSLKNVVNILIVYILFQFIFSVIGVQLFNGKFFYCTDESKHTSAECQGSYFKYEEDELLPKQELRVWKPRAFHYDNVAAAMLTLFAVQTGEGWPQVLQHSMAATYEDRGPIQNFRIEMSIFYIVYFIVFPFFFVNIFVALIIITFQEQGEAELQDGEIDKNQKSCIDFTIGARPLERYMPKNRNTFKYKVWRIVVSTPFEYFIMMLIVFNTLLLMMKYHNQGDMYEKSLKYINMGFTGMFSVETVLKIIGFGVKNFFKDPWNIFDLITVLGSIVDALWMEFGHDSNSINVGFLRLFRAARLIKLLRQGYTIRILLWTFVQSFKALPYVCLLIAMLFFIYAIIGMQVFGNIKLGTVENSITRHNNFQSFIQGVMLLFRCATGEAWPNIMLACLKGKACDDDAEKAPGEYCGSTLAYAYFVSFIFFCSFLMLNLFVAVIMDNFDYLTRDSSILGAHHLDEFVRIWAEYDPNATGKIHYTEMYDMLKNMDPPLGFGNKCPNRLAYKKLIRMNMPLDDELRVQFTTTLFALIRENLSIKMRAPEEMDQADMELRETITNIWPLQAKKMLNLLVPPSDQLNKGKLSVGKIYAGFLILESWRSTRFGQLDSGMPKQSFFNCLLDMAALDKGGSRQGSISFEPNGEGAANSQTHLLASTHHHHANGDAEHNSLATLARRSTIRKRSVRNKKMLELQDASRHPSQESLTGADAGHLHPGHSYMNGHRRSPSLRHNGSPLARSPSPRRRGHQYIHHDIGFSDTVSNVVEMVKETRHPRHGNSHPRYPRGSWSASTSPARSPSPSRYGGHLSRSKRTQLPYPTYGTTSLCQRSRSPSPARLQEMRERDRLGYGIDMGVTHVQHSYPTLASRRAGIGRRLPPTPSKPSTLQLKPTNINFPKLNASPTHTHHSTPHSVHSLPHHRDLLRDPRDMYYSSRERERDRERLRDRDRDRDRDRLHEYDLRYEYRDRERELYERERDREREVERERLEYIAPLSFEQALAMGRTGRVLPSPVLNGFKPKSGLNPRHSDSDEEDWC

>XP_018019172.1|Hyalella_azteca_Cav2

MGGKKGPPLPPGAQGPSSLFILSDTNIFRKYTKFIIEWPPFEYAVLLTIIANCIVLALEEHLPLSDKTVLAQELEKTEPYFLFIFCIESSLKILALGFVLHPNSYLRNIWNIMDFVVVVTGFVTLSMQEDLGVDLRTLRAIRVLRPLKLVSGIPSLQVVLKSIIKAMAPLLQIGLLVLFAIVIFAIIGLDFYCGALHKTCYSLEDLDVIITEGEAATPCYADGLENSNETTSPSGAFLCDPSVSICLERWDGPNFGITSFDNIGFAMLTVFQCITQEGWTSILYWTNDSVGSTFNIFYFVPLIVIGSFFMLNLVLGVLSGEFSNERERVERKEHFRKMRMRKLFTEDFLNYFEWIGRAEEVILAEERTTDEEKAHIMEARRRAAAKRKKLKNLGKSKSTDTDEEDNDIEEEDGQYLKYRMRKQGACKGFWKAEKRFRFRIRRTVKQQWFYWFVIVLVFLNTACVASEHYAQPLWLADFLYYAEYAFLGLFLTEMLVKIYALGPRIYFESSFNRFDCVVISGSIFEVIWSGIYDESFGFSVLRALRLLRIFKVTNSTYGLPGLFLNFNRFAIALLTVFQILTGEDWNEVMYQGIKSQGGTTSTGMIYSMYFIILVVFGNYTLLNVFLAIAVDNLANAQELTAAEEEKEEEDKEKQQAEMEKELEALQQSAAGDGPPTIDASKDADPDIDFADDSGLGPRPMLPYSSMFILSPTNPIRRAAHWVVNLRYFDFFIMVVISLSSMALASEDPVEEGSKWNTYLTYFDYAFTGVFAVEMILKVVDLGVIFHPGSYLRDLWNIMDSVVVICAAVSFCFDMMGSTTGQNLSTIKSLRVLRVLRPLKTIKRVPKLKAVFDCVVNSLKNVFNILIVYILFQFIFAVIAVQLFNGKFFYCTDLSIMTKAECQGEYFWYDNYDQPPEVRPRQWKQQSFHYDNVMFAMLTLFAVQTGEGWPQVLQNSMAATYENHGPQPHFAIEMSIFYIVYFIVFPFFFVNIFVALIIITFQEQGEAELQDGELDKNQKSCIDFAIQAKPLERYMPKERLSLKYKTWKVVVSTPFEYLIMTLIVLNTILLMMKFHNQSKIYQRSLHYLNSFFTALFTIECMLKISAFGIRNYFKDNWNTFDFICVVGSIIDALVVEFGNDSSNSYINLRFLRLFRAARLIKLLRQGDTIRILLWTFIQSFKALPYVCLLIVILFFIYAIIGMQVFGAILLDPMTSYTRHNNFRNFWQGLMLLFRCATGESWQSIMLSCIDGVPCDPRALGDDKEETCGSNIAYAYFVSFIFFCSFLMLNLFVAVIMDNFDYLTRDSSILGAHHLDEYIRIWAEYDPNATGRIHYTEMYEMLRNMDPPLGFGNKCPHRLAYKKLIRMNMPMDENLKVHFTTTLFALIRENLAIKMRAPHEMDTADSELRQTIKKIWPMQAKKVIDLIIPPDDQLVYEKLTVGKVYAALLILENWNTTRFGQIQPTGTMGLELAEVVQAASRRGSLASEDGGAHGLSHFGTPRGKHLRPDEFPSRYPRSRSPSPARGFQSPTVLHRRARSPTPIRRSPSPSRAGARYDALSDAVSDVVDISKRGRLRGRGPSPPGRGGDSFSGRAWNARNVRSASNSPDRFYQTTRLDARSRSPSPQSSHAARRGRLLHYYGTASLERRSRSPSPTHRQRAGSSYPTIPQRRGGGRRLPQTPNKPSTLHLFSQRHTLPKSPGRGQPINFPKLNASPTHIPKIDLPPNLRERRTPPPVDGCARPRIPPTVKVLPPAGPRRPPIAPVREQYPYERPMQEPLSFEQAVAIGRGSRQLPSPAVPNGYKPGRGVRRPRHSDSDEDDWC

>NP_001123176.1|Caenorhabditis_elegans_Cav2

MCIRLKYLKSLFFLERDSTTFDQRSAARRASVLGRSAAESLSAQEAASSSERGEHTNSRSPSTSYSSCIDDERKFSSPHRRRVVDVSDHKTCALLMTRMKEASRQLPSPSQLAAEEARREQKAESGTFVRKTTLSSNAPVKEKGPSSLFIFAEDNIIRRNAKAIIEWGPFEYFILLTIIGNCVVLSMEQHLPKNDKKALSEWLERTEPYFMGIFCLECVLKVIAFGFALHKGSYLRSGWNIMDFIVVVSGVVTMLPFSPATQTANQPVDSVDLRTLRAVRVLRPLKLVSGIPSLQVVLKSILCAMAPLLQIGLLVLFAIIIFAIIGLEFYSGAFHSACYNERGEIENVSERPMPCTNKTSPMGVYNCDVKGTTCLQKWIGPNYGITSFDNIGFAMITVFQCITMEGWTTVMYYTNDSLGSTYNWAYFIPLIVLGSFFMLNLVLGVLSGEFAKERERVENRREFLKLRRQQQIERELNGYLEWILTAEEVILKEDRTTEEEKAAIMEARRRAANKKLKQASKQQSTETEEDFEEDEDEMEEEYVDEGGTVEDEFAERKKRGCCHSVGKFIKQLRIQIRIMVKTQIFYWSVITLVFLNTCCVASEHYGQPQWFTDFLKYAEFVFLGIFVVEMLLKLFAMGSRTYFASKFNRFDCVVIVGSAAEVIWAEVYGGSFGISVMRALRLLRIFKLTSYWVSLRNLVRSLMNSMRSIISLLFLLFLFILIFALLGMQLFGGRFNFPTMHPYTHFDTFPVALITVFQILTGEDWNEVMYLAIESQGGIYSGGWPYSIYFIVLVLFGNYTLLNVFLAIAVDNLANAQELTAAEEADEKANEIEEESEELDEQYQEGDHCTIDMEGKTAGDMCAVARAMDDLDEECEEEESPFGGPKPMVPYSSMFFLSPTNPFRVLIHSIVCTKYFEMMVMTVICLSSVSLAAEDPVDEENPRNKVLQYMDYCFTGVFACEMLLKLIDQGILLHPGSYCRDFWNILDGIVVTCALFAFGFAGTEGSAGKNLNTIKSLRVLRVLRPLKTIKRIPKLKAVFDCVVNSLKNVFNILIVYFLFQFIFAVIAVQLFNGKFFFCTDKNRKFANTCHGQFFVYDNQNDPPRVEQREWRLRPFNYDNTINAMLTLFVVTTGEGWPGIRQNSMDTTFEDQGPSPFFRVEVALFYVMFFIVFPFFFVNIFVALIIITFQEQGEAELSEGDLDKNQKQCIDFALNARPRSLFMPEDKNSTKYRIWRLVTSPPFEYFIMTMICCNTLILMMKYYNNPLFYEEILRLFNTALTAVFTVESILKILAFGVRNYFRDGWNRFDFVTVVGSITDALVTEFGGHFVSLGFLRLFRAARLIRLLQQGYTIRILLWTFVQSFKALPYVCLLIGMLFFIYAIVGMQVFGNIWLNAATEINRHNNFQSFFNAVILLFRCATGEGWQDIMMAAVQGKDCARAGSAEINFEKGQTCGSNVSYAYFTSFVFLSSFLMLNLFVAVIMDNFDYLTRDSSILGPHHLDEFIRVWADYDPAATGRIHYSEMYEMLRIMAPPVGFGKKCPYRLAYKHLIRMNMPVAEDGTVHFTTTLFALIRESLSIKMRPVEEMDEADEELRLTLKKIWPLKAKKNMVDLVVPPNHELCFQKLTVGKIYAGLLILENYRARKSGTEVGGQGLFGGGLRSLVAAAKAAESQHSSHTPQPPEETTPIIPQHAQQFSAAPTMSAQGSLQQMQGTSSGGGQRPYSLFNSFVDTIKSGKQDGDVTDVQYQSVDQQHEKMNSTGRRLSDMFSKIRRGTSADHNPHQTEHLLAQDNRSPSSPRYRSMARASPPSPAERYGHPPRYRTESPPSSRSEYQMSIRDPIIRRNRYNTMEHSRSSHDPQYHQDQQQQQQPHHQQHSQHLQHSHHKTYQNHNQYSRSPIYSDDSSVAESYRREREFRRYQDSTPQDVSEDDDPMPTAVRARRLPLISTMPTHYESAYQPSSYNQHLNDSYGLGTGYQRDYHTSHSHSHHPTSQQQQHQPMYSTSPLISPRSSHSYYTPRSSQYYEIPSPSPDIYPSYRGSASPRRYPTSTVVVAPDREGSSARVIQAQPGSIPLSDSETEDDPRWAIV

>PIC16379.1|Caenorhabditis_nigoni_Cav2

MLGIDQIARLAAEEARREQKAESGTFVRKTTLSSNAPIKEKGPTSLFIFAEDNIIRRNAKAIIEWGPFEYFILLTIIGNCVVLSMEQHLPKNDKKALSEWLERTEPYFMGIFCLECILKVIAFGFALHKGSYLRSGWNVMDFIVVVSGVVTMLPFGSTIQAANQPVDTVDLRTLRAVRVLRPLKLVSGIPSLQVVLKSILCAMAPLLQIGLLVLFAIIIFAIIGLEFYSGAFHSACYNERGEIENVSEKPMPCTNKTSPMGVYNCDVKGTTCLQKWIGPNYGITSFDNIGFAMITVFQCITMEGWTTVMYYTNDSLGSTYNWAYFIPLIVLGSFFMLNLVLGVLSGEFAKERERVENRREFLKLRRQQQIERELNGYLEWILTAEEVILKEDRTTEEEKAAIMEARRRAANKKLKQASKQQSTETEEDFEEDEDEMEEEYVDEAERKKRGCCHSLGKFIKQLRIQIRIMVKTQIFYWSVITLVFLNTCCVASEHYGQPQWFTDFLKYAEFVFLGIFVVEMLLKLFAMGSRTYFASKFNRFDCVVIVGSAAEVIWAEVYGGSFGISVMRALRLLRIFKLTSYWVSLRNLVRSLMNSMRSIISLLFLLFLFILIFALLGMQLFGGRFNFPTMHPYTHFDTFPVALITVFQILTGEDWNEVMYLAIESQGGIYSGGWPYSIYFIVLVLFGNYTLLNVFLAIAVDNLANAQELTAAEEADEKANEIEEESEEVDEQFQEGDHCTIDMEGKTAGDMCAVARAMDEMDEECEEEESPFGGPKPMVPYSSMFFLSPTNPFRVLIHSIVCTKYFEMMVMTVICLSSVSLAAEDPVDEENPRNKVLQYMDYCFTGVFACEMLLKLIDQGILLHPGSYCRDFWNILDGIVVTCALFAFGFAGTEGSAGKNLNTIKSLRVLRVLRPLKTIKRIPKLKAVFDCVVNSLKNVFNILIVYFLFQFIFAVIAVQLFNGKFFFCTDKNRKFAHTCHGQFFVYDNQNDPPRVEQREWRLRPFNYDNTINAMLTLFVVTTGEGWPGIRQNSMDTTFEDQGPSPFFRVEVALFYVMFFIVFPFFFVNIFVALIIITFQEQGEAELSEGDLDKNQKQCIDFALNARPRSLFMPEDKNSTKYRIWRLVTSPPFEYFIMTMICCNTLILMMKYYNNPLFYEEILRLFNTALTAVFTVESILKILAFGVRNYFRDGWNRFDFVTVVGSITDALVTEFGGHFVSLGFLRLFRAARLIRLLQQGYTIRILLWTFVQSFKALPYVCLLIGMLFFIYAIVGMQVFGNIWLNAATEINRHNNFQSFFNAVILLFRCATGEGWQDIMMAAVQGKDCARAGSAEINFEKGQTCGSNVSYAYFTSFVFLSSFLMLNLFVAVIMDNFDYLTRDSSILGPHHLDEFIRVWADYDPAATGRIHYTEMYEMLRIMAPPVGFGKKCPYRLAYKHLIRMNMPVAEDGTVHFTTTLFALIRESLSIKMRPVEEMDEADEELRLTLKKIWPLKAKKNMVDLVVPPNHGIFELCFQKLTVGKIYAGLLILENYRARKSGTEIGGQGLFGGGLRSLVAAAKAAESQHSSHTPQPPEETTPIITHPQQYSTAPTMSAQGSTSQQQQGQAGTSGSVGGGQRPYSLFNSFVDTIKSGKADGDLTPVQYQSVDQQHDKMNSTGRRLSDMFSKIRRGNSTDHNPHQNYSHRFGYSRNRGSRYDETEHLLAQDNRSPSSPRYRSMARASPPSPAERYGSGHPPRYRTESPPSSRSEYQMSIRDPIIRRNRYNTMEHSRSSHDPQYLHHHHQDHYQQSHPQHSHHPQHLQHSHHKMYQNHNQYSRSPIYSDDSSVAESYRREREMRRYQDSTPQDVSEDDDPMPTAVRARRLPLISTMPTHYESAFQPSYNQHLNDSYGLGTGYQRDYHTSHPHHSHHSQSHQQQQQQHHQPMYSTSPLISPRSSHSYYTPRSSQYYEIPSPSPDIYPSYRASASPRRYPTSTVVVAPDREGSSARVIQAQPGSIPLSDSEAEDDPRWAIV

>CDJ96819.1|Haemonchus_contortus_Cav2

MLGIDQIARLAAEEARREQKTESGPFAVRKSTLASNAPVKEKGPSSLFIFSEDNFIRRNAKAIIEWGPFEYFILLTIIGNCVVLAMEQHLPKNDKKPLSELLERTEPYFMGIFCLECVLKIVAFGFIAHKGSYLRSGWNIMDFIVVVSGVVTMLPVSPAAAGGGSGQVETVDLRTLRAVRVLRPLKLVSGIPSLQVVLKSILCAMAPLLQIGLLVLFAIVIFAIIGLEFYSGAFHSACYNDRGEIENVSEKPSPCTNKTTTMGVYNCDVEGTTCLNKWIGPNYGITSFDNIAFAMITVFQCITMEGWTTVMYYTNDSLGSTYNWAYFIPLIVLGSFFMLNLVLGVLSGEFAKERERVENRREFLKLRRQQQIERELNGYLEWIMAAEEVILKEDRTTEEEKQAILDGRRRAETKKKMKDASKQQSTETEEDMEEEEEELDEEYLDEGERRRRGCYYAICKRIRKARIQMRVIVKTQIFYWSVITLVFLNTACVASEHYGQPPWLTKFLQYAEYVFLGIFIMEVLLKLFAMGSRTYFASKFNRFDCIVIVGSAFEVIWAEVKGGSFGISVLRALRLLRIFKLTSYWVSLRNLVRSLMNSMRSIISLLFLLFLFILIFALLGMQLFGGKFNFPTMHPYTHFDTFPVALITVFQILTGEDWNEVMYLAIEAQGGIYGGGMVYCIYFIVLVLFGNYTLLNVFLAIAVDNLANAQELTAAEEADEKANEMDDSEEEEPDGDHCAIDMDGNDQDDDEECEEEESPFGGPRPMVPYTSMFFLSPSNPLRVLVHSIVCTKYFEMMVMGVICLSSISLAAEDPVDEENPRNKVLQYMDYCFTGVFACEMLLKLIDQGIILHPGSYCRDFWNILDGVVVTCALVAFGFAGTEGSAGKNLNTIKSLRVLRVLRPLKTIKRIPKLKAVFDCVVNSLKNVFNILIVYFLFQFIFGVIAVQLFNGKFFYCTDKTKRFAYQCHGQFFIFDNQNEPPRVEQREWRLRPFNYDNTINAMLTLFVVTTGEGWPGIRQNSMDTTFEDQGPSPFYRVEVALFYVMFFIVFPFFFVNIFVALIIITFQEQGEAELSEGDLDKNQKQCIDFALNARPRSLFMPEDKNSIKYRIWRLVTSAPFEYFIMAMICCNTIILMMKFHGNSDFYEKVLRLFNTALTAVFTVESILKILAFGVRNYFRDGWNRFDFVTVVGSITDALVTEFGGHFVSLGFLRLFRAARLIRLLQQGYTIRILLWTFVQSFKALPYVCLLIGMLFFIYAIVGMQVFGNIWLNAATEINRHNNFQSFFNSVILLFRCATGEGWQDIMMACGAQKDCARAGSDEINYDKGQTCGSNVSYAYFTSFVFLSSFLMLNLFVAVIMDNFDYLTRDSSILGPHHLDEFIRVWADYDPAATGRIHYTDMYDMLRNITPPVGFGRKCPYRLAYKHLVRMNMPVADDGTVHFTTTLFALIRESLSIKMRPVEEMDEADEELRQTLKKIWPLKAKKNMIDLVVPPNHELCFQKLTVGKIYAGLLILENYRAKKSGTEIGGGGLFGGGLRGLVAAAKAAGGVNHMAAPQVDETSQLIPNHTPHANKTPAPRPYTLYTPLEETPKSNKSSEDGGETPTRYSPPQDISKTTAGRRLSDMFSRIRRGGGGGAHDHPPQQYAPRFNYYRERSSQYDETEQLLPSQRSRTPSPRYSSLHGRNMSPPSPAERYPSRFRSSSPPSSSDYPMSVRDPAPRRPRPAQLYYSRPLRSGYSYPPQQYARSPYSDDSVTSSYRRHGYTRYHDSTPPEISEDDEAMPNSVRQRRLPIIGSMPPVSGGDLRPAPYAQPPYAGQFASPPYRPPSVISPVRPHHQDYYTPRDNYYDIPSPSPAGNDTYHGYNRTSPSRYPSVVYAHDTGPRTRIIQAQSGAIPLSDSDSDDQERWAVV

>XP_024503488.1|Strongyloides_ratti_Cav2

MVYEEYNENYTKKNYYDCNYMRNVRSHNFTNNDNCKEIRRPSYQRISYINDPLKDISIKQWMLPVTPAFFSPLVSPQKNTSKMSDFSLPLLNSNMHLPTSGNTLVPPAGEAFREMMRLAALPAISGLAAEEARKEQSRLDGGFGTGNHGGVGSGASPFGGIGRKPGIGSAPVKEKGPSSLFIFSEDNFIRKNAKAIIEWGPFEYFILLTIIGNCVVLAMEQHLPKNDKKPLSEMLERTEPYFMGIFCFECLIKIIAFGFILHKGSYLRSGWNIMDFIVVVSGVLTMLPFSPSSNETVDLRTLRAVRVLRPLKLVSGIPSLQVVLKSILCAMAPLLQIGLLVLFAIVIFAIIGLEFYSGIFHSACYNSDGEIENLSEKPFPCSNKSSTTGAYNCDVPGTVCIQQWIGPNYGITSFDNIAFAMITVFQCITMEGWTSVMYYTNDSLGSTYNWAYFIPLIVLGSFFMLNLVLGVLSGEFAKERERVENRREFLKLRRQQQIERELNGYLEWILTAEEVILKEDRTTDEEKAAIMEARRRAATKKLKQATKQQSTETEEELEEEEEEEDEESYIEDHGGKKIKRSFFDNINRKIRNIRTSLRIIVKSQIFYWSVITLVFLNTACVASEHYGQPAWFTEFLKYAEYGFLGIFICEMLVKLFAMGYRTYFASKFNRFDCIVIVGSAFEVLWAEVKGGSFGISVLRALRLLRIFKLTSYWVSLRNLVRSLMNSMRSIISLLFLLFLFIVIFALLGMQLFGGKFNFPNMHPYTHFDTFPIALITVFQILTGEDWNEVMYLAIESQGGIYDGGMVYCIYFIVLVLFGNYTLLNVFLAIAVDNLANAQELTAAEEADEKANEICEDSDDGEDENGDQCIDMEERDYYDDECEEEESPFGGPKPMVPYSSMFIFSPTNCLRVFVHSFVSTKYFEMFVMFVICLSSIALSAEDPVDEENPRNKVLQYMDYCFTGVFACEMFLKLIDQGVILHRGSYCRDFWNVLDGVVVVCALVAFSFAGTDGAAGKNLNTIKSLRVLRVLRPLKTIKRIPKLKAVFDCVVNSLKNVFNILIVYFLFQFIFAVIAVQLFKGTFFYCTDSNKKFAHECHGQFYIYSKQDYEPIVQPREWKLRPFNYDNTLNAMLTLFVVTTGEGWPGIRQNSMDTTEEDQGPSPFFRVEMALFYVMFFIVFPFFFVNIFVALIIITFQEQGEAELSEGDLDKNQKQCIDFALNARPRSLFMPENKNSMKYRIWRLVTSTPFEYFIMAMICCNTLILMMKFHGNSPGYEKVLRFFNTALTAVFTVESILKILAFGVRNYFKDGWNRFDFITVVGSITDALVTEFGGHFVSLGFLRLFRAARLIRLLQQGYTIRILLWTFVQSFKALPYVCLLIGMLFFIYAIVGMQVFGNIRLDPTTEINRHNNFQSFFNSVILLFRCATGEAWQDIMLSCTAGKYCASKDEFTEYNIMKGATCGTNMSYAYFTSFVFLSSFLMLNLFVAVIMDNFDYLTRDSSILGPHHLDEFIRVWADYDPAATGRIHYTDMYEMLRNIAPPVGFGRKCPYRLAYKHLIRMNMPVAEDGTVHFTTTLFALIRESLSIKMRPVEEMDEADEELRQTLRKIWPLKAKKNMIDLVVPPNHELCYQKLTVGKIYAGLLILENYRARKTGTEICSGGLFGGGLRGLVAAAKSAAGGGGTSNFQPPLHHTEHEDNYDDEISSQNNDNKKYTKNDISSNYNHDEGTSLNNNSHINIHRPQQSLFSTIVDTIKPTKSNDSITSNTHEYQQKIRNKPKNDNNDMYYDNSHTSLDTYLQTYDREPMRYGNNYNNGKFSGVFSKIRRREARSDYQPVSFQHYFPRSQRYIPRYCGSSQNNYYSNKRDDYYDETQHLLSNQRHSRSPSPQDIHQNPSLYGRSYSERYSGYQPIHNPPNISYSEDHSGPYNRYSGNMPKGHDNFSMPYYGESHTPKLSSYHNTPNIPNTNNHVLHHQPMGYTNPSNQSYDDIYNDRNYSNDKSIHNAGSSQNIRVNWRYDDQYDEDPRIVPTQRRLPDINKISIPRPRSGFNESSSYENTNNYTRFQPQYHSQGGAQTIVYAQPGNVPLSDSEGEEGGERWAMI

>KRY49212.1|Trichinella_britovi_Cav2

MNLRIDDIARLASERTKLQLHSQSLDSSADKSSQARRIPGKRGDGKGPTSLFIFTENNLVRRYAQISLTVSFFTPFEYFILMTIIANCIVLALDQHLPHNDKMPLSLKLEATEPYFMGIFTIECLLKIIAFGFVMHKGSYLRSGWNILDFIVVMSGVISMLPFTTSGVDLRTLRAVRVLRPLKLVSGIPSLQVVLKSILCAMAPLLQIGLLVLFAIVIFAIIGLEFYSGAFHRIPVAIGNKPFPCTNKSTTGAYHCPEGTTCKEQWIGPNYGITSFDNIAFAMLTVFQCITMEGWTNVMYYTNDSQGDTFNWLYFIPLIILGSFFMLNLVLGVLSGEFAKERERVENRREFLKLRRQQHIERELNGYLEWICKAEEVILNEERTTEEERKAIMEARHLAVSKQLKHLVQQSTETEDDLEEEDLDVEFRTKDLSQTGKCWLVVRRLRVFVRRFVKTQFFYWLVITLVFLNTVCVSIEHYGQPQWLDEFLYYAEWTFLGIFLFEMLFKMFGLGIGTYFQSSFNIFDFVVITGSLFEVIWEELKGGSFGISVLRALRLLRIFKVTKYWTSLRNLVVSLMNSMRSIISLLFLLFLFILIFALLGMQLFGGEFNFPEGRPSTHFDTFPVALITVFQILTGEDWNEVMYLAIESQNDTLLNVFLAIAVDNLANAQELTAAEEAHEQEVQSTYSSADIKNDECEYEKGPKMIENLSQNGQAKLTDACDAVCDVKNTLNIQENKTVTSAPLCYVNSIDESDSNCAFGGHREIVPYSSLFIFSPKNRFRIFVHKIVCTKYFEMAIMVVICLSSISLAAEDPVDESNPRNKYLNYLDYAFTAVFTIEMILKVIDMGVIIHPGSYCRDLWNIMDATVVICALVGFAFVDSTKAGKNLSTIKSLRVLRVLRPLKTIKRIPKLKAVFDCVVNSLKNVFNILIVFILFQFIFAVIAVQLFKGKFFYCTDRTKRFEQDCQGYFFHYDKQGSPPQVVQREWTSFALNYDNTIHAMLTLFTVTTGEGWPGIRQASIDATEENQGPIPFNHIEVALFYVVYFIVFPFFFVNIFVALIIITFQEQGEAELAEGDLDKNQKQCIDFALNARPVCRYIPEDKDSIKYHIWKMVVSTPFEYFIMAMICLNTIILMMSYYQEPPAYRAVLRYLNSTLTAVFTVEAILKILAFGVRNYFKDGWNIFDFITVIGSITDALVTEFGGNFVSLGFLRLFRAARLIKLLRQGYTIRILLWTFVQSFKALPYVCLLIGMLFFIYAIVGMQVFGNIELNGDTEINRHNNFQTFFNSIILLFRCATGEAWQEVTLACIANRKCDPRTGKLNNECGTNFAYVYFTSFVFLSSFLMLNLFVAVIMDNFDYLTRDSSILGPHHLDEFVRVWAEYDPEATGRIHYTDMYEMLRNIPPPVGFGRKCPYRLAYKHLIRMNMPVDEDGTVQFTTTLFALIRESLSIKMRSADEMDRADMELRKTLKKLWPIHAKKNLTDLAVPPNSKLCNKKLTVGKIYAGLLLLENFRSKRCGRVPDRSTFLFGRLVDAALKEKEKIHSSEADPSSCTEFFSDSEPPRSEIRYGFEHQRRMLPTAEQPRIFDIPGKWQTHDEMNSQMQSIYVPQDEYNNERVQHRRKVLRRQQEDDRVDGFASAQEWHNYARQRMQRCTDSKNKALSERANLDHYYVTGLADAKSHHESASNVLFNQLSPVLGHSFSPAFVEKIPSPACYFHQQLPMVWENPVTNKFSEVSHLKAHPMFKSSPELQQCVGKRKRLAAKTARTFCTSTIRPHREVENYAEFLQLSDDESCNYSARSAKQTVFDSNLMSPLTVSPDPAFFSSSSSSSDASFVNVSCPAQQNVNTSSYASYCQISPDNSYIFAGNPTSSGEHPVRSSQSSGATYKLSGYKTAGHCQPALKEPNQQRLYCMRSNGPNADEHHCHSVVHSQPGHIPLSDSEQEDWC

>KRY73966.1|Trichinella_pseudospiralis_Cav2

LASERTKLQLHSQSLDSSADKSSQARRIPGKRGDGKGPTSLFIFTENNFVRRYARTIIEWGPFEYFILMTIIANCIVLALDQHLPHNDKMPLSLKLEATEPYFMGIFTIECLLKIIAFGFVMHKGSYLRSGWNILDFIVVMSGVISMLPFTTSGVDLRTLRAVRVLRPLKLVSGIPSLQVVLKSILCAMAPLLQIGLLVLFAIVIFAIIGLEFYSGAFHSTCYNEEGIYFFIFIFSFFEYAQLIYIVHIIFNFSGIPVAIGNKPFPCTNKSTTGAYHCPEGTTCKEQWIGPNYGITSFDNIAFAMLTVFQCITMEGWTNVMYYTNDSQGDTFNWLYFIPLIILGSFFMLNLVLGVLSGEFAKERERVENRREFLKLRRQQHIERELNGYLEWICKAEEVILNEERTTEEERKAIMEARHLAVSKQLKHLVQQSTETEDDLEEDLDVEFRTKDLSQTGKCWLVVRRLRVFVRRFVKTQFFYWLVITLVFLNTVCVSIEHYGQPQWLDEFLYYAEWTFLGIFLFEMLFKMFGLGIGTYFQSSFNIFDFVVITGSLFEVIWEELKGGSFGISVLRALRLLRIFKVTKYWTSLRNLVVSLMNSMRSIISLLFLLFLFILIFALLGMQLFGGEFNFPEGRPSTHFDTFPVALITVFQILTGEDWNEVMYLAIESQNGIYGGGMIYSIYFIILVLFGNYTLLNVFLAIAVDNLANAQELTAAEEAHEQEVQSTYSSADTKNDECEYENGPKMIENLSQNGQAKLTDACDAVCDVKNALNIQENKTVTSASLCYVNSVDESDSNCAFGGHREIVPYSSLFIFSPKNRFRLFVHKIVCTKYFEMAIMVVISLSSISLAAEDPVDESNPRNKYLNYLDYAFTAVFTVEMILKVIDMGVIIHPGSYCRDLWNIMDATVVICALVGFAFVDSTKAGKNLSTIKSLRVLRVLRPLKTIKRIPKLKAVFDCVVNSLKNVFNILIVFILFQFIFAVIAVQLFKGKFFYCTDRTKRFEQDCQGYFFHYDKQGSPPQVVQREWTSFALNYDNTIHAMLTLFTVTTGEGWPGIRQASIDATEENQGPIPFNHIEVALFYVVYFIVFPFFFVNIFVALIIITFQEQGEAELAEGDLDKNQKQCIDFALNARPVCRYIPEDKDSIKYHIWKMVVSTPFEYFIMAMICLNTIILMMSYYQEPPAYRAVLRYLNSTLTAVFTVEAILKILAFGVRNYFKDGWNIFDFITVIGSITDALVTEFGGNFVSLGFLRLFRAARLIKLLRQGYTIRILLWTFVQSFKALPYVCLLIGMLFFIYAIVGMQVFGNIELNGDTEINRHNNFQTFFNSIILLFRCATGEAWQEVTLACIANRKCDPRTGKLNSECGTNFAYVYFTSFVFLSSFLMLNLFVAVIMDNFDYLTRDSSILGPHHLDEFVRVWAEYDPEATGRIHYTDMYEMLRNIPPPVGFGRKCPYRLAYKHLIRMNMPVDEDGTVQFTTTLFALIRESLSIKMRSADEMDRADMELRKTLKKLWPIHAKKNLTDLAVPPNSKLCNKKLTVGKIYAGLLLLENFRSKRCGRVPDRSTFLFGRLVDAALKEKEKIHSSEADPSSCTEFFSDSEPARSEIRYGFEHQRRMLPTAEQPRIFDIPGKWQTHDEMNSPMQSVYAPQDEYNNERVRHRRKVLRRQQEDDRVDNFASAQEWHNYARQRIQRCTDSKNKALSERTNFDRYYMTGLADTKGHHETAGNVLFNQLSPVLGHSSFSPAFGEKIPSPACYFRQQLPMVWENPVANKFSEVSHLKAHPMFKSSPELQQCAGRRRRLAAKTARTFCTSTIRAPREVENYAEFLQLSDDESCNYSARSAKQTVFDSNLMSPLTVSPDPAFFSSSSSSSDTSFVNVSCPAQQSVNTPSYANYCQISPDNSYIFAGNPTSSGEQQVRSSQSSTNAAYKLGGYKATVHCQPVLKEPNQQRLYYMRSNGSNAEEHRCHSVVHSQPGHIPLSDSEQEDWC

>AAO83841.1|Lymnaea_stagnalis_Cav2

MATFQANNGQQDDGDNTTNQDGPFSHFSRKAALLGLPGMASQSTRSLFIFSEENFIRKYAKIIIEWGPFEYMVLLTIIANCIVLALEEHLPSQDKTPLALQLDDTEVYFLGIFCVEAFLKIVALGFCLHKRSYLRNIWNIMDFIVVVTGFITLFAQGSSTTFDLRTLRAVRVLRPLKLVSGIPSLQVVLKSIIRAMAPLLQVCLLVLFAIVIFAIIGLEFYVGVFHNACYKKGSHTRSEDDIDTGDEDDIRPCLPSSESQGAFQCQVNISNCKAGWRGPNAGITSFDNIGYAMLTVFQCITMEGWTNVLYYTNDALGNQFNFLYFIPLIILGSFFMLNLVLGVLSGEFAKERERVENRRAFFKLRRQQQIERELNGYLEWICKAEEVILSEERTTDEEKLKIIEARRQAAARKMKQLKAEDNENDSEQNDNDLLAAMAPGNSFKSMKKRRTTGKCASFWRAEKRFRYSIRRLVKSQLFYWIVIVLVFLNTASVASEHYNQPEWHVQFLYITEYAFLGLFIFEMSIKMYALGVRMYFQSSFNIFDCVVIVGSIVEVIWSEFKRGSSFGISVLRALRLLRIFKVTRYWSSLRNLVISLLSSMRSILSLLFLLFLFIIVFALLGMQLFGGEMNFEEGRPSAHFDTFPIALLTVFQILTGEDWNEVMYNGIKSHGGIENQGMFYSSYFIVLVLFGNYTLLNVFLAIAVDNLANAQELTAAEEEQEEEEAVRREEIEKEMAEQFAAQGRPPLVNICPPSPQNNEENKTANFNYAGNRVDINLSQNNLKDNRDKKISVDNVLETAAKTSSVTMPLANNTEEDMSDTASTTSSGTADVQPRASSQNDNGGFHGPKPMLPYSSMFIFGPTNPIRRFCHFVVNLRYFDLFIMIVICASSVALAAEDPVIENSRRNEILNYFDFVFTGVFTIELVLKVIDLGVLLHPGSYIRDLWNILDATVVICALVAFVFKDKSDSAGKNLNTIKSLRVLRVLRPLKTINRVPKLKAVFDCVVNSLKNVSNILIVYILFQFIFAVIAVQLFKGRFFYCTDESKSTRDECQGQFFEYDGHSNDPTVRDREWLRQDFHYDNIMMAMLTLFTVTTGEGWPSVLKHSMDSTYEDRGPKPVYRMEMSLFYVVFFIVFPFFFVNIFVALIIITFQEQGENELMDQEMDKNQKQCIDFAINAKPHCRFIPKNKNSIKYKIWRLVQSSKFEYFVMTLITLNTIVLMMKYDGMSDNYKDVLAKLNEGFTVLFTLECLLKIIGLGPRNYFHDPWNVFDFTTVVGSIIDVLITEFSKRQVSFGFFRLFRAARLVKLLRQGYTIRLLLWTFFQSFKALPYVCLLILMLFFIYAIIGMQVFGSIKLDSKTSINRHNNFRTFFSALTLLFRCATGEAWQQIMQSCLAGQPCDPESIRDDDPPDMAESGCGTNIAYMYFVSFIFLCSFLMLNLFVAVIMDNFDYLTRDSSILGPHHLDEYVRVWSMYDPKATGRIHYTDMYEMLRNMEPPVGFGKKCPYKLAYRKLIRMNMPVAEDGTVHFTTTLFALIRECLIIKMGPAEIMDRRDEEMRETIRKLWPVQGKKMMDMLMPPSDELDEGKMSVGKIYAGLLISENWKAYKASQNASNNFKMRPSLFRRLMGGMRTSSARSSQSLDSEQSDDNDGGGGGGGSANAGGGGGSSAGHSFLRRNSSKRRKGGDHGDNSNVQPGTDFSGGLRPEHASSLTSRGDRAGARSPSLPPTPLSPRSPMGAQSPFGSPRASPIPSRRSPSPRRFDVGFASAVANLCEQAHTIADQDRQKRYGVKEDSITSSSPTFRGRSRQRSRPPLQAQSPVLGSPLPSPAHPRVRGGGDSGFYRSTSLETRSRSPSPNLTASPPPRSGSTSLVQRSRSPSPSLVAASPPMTSSAHRRLPVAPSSTSSSGSGVTTMSPAKPVSLNLSEPRYRDNIAIKDTLPVRGAPSPPGRGNINFPRLNASPTRVPKLNIPVSSTSGIAAASPLPPHHHRHAPPPGRLGRPEPYSPTERNNLNKTSDPSSSSARSSTLPIAHRTSGYPRDDSNWGARELPGEGGRDSRSLPRPSPRASSRSPDPRGGSRHDDRFMAASQQEGSPARGTRSRGGILPNGFKPKGRKPEKYEMRSDSHTALQEDSDEDDDDWC

>AVD53847.1|Aplysia_californica_Cav2

MATFQANNSLQDDGDTSNSLDGPFGHLSRKAALFGLPGMAANSTRSLFIFSEENFIRKYAKIIIEWGPFEYMVLLTIIANCIVLALEEHLPEMDKTPLALQLDDTEVYFLGIFCVEAFLKIVALGFCLHKGSYLRNVWNIMDFIVVVTGFITLFASSGSSGAFDLRTLRAVRVLRPLKLVSGIPSLQVVLKSILRAMAPLLQVCLLVLFAIVIFAIIGLEFYVGVFHSACFRKGTHTFTEDDIDLGDEISIWPCDANSDSYGAFRCQTNISSCLPGWVGPNDGITSFDHIGYAMLTVFQCITMEGWTTVLYYTNDALGNWFNYLYFIPLIIVGSFFMLNLVLGVLSGEFAKERERVENRRAFFKLRRQQQIERELNGYLEWICKAEEVILSEERTTDEEKLKIIEARRQAAARKMKQLKGEDTDNDNEQNDDDLLAEMTPGNSFAKNLKKRKRNGKCASFWRAEKRLRYSIRRLVKSQLFYWIVIVLVLLNTISVASEHYNQPEWFVDFLYITEYAFLGLFIFEMSLKMYALGVRLYFQSSFNIFDCVVIVGSIFEVIWSEFKQDSFGFSVLRALRLLRIFKVTRYWASMRNLVISLLSSMRSILSLLFLLFLFILIFALLGMQLFGGKMNFEDGRPSAHFDTFPIALLTVFQILTGEDWNEVMYDGIRAHGGIEKSGMLASSYFIVLVLFGNYTLLNVFLAIAVDNLANAQELTAAEEEQEEEEAVRREEIEKEMAEQFSSSQGRPPLVNICPPSPQNNEENKTANFNYTGNRVDINLSQNNLKDTRDKKISVENVDSGPPTKTSSVTMPLAKNAEEDMSDTASTTSSGTMDAQPRTNSQNDDGGFSGPKPMLPYSSMFIFYPTNPIRQFCHFVVNLRYFDLFIMIVICASSVALAAEDPVVAVSGRNDILNYFDFVFTGVFTIELILKVIDLGIILHPGSYIRDLWNILDATVVICALVAFAFNESAGKNLNTIKSLRVLRVLRPLKTINRVPKLKAVFDCVVNSLKNVSNILIVYILFQFIFAVIAVQLFKGRFFYCTDESKSTRDDCQGQFFEYEDNSEQPVVKDREWLRQDFHYDDLANAMLTLFTVTTGEGWPSVLKHSMDSTQENRGPKPGSRMEMAIFYVVFFIVFPFFFVNIFVALIIITFQEQGENELMDQEMDKNQKQCIDFAINAKPLCRFMPKNKNSVKYKIWKLVQSPKFEYFIMTLITLNTIVLMMKFDPKESRQSRRREKGEDAARILHLINTVFTSLYGLGFLLKLCAYGKNYFHDPWNVFDLITVIGSIIDVVISEFSMGRVSFGFFRLFRAARLVKLLRQGYTIRLLLWTFFQSFKALPYVCLLILMLFFIFAIIGMQVFGSIKLDSKTEITRHNNFRTFFSALTLLFRRATGEAWQQIMWSCLSGKPCDPESIRKSDPPSMAESGCGSNIAYIYFVSFIFLCSFLMLNLFVAVIMDNFDYLTRDSSILGPHHLDEYVRVWSTYDPGASGRIHYTDMYEMLRNMEPPVGFGRKCPYKLAYRKLIRMNMPVAEDGTVHFTSTLFALIRECLIIKMGPAEIMDRRDEEMRETIRKLWPVQGKKMMDLLMPPSDELDEGKMSVGKIYAGLLISENWKAYKASQNASNNFKMRPSLFRRLMGGMRTSSARSSQSLDSEHSDENDGGGGGGGGGGGGGGGGHSFLRRNSSKRRREAGNHGDSSNVQPGNDFSGGLRPEHASALNSRAELGGRGGARSPSLPPTPLSPRSPMGGAQSPFGSPRASPVPSRRSPSPRRFDVGFAAAVANLCEQAHNIADQDRQRKYGMKTEDSISSSSPTFRGRSRQRPRPPLQSQSPVLGSPLPSPAHPRARVAGDSGFYRSTSLETRSRSPSPNLTASPPPRSGSTSLIQRSRSPSPSAAAGSPPMTKRASRRLPVAPSPTGHGSVGAGGGGGHHHHPSSSTSSPAKPASLNLTEPRYRSGDGRGMSHVEKDTLPVRGVPSPPSTRGGNINFPRLNASPTRVPKLNIPVSSSSQPMTSSSSRHAHPPSSASASSQHHPHPQAPPPGRLGRPEPYSPTERNNLNKMSDPSRSSTLPAAHRTSGYTRDDPQWGARGEGGREHYEGGRDSRSLPRASPRPPSRSPDPGRGGAGGGGGGAGGNSAGRHDDRFMAANQHEGSPARGGGGAGRARGVILPNGFKPKGRKPEKYEMRTDSNTALKEDSDEDDDDWC

>XP_019920407.1|Crassostrea_gigas_Cav2

MAQKSGSRSVFLKKLHEMEQQKLRRKKGNYLSSSSFLSGSSGENITDQDKSAIASGGYPNLGYDEEITEDDSEIIEDDEITLDLDEDEGESSRQKAAKWLRECEDADDISRESSAPTSVSVRRSRRPAEMNGKSYLKTEPGLAQKLNPAHGARKTSWLNIFERIRSKKLSYRLKKFLNNEEETNSHQQSEHHHDPFRKQNPLKRKKTKKLKNMTKQERRIYFKRHANSRLRYKAQNMIGATGSRHLSQRRRASLATPAASPSMFSHLEQVNEEDENKPRLPPTRRRAVDTSDYRTCAILQTRLKTVRNYPEMANFTVGGHEGQDGGEYWGPSDGSLRHFAKKAADISLPGTGGSSSRSLFIFSEENFIRKYAKIIIEWGPFEYMVLLTIIANCIVLALEEHLPKDDKTPLAVQLEETEIYFVVIFLVEALLKIVALGFVLHKGAYLRNIWNIMDFVVVVTGIITMAASSQLDLRTLRAVRVLRPLKLVSGIPSLQVVLKSIIRAMTPLLQVCLLVIFAIIIFAIVGLEFYSGAFKNACFKIGTKGNSEDDIYIGDEANIRPCSSGSPSSADLNGAFKCQENISVCRGRWVGPYFGITNFDNIAYAMLTVFQCITMEGWTEVLYYTNDAIGPYINWLYFYPLIILGSFFMLNLVLGVLSGEFAKERLRVENRRSYIKLRRQQQIDMELSGYLEWICKAEEVILNEDRTTDEDKLRIMEARKRAATKMKKIGKEPSDENNEDNDSDLLSDINIGRSNIQNRKQTGRCAAFWKAEKHFRFSLRRLVKSQPFYWTVIVLVFLNTVCTASEHYGQPKWHEEFLYYTEFVFLGLFIFEMLIKMYGLGVRIYFQSSFNIFDCGVIIVSIIEVIWSYYKDGASFGISTLRALRLLRVFKVTRYWSSLRNLVVSLLSSMRSIVSLLFLLFLFILIFALLGMQLFGGEMNFDDGRPPAHFDTFPIALLTVFQILTGADWNEVMYNGIRAHGIEDGDKKGMFYSIYFIILVVFGNYTLLNVFLAIAVDNLTNAQEMTAAEEEEEVGRKEHLEEVRQDVHNQFTDHSQRNKHEKELMEETSGGQQVTVNICPPSPANNEENPKISNFNYISFSKDSNHQNDLKNNKNTANAVTKKSNISQNQFDNDGLDTENNDLVNNNIPRDSNNSEDPPPQAPKQEEEGAFGSGPRPMLPYSSMFIFGPKNPIRRFCHFVVNLRYFDLFIMIVICASSFALATEEPVNEDAFRNKILNYFDYVFTIVFTVEMILKVIDLGVFLHPGSYCRNLWNILDATVVICAVVAFFFDNTNVPSDTASKNLNTIKSMRVLRVLRPLKTINRVPKLKAVFDCVVNSLKNVANILIVYMLFQLIFAVIAVQLFKGKFFYCTDESKSTEEECRGQFFSYEEFEDTPSVENREWLRRDFHYDNLFEAMLTLFTVTTGEGWPGILHNSMDSTYEDQGPKPGNRMEMAIFYVVFFIVFPFFFVNIFVALIIITFQDQGEAELEDAQLDKNQKQCIDFAVNARPTSRYMPKNKKTIKYKIWRLVVSTKFEYFVMTLIALNTIVLMMKFEGMSARYKDILKYLNMGFTIMFSIECTLKLIGCGKNYFHDPWNVFDFITVVGSIIDVLVNEFGSTYSSFNVGVFRLFRAARLIKLLRQGYTIRLLLWTFLQSFKALPYVCLLILMLFFIYAIIGMQVFGNIKLDSHTDLNRHNNFRNFLYALMLLFRCATGENWQAIMIACLSGQTCDPESNLEPPKTCGSSAIAYVYFVSFMFLSSFLMLNLFVAVIMDNFDYLTRDSSILGPHHLDEFVREWAEIDPGATGRIHYQDMYEMLKNIEPPVGFGKKCPIKFAYRKLIRMNMPVASDNTVHFTTTLFALIRESLSIKMGPVEEMDKLDDELRELIRKMWPVQARKKKLLNLLVPPNSELNDNHMTVGKIYVGLIIAENWRAYKSSQSKMNNLKMRPQSFFKRMLGVVKTPARRSDTSLNHDSEHSDDGGYGDQSDRLSWNRSFSFLRRNSSKKKRDNQDATSVQARHKQDGPWYINTTFTMHDEDDAISSISSPVNHKRRVHFTGPNYYQPTESMGQDFSSGLKPEHAAGHTNLTRAGSDLSLRSGNKSPSLPASPISPRSPNMPRAFHSPTSSPLVGRRSMSPRRGGDYGFASAVTNIVDQAHYISEQERLKRFGIGTRHDDSLSMSLPNSPSQRLHSLIRRPPLISQEKVVCSPSPSPQPLRHANRDSTFYRSTSLETRSRSPSPNTTPSQTPMHEYYGTSNLTDRSRSPSPASTPPKKQSRKLPNVPLVKPSTLNLAQPKLKENMPRVLPSPTIPKSVKSPGNINFPKLSASPTHKPKSKNIPPPISHYPPPPGKFGRPEPYSPTERNNLNKVSAPSSSQSKTLPQVGRRSNNGDQWNRDRSNFNKYRSSSHSPDVNKSSSERTKFLASEFEEPSSSASRGRQRPSTSVPNGIKARKKKPEKLEMRSDSNIPLNNDSDESESDWC

>BAA13136.2|Heterololigo_bleekeri_Cav2

MNTFSADTGGRDDGDYTHDGSLQFVAKKAATVSLPGMGSSTNRSLFIFSEENFIRKYAKIIIEWGPFEYMVLLTIIANCIVLALEEHLPNEDKTPLAVQLEATEFYFLGIFCVEALLKIVALGFALHKGSYLRNVWNIMDFVVVVTGFISIFPASNSFDLRTLRAVRVLRPLKLVSGIPSLQVVLKSIIRAMAPLLQVCLLVLFAIVIFAIIGLEFYTGAFHKACFIKPNDDSEDNIEYGDEDTIRPCANEGSGYHCRANIAKCRFNWAGPNYGITSFDNMGFAMLTVFQCVTMEGWTQVLYYTDDAVGDAYNWIYFVPLIVLGSFFMLNLVLGVLSGEFAKERERVENRRAFLKLRRQQQIERELNGYLEWICKAEEVILDEERKKDDGTISDEDKLRIIEGALGFNARRLAAQKVKKMKENKELTIGDEDNDGDLLSGINVGGSFGRGLKNRKAHGRCAGFWRAEKHLRFTIRKCVKTQGFYWFVIILVFLNTLCVASEHYGQAEWHTEFLYVMEFAFLALFMSEMLIKMYGLGVRLYFQSSFNIFDCVVILVSIIEVIWSAIKDGSSFGISTLRALRLLRMFKVTRYWSSLRNLVVSLLSSMRSIVSLLFLLFLFILIFALLGMQLFGGMMNFEEGRPPGHFDTFPIALLTVFQILTGEDWNEVMYSGIRARGGIAGGGMLYCSYFIILVLFGNYTLLNVFLAIAVDNLANAQELTAAEEIQEGVRQEQEAAEKKKLAEEAAEKEKLARMEEIEKDLCPDQFGLTPPQVNICPPSPQNNEDLKTGNFPYNTSRLDSNLNQNNIKEERNKIGADKDLKKDRDPLDNVSLHESNANRNNAQPGSSSNLNIQPQNGNSEEPAFGGPKPMLPYSSMFIFGPTNPVRRFCHFVVNLRYFDLFIMIVICASSIALAAEDPVNDESVNNQILNYFDYVFTGVFTIEMLLKIVDLGIILHPGSYCRDAWNILDATVVICALVAFAFGDAAGGNLNTIKSLRVLRVLRPLKTINRIPKLKAVFDCVVNSLKNVSNILIVYLLFQFIFAVIAVQLFKGKFFYCTDMSKSNREECQGQYFDYEDESDKPRVKNREWLRQDFHYDNVMFAMLTLFTVTTGEGWPMVLKNSMDSTSDDMGPKPGYRMEMAIYYVVFFIVFPFFFVNIFVALIIITFQEQGENELVDQDLDKNQKQCIEFSIEAKPSCRYVPKNKNSIKYKIWQVVVSPKFECVVMVLIALNTLVLMMKYYGSPTEYKLLLQNLNLAFSVLFTIECILKLMGFGIGNYFRDRWNMFDFIIVIGSIIDVVTTNVLPSASSFRTGSFRLFRAARLVKLLRQGYTIRLLLWTFLQSFKALPYVCLLIAMLFFIYAIIGMQVFGNIRLDSKTSINRHNNFRSFFYAVLLLFRCATGESWQQIMLSCLSGRPCDPESKMLDNSCGLDIAYIYFVTFIFLCSFLMLNLFVAVIMDNFDYLTRDTSILGPHHLDEYSRVWAEYDPLASGRVHYTDMYEMLRRMEPPVGFGRNCPYRLACRKLIRMNMPLKEDGTVHFSTTLFALVRESLSIRMSSAEEMDKKDEEMREVIKRVWPVQGKKIVDLLVPPNHELNNGKLTVGKVYGGLLIAENWRAYKASQNQNNSLKTEKKEELFWEDDIKEYRDEEDYREYRDEEDYKDDYQEYQEYQEDCKDYEEDYNQVETLEIESRNKYHSTPLHSPCIAINGQQFQFQQLQYSDSEERPPSIFQRIMGVMRTPSARSSQGIDSEHSDNEMEGGHSIDKSHDKSTWQRSFSFLRRGSSRRRKDSTVQKSETASLQPSEKQDFSWGLRPEHAAHPSAPRPGSGSSRGLNFAQTVPLSPVSPRSPLPSQSPLGSPLASPSMHRRSVSPRRGLDVGFASAVSNIVDQAHSIAEHDRHRKHRAYFHPGKPDDSLSVPTSPQMRGRSRGRHRPPLQQQGQVMGSPLPGPTSRRKEPTFYRSTSLENRSRSPSPNLTPTSTLHQHEYYGSAGLTDRSRSPSPTMTPPRKATRKLPAVPSKPSTLNLAQTRPRENMPRVMPSPTVPQPSKSPGSINFPRLNASPTHIPRVGPTIGQAPPLGRLGRPEPYSPTERNCISKLSPERSRTLPIGQRISNRDFSRGVDLYVSHRSRTFDPRINDRGRYFDDPSLDAHMSDHRSETLPNGFKPKKRKPENLDMRGDGTGGPVRHESDEDDDWC

>XP_022085274.1|Acanthaster_planci_Cav2

MEYSAKPALTNGYTRVRRRLGRSLKGAKLKFGLIALAVSGAAGARAAMLHVPRSRRRSSTASYVDSSGHGSSRTPSIALSESLEYNNRNGRPPDLEGDDSERISDIYMDRRYGAEVNAKDTLIAVEQWKRNQTEGEKLTKAQKRRREYEMAALSSAPTSPFSDALDIPGFGGDGTSPAHAGGGKGGGSKKLKGPVGFFLSVLRSNRSLFIFSEQNFIRRCAKWLTEWPPFEYLVLATIIANCVVLALEVHLPMQDKTPMSQELENTEIYFLAIFCLEACIKITALGLVLHEGSYLRNGWNLMDFVVVVTGFVTFIGGLVEGEGDASSAPDLRTLRAIRVLRPLKLVSGIPSLQVVLKAILKAMAPLLQIGLLILFVIIIFAITGMEFFQGKFHYTCFEADAYGASTGKISETEGEDPQVCGKGNETSGRECPEGTVCSEYWEGPNFGITNFDNMLFAMLTVFQCITMEGWTDIMYNCNDSEGPYFVWLYFIPLIILGSFFMLNLILGVLSGEFAKERERVENRREFLKMRRQQQLDKELNGYLEWICKAEEVMLNDNTTSEEERATIEARRRAAAMLQRELSLGNGRAISGTIEDNFDLETLDKAKLTGSPKSNKSEKKQKQKRCLWLRRREKRLRFQVRHMVKTQAFYWLVIVLVFLNTVCVAIEHYQQPEWLTQFLNHAEYVFLGIFITEMAIKMYGLSPAVYFKSAFNKFDCMVILASLFEVIYTKFQGGSFGLSVLRALRLLRIFKVTRYWSPMRYLIISLVHSIRSIVSLVFLLFLFIIIFALLGMQLFGGSYWSSLRNLVVSLLSSMRSIVSLLFLLFLFILIFALLGMQLFGGSFNYDSESPKPANNFDIFPIALMTVFQILTGEDWNMVMYYGVVSKGGVPEGMVYSLYFVILVLFGNYTLLNVFLAIAVDNLANAQEMTKLDQEDEEEAQAARENLTKLSLAVSKTCFFGLCFCLLFLFVSLVREGSTTVHMNDVKRMNGDHNVENGDPEIKVTVVDPPQSRFRRWYTYLNVHGIPWPCPSWGPVDCPGMDAICNSTCWSACPCSRPRCSGRKTDKDEEAEDTQEDDGFGPKEMVPFSSLFIFSTTNPVRRFCHYIVNLRYFDFLIMVAIGLSSLTLAMEDPVNTGSQYNQVLEYFDYAFTTIFTIEMILKIIDMGLLFHKGSYCRDFWNILDSTVVICALVAFGVTQSQGDSGSSGASKNLNTIKALRVFRVLRPLKTIKRVPKLKAVFDCVVNSVKNVTNIAVVYGLFMFIFSVIGVQLYKGRFYHCTDPSKHTRDECMGNYFLYTGDQITKIEKREWQLYPFNYDNVGSALLTLFTVSTGEGWPDVLKHSIDATEEGRGPEPYNNLQMALFYVVYFIIFPFFFLNIFVALIIITFQEQGDQDVQDGEIDKNQKQCMEFCIHAKPTDGFVPKDKNSVKYKIWKLVVSQPFEYFIMSLIALNTIALMMKTYKAEQTYLDTLKYLNIAFTVLFTIEAILKLIGFGPRNYFRVSWNTFDFITVIGSIADAIISEVGVDDFINLSVLRLFRAARLIKLLRQGSSIRILLWTFVQSFKLVSFVFFLTFMLFFIYAIIGMQIFGTVNIDDETAITRHNNFSNFFLAIIMLFRCATGESWQSIMLACQAGSECHPNSIRPNATDEEKYGCGSSLSVAYFVSFIFFSSFLMLNLFVAVIMDNFDYLTRDASILGAHHLDEYVRVWGELEPVGTGRLHYKEMYEMLRTMEPPVGFGRNCPYRIAYKRLIRMNMPVDEDKTVHFTTTLMALIRTALDIKIGKVADRDRHDRELREAIGNFWPHLNADKLNLLVPPDSELIGEKLTVGKIYAALLIYETWREYKAKLQRDGHARTFIQAAQKRPSLFRQLVGAVRRGSAEKIFKEVEEENESTPLAPNEPKPEIKAIDPPPAKSRERDRERDRDRDRDRERERDHGRPRMAATTLPRIDSSGSEGLTQDQQSRRSSPQVHDKSPDRHRGRSRDRQYDRSQDRPQDRPQDRPSDGPRESRRRPRDTLDSARGIEMADIRYHDAPDIQRSASGSIDRKSMDRNSSASSSHHPIPSLQPPPQSSRSKPNHHERGRSRQRDSLAPQDPYYRERQSSRDRSSLDDPHQSYDPPRTNISRSSSRHNLDRKIGHSSSSVDFQSQKTLAQPPRLSDAYSYDNLRELDSQDGRHRQSFRDVRIDPGLESRESFNRSHSRRPSSLSQSDVRFEDTGTNNAQSPLNKKNALRNSSKHLEANANLRRRGSYNSQEPEKISQKRKEADIRLDKPSSHPRPHRDRHWRSDDRYQDNDYTDSPSLPHRPSPVDSAGIPVSQSLTSFDRSKSGKRYTSPSMPEFATSKPNKNQISTSTPTDNQEINRNSKYLEDERHTRGRSSRRSSKEGVYVDENRNVTMIGETHGQGKQIEKKDGTRGHQRGSVDRETTRRPRRETDNTEADIHYHASGKHRRKHSREEIRPDGDDRYEGDTVPDQRKHTSRGARPRPDEGANRRERLDSEESVSSSQPLLGSGRSPSPIRRRIPKVDLPDPSAHPYLPPSPTYRTNREDQSPRGMSNGKRQPHRYDQRGPEDSPSQEGKTKSRRPSPSQRDISGPVSTPSKRAQSAEHLPETEYSPGMVRRTPGRRLPPVPGEARSSSSLSGTPRPRPRSAGTSRSKDSSPVRQPPSHPRNKEPRHPETRSTPPPFPIDDRPPQYEAVMRTQEQAESSGVDPRLPNGYRPGSVSKGRGHGTQAHSPRRAPQGMSRGHGRRHGREGNGGTLAGYSDTEEDEWA

>XP_011662956.1|Strongylocentrotus_purpuratus_Cav2

MITQGVNFGGVNSASVCNLAFIIQSSEKSPGLTGPSVELPETVKCNDSEGPQFVWVYFIPLIILGSFFMLNLVLGVLSGEFAKERERVENRRAFLKLRRQQQIDKELMGYLEWICKAEEVMLNDKSISEEEREAIEDRRRDAATRTEDLGNGKPSGADVASSYDLDTDKDPNSNSPTKEKKKKKRKKKFVRIRRAEKRLRFAIRHAVKTQAFYWLVIVLVFLNTICVAIEHYNQPHWLEQFLYYAEIVFLCIFIMEMVIKLYGLGPGVYFQSAFNKFDCIVICASMFEVIWTKYKEESFGLSVLRALRLLRIFKVTRYWTNLRYLLISLVHSIQSIVSLVFLLFLFLVIFALLGMEFFGGDFNYDATQAKPSSNFDTFWIALITVFQILTGEDWNVVMYQGIKSQGGVPNGMWASIYFVILVLFGNYTLLNVFLAIAVDNLANAQELTKLDQEEDEENRQAAANELLELQAQQTPNMSRRGSPNPDNPSIEDGTNIQLNDMKDPQRMNGDHNGEMVEEDDPSGDSKSWYRRLYHQLNVHGVPWLCPSCDNCHCPCVCECSCSCQMRKKANNNKEQEAEDENDESFHPKPMVPYSALFIFSTTNPVRRFCHYIVTLRYFDTMIMVVIALSSIALAAEDPIDPDNKRNKVLEYFDYIFTGIFTIEMVLKIMDMGLLLHKGAYMRDLWNILDAIVVVCALFAYAYRGLLPTSMTMLESGGTSGGPQQLNTIKSLRVLRVLRPLKTIKRVPKLKAVFDCVVNSVKNVTNIAIVYLLFMFIFAVIGVQLFKGKFFHCSDLSKVTEEECQGRYFLYEGDQIKESADRVWDKYEFNYDNVALAILTLFTVSTGEGWPDVLKHSIEATEEGMGPTPYNRIEMSLYYVVYFIIFPFFFLNIFVALIIITFQEQGDQDFFDGEIDKNQKQCIEFCITAKPVDRFMPENKNSFRFKVWKLVVSSAFEYFIMTLITLNTFTLMVKYHGQPEIYSAILKNLNIAFTVLFTIEAMLKLTAFGIRNYFKEGWNTFDFITVVGSVADVIISEVGDGFINLSVLRLFRAARLIKLLRQGSSIRILLWTFIQSFKALPWVCLLIWMLFFIYAIIGMQIFGNIAPIDGQQINRHNNFGQFFSSLLLLFRCATGEAWQSIMMSCLPGAACADPDPNNLSGIKEFDTCGSYFSYVYFVSFSFLCSFLMLNLFVAVIMDNFDYLTRDASILGAHHLDEYVRIWGDFDPVGSGRVLYKDMYDMLKNIEPPVGFGRNCPYRIAYKRLIRMNMPVDENKMVQFTSTLMALIRTALDIKIGSVADRDRHDAELREAVKCFWPHLSGDKLDLLVPPDSELIGEKLTVGKIYAALLIYETWREYKAKIQRDGHARTMREVIELVYFKYLYKKYPGFDINDERYQNTFSDEFRRFILAAQKRPSLFRRLVGAVKKNSNENLPGLAEEGGSTSGTVAIKEEKTPSPPPKEKERPRPGKVVTTLPRIDSSGSEGLMVDQPLKQASPKQAERYQRKGRDPDRESRHHRYREPPDNSRTLPLDPSEDCVAPVSGNTDLGLLDLGLDLTGVAQSDQDEFLMSQSTNMTSIYGEYDNHSYEYDEEEEEEVDLGVEGTSALEEQEFEDDKAIEMSDIRYRDRDERGKGGPVDRRERHSMDKHAIQYSNNTTGGNTPYSHSDYSLARGGTGRERERDRERSSERARPRVRHSVAISSDEVPFSDRERFSEERRHYPERDRENASREKRHSLPPVNNFDQMERPRGRPPQSMSQGGLSGRSGSSDGHGSTGNKAIDAILKKGSDIRAQNRQDPTGHGQTPRDGQPRDGQPRDGQPTDRERKMGIPRQRPRQGSLPAPPITGSHSMPAFDDGRPTRQRHSVGSPSMPDFQGRGTPRQPGGGFTSTPAGDDRRDQRRPSGEPSSRKRSNRHDDALIDTRPPQQDIIPEEDEGIPPDGSDVLRRTARNNRKNSGGHLGQYQKFDNSLDSEEPPSQAPGQDVRGERDTVEGSYYNEDAVSSNNRASNYARPTEGASSSSRHERLDSEDSVSSSQPLLSPGRSPSPRRRAPISTRDVSPQRFPPSHSPSSRRRDRHRGPDYVEFHDRDMYDEPSYDDPFGPERSGGSSHRRPSPGWREPRDLLQAGPSQSRASSSERMDPESSQSSPPSRKHTGRRLPTVPGETRGGGGGGGGGSVTPRQRQRPTVPRSRDHSPMRLPPPQHGGRNSGGGSKGDVRLQMDERPPHYDVVMRTQDAPSPDQHVDPRMPNGFRPGGVEKGRGHGTQGHSPRRTPQGVPRGPGRKHVTDNGQGGNGGYSDTEEDEWA

>evg194605+NVE4667(FigShare)|Nematostella_vectensis_Cav2a

MIAYCSLLSLPQLNVGINASSLKALRAARVLRPLKLVSGIPSLQVVMKSIMCAMLPLLQICLLVGFVIIIYAIIGLEFLCGSFHYACHDNSTGVPTLPDEPTICGLKGGFACENNEVCLQKWVGPNDGITSFDNIFAGCLTVFQVITNEGWTDIMYWTFNATDTNGYYFWLYYYSLVIIGSFFMLNLVLGVLSGEFAKERERVENRRAFLKVRRTKQMERQLNGYIDWISKAEDIMLDEEEDDEGERNVDILRKRNLDRVEDGDVAMTAQVLTGKGGRRHRLNTGRKRTCWQRFVRRNKRWRIRVRQIVKHQAFYWAVIICVILNTVITACQHYGQPDWFTQFQDTAEIIFITFFFTEMLFKLYGLGPQLYFKSQFNTFDCVVVCCGIIELVIKNVQGTELGISVLRALRILRLFKFTRYWSSLRNLVTSLLSSVRSIVSLLFLLFLFIVIFALLGMQLFGAQFKSLPGRTGNPRTNFDDFWNAFLAVFQILTGEDWNAVMYDGVLSQGWFDPNKSAQALGASLYFVMLVVLGNYVLLNVFLAIAVDNLANAQQLSADEEEDEKEREERKQQIMAQFQDTKPSPKSSSTGRNGNARRMDDDMPDENEFEEVEEGNKRLTARNFLKSLKNPDNIIEKRNPGEPVPIINTWSLFLFPPGNPVRIGCHYVVNLRHFDNVILVIILISSVLLAAEDPVVEDSYQNQILTYFDYVFTTIFAFEVIFKLIDYGAILHPGSYFRDAWNCIDALVVSCAIASLVLGMNKDASAQSKKTVKVLRVLRVLRPLKAINKAKKLKAVFQCMLYSLRNVLNILIITVLFLIIFSVIGVQLFQGKFFSCNDRSKMTEAECKGHYFVYSDYSDLSKVEVVQREWALADYNFNTVYYAMLALFTSSTGEGWPALMQASIDTTKVDQGPVVDNKIEIFLFYIFFVIVFSFFFLNIFVALIILTFQEQGEKEQGDCELDRNQRDCLHFAMVAKPSERFMPENPNTIQYRVWKIVDSRPFEYFIMTLIALNTLILMMKFYQEPTLYRYYLDLFNSIFTFMFTAEALLKLIAFRTNYFRDNWNVFDFVVVLGSLLDFVLDKALEGNEMPFDPSLFRLFRAARLIKLLRQGYTIRILLWTFLQSFKALPYVGMLIGLLFFIYAVIGMQLFGQIGKNNDSKGEPWTAISGDNHFQSFFPAIQVLFRSATGENWHVIMLACTSGAACEVRANKPADETCGSNVAYVYFVSFIFFCSFLLLNLFVAVIMDNFEYLTRDESILGPHHLDEFVRVWSEYDPGATGRIKHTEVCQLLRQMSPPVGIGKKCPKIVAYKRLIKMNMPLYNDNTVSFTATLFSLVRTSLKIMTEKNNLRENDKELRAMLKRVWPKLTKKTLDKVVPKPPSLINNANKDNSQQQMTVGKIYCAKLIYENYRFVKRKGTGGNQKLMHMQDIPSMYTDGSDLYRSPSQVIYQQNQQTPEPRQRFNSQPIPSTRQSTTSLNDGRGPGVTTAVVSTPGSRGMPRRQLPQQPTYGQSRNGSQISLSGKSPYVENRGDPVLENGYQSSMNQHIAMAIRSGQSPYAIWGLEETGDDDDWC

>XP_020910418.1|Exaiptasia_pallida_Cav2a

MLIVKYFSLLGLQVVMKSIMCAMLPLLQICLLVGFVIVIYAIIGLEFLCGKFHYACFNPGVNGSAPTIAGDDPELCDPDGKGKGCEGSLXCQRYWDGPNDGITSFDNIFAGCLTVFQVITNEGWTDIMYWTFRVYDKDGLAFWLYYYSLVIIGSFFMLNLVLGVLSGEFAKERERVENRREFLKVRRTKQMERMLHGYLDWISKAEDLMLEEDEDEGERNVDILRKRNLDRVEDGDVAMTSQVLTGAGGRRHRLNTGQKRTCWQRFVRKNKRWRIRVRQVVKHQAFYWSVIICVFLNTVITACQHYQQPTWLTQFQEKAELIFISFFFLEMCLKLYGLGPQLYFKSQFNTFDCVVVWCGILELIISHIEGIELGISVLRALRLLRLFKYTRYWSSLRNLVTSLLSSVRSILSLLFLLFLFIVIFALLGMQIFGSQFRNLPGRDENPRTNFDDFWNAFLAVFQILTGEDWNAVMYDGVLSAGWLTPGGNRAGALGYSLYFVALVVLGNYVLLNVFLAIAVDNLANAQQLSADEAEEEEAREERKREIMEQFQDSKYSPKDGGRNGNARRDDDDMGPNEDEFEEVEESGKKLSAKNFLKSLKNPDKIIEQRKPGEPVPIINTWSLFLFPPGNPVRKFCHWLVNLRHFDNVILVIILISSGLLAAEDPVVENSERNKILTYFDYVFTTIFAMEVVVKLIDYGAILHPGSYFRDAWNCIDALVVSCAIASLVLSTNEDSSAGSKKTVKVLRVLRVLRPLKAINKAKKLKAVFQCMVFSLRNVLNILIITVLFLFIFSVIGVQLFQGKFFTCSDKSKMTEQECRGQYFEYSDYRDLSKLEVKDREWANQEYNFDNVFKAMLALFASSTGEGWPALMQASIDTTGIDRGPIVDNKVEIFLFYIFFVIVFSFFFLNIFVALIILTFQEQGEKEQGDCELDRNQRDCLHFAIVAKPSERFMPEDPNTIQYRVWKLVDCRPFEFTIMTLIALNTLILMMKFDNEPKEYRYWLNLFNTIFTFMFTTEAILKLIAFRQNYFRDSWNVFDFVVVLGSLLDFILDKVMEGEGDGQKLPFDPSLFRLFRAARLVKLLRQGYTIRILLWTFLQSFKVRTLINISMSIHFFLFQLFGQIYRENGDRGDPWEQISVENNFQSFTQAIQVLFRSATGENWHVIMLACRYGAXCDPLTGKTGETCGSDITYLYFISFIFFCSFLLLNLFVAVIMDNFEYLTRDESILGPHHLDEFVRVWSEYDPGATGRIKHTEVCQLLRQMSPPVGIGKKCPKIVAYKRLIKMNMPLYPDNTVSFTATLFALVRTSLKIMTEKNNLRENDKELRAMLKRVWPKLTKKTLDKVVPKPPSLINNGKANKENTQQQMTVGKIYCAKLIYENYKFLKKKGQPPQVSSMGGVTRGSFQAPLLGQVVGSPLFAQKIRQEQQPQQDPEWGESIHMQDIRASMPDGPPMYRTHSSQAMNQDYRQRPTLGYQHPGVPTRMSTTSLNDGRGPGVTTAVVSTPNARPPRRQLPQPPMHPGHYQSGSQLSLTGKGQGMPDNRGDPVLENGYQPNMNQHIAMAIRSGQSPYAIYGLEDTGEDDDWC

>XP_020626975.1|Orbicella_faveolata_Cav2a

MADNRRPSSVPSEDFYKTRMANSSNISRKSSMQAGTETSQIVCADRSLFIFSKENFIRKICRTIVESKPFEYFILLTIFVNCILLAANTPLPNNDKSDLNQKLEDAEVYLLAIFCLEAVLKIIALGFVLHADSYLRNGWNVLDFVVVVTGLLSLPQLNVFNAGSLKALRAARVLRPLKLVSGIPSLQVVMKSIMCAMVPLLQICLLVGFVIIIYAIIGLEFLVGRFHYVCNQNVTGRMEITDPDGPQICVAKGTGTRCPEGQFCLKNWEGPNDGITSFDNIFAGMLTVFQVITNEGWTDIMYWTFDAYDAHGYVFWIYYYSLVVIGSFFMLNLVLGVLSGEFAKERERVENRRAFLKVRRTQKMERHLHGYIDWISKAEDLMLEEEEEEDGNVDVSRRRNLDRVEDGDVAMTSVVLTGQSGRRHRLNTGKKKSFCQRFLRRNKRWKIRVRQAVKHQAFYWTVLICVFLNTVITALQYYNQPAWLTQFQDVAEIVFISFFFCEMMLKLYGLGPQLYFKSQFNTFDCLVVSCGIIELIISKAEGTSLGISVLRALRLLRLFKFTRYWSSLRNLVTSLLSSVRSILSLLFLLFLFIVIFALLGMQLFGAEFRDLPGRNGNPRTNFDNFWNAALAVFQILTGEDWNAVMYEGVLSQGYPEKSWTLVWCLYFVLLVVLGNYTLLNVFLAIAVDNLANAQQLSQDEEAEENAREERRREIADQYRDYGSPRSSPGRNGDARRGMGDEAGPTEDEFEEEADDGKPSFLSNLRNPGRMVPHQKPGEAVPMIDTWSLFLFPPGNPVRKACHWLVNLRHFDNFILVIILISSVLLALEDPVDEESKRNEVLTYFDYVFTTVFAMEVLVKLIDYGAILHPGSYFRDAWNCIDALVVSCAIASLVMGNQDQSTIDPGSKKTVKVLRVLRVLRPLKAINKAKKLKAVFQCMVYSLKNVLNILIITILFLFIFSVIGVQLFQGKFFKCDDPSKMTKEECQGQYFSYPDLSDLSNVEIKERSWEGQEYNFDNVFYAMLSLFTSSTGEGWPALMQASIDTTAVDRGPIVDNKIEIALFYIFFVVVFSFFFINIFVALIILTFQEQGEKDQGDCELDRNQRDCLHFAIVAKPSERFMPQDPNTWQYRIWRIVDSSPFEFFIMILIALNTLILTMKFDDEPPLYREILDLFNTIFTFMFTGEAILKLFAFRTNYFRDSWNVFDFIIVLGSLLDFALSRIQSDESDSMPFDPSLFRLFRAARLIKLLRQGYTIRILLWTFLQSFKALPYVGMLIGLLFFIYAVIGMQMFGQITIDNAERGEPWDQIASRNNFQSFPEAIQVLFRSATGENWQLIMLACTADADCQDKEGKCGSAFAYLYFISFIFFCSFLLLNLFVAVIMDNFEYLTRDESILGPHHLDEFVRVWSEFDPGATGRIKHTEVCQLLRQMSPPIGIGKKCPKIVAYKRLIKMNMPLFPDNTVTFTATLFALVRTSLKIMTEKNNLKENDKELRAMLKRVWPKLTKKTLDRVVPKPPSLINNGKSNQVNQQPQMTVGKIYCAKLIYENYKFMRRKGQAQSRGVPMQDVAISLADGPYHSGSAQALYHPNHQQMDSPSRSRYRQNTPVAHRSTTSLNEGRGPGVTTALVGPHRRQLPQTPGSRTGSQLSLTGKAPALPENRGDSVLENGYHPNMNQHIAMAIRSGQSPYAIYGLEEMGDEDDWC

>XP_015773841.1|Acropora_digitifera_Cav2a

MADNRRPSSVPSEDYYKTRMANSNNTSRKSSMQAGGQSSQFICSDRSLFIFSKDNLIRKICRTIVESKPFEYFILLTIFVNCILLAANKPLPKEDKSDLNVELEKAEIYLLAIFCLEAALKIVALGFLLHSDSYLRNGWNVLDFVVVVTGLLSLPELNIGIDAGSLKALRAARVLRPLKLVSGIPSLQVVMKSIMCAMVPLLQICLLVGFVVIIYAIIGLEFLNGKFHYVCHNNETGKIENPDTPQICDPEGKGRSCKPPQQCREFWEGPNDGITSFDNIFAGMLTVFQVITNEGWTDIMYWTFDAADANGYAFWIYYCSLVIIGSFFMLNLVLGVLSGEFAKERERVENRRAFLKVRRTQKMERHLHGYIDWISKAEDLMLEEDEEEDGDRNVDVMRRRNLDRVEDGDIAMTAQVLTGQAGRRHRLNTGKKKSFWQRFSRRNKRWKIRIRQIVKHQAFYWVVLVCVFLNTLITALQHYRQPEWLTQFQDIAEIVFISFFFCEMTMKLYGLGPQLYFKSQFNTFDCVVVCCGILELILSKTQGISLGISVLRALRLLRLFKFTRYWSSLRNLVTSLLSSVRSILSLLFLLFLFIVIFALLGMQIFGARFRELPGRKGNPRTNFDDFANAALAVFQILTGEDWNAVMYDGVLSYDFLKPGKGYALGWAIYFVLLVVLGNYVLLNVFLAIAVDNLANAQQLSQDEEAEETEREERKKNILDQYRERGSPGNSAGRNGDAGMVDDLGPNEDEFESEMDDRPSFISNLRNPGRMVPQQLPGVEVPIIDTWSLFLFPPGNPVRKACHWLVNLRYFDNTILVIILISSVLLALEDPVVEGSYRNRVLTYFDYVFTTIFALEVIVKLIDYGAILHPGSYFRDAWNCIDCLVVCCAVASLVMGQQEGNIDQASKKIVKVLRVLRVLRPLKAINKAKKLKAVFQCMVYSLKNVLNILIITILFLFIFSVIGVQLFQGKFFYCDDASKMTEEECQGQYFEYDDNLSVDKAKNRTWGKHEYNFDNVLHAMLALFTSSTGEGWPALMQNSIDATEVDKGPITDNKIEIALFYIFFVVVFSFFFINIFVALIILTFQEQGEKDQGDCELDRNQRDCLHFAIVAKPSERFMPEDPNTWQYRIWRIVDSRPFEYFIMLLIALNTLILTMKYYKEPKLYRDILDIFNTIFTFMFTAEAILKLFAFRLNYFRDGWNVFDFIIVLGSLLDFGLQQGMQSSGGAKQMPFDPSLFRLFRAARLIKLLRQGYTIRILLWTFLQSFKALPYVGMLIGLLFFIYAVIGMQMFGQISKESEEEPWAQISSQNNFQSFPQAIQVLFRSATGENWQLIMLACAAGANCDERVKDPPETCGSSFTYVYFITFIFFCSFLLLNLFVAVIMDNFEYLTRDESILGPHHLDEFVRVWSEFDPGATGRIKHTEVCQLLRQMSPPVGIGKKCPKIVAYKRLIKMNMPLYPDNTVTFTATLFALVRTSLKILTEKNNLKENDKELRAMLKRVWPKLTKKTLDKVVPKPPSLINNGKSNQANQQPQMTVGKIYCAKLIYENYKFMRRKGQIQAKDAEQGNALSRLVGGHVFQKFRGENDPELGEGLPMQDVAISLADGPYNSGSTQGLYHPNHQVMESPSRSRYPHQVPPVAHQSTTSLNEGRVTTSLVGPHRRQLPHAPGSRTGSQLSLTGKGPVLPENRGDGVLENGYHPNMNQHIAMAIRSGQSPYAIYGLEEMGDEDDWC

>TRINITY_DN6634_c0_g1_i1+evg1115545|Nematostella_vectensis_Cav2b

MYSTFEAIDRDGYLYSVYFVSLIVIGSFFMLNLVLGVLSGEFAKERERVETRRAFFKVRRQQQLDRQVAGYLDWITKADEILIREKSRRESISTVTADTESQAQVISQDTSTNQTSNDRLKQSCMDRFSIEQQRLKLLIRKSVKSQWFYWTVLICVFLNTISLATEHYNQPIWLDEFQDKAEKVFLAIFTLEMVLKMYSLGFDVYFSSSFNVFDCVVVCSGLVEAVLDAVLDQKINLGLSVLRCVRLLRVFKVTRHWKSLRNLATSLVSSIKSIMSLIFLLFLFILISALLGMQIFGGKFIGETKNPRTNFDNFPNAMLTVFQILTGEDWNSVMSYGIMAYGGPKTTWGLVVSLYFVLLVIVGNYTLLNVFLAIAVDNLANAQILTQDEEQEEELKERKKSEHHEKFLPPPPPAETDLRQSKWRKAKPIPVVLAIKNFVRDNKQANGDQLEFDGPPNVGNRRTDRRFSAALGKKIQLYRLEENQTENRINEDGPDGGQSDTGDGSTRKRRVRERRGGRRRTTDKKDSKQNEEDDDEEDDQPITAKNFMNVLHGGRLGNRRRRGLKTVQVIRTRTLFIFGPDNGFRKLCHSIVNLAHFDTAMLVVIGLSSLTIAAEDPLRDDAPRNNILWYFDCVFTAIFAFEVVVKVVDLGLILHKGAYLRNTWNIIDAIVVICNIASLVLSKNDVHAGLAASWIKALRVVRVLRPFKSVHKIKKLQAVFRCMWFSVKNVANILMITGLFLLIFAVIGVQLFNGKFWKCTDEAKLYEKECQGHFFKFKYDKHGVPVGMPTVEERKWEKIKLNFDNVGEAMLTLYTSSTGEGWPTAMHRTMDTTEKDKGPIQDYSTEYAIFYVSFVVVFSFFFLNIFVALIILTFQDLGEKEISNCELDRNQRDCIHFALSAKPAQFYMPRNKHSFQYKVWLVASSRPLDIFIMVLIALNSVVLMMQYYGQSDQYEKACQYLNIAFTSMFTLEAAIKITALRLNYFRDYWNLFDFFIVLGGLLDMAFTIVAHMSEQANSNNGKSYMPIDPSMFRLFRAARLIKLLRQGYTIRILLWTFLRSFKALPYVTLLIMLQFFMYAVIGMQLFGKIALDDDTEINSHNNFRDIFQALQVLFRAATGEDWHLVMTACFSDAKCDKLAGISTGCGSTPLAIIYFCSFIFLCMFLMLNLFVAVIMDNFEYLTRDESILGPHHLDEFVRVWSEYDPAATGFIHHSEVYKMMCSMSPPVGFGKKCPRIVGYKRLIKMNMPLDENGKVTFSTTLFALIRISLNIKLRGNMNANDTELRKLISRLWPNSTSPKVLDKLIPKQSVLSSQQMTIGKIYCAKLIHENYKYSKKKEDSGFQSGLLRRIVGSIRGKGRHSDRISAPSETDSYDQNFLSPRLRSRTSEEPNENSIPRKPVSKKSSDHSSSLVANSIARSHSWSSQANGDIEMSESILRDNLNPASSPSLRNGQARKEEKGMNGPVIPMAYLTSTGSPNPQRKRKNDPHRTIDHSSVHRASRDTVRTGRDGHKPQAPNPTQSRDRSRERPRDQPRDDKYYDDMIVRINEELEQAVKKGQSPYDIYGLEEDDASEWC

>XP_020906771.1|Exaiptasia_pallida_Cav2b

MDGERRYGESSPPKDHYFSTKNSFRMENGSQNNNTTPLAPMETERPRGCCASLLDFIRHDNEHSLYCLST

TNPARRLCRFIVDSKAFEYFILLNIVANCVVLAMNKPLPKNDKIEMAVDLEKAELYFLAIFCIEAFLKIV

AFGFVLHPGSYLRNLWNVLDFIVVLVGIFSLEQLPWNENNXFDVKALRAVRVLRPLKLISGVPSLQVVMK

SIGRAMVPLLQIALLVLFVIVIYAIIGLDFLIGKFHYTCVKNTTXGVKWLSKTNPQPCDGGGKFHGLRSF

LHGRKCNESAGYNCSKVWVGPNHGITSFDNIALSMITVFQCITMEGWTEIMYLTFQAIDLYGYLYSIYFA

TLIVIGSFFMLNLVLGVLSGEFAKERERVETRRAFFKVRRQEQLEREVAGYIEWISKAEEVMTKEARKGD

VRHMSEPSESQAHVISNDLVVNPMAVLTKPVGNGYRLNVPERTSRMDRFHNLESQVRIKLRKAVKSQWFY

WTILLCVFLNTVSLATEHNNQPPWLGEFQEWAERIFLIIFIVEMILKMYSLGMRIYFSSSFNIFDCVVVC

SGIIDTILGEQLNLNLGISVLRCLRLLRIFKVTRHWTSLRNLATSLISSIKSIISLIFLLFLFILISALL

GMQVFGGKFSEKSTPRTNFDNFPNAMLTVFQILTGEDWNAVMYNGMMAYGGPHTLSGVAVSLYFVFLVII

GNYTLLNVFLAIAVDNLANAQILTEDEENEKKEREIKRAKTKEMYSTKSVENRQEPKRISHWGKTKSIPK

ILQLKNKLIQNKELNDVNDIEGGTGRNDAKRFSESLGKKIELYRREDSLYSKKGNGQISGGEGNTLDENS

GRIKRRVREQRGGRLGRGRGGRGKSQSDGEGVKTEEEEDEDGEITASNFMHVLKTGRFGNRRRRTPKDVQ

IIKTRALFIFGPDNCFRRLCHRIVCLPHFDNFMLIVIMLSSLVIAIENPVYDYAEINQYLWYFDCVFTAI

FVFEVVVKVIDMGLIIHKGAYLRSVWNIIDFIVVICNLASLILSKQDKGKTNRIGSSAIRALRVMRVLRP

FKSVHKIKKLQAVFRCMWFSVKNVANIGMITVLFLFIFAVMGVQLFNGKFSYCTDEAKLTQEECQGQYLQ

YQEVDGKLGKPEVKERKWKTRVFNFDDVGHAMLTLYTSSTGEGWPTAMHYTMDTTEKGRGPVQDSNTPYA

IYYVSFVVVFSFFFLNIFVALIILTFQDLGEKEILNCELDRNQRDCVHFALTAKPVQLYMPNNKKTFQYY

VWMLVTSKPFEILIMVLISLNTIVLMMQYDGQHKDYKSVCSKLNIAFTALFVVEAALKLIAFRLNYFRDY

WNDFDFIVVLGGLADIILTFLDREDIPIDPSMFRLFRAARLIKLLRQGYTIRILLWTFFRSFKALPYVTL

LILLMFFMYAVIGMQLFGKIDIHQEDSALTEFNNFRHMPMAVQVLFRSATGENWHEIMRSCFDGAKCDPC

INEENPTSCDPKTAPSLSEDCGNTPLAVIYFCSFIFLCMFLMLNLFVAVIMDNFEYLTRDESILGPHHLD

EFVRVWSEYDPSASGCIQHNEIYKMMCSMSPPVGFGKKCPKIVGYKRLIKMNMPVDDNGAVTFSTTLFAL

IRTSLNIKLKGNMNANDTELRNLMKRLWPNATGKNVLDKLIPKQSVLSSQQMTIGKIYCAKLIHENYKYS

KSKAQGGIERGFLRKLVAGSMRGYHDDNAQGDKVGYNPGPLRPRSRTLDDPHGNPGHYSRAPSDVSSNHH

EYHPNSVSRSQSWSSKANGDLEMIELKPVPTKKSRNGKPRXSDEGSNKPIXPLQYLTVSPKIDRKGRNNS

YSIAVDLNPRDQDSEREASPKKGQPEVSEIRRLSRDSKYYDDMISRINEQLTHAVESGQSPYAIYGMSEN

DENEWC

>XP_020612369.1|Orbicella_faveolata_Cav2b

MDKDKDAATAAKKTGDSSLEMPNPADNERLSYLSQKARSMSLYGSQFLQEARSPTHALFCLSETNPIRELSKSIVVSKVFEYFILLAIGANCIVLALNTPLPNNDRTDMAQQLEDAEYYFVGIFCVEALLKIMAFGFVLHPGSYLRNGWNILDFVVVVVGIISIPEVNKRLNLRLDVKALRAVRVLRPLKLISGVPSLQVVMKSIVRAMVPLLQILLLVLFCILIYAIIGLEFLKDKFHTTCFDIETGARQTNDKPCDAGPFPGVRKIFRGRDCLSGYNCTEYWIGPNDGITTFDNIALAMLTVFQCITMEGWTSIMYKTFDAMDEDGYLYACYYVSLIVIGSFFVLNLVLGVLSGEFAKERERVDTRRKFFKVRRQQQLNRQVDAYMSWIAKAAESAADENYSRSKHLSLNDSQSQLVTNDMVSNPGRVVARREGNCYRVNMPERVGVLQRFIQWQHRFSIKVGRMVKTQAFYWTVLVCVFLNTIVLAVEYYNQPKWLTEFQKYAEIVFLTFFFVEMVLKIYGLGFHVYFSSSFNCFDCAVVCSGFLDLILDTVEGIKLGISVLRCLRLLRVFKVTRHWRSLRNLATSLVSSIKSIVSLIFLLFLFILIAALLGMQIFGGKFTKQPQTNFDNFQNAMLAVFQILTGEDWNSVMYSGVMALGGPHNPGGILASLYFVLLVILGNYTLLNVFLAIAVDNLANAQILTEDEENEKQERELNRARNKQRYTHKGKGWGKAGAKLPVVMAINHFVRKRNGNAQPSAEDPPTTACTTLENGETGVPMSSSLRRKIKSYIREASMYEKAEDRNGTDVAESGEPSGTRSEGGDRTRRIRRMRRGRNEVNGDTEREEDDEDEDDQPVSRRNFMKILKTGGRFGKRRRPSRKKKPIIKTKTLFIFGPENRFRRLCHRIVNLRHFDNFMLVIIMLSSISIAIEDPVNDNSKRNEVLMYFDYVFTAIFALEVIIKIVDVGVIFHKGAYFRDWWNVIDALVVSFNMASLILVQTDTSGDGRSVIKALRVFRVLRPFKGVHKIKKLQAVFRCMWYSVKNVANILMITALFLFIFAVMGVQLFKGKFQNCNDESKLWKEECQGQYFIFDYDEFTQREEVSKVKDREWETKALNFDNVFKAMLTLYTSSTGEGWPSAMQATMATTEVDKGPIPNYSPGYALYYISFVVVFSFFFLNIFVALIILTFQQEGEREIASCELDRNQRDCIQFALTAKPAQRYMPADHKSLQYKIWVLVMSKPFDTFILVLIALNTGVLMSQHYKQGEQFTDILMYLNIAFTILYIVEAGLKFIALRLNYFRDYWNIFDFIVVLGGLLDVVVTVVDRFFKTGGDNTLGIGIDPSMFRLFRAARLIKLLRRGYTIRILLWTFLQSFKALPYVTLLIMLMFFMYAVIGMQLFGKIELDEDGEINANNHFRNLLEALQVLFRSATGEDWHKIMRACYDDAKCDPQVDTEKYGSTCGTTVGAILYFCSFIFLCMFLMLNLFVAVIMDNFEYLTRDESILGPHHLDEFVRVWSEFDPAATGCIHHSEIYKVMCSMSPPVGFGKKCPKIVGYKRLIKMNMPLNEDNTVSFSTTLFALIRTSLNIKLRGNMNANDTELRRMIKRVWPKTSQKVLNSLIPKQLELSCQQMTIGKIHCAKLIHENYKYLKRKGTQPERVEREEERRAANSKRGLFSKLVGSLRRNRRPNTGRENIELGNMERRRNRAYTSDARLNVARRLPRQASMDLKRHSSWGPSPKLNGDVNRNGMQPHPDSSRAVMEASVPCSVATPNVAEQFKDQAVLPAAYLTPTTRRSDKVLVEGASLDLDSSSDSDTGVEMDKDRPQVKQDPYFDDMITRINVQLEEAVARGTSPYAIYGLEDDDADDWC

>XP_022780503.1|Stylophora_pistillata_Cav2b

MDKDRYAATAAKKTGDSSLEMPIASVDNNERLSYLSQKARSMSLYGSQFLQEARSPTHALFCLSETNPLRELSKSIVVSKVFEYFILLAIGANCIVLALNTPLPNNDRTDMAQQLEDAEYYFVAIFCVEALLKIMAFGFVLHPGSYLRNGWNILDFTVVVVGVISLPQVSAILGGNDTLDVKALRAVRVLRPLKLISGVPSLQVVMKSIVRAMVPLLQIALLVLFCILIYAIIGLDFLKDKFHTTCVNKTTNLTASSNPKPCDKGPFPGVRRIFRGRSCENGTQECREYWIGPNSGITLFDNIALSMLTVFQCITMEGWTSIMYDTFDAMDTDGYLYACYYVSLIVIGSFFVLNLVLGVLSGEFAKERERVDTRRKFFKVRRQQQLNRQVDAYLSWIAKAAETTADDNFRTKHLSLNDSQSQLVTSDVVSNPGRVVTRREGNCYRLNVPESTGALQRFLQWNSKLSVKVGHMVKTQAFYWTVLVCVFLNTIVLAVEYHNQPRWLSNFQEWAEIVFLTLFFVEMILKIYGLGLHIYFNSSFNCFDCSIVISGFLDIILSKAVQIKLGISVLRCVRLLRVFKMTRHWRSLRNLATSLVSSIKSIVSLIFLLFLFILIAALLGMQIFGGKFTEEEIPKTNFDNFQNAMLAVFQILTGEDWNSVMYSGVIAYGGPHNPKGIAVSLYFVLLVILGNYTLLNVFLAIAVDNLANAQILTEDEENEKLERERARAANKKMYQPSGKGWAGKGWGKAGNKLPMIMAINRMVRKKNGNAATSAGEGNSTTLENGETSVPPISSSLKRKIKTYRREASLYENKEERNGTAANPKETPGTSSDADEKKHKTRRIKRGKKEANGDTEREDQDDEDEDDPPVSRRNFIKMLKTGGRFGHRRRHPRKKKPILKTKTFFIFGPENRFRRQCHRIVNLRHFDNFMLVIILLSSITIAIENPVNDDAKLNRVLKYFDYVFTGIFALEVLIKVVDMGIILHKGAYFRDWWNIIDALVVSFNIASLILVEIGQSGSGHSLIKALRVFRVLRPFKGVHKIKKLQAVFRCMWYSVKNVANILMITMLFLFIFAVMGVQLFKGKFQYCNDSSKRTKEECQGKYFVFKYDANLQREEVEKVEDRKWSNKDLNFDDVLQAMLTLYTSSTGEGWPSAMKTTMDTTEIDKGPIHNYSPGYALYYIAFVVVFSFFFLNIFVALIILTFQEEGEREIASCELDRNQRDCIQFALTAKPAQRYMPADHKSLQYKVWVIVMSKPFDTFILVLIALNTGVLMSQHYKQDDQFTDILMYLNIAFTVLYMIEAGLKFFALRLKYFRDYWNIFDFIVVLGGLMDVLVTVVDKYAKETQKGGTLGIGIDPSMFRLFRAARLIKLLRRGYTIRILLWTFLQSFKALPYVTLLIMLMFFMYAVIGMQLFGKIALDSETEINSKNNFRNLLQALQVLFRSATGEDWHKIMLACYDAAKCDANVNEKYSCGTTVGAILYFCTFIFLCMFLMLNLFVAVIMDNFEYLTRDESILGPHHLDEFVRVWSEFDPSASGCIHHSEIYRVMCSMSPPVGFGKKCPKIVGYKRLIKMNMPLNDDNTVSFSTTLFALIRTSLNIKLRGNMNANDTELRRMIKRVWPKTSQKVLNSLIPKQLELSCQQMTIGKIHCAKLIHENYKYLKRKGTQTERVEREEERSATNAKRGLFSKLVGSLRRQRRPNMIREDIELGNVERRRERAYTSDARLTMARDLPRERVRRHSSLGPTSKSNGDVNKNGLQTHPDSSRGVVGAPVPCSVATPRVAERFRDQAVLPALYLTPTTRRNERALMEGKSLDLDSSSDSDTGVEVDKDRPHLRRDPYYDDMITRINSHIREAVERGTSPYAIYGLEDDDADDWC

>TRINITY_DN35757_c0_g1+evg1243599|Nematostella_vectensis_Cav2c

MADESTLSNRSAESKLQAATEFFQRKYKSVSNAENSQRAYKPNALFCLKENNKLRVNCKKIVDSKFFETSILLIIAANCIVLILDTPLPKGDSTDLNKTLEQAEYVFVVIYCLESALKIIAQGFLFHEQAYLRNGWNILDFAVVVVGLVGMVWDLDGAGNRNLNQQKDSLKVLRAVRVLRPLKIVSGIPSLQVVMKTIWRAMIPLLQILLLIIFVIVIYAIIGLELLKGKFHSTCYDANDRMEKGYLFPKICSNDSAGRQCSDGHTCKTLDHIWPGPNKGITNFDNIFLSMLTVFQCITMEGWTDIMYHSYDARDYKQGVVTSIIYISLIIIGSFFMLNLVLGVLSGEFAKERERVENRRKFFKLRHQQQMERQLSGYMDWIARAEDIMLKEDMKQHGVGEGGPRDPLRKRTSLSDSLAHLVEDQKMFLNRLKEKKKKDSQDRSSSIMEKIKNQLLRRKVKVMVRSQIFYWAVLVCVFLNTVLMSFEHYGQPDWLERTQTIAEKVFLGIFIAEMLLKLYGLGATQYFKSSFNRFDFVVVLSGIVEMFLQKYLHISFGSSVLRSLRLLRIFKFTRFWSSLRNLVTSLLSSMRSILSLIFLLLLFIFIFALLGMQLFGGRFSTTLDAPRTNFDNFVKAMLAVFQIMTGEDWNTVMYNGIEAAVGPKNVFGILGSLYFVALVIVGNYTLLNVFLAIAVDNLANAQALTRDEEQEVRMREQIKKRRNEKRRNGWAKAKQLPMIMAIAKLNHNSKNNPFPEVKPVYHSPRGAGGWNKKSALKLKRQTTEDTTSDALQNGRISRQDTQDNALTDPVTDREDEDEPADMPTPEMVRTPRRSLAISSLRSAGRIIRRRNIHRTSPIIRKSSMFIFGPDNPIRRLCHWVVNLRYFDTFILFIILISSVLLVFEDPVSTNSQTNTILGYCDYVITAIFGLEVLFKVIDLGVILHKGSYFRDAWNVIDAFVVACNIAALVLNAKAVADKTSNTSVQEAIKSFRVLRVLRPVKAINKSKKLKTVFQCMVYSLKNVRFILLINLLFYYIFAVIGVQLFKGKFFYCTDMSKMQKSECKGHFFRYMVGVDSQVSLDNFELGERNWTRWHFNFDDVPSAMLTLFSASTGEGWPTAMYHTVDATHVDRGPRRDNNIQMSIYMVCIVVIFSFFFLNIFVALIIVTFQEQGEKEMVGCELDRNQRDCIQFAMTARPRQRYMPENQKTCFYKVWCVVDSKPFEIFIMTMIVLNAIVLMMTYHGATQEFNNIIEYVNMAFTFVFLFEAILKLIAFKLNYFRDYWNVFDFIIVVTTLVGVLLELVQDTKNQLDIDPSFFRLFRAARLVKLLRQGYTIRILLWTFLQSFKALPYVVMLIAMLFFVYAVIGMQLFGRIAKGIPNREINIHNNFQSFFQALLVLFRAATGENWHLVMLACFDNAPCEKGGKACGNTAASIIYFITFYFFCSFLMLNLFVAVIMDNFEYLTRDESILGPHHLDEFVRVWSEYDPGATGRIKHTDVYHLMCDMSPPVGFGKKCPKFIAYKRLIKMNMPIMGDNTVLFTATLFALIRTALGIFSTGDPAYADSELRRTIRRLWPKTSKKTLEKMIPLQSVLSSQQMTIGKIYCAKLIYENYKHAKKKRMERKKKGRRPSLFRRFVGAFRNNPSNSEAEDEESELRAPPDYLSHRRRSKTLSSLPTFIHNEPETTTNNKRRMHRSLKFFRRPFGRTESNSDTDEARSARKTVHFKDAPMDLTDINPTIISTDTEDDDQGEPVNRKRKPSKLGYKNPVALEDAQSEISSSMESHPSRSSQDPQSPLSPPMVHIELHPPLTLEPQSLVVNDDRGMMRLDPFPDVIKSAANSPITEFFPRVAPANPAERRSREIINQINEEITQAVRSGQSAYFIFGMFDNDEETWC

>XP_020613420.1|Orbicella_faveolata_Cav2c

MADESKDPIKNSENKLRAATEVFQRKYRSMSTYGEQVIDDANRSKKALFCLPEDNPVRFYCKKIVESKKFEYFILLTIAANCVVLMLEEPLPNGDTTDRNKKLEESEKYFVIIYCIEAATKIIANGFLLHKDAYLRNGWNILDFVVVVVGLVGMISDLEISGDGDSNSIKESESLKVLRAVRVLRPLKIVSGIPSLQVVMKSIARAMIPLLQILFLILFVIVIYAIVGLELLHGKFHLTCYDAITGELDTKFTSPRVCSPPGKGGRPCEPGLNCTSKESVWSGPNKGISTFDNIFLSMLTVFQCITMEGWTDIMYHSYDARDYNARVVTSIIYISLIIIGSFFMLNLVLGVLSGEFAKERERAENRRTFFKYRSREKIERQVNAYTDWIGRAEDILLREEREKHGVGEGGPRDPLKKRYSLSDSIMHLIEDHGEMVKYLRNKSESSSTMRKVKKKEKLLRIQVRHMVKSQVFYWSVIVCVFLNTVLMSVEHYGQPDWLEKFQEISEYVFLSIFIAEMLLKMYGLGPRVYFKSAFNRFDCAVVLGGIIEIVVQTFTDYSFGISVLRSLRLLRIFKFTRFWASLRNFVTSLLNSMRSILSLIFLLFLFIFIFALLGMQLFGGKFSERHDAPRTNFDNFLKAMLAVFQIMTGEDWNAVMYDGIVASKGPHTITGMLSSLYFVSLVILGNYTLLNVFLAIAVDNLANAQAVTQDEKEEQRQLEAMRKKRLEKKRDGWAKARQIPVLMAIKNINHSKNDNDNPFHNMKPVYPSLDHVSPRGARWNKKSALRVKISQDGKFPQATNGQMSNGNSKENPDKEEEGEDTENSVPRRTLLSLRDAGRIIRRRNIHRNVPIIRKSSMFIFGPDNPIRQACHWVVNLRYFDDFILAVILISSVLLAIEDPVHPDASRNKVIRYFDYGITGIFALEVLVKMIDLGVILHKGSYLRSGWNVIDAFVVGCNIAALLLDIGTDDNLQKDAIKSFRVLRVLRPLKAINKSKKLKAVFECMMYSLKNVRNILLITLLFYFIFAVVGVQLFKGKFWYCTDLSKMTNATCQGQYFKYNVAINSKISLDNFEVKDREWKKHKFNFDNVPHAMLALFSSSTGEGWPQGMYHTVDATKEDQGPIKDYQIQMSLYYVCFVVVFSFFFLNMFVALIIVTFQEQGEKEMDGCELDRNQRDCIQFAMTAKPRQRYMPEDKNTCVYKVWKVVDSKPFEIFIMATIVLNAIVLMVSYDDASPQYERILINLNAAFTFVFLSEAILKLIAFRQNYFRDFWNVFDFIIVITTLVGVILELNSTTQKASDALPVDPSFFRLFRAARLVKLLRQGYTIRILLWTFLQSFKALPYVVILLGMLFFVYAVIGMQLFGRIRPSEDWSKQINHHNNFRNFFMALQVLFRASTGENWHKIMLDCFDDAKCDSDENRSCGSTVASIIYFCTFYFFCTFLMLNLFVAVIMDNFEYLTRDESILGPHHLDEFVRVWSEYDPGATGRIPHTEVYRLMCDMSPPVGFGRKCPKFIAYKRLIKMNMPIMGDNTVLFTSTLFALIRTALGIFSTGDPAYADSELRRTIKILWPKTSKKILEKMIPLQSVLSSQQMTIGKIYCAKLIYENYKHMKKKRLEKKKKRPSLFRRLVGALRSGNSNSDDEEGEIDTPSDYLTTTRRRSKTLTALPTMSKEKEEVLNSKRRMHRSLKFFSRRPFGRSDSYSEDEDFRGINKSVHFKDAPMDLTDINPTIISTDTEDEDQVDGACPKLKRKPSKLSFKNPVALEDEMETASSESRSRSSTGLSSPLSLPMVKIEVTSPQPSGAQGPGEEPAQRLAGDRSLMLLDFPDVIKSAESSPRHSLIPLSEFYESQGRTLNDRRTREIINQINVEVAQAVNRGQSPYYIYGIMDNDEETWC

>PFX33508.1|Stylophora_pistillata_Cav2c

MADEAKDLTKNSENKLRAATEVFQRKYRSMSTYGEQVIDDANRSKKALFCLPEDNPIRFYCKKIVESKKFEYFILLTIAVNCVVLMLDEPLPNGDTTKRNEQLEKSEKYFVIIYCIEAATKIIASGFLLHKDAYLRNGWNILDFVVVVVGLVGMISDPEISGGSDSNSLKESESLKVLRAVRVLRPLKIVSGIPSLQVVMKSIARAMIPLLQILFLILFVIVIYAIVGLELLRGKFQWTCYNITTDGLDTKFITGSRVCSVPGNGGRPCDVGLRCKNNQSVWRGPNKGITTFDNIFLSMLTVFQCITMEGWTDIMYHSYDARDYHTRVITSIIYISLIIIGSFFMLNLVLGVLSGEFAKERERAENRRTFFKYRSREKIERQVTAYTDWIGRAEDIILREERERHGVGGGGPRDPLKKRYSLSDSIMHLIEDHGEMVKYLRNKSENWSESSSTMRKLKKKEKLFRIHVRQMVKSQVFYWSVIVCVFLNTVLMSVEHHGQPDWLERFQAISEYVFLSIFIVEMLLKMYGLGPRVYFKSAFNRFDCAVVLGGIVEIVVQNFTNYSFGISVLRSLRLLRIFKFTRFWASLRNFVTSLLNSMRSILSLIFLLLLFIFIFALLGMQLFGGKFSERLEAPRTNFDNFLKAMLAVFQIMTGEDWNTVMNDGIVASGGPHNIWGILSSLYFVSLVILGNYTLLNVFLAIAVDNLANAQAVTQDEKEEQLQPEAMRKKRMEKRRDGWAKARTIPVLIGLGKINHNKNDNDNDNPFRNMKPVYPPLDHVSPRGARWNKKSALRVKISQDANGYLEANGKFLNGNSREGGNTDREEEGDMADQTVPRKTMLSLRDAGRIIRRRNVHKNVPIIRKSSMFIFGPDNPIRQACHWVVNLRYFDDFILVVILLSSILLAVEDPVNPEARRNKVIRYFDYGITGIFALEVLVKMIDLGVILHKGSYLRSGWNIIDAFVVACNIAALLLDIRGSGEDNLQKDAIKSFRVLRVLRPLKAINKSKKLKGKFWYCNDRSKMTRETCRGSYFKYNLGLNSNIDLEKFKVTQRNWTKHKFHFDNVPNAMLALFSSSTGEGWPQGMHNTIDATKEDHGPIKDYQIQMSLYYVCFVVVFSFFFLNMFVALIIVTFQEQGEKEMDGCELDRNQRDCIQFAMTAKPRQRYMPENKNTCFYKVWKVVDSKPFEILIMATIVLNAIVLMVSYDGESSEYEKVLDYLNYAFTFVFLIEAVLKLIAFRQNYFRDFWNVFDFIIGVTTSVGMILEFTKALPYVVILIGMLFFVYAVIGMQLFGRIDLSENWSRQINHHNNFRSFLMALQVLFRASTGENWHKIMLDCFDDAPCDSDKSKTCGNTVASVIYFCTFYFFCTFLMLNLFVAVIMDNFEYLTRDESILGPHHLDEFVRVWSEYDPGATGRIPHTEVYRLMCDMSPPVGLGRKCPKFIAYKRLIKMNMPIMGDNTVLFTSTLFALIRTALGIFSTGDPAYADNELRRTIKRLWPKTSKRILEKMIPLQSVLSSQQMTIGKIYCAKLIYENYKHMKKKRLEKKKKRRPSLFRRLVGALRNGTSYSDDEEAHIDTPPEHLTIRRRSKTLTSLPSISKEKQEVINSKRRMHRSLKFFSRRPFGRSDSYSEDEDFRGISREVHFKDAPMDLTDINPTIVSTDTEDEDHVDGACPKLKRKPSKLSFKNPVLKVLSSFPWGNESQPNTFHCLANTALEDEMETESFESRSRSSTGLSSPLSLPMVRIEVTSPQPSSVQDPEEETAPQIARERLMVLDFPDVIKSAESSPRHSLIPLSEFYESHRRTLNDRRTREIINQINAEVAQAINKGKSPYFIYGIMDNDEETWC

>evg1041627|Trichoplax_adhaerens_Cav2

MASSSFNSSVTLKTNNLSETVRHLVHEKIAANNANKQQGIIRAYFRRIWQYDRDKSLCCLPANNLLRKYAKKLVDWTPFEYLVILTIVANCVVLAMDVPLPDNDSTEISLLLERNAEIGFLVVFCIEAALKIIAKGFFFHPQAYLRSGWNILDFLIVVVGLVNAFYINAASNEIDVKILRVVRVLRPLKLVSGMPSLQIVLRSLLTAMGPLFQISLLVLFVIVIYSIIGMEFFLGKFHLGCRDPRTGQLLTQNHMSPCNNQPNSSAGFRCVINTTQYQVFGTCDYKYDGPNYGITGFDNIFMALLTVFQCISLEGWTNLLYDTNNAVGSTFTWFYFLTLIIWGSFFMLNLVLGVLSGEFAKERDRVEKRREYKKFQENRKIERDFLGYLEWIGRAEDLILGEQRLKEEETAPHFDTRSEIFQFAEQDQVAELAEITLSENPINTKAYATTTHSPRANCLHMVQRSEKFLRLAIRRTVKSRPFFWIVILLVFLNAVTIASEHSGEPLWLKDFREATNIVFVALFTLELILKLYGLGAVFYFSSTFNCFDFAGVIASIAELKVRNVGGPKLGIRGFPCIRLLRIFEITKHWKSLSNLVASLISSLRSILSLLFLIGLCIMVFALLGMQLFGGRFNFAEGVPRSNFNDFGHAVLSVFQVLSGEDWNEVMYNGIRAYRSSGEFVSYAVSLYFVVLVCLGNYTLLNVFLAIAVDNLTKAQEISKDEDEEIMLQKRLSIKRKNYSKDSGCQSALGDGQYGTDDRPRESIRTLSIASTDGRSNQRSRRSTLRIPMPSSPSINDDNAFPEFCDNNTSQQETRIDEINDIENPGNENDDEEPNFRQKRLGSTKMNFQDAHQEPELIIGKEIFAQNVPILQVNSLFIFSPQNRFRRFCHYIVHLRHFENFMIAAIIISSGLLAVEDPMNEDPVLNYVLRIFDSIFTGIFLIELILKVVDFGFILHRGSYCRNLWNILDMIVVVTAVTSFIYFEIGVENQSNRNISAIKAIRTLRVLRPLKAIRTAKKLLACFQCMVNSLKNVLNVCIVMLLFLFMFAVIGVQLFKGKFHYCTDQTKHTKSQCRGNYYHYTDGHVYNVEVLERRWEQHPYNFDDLPRAMLTLFTMSTAEGWPRILYWSIDATNENEGPMRDYNLAVALYFCIYIVVFPFFFINIFVALIIVTFQEEGDKDIANYQLNRNQRDCIEFALNAKPIHRHMPKDKKSYAYKVWRIVTSTPFEFIIMVLIIVNTIVLMMEYNGQSKDYKDMLQIINITVTILFTVEMLLKVIAFSPRNFIKEWWNIFDLIVVIGSWTDIIITYASINGTSTVSISFFRLFRAGRLIKLLRKGYTIRVLLWTFLKSFQALPYVGLLIGMLFFISAVLGMQLFGQIQSDPTTAIFRYNNFQTFTGALIVLVRCSTGENWPEVMLACLPGRAKCSTKFPDCGSYVAYPYFVIFVFLSTFLMLNLFVAVIMDNFKYLTRDKSILGPYHLDHFLHTWAEFDPEASGRIKHQELCTMLCRIPPPLGFGSSCPMRTAYRRLMKLGIPINGDNTIRFKATLFALIRTSLKIKVREDQHEADAELCQIIRKIWPQVTSRTLERILPQVEHGARHLTIGKIYAALIIYEYYKRYKKQQLREEDEQVRMRTKSLFHRFIDVVRTPIRTGAHKDLEIPTEYPTKQLRSKHFHSHSFKIPRPSLRRSRKRDKMKSKHAVSAKPSDSSVSGENNRSRSLPGLIFESIDDNAGKVQDINLYLANRSSKETDFCIAEVHHEDDEGYLQSEYRNNTDERSQTLDPLLSNRRQGSRFNIYNTSGSNLQPSPLTFEPGRPLHRMIRTTPIGTPKSPQKISIVQSTYNSANNSPTTIHDRYHSGSANNSNNINGGSYSPTRNQHQFKLKLQHEDSNDSCTGSLGLRLQKSGSSCGSTDDDLDLNDAIQFNNLQDLLPVDIPDVTISISDSPYGSTDGLDKQYASDLNGATEYKRLPSNAFWSSVQRNYNKVLSKSSNNSLNNCNGASNLSQDYSNNHTNSTGNNASSRGTNNLHHQRHFSNNHNSTLNHSGHQSDYYKNDYNGIRNDDYDELELNRNLMLSNFSFNSDDLTNISKCTAV

>XP_004989719.1|Salpingoecca_rosetta_Cav1/2

MTNDDLSSRETDSLEYVFLAIFTLEALLKIIATGFLFCGPPSYLRNKWNILDFIIVAVGLIGVVVEQSGSSVADVKALRALRVLRPLRLITSVQSLQIVLNSILLSIPALADVAMLLGFLIVIYAIIGLEFYRGVLNHQCFLPSSDVGANVTADRYINNTPYFLAPDTAPCDPAGRGRVCSTDGLCLAGSSPNSNITAFDHAGSFLKFDAAESMAEHEQNFFNSSAGADGNDGDDDDDDIDDDATTVLVLGLSLPSGNRDFVEAEPTEPISNIRTRIMQKFAEQGEDRDKLMSFVLAHPHDGHILDETRTLGEQGVQVKSLREYNEILTMSQHNQHELLKPATVQMLSKLGAVVKSRWFNLVVTFMVLVNTVLLAVQTDAGATDEAAFAFTIVEASFVGLFVLEMLVKLAGLRPHMYFESKFNRFDLTVVLLSLLELILVHTTGLRSIGISALRSLRLLRIFREMKQYWEDINDFVVSLLNSIASIVSLLLLILIYMVIVALLGMQIFGGRFDFEDPKPRINFDDFFSALLTVFIVIVGDDWNSVMYNGILAYNGVNKDGWVAIVFFCVVVILGMFVLLNVFLAIAVKSLDDARDLKAARDEHKERWKAEAAVSDESEDDREDRRRHRQYANPLVGAAEQEKEEVELQNTVLCDVDNVPLRKHLTRAVANNKSLFCLGPRNSFRKFCNNIAYDNRFESVILLLILISSALLAAEDPVNLDAQINKDLETADIFFTSVFSLEMALKIVALGFIPYITDPWNDLDAVVVLASVVSLAISSDDAAVVRVLRVFRVLRPLRAIKRAPGLRKVVSCMVVSIKTIGNVFIVTFLLTFIYAIIGVQSFKECFGRCNDPDVMFKSQCNGTFLVEDDAGLFSNATRAWSTPYFNFDNVGKGMLTLFTVSTLEGWIDVMNNAIDCTAENRQPERNNNPVAALFFVTYVILVAFFMLNIFVGYVIITFSSEGESYEAVDGLDKNQRKCLAFCLNAQPIRVHRPLYRAQISIFRFVSSKHFEWFIMAAIIGNSIVLLMAYEGMPSDYEMGLQLCNIVFTGIFTVEALLKLFALNPTGYFHDSWNFFDFIIVVGSLVDVFLSATQSSGDSGVNIGFLRLFRVARLLKLVSRGKGMKRLLWTFAKSFQSLPYVAALIMMLFFVYAVIGMQLFARTGFREDGDINEHNNFRDFFGALLLLFRCATGENWQNMMRDIHLGPPNCDPATEPGVCGSVVAVPFFCTFLVLCSFLILNLFVAVIMDNFEYLTQDNSLLGEHDLPQFIDRWSEFDPACTHRISHHDLMELLRSEEPPLGFGRKCPPKTVYSKMMRLNVPLHLDGTVDFHACLLAIVRNQLHIKTSYLDGSWEQRNYDLRKLLEKLYDPPKEQLDRMLPPPSKRNVTIGVLYAVYLLQEIYRENKRNAARQARETAEVLDAGSNEEKPQPSENEEVEDDLRLVEHAF

>Aqu2.38198_001(Fernandez-Valverde_2015)|Amphimedon_queesnlandica_Cav1/2

MATDRYASSLSPRPSLLARQSSMLGEEDIERSLSPTTMQTRLRHMSKGYVGGIDLKSTLLDEDEGVWSHLMKSRHVVKFRSYLRQRARNDYILCCLPKKNRWRKRIKTLVYSRYFNGVIGLLVIIHCIVLATYKSYSANDLKGYNKNLLYSTIAFLLVYIVEACLRIIADGLIMHPSAYLRKIPNLVDVFVIFVSFIYLVTPFGLVIGEVFAALITLRLFRVMLQLKTVRFLFRALGSSLFPLLYVAWVTFCIMMFFALVGLELFSGGLHKACYVNYTDTDTLVMRQFQVSQFPCHDPDYFGGHNCSLATDKPEDAFCENWPEGPHSGLVGFDNIGIGLLTVFQCITLEGWTSILYQYESVYGTSFVWIYFISLISFGSIFMLNLLLGVLTSVFIGVSDHQEVEGHLRVLKKSRKIKEDYDGYNEWMKRGGYCHHGGGANENDDGDVDDDDDQEIDEEFNKHANSEMNNSTCNSTKWGPFLFFNKRLRSVIRHVVKSHVFFWSMIAIISINFMFLSADFYPIDKEWISSLIIINYIFTGIYIIECFLKLYSLGPRRYFTSQFNRIDFLFTLINIIDILALVHISDIYFSVSSIVNACRAIRLISAFKYTRYWKGMRAIISTFAGVGVVILSVMALLLSFIFTASLLGMRMFGTRLFDITHLHPTFNDFVDSFLLVFQLTTTEDWNTIMYRSGLTSDERFTYNIGNFAIIIYYLYIIIIGAFNIVNIFLAIAIDKLSEVKAVNEESTHRLEQREEERKELEEQLNALKDPLQYFKLSQTLRKVLIFYSSNLKEEEELQETRKRRRERSVSEAHVKVDRSVSLPLAEIKRRSFASAHAVSDPGRMTEEEGERREQLKHYLTGMVPQKVSIANPLGKFFKLQHSASSMATASQLNTGVKAMTLELTETTPPTSRADQSKSKKPDIDKDEEEVTDFCRNNSINSLGDCSLELNTPASLVAPPSPLDPPTDASVARLVRKSSLEDIDLIIPPESERSGMANGDKGLAMKPTKFKKIRRAWSIFKEKGSLLLHWLSNYFGSIIIDPRLELKEIPAHSAFFILAPDNRLRVYFYNVVKSKPFQTVTFTIIILSSLLLTLEIPVRSESSLLCGIQGTIFFFDIFCSVWFLLEFILKIISLGAIIHRGSYFHCLFNIIDFFVVVTTIFPLILHLATANLNDDGICLPYSQRLYHEQNSYMYIEVILVFRVLRPLRLWRVEGLFLVTKGLFSSLRRMGYVFFVGTVLLLIFAVIGVQLFKGRFFYCTDFVSHTEEECRGQYFIYPSNDLNCPKVLNREWKKWDLHFDNIGWSFLSIYTMITKEGWQDIMYHAIDSHDVGEGPVYNFSRWALVYHVLFMVLVTFFLINLLVGFVIVTFQQNGIKRYDEANLDRNQRNCLYFSLTSVPRKKYIPQFSFHKRLYSVINSWLWRLVINILVAINVIILSCQYYTNNNTNNMDSPSKLEVTIYWINFGFTILFTLEAVFKIIILTPPHYFRSSERGFEFLLVIGSVLELTLQSVLVRDSNSIWRYSQIASCLRVLRLVQISKNTRLIVWTVLRSLEIFPWVGVLLLAVLFTYAIVGMQVFGRIKPVTIDNSTHNAIHQYNNFANFPQALLVMIRCFTGENWEQIMLGSVNALCVDEVQNQTTTCGSPFTYFFYPSFLLISSILVLNLFVAIIIDNFDYFVRDKAILGSHDLTYFYQLWAKLDPSASGKIHHTELIKLMRSANPPLGWGRLCSQVSCYKRMISLSIPVDEDGMIAFNATLFAIVRVSLKIDSSCGKNNNNNYNSRSADLRLKLQSVFPNCPKKILDIVLPERPVRTTSEEYAALFIQQFWRRWKLKVEKQREGKPQPHQQRMSNVLRRVPDMGPTLLQRRLTTINMVNVSSRLAVSPSDTSADDTSHGLREIASTDVPVLKPIRSLHLIRLEPITEGNDQESASVDQNSASRDQKPAPVPRKSVLFDQESSALDQKSASIITKAKSVDSDIFYDPEVDRSPSLIAKRQSVISDSDALFSTPTGSMISLDSAKAITETTV

>m.28368+m.28207(Compagen)|Haliclona_ambioensis_Cav1/2

MAAPFERPSSMVLEDGNVSDYDLERSLSPTTRQTRLRHMSKGYVEVNLRQFSYASDEGTGVWSRIVQSKRAQHIRSYLRQRARNDYILCCLPKKNRWRKRIKTLVNSRWFNWFIALLVVAHCIALATYQTYMAGDYKPRNINLIYITVVFEVLYIIEAGLRIVAAGLIMHPSSYLRKLSNVVDVIVIVISFVFMLRPDNRIFETLSAFISLRLLRVMLPIKTIRFLLRALAKSLLPLLYVAWMTFCIMVFFALVGVEIFRGGLHSACFVNYTDSQSLSPKQYQINEFPCSKVDRPGSHDCARISERGYCDYWEGGPHSHLVGFDNIGSALLTVFQCMTLEGWTSVLYLYESAYGTHYVWLYFILLISFGSIFMLNLLLGVLTSVFIAVSNQQEAQGHLRVLKQKRKIKEDLEGYRQWLKKGVLEEEHSTNITRSRSLQRQLTHYGSMDSSWSDSEEDEDDVEDEEAATTSSKWEPLLTFNRRLRSVVRHIVKSNEFFWFMIVVVLTNFILLAADFYPSPSKWKLALLIINYLFTAVYILEFVLKFYSLGPKVYFYSKFNRIDFVFTMINILDILVLAYISAIYFQVSAAVNACRAIRLVSAFKYTRHWQGMKALTSTFVDLWLIILSVLILLGSFLFIASLVGMRLFGTRLSAYGSLYPSFFDFLDAFLLVFQLSTTEDWHAIMYRTAIKEDKASNEALFRIGNFITAAYYVFVIILGGFIIVNIFLAIAIDKLSEVKAVKEDSSYRSEQREEQRREREEQLLALKNPTQPSGLSQTLRRVLVFYSANLKQQEELQMTRRRRSRKIQRGSLHYPAPKTDRSISVPHKIASTEFRRQSYAASRATSDPVSPTGRHEEERKDSAATNVVTHLQNLVPQKVSVLNPLTKFFKTQSQHSSTVLKTNAVIEMDRLETTTPSNEAKSHDQACDDHKLELSPALPSESVENALEDDMMIVEEESTAFDNKSHSNSLDSVSLPEIGEQSLPPPPPPLLSSHSQSMTPPPADVSIARLVRKSSLEEIDQIALPRDEERDNQPKKRTKFIKAWQSFKSKALIVIRWLSNYLGTIELDPRNELKEIPNHSSFFILAPDNRLRVRCYTTIKSKPFQTITLTVICLSSILLALETPLPVAEIGGIFCGIQKTIIFFDVFCLLWFLLEFCLKIIALGVFLHKGSYFHCLFNIIDFFVIVTTIVPIGTYYYHSSIDVCLSYSQTLLIYYPNLPAIIDVILILRALRPLRILRVEGLYLVTKGLVASLRKMGYVFFVGVILLLIFAVIGIQVFKGRFFYCSDLVKVKEEDCRGQYFTYPSNDLNCPRVLNRTWNRRALHFDHIGWSFLSLYTMITKEGWQDIMYHAIDSTGINTGPMYHANRANLIYFVLFMVLVTFFLINLLVGFVIVTFQQVGMKSHKEAHFDRNQENCLYLALTYSPKKRYIPRFTCHKRLYRIAKLKFFSIIIYVAVLINIIILALQYDNMDPKLTQSIYWANVGFTVFFTLEALFKLIILTPAHYFRSSERSFEFLVVIGSILEFILQTSLDLGNVMWRYSHIVSSLRVLRLIQITRNTKLIVWTVLRSLEIFPWVGVFLILLLYTYAVIGMQVFGQIEPVPIDYSATSAIHQYNNFENFPQALLVMIRCFTGENWERIMLTSANATCADNLNRTYCGSEFSYFFYPSFFFLSSILVLNLFVAIVIDNFDYLIRDRAILGSHDLTHFSPMWAKVDPAATGKIHHTDMFKMMHSINPPLGWGRLCPQVTCYGRMMALGIPLDEDGMITFNATLFAIVRQALNIETPGSACRNAELRQKLLNLFPNCQKIIDSVLPERPEWTTSEEYAALFIQQYWRKWRLKIERQAEAKPYLHHRISTVLRRVPDMGPALLQRRLTSMNVGISQVLPSPSASPNLLVATRTSSYSPSHSPSRSPVQLSPSRSPLINVRPVALDPRRASHLVRLETIQETDPRHSPLLESVPLVIPPSQSPAAFAKRQSTLSESDVLFVTPTGSLNASLNSEEEEEEQLHLETTV

>m.43115(Compagen)|Haliclona_tubifera_Cav1/2

RSQTSTYLLTSSFTLYPSSCLVMNLNPNSLQSPRQSFDLLEEGFPDGRTLSIVSEGERKSSFANALPTSPRSASQLHFHHHNDEVNLIDWISHTALVTRVRRLINHHSKPDYILCGISKKNKWRKRVKTLVNSFGFFAFISFIVIVHFIALTLYLPYINDDYEDRNELLAYINVIFILFYIVEAGLRIFAAGFILHRSSYMRSWDNALDVFVILIGIAFIGTRIVWASSERPLPGRFEFLSSLMALRLLRILSYFKSVQFILRAIGRSVKPLVHLGWFIFCVMMFFALIGIELDTQGNLHKGCFVDYTDKFDSVKTVRVSDYPCSDYGSHKCDQVGNRSYCEEWSRGPYNGMVGFDNLPVAMLTVFQCMTLEGWTSVLYLYESSVPAAIAWIYFIALIIFGSLLILNLFLGVLTSVFIGVSDRQEELGNTRALKEKRKLNEELYGYREWIRTAIEDEESSGPGGSTTTPGLTKSHSETKRTSSHSVEVLDADSENEVEIEENLAGSDERHWILSAIRQARKKVWKLVKSHAFFWFMIVIIQLNLIILLFSHYKAPIWLDSTIYNLNYFLTGIYILETLLKVFAMGFHNYFASNFNKLSFVLCCVNVLDIVVFMHIQEIRIPISQIVNAIRALRLITIFKYTRYWHDMKAVVNAFANIWSTMLSLLGLLGVYLVLVALFGMMMFRTRLTIPNLKHPNFNTFINAILLAFQLATTEDWHVVMYSTANSYINNPIPNINLNWLVGLFYVSSIMIGGLLLANIFLAIAVDKLLEVNEVEKDITKRVEQMEEQRREREKQLTALKNPTEQTGLGRTLKKIVMLYTLSVSHEEEIKRRKRNQRRRSGSEIKHDRPNSTSAMSDVERRRSSFGRAISDPMQYSGGSDPEGSPRGEREGVDGRDHPRSSVTSMSDRLFPRRVSLNNPLRKAEESIELQGYRLRDHSEGRRGSKRFNKTSSFVSISDDFEEFDFFGKRRMSQVTNSDQSTYSESCYEGSEGTAEYSKTPPLSALKNRQFSIDSDSSSVTVSTDSSDPSLRHIRSSPVIKTLQVPNPIQVPGNSPILKPGSTPMVINDEMGGVITEDATVSRLVRQSSLEDLDTSGDQLSDERQEVDIVTQWKDKRRKKVKQCWNSFTDKLTQVKQWAKSYFGSIELDPRNELKSIPAHSTFFIIAPDNRLRSWCFEVVRSKAFQSIAYTVTVISSLILAVESPIPVSSGLYCTLQDTVIYFDFAVLVWFLIEMVLKVISLGFIVHRGSYLRSLFNIIDFLIIIVTMIPLVVYFTNDNVSNPIQCYSYSQQIPRIYIWILVLRVSRPIRIIRITGLYLVCRGMISSVKRMGHMLLIGVLCLLIFAIIGVQCFKGRFFYCSDFVTEYEYDCGGDYFTYLTDDNNCPMVLPRKWKEYRLNFDNIFWSMLTLFTMLTKEGWQDVMYSAIDSTGQDRGPKYNTSQESLIYFILYMIFVTFFMLNFIVGFIIVTFQRVGIKTYTDSGLDKNQRNCLYIALTAHYKKGFVPRNHFHKFLYAIITSTVFDTVIFILVFMNTAILAIQHYQMGETLQEAIYWINLGFTVLFTMEAILKLVILTPPNYFRSFSRVFEFFVVIGSILELILQNTTDFLMENGFRYSHVISAFRVLRLTLVSKSTRLVFWTVIRSIEIFPWIGVLLVLVLYMFAVSGMQIFGQIDVVSIDSEPVTAIHQYNNFENFFQAMKVMIRTFTGENWEKIMLSCVNSDCDERIVKVTNGTTTTCGSPAAYAFFPTFFILSSILILNLFVAIIIDNFDYYVRDKSILGSHHLRDFPILWAKFDRNRSGTLRHGTIISMMKSLEPPLGWGKLCPQMTAYKHMMRLNIPLLEDDTVSFNSTLLAIVRHSLKINTSGNLTADNAELKNKLRSMFPSVPKKMLDSVIPERCDRTTGQLYGALVIQYHWRSTKISQQKLEQYKLQKRMSSMTIFPMGRELARRGTTLSIGAPSDSQAHPSSPPTTPTFASTSTRSFNLPPVMEETEEAATPTPSVRAHLLQRLETIEETDDQSVGMVNEDSISNTPSNKLYLHRLKSNATFSVTSESDMYATPTGSLTSLNDDDNLDDNGDSKSLKETSV

>XP_019398025.1|Crocodylus_porosus_Cav3.1

MVTFFIKLIGVHRWQISEKSWISLLFPVCALASLIPAGEKDGLSPPRSFPASAWKGLLVSLTERLCTWFERVSMLVILLNCVTLGMFHPCEDTACGSPRCRILQSFDDFIFAFFAVEMIVKMIALGIFGKKCYLGDTWNRLDFFIVIAGMLEYSLDLQNVSFSAVRTVRVLRPLRAINRVPSMRILVTLLLDTLPMLGNVLLLCFFVFFIFGIVGVQLWAGLLRNRCFLPENFSIPYTVELERYYQTENEDENPFICSQPRENGMRYCRNIPTRREEGLECTLDYYSYNDTTNTSCVNWNQYYTNCSAGEHNPFKGAINFDNIGYAWIAIFQVITLEGWVDIMYFVMDAHSFYNFIYFILLIIVGSFFMINLCLVVIATQFSETKQRESQLMKEQRVRYLSNASTLASFSEPGSCYDELLKYLVYVTRKASKQLVEAYRAAGLKMGLLSSPGNKNGADRQPCKHRQRKRSSVHHLIHHHHHHHHHYHMGNGNLRAPRASPEISDVETSSLHNSTNRLMLPPSTPNLHGASSNTESVHSIYHADCHFEPIRCRSSLPQPGLSLPSPEGLPKSMVGSKVYPTVHPSTSHEMLKEKSLGELAANSGAGTLTNLNIPPGPYSTMHKLLENQSTGACQSSCKITSQCGKLDSGSCNPDSCPYCIKTLANDLEPTDNETVDSDSEGVYEFTQDARYGDQRDPQRGEVGGKKMSRFLVFWNVVCETFRKIVDSKYFGRGIMIAILINTLSMGIEYHEQPEELTNALEISNIVFTSLFALEMLLKVLVYGPFGYIKNPYNIFDGIIVVISVWEIVGQQGGGLSVLRTFRLMRVLKLVRFMPALQRQLVVLMKTMDNVATFCMLLMLFIFIFSILGMHLFGCKFASERDGDTLPDRKNFDSLLWAIVTVFQILTQEDWNKVLYNGMASTSSWAALYFIALMTFGNYVLFNLLVAILVEGFQTEGEVSKSDSEGDVFPPSLEEEGGLKKHLSNPALMALSDHPELKKSLTPPLIIHTAATPMPMPKSAMFGDAAQGYESRRASSVSMDPSAHELKSPSSIRSSPHSPWSAASSWNSRRSSWNSIGRAPSLKRRGQSGERKSLLSGDGKESSEDGESSDEEQSSRAGSVNDSLPHRMGSLETKGSFDLQDTLQVPSLYRTSSMYSSRTSASEHQDCNGKTSAGALLHQFHLDDPRQDCDDCDDEGNMSKRDRAKAWIQARLPTWCKERDSWSIYIFAPHSKFRLMCNKIITHKMFDHVVLVIIFLNCITIAMERPKIEPHSAERIFLTLSNYIFTVIFLAEMTVKVVALGLCFGEKAYLKSSWNVLDGVLVLISVIDILVSMVSDSSTKILGMLRVLRLLRTLRPLRVISRAQGLKLVVETLMSSLKPIGNIVVICCAFFIIFGILGVQLFKGKFFVCQGEDTRNITNKSDCAEASYKWVRHKYNFDNLGQALMSLFVLASKDGWVDIMYDGLDAVGVDQQPVMNYNPWMLLYFISFLLIVAFFVLNMFVGVVVENFHKCRQHQEEEEAKRREEKRLRRLEKKRRNLMLDDVIMESSASAVQEAQCKPYYSDYSRFRLLIHQMCTSHYLDLFITGVIGLNVITMAMEHYQQPKVLDEALKICNYIFTVIFVLESVFKLIAFGFRRFFQDRWNQLDLAIVLLSIMGITLEEIEVNASLPINPTIIRIMRVLRIARVLKLLKMAVGMRALLDTVMQALPQVGNLGLLFMLLFFIFAALGVELFGDLECDDTHPCEGLGRHATFRNFGMAFLTLFRVSTGDNWNGIMKDTLRDCDQESTCYNTVISPIYFVSFVLTAQFVLVNVVIAVLMKHLEESNKEAKEEAELEAELEMEMKTIAPGQHPSSDLFAWTGGNGGDRPESPKGCTNPMQIKVDSQLSLFYPMERHLFDTLSLLIQESLEGELKLMDNLSGSVCHHYALPAPEYYNSEKQTNFHSKNDTLTLSPSKDLLSVRKPSVGRTHSLPNDSYMFQPPYSGPCADTPGERKPSYLKSQSGSKTSVQSQPADTSSLLQIPKVNFHCIRPHDNLDGEGRPKTSRPVHSPSAERLLRRQDSNITVQTDNPDLTPCFQNTVLAWMPSIYLWTAFPFYILYLKHYKRGYIVLSVLSRFKTFLGVLLWCVCWADLFYSFHELLQSRTPHPVHFVTPLILGITMLLAAILIQYERLRGVQSSGILIVFWFLSILCALGPFRSKIMTATTQGQVKDRFRFVTFYIYFVLIIIELILSCFKERPPFFSPVNTDPNPCPESNSGFLSRLTFWWFTSMAILGYKKPLEEKDLWSLNEEDTSKVVVGQLQKEWDKQQEECNQKEARAYMNKSSHVLNHVGDDPNEAEPWIDNKKQHKQPSFLKALLWAFGPYFLIGSFYKLIQDLLAFVNPQLLSVLIAFIKNKDAPSWWGFFIATLMFICAMLQTLILHQHFQYCFVTGMRLRTSITGLIYRKSLVITNSAKRTSTVGEIVNLMSVDAQRFMDLTTFLNLLWSAPVQIILAFYFLWQTLGPSVLAGVAVMILLIPFNAAIAIKTRAFQVEQMQHKDSRIKLMNEILSGIKVLKLYAWELSFNEKVLEIRKNELRILKKAAYLNALSTFAWVSAPFLVALTTFAVYVSVDENNVLDAQKAFVSLSLFNILRFPLNMLPQVISSIAQASVSLKRIQQFLCHDELDPNCVETKKITPGYAITVTNGTFSWAKELEPALKNVNLLVPSGSLIAVVGHVGCGKSSLVSAVLGEMEKLEGEVAVKGSVAYVPQQAWIQNATLKDNILFGQPSNEHKYQNVLEACALKTDLQVLPGGDQTEIGEKGINLSGGQRQRVSLARSVFSDADVYLLDDPLSAVDSHVAKHIFDKVIGPEGALKEKTRILVTHGISFLPQVDHIVVLIDGRVSETGSYQELLKQNGAFAEFLRNYAPDEDTEEDEPTMLEEEEVLLAEDTLSNHTDLTDNEPVTNEVRKQFLRQISVISSEVGECPSKMSTRRRVCEIKPVETLPTKKKDAKKLIEAETSETGTVKLTVFWQYMKAISPIACVIICFLYCCQNAAAIGANVWLSDWTNEPVINGTQHNTSMRLGVYAALGLLQGVLVLISSFTLAMGGISAAQKLHAALLENKFHTPQSFFDTTPTGRIINRFSKDIYVIDEVLPPTILMFLQTFFTSLQTMIVIVTSTPLFAVVIIPLAILYFFVQRFYVATSRQLKRLESVSRSPIYSHFSETVSGTSVIRAYGREKSFINISDIKVDENQKSYYPGIVSNRWLGIRVEFVGSCVVFFAALFAVLGKNSLNAGLVGLSVSYALQVTVALNWMVRMASDLESNIVAVERVKEYSETETEAPWIIEDRRPPEDWPAKGEVEFVNYSVRYRKGLDLVLTDLNLRVNGGEKIGIVGRTGAGKSSMTLCLFRILEAAKGDIKIDGVRISEIGLHDLRSKLTIIPQDPVLFSGTLRMNLDPFNSYSDEEIWTALELSHLKRFVNSQPAMLDYECSEGGENLSVGQRQLVCLARALLRKTRILVLDEATAAIDLETDDLIQMTIRTQFEDCTVLTIAHRLNTIMDYTRVLVLDKGTIAEFDTPTRLIASRSIFYSMAKDAGLA

>XP_015150965.1|Gallus_gallus_Cav3.1

MDEDGPRAADEEPEPGRAKTFIRLNDLSGAGGRPGPGEREVGSGDSEAEALPYPALAPVVFFYLSQESRPRSWCLRLVCNPWFERVSMLVILLNCVTLGMFHPCEDIACDSPRCRILQSFDDFIFAFFAVEMIVKMIALGIFGKKCYLGDTWNRLDFFIVIAGMLEYSLDLQNVSFSAVRTVRVLRPLRAINRVPSMRILVTLLLDTLPMLGNVLLLCFFVFFIFGIVGVQLWAGLLRNRCFLPENFSIPYTVDLERYYQTENEDENPFICSQPRENGMRYCRSIPTRREEGLECTLDYYSYNDTTNTSCVNWNQYYTNCSAGEHNPFKGAINFDNIGYAWIAIFQVITLEGWVDIMYFVMDAHSFYNFIYFILLIIVGSFFMINLCLVVIATQFSETKQRESQLMKEQRVRYLSNASTLASFSEPGSCYDELLKYLVYIARKGSKQLVELYRVAGVRMGFLASPASKVERHAGKRRSRRRSSVHHLIHHHHHHHHHYHLGNGNLRAPRASPEISDVDTGSLHNGTNRLMLPPSAPNPHMAPTATASGTESVHSIYHADCHVEPLRCRVALPQPSPEDIPRSVVMGSKVYPTVHPSTSHEVRKEKSLGEEAMGAGSSTLSGLNIPPGPYSTMHKLLETQSTGFLSVHVTDKGDGFRGPCPSSCKIASPCTKLDGGSRSPESCPYCLKALANEAEQTDNETDSDSEGVYEFTQDAHYSDQRDPQRGRAWARSSSRVLAFWRVVCETFQKIVDSKYFGRGIMVAILINTLSMGIEYHEQPEELTNALEISNIVFTSLFALEMLLKVLVYGPFGYIKNPYNIFDGIIVVISVWEIVGQQGGGLSVLRTFRLMRVLKLVRFMPALQRQLVVLMKTMDNVATFCMLLMLFIFIFSILGMHLFGCKFASERDGDTLPDRKNFDSLLWAIVTVFQILTQEDWNKVLYNGMASTSSWAALYFIALMTFGNYVLFNLLVAILVEGFQTEEISKREDAGGQLSCIQLPGDSAAGDASKSDSEGELFLRSLEEEGGLKKNLSNPVLMALSEHPELKKSLTPPLIIHTAATPMPMPKSAMFGDAAQGYESRRASGVSMDPAAYELKSPPSTRSSPHSPWSAGSSWPSRRSSWNSIGRAPSLKRRGQSSERRSLLSGEGKESSEDGESSDEERSSRAGSVNGSLPHRMESLEAKGSFDLQDTLQVPSLYRTSSMHSSRTSTSEHQDCNGKTSPGLLVHQLHLDDPRPDCDDADDEGNMSKRDRMKAWVRAHLPTCCKERDSWSIYVFAPHSRFRLMCNKIITHKMFDHIVLVIIFLNCITIAMERPKIEPHSAERIFLTLSNYIFTVIFLTEMTVKVVALGLCFGEKAYLKSSWNVLDGVLVLISVIDILVSMVSDSGTKILGMLRVLRLLRTLRPLRVISRAQGLKLVVETLMSSLKPIGNIVVICCAFFIIFGILGVQLFKGKFFICQGEDTRNITNKSDCAEASYKWVRHKYNFDNLGQALMSLFVLASKDGWVDIMYNGLDAVGVDQQPVMNYNPWMLLYFISFLLIVAFFVLNMFVGVVVENFHKCRQHQEEEEAKRREEKRLRRLEKKRRNLMLDDVLMESTASVVPEAQCKPYYSDYSRFRFLIHQMCTSHYLDLFITGVIGLNVITMAMEHYQQPKVLDEALKICNYIFTVIFVLESVSKLIAFGFRRFFQDRWNQLDLAIVLLSIMGITLEEIEVNASLPINPTIIRIMRVLRIARVLKLLKMAVGMRALLDTVMQALPQVGNLGLLFMLLFFIFAALGVELFGDLECDDTHPCEGLGRHATFRNFGMAFLTLFRVSTGDNWNGIMKDTLRDCDQESTCYNTVISPIYFVSFVLTAQFVLVNVVIAVLMKHLEESNKEAKEEAELEAELEMEMKTISPGQHNPSDIFTWMGGVGGERPESPRGCPHPLQIQVDSQLSLVYPMERHLFDTISLLIQESLEGELKLMDNLSGSVCHHYALPAPEYYSSENQIPLAEMEALSLTSDILSEKSWSLALTDDSFPDDTNTHLLNALESNAELRCRDAALALSPCQDLLSVRKPSVGRTHSLPNDSYMFHTSQPRSRPSTASLRDGNPAQHKVHSGSKASVQSQPADTSSLLQIPKDHFHHMRAHDPLVRESKAPAAAPVHSPSAERLLRRQMAIRNDSSDSKENLHAELSELPNPDIPSAPQEESAAALTPSEGEDLAAWSRASVHTQHHSHNQYNVSKQAPASCACADSYQETPGDSMDQEVSETNSSLEPFPSETCTASSACSEAQPLTPKRNIGSTGSVTLKDLKKYHSVDTQGLLKKPPSWLDDQRRHSIEICSMENSPQHHSTSSSSGFISQVVSEMEGLQGTRQKKKLSPPCISIDPPSEQGLLPRGPHSVSPARGDTCLRRRAPSCDSKDSVDIGDSLLPDSMSASPTPKKDLLTLPSFSFDQNEVEP

>XP_020670763.1|Pogona_vitticeps_Cav3.1

MLVILLNCVTLGMFHPCEDMGCDSPRCKILQSFDDFIFAFFAVEMVVKMIALGIFGKKCYLGDTWNRLDFFIVLAGMLEYSLDLQNVSFSAVRTVRVLRPLRAINRVPSMRILVTLLLDTLPMLGNVLLLCFFVFFIFGIVGVQLWAGLLRNRCFLPENFSFPYTVDLEPYYQTENEDESPFICSQPRENGMRYCRSIPKRREEGLECTLDYYSYNDTSNTSCVNWNQYYTNCSAGEHNPFKGAINFDNIGYAWIAIFQVITLEGWVDIMYFVMDAHSFYNFIYFILLIIVGSFFMINLCLVVIATQFSETKQRESQLMKEQRIRYLSNASTLASFSEPGSCYDELLKYLVHVLRKATKQLVEVYQMAGVKMGLLSSPGSKSSVERQSCKHLPRKRSSVHHLIHHHHHHHHHYHLGNGSLQAPRASPEISDVDTSSLHNGANRLMLPPTANPHAASSNTESVHSIYHADCHFEPIHCRSSLSQPGLSLPGPEVLPRTLVGTKVYPTVRPGSSHELLKEKGLGELTLNPGTSTLMNLNIPPGPYSPMQKILETQSTGPCQSSCKISSQGGKSDSPSNPDSCPYCIKTLANKPEQTDNETGDSDSDGVYEFTQDAHYADQRDPKRWKAQGRKALKLLAFWRVICETFRKIVDSKYFGRGIMIAILINTLSMGIEYHEQPEELTNALEISNIVFTSLFALEMLLKLLVYGPFGYIKNPYNIFDGIIVVISVWEIVGQQGGGLSVLRTFRLMRVLKLVRFMPALQRQLVVLMKTMDNVATFCMLLMLFIFIFSILGMHLFGCKFASERDGDTLPDRKNFDSLLWAIVTVFQILTQEDWNKVLYNGMASTSSWAALYFIALMTFGNYVLFNLLVAILVEGFQTEEITKREAASRQLSCIQLPVDSSVGDATKSDSEGELFPHNLEEDCGFKNSFSNPALMALCDHPELKKSLTPPVIIHTAATPMSMPKSANCGEAAQGYDSRRASSVSMDPHAYDLKSPMSARSSPHSPWSAGSSWNSRRSSWNSIGRAPSLKRRHQSGERKSLLSGDGKQSSEEGESSDEEQSSRTGSINGSIPHRMESLETKGSFDFQDTLQVPSLYRTGSIHSSRSSASERQDCNGKTTSRVLVQQLHLEDPVPECDDMDDEGNLSKKERMKAWIRARLPACCKERDSWSIYIFAPQSKFRMICSKIISHKMFDHIVLVIIFLNCITIAMERPKIDPHSAERIFLTLSNYIFTAIFLAEMTIKVVALGLCFGEKAYLRSSWNMLDGVLVLISVADILVSMVSDSGTKILGMLRVLRLLRTLRPLRVISRAQGLKLVVETLMSSLKPIGNIVVICCAFFIIFGILGVQLFKGKFFVCQGEDTRNITNKSDCAEASYKWVRHKYNFDNLGQALMSLFVLASKDGWVDIMYDGLDAVGVDQQPSMNYNPWMLLYFISFLLIVAFFVLNMFVGVVVENFHKCRQHQEEEEAKRREEKRLKRLEKKRRKAQCKPYYSDYSRFRLLIHQMCTTHYLDLFITGVIGLNVITMAMEHYQQPKVLDEALKICNYIFTIIFVMESVFKLVAFGFRRFFQDRWNQLDLAIVLLSIMGITLEEIEVNASLPINPTIIRIMRVLRIARVLKLLKMAVGMRALLDTVMQALPQVGNLGLLFMLLFFIFAALGVELFGDLECDDTHPCEGLGRHATFRNFGMAFLTLFRVSTGDNWNGIMKDTLRDCDEESTCYNTVISPIYFVSFVLTAQFVLVNVVIAVLMKHLEESNKEAKEEAELELELEMEMKTITPGQHPLSDPFVWAGGNGGERPDSPGGYTNPMQIKMDSQLSLVHPMTNIPSRNDSLIFSPSQDHLSVRKTSVGRTHSLPNDSYMFQAPNTGVYDGSSGEKNSPHQKSQSGSTTSVQSQPANINALLQIPRDPFHHLKPQYNMDWENKSKVLPAVHSPSAERLLRRQMAIRNDSLDTPSSESRENIRVEVSELSDPNLSAPFSEDLMPSEANEAASWRRSSVRTQQHSCNQYNISKHVPDLCACTDSLQELPGDPMDQEVSEINSSFTSGTCSAMSASSVVQPETPKRNLGNSGNVTLKDLKKCYSVDTQGLLKKPSSWLDDQRRHSIEICSMENSPQHHSTTSSSSGFLSQVLSEMEGLPGTRPKKKLSPPCISIEPPDEQRLFLRGSRTASPAPTADVCLRRRAPSWDSKDSMDAGDSALPDSMSVSPAPKKDLLMPPSFSFDQTDTDP

>XP_015266211.1|Gekko_japonicus_Cav3.1

MDEDGPRAAEEQDPEEDHSKTFVRLNDLSGDGGRSGPGDKEKDPGSGDSESEGLPYPALAPTVFFYLSQHSRPRSWCLRMVCNPWFERASMLVILLNCVTLGMFHPCEDTDCDSPRCKILQSFDDFIFAFFAVEMIIKMIALGIFGKKCYLGDTWNRLDFFIVIAGMLEYSLDLQNVSFSAVRTVRVLRPLRAINRVPSMRILVTLLLDTLPMLGNVLLLCFFVFFIFGIVGVQLWAGLLRNRCFLEENFSFPYTVELEPYYQTENEDESPFICSQPRENGMRYCRSIPTRREEGLECTLNYYSYNDTTNTSCVNWNQYYTNCSAGEHNPFKGAINFDNIGYAWIAIFQVITLEGWVDIMYFVMDAHSFYNFIYFILLIIVGSFFMINLCLVVIATQFSETKQRESQLMKEQRIRYLSNASTLASFSEPGSCYDELLKYLIYILRKASKQLVEAYRTTGVKMGFLSSPGSKTSAERQPCKRRPQKRSSVHHLIHHHHHHHHHYHLGNGSLQAPRASPEISDVETSSLHNGANRLMLPPTANPHAASSNTESVHSIYHADCHFEPLRYRSSLPQPGLSLTGPEVFPRTVVGSKVYPTVHPSSSHELLKDKCVGEPAINPGANTLMNLNIPPGPYSTMQKLLETQSTGPCQSSCKISSQCGKPDSTCNPDSCPYCVKTLANEPELTDNEMGESDSDGVYEFTQDAHYNDQRDPQRWKAPGRKTLKCLAFWNVVCETFRKIVDSKYFGRGIMIAILINTLSMGIEYHEQPEELTNALEISNIVFTSLFALEMLLKLLVYGPFGYIKSPYNIFDGIIVVISVWEIVGQQGGGLSVLRTFRLMRVLKLVRFMPALQRQLVVLMKTMDNVATFCMLLMLFIFIFSILGMHLFGCKFASERDGDTLPDRKNFDSLLWAIVTVFQILTQEDWNKVLYNGMASTSSWAALYFIALMTFGNYVLFNLLVAILVEGFQTEEITKREAASRQLSCIQLPVDSSAGDASKSDSDGELFPRSLEEECGLKKNFSNPAMMALCDHPELKKSLTPPLIIHTAATPMPMPKSAMYGEAAQGYDSRRASSISMDPNAYDLKSPMSTRSSPHSHWSAGSSWNSRRSSWNSIGRAPSLKRRNQSGERKSLLSGDGKQSSEEGGSSDEERSSQTGSINGSIPHRMQSLETKGSFDFQDSLQVPSLYRTSSMHSSQTSASEHQDCNGKTSARVLIQQLHLGDLPPECDDLDDEDNLSKKDRMKAWIRAHLPTWCKERDSWSVYIFAPQSKFRLVCNKIITHKMFDHVVLVIIFLNCITIAMERPKIEPHSAERIFLTLSNYIFTAIFLAEMTIKVVALGLCFGEKAYLRSSWNMLDGVLVLISIIDILVSMVSDSGTKILGMLRVLRLLRTLRPLRVISRAQGLKLVVETLMSSLKPIGNIVVICCAFFIIFGILGVQLFKGKFFVCQGEDTRNITNKSDCAEASYKWVRHKYNFDNLGQALMSLFVLASKDGWVDIMYDGLDAVGVDQQPIMNYNPLFVGVVVENFHKCRQHQEEEEAKRREEKRLRRLEKKRRNLMLDDVIMESSASAVPEAQCKPYYSDYSRFRLLIHQMCTTHYLDLFITGVIGLNVITMAMEHYQQPKVLDEALKICNYIFTVIFVMESVFKLIAFGFRRFFQDRWNQLDLAIVLLSIMGITLEEIEVNASLPINPTIIRIMRVLRIARVLKLLKMAVGMRALLDTVMQALPQVGNLGLLFMLLFFIFAALGVELFGDLECDATHPCEGLGRHATFRNFGMAFLTLFRVSTGDNWNGIMKDTLRDCDEESTCYNTVISPIYFVSFVLTAQFVLVNVVIAVLMKHLEESNKEAKEEAELELELEMEMKTITSGQHPSSDAFMWTGINTGERPDSPGGYANPMQIKVDSQLSLVHPMERHLFDTISLLIQESLEGELKLMDNLSGSVCHHYALPTTENYSSEKQIPLAEMEALSLTSDILSENSWSLALTDDSFPDDNNTFLLNAPENNTDLPSGNDPLTLSPSKDLLSVRKPTVGRTHSLPNDSYMFQSPDTGTYEVSSTEKNPSHQKSQSGSTTSVQSQPANIDALLQIPRDFFHPLKLHSNLDWRSKSKAPTSVHSPSAERLLRRQMAIRNDSSETHSSESKENIHAEVSELSDPNLSAPSTEDLSVILRPSEAQDSASWGRSSVHTQQHSLNQYNICKHVPDSCACTDSLQELPGDPMDQEVSEINSSFTSGTCSAMSASSVVQPETPKRNVGNSGNVTLKDLKKCYSVDTQGLLKKPSSWLDDQRRHSIEICSMENSPQHHSTSSSSGFISQVLSEMEGLQGTRQKKKPSPPCISIDPPNEQSLLRRGSHTASSSSAEVCLERPAPSWDSKDSMDAGDLALPHSMSVSPAPKKDLLTLPSFSFDQPDADP

>NP_061496.2|Homo_sapiens_Cav3.1

MDEEEDGAGAEESGQPRSFMRLNDLSGAGGRPGPGSAEKDPGSADSEAEGLPYPALAPVVFFYLSQDSRPRSWCLRTVCNPWFERISMLVILLNCVTLGMFRPCEDIACDSQRCRILQAFDDFIFAFFAVEMVVKMVALGIFGKKCYLGDTWNRLDFFIVIAGMLEYSLDLQNVSFSAVRTVRVLRPLRAINRVPSMRILVTLLLDTLPMLGNVLLLCFFVFFIFGIVGVQLWAGLLRNRCFLPENFSLPLSVDLERYYQTENEDESPFICSQPRENGMRSCRSVPTLRGDGGGGPPCGLDYEAYNSSSNTTCVNWNQYYTNCSAGEHNPFKGAINFDNIGYAWIAIFQVITLEGWVDIMYFVMDAHSFYNFIYFILLIIVGSFFMINLCLVVIATQFSETKQRESQLMREQRVRFLSNASTLASFSEPGSCYEELLKYLVYILRKAARRLAQVSRAAGVRVGLLSSPAPLGGQETQPSSSCSRSHRRLSVHHLVHHHHHHHHHYHLGNGTLRAPRASPEIQDRDANGSRRLMLPPPSTPALSGAPPGGAESVHSFYHADCHLEPVRCQAPPPRSPSEASGRTVGSGKVYPTVHTSPPPETLKEKALVEVAASSGPPTLTSLNIPPGPYSSMHKLLETQSTGACQSSCKISSPCLKADSGACGPDSCPYCARAGAGEVELADREMPDSDSEAVYEFTQDAQHSDLRDPHSRRQRSLGPDAEPSSVLAFWRLICDTFRKIVDSKYFGRGIMIAILVNTLSMGIEYHEQPEELTNALEISNIVFTSLFALEMLLKLLVYGPFGYIKNPYNIFDGVIVVISVWEIVGQQGGGLSVLRTFRLMRVLKLVRFLPALQRQLVVLMKTMDNVATFCMLLMLFIFIFSILGMHLFGCKFASERDGDTLPDRKNFDSLLWAIVTVFQILTQEDWNKVLYNGMASTSSWAALYFIALMTFGNYVLFNLLVAILVEGFQAEEISKREDASGQLSCIQLPVDSQGGDANKSESEPDFFSPSLDGDGDRKKCLALVSLGEHPELRKSLLPPLIIHTAATPMSLPKSTSTGLGEALGPASRRTSSSGSAEPGAAHEMKSPPSARSSPHSPWSAASSWTSRRSSRNSLGRAPSLKRRSPSGERRSLLSGEGQESQDEEESSEEERASPAGSDHRHRGSLEREAKSSFDLPDTLQVPGLHRTASGRGSASEHQDCNGKSASGRLARALRPDDPPLDGDDADDEGNLSKGERVRAWIRARLPACCLERDSWSAYIFPPQSRFRLLCHRIITHKMFDHVVLVIIFLNCITIAMERPKIDPHSAERIFLTLSNYIFTAVFLAEMTVKVVALGWCFGEQAYLRSSWNVLDGLLVLISVIDILVSMVSDSGTKILGMLRVLRLLRTLRPLRVISRAQGLKLVVETLMSSLKPIGNIVVICCAFFIIFGILGVQLFKGKFFVCQGEDTRNITNKSDCAEASYRWVRHKYNFDNLGQALMSLFVLASKDGWVDIMYDGLDAVGVDQQPIMNHNPWMLLYFISFLLIVAFFVLNMFVGVVVENFHKCRQHQEEEEARRREEKRLRRLEKKRRNLMLDDVIASGSSASAASEAQCKPYYSDYSRFRLLVHHLCTSHYLDLFITGVIGLNVVTMAMEHYQQPQILDEALKICNYIFTVIFVLESVFKLVAFGFRRFFQDRWNQLDLAIVLLSIMGITLEEIEVNASLPINPTIIRIMRVLRIARVLKLLKMAVGMRALLDTVMQALPQVGNLGLLFMLLFFIFAALGVELFGDLECDETHPCEGLGRHATFRNFGMAFLTLFRVSTGDNWNGIMKDTLRDCDQESTCYNTVISPIYFVSFVLTAQFVLVNVVIAVLMKHLEESNKEAKEEAELEAELELEMKTLSPQPHSPLGSPFLWPGVEGPDSPDSPKPGALHPAAHARSASHFSLEHPTDRQLFDTISLLIQGSLEWELKLMDELAGPGGQPSAFPSAPSLGGSDPQIPLAEMEALSLTSEIVSEPSCSLALTDDSLPDDMHTLLLSALESNMQPHPTELPGPDLLTVRKSGVSRTHSLPNDSYMCRHGSTAEGPLGHRGWGLPKAQSGSVLSVHSQPADTSYILQLPKDAPHLLQPHSAPTWGTIPKLPPPGRSPLAQRPLRRQAAIRTDSLDVQGLGSREDLLAEVSGPSPPLARAYSFWGQSSTQAQQHSRSHSKISKHMTPPAPCPGPEPNWGKGPPETRSSLELDTELSWISGDLLPPGGQEEPPSPRDLKKCYSVEAQSCQRRPTSWLDEQRRHSIAVSCLDSGSQPHLGTDPSNLGGQPLGGPGSRPKKKLSPPSITIDPPESQGPRTPPSPGICLRRRAPSSDSKDPLASGPPDSMAASPSPKKDVLSLSGLSSDPADLD

>XP_021336020.1|Danio_rerio_Cav3.1

MMAEDGGIISEDNQLNEAEGSSSLPRTFIPLNDLSGGGVRSELESDGGERRESAAGGAAEETSASDGEALPYPSLAPVVFFYLKQTTRPRSWCLKMVCNPWFERASMLVILLNCVTLGMFHPCEDSHCDSERCKILEDFDDFIFAFFAVEMVIKMVALGIFGKKCYLGDTWNRLDFFIVLAGMLEYSLNLQNVSFSAVRTVRVLRPLRAINRVPSMRILVTLLLDTLPMLGNVLLLCFFVFFIFGIVGVQLWAGLLRNRCFLPENFSLPTSLELRKYYHTENDDENPFICSQPRENGMRLCTSVPTLHEEGRQCQLDMASYNSTDNTTCVNWNKYYTNCSAGEANPFKGAINFDNIGYAWIAIFQVITLEGWVDIMYFVMDAHSFYNFIYFILLIIVGSFFMINLCLVVIATQFSETKQRESQLMKEQRVRFRSNASTLNSYSEPGSCYDELLKYLVHVIRKGTRQIAHLIRAAGRRAGLRICASPPLEAPPTKRRRQKQRQGSIRHIMHHHHHHLHHHYHLGNGSVRSDGGREVDNPSQSGIVNKSGNVNLMLPVIVPQQDAFSGSFGHSSAESVHSVYQVSGHLEPLRCGPSPSPTVLHAYKRNSVPFAAPVHKNYPTLQSSLALEQLRQRILDPASVLTSLNIPPNPINTSQCLVETQGPPGKVISKDALMNCSSTNIDTALTYDPETCPYCAKSMANDSEGIDGNETADSDSEGVYETQVAHYRDSRGPKKKKKFALGARATKVIDFWRLVCGTFRKIVDSKYFGQGIMIAILINTLSMGIEYHEQPDELTNALEISNIVFTSLFVLEMLLKLLVYGPFGYIKNPYNIFDGIIVVISVWEIVGQQGGGLSVLRTFRLMRVLKLVRFMPALQRQLVVLMKTMDNVATFCMLLMLFIFIFSILGMHLFGCKFGSERDGDTLPDRKNFDSLLWAIVTVFQILTQEDWNKVLYNGMASTSPVAALYFIALMTFGNYVLFNLLVAILVEGFQTEEITKRDDLHGQLSCIQLPIDSGGDASKSDSEADIYARSLEDVSGHKKDMSASSVVPINGHVDLKSSLTPPLITHTAATPMPIPKISGGGDSALSQESRRGSSVSMEPNGQDKSPSSARSSPNAPLSSASSWNSRRSSWNSLGRAPSLKRQKHQSGERRSLLSGDGQSSSDEGDAGESMGGMSEGDDASLARTDSLGQRPRHRRMESLETRSSFDLHPDTLQVPYPHRSGSIHSARPPNFLSNGKTSPSAATTQLSLDEHHSEDDNADEEGNLSRRARLYRWLEHKQPEWCRQRVTWSLYLFPPESRFRVTCNKIITHKMFDHVVLVIIFLNCITIAMERPRIDPSSAERIFLTLSNYIFTAIFVTEMTIKVVALGFCFGEKTYLKSSWNILDGMLVMISVIDILVSLISNSGTKILGMLRVLRLLRTLRPLRVISRAPGLKLVVETLMSSLKPIGNIVVICCAFFIIFGILGVQLFKGKFFVCHGDDTRNITNKSDCLLANYKWVRHKYNFDNLGQALMSLFVLASKDGWVDIMYDGLDAVGVDQQPVMNYNPWMLLYFISFLLIVAFFVLNMFVGVVVENFHKCRRNQEEEEAKRREAKRQKRRDKKRRNIMLTGVSWSSPDHTQEAQSKPFYSDYCPTRRLIYNMCKSQYLDLFITIVIALNVITMSMEHYHQPKVLSDSLKICNYIFTVIFVLESIFKLVAFGFRRFFKDRWNQLDLAIVLLSIMGITLEEIEVNASLPINPTIIRIMRVLRITRVLKLLKMAVGMRALLDTVMQALPQVGNLGLLFMLLFFIFAALGVELFGDLICDELHPCEGLGRYATFKNFGMAFLLLFRVSTGDNWNGIMKDTLRDCSQDTGICYNTVVSPIYFVSFVLTAQFVLVNVVIAVLMKHLEESNKEAKEAEAEAELELEQEMEALGRDREMQRGHMPRLGSGDLGGPGGSPWISRDHRDLSGPLYPIDSPTADIRRDFEDHLRTDPDPPPNLEPPLERRPMFDSVSLVIQGSLEGELSLMDNLSGSICHYYALPPLPNKHCTEKQIPLAEMEALSLASEKSWSLALTDDSVPDDFNPLLLTSQECNTTDPSDPPDTRQADEPTDEHLLSVKKTSVGRTHSLPNDSYMFLPLQSNSNHSTHTTSPHLAQTGSTASVQSQTEEASQHLTVPSDLFRPISPHSLSDSESIPRIPPPRRAHTLSRTLRRQVAVSADSQETLYTEGGEAEGPLGLRELSDPLPPQQQQHQPSLFLVPATPGASPKPSRPSVHTQHNPYDQYNVFSRSSHSPPVPPPPPDYKKQEDVDSVDQEVSRIIRAGLTGRSDDVNREGTGDQCAKRQLKKYHSVDTQGQRALLLPRPLSWLDDPRRHSIEVCSSVESSPQRSSISSGFVSRADSLQQQSSPRGRKKKMSPPCISVDPPEDPELPGGFHPALGIVPPPLPSRDTCLRRRAPSSESKDSFDLGGGEALPQEGVPNSKLLTLPSFSFEKTSSEH

>XP_019344978.1|Alligator_mississippiensis_Cav3.3

MMVILLNCVTLGMYQPCEDMDCLSDRCKILQVFDDFIFIFFAMEMVLKMVALGIFGKKCYLGDTWNRLDFFIVMAGMVEYSLDLQNINLSAIRTVRVLRPLKAINRVPSMRILVNLLLDTLPMLGNVLLLCFFVFFIFGIIGVQLWAGLLRNRCFMEENFTIQGDIVLPPYYQPEEDDEMPFICSLSGDNGIMGCHEIPPLKERGHECCLDKDDYYYYNSVRQEFNVSGMCVNWNQYYNVCRTGNTNPHKGAINFDNIGYAWIVIFQVITLEGWVEIMYYVMDAHSFYNFIYFILLIIVGSFFMINLCLVVIATQFSETKQREHQLMQEQRARYLSSSTVASYMEPGDCYEEIFQYICHIVRKAKRRTLSLYNNLQNRRQGRMEPGNMKPDMNGKGQRCYRHCQKHNSLDYTPQALVQPIAVTLTSDPNNCPRCHRVQREAARRLSVLDSADSDQENGGAGEEEGEGSRESNGVAELEKEEEEGEEGKQMRLCSDMWQEVRVKLRGIVESKYFNRGIMIAILVNTISMGIEHHEQPEELTNILEISNVVFTSMFALEMILKLAAFGLFDYLRNPYNIFDSIIVIISIWEIIGQSDGGLSVLRTFRLLRVLKLVRFMPALRRQLVVLMKTMDNVATFCMLLMLFIFIFSILGMHIFGCKFSLRTDTGDTVPDRKNFDSLLWAIVTVFQILTQEDWNVVLYNGMASTSPWASLYFVALMTFGNYVLFNLLVAILVEGFQAEGDANRSYSDEDQSSSNAEELDRFQDVQDGSDPKLCAIPVTPNGHLDPSPPSSNQPVVSAAVTNSSRNSLQPDQVLIAVGSRKSSVMSLGRMTYDQRSLSSSRSSHYGAWGRSGAWGSRRSSWNSLARGHSLKHKPPSAEHESLLSGERHRTADGERAGMGRTPHHRRTLSLDTKGSCDLVELASVPPSHRATRKVGVTSGTTEHQDCNGKMPSIAKEIFPKMNNRKERGEEEEEIDYTLCFRIRKMIEVYKPDWCELREDWSIYLFSPQNRFRILCQTIIAHKLFDYIVLAFIFLNCITIALERPQIEHRSTERIFLSVSNYIFTAIFVAEMTLKVVSLGLYFGDQAYLRSSWNILDGFLVFVSLIDIVVSVASAGGAKILGVLRVLRLLRTLRPLRVISRAPGLKLVVETLISSLKPIGNIVLICCAFFIIFGILGVQLFKGKFYHCLGVDIRNITNRSDCVAANYKWVHHKYNFDNLGQALMSLFVLASKDGWVNIMYNGLDAVAVDQQPVTNNNPWMLLYFISFLLIVSFFVLNMFVGVVVENFHKCRQHQEAEEARRREEKRLRRLEKKRRKAQRLPYYASYCPVRLLIHSVCTSHYLDIFITFIICLNVVTMSLEHYNQPVSLETALKYCNYMFTTVFVLEAVLKLVAFGLRRFFKDRWNQLDLAIVLLSVMGITLEEIEINAALPINPTIIRIMRVLRIARVLKLLKMATGMRALLDTVVQALPQVGNLGLLFMLLFFIYAALGVELFGKLVCNDENPCEGMSRHATFENFGMAFLTLFQVSTGDNWNGIMKDTLRDCTQDDRSCLSNLQFISPLYFVSFVLTAQFVLINVVVAVLMKHLDDSNKEAQEDAEMDAEIELEMAHGLCPRSLGSPSSGQGGSGGGGGGGGGGRVDPEGRLCMRCYSPAQVQLAEMEALSLNSDKSSSIFLGDDMDNRSIRQLSPKETKNGQDSPDSVEGGGLGECFLPLSNTDASLNPESYLCEMEKIPFNPVQSWLKHESHQVPPSPFSPGTSSPLLPVPAEFFHPALSATQAGQEKVLCPGSLPKISLQGSWASLRSPSVNCSLLHQATESDTSLDDSRSNSSEGSLQTTLEDSLNLSDSPQCTLDLPVALCPMPAAATQRTMVAPLSPASRRRSLRTRSIFSLRNIRRHQRSHSSGGSTSPGCTHHDSMDPSDEEGVGSSLRGGGNNSEQSETLSSLSLTSLFSPPPPPPGLALVKKCNSTSSLNTSRGPATNEARQFYSVDPRGFLSMPSWVTDFCKDPPSAGEGEGPDGPSKLGTAEHSSPLPSELELGNGVSKRNR

>XP_015144216.1|Gallus_gallus_Cav3.3

MAECTPQPPSTWPASPQPHNTSAACPLEPMNMGVSEQQMPLEQPPSSPELGEDEEVTSPPDPDVPYPDLAPVVFFCLKQTTSPRSWCIKMVCNPWFECVSMMVILLNCVTLGMYQPCEDMDCLSDRCKILQVFDDFIFIFFAMEMVLKMVALGIFGKKCYLGDTWNRLDFFIVMAGMVEYSLDLQNINLSAIRTVRVLRPLKAINRVPSMRILVNLLLDTLPMLGNVLLLCFFVFFIFGIIGVQLWAGLLRNRCFMEENFTIQGDIVLPPYYQPEEDDEMPFICSLSGDNGIMGCHEIPPLKERGHECCLDKDDYYYYNSVRQEFNVSGMCVNWNQYYNVCRTGNANPHKGAINFDNIGYAWIVIFQVITLEGWVEIMYYVMDAHSFYNFIYFILLIIVGSFFMINLCLVVIATQFSETKQREHQLMQEQRARYLSSSTVASYMEPGDCYEEIFQYVCHIMRKAKRRTLGLYNSIQSRRQGHVEPSEAKRGASRKKQRHYRICQQHNPLDCPPQGLVQPIAVTATSDLTNCPRCHRGEHDTSRRLSVLDSADSDQEDGGDSEAEGEGCRGDHSASALEKEEEEVEEGGRMKLCSDMWREVRVKLRGIVDSKYFNRGIMIAILVNTISMGIEHHEQPEELTNILEISNVVFTSMFALEMILKLAAFGLFDYLRNPYNIFDSIIVIISIWEIIGQSDGGLSVLRTFRLLRVLKLVRFMPALRRQLVVLMKTMDNVATFCMLLMLFIFIFSILGMHIFGCKFSLRTDTGDTVPDRKNFDSLLWAIVTVFQILTQEDWNVVLYNGMASTSSWAALYFVALMTFGNYVLFNLLVAILVEGFQAEGDANRSYSEEDQSSSNMEELDQFQEIQEGSDPKLCAISVTPNGHLDPSLPSSNAPVAAGGVSNSSRNSLQPDQIRVAVGSRKSSVMSLGRMTYDQRSLSSSRSSHYGAWGRSGAWGSRRSSWNSLARGHSLKHKPPSAEHESLLSGDRHRVGDREPDGAERVFHHRRTLSLDNKGSCDLLELASIPSGHRGTCRPGTTSGGSEHQDCNGRMPSIAKEIFPKMNNRKERGEDDEEIDYSLCFRIRKMMEVYKPDWCELREDWSIYLFSPQNRFRLLCQTIIAHKLFDYVVLAFIFLNCITIALERPQIEHRSTERIFLTVSNYIFTAIFVAEMTLKVVSLGLYFGDQAYLRSSWNVLDGFLVFVSLIDIVVSVASAGGAKILGVLRVLRLLRTLRPLRVISRAPGLKLVVETLISSLKPIGNIVLICCAFFIIFGILGVQLFKGKFYHCLGVDIRNITNRSDCVAANYKWVHHKYNFDNLGQALMSLFVLASKDGWVNIMYNGLDAVAVDQQPVTNNNPWMLLYFISFLLIVSFFVLNMFVGVVVENFHKCRQHQEAEEARRREEKRLRRLEKKRRKAQRLPYYATYCPIRLLIHSVCTSHYLDIFITFIICLNVVTMSLEHYNQPVSLETALKYCNYLFTTVFVLEAVLKLVAFGLRRFFKDRWNQLDLAIVLLSIMGITLEEIEINAALPINPTIIRIMRVLRIARVLKLLKMATGMRALLDTVVQALPQVGNLGLLFMLLFFIYAALGVELFGKLVCNDENPCEGMSRHATFENFGMAFLTLFQVSTGDNWNGIMKDTLRDCSHDDRSCLSNLQFISPLYFVSFVLTAQFVLINVVVAVLMKHLDDSNKEAQEDAEMDAEIELEFAHGLCSRKSGSSKSRQGKGAGSGSGGGRTEPEGRLCTRCYSPAQENLWLDSVSLIIKDSFDGELMIIDNLSGSVFHHYSSPAMCEKCNHDKQEVQLAEMEALSLNSDKSSSILLGDESDNRSTSQLSPKETKDGQDSPDSTEAGDVGECLLPISSTDGSTNPDNYLCEIEKTPFNSVQSWLKHEGTKVPPSPFSPGTSCPLLPVPAEFFHPAISATQTGPEKVSCAHNLPKISLQGSWASLRSPSVNCSLLHQAPESDTSLDSRSSSSAGSLQTTLEDSLNLSDSPQCALDLPVALCPMPAASDPRPLAAPLSPTSRRRSLRGRGLFTLRSIRRHQRSHSSGGSTSPGCTHHDSMDPSDEEGAGSSLRGGGNNSEPSETLSSLSLTSLFSPPPPGLTLGKKCSSTSSLHASPSPRRPAAHHTKPFYTVDPKGFLSMPSWVTDFCKDTPPPVQGAEPDGSSRLGTGECGSPLPSELELGDGVSKRNR

>XP_015274652.1|Gekko_japonicus_Cav3.3

MAEQGSHPSSAWHSGNPSAHNPTLCPLERVDLGITEQLTPLEQPPPSPDLGDDDEDDDVEPLPDPDVPYPDLAPVVFFCLKQTTSPRNWCIKMVCNPWFECVSMMVILLNCVTLGMYQPCDDMDCLSNRCKILQVFDDFIFIFFAMEMVLKMVALGIFGKKCYLGDTWNRLDFFIVMAGMVEYSLDLQNINLSAIRTVRVLRPLKAINRVPSMRILVNLLLDTLPMLGNVLLLCFFVFFIFGIIGVQLWAGLLRNRCFMDENLTIQGNITLPPYYQPEEDDEMPFICSLSGDNGIMGCHEIPPLKERGHECCLDKDDYFYYISVRQEFNVSGMCVNWNQYYNECRTGNSNPHKGAINFDNIGYAWIVIFQVITLEGWVEIMYYVMDAHSFYNFIYFILLIIVGSFFMINLCLVVIATQFSETKQREHQLMQEQRARYLSSSTVASYMEPGDCYEEIFQYVCHIVRKAKRRTLGFYNSLQDRRLGQTEPSTAKRSISGKDQRQYRRCRKHNSVDYMPQNLGQPIAMTLTSDPSNCPSCHKTQREATRRLSVLDSADSDQEEGGEASEDEGDGRKGSEGVADLEKEEEDGEEGGQRTHCSYIWQEVRVKLRGIVESKYFNRGIMIAILVNTISMGIEHHEQPEELTNILEICNVVFTSMFALEMILKLSAFGLFDYLRNPYNIFDSIIVIISIWEIIGQADGGLSVLRTFRLLRVLKLVRFMPALRRQLVVLMKTMDNVATFCMLLMLFIFIFSILGMHIFGCKFSLRTDTGDTVPDRKNFDSLLWAIVTVFQILTQEDWNVVLYNGMASTSPWASLYFVALMTFGNYVLFNLLVAILVEGFQAEGDANRSYSDEDQSSSNIEEFDKCQDSLDPKLCAVPMTPNGHLDPSIPCLNQPGAVVGIANPSRHTLQPDQVLVALESRKSSVMSLSSCRSSHYGAWGRSGGWGSRRSSWNSLARGHSLKHKPPSAEHESLLSGERHRISEGEREGLVRVRHHRRTLSLDTKGSSELVELAPAVPSGHRPTRRIGVTTGPIEHQDCNGKMPTITKDIFPEMNNRKDHGEDEEEIDYSLCFRIKKMIEAYKPDWCELREDWSIYLFSPQNRFRILCQTIIAHKLFDYIVLAFIFLNCITIALERPQIEQRSTERIFLTVSNYIFTAIFVAEMTLKVVSLGLYFGDQAYLRSSWNILDGFLVLVSIIDIVVSVASAGGAKILGVLRVLRLLRTLRPLRVISRAPGLKLVVETLISSLKPIGNIVLICCAFFIIFGILGVQLFKGKFYHCLGVDIRNITNRSDCMAANYKWVHHKYNFDNLGQALMSLFVLASKDGWVNIMYNGLDAVAVDQQPVTNNNPWMLLYFISFLLIVSFFVLNMFVGVVVENFHKCRQHQEAEEARRREEKRLRRLEKKRRKAQRLPYYATYCSVRLLIHSVCTSHYLDIFITFIICLNVVTMSLEHYNQPMSLETALKYCNYMFTAVFVLEAVLKLVAFGLRRFFKDRWNQLDLAIVLLSVMGITLEEIEINAALPINPTIIRIMRVLRIARVLKLLKMATGMRALLDTVVQALPQVGNLGLLFMLLFFIYAALGVELFGKLVCNDEHPCEGMSRHATFENFGMAFLTLFQVSTGDNWNGIMKDTLRDCTQDDRSCLSNLQFISPLYFVSFVLTAQFVLINVVVAVLMKHLDDSNKEAQEDAEMDAEIELEMAHGLCPRSSSSPSSGQDGGGGRVDPEGHLCMTCYSPAQENLWLDSVSLIIKDSFDGELMIIDNLSGSVFHHYSSPAMCEKCNHDKQEVQLAEMEGLSLNSDKSSILLGDDGDNHSVDQLSPKETQDGLDSPSSLGMGDLEECLLPLANSEASPAPESYLCETEETPFSPVQSWLKNESNQGPSTPFSPDIPNPLLTVPAEFLHPVVSADRPGQEKASCPENLPKISLQASWASRQSPNVNCSLIRQTTESDTSLDDSRSNSSEGSLQTTLEDSLNLSDSPQCTLDLPVALCPVPATATQRALVAPLSPASRRQSLRGRGIFTLRNIRRHQRSHSSGGSTSPGCTYHDSMDPSDEEGAGSSLRGGGNNSEQSETLSSLSLTSLFSPPPPPQGLTLVKKCNSTSSLHANPSSRGSAATEAKQFYSVDPRGFLSMPSWVTDFCKNSPSPREGKVPDLGSDKLGTVDSSSPVPPELELGNGVSKRNR

>XP_020665783.1|Pogona_vitticeps_Cav3.3

MAEHGTHTSSAWTSGNPPTYNPILCPVEHVDLGISEQSTPLEQPPPSPDLGDDDDDAEMEPAPDPDVPYPDLAPVVFFCLKQTTSPRSWCIKMVCNPWFEFVSMMVILLNCVTLGMYQPCDDMDCLSNRCQILQVFDDFIFIFFAMEMVLKMVALGIFGKKCYLGDTWNRLDFFIVMAGMVEYSLDLQNINLSAIRTVRVLRPLKAINRVPSMRILVNLLLDTLPMLGNVLLLCFFVFFIFGIIGVQLWAGLLRNRCFMDENLTIQGNITLPPYYQPEEDDEMPFICSLSGDNGIMGCHEIPPLKERGHECCLDKDDYYYYTSVRQEFNVSGMCVNWNQYYNECRTGNANPHKGAINFDNIGYAWIVIFQVITLEGWVEIMYYVMDAHSFYNFIYFILLIIVGSFFMINLCLVVIATQFSETKQREHQLMQEQRARYLSSSTVASYMEPGDCYEEIFQYVCHIVRKAKRRTLGFYNSLQDRRLGRTETSAAERSANRKVQRQYRRCRKHNSMDHMPQGLGQPIAVTLTSDPSNCPRCHKAQQEAARRLSVLESADSDQEEGGGANEEEGGSRKGSVEAADLEKEEEDGEKGRQPTLCSDMWQEVRVKLRGIVESKYFNRGIMIAILVNTISMGIEHHEQPEELTNILEICNVVFTSMFALEMILKLAAFGLFDYLRNPYNIFDSIIVIISIWEIVGQADGGLSVLRTFRLLRVLKLVRFMPALRRQLVVLMKTMDNVATFCMLLMLFIFIFSILGMHIFGCKFSLRTDTGDTVPDRKNFDSLLWAIVTVFQILTQEDWNVVLYNGMASTSPWASLYFVALMTFGNYVLFNLLVAILVEGFQAEGDANRSYSDDDQSSSNTEDFDKFQDSSDPKLCAIPMTPNGHLDPSLPCLNQPGAAAGIANPSCNTLQPDKVLVALESRKSSVMSLSSSRSSYYGAWGRSGGWGSRRSSWNSLARGHSLKHKPPSAEHESLLSGERHRILEGEREGPVRVHHHRRTLSLDIKGSSELVELAPSIPPGHRPARKAGVTTGPNEHQDCNGKMPNLAKDIFPEMNNRKDHGEDEEEIDYSLCFRIRKMIEAYKPDWCELREDWSIYLFSPQNRFRILCQTIIAHKLFDYVVLAFIFLNCITIALERPQIEQGSTERIFLTVSNYIFTAIFVAEMTLKVVSLGLYFGEQAYLRSSWNILDGFLVFVSIIDIVVSVASAGGAKILGVLRVLRLLRTLRPLRVISRAPGLKLVVETLISSLKPIGNIVLICCAFFIIFGILGVQLFKGKFYHCLGVDIRNITNRSDCMAANYKWVHHKYNFDNLGQALMSLFVLASKDGWVNIMYNGLDAVAVDQQPVTNNNPWMLLYFISFLLIVSFFVLNMFVGVVVENFHKCRQHQEAEEARRREEKRLRRLEKKRRKAQRLPYYATYCSVRLLIHSVCTSHYLDIFITFIICLNVVTMSLEHYNQPMSLETALKYCNYMFTTVFVLEAVLKLVAFGLRRFFKDRWNQLDLAIVLLSVMGITLEEIEINAALPINPTIIRIMRVLRIARVLKLLKMATGMRALLDTVVQALPQVGNLGLLFMLLFFIYAALGVELFGKLVCNDENPCEGMSRHATFENFGMAFLTLFQVSTGDNWNGIMKDTLRDCTHDDRSCLSNLQFISPLYFVSFVLTAQFVLINVVVAVLMKHLDDSNKEAQEDAEMDAEIELEMAHGLYPHSALNPGQDGGGGRMDPEGRLCMACFSPAQENLWLDSVSLIIKDSFDGELMIIDNLSGSVFHHYSSPAMCEKCNHDKQEVQLAEMEGLSLNSDKSSILLGDDGDSHSVDQPSPKETKDDEDPPLMGTEGLEECHFALTNTEAPPAPESCLCEMENTPFSPLQSWLSNENNQGTPTVLSPDASNPLLPAQAEFFLPSASANQPSQEKSSGPESLPKISLQASWASLQSSKVNCSLIHQTTESDTSLDDSRSNISEGSLQTTLEDSLNLSDSPQCVLDLPAALCPVPPTATQKALVVPLSPASRRQSLRGRGIFTLRNIRRHQRSHSSGGSTSPGCPYHDSMDPSDEEGAGSSLRGGGNNSEHSETLSSLSLTSLFSPPPPPQGLALMKKCNSTSSLHASASVRSSAATEAKPFYSVDPRGFLSMPSWVTDFCKNSPSPREGCVPEGSNRTGTPDHSSPVPSELQLESGVSKRNR

>NP_066919.2|Homo_sapiens_Cav3.3

MAESASPPSSSAAAPAAEPGVTTEQPGPRSPPSSPPGLEEPLDGADPHVPHPDLAPIAFFCLRQTTSPRNWCIKMVCNPWFECVSMLVILLNCVTLGMYQPCDDMDCLSDRCKILQVFDDFIFIFFAMEMVLKMVALGIFGKKCYLGDTWNRLDFFIVMAGMVEYSLDLQNINLSAIRTVRVLRPLKAINRVPSMRILVNLLLDTLPMLGNVLLLCFFVFFIFGIIGVQLWAGLLRNRCFLEENFTIQGDVALPPYYQPEEDDEMPFICSLSGDNGIMGCHEIPPLKEQGRECCLSKDDVYDFGAGRQDLNASGLCVNWNRYYNVCRTGSANPHKGAINFDNIGYAWIVIFQVITLEGWVEIMYYVMDAHSFYNFIYFILLIIVGSFFMINLCLVVIATQFSETKQREHRLMLEQRQRYLSSSTVASYAEPGDCYEEIFQYVCHILRKAKRRALGLYQALQSRRQALGPEAPAPAKPGPHAKEPRHYHGKTKGQGDEGRHLGSRHCQTLHGPASPGNDHSGRELCPQHSPLDATPHTLVQPIPATLASDPASCPCCQHEDGRRPSGLGSTDSGQEGSGSGSSAGGEDEADGDGARSSEDGASSELGKEEEEEEQADGAVWLCGDVWRETRAKLRGIVDSKYFNRGIMMAILVNTVSMGIEHHEQPEELTNILEICNVVFTSMFALEMILKLAAFGLFDYLRNPYNIFDSIIVIISIWEIVGQADGGLSVLRTFRLLRVLKLVRFMPALRRQLVVLMKTMDNVATFCMLLMLFIFIFSILGMHIFGCKFSLRTDTGDTVPDRKNFDSLLWAIVTVFQILTQEDWNVVLYNGMASTSPWASLYFVALMTFGNYVLFNLLVAILVEGFQAEGDANRSYSDEDQSSSNIEEFDKLQEGLDSSGDPKLCPIPMTPNGHLDPSLPLGGHLGPAGAAGPAPRLSLQPDPMLVALGSRKSSVMSLGRMSYDQRSLSSSRSSYYGPWGRSAAWASRRSSWNSLKHKPPSAEHESLLSAERGGGARVCEVAADEGPPRAAPLHTPHAHHIHHGPHLAHRHRHHRRTLSLDNRDSVDLAELVPAVGAHPRAAWRAAGPAPGHEDCNGRMPSIAKDVFTKMGDRGDRGEDEEEIDYTLCFRVRKMIDVYKPDWCEVREDWSVYLFSPENRFRVLCQTIIAHKLFDYVVLAFIFLNCITIALERPQIEAGSTERIFLTVSNYIFTAIFVGEMTLKVVSLGLYFGEQAYLRSSWNVLDGFLVFVSIIDIVVSLASAGGAKILGVLRVLRLLRTLRPLRVISRAPGLKLVVETLISSLKPIGNIVLICCAFFIIFGILGVQLFKGKFYHCLGVDTRNITNRSDCMAANYRWVHHKYNFDNLGQALMSLFVLASKDGWVNIMYNGLDAVAVDQQPVTNHNPWMLLYFISFLLIVSFFVLNMFVGVVVENFHKCRQHQEAEEARRREEKRLRRLEKKRRKAQRLPYYATYCHTRLLIHSMCTSHYLDIFITFIICLNVVTMSLEHYNQPTSLETALKYCNYMFTTVFVLEAVLKLVAFGLRRFFKDRWNQLDLAIVLLSVMGITLEEIEINAALPINPTIIRIMRVLRIARVLKLLKMATGMRALLDTVVQALPQVGNLGLLFMLLFFIYAALGVELFGKLVCNDENPCEGMSRHATFENFGMAFLTLFQVSTGDNWNGIMKDTLRDCTHDERSCLSSLQFVSPLYFVSFVLTAQFVLINVVVAVLMKHLDDSNKEAQEDAEMDAELELEMAHGLGPGPRLPTGSPGAPGRGPGGAGGGGDTEGGLCRRCYSPAQENLWLDSVSLIIKDSLEGELTIIDNLSGSIFHHYSSPAGCKKCHHDKQEVQLAETEAFSLNSDRSSSILLGDDLSLEDPTACPPGRKDSKGELDPPEPMRVGDLGECFFPLSSTAVSPDPENFLCEMEEIPFNPVRSWLKHDSSQAPPSPFSPDASSPLLPMPAEFFHPAVSASQKGPEKGTGTGTLPKIALQGSWASLRSPRVNCTLLRQATGSDTSLDASPSSSAGSLQTTLEDSLTLSDSPRRALGPPAPAPGPRAGLSPAARRRLSLRGRGLFSLRGLRAHQRSHSSGGSTSPGCTHHDSMDPSDEEGRGGAGGGGAGSEHSETLSSLSLTSLFCPPPPPPAPGLTPARKFSSTSSLAAPGRPHAAALAHGLARSPSWAADRSKDPPGRAPLPMGLGPLAPPPQPLPGELEPGDAASKRKR

>XP_021329632.1|Danio_rerio_Cav3.3

MNVGLNSCFLTWFERISIMVILLNCVTLGMYQPCENIDCTSERCQVLQAFDAFIYIFFALEMVVKMVALGIFGRRCYLGDTWNRLDFFIVMAGMVEYSLDLQNINFSAIRTVRVLRPLKAINRVPSMRILVNLLLDTLPMLGNVLLLCFFVFFIFGIIGVQLWAGLLRNRCYPEENFTLTSGLTLPTPYYQPEEDDERPFICSLAQDNGIMSCSDVPARREGGRECCLDKEDALHRQALGLSAEPLVNGSASAMGLCVNWNQYYTRCHTGHTNPHKGAINFDNIGYAWIVIFQVITLEGWVEIMYYVMDAHSFYNFIYFIFLIIIGSFFMINLCLVVIATQFSETKQREHQLMQEQRARYLSSSTLASLAEPGDCYEELFQLVCHILRKARRRSAALYYMLRGKAPPPGGGRGRGKGGAGSIGGGANVNGGEKHHCHSAQTSHCIHETKLDHSSENAVSLSISPNPEECPQCAAALSLKEGEESTGHSANGEEEEGALEETDRDENRLDKKRSSDSDDTLKKKRTCLGKCKDVWDEIRVKLWGIVESKYFNRGIMIAILINTISMGIEHHNQPDELTNVLEICNIVFTSMFTLEMILKLTAFGFFEYLRNPYNIFDGIIVIISVCEIIGQSDGGLSVLRTFRLLRVIKLVRFMPALRRQLVVLMKTMDNVATFCMLLMLFIFIFSILGMHIFGCKFSLKTEAGDTVPDRKNFDSLLWAIVTVFQILTQEDWNMVLYNGMASTSPLAALYFVALMTFGNYVLFNLLVAILVEGFQAEGDANRSYSDDDRSSCNLEETEKKDSLQLSDPKISTLTPNGHLDLAPAPNARGFYPSERLSFALGSRKSSVSSLGRASLEQTTLCPGRASLYHNWGRPARPGIWSRRSSWNSLGRSSRCLGMGGGGSLRVRSPHCHPAEQESLLSPPPPPLHPPLLPRHFPPRRERRALSLELPELLQVPGPPLPPLHPRQRKKSFSGGLGSVGEHQDCNGKTPSVQPQIINEVYPQVNTRKDREDLEDELDYSLCFRIQKMMEVYRPDWCETREDWSVFLFSPQNKFRLLCQSIIAHKLFDYVVLAFIFSNCITVALERPKILQGSLERLFLTVSNYIFTAIFVGEMTLKVVSMGLYIGEQAYLRSSWNILDGFLVFVSLIDIVVSMAGGAKILGVLRVLRLLRTLRPLRVISRAPGLKLVVETLITSLKPIGNIVLICCAFFIIFGILGVQLFKGKFYYCLGLDVKNITNKSDCLLANYKWVHHKYNFDNLGQALMSLFVLASKDGWVNIMYHGLDAVAVDQQPITNNNPWMLLYFISFLLIVSFFVLNMFVGVVVENFHKCRQHQEVEEAKRREEKRQRRMEKKRRKAQKLPYYASYSHVRLMIHTLCTSHYLDIFITFIICVNVVTMSLEHYSQPHSLEIALKYCNYFFTSTFVLEAVLKLIAFGFRRFFKDRWNQLDLAIVLLSVMGITLEEIEISAALPINPTIIRIMRVLRIARVLKLLKMATGMRALLDTVVQALPQVGNLGLLFMLLFFIYAALGVELFGELVCNEDYPCEGMSRHATFENFGMAFLTLFQVSTGDNWNGIMKDTLRECPPGEYTCNPSLQFISPLYFVSFVLTAQFVLINVVVAVLMKHLDDSNKEAQEEAEMDAEIEMELAQGTLCCIGGGVAVADRTGMGHQGAASCSVVPPHSPITHPVAHEPNSIRRLYSPAQENLWLDSVSLLIKDSFEGEMLMIDNLSGSVFHHYSSPPVCKDCRTHPQEIHMAELEQASLRSEQLSDKSSSPALPDDLSLDEQSVYQMAAKEGKERVCDECHHSEEPGGRGNSRLRRSSRVHSSGTEDGGTCQSPHHNTPARHSIGGVPCSSSSQEHPGVSGGSSRSNTPVCLPAEFFHPAAAALPAPPRGRTGHKPRGLRLTSPASWASLRSPGANTRLLTTQYPSHSDSSLATGSSEGSLPTTMEEGLSFIVSTPQPLLDTLLDERTSLSTETLRPSPPATHILQTTRGHQRSRSSCETDINTGSIRQGSEADVGLVWCSSEQTCDSQQLSETQSSLSLTSLLLPSSLVPLSVKKCNSTGSLDQGTLTAQEKVFTKEPQGNITSPWKERKERQSEAAQGTDGESTSQITIAGSRKNR

>XP_019397684.1|Crocodylus_porosus_Cav3.2

MTDGGRSAPFPPPAGGEVRVPIACRPARDEDALSAGKPVRGSPGSRSRDAGSEEEEQVPYPALAPTVFFCLQQATRPRSWCLRLVCNPWFEHVSMLVILLNCVTLGMFQPCEDVKCKSERCTILEAFDHFIFAFFAVEMVIKMVALGIFGQKCYLGDTWNRLDFFIVMAGMLEYSLDGHNVSLSAIRTVRVLRPLRAINRVPSMRILVTLLLDTLPMLGNVLLLCFFVFFIFGIVGVQLWAGLLRNRCFFDSNFAMTYNLTFLHPYYQPDDGEDNPFICSSHRDNGMQKCSNIPNLKHLKVECTLSMDPYTPNLYNPDFSNRNACINWNQYYNVCMAGDVNPHNGAINFDNIGYAWIAIFQVITLEGWVDIMYYVMDAHSFYNFIYFILLIIVGSFFMINLCLVVIATQFSETKQRENQLMQEQRARYLSNDSTIASFSEPGSCYEELLKYICHIFRKVKRRTLRLYNNWQSKRRKKVNPNSTTNGQSRRGRKRITSIHHLIHHHHHHHHHHYHISNGSPRGPRSNPEICGLELKVMKPGGQLMLPPPSPNLQSPPAPDSESVHSIYHADCHVEAPQVKCKSSNTPASIKLAAGLTSHGNMNYPTILPSPASKANSVPGPKGKKNGSSPVPMVSSPVRLNADSFGKLQHLMGEHGLRRASSRLSGLNVPCPLPSPQASMLTCELQNCPYCASILEDPEFEFSESDSYDSDNNGVYEFTQDLRHGDQRDQIQQQRNKRRKKKKKPKERNKAARLWSAFGNKLKKIVESKYFNRGIMIAILINTLSMGIEYHEQPDELTNALEISNIVFTSMFALEMLLKLLAFGIFGYIKNPYNIFDGIIVIISVWEIIGQSDGGLSVLRTFRLLRVLKLVRFMPALRRQLVVLMKTMDNVATFCMLLMLFIFIFSILGMHLFGCKFSLKTKTGDTVPDRKNFDSLLWAIVTVFQILTQEDWNVVLYNGMASTSSWAALYFVALMTFGNYVLFNLLVAILVEGFQAEGDANRSDTDEDKTSANFDEDFEKLKDLRAAEMKMYSLAVTPNGHLEARGSMPPPIIMRTAATPMPTPKCSPHMDSVHTFVDSRRGSNASIDPLSYDQKSLSSLRSSPCANWGPSSNWGSRRSSWNSLGRAPSLKKKNQSGERESLLSGEGKGSTDDESDDAKSSTVSRASLHRRAESLDYRSSLDLPELLQLPSMRHSLSINPMAMLPTEYQDCNGKMMHVPNEFFLRIDSHKDDPLDYDDDMEDSYCYRIRKMLEPYKPEWCKNHEDWSLYLFSPQNRFRAMCQKVIAHKMFDHVVLVFIFLNCITIALERPDIDPHSTERIFLSVSNYIFTAIFVAEMMVKVVALGFFSGENTYLQSSWNVLDGVLVFVSIIDIIVSMASAGGAKILGVLRVLRLLRTLRPLRVISRAPGLKLVVETLISSLRPIGNIVLICCAFFIIFGILGVQLFKGKFYSCEGSDTKNITTKADCTNAHFKWVRRKYNFDNLGQALMSLFVLSSKDGWVNIMYDGLDAVGIDQQVATKDILRFFPGIQCESAWSTESEEVICGPEETLAVGSYVPRAGDREPCSLLCLGFSLSTEAQRRPYYADYSPARRYIHTLCTSHYLDLFITFIIGVNVITMSMEHYNQPKSLDEALKYCNYVFTIVFVFEAVLKLVAFGFRRFFKDRWNQLDLAIVLLSIMGITLEEIEMNAALPINPTIIRIMRVLRIARVLKLLKMATGMRALLDTVVQALPQVGNLGLLFMLLFFIYAALGVELFGKLDCSEDNPCEGLSRHATFTNFGMAFLTLFRVSTGDNWNGIMKDTLRECTREDKHCLSYLPVISPVYFVTFVLIAQFVLVNVVVAVLMKHLEESNKEAKEDAEMDAEIELEMSRGAATPASSGGGGGTAASDGRAPYGSQEGGKLETPLRVKQQALEALSCENVSLAMPGSDPLEGGETPVMPHTSFCSVPHHHVPTPSHKAPGHESEEGAACLWAKKESQNFLTVRKISVSRMHSLPNDSYMFRPVMPAKPPYPLHEVEMESHPCKSQRGSITSTRSQPVGTSSSLRIPTECLQLTPGPSYRNARDLWKIHPPTAAPRSPSLNRLLCRQEAIRTESVDGQKAEAKDNVQVESGESLQPKKPSQSTLSPPPLHPAPRSLPGTPPHSPASPSPHRHSSVCTRKHTCSQHSIPSWASSESSDPLRPENSLFESSDLADEEVSHINSSAKDWREAPGSRSLSVSPFASLASSPAHHKNCSTTSLAGSGGSREKDLKKFYSVDTKGFLDKPSWADNQRRHSIEICPSLDDGDCSFEQPAEGTPRSPVHVQLESEYVHGARRKKKMSPPCISIDPPMEDESGAASRTKASENSLLRRRTPSCESAAYRDSLDLAENPPGEQPSKAERRVQPLCRGEHLAIPNFSFEQSDASSLSSLSELLLDSGQSSPSSGESRPDPAALELTQLEPLQTQCNHPETNKELLVVTKSPLQKHGLVSTTVATDEDVEEPV

>XP_015149910.1|Gallus_gallus_Cav3.2

MSDGESREPRPAGGEVRVPITVPPRRAAEQPLDDAGSAPSGSPSRDPPPDLASEEEEQVPYPALAPTAFFCLKQTTRPRSWCLRLVCNPWFEHVSMLVILLNCVTLGMFQPCEDVECQSERCTILEAFDDFIFAFFAVEMVIKMVALGIFGQKCYLGDTWNRLDFFIVMAGMMEYSLDGHNVSLSAIRTVRVLRPLRAINRVPSMRILVTLLLDTLPMLGNVLLLCFFVFFIFGIVGVQLWAGLLRNRCFLDRNLATTYNLTFLHPYYRTDEAEENPFICSSHRENGMQKCSNIPTRKEYKVECTLSMDSYTPSLYNPDFSSKNACINWNQYYNVCMAGDVNPHNGAINFDNIGYAWIAIFQVITLEGWVDIMYYVMDAHSFYNFIYFILLIIVGSFFMINLCLVVIATQFSETKQREHQLMQEQRARYLSNDSTIASFSEPGSCYEELLKYICHIFRKVKRRTIRLYNNWQSKRRKKVNPNNTTNGQSRRGKKRITSIHHLIHHHHHHHHHHYHISNGSPRGPRSNPEICDLELKVMKPGGQLMLPSPSPNLQSPVPPPDSESVHSIYHADCHVEGPQVNCKSSNAPASIKLTTGLSNHGNMNYPTILPSPTSKASSVPVPKGKKNGSSPVAVVNSPVTLGTDSYGKLQQLVGEHGLRRTPSRLSGLSVACPLPSPPGGTLSCELQDCPFCASLLEDPEFAFSESDSCDSDSNGIYEFTQDLRHGDHRDQLQQQRGRRKRKKKKPKERNKVTRLWKAFGSKLKRIVESKYFNRGIMIAILINTLSMGIEYHEQPDELTNALEISNIVFTSMFALEMLLKLLAFGLFGYIKNPYNIFDGIIVVISVWEIIGQSDGGLSVLRTFRLLRVLKLVRFMPALRRQLVVLMKTMDNVATFCMLLMLFIFIFSILGMHLFGCKFSLKTDTGDTVPDRKNFDSLLWAIVTVFQILTQEDWNVVLYNGMASTSSWAALYFVALMTFGNYVLFNLLVAILVEGFQAEGDANRSDTDEDKTSANFDDDFEKLKDLRATEMKMYSLAVTPNGHLEGRGSMPPPIIMRTAATPMPTPKCSPHMDSVHTFVDSRRGSNASLDPLTYDQKSSSSLRSSPCANWGTNSNWGSRRSSWNSLGRAPSLKKKNQSGERESLLSGEGKGSTDDDSDDAKSSTVSRPSLHRRAESLDYRSSLDLPELLQLPPMRHSLSISPMAVLPAEYQDCNGKMVHVPSEFFLHIDGHKEEAVDYEDDMEDSYCYRIRKVLEPYKPEWCKSHEDWSLYLFSPQNRFRVMCQKVIAHKMFDHVVLVFIFLNCITIALERPDIDPHSTERIFLSVSNYIFTAIFVAEMMVKVVALGFFSGENTYLQSSWNVLDGVLVFVSIIDIIVSMASAGGAKILGVLRVLRLLRTLRPLRVISRAPGLKLVVETLISSLRPIGNIVLICCAFFIIFGILGVQLFKGKFYYCDGPDVKNITTKTDCTNAHYRWVRRKYNFDNLGQALMSLFVLSSKDGWVNIMYDGLDAVGIDQQPIQNHNPWMLLYFISFLLIVSFFVLNMFVGVVVENFHKCRQHQEAEEARRREEKRLRRLEKKRRSKEMYLSEAQRRPYYADYSPARKYIHTLCTSHYLDLFITFIIGVNVITMSMEHYNQPKSLDEALKYCNYVFTIVFVFEAVLKLVAFGFRRFFKDRWNQLDLAIVLLSIVGITLEEIEMNAALPINPTIIRIMRVLRIARVLKLLKMATGMRALLDTVVQALPQVGNLGLLFMLLFFIYAALGVELFGKLDCSEENPCEGLSRHATFTNFGMAFLTLFRVSTGDNWNGIMKDTLRECTREDKHCLSYLPVISPVYFVTFVLIAQFVLVNVVVAVLMKHLEESNKEAKEDAEMDAEIELEMSRGASTTSGSRGGAVALESRASCRSQEAMKSEPAAQAKHQALGGTAIPQPQDSQNMLTVRKISVSRMHSLPNDSYMFRPVMPAKAPFPLHEVEMEAYPCKSQRGSITSIRSQPVGTCSSLRVPPESLHVAVRSPGREGWHRWKVLPPHVVPRSPTLSKLLCRQEAVRTDSVDGHPPDPKDNLQAERGEMPPSPKHPQRTLTPSPLHPAASSLPGSPPRSPPTSPHRHPSLTRKHPCSQRGVPSPTESPPSPQGCLLEPSDLADEEVRHINSSAKDWGEHSEAGSCSTSPFVSPAASPVPLKNCSTSSLTAGSGKERDLKKFYSIDTKGFLTKPSWADDQRRHSIEICPSVRDGDCGFEERAEEKPRSPAHVQPESEYIHGARRKKKMSPPCISIDPPMEDDSGAASRTKPSENSMLRRRTPSCEFPAYREPLEPAEAQPGEPGCKAERRAQHLAIPGFSFEQSDGGGSLSSLSDLLDSGRSTPDCELRKPEPLKTQRSNAEKSKELLSVAKSPVRMQDLVTVATEEDIDEPV

>XP_020633697.1|Pogona_vitticeps_Cav3.2

MVISTMTRLSGSDGVGSSEGRWLGKSFNPLVSQPEELTNALEISNIVFTSMFALEMVLKLLAFGIWGYIKNPYNIFDGIIVVISVWEIIGQSDGGLSVLRTFRLLRVLKLVRFMPALRRQLVVLMKTMDNVATFCMLLMLFIFIFSILGMHLFGCKFGLKTDTGDTMPDRKNFDTLLWAIVTVFQILTQEDWNVVLYNGMASTSSWAALYFVALMTFGNYVLFNLLVAILVEGFQAEGDANRSDTDEDKTSLNFEEELERLKELASEMKMYSLAITPNGHLESKASMLPPIIMRTAATPMPTPKCSPHMDSAYPFADSRRGSNASVDPLSLSSPRSSPGTHWGGGGGSNWGSRRSSWNSLGRAPSLKKKTQCGERESLLSGEGKGSTDDEAEEAKLSAVSRPPPHHRAEPLDYRGAVELPELLQVPGIHPSLSLSPLRPTEYPDCNGKMLHVPNEFYLHMDGHKEDLLDLEEEEDSYCYRIRKAFEPYKPTWCKTHENWSLYLFSPQNRFRASCQKVIAHKMFDHVVLVFIFLNCITIALERPDIDPHSTERVFLSVSNYIFTAIFVAEMMVKVVALGFFSGENTYLQSSWNVLDGVLVFVSIIDIIVSMASAGGAKILGVLRVLRLLRTLRPLRVISRAPGLKLVVETLISSLRPIGNIVLICCAFFIIFGILGVQLFKGKFYHCEGPDIRNVSTKADCTNARYKWVRRKYNFDNLGQALMSLFVLSSKDGWVNIMYDGLDAVGIDQQPSQNHNPWMLLYFISFLLIVSFFVLNMFVGVVVENFHKCRQHQEAEEARRREEKRLRRLEKKRRKAQRRPYYADYSPARRYIHTLCTSHYLDLFITFIIGVNVITMSMEHFNQPKSLDEALKYCNYVFTIVFVIEAVLKLVAFGFRRFFKDRWNQLDLAIVLLSIMGITLEEIEMNAALPINPTIIRIMRVLRIARVLKLLKMATGMRALLDTVVQALPQVGNLGLLFMLLFFIYAALGVELFGKLDCSEDNPCEGLSRHATFNNFGMAFLTLFRVSTGDNWNGIMKDTLRECHRDDKHCLSYLPIISPVYFVTFVLIAQFVLVNVVVAVLMKHLEESNKEAKEDAEMDAEIELEMSRGVNTPVRVPEGSPALSKTGSPLLLKHQAPASLSCESISLAIPDLDLLLEGGKDPMMMEFAGFRSSPHRRLLTPSRQAPAGEAEVTEAASGDAAQNLLTVRKISVARMHSLPNDSYMFRPVAPAKTPYPLHEVELEAYSFKSPRGSISSVHSQPVGTCSSLRVPRAAFRPTTPPPRHEGGGPWKLRPPHKAPPSPSINRPLCRQEAVQRDSLDGPEAELKENPLTEPCESLPPKMLSQSILGGPLLQASPTSLPGTPPLPHASPQKPAGTHTCKPPFSQHKLASRPPSCSSHPEGSLFESSDLADEEVSHINSSARGWGQRPDAGDRSATPLPSPAQSPAPTKNGSTTSLTAGGGKEKDLKKFYSVDTRGFLAKPTWADDQRRHSIEICPSVIDRERGFAEAAEERKRTPLHLPSESEYLHGVRRKKKMSPPCISIDPPMEEENNPSTRTKASENSLLRRRTPSCESTAYRDSLDLADSQPGEPASKAERRAPPPACRGEHLTIPNFSFDQPDTTTGTPSDRLSDSGLATALSADSRPDDGAGLEVPKADLLKAQCGQAERNREVPGGPRSPLRKHELVPVIRATEGGVDEPV

>NP_066921.2|Homo_sapiens_Cav3.2

MTEGARAADEVRVPLGAPPPGPAALVGASPESPGAPGREAERGSELGVSPSESPAAERGAELGADEEQRVPYPALAATVFFCLGQTTRPRSWCLRLVCNPWFEHVSMLVIMLNCVTLGMFRPCEDVECGSERCNILEAFDAFIFAFFAVEMVIKMVALGLFGQKCYLGDTWNRLDFFIVVAGMMEYSLDGHNVSLSAIRTVRVLRPLRAINRVPSMRILVTLLLDTLPMLGNVLLLCFFVFFIFGIVGVQLWAGLLRNRCFLDSAFVRNNNLTFLRPYYQTEEGEENPFICSSRRDNGMQKCSHIPGRRELRMPCTLGWEAYTQPQAEGVGAARNACINWNQYYNVCRSGDSNPHNGAINFDNIGYAWIAIFQVITLEGWVDIMYYVMDAHSFYNFIYFILLIIVGSFFMINLCLVVIATQFSETKQRESQLMREQRARHLSNDSTLASFSEPGSCYEELLKYVGHIFRKVKRRSLRLYARWQSRWRKKVDPSAVQGQGPGHRQRRAGRHTASVHHLVYHHHHHHHHHYHFSHGSPRRPGPEPGACDTRLVRAGAPPSPPSPGRGPPDAESVHSIYHADCHIEGPQERARVAHAAATAAASLRLATGLGTMNYPTILPSGVGSGKGSTSPGPKGKWAGGPPGTGGHGPLSLNSPDPYEKIPHVVGEHGLGQAPGHLSGLSVPCPLPSPPAGTLTCELKSCPYCTRALEDPEGELSGSESGDSDGRGVYEFTQDVRHGDRWDPTRPPRATDTPGPGPGSPQRRAQQRAAPGEPGWMGRLWVTFSGKLRRIVDSKYFSRGIMMAILVNTLSMGVEYHEQPEELTNALEISNIVFTSMFALEMLLKLLACGPLGYIRNPYNIFDGIIVVISVWEIVGQADGGLSVLRTFRLLRVLKLVRFLPALRRQLVVLVKTMDNVATFCTLLMLFIFIFSILGMHLFGCKFSLKTDTGDTVPDRKNFDSLLWAIVTVFQILTQEDWNVVLYNGMASTSSWAALYFVALMTFGNYVLFNLLVAILVEGFQAEGDANRSDTDEDKTSVHFEEDFHKLRELQTTELKMCSLAVTPNGHLEGRGSLSPPLIMCTAATPMPTPKSSPFLDAAPSLPDSRRGSSSSGDPPLGDQKPPASLRSSPCAPWGPSGAWSSRRSSWSSLGRAPSLKRRGQCGERESLLSGEGKGSTDDEAEDGRAAPGPRATPLRRAESLDPRPLRPAALPPTKCRDRDGQVVALPSDFFLRIDSHREDAAELDDDSEDSCCLRLHKVLEPYKPQWCRSREAWALYLFSPQNRFRVSCQKVITHKMFDHVVLVFIFLNCVTIALERPDIDPGSTERVFLSVSNYIFTAIFVAEMMVKVVALGLLSGEHAYLQSSWNLLDGLLVLVSLVDIVVAMASAGGAKILGVLRVLRLLRTLRPLRVISRAPGLKLVVETLISSLRPIGNIVLICCAFFIIFGILGVQLFKGKFYYCEGPDTRNISTKAQCRAAHYRWVRRKYNFDNLGQALMSLFVLSSKDGWVNIMYDGLDAVGVDQQPVQNHNPWMLLYFISFLLIVSFFVLNMFVGVVVENFHKCRQHQEAEEARRREEKRLRRLERRRRSTFPSPEAQRRPYYADYSPTRRSIHSLCTSHYLDLFITFIICVNVITMSMEHYNQPKSLDEALKYCNYVFTIVFVFEAALKLVAFGFRRFFKDRWNQLDLAIVLLSLMGITLEEIEMSAALPINPTIIRIMRVLRIARVLKLLKMATGMRALLDTVVQALPQVGNLGLLFMLLFFIYAALGVELFGRLECSEDNPCEGLSRHATFSNFGMAFLTLFRVSTGDNWNGIMKDTLRECSREDKHCLSYLPALSPVYFVTFVLVAQFVLVNVVVAVLMKHLEESNKEAREDAELDAEIELEMAQGPGSARRVDADRPPLPQESPGARDAPNLVARKVSVSRMLSLPNDSYMFRPVVPASAPHPRPLQEVEMETYGAGTPLGSVASVHSPPAESCASLQIPLAVSSPARSGEPLHALSPRGTARSPSLSRLLCRQEAVHTDSLEGKIDSPRDTLDPAEPGEKTPVRPVTQGGSLQSPPRSPRPASVRTRKHTFGQRCVSSRPAAPGGEEAEASDPADEEVSHITSSACPWQPTAEPHGPEASPVAGGERDLRRLYSVDAQGFLDKPGRADEQWRPSAELGSGEPGEAKAWGPEAEPALGARRKKKMSPPCISVEPPAEDEGSARPSAAEGGSTTLRRRTPSCEATPHRDSLEPTEGSGAGGDPAAKGERWGQASCRAEHLTVPSFAFEPLDLGVPSGDPFLDGSHSVTPESRASSSGAIVPLEPPESEPPMPVGDPPEKRRGLYLTVPQCPLEKPGSPSATPAPGGGADDPV

>XP_009297960.1|Danio_rerio_Cav3.2

MLVILLNCVTLGMYQPCEDLKCQSEWCIVLQAFDDCIFAFFAVEMVIKMIALGIFGINGYLGDTWNRLDFFIVMAGMMEYSLDGHNASLSAIRTVRVLRPLRAINRVPSMRILVTLLLDTLPMLGNVLLLCFFVFFIFGIVGVQLWAGLLRNRCFMPSKVRDQYNLSYMSLYYENEDGEDNPFICSSSKDNGMKRCSAVPPLKEGVECTLNASFLGHVYSGLSGTGNYSCVNWNQYYSECKPGDLNPHKGAVNFDNIGYAWIAIFQVITLEGWVDIMYYVMDAHSFYNFIYFILLIIVGSFFMINLCLVVIATQFAETKQRENMLMKEQRARYRSNDSTLASYSEPGSCYEEMLKYVSHLYRKVKRRLSRIYNSWQSKRRKKVNPNTGAGGGANGHSRHRGYGHWVQSIHSLIQQHQRQHCHLSNGSPSPVATTGSSEALEMRSITAGQQLTVSSQPTATHNLSSSTQSVHSIYQGDFREAPQERTPTNVAVPALSRLNGGMNYPTILPSLICNYSGGGSLGKERGKSDANTGHEVIQDFDKLHQQIEAHSHSRVLLQMAGVVPTQLLLEVMSCPYCVRALESMDLEIGDSDYPRSEGDMVYEFSYDAGFITGRSAHHKMQEKQNRLVQFWEDFRERLTRIVDSKYFNRGIMIAILINTLSMGIEYHEQPEELTNILEISNIVFTSMFVLEMLFKLLAFGIFGYIRNPYNIFDGVIVVISVWEIIGHADGGLSVLRTFRLLRVLKLVRFLPALRRQLLVLMKTMDNVATFCMLLMLFIFTFSILGMHLFGCKFSLKMENGDTIPDRKNFDSLLWAIVTVFQILTQEDWNVVLYNGMASTSPWAALYFVALMTFGNYVLFNLLVAILVEGFQAEGDANKSDGDEEKTSVNSEEKMEQLQSSDLKFYSLMLSANGHVDPCGTLPPPIIMRTAATPMPTPKSSLGPESVFDLTESRRGSAVSIDPNAFDQKSLSSPRRSPCHPRGSGSNWGSRRSSWNSLGRAPSLKRKDTSGERESLLSGEGRGSFEDDDIEAEKLTNISGGSLQHPSSLDIPELPQMGEYLDCNGRSQHILADLSASLNKEDSIAEEDLDDSFCFRLRRTLAAYKPKWCKDHEEWSLYLFSPHNKFRMMCQKLISHKMFDYVVLVFIFLNCITIALERPHIQQSERLFLLVSNYVFTVIFVAEMTVKVVALGFYSGNQSYLKSTWNVLDGVLVFVSLIDILVSLAWTGNRIFGILRVLRLLRTLRPLRVISRAPGLKLVVETLITSLRPIGNIVLICCAFFIVFGILGVQLFKGKFFHCEGGDTRNITNKSDCLQANLKWIRRKYNFDNLGQALMSLFVLSCKDGWVNIMYDGLDAVGVDQQPERNHNPWMLLYFISFLLIVSFFVLNMFVGVVVENFHKCRQDQEEVEARLLELKRQKLMEKKRRSKENCGSEALRRPYYADYSPARLYIHTLCTNHYLDLFITGIICINVVTMSIEHFNQPSYLDEALKYCNYVFTIIFIIEALLKLVAFGIRRFFKDRWNQLDLAIVLLSIMGITLEEIKMNAALPINPTIIRIMRVLRIARVLKLLKMATGMRSLLDTVMQALPQVGNLGLLFMLLFFIYAALGVELFGKLECTENNPCEGLSPHATFENFGMAFLTLFRVSTGDNWNGIMKDTLRECLPSETQCLSYLPWVSPIYFVTFVLMAQFVLVNVVVAVLMKHLEESNKEAKEDAEMDAAIKLAILKETRRLSTISASATAGPEGRATYIEPHPDSPEQDEGMQRTQGNLLSPRKMSVSRMHSLPNDSYMFRPVRPASAPYPLEEVETCQDYLELGSITSVHSQPCESLSLLQVPGVPPHSQFNCLPKIPPPMPSPTHSIDRILRRQRAIKNDSIDMHNFDSKENVERQKGITTGSFKLHVPQITGISQGSPLLPPRGPSVHTFQQPHSQHNISSRPPSLASSSRSTSPSGSMFDSADPTDEEVRHINSTARLWIGVHAEARSHSLSPYSTSGTLLPIPVKNCSSAANLTIKDPKKCFSVDTEGFLERPYASDDHRRHSIEICSSGDDGLFGQELCEKRHVPTRGKSLHIPRKTKMSPPCISIEPPFEKDVSPSHASMKKLSDSMLLRRRTPSYDLALQRDSLDLPENQLSTQPLIKVERHPSHCAEYLSLPHPAPMSGLAGVLCESTQPPPFDSRYEESTDFGSVNRTAQ

>XP_018667817.1|Ciona_intestinalis_Cav3

MYAETDFNRSIPVTSRFRAVRNAERRRKRSRVGNTNEADGINISQTLSSKMESGKTVAFQSRSEDDSPEDTKKDGSEDEDSGRSYEGPLFPALYPVVLLSLKQTSVPRIWCLRMVANPWFERVSMLVILINCVTLGLYQPCQHRTGLHCESERCMVLEMFDHFVFAFFALEMLIKMLAMGVWGKLGYLGEAWNRLDFFIVLCGMLEYTLQMEDTMNFTSVRTVRVLRPLRAINRVPNMRILVMLLLDTLPMLGNVLMLCSFVFFIFGVVAVQLWEGTLRQRCFLDPHVFNITGDMVTLKHSNISINLPPYFTLDDSDDEVAICSLPIDNGDFKCMDRNIITPNKVNNVTCSLNISSFIPGQAMDHIYTDLTGATNCIDWNQYFNTCKAGDQNPYLGSINFDNIMYAWVAIFQVISLEGWVDIMYYLMDGYSFYSFIYFILLIVIGSFFMINLCLVVIATQFSETKQREQRLIEEQRLRFKSNDSTLASYSPPGNCYEEMLKYISHLYRRNKRKIRKRWNKWKLKKKMNESVEIKRESSTCQSKKKKKSIHLHHHHHHHHHHYHISETLDRTEGIIRTLEKQDNTVDTVSEPRARTPQNYPSILPQRLMVTDGTSEIGDPIKTGYKLTVPIPTPGLSTSLHSVSGSGVGQRSPSLCSDRRPSAEINVNPGSVYFKRSTLDCGFPTERFCTGVRIHDDIIVANPVEVLGKTSPCPQSFVTSELNCSCAEYEVNPCIFEPTAEVEVENTSRFRRCLCCCGSSQNEEDKENSESILSKFQTQTKVVVDSNYFNRGIMVAILINTLSMGIEHHNQPTGLTEVLEISNVVFTTLFALEMLSKIVAYGFAGYIKNLYNVFDALIVIISVWEIAAGTQNSGGGLSVLRTFRLLRVLKLVRFMPALQRQLVVLMKTMDNVATFMMLLTLFIFIFSILGMHLFGCDFCWVNQHGRTECDRKNFDSLLWAFVTVFQILTQEDWNIVLYNGMAATSPFAAIYFVTLMTIGNYVLFSLLVAILVEGFQAECYSTEAENQTSQDEEDSDAEEPAILAENYTKRVAKALGITGPIAGKSMSPKIKEKKKTAPHDGGDNLLEDTSSSSDDDEEVELLTRSSNPRIKITSSKDDILSANSSFSETTGENRLLPVITRTAATPLHTPTLHSPIQRHEIEKSFMEKTVENQTLKSGSSTESEGHSGQLNKKHSLLSRKSSTESCLSVSSVIKFAGGSPISRTGSWRLRRFKGLPLDRQSLVCSEEEDQDNEIEDDMYSNASFENEPEQEIEIKTPNVITESQDSSEIESTYSPINNNNNNNINIEQDTHSNSKADMQLKETDSNSASSNKSRCNCIRSCKCPSIRRRSPDWVQRKKDWSLYLFSPELKFRKAVQKITEHKLFDYMILLLIFGNCITIALERPSLKEEDHERKVIDGFNNVFTFVFLLELILKVIASGFYIGHKAYLKSGWNVLDFFLVASSLIDVIMTLTYSSGSKLLGILRVFRLLRALRPLRVISRAPGLKLVVQTLISSLKPIGNIVLICCAFFLIFGILGVQVLKGKFYYCDGPDLRNVTNKTDCLLSSNNQWVNRRYNFDDVGQALMSLFVISSKDGWVEIMYHGIDATGIDQQPIRNSNPWMLLYFVSFLLIVGFFVLNMFVGVVVENFHRCREEHELEEQKRREERRKRKQERLDYKRSMRNQRGDKYLTFRRKTLSKAKNHVTGVRGRCTNCWRKITTFFCGSPSFQANTKYYEDYGRTRRALHAFCLNKYFEIGVSIVIGINIFTMAAEHYQQPKVLDQALKIANYFFTAVFVLEAILKLIALGVRRYFRDKWNQVDMIIVILSLVGIAVEALMSAGDRSLLINPTIIRVMRVLRIARVLKLLKVSKGIRSLLETVANALPQVGNLGLLFLLLFFIFAALGVELFGTLSCDELHPCNGLSRHASFSNFGIALLTLFRISTGDNWNGIMKDVMKRPLSGPLYNDTIRETCDSSDACVTNCCGSSIISPIYFVLFVMTAQFVLVNVVVAVLMKQLEDNRAEDDEEDEVFEDDSNNFTDNDDQVIIVPETVDRDCRSNATGPVEVLIDTHTHKADDDIVSPSICEVKQSRVNSAPRKLVKQQKIDLADSTEHSLHWTPTETPEEMTTKSLVKHVLDDPNFVTLAQSMPCLRPTPIHTRQLLPLTTHFSGCRSVEVTPQQSPNFRVQRKESVFILPQSTSSVPTCSLDIPQSTSNVIVTSQTNGYVRADSVTSLTSSVHISMTSPRNDVKPSTPNCDTTPPRTSTITQSGKNTFFDDVTPSRSVGDAISESFDNVTKPQRPSDVTIYKNDEKPSGNTPVSRTTICDNPSVTSKVDDVTHCDVINKPRWPPSQQGASPKSTLQSCGKRHLVRTFSFNPNLLRHNDVTQPTIYATSSIDDVRKSYSDS

>NP_001314887.1|Apis_melifera_Cav3

MSLHYPQGYRYRPASKAVGSGNGEQGSDSDVAELSDLEDDEDEDGGGQSTNRCVEQNVEKEEKEDEDDNEDEDDDDNEDDEDQEDEEEDENEEDNGVDDVDGDELPYPGFVPVALRYLDQNTRPRNWCLALITNPWFERVSMMVILLNCITLGMYQPCVDDQCVTNRCKILQMFDDIIFAFFSLEMTIKMVAMGIYGKGTYLADSWNRLDFFIVIAGALEYCLNVENMNLSAIRTIRVLRPLRAINRIPSMRILVMLLLDTLPMLGNVLLLCFFVFFIFGIVGVQLWEGILRQRCFLKALPNVKYPDDLEKYFEYQGQDYICSRPDDNGMHSCSNLPPLKLGNVVCNNTALPNNNTTFITNDTCVNWNYYYTECKGQGNNPFQGTISFDNIGLAWVAIFLVISLEGWTDIMYYVQDAHSFWDWIYFVLLIVIGSFFMINLCLVVIATQFSETKKREMERMRLERARFHSTSTLASSTNTSEPTTCYAEIVKYIAHLWRRGKRRLMKRYRVYLYKRQQKREQNLLKEQQQHGHPFRGGAPNSDSNRLPGDRRLHHGRCPRLLAALEYAEQQQQQQQGIGSGGGSLTDFPANSPPTQTAILATAIGNIGGSNGNLAIAPRASPEVSEADVSLNVYNRIGLHRTSSVSCNGSDNVLSNATEASIQTNNVLLSPPCTHYRRRSSVMFSDVVLLHGSNNIGNTLQAGMAVSPGERNVCSSEKMTQTGDGNVWSSPLPDHIQMQAELGGNEAMTCQELLALSGALSAALPTGQLALDSFLNSFTKGITDRHITLEDRTQWLASDIDNCSCCCELQGIDQWPDEGDKWTKNSRAKRFLRSCGNSCICAIRCIRRLIKKLVEHKYFQQGILLAILINTLSMGIEYHNQPEQLTIVVEVSNIVFSAVFAVEMLLKIIAEGPFGYISNGFNVFDGVVVVLSVVEICQAFVEERGGSSGLSVLRTFRLLRILKLVRFLPNLRRQLFVMLRTMDNVAVFFSLLVLFIFIFSILGMYLFGGKFCMWADRSRPCTCAEVVSRHPLCRCDRKHFNDIVWALVTVFQILTQEDWNVVLFNGMQKTSHWAALYFVALMTFGNYVLFNLLVAILVEGFSSERNERREREQREMARLAAKETGIGSDDGSSRISRSHSITDSDTYTQDRKNSWQSAEELHKYKDNNSKEKQNTVWKHQIDEPKCNIQKVNKMKGSTGQPPIITHTAATPQDSPNTTLDVGRVVYPTAALSIESIDRSGSQCSISSGLLKLPDVSNKIPTKNLIAGQFPRRINLVTAVVNPTMRESSNSSSPRIQRGYSWKLSRPSLRKKRWLQTEDESPRRATVLNNGRSTILGSNSTFNGGYLHGSIRNDTQSDTPNNRTTVLTSNNRSLSPNNSIESRSSSIRRYTATPNQMRWISDLSRRNSLRENENVQSPTRKTLPLDEVPMQCSTARTINNLSMETGPLPRIKRLPDQDDDNPRTDEQTPPLNGHGSASSIERIKKIFMFFEPKGCLKERDDYSLYIFPPNNRFRVLCRLLVDQRWFDNVVLFFIGLNCITLAMERPNIPPDSGERLFLSTANYIFTGVFAVEMFIKVVASGMLYGSDAYFTSGWNIMDGVLVIISIIDLSMSLLSSSSPRIFGILRVFRLLRSLRPLRVINRAPGLKLVVQTLLSSLRPIGNIVLICCTFFVIFGILGVQLFKGAFYYCEGPDIKNVRNKTDCLADKRNVWLNRKYNFDDLGKALMSLFVLSSRDGWVNIMYTGLDAVGVDQQPIENYSEWRLLYFIAFILLVGFFVLNMFVGVVVENFHRCREEQEKEERVRRAAKRALQMEKKRRKMHEPPYYTNYSKSRLFVHNVVTSKYFDLAIAAVIGLNVVTMAMEFYMMPKALTYALKIFNYFFTAVFILESFMKLLALGLHLYLKDKWNQLDVGIVILSVVGIVLEEVESKIIPINPTIIRVMRVLRIARVLKLLKMAKGIRALLDTVMQALPQVGNLGLLFFLLFFIFAALGVELFGRLECSDDMPCQGLGEHAHFSNFGMAFLTLFRVATGDNWNGIMKDTLRDDCDEAADCVKNCCVSTIIAPIFFVIFVLMAQFVLVNVVVAVLMKHLEESHKQMEDELDMETQLERELAAEQEELLEVEDEEDDETIKRERDDGDIDEDDEDVREHESILVANEKIPASRPGLAKVRSLPANFIYNPPRERNADDGTISVSLARRSSYHRSSSRPSKFKSKRRQTFHSGHHQRRSLLPMHFEVAEVFEKPVSNLSVPRIIPQHNYDATSRDITHVQRQRLQATSDGQENKSPLNVSKPPPSSSSPMSLAGSVTTLTCPKISSERYLMPTFNIYPSKATLSCRPSSEAMQSQFDTNVSIGSIVSKTDSTMNGYMETGVPRSNGDRSNDSKEKNERRVSAPPTDDLDVQSIINERRPSKLKSGNLTETMRIVSDQSSSTRIESYGQVYVTEERFEEVSPMSTGNMSDSIIGSSGTNGSESVTGSGTGSASGSGSRNESAVMHVTEGVCATIGSDVRIYVDDTDSASSNNHQRRKANEGTPEVSMTISSVIAIPDVTIEEERSERFDVPSSDGPSDPS

>XP_021706157.1|Aedes_aegypti_Cav3

MLNAYSESQSDLENEQNFGAKQKPRPSVNRQCQDRRSSLVKRSVSAKSNRRPSINSVYNEGQFSESCSMRKKPNKSVGSVSSTSGTTMDSSGSSCSDGDTSSSYGEPNLPYPGFAEYSLKYLSQDTKPRIWCLQLITNPWFERVSMLVILLNCVTLGMYQPCVDDACVTNRCKILQIFDDIIFAFFSLEMTIKIVAMGAWGKGTYLADSWNRLDFFIVLAGALEYCLQVENLNLTAIRTIRVLRPLRAINRIPSMRILVMLLLDTLPMLGNVLLLCFFVFFIFGIVGVQLWEGILRQRCVIKLPDNVSAPDISFYYEFSKEQDYICSKPDDSGMHLCNNLPPYRIGPLVCNASAMPFSNNVPTARSCVNWNQYYTNCTQLGNNPFQGTISFDNIGLAWVAIFLVISLEGWTDIMYYVQDAHSFWDWIYFVLLIVIGSFFMINLCLVVIATQFSETKKREMERMRQERARFTSTSTLASSTNNSEPTTCYAEIVKYIAHLYRRLKRRIIKKLRQYKYQYQHRKEGLLPQTPETLTLSPNKIKVHHPKCPKMHALAVSTVHRNNQGSNTPCAIIPRNAPSAVGTEKQMSSPDISEMVSVENIKNNALNNCTSYSNLDDKQKIVLLKIPNSENNQSSSNSLSPTSSGRRRSSVMFNEYVIHHTPPTIPAPTTDKNVYCLEKMTQTSDSGIWQVSLPQSIQTMATVFNDYSDLCMTDAMTCQELLAFSVAFSAALPTGHTTLESFYSTLTRNRTNQSNNKNASITNIQGYSGDNVQQKSPPNVAKNLSQNVSNINNINNEEFACCYDLYQNVALEEPPKKHSKPYRCITGTYRFFKRVLHCIRVYIKKLVEHKYFQQGILLAILINTLSMGIEYHNQPEELTAIVETSNIVFSGIFAVEMVLKVIAEGPFGYVANGFNVFDGVIVILSVVELGQAYLGEGQGSSGLSVLRTFRLLRILKLVRFMPNLRRQLFVMLRTMDNVAIFFSLLILFIFIFSILGMYLFGGKFCKFVDETGQERECTCPEIVSKHPLCECDRKHFNNILWATVTVFQILTQEDWNVVLFNGMEKTSHWAALYFVTLMTFGNYVLFNLLVAILVEGFSSERNERREREQRELVKAKLNAESLAAQEQGSEIYDDPKSFSESTTSDSYNESKGKWFSAEELRKIRDIKCNIQKQRLLQPGYDDINLSIPKLNSEKDALNSIDLYSNRKKKCEPLKLYNIQVTDPPIITTTAATPQDSPNATLEGGASFKDWENLDFEQLEKSSSSSLLRPPPILGSLKALDDRTYFEGMPVLSEMRRKSDKGTGSQGDSKSADRMQQSTQDGSRKSETTRKPSLKSRKDAPREGEVLNNGKCVSRSGGITPLDARKIDPLSQRSNRIQNDTQRDRRNSSRRSSVRRSSSIKIDSGPGSIASNISAPSYSKYIYNDKNKYYIDRKSSLRLSDIKSPNNARRMSSIEHTYHKHSRVSVHNLELRKIQEELTKNKLNNASVTSVQDVLPDAPDTRIKNKSKGRLKQFFRYVTPKNFVEERDLYTLYIFPEDNRFRQICSWFVNQKWFDNVILLFIALNCITLAMERPNIPPTCTERYFLSTANYVFTVVFAVEMFIKVVATGMFYGRDAYFTSGWNIMDGSLVIISIVDLLMSLISESSPRIFGILRVFRLLRSLRPLRVINRAPGLKLVVQTLLSSLRPIGNIVLICCTFFIIFGILGVQLFKGTFYYCEGENIKGVKNKTECLSIEGNVWVNRKYNFDDLGKALMSLFVLSSRDGWVNIMYTGLDAVGVDQQPVVNYNEWRLLYFIAFILLVGFFVLNMFVGVVVENFHRCREEQEKEEKIRRAAKRALQMEKKRRRMHEPPYYTNYSPLRMFVHNVVTSKYFDLAIAAVIGLNVVTMAMEYYMMPRALEYALKIFNYFFTAVFILEAIMKLVALGLKIYMKDKWNQLDVAIVILSIVGIVLEELETNIIPINPTIIRVMRVLRIARVLKLLKMAKGIRALLDTVMQALPQVGNLGLLFFLLFFIFAALGVELFGRLECSEEVPCQGLGEHAHFANFGMAFLTLFRVATGDNWNGIMKDTLRDDCDDAADCVKNCCVSTIIAPIFFVIFVLMAQFVLVNVVVAVLMKHLEESHKQMEDELDIETELEREFEREQEFEEEQALCMQLNDDQKQLQKRPLTKVSSLPSNFTYSTPVAEKKLNIHRRQTIQYINQNLNVQHSYTNSNSYDETAENDIPNEDDESFNSVNESSSTKNTNKKIHINKSFFSKNNEDKRASLDEFSKPHKIVEESGKPDKPSSTKGSNKWKDNQSDQPSKEAESNTKLSTIISSNKSLTKSNKPDYRQLSLDQDAKHDNVTLAPNTSSSNFSNASTALNNLGGNKSSFLSVPKLQPKSRSGSTKQLFKQVALDEDNETNESSLLLPSVSNAAGSGGDHSYDLLKVQDESYDAVRKSESCEILRIISERRKI

>XP_018025902.1|Hyalella_azteca_Cav3

MITWFERVSMMAILLNCVTLGMYQPCHDEECVTTRCKTLQIFDDLIFAFFAVEMLIKVMAMGFHGKGTYLAESWNRLDFFIVVAGAVEYCLSVENMNLSAIRTIRVLRPLRAINRIPSFNLQEIPYPPTGEISILSVISLEGWVDIMYYVQDGHSFWDWIYFVLLIVVSMDVLLIVVSMDVLLIVTKKREMERMKQERARYHSSSTLASGSDQPGCYAEILKYIAHLARKFKRKVYARYRIYRRKRNQGSKHLSLKKQRKKRHQFELTDVAEEREMVEAIPTLTIINNTGSSTLDDQLEARQASEEPRRSNTNLMPNLLPLPSSDGSCTPSTSRTRRSSVMFNDVVQMHGGEGDTKNVCLSEKTTQTDPEPPEPLVSRESPKPIRYLPRGRGGSGLTCGELLALSGALSAALPTHMGVDSRSVHTLYSSLAKGVKHFSAPSLFFSNDPSLPYDDSDDSSRTDSDWSESEEEWSEETMRNRPIRRWCSAGRKKVQQLVNSNYFQRWILCAILVNTLSMGVEYHNQPEELTQIVETSNMIFSGVFAFEMFLKIISEGPFGYISNGFNLFDGVIVVLGIVEMCQTYIVQSSSDLLADAAGSGSSLSVLRTFRLLRILKLVRFMPQLRRQLFVMLRTMDNVAIFFSLLVLFIFIFSVLGMYLFGGKFCMRPDGVTECSCKEILAPNSYCRCHRMHFNSLLWATVTVFQILTQEDWNVVLFNGMEKTSHWASLYFVALMTFGNYVLFNLLVAILVEGFSAERHERAERDQKKEERRLRRLQKEQERLEQQQQLQLQQEQTEVQAAPENDQSSPANVVNPPVAPKTKKKRSSSSDQNPIKDVAKIEAPLVANIAIEQEEKKLSTASLESDEERRQLKTNQERVKENQLRQKLILGGAAPETKCNIEKEHQLLQLPSQLAASPPECVALPPPVQMPRPLITHTIATPQGSPHTSLDSGSRDLNNRLSPHLLPPAHAQGLLRSSSVRSNASHRSRHTRGRDSLSPHDLPRRYSLSPSLCYRERSHSGERSSSRRSFHRTSSRSRKRQNAASLSCPTSSSTPTSHHHHHHHHVHSYPPNNSPNYHNIYHNHHHQPHHHRHNHPTSTSLSPTSSFRQSRRLSASKPTSASALAVTTSVTSGAIVKVTVSGERTAVGNGERDAEDGGRHVEEETLNNNVTYNNNNLNNNQQIRRAILTQQNSIGSPRSPLSPQPSIRSAASRQPSLRGRTSVHGSIPERDFRPNNLSDVIKLEESDSGDSDSPDAEATSVTANEVYTYKIFGYFEPKGCFLERVDYSLYIFSETNWIRRVCKYLVSKKYFDSTVLFFIGLNCITLAMERPNIPPDSTERAFLSTCNDVFTVVFGLEMLIKVIAQGLLYGKDSYFSSGWNIMDGILVIVSVIDVLMSLIARSSPRIFGILRVFRLLRALRPLRVINRAPGLKLVVQTLLSSLQPIGNIVLICCTFFIIFGILGVQLFKGAFFYCDGPGLDGVETKADCLKDKRNQWVNRKYNFDNLGHALMSLFVLSSKDGWVNIMYTGLDAVGVDRQPKENYSQWRLLYFISFLLLVGFFVLNMFVGVVVENFHRCREEQEKEEKARRAAKRAKKLEKKRKKMREPPYYAGYSKGRLFVHNLVTSKYFDLAIAAVIGLNVVTMAMEFYKMPKIVARAIKCPRWNQLDVVIVLLSVAGIVLEEMKSEIIPINPTIIRVMRVLRIARVLKLLKMAKGIRALLDTVMQALPQVGNLGLLFFLLFFIFAALGVELFGRLECDDKHPCQGLGEHAHFKNFGIAFLTLFRVATGDNWNGIMKDTIRDECSDESDCLKNCCLYPFVAPIFFVIFVLMAQFVLVNVVVAVLMKHLEESHKLMEDEYNLEVEIERQLALEEEEINELQETLANHKLSRVSTTAKIINEVAQNRSLAKGFGLTTKGGVSSIMLKGGIGIRGLGAHGLAGQFFHKPLGKMSSLPDNFIFRGHRGSDPGIPRIREPSPIVTISELQAKVPSININGCNGDPTSKDAKEFPSHLKSTHNDPNLKKPSPEDPEHKISITDLRTSLRSSKPPPPGENVRVASPTSSPANSSPRSMSPIVLSSIATATVSFINSPRSERKVPPSPLVAAASVSPPSSPSLSLSQISPRSTTVLGPITRFTFGDSSSGTSIDAQTTGYSPESPRPPTDRKPSSPSKQRDSPSPSKIDNQSKAGEMSRGNASPLSQRAFKQLRWCTSSRKSAGAFIVKRKISREDSMVATLMTSLDKVPELGGHLDMDSTSCSLCSSGDDDLPSLPRGQSPSLFPRATPSSNTPSDSDAAPKKAPTNIQIHRDDDDDTLGSNSDYTIEGELDDQEDFLPSPSDPSYKEHPELQLGSGDVPSTNNLYSSELPTPCDNSSNCDASTNFATFRIPTFSDASFLQLSVQSSDNTSTLTPPPGLDAIDEGVKNVGSKLERALQRSASSPYGEEPRASCSSHYSGSKSESDSMTNRKSKISKKAPTSIAEFCSLREMPQSSSGALESQVKRDKCQSSSRKSSVENPTTESCASPEVRTDVKDEESSLPES

>AAO83843.2|Lymnaea_stagnalis_Cav3

MAEESSGAGYMPVPGAGYMPVPVIVEPVDDDNDDDDEVRTHVTGGQASSADQTTPEQSEPAEGHVDEMTPLRQGSHDGVRLLKRGQDHGDNLEKDVSDSAKVSGQQGLDDPGQSQLEHPTDQQVDHQVDQQNDQQNDQQNDQQNDQQNDGEMDEDLLFPGFVPKTFYIFTQRNYIRFWCLRCITWPWFERISMFVIILNCVTLGMYQPCNDKECVTLRCRILEGFDHFIFAFFAVEMIIKMIAMGVIGKETYLADSWNRLDCFIVVAGLAEYIVNKTIEYAVDTENLSLSAIRTIRVLRPLRAINRIPSMRILVMLLLDTLPMLGNVLLLCFFVFFIFGIIGVQLWSGVLRHRCYMDLNASYGIPSYVSEFYKQKNKQEYICSPMNESGIRKCEDLPPYEYEGILCNASAKMDSNNTPTNESCVNWNQYYTNCIPKAPNPFEGAVSFDNIGLAWVAIFQVISLESWVIIMYHVQDAHSFWDWIYFVALIVIGSFFMINLCLVVIATQFSETKKRETERMMQERKRFQSCSTLASNSEPGGCYSELLKLVAQVYRRVKRKVIKTYYKTRGLQKINPEKSLSLRRKKSKKKGTNNLRSQQQYPSVVRQLSLSHQPPVHHHQHQHLLPLPSPLPPQHLHHINLVTPDIRYQSPPRPSSSPQAPRASPEQSDIDSMSSPRRPNYLVLPSSNYSLNPSSESLAMSHLSMDPFTPVLFKSQQNSPNHLTTNFGFAFPLLLSRASSFNSGACGPGKIMPSLPEVLAAQGAKNAVLAASNMLLNVDYEPSKTQSLADKGLFADNLLMDIVGDKQLTTQLSMQSDTDVLQLGTGQTQADQEASSCGGGGVIGAGDKELAGEHGTLSYNPTAIIIAGGDTLDSELETSKDESGVSMRSKMLNCLQKKLKTFVESNFFQRSILIAILLNTLSMGVEFHNQPDLLTTILEYSNVVFCVLFGTEMAFKISAYGLFGYISNGFNVFDGFIVILSIVELAQGGASGLSVLRTFRLLRILKLVRFMPALRRQLVVMLRTMDNVATFFALLVLFMFIFSILGMSLFGGTFCETEEKKPCSCKDRLNASLCSCDRANFDNLLWSLVTVFQVLTQEDWNTVLYNGMAKTSNWASLYFVALMTFGNYVLFNLLVAILVEGFSTEDEEKKKEKMKELEDVDKEDEEEEEEKEKQRLAENNNIDESQSAKSCLNSGDLDVKNNKEKEKKMLALPPISGTSIKTDSPPQIQGQPPSDGPLNPPIITHTAATPMATPQGSPNENAIKDTNNKLSVSVKRSALKSTLSVDSDKSPNFRGSSPRPSPCLRRTNSRGGHHQRTASWRARNRRLRGDKTSLVVDSMSDSADDVDDDVFSSSHSQSPTYSAGTPRTPSVNTHGHAVTYVFSDCNGHPPLNNQRALSPQSSLKGRALTPRNSFKSHHTWSRNNSVGSGRSICNSFELSRQNSFTSHRTLNSLGSGNSKDDNKSNKNYVDLPDVKIALDKEEGREDDIDEPDPDAIDETEWNCSWCPEPKGCFLERHEYAFYLLSNENRIRKFALHLISRKWFDNAVLFFIALNCITLAMERPDIPPDSIERYFLTYTNYIFTFVFALEMMIKVIGKGFFVGKHAYLKSGWNVMDGFLVIISLIDILISMSASSSPRIFGILRVFRLLRTLRPLRVISRAPGLKLVVQTLLSSLRPIGNIVLICCTFFIIFGILGVQLFKGTFYHCKGPNILQITNRSQCEADKKNSWINQKYNFDNLGQALMALFVLASKDGWVQIMYTGLDAVGIDQQPIENYNEWRLIYFISFLLLVAFFVLNMFVGVVVENFHKCRESQEIEERAKRAAKRQEKLDKKRKKMREPPYWAHYSHSRLLIHTVINSKYFDLAIAAVIGLNVITMAMEYYMMPEELEFALKIFNFFFTSVFILEAVMKIIALGFYRYIRDRWNQLDIMIVILSIVGIVLEEMRTNVIPINPTIIRVMRVLRIARVLKLLKMAKGIRALLDTVIQALPQVGNLGLLFFLLFFIFAALGVELFGRLDCIDHPCEGLGVHAHFRNFGMAFLTLFRVATGDNWNGIMKDTLREDCDRADNCLKNCCVSPLIAPVYFVVFVLMAQFVLVNVVVAVLMKHLEETYKYKELDDELEDELRAEMCALALAGKAGPDGIEIKESHRMMDDDSILLRDALITITKANDDPESKESGVEDDNPGGLQVSTLPSFTFSVRPPSEEQSRSEDATGMSPVNASLTLENMGSSWSSMDSNRLAVTPSMTGSSPSSERSVRISSPPIRRAHSPYLLNLQGSSGSIHRRPLVKQYSMDVGSDPQLNNMNTSMPDVWDKFKEAHKSVCKTDNGISRSQPSLKDLKSGSLDGKTEGSLDSKDGALPGEHLAARDRIKHKLLRSRSSGAKYSRFRDKYKQSNLNKAVHLNNSETNVSKERMKRSNQTGSNETGSDIDESLCSDWGSEEAACDLEVKMITGGTEDEEDDIHPTLSPYICEDGDGGHLAACRTAVDSAGSSGLSTPRGESTPRAMKPSYPKCKSPMPPPLQSSRNNPGIDNTSFELGEDVRLPPCCKARETAAVRRSNSFSSSKLIPKAFSPPRTISPRSDRSLSPYTFREHTTSVSPLALPPSPLALEDSAYKLRCSDGGMREMAVSPLPFSHSHMLTVPSPHSLYSNPHMRSPSPTPPKHVHLVPPSSDCPFSSRRQPKGSPAPIRRQQTLSRSSAFDQDNSPAVNSPLRHRIPHTLSMDSAFLIPPGISQSYSINEPYMELHAGCRASSPLPSPRNSQQTLPGMFTRSASPQAHHIQSPSHQPQQPSQLCHHYQSPLSDTVPSISATVSIVQTSPEAASTCLSIPVPAPNAKDVTALTGEELEPLACSSPIQDSDGKELLPLTDDNTNQDSNSRAQPPADPDDKL

>XP_013096433.1|Biomphalaria_glabrata_Cav3

MFVIILNCVTLGMYQPCNDECDTFRCKLLENFDHFIFAFFAVEMGIKMIAMGVAGKDTYLADSWNRLDCFIVVAGLAEYIVNKTIEYSVNTENLSLSAIRTVRVLRPLRAINRIPSMRILVMLLLDTLPMLGNVLLLCFFVFFIFGIIGVQLWSGVLRRRCFLELPENVSVPAFVPLYYIKSPKIEYICSPFNESGISKCQDFPPFHYEGIACNATARNDSDNVPVNGSSCVNWNQYYTSCEERGENPYQGAVSFDNIGLAWVAIFQVISLESWVNIMYFVQDAHSFWDWIYFVALIVIGSFFMINLCLVVIATQFSETKKRETERMLQERKRFQSCSTLASNSEPAGCYTELLKLIAQMYRRVKRKLIKTYYKTRGMQKINPEKSLSLRRKKSKKKGTNIQKSLQQRFSIVHPHTSSHPATVHHHQHMHLVPSHSPQPQHPVHHISLVTPDIRYQSPPRPSSTPQAPRASPEQSDIDSMSSPRRPNYLVLPSSNYSLNPSSESLAVSHLSIDPFTPVLFKSQQNSPNHLTANIGFALPLWFSRASSYSSGASGPGKILPSLPEVLAAQGAKNAVLAASNMLLNVDLEPSKTQSIADKGLFADNLLMDIVGDKHLTTQMSMQSDPEVLQLGSGQSQVDQDIASCAGSGVLCAGDKEVSGPHSTSSFNPTAIIIAGGDVADSELDISKDDSGVSFRSSNLCELLNGMQKRLKYFVESNFFQRSILVAILLNTLSMGVEYHNQPEELTIVLEYSNIVFCVMFGTEMTFKIFAYGLFGYISNGFNVFDGFIVILSIVELAQPGENGASGLSVLRTFRLLRILKLVRFMPALRRQLVVMLRTMDNVATFFALLVLFMFIFSILGMNLFGGSFCEMEDGTACSCKERCNASLCKCDRANFDNLLWSLVTVFQVLTQEDWNTVLYNGMAKTSTWASLYFVALMTFGNYVLFNLLVAILVEGFSTEDEEKKKEKIKETDAADKDEEDEEEKEKQRLAENNNIDDVQSTKSCLNSVDLDVKNNKEKEKKMLALPPSSAPIIKTESPPPNHGFTSSDGPLNPPLITHTAATPMATPQGSPNENAIKDTNNKLSVSVKRSALKSTLSVDSDKSPSFRGGSPRPSPCLRRTNSRGSHHQRSASWRARNRRLRGDKTSLVVDSLSDSADDVEDDVFSSSHSQSPTYSAGTPRTPSVNTHGHAVTYVFSDCNGHPPLNNQRALSPQSSLKGRALTPRNSFKSHHTWSRNNSVGSGRSICNSFELSRQNSFTSHRTVNSLGSGNSKDDKSNKNYVDLPDVKIALDKEEGRENEADEADPDAIDETEWNCSWCPEPRGCFLERHEYAFYLLSNENSLRKLAHHLISRKWFDNTVLIFIALNCITLAMERPDIPPESVERHFLIYTNYVFTFVFTIEMLIKVTAKGFFIGKHAYFKSGWNVMDGFLVIISLIDIFISITASSSPKIFGILRVFRLLRTLRPLRVISRAPGLKLVVQTLLSSLRPIGNIVLICCTFFIIFGILGVQLFKGTFYYCRGLNVTHVTNRNQCLEDKKNEWVNQKYNFDNLGQALMALFVLASKDGWVQIMYTGLDAVGVDKQPIENYNEWRLIYFISFLLLVAFFVLNMFVGVVVENFHKCRESQEIEERAKRAAKRQEKLDKKRKKMREPPYWANYSHSRLLIHTVINSKYFDLAIAAVIGLNVITMAMEYHTMPEELTFALKIFNFFFTSVFILESVMKIIALGFLRYIKDRWNQLDILIVILSVVGIVLEEMRTTFIPINPTIIRVMRVLRIARVLKLLKMAKGIRALLDTVIQALPQVGNLGLLFFLLFFIFAALGVELFGRLDCDREPCEGLGKHAHFKNFGMAFLTLFRVATGDNWNGIMKDTLREKCDRSDNCLKDCCVSPLIAPVYFVVFVLMAQFVLVNVVVAVLMKHLEETYKYKELDDELEDELRAEMCALALAGKAGPDGIEIKESHRMMDDDTILLRDALITITKAADNPDSKESGLEEDNLGGLNVSTLPSFTFSVRPPSEEQSRSEDAATGVSPVNASLTIEGMGSSWSSMDSNRLAVTPSMTGSSPSSERSVRISSPPIKRAHSPYLLNLQGSSGSLQRRPLVKQYSMDVGSDPQLNNMNSSMPDVWDKFSDTQKAVSKPESGIFRSQPSLKDTKMGSLEGKSGGSLDLKEGVLPGECLPARDRIKHKLLRSRSSGAKYSRFRDKYKQSNLNKVVHQASSDVASKELLKRSNLTGSNETGSDVDESLCSDWGSEEAACDLEVKMITGGAEDDDDDIHPTLSPYLSEDRDSSQLATCRTVVDSAASSNMSTPRGDSRVIKAPYPKCKSPLPPTIQSPRNNPGIDNTSFEFGEDVRIPPCCKARETVAVKRSNSFSSSKLVPKAFSPPRTISPRSDRSLSPYSFREHTTSVSPLALPPSPLALEDSAYKLRNIDGGVSPLPFAHSHMLTVPSPHSLYPNPHIRSPSPTPPKYVHLLPPSADCSYGGNRQSKGSPAPVRRQQTFSRSSAVDQESFIPAAMSPLRRRIPHTLSMDSAFLIPPGISQSYSINDPYVELHAGCRATSPLPSPRNSQQTLPGMFTRSASPQPPQQPSRSCHHLLTQQDGSVLYPSVSVISTSPDSDQNNVSLIVPAGNANSIDSTASDKRETNFSSTVLHNDDNKQALDDEGTMKDSQNEDKYVDKSL

>XP_011439125.1|Crassostrea_gigas_Cav3

MADEAVSDAAISSPEVAAVGGDKQTTATTEPEKKVTMARLNFSEECGGDGEEEEGEGSDSDSPEDEEELLFPGFVAKSLRCLPQTNKLRFICLKIITWTWFERISMAVILLNCITLGMYQPCSDLECTTTRCKILEKFDHFIFAFFAAEMIIKIIAMGLYGKYTYLDDSWNRLDCFIVIAGAIEYGVDSENLSLSAIRTIRVLRPLRAINRIPSMRILVMLLLDTLPMLWNVLLLCFFVFFIFGIIGVQLWAGVLRNRCFLSINENISYSYANLSPYYVPKKIGSMPPGDFICSKESESGMRYCSGLPRFEYEGIKCNASARLYSNNTPTNDSCVNWNQYYTNCTAGDKNPFQGGVSFDNIGLAWVAIFQVISLESWVNIMYYVQDAHSFWDWIYFVALIVIGSFFMINLCLVVIATQFSETKKRETERMMQERKRFQSSSTLASNSEPGGCYAEILKYLAHLWRRAKRKVLKAYYNARGKTHKNRKVKPELSLRRKRHKGTSSSCSHKRPQSRSSCPSCRYNYYYQLISANENLYAQPDEFHRNSVRNLRRNSSPLAPRASPEMSDLESISSPRRSNFLTVPNSLHPTSFDSLHSLSVSSTSDNLSSSLPKYRGSSNHLAAPTSHASISRASSFNSSSRRHLPSLPESLTMHQVGCPSHGSCHKLAANSLLHVDSGSRVRIRHASTRSEIPEDLLDAINDKQIQNLVNSIEKRGSGPRCVHVSHDVPDSDDSESKDSDDRPKTICGMLAACKKNFQQVMKTLVEHRFFQRGILTAILINTLSMSVEYHNQPRELTEAVEYSNMVFSILFAVEMFFKLCAYGIIGYIQDGFNVFDGLIVILSMIEFTQQGGASGLSVLRTFRLLRILKLVRFMPALRRQLVVMLRTMDNVATFFALLVLFIFIFSVLGMNLFGGKFCWRGNGTLCTCLDALDPAMDCQCERANFNSLLWSVVTVFQVLTQEDWNTVLYNGMETTSSWASLYFIALMTLGNYVLFNLLVAILVEGFSTEEEEKRKMKLGNKDREHQSLLSETNNNEEKSENIKLKSLDLVQGKINNTERAKLLEVASQTPTAKDDNAKTSPPVTQSDLNPPIITHTAATPMPTPQGSPNDHGPKECNKVTLCVHPPVEMSTLSVRSSSTNSSIRASPRLSPRLSPRISPRPSPCLSPCLRRSLSSGSRSNSWKRRRSQDGDRKCLVPECKIDSMDTDDDDVFQPNCPTPNRSDCNGHLPLNNQRTLSPQNSIKRPVPSPRNSFKIRNNSCSSSQSNCNSIILSRQNSFNSRRTLNSLSSMSAVLKEEVKQNNLMEGTTSPTQVEVGMGAEEEEEDPDAIDETNCTCSWCPEPHGCFKTRHEYALYLLHPNNRLRRVCHHLMAQRWFDNTVLFFIALNCITLAMERPNIPPDSVEREFLNYSNYVFTFVFSVEMMIKVLAKGLVIGQHAYLKSGWNVMDGFLVGISLVDILISLSADSSPRIFGILRVFRLLRTLRPLRVISRAPGLKLVVQTLLSSLRPIGNIVIICCTFFIIFGILGVQLFKGTFYYCKGPTVRNIKNKTQCLMDNKKNVWINQKYNFDDLGQALMALFVLASKDGWVSIMYTGLDAVGIDQQPIENYNEWRLIYFISFLLLVGFFVLNMFVGVVVENFHKCRQDQEKEERERRTAKRRYKMEQKKMKLNRPPYWAEYSPSRLIIHQVVNSKYFDLAIAGVIGLNVITMAMEYYMMPEELEFALKIFNYFFTSVFIIEATLKILALGFMRYIKDRWNQLDIFIVILSIVGIILEEMKTNVIPINPTIIRVMRVLRIARVLKLLKMAKGIRALLDTVIQALPQVGNLGLLFFLLFFIFAALGVELFGRLECSEAHPCEGLGEHAHFRDFAMAFLTLFRVATGDNWNGIMKDTLRDDCDASEKCLTNCCVSSFIAPVFFVVFVLMAQFVLVNVVVAVLMKHLEETYKHKEIDDELEAAFKMECSVNNEMLRLAVKESHRFLDDDGGNAEENKMDEAGDGGDEEDGNSVKKEEEQMLPSIVGTPRPPFSRRLRTPVCKQISLPPSFTFRPPSNEPPYEDDQGRLVPVDITISNYGSSMEVSLDMSGSSLDAPPSPYSSSYSDNSSYVDTTIDENTLGIPRIAEEPRRKSTSPQQKQRLRDRKLVRQKSCEYESPPATAHVEPLQTQNISECKAAACDNPVSLSQPTLNKNLYDGEEGSKVNKSHNPIIKLKCLHRSQSSGSEKVRLREKHIAGARVQLRPTTLHALVTALSNYSHEQSTLALPSTQVTHNSPVICKKKLSQQRRCEGHHGYSLNPSLPRSNVSSKHSLPQVQRNSTCESDSYELSGPDWAAEEEACDEEVRLITGDSTDHDDIDELECEDEVTESCPLVTERCTDPHLDHSRTENSSVSQDTLVGDDRPSRTPRKSEVCGSVKYERVPSPPSSSSPALTSVPSQDPQIVVLEDPRSVDAKTRKISLNESQNSDTDLSGGECPYDSEVMEPMSENSGEATLAAISGSACLSVPGMTLDDSCVRRRTPNLFQPSSES

>CCD68017.1|Caenorhabditis_elegans_Cav3

MLRQPVPELRSFQSLSKYAGGPRSVLGRRTSAITVNRRQSQSTRRHEDVEALGSIEGSKETLQLSEHGRLASSSEASPSRWEGRQIEWGNEEQIEEESELPYPGFAEPALRCFYQARPPRKWALQMVMSPWFDRITMAVIMINCVTLGMYRPCEDGPDCDTYRCQILDIIDNCIFVYFAFEMVIKIMALGFYGPAAYMSDTWNRLDFFIVMAGIAEFVLHEYLGGNINLTAIRTVRVLRPLRAVNRIPSMRILVNLLLDTLPMLGNVLLLCFFVFFIFGIVGVQLWAGLLRNRCVINLPKTISENQSALFNNVKLTRFYIPEDTSLEYICSQPDANGLHTCSNLPPYTVDGVKCNLTLDEYDKVTNDSCINWNIYYNECQVMQRNPFQGSVSFDNIGFAWVAIFLVISLEGWTDIMYYVQDAHSFWNWIYFVLLIVIGAFFMINLCLVVIATQFAETKRRETERMLQERKMLLNRDSISCTGSEIGGASSKEEGDTVYAAFVRFIGHTFRRTKRAAKKKYTAYMEERAERKSSERQQRRKSKLDDMATLSRIEEKAEDEEDETTITRENGDDQIEQNGDGVRIKRVKIEEEPKIKIGNGNSNGPHYKHSSSDEESDEDGEEDQVYDGEEAKKKSTPSKLWWFREKIQKFVICDHFTRGILVAILVNTLSMGVEYHQQPEILTVILEYSNLFFTALFALEMLLKIIASGLFGYLADGFNLFDGGIVALSVLELFQEGKGGLSVLRTFRLLRILKLVRFMPALRYQLVVMLRTMDNVTVFFGLLVLFIFIFSILGMNLFGCKFCKVEEKFLGGLAKKCERKNFDTLLWALITVFQILTQEDWNMVLFNGMAQTNPWAALYFVALMTFGNYVLFNLLVAILVEGFQESKEEEKRQLEEDARKQAVEEEDERKRELELIIAKTTSPAFNNGVAPAECTCQRPSSPEESPSPRLLSANYHPSPERKHSANLDAIIDKRLVLRNSAPFDRSPVSEGRDDSRLNRHASLVLPVANGVPYRRQRVHSWSGLCHHFNPNCPVHGRRALIETYAREKFLEASQELKQALAEEEKRNEAKQNTFVRKLLKKTCLHNRTEFSLFLMGPKNPLRIKCLQTTQKKWFDYTVLFFIGINCITLAMERPSIPPDSFERQFLHISGYIFTVIFTGEMMMKVIANGCFIGQAAYFKDGWNILDGILVVISLINIAFELLATGDSPKIFGVIRVLRLLRALRPLRVINRAPGVKLVVMTLISSLKPIGNIVLICCTFFIIFGILGVQLFKGMMYHCIGPEVGNVTTKADCIEDYRNKWVNHRYNFDNLGQALMSLFVLSSKDGWVSIMYQGIDAVGVDVQPIENYNEWRMIYFISFLLLVGFFVLNMFVGVVVENFHKCKEALEKEMREKEKEKRLKRKLKRQKFEESMAGKRKKMERNYPYYHDYGHTRLFLHGIVTSKYFDLAIAAVIGINVISMAMEFYMMPMGLKYVLKALNYFFTAVFTLEAAMKLIALGFKRFFIEKWNRLDMFIVILSIAGIIFEEFEALELPINPTIIRVMRVLRIARVLKLLKMAKGIRSLLDTVGEALPQVGNLGSLFFLLFFIFAALGVELFGKLECSEDHPCDGLGEHAHFKNFGMAFLTLFRIATGDNWNGIMKDALRDDCDSSDHCETNCCVDPILAPCFFVIFVLISQFVLVNVVVAVLMKHLEESNKRDAEGPAEPTGENIENEITKSDDDEIVEEHEPLAIEHVKEGELDEEEETEEGPTTQIPDGHGGIKRLSMQVLEQELIEVERHLEERYRRASECLGGELQPLNPGEIEDLDDPEFRPRSRSHRPRARTNSALSNKSRGSHKSAL

>PIC19332.1|Caenorhabditis_nigoni_Cav3

MLRQPVPELRSFQSLSKYAGGPRSVLGRRTSAITVNRRQSQSTRRYEDVEALGSIEGSKETLQLSEHGRLASSSEASPSRWEGRQIEWGNEEQIEEEDSDLPYPGFAEPALRCFYQARPPRKWALQMVMSPWFDRITMAVILINCVTLGMYRPCEDGPDCDTYRCQILDIIDNCIFVYFAIEMVIKIMALGFCGPAAYMSDTWNRLDFFIVMAGIAEFVLHEYLGGNINLTAIRTVRVLRPLRAVNRIPSMRILVNLLLDTLPMLGNVLLLCFFVFFIFGIVGVQLWAGLLRNRCVINLPKSISENQSALFNNVKLTRFYIPEDTSLEYICSQPDANGLHTCSNLPPYTVNGIKCNLTIDDYDLVTNDSCINWNIYYNECQVMQRNPFQGSVSFDNIGFAWVAIFLVISLEGWTDIMYYVQDAHSFWNWIYFVLLIVIGAFFMINLCLVVIATQFAETKRRETERMLQERKRLQNRDSISCTGSEMGGASTKEEGGDTVYAAIVRFIGHTFRRTKRAAKKKYNAYMEQRAERKSTERMQRRKSKLDDMATLSRIEEKAEDEEDEIATPPEIKEEPLEHNGDGVRKKRVKIEEEPRVKIGNGNSHGAHYKHSDNEDESDESQSCDESYDGEKAEKKRKPSKIRWFRDKVRKFVVCDHFTRGILVAILVNTLSMGVEYHQQPEILTVILEYSNLFFTALFALEMLLKIIASGFFGYLADGFNLFDGGIVALSVLELFQEGKGGLSVLRTFRLLRILKLVRFMPALRYQLVVMLRTMDNVTVFFGLLVLFIFIFSILGMNLFGCKFCKVEEKFLGGLAKKCERKNFDTLLWALITVFQILTQEDWNMVLFNGMAQTNPWAALYFVALMTFGNYVLFNLLVAILVEGFQESKEEEKRQLEEEARKQAVEEEDERKRELELLIAKTTSPAFNNGVPPAECTCQRPSSPEGSPMLLSANYHLSPERKHSVNLDAIIDKRLVLRNTITPFDRSPVSEGRADDSRLNRHASLVLPVANGVPYRRQRVHSWSGLCHHFNPNCPVHGRRALIETYAREKFLEASQELKQALAEEERRNEAKQNTFVRKLLKKTCLHNRTEFSLFLMGPKNPLRIKCLQTTQKKWFDYTILFFIGINCITLAMERPSIPPDSFERRFLQVSGYIFTVIFTGEMMMKVIANGCFIGQSAYFKDGWNILDGILVVISLINVAFELLATGDSPKIFGVIRVLRLLRALRPLRVINRAPGVKLVVMTLISSLKPIGNIVLICCTFFIIFGILGVQLFKGMMYHCIGPEVGNVTTKVDCLKDTRNKWVNHRYNFDNLGQALMSLFVLSSKDGWVSIMYQGIDAVGVDVQPIENYNEWRMIYFISFLLLVGFFVLNMFVGVVVENFHKCKEALEREMREKEKEKRLKRKLKRQKFEESMAGKRKKMERNYPYYHDYGHTRLFLHGIVTSKYFDLAIAAVIGINVISMAMEFYMMPMGLKYVLKALNYFFTAVFTLEAAMKLIALGFKRFFIEKWNRLDMFIVILSIAGIIFEEFEALELPINPTIIRVMRVLRIARVLKLLKMAKGIRSLLDTVGEALPQVGNLGSLFFLLFFIFAALGVELFGKLECSEDHPCDGLGEHAHFKNFGMAFLTLFRIATGDNWNGIMKDALRDDCDSSDHCETNCCVDPILAPCFFVIFVLISQFVLVNVVVAVLMKHLEESNKRDAEGPAEPTGENIENEITKSDDDEIAEEPEVLAIENGEDKEEENAEDEETEPSTQIPDGHGGIKRLSMQVLEQELIEVERNMEERYRRASECLGGELQPLNPGEIEELEDSEFRPRSRSQRPRARTNSALSNKSRGSHKSAL

>PAV61762.1|Diploscapter_pachys_Cav3

MLKHPVSELRSFQSLSRYAGGPRSVLGRRTSAIPPSRRRSSVAKRRDELLEAGGSSEGSKEETLHFPETARAGASSSEVSPVRQVEFADENEQIEVEGEGQLPYDGFADPVLHCFYQSRPPRSWALKLVMSPWFDRITMAVILINCITLGMYRPCDDGPTCSTYRCHILDIVDNCIFVYFTLEMMIKVIALGLVGPTGYLADTWNRLDFFIVIAGIVEVVLADVMGGSINLTAIRTVRVLRPLRAVNRIPSMRILVNLLLDTLPMLGNVLLLCFFVFFIFGIVGVQLWAGLLRSRCIINLPTHIPENQTALFKDVKLTRYYIPEDTSLEYICSQNDANGLHKCDNLPPYTQNGVKCNLTLDQFDQISNTSCINWNIYYNLCQVEERNPFQGSVSFDNIGFAWISIFLVISLEGWTDIMYYVQDAHSFWNWIYFVLLIVIGAFFMINLCLVVIATQFAETKRRETERMLEERKRMARHDSICTGSEQALDSSKVEGGSVYAAIVRMISYTFRRAKRQARRKYKTYMDARKEQRQNAEIYRRKSKLDEMATLSRIEERMEEEEDDEKPNSNAGNQQNAIVLVKKLGNGSVRDYDNDETATDDDINESEFSSDEQEEEKPRSSSKWQWFRFHVKRFVDSDHFSRSILVAILVNTLSMGVEHHQQPEIFTTILEYSNLFFTGLFAFEMLLKVIAYGLFGYLADGFNLFDGGIVALSVLELFQDGKGGLSVLRTFRLLRILKLVRFMPALRVFGMILFGRKFCNHKETGEQCNCLQIGRGECICDRMNYDTFMHATLTILTQEDWNMVLFNGMSQTNPWAALYFVALMTFGNYVLFNLLVAILVEGFQESKEEEKRQQEEEARKHALEEEEQRKAELEALLAKSTSPGFINGAANGACICFDRKHTPLHLAISVVSEPDPFARNANNRHRSNEERQINSDDGVLALIPSSPTPIPLKNSPLSERSLPMSPILKGTDKLHRHTSLTLPTQNGSVGRKPRERTHSWSGVGHIFNENCPIHNKRTLIEIYAREQLAKAGEELKQALAEEERKKELKQNTLCRKLLRKTCLYRRSENSLFIFSPKNPLRVKFITLTQKKWFDYTILLFIGINCVTLAMERPSIPPYSIERKFLDISGYIFTIIFTVEMLMKVIANGCVFGDGAYFKDGWNVLDGILVIISLINVGFEIFVTGDSPKIFGVIRVLRLLRALRPLRVINRAPGVKLVVMTLISSLKPIGNIVLICCTFFVIFGILGVQLFKGMMYHCTGPDIANVTTKEECLQDSRNKWVNHRYNFDNLGQALMSLFVLSSKDGWVSIMYQGIDAVGVDIQPIENYNEWRMIYFISFLLLVGFFVLNMFVGVVVENFHKCKEALEKEMREKAKEKQLQRQLKRQKYEESQAGRKKRAQELARRERPYYASYGRTRLFLHGIVTSKYFDLAIAAVIGINVISMAMEFYMMPVVLKYVLKALNYFFTAVFTLEAIMKLVALGFRRFLKEKWNQLDMFIVILSIAGIIFEEFEALELPINPTIIRVMRVLRIARVLKLLKMAKGIRSLLDTVGEALPQVGNLGSLFFLLFFIFAALGVELFGRLECSEDHPCDGLGEHAHFKNFGMAFLTLFRIATGDNWNGVLKDALRDDCDSSDRCESNCCVDPILAPCFFVVFVLISQFVLVNVVVAVLMKHLEESNKKDAENEEKDLHTDGDLETAASNDILEDVQEPEEQIEIAAGNKRLSLQTLERELAEVEEKVEERMRKESAILGDSNCNADRGDTSDENEYCGRRSEHHHTPCKKKRKAAR

>PDM63609.1|Pristionchus_pacificus_Cav3

MLQQPVAPELRSFQSLSKYSGPRAVVARRQSQIIARRSSQCRKRTETDDEAMEAGTYQLGSRENEANTMPSASPKTSIADQSSIRDGGGSRQVEWGDDEEFEEDFDPNLPYPGFVEPSLFFFKQAKAPRSWALQAVMSPWFDRVTMGVILINCITLGMFRPCEDGADCNTYRCQMLQLADHVIFAYFAFEMCVKVIAMGFTGPAGYLSDTWNRLDFFIVIAGCAEYVLQEYVGNINLTAIRTIRVLRPLRAVNRIPSMRILGKEENERKARGFTLCEGAKINLLLDTLPMLGNVLLLCFFVFFIFGIIGVQLWAGLLRNRCTLNLPNTTTGIPEDFFLDVSLSRFYIPEDTSMEYICSQADAGGIHTCHDLPPFSKNGIRCNLTIEEWERADNVSCVNWNRYYDKCAVIHKNPFQGSVSFDNIGFAWVAIFLVISLEGWTDIMYYVQDAHSFWNWIYFVLLIVIGAFFMINLCLVVIATQFAETKRRETERMLQERKRLRSAGSISGSEAGAAASSKDGGGDSVYAAMVRFIGQTFRRTKRRVKKEWKLYKIRRRDRQEARVTKKEQKNNNRQHSLPAIEERPDEENPSSENNFLAIPESTSKLRRNSAKKANGKVRLSVDVGRDSDKSSSDCSSGSETDDECDEETYRLDKIEDSTSRKKRKRNKRGPIGALRDRIRAFVVCDHFTRGILVAILVNTLSMGVEYHQQPLWLTKILDISNYFFTALFAFEMILKVIADGLFGYLADGFNLFDGGIVALSVLELFQEGKGGLSVLRTFRLLRILKLVRFMPALRYQLVVMLRTMDNVTVFFGLLVLFIFIFSILGMNLFGCKFCKEEANPFLPFAPVTKKCERKNFDTLLWALITVFQFSVVSFSIFGMILFGCQFCNHVETGVKCTCEQIKQTPDLCYCDRSNYDNLLHATFTTFQILTQEDWNMVLFNGMSQTTPWAALYFVALMTFGNYVLFNLLVAILVEGFQESKEEEKRQLEEEARKKADEEEEKRKHELEMLIAKTTSPAFMASHPKEAQECTCTKGEHFLRITAACAASMSSRSPSLTPAATETLGSLPHSPVRQPTPSRIVGNNVYQPVPKREDSEETETRSPRMSTNYDSDSSDDSKELLTVQKTYRLNRHKSLFLPAGSRSPLNIQPRQRLHSWGGVHVHFNPNCPVHGRKALIETYAKEKYLQASQELKQAMAEEEKLAEAKNNAMWRRVLRKTCLKGRTDYSLFMFSPKNWLRIKCLQITQKKWFDYTILFFIGINCVTLAMERPSIPPMSPERVFLDISGYCFTVIFSLEMLMKVISNGCIIGEGSYFKDGWNILDGILVIISLVNVLFEFFASGDSPKIFGVIRVLRLLRALRPLRVINRAPGVKLVVMTLISSLKPIGNIVLICCTFFIIFGILGVQLFKGVMYHCVGPDIANVTTKWDCLLDTRNKWVNHRYNFDNLGQALMSLFVLSSKDGWVSIMYQGIDAVGVDIQPIENYNEWRMIYFISFLLLVGFFVLNMFVGVVVENFHKCKEALEKEMREKAREKRIQKQFKRKLKRQQFEESMANKKKKSVKTRPYWEEFGPTRLFLHQVVTSKYFDLAIAAVIGINVISMAMEFYMMPMGLKYVLRALNYFFTAVFTLEAGMKLTALGFKRFFRETWNRLDMFIVILSIAGIIFEEFEALELPINPTIIRVMRVLRIARVLKLLKMAKGIRSLLDTVGEALPQVGNLGSLFFLLFFIFAALGVELFGKLECSEEHPCDGLGEHAHFKNFGMAFLTLFRIATGDNWNGIMKDALRDDCNPSERCETNCCVDPILAPCFFVIFVLISQFVLVNVVVAVLMKHLEESNKRDAAETGAAEQNGAAQSGVDTDAKATDTEAEDVERDTPCPPKDEIMSMENLEKELIEVEERMLLSNRITIDDGSLVFDEESEELLSPVSNIHLSASFKRRRRDSLIIESPSKEYHPMAHQFHSFRTKRDGSTRRKRLTSKASHTSSADEGISPMREGLEEETVDA

>XP_024510580.1|Strongyloides_ratti_Cav3

MLQQPVSKELRSFQSLSKFSGTAQKGSVPHRRQSAATNNSITKSSYGRRRLPDSLNDDSMTIRSVDNTLHRSPSSIQQMPLNVKSFHTKSGIISSHTKSEELFHKDALSGEYNKSNSLTYDASPIHKWDKKEQSSNECHRQPQVEWNDEGEFVFDDECNLPYPGFVEPALHCFSQTKIPRKWCLKMVTNPWFDRLTMIVIIINCITLGMYKPCEDGPGCNTYRCQLLSMIDHMIFIYFFLEMVIKVIALGFTGQAAYLSDTWNRLDFFIVIAGIAEYLLQEYLGNINLTAIRTIRVLRPLRAVNRIPSMRILVNLLLDTLPMLGNVLLLCFFVFFIFGIIGVQLWAGLLRNRCTLNLPKKNISLENLNNILKDVQLSRYYIPEDTSLDYICSQNDASGIHTCNDLPPYVYNGIKCNLTIDEYNKIDNSNCINWNIYYDECTVAPMNPFQNSVSFDNIGFAWVAIFLVISLEGWTDIMYYVQDAHSFWNWIYFVLLIVIGAFFMINLCLVVIATQFAETKRRETERMLQERKRLSSYSNSIYGSEGEKLSQERDENNGDTVYAALVRFISQTARRVKRHFRKQGPVYKTRILRFFGRKTIEIPKKNDNKGISESDDLLEKKDKNSDNNNSICALQIDITKEDNDIKSQKSESFNSNKRKNKENGKIKKKNSNISKFTEIRNIVKKFVICDHFTRGILVAILLNTLLMGVEYHQQPEWLTIILEYSNYFFTGLFAFEMLLKVFADGLFGYLADGFNLFDGGIVALSVLEIFQEGKGGLSVLRTFRLLRILKLVRFMPALRYQLVVMLRTMDNVTVFFGLLALFIFIFSILGMNLFGCKFCKIDEGLSGPDSKKCERKNFDSLLWALITVFQILTQEDWNMVLFNGMAQTTPWAALYFVALMTFGNYVLFNLLVAILVEGFQESKEEEKRQLEEEAKKRAEEEEKERNKELELLIAKTTSLSFMGKSNTKTATCTCSKDILNKESSQLTNENTLKIPDSRPRAYSSSIESCKSFNNIKLEHDCNGDIKNDDELFINNDETNKKFDFCKNRNIDKKKFSKSQENLIERSSSDIENSCKKLSKKDKMYDSAKDVYSEKSIIHFQNNSNEIKDNNSYLENSRLRTNSWCGIQTLFNPQCPIHSRRALIEAYARDKLIQASQELQQVLVEEERKAEERKNTFCRKLFTKTCLYKRKDYSLFLFSTKNKLRISCLKLTQKKWFDYTVLVFIGINCITLAMERPSIPPDSLERKFLTIIGYIFTIIFTLEMSLKVIANGCLFGRGAYFKDGWNILDGILVIISLINIIFEMLVHIDSPKIFGVIRVLRLLRALRPLRVINRAPGVKLVVMTLISSLKPIGNIVLICCTFFIIFGILGVQLFKGMMYHCIGPDVSNITTKNECLAVNGNKWVNHRYNFDNLGQALMSLFVLSSKDGWVSIMYQGIDAVGVDMQPIENHNEWRMIYFISFLLLVGFFVLNMFVGVVVENFHKCKEALEAEMREKARQKRLARKLKRQQYEDHCALKKKQKEKSYPYWYNYGPARMYTHNIVTSKYFDLAIAAVIGINVISMAMEFYMMPSGLRYVLKALNYFFTAVFTLEAAMKLYALGLKTFFMEKWNRLDMFIVILSIAGIIFEEFEALELPINPTIIRVMRVLRIARVLKLLKMAKGIRSLLDTVGEALPQVGNLGSLFFLLFFIFAALGVELFGKLECSDDHPCDGLGEHAHFKNFGMAFLTLFRIATGDNWNGIMKDALRDDCDPSDHCDTNCCVDPILAPCFFVVFVLISQFVLVNVVVAVLMKHLEESNKKNEITESEVVNSGIEVDVNKIDVNIGDENDFEHKPLTVVALEQQIIDLEQKMFEDGRLEITANSNATNDKTTNNDCIEKNIK

>KRY57796.1|Trichinella_britovi_Cav3

LVKKLKMWEGPLPAELRSFRSLSRVSRASIRRKSRSSVAAPPSPTGKFGESSSPGHNVDFGRKRSASTATSDAPEHNNDHIQHIDDVEDVPYPAYVTTALRCLDQRTPIRYWCLRIVNNSWFEKISMMVILINCVTLGMYKPCDEDMELAVCNTTRCITLSVIDHLVFAFFAVEMIIKIIAMGFYGPDTYMSDTWNRLDFFIVIAGCAEYIFKERMGDINLTAIRTVRVLRPLRAINRIPSMRILVNLLLDTLPMLGNVLLLCFFVFFIFGIIGVQLWAGLLRNRCLLDLPSSNDSSLELMGLVRYYMPDETSMEYICSLEKDNGMHSCNNLQPSTYNGHICNLTIDQWHAGNYTLSNSSCINWNQYYTKCGPTSHNPFQGSVSFDNIGFAWVSIYLVISLEGWSDIMYYVQDAHSFWDWIYFVLLIVIGAFFMINLCLVVIATQFADTKRRETERMLAERARYPSTSTLTTIDQQSDSDGVYRAIVKYIAHLGRRAKRKAIRYYREWKKRRANVAEIPSPHLITNSPKPAISQEADPSQICLVPKRFTYNSPRGDSLTIPCNSPTGLHSRGSSVSYYSTLEDSGLTIPAADYLPYSPARRYSSHRRSRLVRSARTSAGSADINRNGRKIKHHRLASSKRYDPRFALNYTESAPVCRPSFTGIRLRIHREASTSSNSLSLDSEDEQTALYLDDESSNRDEMDLELRKPYTDTDSELDHDRVRFHTDQLHLRNKRQNSSSDLLVRCLAFLKVCIKRFVDSDHFTRGILVAILINTLSMGVEYHNQPEELTIILEYSNVFFTALFSIEMALKIIADGPFTYVSNGFNLFDGGIVILSIMELLQGGNGGLSVLRTFRLLRILKLVRFMPALRYQLVVMLRTMDNVTVFFGLLCLFIFIFSLLGMELFAGRFCQMPDGTPCHCKSKIPACKCPRRNFDTIINAAFTVFQILTQEDWNVVLFNGMAQTSPWSALYFIALMTFGNYVLFNLLVAILVEGFQESKAEERRLMQEEIDKTEIDPSTKAKENMFEVYKSENEKGEKSAITCTCAAGVAAATLLSSRSRDRLLSLPIMPVYVSGSELVNAPPPIIQTLPTPVGSPVMSSPSARNCRKSGSKFDQNNELSNGNSLYLPNSNAMSNSTISVKSHPLYKAGSHRDGQSKRKLTGQQSITYLDIPNTKSGSCLRRHWSSSSSTTTFYSCVNTNTESVATSSYKSLESSQQEPVSVSTDASTNNQCNGSVMMQSEVEFSPRKVSSQLSQQIYSSVNPHIFYPYCKVHGYRLSLIGDGQKKEALLQNSPLFIHDELKAQLVARFHQWFQQSWFQTRMDASLFIFMPKNKFRAQCVYLSQQKWFDFCILTFIGINCITLAMERPGIPPNSLERLFLDISGYIFTVIFALEMFIKVVAKSLVLGDGAYFKNGWDFMDGTLVLISLSNLIFDLLVKSHAPKIFSVVRVLRLLRALRPLRVINRAPGVKLVVQTLISSLQPIGNIVLICCTFFIIFGILGVQLFKGKMWHCVGPNVAKVINKTDCLADPHNRWINHRYNFDNLGQALMSLFVVSSKDGWVSIMYQGIDAVGVDMQPVVNYFEWRMLYFISFLLLVGFFVLNMFVGVVVENFHRCKEALEKEMKEKEREKKMRKRLQKQLSRQQLLIKRSKKLPYWYHYGPVRMYLHGIVTSKYFDLAISAVIGVNVITMAMEFHMMPPELTYALKVFNYFFTAIFTLEAILKVFALSIPRYLKDRWNQLDVIIVLLSIGGIVLEEMESNILPINPTIMRVMRMLRIARVLKLLKMAKGIRSLLDTVMQALPQVGNLGLLFFLLFFIFAALGVELFGRLECSDEHPCDGLGEHAHFKNFGMAFLTLFRIATGDNWNGIMKDTLRDDCDNSDNCTSNCCVSAIIAPVYFVVFVLMAQFVLVNVVVAVLMKHLEESSKKMADGSSEAGGSNATDTDEKSCTLRGDSDDERALHAISSTTSLPTPQDLSKIELPDNPVKFWIPDQSITQSDTLKKRNDKIDFDHQLDGLIENIV

>KRZ43346.1|Trichinella_pseudospiralis_Cav3

LVEKLKMWEGPLPAELRSFRSLSKVSRASIRRKSRSSVAAPPSPTGKFGESSPGHNVDFGRKRSASTAASDVPEHNNDHIQHIDDVEDVPYPAYVTTALRCLDQRTPVRYWCLRIVNNPWFEKLSMMVILINCVTLGMYKPCDEDMELAICNTTRCITLSVIDHLVFAFFAVEMIIKIIAMGFYGPDTYMSDTWNRLDFFIVIAGCAEYIFKERMGDINLTAIRTVRVLRPLRAINRIPSMRILVNLLLDTLPMLGNVLLLCFFVFFIFGIIGVQLWAGLLRNRCLLDLPSSNDSSLELMGLVRYYMPDETSMEYICSLEKDNGMHSCNNLQPSTYNGHICNLTIDQWHAGNYTLSNSSCINWNQYYTKCGPTSHNPFQGSVSFDNIGFAWVSIYLVISLEGWSDIMYYVQDAHSFWDWIYFVLLIVIGAFFMINLCLVVIATQFADTKRRETERMLAERARYPSTSTLTTLDQQSDSDGVYRAIVKYIAHLGRRAKRKAIRYYREWKKRRANVSEIPSPHLITNSPKPAVSQEADPSQICLVPKRSTYSSPPGDSLTIPCNSPTGFHSRGSSVSYYSTLEDPGLTIPAADYLPYSPARRYSSHRRSRLVRSARTSAGSADINKNGRKIKHHRLTSSKRYDPRFALNYTESAPVCPPSVTGIRLRIHREASTSSNSLSLDSEDEQTALYLDDESSNRDEMDLELRKPYTDMDSEFDHDRVRFYTDQLHLRNKRQNSSSDLLVKCFAFLKVGIKRFVDSDHFTRGILVAILINTLSMGVEYHNQPEELTVILEYSNVFFTALFSIEMALKIIADGPFTYVSNGFNLFDGGIVILSIVELLQGGNGGLSVLRTFRLLRILKLVRFMPALRYQLVVMLRTMDNVTVFFGLLCLFIFIFSILGMNLFGCKFCKQEMNADGELVKSCDRKNFDSLFWATLTVFQSIRQNALFKKTILWIAFSICFFNAFILLGMELFAGRFCQMPDGTPCHCKSKIPACKCPRRNFDTIINAAFTVFQILTQEDWNVVLFNGMAQTSPWSALYFIALMTFGNYVLFNLLVAILVEGFQESKAEERRLMQEEMDKTEIDPSTKAKENMFEVYKSENEKGEKSAITCTCAAGVAAATLLSSRSRDRLLSLPIMPVYVSGSELVNAPPPIIQTLPTPVGSPVMSSPSARNCRKFGSKFDENDDHSNGNSLYLPNSNAMSNSTISVKSHPLYKAGFHRDRQIKRKLTGQQSITYLDIPNPKSGSCLRRHWSSSSSTTTFYSCMNTNTESVATSSYKSLESSQQEPVSVSTDASTNNQCNGSVMMQSEVEFSPRKVSSQLSQQICSSVNPHIFYPYCRIHGYRLSLIGDGQKKEALLQSSPLFIHDEINQLNAALRQRKKSLKAQLVARFHQWFQHSWFQTRMDASLFIFMPKNKFRAQCVYLSQQKWFDFCILTFIGINCITLAMERPGIPPNSFERLFLDISGYIFTVIFALEMFIKVIAKSLVFGDGAYFKNGWDFMDGTLVLISLSNLIFDLLVKSHAPKIFSVVRVLRLLRALRPLRVINRAPGVKLVVQTLISSLQPIGNIVLICCTFFIIFGILGVQLFKGKMWHCVGPNVAKVVNKTDCLADPHNRWINHRYNFDNLGQALMSLFVVSSKDGWVSIMYQGIDAVGVDMQPIVNYFEWRMLYFISFLLLVGFFVLNMFVGVVVENFHRCKEALEKEMKEKEREKKMRKRLQKQLSRQQLLIKRSKKLPYWYHYGPVRMYLHGIVTSKYFDLAISAVIGVNVITMAMEFHMMPPELTYALKVFNYFFTAIFTLEAILKVFALSIPRYLKDRWNQLDVIIVLLSIGGIVLEEMESNILPINPTIMRVMRMLRIARVLKLLKMAKGIRSLLDTVMQALPQVGNLGLLFFLLFFIFAALGVELFGRLECSDEHPCDGLGEHAHFKNFGMAFLTLFRIATGDNWNGIMKDTLRDDCDNSENCTSNCCVSAIIAPVYFVVFVLMAQFVLVNVVVAVLMKHLEESSKKMADGSSEAGGSNATDTDEKSCTLRGDSDDERALHAVSSTTSLPTPQDLSKIELPDNPAKFWIPDQSITQSDTLKKRNDEIDFDRQLDEMDSQCIKPCKNDKRRL

>XP_022091428.1|Acanthaster_planci_Cav3_

MAEEDREAQRLPRVRGGSVRIRAPGEQSMGRRSSPPPSRSPSSSQGDMGDHDAEQPSPTGEGSVGFASRDGVDEDGMSTGSCEEELPFPGLNDKVFYCLSQTSKPRVWCIQLVCWPWFERISMAVIILNCITLGMYEPCEKECTSTRCVVLEGFDHFIFAFFAAEMVVKILALGVRGKSGYFFETWNRLDCFIVVAGITDYTMQALEYSLQLENMNLTAIRTIRVLRPLRAINRIPSLRILVMLLLDTLPMLGNVLMLCFFVFFIFGIIGVQLWKGLLRNRCFLDLPDNTSIGAGYDVTTFYTLPDMYQDYVCSFKTDAGDLHCDSDHKDPFLIGEQECNSTALPYINNSYDSNLTDCVNWNQYYTMCKTSDVNPFLGSISFDNILYAWVAIFQVITLESWVEIQYYLQDVHSQWVWIYFMLLIVIGAFFLINLCLVVIATQFSETKQRESKLMAEQRKRFRSTSTLASNSEPGSCYDEILKYIGHLVRKVKRKVRRWHKGIRGKRQRKVMPAISLRRKKRRTKSVHVHCHHHHHHHHYHFGSNSPQAPRASPENSDIGSSPTRQNRLVVPSANGSLGPSVDSLHSITYCTENLSRLEPLPSRSRCKSSPHHHVVTTLSIHRASSINYPTTNSKDARPSSALAKLTAETALNSMMDVPPNKLQERWKDVCLGETLNKYSKLPPVKNGNKISGDGLAPAGAHPPLAQRLSAPPLTHPPSSYPGGHPHLAPPSPCLSIHSAGCPVHQPCPVHTPCRPSTPCPSHSSCPTHSPCPSRGSHTCPHACPEGHNDYDWTESDLESDEDSDADDSDYEEEQKGNLKFGGSGCKSFYRKVSDKIGEIVESKYFMRSILICILVNTLSMGIEFHNQPDELTEALEISNRIFTSLFALEMLLKLMAYGFVGYIRNGFNVFDGIIVIVSVVEIVQQGGGGLSVLRTFRLLRILKLVRFMPALRRQLLIMLKTMDNVATFFSLLSLFIFIFSILGMHLFGCNFCRYVDGVKVCDRKNFDNLLWALVTVFQILTQEDWNIVLYNGMHNNSAWAALYFIALMTFGNYVLFNLLVAILVEGFAAEPYEQKESSLAAASRESLEEDEGYSGEEHEKKEEGNDVQSVTSGKDTSDEKQLALPPPESTQSPPPASLMSPPIITRTAATPQGSPMCEQPKGFKGPFLDNQSIDSDTQSLSSFHIPGSPRLPRSNLSRNGSGRNRNVLSANLPHVRDRNYLRPFQPPEADHGADSDSHSNISSRISSRRSSSDHNGYVPDSDSRRTSVASCNGDSRRTSYISCNGGSFTAQLEKHSLEAEKRSSRGSVGRRSSRHEDEESIEDPDQVDEAKKQSAELGRKCFCLPGDCNCTKCCPEPKGCFKTRIEYSLYLLSPHNRFRRRLQSLIAHRWFDYVILLIIMINCITLAMERPDIRDDSVERTFLSISNYIFTGIFTFEMVVKVLAKGLFIGEHAYLYSGWNIMDGSLVVISWIDITISLVSSNSPEIFGILRVFRLLRTLRPLRVISRAPGLKLVVQTLLSSLRPIGNIVIICCTFFVIFGILGIQLFKGTFYYCTGPSVKNVMNKTHCLQAHPNNQWVNQQYNFDNLGQALMALFVLASKDGWVEIMYNGIDAVGVDKQPKENNNEWLILFFISFILIVGFFVLNMFVGVVVENFHKCREQQAAEETARRQAKRLRKMEKARQSMSRVREGDEIISNPKKRQARARERPYYIDYSRSRRFIHNCVINKYFDLGVAAVIFINVISMALEHYGMSKTLQDILRYLNYFFTVVFILEAVLKIAALGFKRYIKDRWNQLDMIIIILSIVGIFLEELQTDIIPINPTIIRIMRVLRIARVLKIMKTMKGLRELLAVLMGAIPQVGNLGLLFFLLFFIFAALGVELFGRLDCTKENPCQGLGRHASFKNFFIAFLTLFRIATADNWNGIMKDTLRQEKCNKSSDCTYNCCASSILAPIYFVCFVLMAQFVLVNVVVAVLMKHLEESHKMEQDEEDDQLALEEEMREEGREAARREEEAVEQQQQGQEAAEQANSSDDNDPETPLTPLLKPLQSSDIVIAMPSSQDSVRVERETNLDMGKSLVPPTSLQLSPYKVPVDRVQSVPGSFHSQKDSAVEVPEDGGEPSQQTASTERKQQLPCQVPHIQLPSDNEHSSSTSSSSHHLAGDRPRISSVSLESEIGSGQQSPASRSSFFTASPRSDSQAKLDSDPDRTGPSSKDAARHLSLPTGPAHPDRDMSSDLAFRTGEAESNAKPEEVPLQQWPRKSSDSSTGDAAKQPPSKSFVDDVVRQKSRDLDPRAHPRVRRGARPAGPSNLNNLASSNKPHQAHRRASKSSVATSSSDSDSSSPPNRKRTRRRQRHSPSKTLPQKREDGPSLPSSREPSVSGEIPEIQVKQPKEILQDCELQNDGTDSGSHPGKTDQDCTPNKPEKTKGNYSCDPAEKV

>NVE5017(FigShare)|Nematostella_vectensis_Cav3a

MLCKWLAMGVFGKQGYLAENWNKLDCFIVAAGTFELCYDQGKYMTAVRAIRVLRPLRAINRVPSIRILVTLLLDTLPMLGNVLAMCSLIFSIFGIVGVQMWQGVLRSRCVAQLPGNLSANQLNISAFYQPGGSSDLVCSLPENSGDKHCTSAFIPASTAHGRECGLTYEQYSNRSNYTNVCANWNQYYTLCADMGINPLYDAVSFDNILIAWVAIFQVITLEGWTDIMYYIQDAHGFWNFIYFVVLIVIGSYFMTNLCLVVITTQFQETKQRENDLMRQSRRKHTTSTSTIGSSRYGREGCWLEILKWIGHVYRHTKRRLGKKLNIKCAEPVSKRTRVTRKKKRRKKKLVYHHHHHHHHHHHYHHHVHCPGNCQYNTVGTFCPIHDVSLPPTPNASNADFNSGASTSEPPVSGVSGNSLAVPALVRATVHNGSEEAEPKAPELPGAVPSVSIVTGSACSVRPAASPLIGEMLTTATATVSVNGQPAIADVTAHSFVSAASLTGAAATATASAAATAVIQNACACAVGQHLDKVEKFEAEYVYEEGEESETDLSDDEDTSESSPRRCAKFRDGCASSVDSKWFMYVIMGAIFVNTLTMGIEYYGQPQKMTDVLEIFNYIFTAIFGIEMIMKLIGLGFYGYIKDAFNIFDGTIVIISVVELFGDDDSGISVLRSFRLLRVFKLVRFLPALRRQLLVMIHTMDNVMTFLALLVIFMFTASILGMNLFGGKYRFPNDEGVMETSRANFDDLFWAIVTVFQVLTQEDWNIVMYDGMRATSKWAGLYFILLMTIGNYILFNLLVAILVEGFASAPARGESTASLRPTPCPRTEYQLAEINYKERLESQRLKCHSKPKVCISPTPSVMYAEQAGYDSEGNYDYPPKSPRRSSLPELFTKKCPIPELDLSSVPVTPCMSPTVTEKPNPFCPPAGRTQVFVNSRTPSENDMEKTGEDEAVTDNPTVDIKVPTSCCSKRADWSLYVFAPDNRFRMLNKELYQNKWFDRTVLLFILLNCVVMALEGPSVLPGSLERRVIDICMYVFLGIFTIEMMVKVIALGLWIGPDAYLRSGWNVMDGFLVVISWVDVIVTATTNGQNSILGVLRVFRALRTLRPLRLVCIQFSSTEMYKLVIALGLWIGPDAYLRSGWNVMDGFLVVISWVDVIVTATTNGQNSILGVLRVFRALRTLRPLRVISRAPGLKIVVETLISSLKPIGNIVLIAATFFIIFGILGVQLFKGKFHYCTDATVPVTTKTECLENGGSWVNREYNFDNLAKALLTLFVFSTKDGWVNIMYDGIDAVGIDKQPIRNHARYNVIYFVGFLLLAGFVVLNMLVGVVVENFQKCRTLIEMEKKIDEEKKNKKEEKMRREIAEDEALSEHYPRPRKFIHAVCTHGYFDLGIAAVIALNVLCMALEHYQQPDGLTAFLKCANYVFTAVFILEAILKIFALGIKRYIKDRWNQLDMLIVILSIVGIALEEMTMELPINPTIIRVMRVLRIARVLKLLKTAEGIRKLLDTVAQALPQVGNLGMLFLLMFFIFSALGIELFGKIDCEKPGITCQGMDEHANFRSFGIAMLTLFRISTGDNWNGILKDTISPDVCRVNPEIDCSMLEHVAPIYFAVFVLATQFVLLNVVVAVLMKHLEEAKDTKTPVNTPPLSSSSRESSKDNNGASLKVPSSDRKVSFDDEADNYSKPESRSSRCLPMVAVNGQGYNSFDNEFGLQGITRQDAIKPPAMLSRSAPSLRPAITDRRRPTSAKPSIESSRMVGSTGSLTRLSPLIGRRTKSQGKHQEEAGGPELTFTPSWDSGLKPDSIGGQVEPRTMSPRAPIYTMSEDSDLYDSNLDTDGGSDVRAFSPRPSLKSTSGSSESEDAQYRHKMPPALRSGAELAWGTPDAQEFDSNEGRPSSSKSAPVGAEVGENVQLQNMRPATATEDKSHRMQSYV

>XP_020900273.1|Exaiptasia_pallida_Cav3a

MLERKTTDDSTTTDNVIELGKLKQENTNQCEEKLEIPGCAEDDGSEEDLLESFKPVAFYFLKRXKYPRLLFVKLVSWSYFERISIFVILCNCVTLGLYDPFDPDCLTQRCQILEKIERAIYAFFVVEMLCKWIAMGIFGKLGYLSDNWNKLDCFIVAAGTFELFYDKGKYMTAVRAIRVLRPLRAINRVPSIRILVTLLLDTLPMLGNVLAMCSLIFSIFGIVGVQMWQGLLRNRCMLELPDNISPQRYNLSSYYTPSKGGDLVCSLPKNNGMKRCTKEWLDPYRIDGRECLLDYRSYLNITSQSSNTNSCVNWNQYYSKCGNAGENPDHDTISFDNILIAWVAIFQVITLEGWSDIMYYIQDANSPWDFIYFVVLIVIGSYFMTNLCLVVITTQFQETKQRENDLMRQSRRKHAASTSTIASSRFGRDGCWVEILKYIAHVFRRTKRRWGRTLNIKCCRPATHKTRVSKKRKRRKKKLVYHHHHHHHHYHHFHHHVHCPGHCQHAEGYCPVPDGRYTPNTSNADLNSSLCQQAEIPVGNTLTVPGQIPVNGTLQGEGRDSNGYAVDLPGTVPNVSIVTGSACSIRPSSASKVAGEMXTTATAAVNINGQSASSDVAAHSFVSAASLTSAAATATASIAATSVIQNACECAVGHASGDKGVVAEEFESESVYEGEESETDYSDDEMNKDKQKMTAFSKFRHALRDWVEGKMFMYFIMGAIFINTLSMGIEFYGQPQEMTDVLEILNYIFTGIFALEMLVKLIALGLYGYIKDAFNLFDGAIVIVSIVELFGDGNNNISVLRSFRLLRIFKIVRFLPALKRQLLVMIHTLDNVVTFLALLGIFIFTASILGMNLFGGKYSFPNDKGVMVKSRANFDDLFWALVTVFQILTQEDWNVVMVDGMRATGKWAALYFILLMTIGNYILFNLLVAILVEGFANQPDKNGSTATIRQSAVVLSKTEYNQLAKRTEYPMDQSATKFYGGSKPRVCISPTPSIFYVDQLQCTAYSGCVGENPKSPRRSSMPNLPKNRYADVTPSRSCLSPSPSRKIGFFLNDRNEVFMHSLKRSDTQEQKTGMEIVASNKKQGSSATATSSSSCCAKRVDWSLYIFSPENSFRKWNVALYKNKWFDRIVLVFILLNCVVMAMENPSVKDDSTERQAIDICMYIFLGIFTLEMLIKIIAMGFWVGEGAYLRSGWNVMDGFLVVISWVDVIVTLSTDKENRILGVLRVFRALRTLRPLRVISRAPGIKIVVETLISSLKPIGNIVLIAATFFMIFGILGVQLFKGKFYYCDGISLEVDTRQQCEANGGSWKNKEYNFDNLAKALLTLFVFATKDGWVSIMHDGIDAVGIDKQPKENYARVNVLYFVAFLLLAGFVVLNMLVGVVVENFKKCRDLIEMERDEEEKQREKDEKKKEKECREGDEASTHYSRHRRFIYHTVTHAYFDLGIAVTIGLNVICMALEHYDQPKGLGDFLQSANYVFTAIFILEAILKIYALGVKRYFSDRWNHLDLVIVILSIAGIILEEMNNDLPINPTIIRVMRVLRIARVLKLLKTAEGIRKLLDTVLQALPQVGNLGMLFLLLFFIFAALGIELFGGIDCKNLECEGMNEHAHFQRFDIAMLTLFRISTGDNWNGILKDTINKKMCESNPEVDCNVLEHVAPIYFAMFVLATQFVLLNVVVAVLMKHLEDAKDDDSSSSGARESITSGERDSNYKDFNGGRGXVSLDVPQMEREVKSSSDDDTENKQITTAEINGIQNGGNNVPSVAINGEDLNSDSEYSTTSPLPXVKSKGRKINMLSRSVPTLRPALRTRNRSSSAAPRIAKEVSRSGSLVKLSPILGRLSKSNRYDDSDELVSHVSPVYEFTAMTNEASEHDVSPEREKPPLSPRAPLYKMSQDSDLYDSALEGDHGGSTGSVRRIPKLSQASRPSLXSTQSTSTSSSGDDERFHLPKRFKGKHTAWGSPXVEDKQXKSESSKQSSSNEPQVPKSAPLASDKRKNVDTVELTHRSLSTGSMPEKKMKKHYKEQSYV

>XP_015765864.1|Acropora_digitifera_Cav3a

MIADNKSNNNLSSEEAFEQIQNNPRLSCKLSSDAGACYDNKIEQEDGGGQNVFKDFQPVACFFLRSESPPRSWFIRLVTWPYFERVSIFVILLNCVTLGLYDPFDPECSSQRCQTLDTMEKIIYAFFLVEMLCKWMAMGLFGKMSYFADPWNRLDCFIVAAGTFELLYDKGEYLSAVRAIRVLRPLRAINRVPSIRILVTLLLDTLPMLWNVLAICFFIFAIFGIVAVQLWQGALRGRCFMSDSNHLLMQKLKSPGILCACEICINEFFNYNVSDMFVCELPNVPDMQKCTSEFISPFENEMGLQCSLTLEEYVNKSRLSNNSNECIDWNQYYTQCKQEGDNPAWGAIGFDNIFIAWVAIFQVITLEGWADIMYFVQDAHGFWNWIYFVILIVIASYFLTNLCLVVITTQFQETKQRENELLRASRRREGTSTSTIASSRYGKDGCWVEILKYIEHVCRRLKRRFHKRFKWKYCEESNKGTKTSKRRKRRKRTKKLVYHHHHHHHHYHHYHHHVHGTDPGCLSGNVHPGATSLTPDASCVDVQQLQDKQDQPNRNTLTVPGQVPSNDQQDSAEKSGAIPTISIVAGSSCSLRRDPICSSQGEMITTATAAVSLNGKSAVADVTAKTVLSAASLSSAAATATASTAAASVIHNACACAVERVEEDLKMNYESECVYEESDDSDFTDDEETADQTQNNGDRDKLFARFRRYCRRSVDSKWFMYIIMGSIFLNTLSMGIEYHGQSSIVELFGEGDSSISVLRSFRLLRIFKLVRFLPALRRQLLVMIHTMDNVVTFLALLALFIFTASVLGMNLFGGKYTFEDEDGGKVTARANFDDLFWALVTVFQVLTQEDWNTVMYDGMRATTKWAALYFILLMTIGNYILFNLLVAILVEGFANQPERTASTWSIKSKQSKQDQDHVTGVVMSPHKKPSLTIEPPLVCVSPSHSSPDMDLACTHDGKTIDGETTEPMERMRCSFPSLSHRCPAFRDVSFNSYAFLVRSPKDGELLGLEWGMILQISYTIFKCQMSIFPRVGVLTQEDWNTVMYDGMRATTKWAALYFILLMTIGNYILFNLLVAILVEGFANQPVPLTCNFYCFRGSLALFVLLVVVSFSSLRQCLERFRKLMIATYSNKWFDRVVLVFILLNCVVMALERPDLPKDSELQKIIDICMYIFLGIFTLEMFIKVMALGLWVGPGAYLRSSWNVMDGFLVVVSWIDVIVTLTTDTESSILGVLRVFRALRTLRPLRVISRAPGLKIVVETLISSLKPIGNIVLIAATFFIIFGILGVQLFKGKFYHCKGASDVETRAECTSTDGSWVNKEYNFDNLARALLTLFVFSTKDGWVTIMYDGLDSVGVDKQPKRNNNKWNVLYFVAFLLLAGFVVLNMLVGVVVENFQKCRDIIEKDRQVEKEKEKEQERARKRQEGEMMTKLAEEFLQPRRFFHRICTHGYFDLGISAVIVLNVICMAMEHYNQPQEMRDFLKYANYVFTAVFVMEGILKIFALGFKKYIKERWNQLDLIIILLSIVGIVLEEMDASLPINPTIIRVMRVLRIARVLKLLKTAEGIRKLLDTVAEALPQVGNLGLLFLLMFFIFAALGMELFGQINCDDVPCEGLDNHAHFRNFGFAMLTLFRVSTGDNWNGILKDIINKERCEHNPPPGCAALEHIAPIYFAFFVLVTQFVLLNVVVAVLMKHLEDAKEEIPVTSSRAGELENIVTTRTGEEDRQDGQGNQGVGDPVSGPNKKATCVIPVSADLHRQRPNDSESSLSSFEYTVKSQPTSASPRHRLPPVSKTQSSCCTGQATQLRLPPIGSQPRDLNEINGGSEKIDSFTSELPEKSPESARSESKSTDGHVSPRAPIYVMSEDSTEIYDSNMEVDREPSKVFRPVENRQAPRTPSPHSSSEEEGKKKFFKKPFRKSETKRNSSKIEPEAAVGVPDTKPAKQGKQEVKLPKHDAQWASPVTSSNVQQQKGQEIQLNPVTKPAKDEEQDKSYQVQSYV

>XP_020622701.1|Orbicella_faveolata_Cav3a

MQGEEKLNAEVSREDDAEKIRSNPSLVVECKLNRGTEPRYGNEIGNIEGKHNVFQDFMPVAFFFLHREKRPRKWFIRLITWPYFERLSILVILVNCVTLGLYDPFDPECKTQRCQTLDAMEKVIYTFFLAEMLCKWIAMGLFGKLAYFAESWNRLDCFIVAAGTFELLYNQGEYLSAVRAIRVLRPLRAINRVPSIRILVTLLLDTLPMLWNVLAICFFIFAIFGIVAVQLWRGALRGRCFLELPANVSRDSTAVTQRIGKDLSDSFWPAVGFKAFDYISLILFNYFERLSILVILVNCVTLGLYDPFDPECKTQRCQTLDAMEKVIYTFFLAEMLCKWIAMGLFGKLAYFAESWNRLDCFIVAAGTFELLYNQGEYLSAVRAIRVLRPLRAINRVPSIRILVTLLLDTLPMLWNVLAICFFIFAIFGIVAVQLWRGALRGRCFLELPANVSRDSYNLTDFYVPSFDEPDFICAMKGIPGMQSCNSESIPPLKDEISGRTCTLSYEAYLNETRRSNNSNACVNWNQYYSQCRKEGDNPSWGAIGFDNIFIAWVAIFQVITLEGWTDIMYYVQDAHGLWNWIYFVILIVIASYFMTNLCLVVITTQFQETKQRENELLRASRRREGASTSTIASSRYGKDGCWVEILKYIEHLIRRFKRRIHRRFNWKYCDSESKRKKTLRHRKRRRRKKKLVYHHHHHHHHHHHYHHHVHCTDPGCQCGKIGQQFSNVHPGSEQVTPSTSSVDIRRVQSKPVKPAGNTLTVPGQVPNIDKQENSNEAGGIPTISIVTGSSCSVKKEPSSSAPGGEMITTATAAVAINGKSAVADISAHTFLSAASLSSAAATATASAAAASAIHNVCTCAVEHVEEDIKVDYESECVYEEDAGSESEYTDDEVEEGAQRKKTPFSRFRHMCRNSVDSKWFMYIIMGAIFLNTLSMGIEYHGQPLKMTEVLEILNYIFTAIFGFEMLLKLMGLGPYGYIKDPFNLFDGFIVVMSIVELFGGGDSSISVLRSFRLLRIFKLVRFLPALRRQLLVMIHTMDNVMTFLALLVLFIFTASILGMNLFGGKYMFPDEDGVETAARANFDDLFWALVTVFQVLTQEDWNTVMYDGMRATSKWAALYFILLMTIGNYILFNLLVAILVEGFANQPEKSSSSWSVKSRSYSIKHDGDYETCKGMAPYPMVTPRVCVSPTPSHDVTTRDDYKETAEPVHHMRCSFPSLSPKCSLFRDECHASAPASPASTSPNVTRKTLSFGDTKTMVIANPAQAERCITSDDENETSAKEDDEAVVRIQDGSTCCHKRRNWSLFLFSPSNRFRTWMVTVYRNKWFDRVVLVFILLNCIVMALERPDLPSDSTMKKVITICMYIFLAIFTLEMIIKVIALGFWIGRDTYMRSAWNVMDGFLVIVSWVDVIVSLTTDTQSSILGVLRVFRALRTLRPLRVISRAPGLKIVVETLISSLKPIGNIVLIAATFFIIFGILGVQLFKGKFHYCKEAEYPTVITKEDCLHRGYTWENKEYNFDNLAKALLTLFVFSTKDGWVTIMYDGIDAVGIDKQPIRNNNKWNVLFFVAFLLLAGFVVLNMLVGIVVENFQKCRDMIEKDRLAEKQKKKLKNCDQPIRNNNKWNVLFFVAFLLSAGFVVLNMLVESNKEEGGEFPQPRRFFHRICTHGYFDLGISAVIVLNVICMAMEHYNQPEDMSVFLKYANYVFTAIFIVEGVLKIYALRFRKYIKERWNQLDLFIILLSIVGIVLEEMNSEVPINPTIIRVMRVLRIARASIPSYIMVTQPSLVENTNNVNNACQRKGG

>XP_022794522.1|Stylophora_pistillata_Cav3a

MQREEKQNAERPSREDEAEKIRSNPSLVEECKLTRRSEPRYETEIVNVDERYDMFQGFMPTAFFFLHREKRPRRWFIRLVTWPYFERLSILVILINCVTLGLYDPFDPECQTQRCQTLDAMEKVIYTFFLAEMLCKWIAMGLFGKLAYFGDAWNRLDCFIVAAGTFELLYNKGEYLSAVRAIRVLRPLRAINRVPSIRILVTLLLDTLPMLWNVLAICFFIFAIFGIVAVQLWRGALRGRCFLDLPQGVSQNSSNLTDFYIPSFDKPDFVCAVDGVPGMQTCIKEFIPALKNENRECSLNYGGFMNQSRRSNNSNNCINWNQYYTDCRKKGDNPSWGAIGFDNIFIAWVAIFQVITLEGWSDIMYYVQDAHGFWNWIYFVILIVIASYFMTNLCLVVITTQFQETKQRENELLRESRRREGTSTSTIASSRYGKDGCWVEILRYVEHLIRRFKRRLYKRFHWKYCDGGSKRKASRHRKRRRRKKKLVYHHHHHHHHHHHYHHHVHCTDPDCPCSKVAPQFSVHPGSADATPNTSTVDIQPIQSKDVQPSSNTLTVPGQVPSNGTSGRENDNGNGKGKIPTISIITGSSCSVRKEPSSSTRGGEMITTATAAVAINGKSAVADMATHTFLSAASLSSAAATATASAAAASVIHNVCSCAVEHVEEDIKVDYESECVYEEDGGSESEFTDDEVDEEEHGKDTACSRLRQVCRRSVDSKWFMYIIMAAIFLNTLSMGIEYHGQPTKMTHVLEILNHIFTAIFGVEMLLKLVGMGLYGYIKDPFNLFDGFIVIMSIVELFGVGDSNISVLRSFRLLRIFKLVRFLPALRRQLLVMIHTMDNVVTFLALLVLFIFTASILGMNLFGGKYMFRDENGIYSAARANFDDLFWALVTVFQVLTQEDWNTVMYDGMRATTKWAALYFILLMTIGNYILFNLLVAILVEGFANQPEKTGSTWSLKSRSSSSRNNGDYETCREVSSYPMTTPRVCVSRIPSPEDALHGHYHEPDEPVQHIRCSFPSLSPKCSLFRDECHASAPTSPASTSPHVARKTLSFGDTTTMVIANPAHAQRCITSDDENEASAKEDDESVVANESVSRCCPRRRNWSLYLFSPSNRFRQWNIALYKNKWFDRVILFFILLNCIAMALEKPGLKSDDKLKKAIDISMYIFLGIFTLEMMVKVIALGFWVGKDTYMRNSWNVMDGFLVLVSWIDVIVSQTTKTESSILGVLRVFRALRTLRPLRVISRAPGLKIVVETLISSLKPIGNIVLIAATFFIIFGILGVQLFKGKFYHCKDSEHPTVATKNDCQKLGLTWENKEYNFDNLARALLTLFVFSTKDGWVTIMYDGIDAVGVDKQPIPNHNKWNVLFFVAFLLLAGFVVLNMLVGVVVENFQKCRDMIEKDRLAEKDKERAEKLQSEADRDEAEEFPQPRRFFHQICTHGYFDLGISAVIFLNVICMAMEHYRQPEEMTLFLRYANYVFTAIFIVEGVLKIYALGFKKYIKERWNQLDLLIILLSIVGIVLEEMKSKLPINPTIIRVMRVLRIARVLKLLKTAEGIRKLLDTVAEALPQVGNLGLLFLLMFFIFAALGMELFGQVECTDEVPCEGLDHHAHFKDFGFSMLTLFRVSTGDNWNGILKDIINKKRCELNSTSGCAALEHIAPIYFAIFVLATQFVLLNVVVAVLMKHLEDAKEESPTASSEEGSDDVLAIQVDRVSDPKHMQGGSSKDHEYCMVPSGVKGKTESNNNPLVVVPVVGVVRQTNGSDSSLSSIGPVQPEGASSYWLPAIDQAKPEHNKSLASIRLPPIGKSSDKIDDSSLSEMSSPENDKPISKRLDGQISPRAPIYVMSEDSDLNDSNMEVDHDHSKVFRPVETSTKSKGSVRSPSPQSSSEDEGKNTKIHLRKSHYKSESPRGKTRQKQPKTSIVSPKPDGKPVTPGAKPVKPDARWASPVINVEKGKETIKLDSPKHDENKGTDYQMQSYV

>NVE7616|Nematostella_vectensis_Cav3b

MVVKMVAMGVFGSHGYLQDTWNKLDFVIIVMGVIEKSLQGSDYLTIIRAFRVLRPLRAINKVPSIRILVTLLLDTLPMLGNVLLLSFLVFFVFGIIGVQLWQGKLRNRCFASTNDTALLLSFNQSVFYKPSFIQPDFVCSGPTNTGMTSCLSDIPHSYLHGRQCTLPPENLTSNSSNCADWNSLYRDCREDGPNPFWGTISFDNIAIAWMVIFQVITLEGWSDVMYLVQDTHSFWNWIYFIILIVIGAFFLVNLCLVVITMQFQETKAREIELMEENRRRQRYRSSRFSLSCLEKLRRTFCPGRCQGEPKEHIHHHHHHIYHHHHFHHHLYDCYTPSSPGLALSSDEDKFFGQDICQPDTLQIPEITIDEGDRKAFLRPPEQLTLSRRESDTSVCSHSFSLHAEAKSSVSIDEILTTSSVTLTVPGSPSGFIAAAHTSVRATTSLPEDSILKENRASFQTRAQALATAEVNADQRCLRAQGARQRSRNAESKERIRESPTLSGAKIQGTSRNIEAVSVCSPGLSRSYTGSPTISARSYEMTDSEGLMCDVSPSLPRLRVVSCHGSPTMSRNATPTAQSPNLSRSNDQTNESHNESRGESPNQSRHSRRYSQPFGLKEVNWTRKTSEKRQKNRRLDLKELAIEELQDEEPESKFSLRKFSIKNKQKGAKNSLNACTSRDESKVTRARWGSSEDLTQDLKRDLQLSLSSDKGSLDSLSKHADIKHEGRIKSTDNGMPSKVPRDASRDKQKSSDKVTSSKWDMLRDASRDAGDFFLNRNAVKGDMFQDEPLALATTDVAEPRPNPFLLDDDVDDSVFDFHQSDSYDVSRDMGVVHRFRAFCRRMAESRHFSRLIMIAILLNMICMGLEHHNQPQALTVTLEKTNIVFVTIFVLEMIINVISFGIMGYLSQLQNIFDGFVVVLSVTELLLENGYARLSVFRSIRLLRIFKLVRPVRYQLLVVIRTMTSVVTFFGLLFLFMFAFAILGMNLFGGEFYFPNAENISVPARTNFDSFLWAMVTVFQILTQENWNQVMFKGMRATSYWAALYFIALMTVGYYVLFNLLVAILVEGFTSSGCKTSAISQDTAEASRQRHHNQADHLKAPCHRNPLTLVNNEAGSYDRDQHAEPAMDYCGHALRNALLSGASPQTRDSSKKSGKTCNSRKTSLERCLSSGPVGELSFTLPELKQMRQTSISLLRKTRRNPGSSRIPGQPVYNPAFVDDDVIKTQVQMKGFSPRKSSNLDDEDNVDDEKITEKLELETPEGNIPEACKTVFFKPEVKVKEVNKPDIKSSKIIQPKVIEPEVTYEESCEEIPYMAYYTQVGDGMPSVMKGRRQSTLVQPRKKVSVFAVRHDWSFFMFAPDNRFRQRMRAVCSHRAFDYVILVFIIFSCAVLAIEAPDIAEQGLKRQIIDISMLVFTIIFTIEMLIKLVAMGLVLGPGTYLRDGWDVLDGFLVMVSWIDIIVTYTSHVSPEVLGTLRVFRALRTLRPLRVIRRAPGLKLVVQTLLYSLKPIGNTVLIAAIFFVMFGILGVQLFKGKFYYCEGDSHVISKQECANSTRGQWVNRRYNFDDLLQALISLFVVSTKDGWVEIMHHGIDAVDVDVQPIVNYAEWRLVYFIPFLLLGGFLVLNMIVGVVVENFQRCRERMDEEEQARPHRKGKKARNQMQAEDSLYYESYGPLRLKIHFICTHRNWDITIAAIICINVICMSLEHYKMPQSYEVFVETTNYFFTSVFVIEVVVKVIALGFVRYPKDRWNLIDLAIVLLSVTGIVLELLVKVDHLFNPTVIRTLRVLRITRVLKLVKLAKGVRSLLDTLFEALPQVANLGLLFFLLFFIYSCLGIQLFGSLECSHDYPCQGFNRHAHFRDFGTAMLTLFRIATGDNWNGILKDTLRSKCEDSADCSKNCCLSRYTSPLYFVTFVLAAQFVLVNVVIAVLMKHLKESKEKIAATMAAKDIEKKLKLLTLVARNFIRAKAARKSSQPSQDLNRRRSVVELGLGGIELAQLKSHTQKWKSATLYRDVAKADAIFQEYMHLNASLRKAKMKAMRTTSNKIERTSSLVERSTKRPLQRRHTISDLNSFSPQASPSIAKRTVWKSGIEL

>XP_020912298.1|Exaiptasia_pallida_Cav3b

MLPSLPADVTVAVLTDQYTVKKESMEGGHNDMVALEAIRDEASNDQIDVEEISCMFLGVRTLPRRWLLVVYSNRWFERVSMFVILVNCVTLAMDDPYDRQCESLRCQILQSIEHFIFAFFLLELVVKMIAMGVWGKKGYMQEPWNRLDFVIIAVGTVEKLMEGSDYLTIIRAFRVLRPLRAINKVPSIRILVTLLLDTLPMLGNVLLLSFLIFFVFGIIAVQLWQGHLRNRCFIDLPLTYRTNESLFYKPSFDKPDYICSLPQDNGMTSCNHIPPNYANDRTCTQNFRTVHHNHTMINSSDCIGWLYKTCRREGPNPFYGTTSFDNIVIAWIAIFQVITLEGWSEIMYLVQDSHSFWNWIYFVILVVIGAFFLVNLCLVVITMQFQETKAREIELIEESRRRQGINISSMQAYSCFQTLKRYVQEFCCSRQAFKRKQSQIHHHHHHFHHHHHFHHHLYDCYTSSPVLSPEIKSKPYIIEVEDESSTGPCIPAITIEKVDDEISDLQVHPSRSTFLFPKSESAAFSKGISIHTEASSSIFSKEILATSSVSLTVPSGAANAHATASASASLPIIRISQDHGFAVAGTSDSSSADFESRPRSSSDTCGKRSNTEVVTNHSVHQRRHSQPFGDQEIAMSVKNSQICSRKWQDLQELELKKLEIETSKLEKESFKSKYSFDKRNGTSRLSEVPSLQMYNSSVHGTKSGDDTSQEKGQNVSPNNMVDIGIVTDTSDHIGDNNEGYVADTDERKSRGIHTRKDNDIYDSIFDLRQCDSYDVARKNGPCSRLRNFFRRLAESKQFVHFIMISILVNMICMGLEHYKQPERLTAALEQSNIIFVSIFSLEMLINLMAYGFTGYLSQVQNVFDGFVVVVSVAELLVNQDYARLSVFRSIRLLRIFKLVRPVRYQLLVILRTMNSVVTFFGLLFLFMFAFCILGMNLFGGKFFFYDANNEKVLRRSNFDSFLWAMVTVFQILTQENWNHVMYDGMRVTNNWAALYFIALMIVGYYVLFNLLVAILVEGFTNTEDTLIAQSEARRSSIKQGQTSKYSKHTYNLHSDMKPKDSVKKDVFEENCSLSRDSTLDYCGHAVRGAVWSNRQARTKAKNSLGIQKNWAGMNSCKSTSVNKCGKSFLTMPELKQKRKTPTNVWEMTLRLPGRESRDVSDRLSTGDFKWNPSFVPNDNDRSSEIAMNSGEVGQCSLTDXQYCQFIPKHTTSATQESIREFVNNQVSHENKQDLKTIFDSNVVKQQATRDIIAISDPETGVPQATPRYRITGRQGFFSSRREWSLFMLSPDDRFRTTLLTICSHKIFDYVILVFILMSCVVLAMEGPGTDNNATVREIIDISMLLFTVVFTIEMMMKVVAMGFIIGSDSYLKDGWNVLDFILVIISWLDVIITYSPQAASVDVLGTLKVFRALRTLRPLRMIRRAPGLKLVVQTLLYSLKPIGNTVLIAAIFFAMFGILGVQLFKGKFYHCEGDNHVIDRAQCLNSTHGRWVNRRYNFDNLFQALISLFVVSTRDGWVEVMHHGIDAVDIDMQPKVNHAEWCLVYFIPFLLLGGFLVINMIVGVVVENFQRCREQLDEEQQNSKKKRHKSETNYNEQDDSSDYSVHYNVARRWIYHVCMHQYWDIAIAIIICVNVICMSMEHYQMAPTFIIFVETSNYFFTAVFVIEALLKVVALGWLRYIKDRWNIIDLLIVVLSVSGILLDDLVKKELPINPTVIRTLRVLRIVRVLKLVKLAKGVRSLLDTLFEALPQVANLGLLFLLLFFIYSCLGIQLFGSLECSYENPCEGLGQHANFNSFGSSMLTLFRVATGDNWNGILKDTLRSECDASAGCRKNCCISKYTAPLYFVTFVLSAQFVLVNVVIAVLMKHLKESKEKLAATLAAKNLEKRLKLLTVIAKNFIRAKASRKPSPPITRRRSLVELSLGGIELAQLKNHKKKWVNESFVQDVVKADTIFQQFIQLNNSLRSVKNLRAHRTTSNSLEQMSSGWESAKTTKRRHTLGGIWDLEMHQSTPQNKHIVWKSGCSP

>XP_015766817.1|Acropora_digitifera_Cav3b

MFVIFVNCVILAMYNPLDENCTETRCRVLENVEHFVFAFFCAEIVIKMVAMGVTGNRGYLQDKWNRLDCLIVMIGLIEKVISNNNYLTIIRAFRVLRPLRAINKVPSIRILVTLLLDTLPMLGNVLLLSFLIFFVFGIISVQLWQGKLRNRCFTNFPENSTFFTRSTFYKPSFDEPDFVCSLPEDGGTSRCSNISPKDTNCSGSFEDNSINWNKCYNICKEVGPNPFYDLVSFDNIGIAWIVIFQVITLEGWSDIMYFVQDAHSSLSWIYFVVLIVVGSFFLVNLCLVVITMQFQETKAREIELIESTKDEAQPNSSEPLRAVMKFAKNLCWCRTEDEPQVHHHHHHHFYHHHHFHHHMYNCYQPASPGIFHSTSTNEPSFVFPSSDIPDIQVGGDAAMAFPSNAEHCEINPDFLAPPDRVSRRNSRTSICSHTLSIHSETYLLKHRRLSQPAHLIEMNSSPQSANVSSYSPSEMLDQNMSCIDFNQNERNRDQSEIEKVLSCADISISEFPLKTQGSDNQTKVTSSKPKKYPQIQRGARTQSEDDMFDSLFNLRQSASYDVTARRGAWQSLRALCRKVAASKHFTFFIMVVILLNMICMAPEHYDQPEFLTDAMEITNKIFVCIFSFEMVIKLLGDGIAMYLSSGQNVFDGIIVIVSVCEILLSRQTSALSVFRSIRLLRIFKLVRPVRYQLLVVIKTMTSVMTFFGLLFLFIFAFAILGMNLFGGKFEFPNKEGKSVTSRSNFDSFLWAMVSVFQILTQENWNLVMYDGMRTTNKWAALYFTALMAIGYYVLFNLLVAILVEGFTNTGSALSVFRSIRLLRIFKLVRPVRYQLLVVIKTMTSVMTFFGLLFLFIFAFAILGMNLFGGKFEFPNKEGKSVTSRSNFDSFLWAMVSVFQILTQENWNLVMYDGMRTTNKWAALYFTALMAIGYYVLFNLLVAILVEGFTNTGEPKQTPKTEEQKGQNSNAGKKDTPKISKTFYGSQNCCPGGYYHDDLPEFCGHGLRNALGIMKKKRNLYVDIFLELKNETKKRQWAKFTVTRQNWSLYLFAPSNRFRCRMASVCEHKYFDYVVLMFILISCIVLAMEEPNILPKKRQIIDATMLLLTVIFTCEMMMKDVTPSVLCLLTSFHHRMIRRAPGLKLVVQTLLYSLKPIGNTVLIAAIFFVMFGILGVQIFKGKFHYCKGNDNVTNKAECEQGGNKWENRMYNFDNLAQALISLFVFSTRDGWVKIMHNGIDAVGIDQQPITNYAEWRLVYFIPFLMLGGFLVLNMIVGVVVENFQRCRERLEDEEKQKKRKKLLGKVKTEKQEDDKRNYSEGYSKWRLRILGICLHPYWDVSIAIVILVNVICMSLEHYQMPKSLEVFVEIANYFFTGVFVLEVVVKFIAFGFARYFSDRWNLVDLVIVVLSLAGIIIESLNSARNVLKIMINPTVARSLRVLRFIRVLKLVKLAKGVRSLLDTLFEALPQVANLGLLFLLLFFIYSCLGIQLFGNLKCCPSHPCQGLGPHAHFRDFGTAMLTLFRIATGDNWNGILEDTLRCSSEDKCEEYCCVSKYTAPLFFFTFVLAAQFVLVNVVIAVLMKHLKESKEKIAANMAVKDLEKKLKLLTLSAKNFIKAKARRSDPVTTSRRQSIVQLGLSGIELVQLRQSEAPETVKESGFCTDLARADAIFQEFLRLNDNLRMVKMTIVRTSSHNTEQITPTPVTPYPRQRRNTLSGIRELHKTSSNLKKYTVYKSGTEL

>XP_020626608.1|Orbicella_faveolata_Cav3b

MEEDRETSDTTLPTQHDIDGSSEFEALSCFIISKDSKGRRWMIQLIKWPWFERISMFVILINCVTLAMYNPLDPQCISTRCQVLEHVEHVVFAFFFAEMVIKMLAMGVTGKKGYLQDKWNRLDCFIVVIGLIEKVIKSGNYLTIMRAFRVLRPLRAINKVPSIRILVTLLLDTLPMLGNVLLLSFLIFFVFGIIGVQLWQGKLRNRCFTTFSKDSAFFNRYNRSTFYQPSFYFPDFVCSSPEDRGTSKCSDIKPYYLDGLECHGSFENNTVANKSVCVNWNQYYINCTPNGPNPFYDTTSFDNIGIAWIAIFQVITLEGWSDIMYFVQDAHSNWNWIYFVVLIVMGSFFLVNLCLVVITMQFQETKAREIELMENSKDESPLRSSKPLKTLFKLLGTVCNYNSQEKEHVHHHHHHIYHHHHFHHHLYNCFVPPSPGPNSFPSANVPKIQVQEEGTLSSSPNFLEEPGLFQDFLTPPDQPSRRGSKTSVCSHTLSIHAETSSAVSIEEIVAQTKTSVKLSIPSPTTGFFAFSSSNTASRTNSISGKTEVSASTTVSAEINPLVARSSTSEDEVTEPSKESKEVQSKDRDALKYRRFSQQAHVKLARTRKLGISSYNSSEHFDEDVAELSKNEINPEMADVDFLSSLDGDIENEHMEGDAAFKDLFLERGNDECQPENTSGRSLKRPSLKRRNISTQSEDDMFESLFNLRQSASYDVTANHGAWQKLRTLCRKITESKQFTFVIMSAILLNMICMGLEHYQQPERLTVALETVNIIFVSIFAVEMIIKLLGFGVTAYVSQGQNVFDGFIVIVSVCEILLSQGNSALSIFRSIRLLRIFKLVRPVRYQLLVVVKTMTSVMTFFGLLFLFIFAFAILGMNLFGGKFKFENAEGKQVVSRSNFDNFLWAMVSVFQILTQENWNQVMYDGMRSTRKWAALYFIALMAVGYYVLFNLLVAILVEGFTNSGKRKASPKPKEETGGKTLGEPVVRKFEMPHILISKEDSEEPCPGNYNPESLPDFCGHGVRNALGTTFQHKTNFTPSLSLATLEPLETKNSVAASQIKDANLTMPQLKKIRQESKLAKQRVRRTSLNAVCGDFELANSAAYNSGFLCDELVTEDTSNLDTALESERHSEASSKRTPELTWKIESSTIAGRRSRMNQSEVSLATSRPESPQIVFQNNHNTRDVYQLRECAAPESKTRKVRVMKITDIRSDWSLFLFSPSNRFRRFLTAVCGHKYFDYAVLFFILISCVVLALEEPNIPSDHQKRRIIDIAMLTLTIIFSLEMMIKIIAHGLVLGPGTYLKDGWNVLDGILVLFSWIDVIITYSSATAPEVLGALRVFRALRTLRPLRMIRRAPGLKLVVQTLLYSLKPIGNTVLIAAIFFVMFGILGVQIFKGTFHYCEGNRGHVTNKSECQNDSGRWENHMYNFDNLPNALVSLFVFSTRDGWVEIMHNGIDAVGIDKQPIKNYREWRLAYFIPFLMLGGFLVLNMIVGVVVENFQRCREKLEDEEKQRRRRKLLEKTKNRNQEG

>XP_022784214.1|Stylophora_pistillata_Cav3b

MEENRERITDHTDHSAEFETLSCFLISKDNKARRWMIQLIKWPWFERISMSVILINCLTLAMYNPLDKDCKSTRCQVLENVEIIVFAFFSAEMIIKMIAMGVTGKKGYLQDKWNRLDCFIVIIGVIEKAVVRVNGNYLTIIRAFRVLRPLRAINKVPSIRILVTLLLDTLPMLGNVLLLSFLIFFVFGIIGVQLWQGKLRNRCFTEFPENSTFLTRYNKSQFYQPSFSNPDFVCSLSKDYGTRLCSDIEPKEENVSTKCPDSSGNVSAEIKRVTFNWSQYYNKCKTSDQNPFYNTTSFDNIGIAWIAIFQVITLEGWSDIMYFVQDAHSFWNWIYFVVLIVMGSFFLVNLCLVVITMQFQETKAREIELMENSKGEQSLESPKLLKTLMKYLQHFCNCKPQEKEHVHHHHHHFYHHHHFHHHLYNCFVPPSPVIGTDHEAAAGDPSFFPDTTNFPKIQVEAEETTCSHSCVEESPILQDFLAPPFTRQSSRRSSRTSICSHALSIHAETSSAVSIEEIVTQAKTSVTLNIPSSPVATFFAFSSDNNASRKNSTQSRKREFSASTLTSAEINPHHARTSTSQSEDVFSKTSKEATESSIQRKDKEPLKYRRFSQPAHAKLARTPASEISFQPSKSLETKEADDYRDVEANPEMVNFDVISSLESDLEKSDATQDPPSNDLVSTSQANDCQIDDTVNTTLKRPSLKRRDIPAQSEDDMFESLLNLRQSATYDVTGNHSAWRKLRIICRKTIESKQFTISIMGAILLNMICMGLEHYEQPDRLTVALENINIIFVSIFGIEMVIKLLGYGVTSYLSEGQNVFDGLIVTVSVCEILLTKDASLSIFRSIRLLRIFKLVRPVRYQLLVVVKTMTSVMTFFGLLFLFIFAFAILGMNLFGGKFIFKNAEGKQVTSRSNFDDFLWAMVTVFQILTQENWNLVMYDGMRATSHWAALYFIALMAVGYYVLFNVLVAILVEGFTNSGKRKTTPKNEPGASKNEDAPKPAFKEVEKPHILISCKDSEESCPGHYQEEPVPEFCSHGVRNALGSTFQLRSNYTPSLSLATLEPLQDKKPPTAAQPTEANMTMPQLKRMRQASASIIRKTGQMAHSTDWRHDEPIFSVHNRGFLGDDMETRGKLHLEVSERERCNKAGGKTLEEIVSPTMACSLSFPNQSEGTTSYQSRPESPHMIFQNNSSDEMYQPRECHLPEAQECTKKVNKITRKRSDWSLFLFSPSNSFRRAVMAVCEHKYFDYVVLAFILISCVVLALEEPNIPRNSKKRQIIDIAMHILTVIFTIEMLMKIIAQGFLLGAGTYLKDGWNILDGILVLFSWIDVIITYSPADAPEVLGALKVFRALRTLRPLRMIRRAPGLKLVVQTLLFSLKPIGNTVLIAAIFFVMFGILGVQIFKGTFYHCEGDSEVKDKKDCRGEGCWQNRMYNFDNLFNALISLFVFSTRDGWVEIMHHGIDAVGIDKQPIKNYREWRLAYFIPFLMLGGFLVLNMIVGVVVENFQRCRERLEDEEQQRRRRKLLEKARDKNQEEVDRTYYESYSPWRRHIHDMCLHLYWDVAIAIVICLNVLCMSLEHYQMSKAFQRFVDTANYFFAAVFVLEVIFKFIAFGFVRFFKDRWNIIDLVIVILSLTGILIQSLRTTKALGAKVPINPTVARSLRVLRIIRVLKLVKLAKGVRSLLDTLFEALPQVANLGLLFLLLFFIYACLGIQLFGKLDCTRSSCQGLGPHAHFKDFGTAMLTLFRIATGDNWNGILEDALRLSCGESAECVSRYIAQLFFVTFVLAAQFVLVNVVIAVLMKHLKESKEKIAANMAAKDLEKKLKLLTTTAKNFILAKAKKNDLSISRRRSIVQIGLGGIELTQLKHGTPMKNIDSSFRAEFAKADAIFQEFLLLHENLQKVKMTICRTASHNIEVTPAPVTPSLRQRCNSFCGVRELHSSEYDLGKFTVYKSGAKL

>evg954676|Trichoplax_adhaerens_Cav3

MLKVDMSSNGHRCHQRFAASSSYQSDQPSHHAIDRNKHQDLPYHGKAKSPDLLSPISAYDRYQQGRQQQDGTNQNYKADNYRQPHDIIREFQSEQQINSNDDADIEEEIHPVVCWYLHKDQWPRKYFVKLSQWRWFERITIIVILINCVTLGSYNPAGQFKNNTCVDSTCQITSVVDNIIFGYFVVEMIIKMVALGVFGKYAYFSSGWNRLDFIIVLTGCLEYLINEGEFLTIIRTVRVLRPLRAINRVPSMRLLVNLLLDTLPLLGNVLMLCFIVFSIFGIVGVQLWKGILRSRCTLQQNITAPPGIFLYRYYLPSYEDPDYVCSLPEYNGIHHCRDLTLSNLTLDLTCKITNQTLASQYGDGTCTEYLQFYDTCSTSGPNIFYDNISFDNFAMAFIAIFQVITLEAWVDIMYAIQDGHASIDWIYFVILILIGSLFLLNFTLVVMATQFSETKRRVTKKARNSPSAIRSRRGSFSTLSSSAESRTCYNETIRLIRRIILYIAYLIQQFYSRCLKKIWNRLYNRCLTCKRKWSSRHQRSTSMSESRVNGNAKFTSDDQRLDRLHQNFFLPDIDGSCLSDGTERSTYLAFRLDNWRQRSYSQNTTTTSQIYQESALRDRLGKRNGDLAINNSLYDGTTDNDGDTPIHPLESNTELQLFNFQLRHQPDGVSLTTDLSNKPTNYQKLKQRIRRLRVRCCIFTKSQKFSLIVLFAILANTIVMAIEHHNQPTYQIQALEVCNIIFTIFFTLEMVFKLFALGLLRYAKDSFNVFDAIIVIVSLVEIATDGKGLSVLRSFRLLRIFKIVRFLPTLQRQMMVMAQTFDNVVIFLGLLFLFMFTFSILGMHLFGNRFCLARVKDGPVVCSRKNFDSLLWAFVTVFQILTQEDWNVVMYDGMLARGKWAAIYFLALVTLGNYVLLNLLVAILVNGFQEQEKDEKNRKKDLTAKMGRALKHYCDSITDVRSEDQPANRQVNAPDENSGYQVGKDLQGIPTTQIVEGYYGKSTPICDHQSCAMESSALNQSNKREEDGDSQTEFYIRTFQSSSLSCNHNQEEEQQQLPNLVDPNEVQIFPCRVPIGPRDPPQSKNKFAKLRRLMPWKSKQKWQAIRRKYWICPEEVNNPNWLTRHRNHSLLILPKENRFRKWCKNVVKNPYFDRIILVVIIFNCVTLAMERPGIDPNSMERHFLNLMVIIFTFIFTSEMIIKVLALGLVTGDKSYLRNGWNVLDLLLVIISWVDLIITYTGGASNILGVLRILRGFRTLRPLRVINRAPGLKLVVQTLFSSLKAIGNIVIICVAFFVIFGILGVQLFSGKFYYCKATDDVENKTQCVEQYGLSEWVNRPYNFDDLVNASLSLFVISSKDGWMDITYHGIDARGVDLQPKKNHNVVVLLYFISFLLLVGFFVLNMFVGVVVENFHKCQEEHERRQREKKRNRKLQRAKKNQNKANAKKVIIKKKVKEKKKNHIPPVEDYPAWRRKLYRFCVHRYFDITITIVIAVNIIFMATEHYKMSQAWEEVHKYANYFFTVVFTLEAVIHLVAFGVVYYFRDRWNIFDLLIVILSWTGIIIESQVLSNPTINPTIIRVMRLLRIVRILKLIKAAKGIRALLRTIMNAMPQVVNLGMLFFLLFFIFAALGIELFGRLDCTTDNPCNGLSQHANFKTFGMAMLTLFRIATGDNWQGILQDTLRENCDRSENCQQNCCSNPILSSLFFVIFVLAAQYVLTNVVVAVLMKHLEESDEKENNDLDPGSSSAKESQGLDDGMTESNNYYPPGLTDIHLSSNHKNLDKIGEDYTTHQFDQNHTLKSPLPYQNKYARITVSRSITAIVPQKSVLDQAENEVVMQDKFQVHDIRTKLSYPLDDHDDVEDQMELTASIHCENRKLHRQGQNSDDGHPYLQQHSPLFRHQSYESNCHHGSIQSDFNSFSNLSSDTSSFSENPLKSNNSKDNPTRDRIYTADDLNRILSKKRKSLKRSNSWDFSFTSEFSNHRRYPCSKKKTIYPFHYPSTKLSKDDSTISDQPMVVIAKEKKVTLPRKMIQTLV

>XP_004995501.1|Salpingoeca_rosetta_Cav3

MSTTATETTAATTSTLQSASVSTQRKQRTWLAQAGATVRDACAALSRSFIFEQAVLLVILFNCITLALYDTSDSTCSTRRCKILEVCELVVTVAFTVEMLIRMIATGVRQYFGSGWNRFDAIIIVFGFVDFIPTVSGGTITLVRLVRILRPMRMVTRFQSLQLLVALLLDIIPMLGSLAILTLFLFCTFGLVGVQMWKGMLRQRCYDSNLTAHFVPSTHATPPWTTHYICGAPDFTCPPPSSVVGGPNVTTTTTTSSITASVVDAFPVCAARAPNPFAGAVSFDNIAIACNTVFQVITLETWGNIMAAVQKAHSFWAFIYFVLLIFAGSWFALNLVLVVIATQFKLTKKRVMAERALGALLRPVRPRTRRWEEVVADFARRWLKMDIVHINETPEEVQHDIMHSDALTEEEKEDIVATMQVFRMFDTDGDGTISVGELTSALNSTSDDMIDSEQIRDLMCQIDADGDGTIDFAEFMEMLKRSRRHSTAIMADAPDMCGDDQGHDTQTGHPDDDGDDDDDDDDGGHGDSDGVDNEGNGRRDEHDTGNFDSNHANSRHRSRKLGRHSSLSHEVRRASIRLSQTLRQPHITVGGVAVGHLSASAVPLTTHQHHRHFTHATAAVAGHASTPAVNDSKFLRPRIAARGRPVLRSRHTVAVAPIPPTPPPSTTTTPAATTPTAATALDARQAALAEATTATASALASSPTSAATVATLLSTHASGTAVDAPPAARAQSLQSASASQVTSVTAAAMAARLAAWQSSGGGGGGGVNGAGFKVPSASAQRQNPRADQAELQRHHFHGETAAASFDLTQLPKRIATRTIPRPASTAMRTCTSDDEENKQTLNECHCNACERTKASAQDCGDGDEQTRDERVRDGPHNENMADVVASSAAVTSSPPTATAATTRPATSVAETQLRPGTAKVISGFGRAVAQIRRACARIARHPRFSNFVVICIFVNMAVMSLEHMGQPHALEEFNRITNAVFTVVFAGEMVIKIVGLGPIAYLINKANVFDFVIVLISLSEFGTGTNGLTVLRSFRLLRVFTALQVLPTMRRQLAVMIKTLDSVLTFLFLLGLFVFMAAIAGMRLFGQRLELPEWEEDGTGVPRANFSTFWNAVILVFQVLTTEDWTLVMYKAAHATSPTACIYFVLVLVLGTYILFNLFIAILVEGFATDLEALKRFKEAVVKTRDVLHDIPSTSTLAYATSPIHGGLGQAQQERQEQREQQGDIVLYDAAMQRCDCPHMTVSANLDASVRMPTSATTTTATAVLGSACGTDSGGDANSAGDGKWCSRDGLPAVSRHRSQQKPSAWRQFRAVVDAMVCACEPVRRLVCCWQSRRVAPAPTPPLPPSPQSPSSTARAARAGSAATTSLATTPDAITPEEERINEDDNRPPLFVRVCPGCNGWRHGEARGHGPRQHVLGERDQPQHWQPHHEHTHRRHELLWRLRVARARACVRRVLQNPRCEAAMFVLILMSALTLLWETPRAGEKRKRILQGVYMFFNTAFLIEVLCRVFAEGFLPRKRPSQEAAQQDHQGHQQQMQHTQQQSLPHGSQISNLEQSAAAAAATPTTASIDAGTAGTPTTTAATATEAIAGSSIKPQSARAGVSSAAVSSAPAPQLAPTDASGDAWAEESDGFLASGWNILDGFVVLTSWISMGLELGGLSSSVLRTIRVLRALRALRPLRLVHRMPKLKQTVGTLFTSVRPLTNVLLIEIIFFLIFAILGVQLFKGSLHACTMTHGVVINASTCNNQSYAATLGYTLPTTMAEVPTTAVATVTASALDQAGVPLATAVAVTAASELIGLNSASSTPHAFSGGGTNTTTGPVPGAGLCGHVRVTDKASCIAVGGEWEAFQYNFDNLAMALLTLFVVSTRDGWVLVMDRATDAVGPGRQPQRNANPLAALYFVAFVLVVGYFVINMFVGVLVENFQRSMPLDEGDEDNKDDKGGDIEDEEKEHQHQHQQEQEQEQEGEEREGEERDGGVEAVQQHGEGEKRFAAVCAHERSEATVSRLDAGVQSSDGDDDGDDDDAGGDGGDGDGDGLAAGWHESGHAVRCPTDLLIPQVLVSSPSPSLLEETLKNDQIEDNGSGNGNGNATAIAGTGCRDIVSHHHSIVSGASSDTDVALTLNAREDSSTSQARVPDITSQTSTHVAINTTAGADDPACATTPHRRRKGHTGEWTRFRLACCALVHHPTFDLVTSSLNASNVVLMMAEHHGQPAWVGHMLSTAELFFTACFTLEVIVKLVASGARVFAQSAWNKLDLFVVVTSIAGIVVEWSTDGLPVNPTVLRVLRILRVARVFRLAKMTRGMRSLLATVTQSASQVGSLALLLLLLFYISAAIAVELFGRMSCSVDAPCSGLSSHTNFRNLGMAMLALFQVATGDNWTGILSDALRRPPKCNDAPDCEIGCCANPHIASLFLVTFVILAQFILLNVVVAVLMKHLTQVLEHDQVDRRRRSKM

>XP_001701475.1|Chlamydomonas_reinhardtii_CAV

MTYPEYSLLFLRKSNFVRKWCVYTTHTRVFEWTILAAIIANCVTLAVSSNRQDFDETPLGRTLVNLEYLWVAIFTTEALLKIVAMGFVLAPGTYLRDGWNIVDFTVVALGFVDIFSSGNLTALRTVRVLRPLRAITRIRGMRILVTTMIAALPMLIDVFALCAFTFFIFGLVAVQLFSGRMTHRCAVPVFDGSYTEVVGGSTFYRNVTYVVQDEESGDGCSGPMSEGLEWVDVNGTAVALAGGDGLGRACPDGLYCTNYGNPNYGITSFDHILWAWLTIFQMITQEGWTDIMYFTSDTITWWVWPFFVALVAFGSFYIINLALAVLFMQFSNDNHADDPSKKKSGRQGSGDGQRQGDKAGAGGGGAANSGGEHKDSDSESSMSDVGSEEDERLRPTMAEVTGLRTVGQHSTKLKSATSSPLTMDEVERLSPFRRRLYRVAASRWLELFTMSLIILNTVLMCINWFEMPESVERATNYINYVFTVYFLVEMILKMTAFGLVRYFRDGMNIFDCLVVVISVTEMVLDIIPSVSGLGPLSVLRAFRLLRIFRLARSWKELNLIIRAIFKSITSTTYLLLLMLLFMFIASLMGMQLFGYKFMFCDYVEGAAPTCPLGWNVWGQCPDHFHCYLPCMSADYGSWVNATGSFYNDLAYCERFCANDQAAAAADANATSPDVAAAGGCEYLAMVGKSEVPRANFDNIFWSMYTVFQILTMENWNNIMYDGMRSTTPWCAAYFVAVVLIGTYLVFNLFVAILLDNFSSVFGESDDSDGSGSGSLKADGQAAGSGKHKSRRSSSSSSDTDSLHDSDLDAGEAGDAFGSMRSHCEYAGNEFADSPHADDASRPSSPGGTYYSTEGYHTRFSREYERAHGAGTGRGSGEGGGAGYTTHEAHAQRATGGPGGGHAPTGASPAPQQGEGEDVAAAGGADGPGAGAGPGIGIGIGGSRRAPPTLHLPDSPSAVGALPPIGPAHQPVPSPKTPFSAGIGYAHPSPGAKGGHPVTQSPSDLSRGLRGVLRKVVPVDDEGRPGSSHSGSQPPRPSLRPVSAAPRPLSSVRGGGGNRVRPVSGRPPVKFDGVDSGREGGGGGGVTFANDAVSRRASFIAAHQTAAAGSQGAGEANGGSTSQGGLAASISRSIGRSIGRSVSRTLQRAMSSIRSIREVRHNEHLAKQIKGRSLLLLDPEHWVRWRAARMVHHTHFETVILGLIVLSSITLALDSPGLDPDSQLAQALRYLDYIFLGAFTLEAALKIITFGFAFTGKHAYIRNGWNVLDFIIVLAGYALLAVELSGANGEDLKMLRILRTLRALRPLRAASRYEGLKLVVNTLFAVLPAMADVALVCALFYIIFSILAVNLFKGQLYNCIDADSGERLDPYYLLPPGQTLERGMCEAGSITVNSSVYTAARNISLMPYNITTSWVNPVANFDNVAISMLTLFQIATLELWVDIMFTAVDVAGVGKQPLWNNHPVVILFFILVVIVCCFFVLNLFIGVTLDKFTELQQAQTASSVFVTPQQQNWVDVQKLLLRTGMTSRPARFEEPAWRAGLYDFVMGSVFEEFILITIVVNVLFMAMVHADMSPQWQACMTYTNLIFTCVFVIEAALKIMAFGAFAYFRDRWNAFDFFVVVISVASVVLDFSGTQNLSFMPVLRVLRVVRVVRLIRRAVGMRRLLLTLVQSLPALGNVGGVMVLFFFVYAVIGMNLFGGIKFGDYISRHANFNNFGKAMLLLFRQASMARYYEARFLAVSPGRMITGESWSGVMQDCMITHNCVLITANVTAPATNATLVEGTYLSPNDPSMSGLPPDVHQNQCSLSPWAAVVYFPTFIVLCGFILLNLVIAIILENMITSENDEGLTVSKSLVGQFVDAWSQVDNCATGYVHASKLPLIISHIEPPLGTRGQGNARAVTQAVIMSVDVPIHENNTVTFVETLHALAGRVAGTELPEIEEEWLVEKYSKRLPDGGAAFPKLTAAHFHAALYVQAAIRGFMARHKMRGMMQTLAGGPGGAGGGTGSGSGADADGLHNGGKAFEKGGKVA

>KXZ52368.1|Gonium_pectorale_CAV

MSSNRQDFEDTTLGSTLNKFEFLWVAIFTVEALLKIVSMGFALAPGTYLRDGWNVVDFMVVALGYIDIFTAGNLTALRTVRVLRPLRAITRVRGMRVLVTTLLGSMPMLLDVFLLCAFTFFIYGLIAVQVFAGALRYRCGTPVFDNAYGGGLDASGQQVLHNVSYVVSDDDAGELCGGPMAADVTWYNVSGTPTASHGSAGDGHVCPAGMYCTSYGNPNYGMTSFDHILWAWLTIFQCITQEAWTDVMYFTSDSLSWWVWPFFVVLVVFGSLYIINLALAVIYAQFMSDSAAAEEKRKAEVALKAAEAAEGVQREAERAAEEQRKAGEQGGGEGKAALAAAAGKPGKEGDRHSSQGHGLAYNHPGRHSDTGSEAESLASVEHGGLQASGAVAGSKAALQRKPTMTLGSADAIFTAEEVEAMHPLRRAACRVAVGKRLEYTTMVVIILNTALMCINWFRMPASVENSANIVNYVFTMYFFLELLLKLFAFGVVRYFRDGMNIFDFIVVLISMVEFIMDVIPSVSGVGPLSVLRAFRLLRIFRLARSWKELNFIIRALFRSVTSTTYLLLLTLLFLFISALMGMQLFGYKFMFCDYVDGARPVCPLGLRVWGDCPEHFHCYLPCTAEEYGSWIPAPGSFYNGQAYCERFCATPAAAEAADANATVPAVAVGGGCEYLGMVGKSDVPRARFDTVFWGIYTVFQLLTTENWNNIMYDSMRSTTPWGAVYYVAVLVIGTYLVFNLFVAILLDNFSGSVTDSLDTSARSDPALKQQEDQLGAKAKRSSDSSSEDEYDEYGNDYYGAGSEYSAYSLGNLELGVTETGDAGSELAPGEHPLQYVSRRHMHPDLAASAPHGHGVSTSGGQAAVDWSDGQSEAAASSGTGFAQPQGFSPLKGVKRKVVPVDRDGAPESSSRPVSARPHSARAGRYAGFLRVSAATKASADGGRRELPQRDPSGRGVAFARDPAASASARAAVTVESRPGASGVQGTLKGVAASLRAITEVRRNEHLARFIRGRSLLLLDSNHWLRWRAARLVHDTRFETVVLVLIVASSVTLALDTPSLQRGSRLELAIRYCDYVFVGAFTLEALLKIITFGFAFTGKHAYVRNGWNVLDFLVVLIGLTMIALESSGLDANNLQMLRVLRTLRALRPLRTASRYEELKVVVNALFAVIPAMGNVALVSLLFYLIFAILAVNLFKGQLYNCVDADTGERLDPYYLLPPGETMTRQWCEAGAATINASVYTAERNITLPAYGLNTSWVNLRSNFDNVGSAVLTLFQLSTLELWVDIAFSAVDATGVDQQPLWNHQPQMLLFFGLFIVVCAFFILNLFVGVTLDKFMEMHEAQTARSVLITPQQTAWVDVQKLLLRTSMDQRPAQPDGPPWREGLYALATSNAFNNFIMGTITVNVLFMAMVHADMSNSWQAVMSISNVIFTAIFAVEAALKLAAFGPTDYFRDKWNCFDFSVVLISMASIALDFSDTQNLSFMPVLRVLRVVRVIRIIRRAKGLQRLLVTLLYSLPALGNVGGVMLLFFFCFSVIGMNLFGGIKFGDFLNRHANFNNFPNAMLLLFRMITGESWNGIMHDCMITKGCVLLTSDFTSPATGAALAAGSYFDPGDPQLSGVPPDLLNDRCALTPAAAVIYFPVFIILCAFIMLNLIVAVIVENMIMAGADEGLPVNKSMTAQFVEAWSRVDPAATGLAAASSLPLVVQALDPPMGTRQARGVNRQATQEVIMSVDIPLHPQNMVSFIETLHALSGRMAGAELPDIEEEWLMERFARRLPGGDGPFPKYTAAHYHASQYVRAAIRAFMVRHRLKEMVQALASEPPVKTPPTLLESGEGMPSGKAGA

>GAX86028.1|Chlamydomonas_eustigma_CAV

MITYRVRRPITAYPRVDWRKSLHPYWLFCVDCVTFPPWEIVRSVNLAPKALLFLQVNSFIRKPCIRLVRWMFFDALMLLRMVFVFGKYTYLRDGWNILDFIVVVMGVLELTSLGNYTFIRSFRALRPLRAITKIASLKIIVESLFRSLPMLGDVMILAMFYFSVFGIFCTELFKGQLYGRCGAPAFDQAYDVVQEGDTKMTTQVVIMNVSYVVSTTAATQVCKGPLSSDQIWYNVSNTAVAAPFSYIGGLQWGYACPYQPSSNPNDINYPSGVFCTNYGNPDIGGYRNFDNILITWVQLYQHMTWQDWSYIMYATQAAMSWWTWPLHIFLVIVGGLLLANLALAVIFLHFSKYYNEAKLNSASSLDSSKSAKMLAVELNVVTDIPSAGDGLRQPILDFPPGPSLQKPLVVTGISNIAWQQFRDLNYTICYSTWFLHLTTFMIVSNAIVLAIYWYEMPQEWVTGTTNANIAFSCYFVLEMLIKIIGMGPRQYAADSFNIFDFFVTLLGVVDMSLTLAPGVSSPGALSVFRTFRLLRVMRLARSWTGLNRIIQVLLSSLVSVGWLTVLLFMYIFITGLLGMAFFGFKLDSCPQVPNAIQLCPPGLTWMDCPPHFDCYVPCNSSVALSWFSVSGSPYGNQAYCEVFPRSQVASFMQQESTLQVVNNTLQDGNAITTTPVPNKTKLEQLQFWAQVGQSTTFMPNYDNIFQAMLSTFIILTSDNWDSNMKTIMVLTQSPWLPAFYTIITMTLGIFTVLNLFLAILLNNLDDLVVISSQSSNVERVQDEETRYIEDLYGAAVTSINKQLSGSSINRTGSSAREYSRNKMVPQALAVEVQVRSHIPVGMIMDVAEEGNKTSVLSSNTNNGGLEITGYGTVAGEVAASAAGLLTQMQGLNAAPVTQVVVNSHRLNSPESGARKEAVLAPHLGTTGGIGDGPQSVLSDAQQPVAAAVYLDPSQLKESDNGSMSSSMMERLSKTRTESFWPRKINRVSPLSQGQYPVQGLHQTEHSDDELFSVNGHAKPSSVNGHAMSMTSIGKSHAGHEPLIWSTHRLRAESGFSLHQQGPERLKSTRRSSAFRQPPVETLEGRSLFMFAPTNQVRKFLLLVTSNVHFEYAMLFLISLSSLELCFDDASSVPGTTKFAALRALDVFFTITFGLEALMKIFTYGLLFNGKDSYLRNPWNILDMFVVIVNVLVLALDTVTNPNYIIWLRAFRALRALRPLRVASQLDGIRVIVMAMAKSLPAMGEIFLVGALFFYIFAVLGVNLMCGLFLGCYSQGNLLNPAYYVGLGEGINRTWCEADGGIHNITHSYYHDMINVAVPKWQLSTSWGANGQLARFDNLIMALWVLFWMTSLENWSPIMIQAMDITSLDDQPVFNNNIYITFYFIVFIVIGVYFIMNLVIGVAISTFGKMREQLGRSALLTEAQQEWLTIQRMLATLQLTKKYKRPSGRFRLSVYKMVMTERFEKIMMCIIIANLLPLFMSYQGESDTWAAGLGVVNVVFTALYVIEMILKWISIGACAYFKDKWCLFDFLVVVVSVMGVIIDYVLHDNLTILTVLRSLRVLRIFKIIPKARGLKMMMTTLLWSLPALMNVATVLLLFMYTFAIIAMNIFGNMKWTGEIDLYANFESFPTAMFTLFRMQTGENWNYVMTACMNLQQCIQVTNDVDIIIPGNTTASVIYRGTYLDTESDALTLSVVPSDLQNNRCSPSPAVAAIFFTLYMALCTYLVLELVVAVIIENIEYQSQIENMAVKQRHIEDFCTAWEELDQTSCGFTEAAGLTTILTKVSPPMGVKGLDCQSQYIQDIVMSCNIPLRGLKIHFMETLHELTGRVAAAPLPAEQEWVVHDKIMQKLPQDKVLPKYTVGDYYSALYVKASIKGYLIRSKFNNVGRLPLFGSEAGAKEAEVETGAKQAVAQVGIREADIEAYTREDVAQVGAKEAEVEGLLSIQDQPDAPKVALHPIYPGETLHQTSLAMVQSPEHCLQSNSTPKESMREEDTGLRPSALFVTADKQGQALGVGKEKEAQSM

>GSPATP00010323001(Lodh_2016)|Paramecium_tetraurelia_Cav1a

MSKEVAVTSTLKQRLSAIEDKEIYFEDRDSFSEDGKYGEYVQKQNREILKKVQELMHDGQSTVNQQEDQSFAVMNENQKQEILEDLEEEFGQSYTETKVKVNTYIKLIPKNELQLAYIYSDKYPFLVIRKQILIFTKLISSYAQKITTHPLFELMTLLMIIFNSAMLAIDDPTTNVQTSFQDLTDIIFLAYYTAEAVLKIVALGFILPKKAYLKDTWNILDFSVIVTAYIPYFLSSNSVNLNALRSFRVLRPLRTVSSIKALRTILLALFASIAQLRDAVVVLIFFYSIFAIAGVSLFSGYLKRRCIGEMSGITWISDEILFCADDNNCPFPEDTIYNENFICGKQIANPQNDLVNFDTFGYSFLQVFIITTLEGWTQIQTAVMLTFSQYVVLYFIIVVIVGAFFLVNLTLAIIKLNFKPEKIQEELAQIKEEIEEYDYRQLRQLKLYDPQRPIVDTDGYGISWDKHHDFDQNAIMNRRKNSRRGASQLNMIRSQNMKKLRGKNKVSFQPIKSAVYYSNPIVLKNKQLKIEGIYGMGNQSKLNQQTLGVSGQNNNNRSSVVRRSSQSSNNDERTSSKKEDKNNINENESQSQSRPTKVIPRKSIQFGDQQISSEFLNSPMMDLQSENIRPLNGGTMIVHHQGSMESSKSGGSKQDKTPTNNQLEIPRLGEIRQQSQNVKSNQKFSLVAGMKYQKPPRNSPKPDSQKSLDQNEFDSVNLSELNTISDDDLDQRLEEMDIFVQDKNDDSESEKKKKQNQKELNRQKRMENRRLRDLNKNAQQHLDDASLKLKSKLYNTKFYPIIIVDQEFGSVNDVLVSQMLLKLEKEKLEQEEKIKEMDIKITYCFKNQKQSLELKKSSSGKSKSMTKSGYLKSGLKSKKNNLSLRRVQPIMDELHPFEDLQFPTDKNFENQDDQNKENENSDSDDEPDEQNNEKNNNFNGSASQKIKMKKKKKDQEQNEKLDLQQIGEKFSESSDCIVDLNRIRQVETDKQFNQQELALIPASELEQQKGVFYQEFQDIKQRDIEESKGAIVAQASIEDVLLIADYTFYDSKFAKQMNSVMKALNYSKRETFIYLQGFIGFLKVCQNHLLYFVQSGYFEAAMNLAVALNTVILALDGLLPDSSANTLQQFNLGFTILFTIELGLKVIGMGPKNYISDTMNVFDAVIVALSLVELFFLGGGTSGKSSLSAFRSVRIFRAFRVLRVTKLMRSLQFMGFLIKVLGNAFQSFMYIMVLLLLFIFIFTLLGMAFFGGQLSKTPSRQSYDDIQSAFLVVFQVLTLENWNSILWDLLIQDVSAFITVPYLVFWIMIGNYVFLNLFLAILLENFEEEYKNDKAGLDTNIEIGQDSIMDNTTQQVNSTSTLKSTMKTKKHTVAQQLENNGDPESKKHKQFQQIFQYFVEPGLCQFSLYLFSQENIVRRICYRIVKDDKFETLIFFMIFLTSTKLVFDTYIPDTGKLKETSLQIDIFFAVFFGVEMIMKIIAFGFVQQESSYLRESWNILDFFIVIASFIDVSVSTINLSFVKILRLLRTLRPLRFITHNRSMKILVSALLQSINGIFNVAIVVILVWMMFAILGINLEKNKMHYCDTGDDEIWYHYGPDECAKHGGVWANRKVNFDNILNGMLTLFILSTLEGWPDQMYWFIDADESGPIKGAQLQFSWYFIVFILIGSILLMNLFIGVILVNYHLAEEASRDKILTQPQVDWIELQKLIVHANPNLAMFFSPENPFRAKVFIIIKHRYFDPTILMIIVCNIVTMGLSQDDSPIAYDNALQSLNTAFTFVFITEALLKIIALGPVGYMRNSWNQFDFFVVCASILDLVLQFTGNSFISFLIFRVLRVTRLFRLIKSFEGLQKLIETAIYSLPAMLNVTALLFLVFFIFSILGVFLFGSIRSGWAIDDVNNFSDFHHSFELLFRCSTGEDWYKVMFDITMQDGQGYYCIFFIIFIVIQQYIMLNLFILIILDQYEINYFNSDNPLNKFQEYENMFIESWSKFAKEDKGMKMSQQLLVPLLLDMEKPIGYDLKQKLNDEISEWRRINPQLDTKENVLKQTLILKAQAKRAVSTQIMKMNVYADVAGQVKYHQILFSVMKSYMWKKVQLNLSPEGAEKILQKEDETQKRLKKKQVGVQSKEVNLVNPIVHFLFVHMAFKTLKRYGEKKKQQKELQAQLLLEQEQEHYSEGFSSDSSFDNKIEILSRVSKDTEHPKRPDYGKTKYLTLPNTEIYKEEIYINQETPVVNDSDRSSKEEEEEPDEDVMQQYTKDFKKHLINPNGSNGNIQDDKSASNQHQGGGATRSSNSRTSQINKQPKRLTLKPNDMIQGLGLSKKSIQTNQSPANQNPNVSQDNPSIMNQSNGNSSSNK

>GSPATG00033414001+GSPATG00033415001(Lodh_2016)|Paramecium_tetraurelia_Cav1b

MSKDEAVTATLKQRLSAIQDKDIYFEDRDSFSEDGKYGEYVQKQNKEIMKKVQELLNDGQSTVNQQEEQSFAVMNENQKQEILEDLEEEFGQSYVEAKVKVNTYIKLIPKNELQLAYIYSDKYPMLAIRKQILIFTKLISSYAQLITTHPLFELMTLLMIIFNSAMLALDDPTTDVQTSFQDLTDIIFLAYYTAEAVLKIVALGFIFPKKAYLKDTWNILDFSVIVTAYIPYFLASNSVNLNALRSFRVLRPLRTVSSIKALRTILLALFASIAQLRDAVVVLIFFYSIFAIAGVSLFSGYLKRRCIGEMSGITWISDEILFCADDNNCPFPEDTIYNENFICGKQIANPQNDLVNFDTFGYSFLQVFIITTLEGWTQIQTAVMLTFSQYVVLYFIIVVIVGAFFLVNLTLAIIKLNFKPEKIQEELAQIKEEIEEYDYRQLRQLKLYEPERYIVDTKGYGTTWDKHHDFDQNAIMKRRDNSRGGASQLNMIKQNNLKRLRGKNKVSFQPIKSAVYYSNPIVLKNKQLKIEGIYGMGNQSKLNQQHQGTTQNNNNNRSSVVRRSSQSSNNDDRTSSKKEEKNNSNSNNNNNENESLSRPTRFAPRKSMQFGEQQISSEFLNSPMMDQLTENIRPLNGGTMLVHHRGSMESSKSGESRPDKTPINNQLEIPRLGEIRQQSQNVKSNQKFSLVAGMKYQKPPRNSPKPDSQKSLDQNEFDSVNLSELNTISDDDLDRRLEEMDIFVQDKNGDSESETKKKQNQKEMIRQKRMENRRLRDLNKNTQQHLDDTSLKLKSKLFKTKFYPIIAIDQDFFSVNDVLVSRMLLQLEKEKLEQEEKIKEMDIKITYCFKNSKQSLELKKSSSGKIKSMTKSGYLKSGLKSKKNNLSLRRVQPIMDELHPFEDLQFPTDKNFENQEDQNQENVNENSDSDDEPEENNEKNNNFNGSASQKIKVKKKKKDQEQNDKLDFEQMGEKFIESSDCIVDLNRIHAVEIEKQFNQQELALIPAWELEQQKGVFYQEYQDIKQKDIEESKGSITAQASIEDVLLIADYTFYDQKFAKQMNSVMKALNYSRRETFIYLQGFIGFLRVCQNHLLYFVQSSYFEAAMNLAVALNTVILALDGLLPDSSANTLNQFNLGFTILFTIELGLKLIGMGPKNYISDTMNIFDAIIVALSLVELFFLGGGTSGKSSLSAFRSVRIFRAFRVLRVTKLMRSLQFMGFLIKVLGNAFQSFMYIMVLLVLFIFIFTLLGMAFFGGQLSKTPSRQSYDDIQSAFLVVFQVLTLENWNSILWDLLIQDVSAFITVPYLVFWIMIGNYVFLNLFLAILLENFEEEYKNDKAGLDTNIEIGQDSIMDNTTQAVNSTSTLKSQMKTKKATVAKNLENNEDPESKKHKQLQQVFQYFVEPGICQFSLYMFSQENIIRRICYRIVKDDKFETLIFFMIFLTSTKLVFDTYIPDTGQLKETSLQIDIFFAVFFGVEMIMKIIAFGFVQQESSYLRESWNILDFFIVIASFIDVSVSTINLSFVKILRLLRTLRPLRFITHNRSMKILVSALLQSINGIFNVAIVVILVWMMFAILGINLEKNKMHYCDTGDDEIWYHYGPEECAQHGGVWANRKVNFDNILNGMLSLFILSTLEGWPDQMYWFIDADESGPIKGAQLQFSWYFIVFILVGSILLMNLFIGVILVNYHLAEEASRDKILTQPQVDWIELQKLIVHANPNLAMFFSPENPFRAKVFIIIKHRYFDPTILMIIVCNIVTMGLSQDDSPIAYDSILQSLNTAFTFVFITEALLKIIALGPVGYMRNSWNQFDFFVVCASILDLILQFTGNSFISFLSAGPQLARVFRVLRVTRLFRLIKSFEGLQKLIETAIYSLPAMLNVTALLFLVFFIFSILGVFLFGSIRSGWAIDDVNNFSDFHHSFELLFRCSTGEDWYKVMFDTMQDGQGYYCIFFIIFIVIQQYIMLNLFILIILDQYEINYFNSDNPLNKFQEYENMFIESWSKFAKEDKGMKMSQQLLVPLLLDMEKPIGYDLKLKLNDEISEWRRINPQLDTKENVLKQTQILRAQAKRDVSTQIMKMNIYADNVGQVKYHQILFSVMKSYMWKKVQVNLSPEGAEKILIKEDETQKRLKKKQVGVHSKEVNLVNPIVHFLFVHMAFKTLRRYGEKKKQQKELQAQLLLEQEQEHYSDGFSSDSSFDNKIEILSRNSKDTDHPRRPDYGRTKYLTLPNTEIYKEEICLNQETPIVNDSDRSSREEEEEPDEDVMQQYNKDFKRHLVNPNGSNGNIQDDKSASNLGGGGGNTRSSQSRTSQIPKQPKRLTLKPSDLISGLGASKKSIQSNQPSANQNPNASQDNPSIMNQSSGNSASNK

>GSPATP00010323001(Lodh_2016)| Paramecium_tetraurelia_Cav1c

MSKEIKVTAGVKKRLTALEDQEISVQNQDSFSDDGKYGEYVKKQNREILKKIKELIHDGQSTTNQQEEQSFVIANDNHKQDLLEDLEEEFGQSYQESKVKINTYVKLIPKNELQLAYIYNDKYPFLVIRKKILISTKLVSFYAQLVTTHPIFEVITLIMIVFNSVMLAIDDPTTNVQSPFQNLTDLIFLAYYTFEAVLKIVAQGFIIPKKSYLRDTWNILDFSVIITAYIPYFLASNSVNLNALRSFRVLRPLRTVSSIKALRTILLALFASIAQLRDAAVVLMFFYSIFAIAGVQLFSGYLKRRCIGEESGITWVSEEILFCADDNNCPFPEQTIYNENFICGKQIANPQNNLINFDTFGYAFLQVFIITTLEGWTQIQTAVMLTFSQFVVLYFIIVVLVGAFFLVNLTLAIIKLNFKPEKIQEELAQIKEEIEEYDYKQLKQLKLYVPEREKVDTAGFGITWDKPNDFDQNIIMNRRKHTRKGGSQMNIIKNTYQRRFKGKNKVHFEPIKSAVYYSHPVMLNKKYVKIEGIYGMGNQTKLNQQQQITSNIQSSIAAKRSSGVRRSSQSSNNDDKTSSKQMFDNESGSGHKQPKKSLQLCDQQLSSEFMNSPMMDLQSENIRPLNGGTMLVHRGESIDSSRSGESKLGRTPVNNKLEIPKIGDIKQQSQSIKPNQKFSLAAAMKFQKQPRNSPKPDSQKSLDDHEFDSVNLSELNSLSDDELDQRLEEMDIEVEEKVTNVNSVNQKQIQKELNRQRKMENRRLRDLNKNSQQNLDEASLKLKSKLFDTKFYPIIVVDAEFNSVNDILISQMLQKLEQKRLEQEKKIKELDFKITYCFKNSKQSLELKKTSNGKTKSLTKSAYLKSGNKSKKTNLSLRRVQPILDEIHPFEDYQLPTEKIAEGQEEQIQENQNGNSESDEDNENENNIDRDQNNVNGSANQKLNQKKRKKRDEQNQKLDYEQLGEKFNDSSDCIVDLNKIRQVELHKQFNQDELALLPPQEIELQKGVFYQEFVDIKQKDIHESKGIITAQASIEDVLLIADYTFYDQKFARQMNSVMRALNFSKRETYIYLKGFIGLLLVLQNQLLYFVQSGYFEASMNLAVALNTVILALDGLLPDSTSEITNQFNFGFTILFTIELGLKMLGMGPRKYLRDTMNIFDAVIVALSLVELFFLGGSTNGKSSLSAFRSVRIFRAFRVLRVTKLMRSLQFMGFLIKVLSNAFQSFMYIMILLLLFIFIFTLLGMAFFGGQLSKTPSRQSYDDIQSAFLVVFQVLTLENWNSILWDLLVQDVSAFITIPYLVFWIMIGNYVFLNLFLAILLENFEEEYKNDKAGLDTNIEIGQDSVMDNTTQAVNSTSTLKSTMKTKKHTIAQQLENNEDDHDQKKHKQINQAFSYFVEPGMCQYSLYLFSQQNIVRKICYRIVKDDKFETLIFTMIFLTSAKLVFDTYIPDTGQLKEVSLDIDIFFAAFFGVEMCMKIIAFGFISQESSYLRESWNVLDFFIVIASFIDVSVSTINLSFVKILRLLRTLRPLRFITHNRSMKILVSALLQSINGIFNVAIVVILVWMMFAILGINLEKNKMSFCNMGDDEIYYHYGVQECKENGGVWENRKTNFDNILNGMLTLFILSTLEGWPDMMYWFIDADESGPIKAAQLQFSWYFIVFILFGSILLMNLFIGVILVNYHLAEEASRDKILTQPQVDWIELQKLIVHSSPNLAMFFSPENPFRAKIFTIIKHRYFDPTILMIIVLNIIIMGLSQDDSPILYDQVLTQFNTAFTFVFIGEAILKIIALGPVGYMRNSWNQFDFFVVCASILDLILSFTGNSFISFLIFRVLRVTRLFRLIKSFEGLQKLIETAIYSLPAMLNVTALLFLVFFIFSILGVFLFGTIKSGWVIDDTNNFSDFHHSIELLFRCATGEDWYKVMFDTMQDGQGYYCIFFIIFIVIQQYIMLNLFILIILDQYEINYFNSDNPLNKFQEYENMFIESWSKFAKEDKGMKMHQQLLVPLMLQMEKPIGYDIKQKLQTEIAEWKRINPQLDNKENVQKQTLFLKAQAKRIVSTQLMKMNIYSDNNGHVKYHQILFCVMKSYMWKKVQVNLSPEGAEKILIKEEETQKRLKKKQVQQSKEVSLVNPIVQFLFVQMAFKTLKRYGEKKKQQKELQVLIEQEQEHYSDGFSSDSSFDNNIEILSRNSKDTEYPRRPDYGKPKFISLPNTEIYKEDEYINKGIPVQNDSNKNSEEEEENPDEDVMQQYTKDFQKHLINNGGSNGNIQDDRSGGAATKSSISRQSQINKQQKKLTLKPNDLLSGMSKKSIQSISQNNNLAAIYDNPLQ

>NP_011733.3|Saccharomyces_cerevisae_CCH1

MQGRKRTLTEPFEPNTNPFGDNAAVMTENVEDNSETDGNRLESKPQALVPPALNIVPPESSIHSTEEKKGDEYNGNDKDSSLISNIFRTRVGRSSHENLSRPKLSLKTASFGAAESSRRNVSPSTKSAKSSSQYIDLNDERLRRRSFSSYSRSSSRRVSNSPSSTDRPPRSAKVLSLIAADDMDDFEDLQKGFKSAIDEEGLTWLPQLKSEKSRPVSDVGEDRGEGEQESIPDVHTPNVGASATPGSIHLTPEPAQNGSVSEGLEGSINNSRKKPSPKFFHHLSPQKEDKDQTEVIEYAEDILDFETLQRKLESRPFVLYGHSLGVFSPTNPLRIKIARFLLHRRYSLLYNTLLTFYAILLAIRTYNPHNVVFLYRFSNWTDYFIFILSACFTGNDIAKIIAFGFWDDSEMFKAYGREYKSILQRSGIMKLYIYLREKYGRKLIDFIIPFRIISPGEETKYQRSSLSTSLTKPYGAKENQRPFGTPRAFARSSWNRIDLVSSVSFWLGMFLSIKSYDTKTGIRIFKPLAILRILRLVNVDTGMPSILRGLKYGIPQLVNVSSMLVYFWIFFGILGVQIFQGSFRRQCVWFNPEDPTDTYQYDMQFCGGYLDPVTKRKQNYIYEDGSEGSVSKGFLCPQYSKCVSNANPYNGRISFDNIVNSMELVFVIMSANTFTDLMYYTMDSDEMAACLFFIVCIFVLTIWLLNLLIAVLVSSFEIANEEYKKKKFIYGSRKTGYVARIVTGYWKYFKLKANQTKFPNWSQKGLAIYSHVEFIFVILIICDIGMRASVKVSTSANCNNILLKTDRGISIVLFIESLARLVLYLPNMWKFLTKPSYVYDFIISIITLVISCLAVEGVLGHMYAWLSIFHISRFYRVIISFNLTKKLWKQILSNGVMIWNLSSFYFFFTFLVAIIMAVYFEGVIPPEEMADQPFGMYSLPNSFLSLFIIGSTENWTDILYALQKHSPNISSTFFCSVFFIIWFLLSNSVILNIFIALISESMEVKEEEKRPQQIKHYLKFVYPQKIQEYTHASLVARIRKKFFGGHRNEDTRDFKQFLMRGTAIMNIAQNMGELADEFKEPPSENLFKKGLSKLTIGVPSLKRLRMFANNPFYKNSDVVFTETNDINGRTYILELNEYEDEKLDYLKKYPLFNYSYYFFSPQHRFRRFCQRLVPPSTGKRTDGSRFFEDSTDLYNKRSYFHHIERDVFVFIFALATILLIVCSCYVTPLYRMHHKMGTWNWSSALDCAFIGAFSIEFIVKTVADGFIYSPNAYLRNPWNFIDFCVLISMWINLIAYLKNNGNLSRIFKGLTALRALRCLTISNTARQTFNLVMFDGLNKIFEAGLISLSLLFPFTVWGLSIFKGRLGTCNDGSLGRADCYNEYSNSVFQWDIMSPRVYQQPYLHLDSFASAFSSLYQIISLEGWVDLLENMMNSSGIGTPATVMGSAGNALFLVLFNFLSMVFILNLFVSFIVNNQARTTGSAYFTIEEKAWLESQKLLSQAKPKAIPNLIELSRVRQFFYQLAVEKKNFYYASFLQVVLYLHIIMLLSRSYNPGNLIGYQGVYFMFSTSVFLIQEALHMCGEGPRLYFRQKWNSIRLSIIIIAFIMNAVAFHVPASHYWFHNIKGFFLLVIFLFIIPQNDTLTELLETAMASLPPILSLTYTWGVLFLVYAIALNQIFGLTRLGSNTTDNINFRTVIKSMIVLFRCSFGEGWNYIMADLTVSEPYCSSDDNSTYTDCGSETYAYLLLMSWNIISMYIFVNMFVSLIIGNFSYVYRSGGSRSGINRSEIKKYIEAWSKFDTDGTGELELSYLPRIMHSFDGPLSFKIWEGRLTIKSLVENYMEVNPDDPYDVKIDLIGLNKELNTIDKAKIIQRKLQYRRFVQSIHYTNAYNGCIRFSDLLLQIPLYTAYSARECLGIDQYVHHLYILGKVDKYLENQRNFDVLEMVVTRWKFHCRMKRTIEPEWDVKDPTVSSHISNINVNLEPAPGILEREPIATPRMDYGVNNFMWSPRMNQDSTMEPPEEPIDNNDDSANDLIDR

>NP_593894.1|Saccharomyces_pombe_CCH1

MSSSSNSDPSSSPDNTDFIPLKDNPKDTSSYINKKNPFVNDSFDSHDDSYLDNVNPFEYNVADDEDTISMTSDGVEFHNLGITETVDEQDKILSELQLAYPTVSRYSETLDKGLLNGDKTTKGDYTSLRRFRKPFLFDKLSPYFTHFFSEIYAAVLRVVGPPRANETPGRNLPFSPILRQIYNHPLYNIFIFVVIVLHAVLLMIRSDDPHDKSQTIDYLIIVIGILYTLEMLLKIYLFGFLYDGSSSFYDFINSYTKKTPRTMSYLRHSWNRVDFVAIVALWISVIGKKQQGIFRLFSMIACLRLTRLLNITRKTETILKSLKESSTPLVQVVSFNAFFGVMIAILGVQFFKASLNRQCVWLGDYGDQYLPTGQFCGGHWENGIKRAYLDKNGFPSNVNPRGFICAEKSVCRVVENPYSNTVSFDNFFNSLELIFVIMSSNGFTDIMYDIMDAEYFVSCLLFIISAYFLTLWLMSLVIAVVTSSFIDLQHSGKNQKEQKSVDKHLIRNKLCEKYLFYSNFIWISFIVAQFVTLCTQTYDQTSSTANRYLIFYACVDFLLAAEVILRFFAYLPDYRLFFRRYTNLVDIVLAVLNLVTLLPSIRKNPVAFGWLSIFAIARIYRCILLIPYTRKIAKLLFSNFKQLLNLMLFLVIVLFIASLCAVRLFQDLPNDGDSDDDAISFATTYESFLYMYQILTSENWTDVMFAIQARLAHLHLSWIPGAFFTLWFLFSNNVVLSMFIAVIQVNFAPSESDLKMEQLKMYLARLLRNYNPFQASLTLNALIKRNGSRKGKTVDESHYEWLYEDNIIKGFLRVTNIPPKEPLKPSLDTSPELSKYSLAKLSNNFKRFVMRDDPFSQAYFKRVIGIRWEKDMNLKTAAKDMQAAKAFVRIKQTEFLKNHPKYNDVFWVIKPSNRIRRFCQRLVMPGVNERYGGVEPYQWVYRVIQVFIYACILTAVIIECIATPIYERDHLLNDKQHAWFVWTEVAFATIFTIEAAIKIIADGFCITPNAYLRSTWNCIDFFVLVTLWINLYAVLTSHALLSRAFRAFKALRVLRLINLTQTSQRMFHDALISGFFKIFSAAVVSATLLIPFALWAKNVFGGLLYSCNDDNVLSASQCVLEYASTPNNWEVWAPRVWSNPPDYDFDRFPHALLALFEIASIEGWVDIMRSVMDITGFNNQPQTNASSGNAMFFVLFNLVSMIYILTLFIAIIISNYAERTGSAFFTAEQRAWLELRRKIKSMRPSKRPAIRPLGLRGLCYDFAVQKHGIWRRTFTGLYIVHLLFLLTIFYPCPIAYTYVRNSIFLILSICYTINICVKVYGLSFYYFFHSFWNMFDVVVTLGSLTCNIAILAKFENRSLTLLQTTLLVLVTVHLIPKFDNFDQLSKTVVASLPSIFSLIATWIVLYITFAIAFNQIFGLTKLGLNGGPNKNFRSIRNALVLLFTMTFGEGWNDVMHDYTISYPNCVNGDDFYNSDCGNKPWAYGLFIAWNIISMYIFVNMFITVVFDNFSYIHTKSSSFTNLQRNDFRQFKDSWAPFDPMVTGYIPKRNAVKFVLSLRGVYDFRIYRDEHTLRSIISKVQSKTGQQSVPMLENPLMGSEINLEALDQIIDTIDVNVVKERRNVLNSLCTEIMTLPGNVISFSNILMLVVLHKIVDHREAFPISDYIRRAYVLSELEKSIRMEKLLGLVETSIIRKTFLQHMEEKKKALENPFILLSEVSEILPETPIQEVLRQHNADPEQLLMSTRTPSVSDRSFSIVESTVPTIASGEGDDNHSVEDHLKVPTDNEPRRSPSLKEVLLRGSHSLHSNNDRTSFDIEAGFGTAESDFQFGGATEDINRIADRIDDYLDRDSFKG
