## Supplementary material for "Early metazoan origin and multiple losses of a novel clade of RIM pre-synaptic calcium channel scaffolding protein homologues": File S3

>Trichinella_pseudospiralis_Cav3_KRZ43346.1

ALRCLDQRTPVRYWCLRIVNNPWFEKLSMMVILINCVTLGMYKPCDEDMNTTRCITLSVIDHLVFAFFAVEMIIKIIAMGF-YGPDTYMSDTWNRLDFFIVIAGCAEYIFKERMNLTAIRTVRVLRPLRAINRIPSMRILVNLLLDTLPMLGNVLLLCFFVFFIFGIIGVQLWAGLLRNRCLLLVRYYMPDETSMYICSLGHICNSHNPFQGSVSFDNIGFAWVSIYLVISLEGWSDIMYYVQDAHSFWDWIYFVLLIVIGAFFMINLCLVVIATQFASDSDGVYRAIVKYIAHLGRRAKRKLTSSKRYDPRFALNYTESAPVFLKVGIKRFVDSDHFTRGILVAILINTLSMGVEYHNQPEELTVILEYSNVFFTALFSIEMALKIIADGPFTYVSNGFNLFDGGIVILSIVELLQGGNGGLSVLRTFRLLRILKLVRFMPALRYQLVVMLRTMDNVTVFFGLLCLFIFIFSILGMNLFGCKFCPACKCPRRNFDTIINAAFTVFQILTQEDWNVVLFNGMAQTSPWSALYFIALMTFGNYVLFNLLVAILVEGFQEKARLMQEEMDKTEIDPSTKAKPRKVSSQLSQQICSSVNPHIFYPYCRTRMDASLFIFMPKNKFRAQCVYLSQQKWFDFCILTFIGINCITLAMERPGIPPNSFERLFLDISGYIFTVIFALEMFIKVIAKSLVFGDYFKNGWDFMDGTLVLISLSNLIFDIFSVVRVLRLLRALRPLRVINRAPGVKLVVQTLISSLQPIGNIVLICCTFFIIFGILGVQLFKGKMWHCVNVAVVNKTDCLAINHRYNFDNLGQALMSLFVVSSKDGWVSIMYQGIDAVGVDMQPIVNYFEWRMLYFISFLLLVGFFVLNMFVGVVVENFHRCKEKEMKEKEREKKMRKLQKQLSRQPYWYYGPVRMYLHGIVTSKYFDLAISAVIGVNVITMAMMPPELTYALKVFNYFFTAIFTLEAILKVFALSIPRYLKDRWNQLDVIIVLLSIGGIVLEEILPINPTIMRVMRMLRIARVLKLLKMAKGIRSLLDTVMQALPQVGNLGLLFFLLFFIFAALGVELFGRLECSDDGLGEHAHFKNFGMAFLTLFRIATGDNWNGIMKDTLTSNCCVAIIAPVYFVVFVLMAQFVLVNVVVAVLMKHLEESSK

>Trichinella_pseudospiralis_Cav2_KRY73966.1

SLFIFTENNFVRRYARTIIEWGPFEYFILMTIIANCIVLALDQHLPHNDKMPLSLKLEATEPYFMGIFTIECLLKIIAFGFVMHKGSYLRSGWNILDFIVVMSGVISMLPFTTSDLRTLRAVRVLRPLKLVSGIPSLQVVLKSILCAMAPLLQIGLLVLFAIVIFAIIGLEFYSGAFHSTCYNFSGIPVAIGNKPFPCTNAYHCP-IGPNYGITSFDNIAFAMLTVFQCITMEGWTNVMYYTNDSQGTFNWLYFIPLIILGSFFMLNLVLGVLSGEFARRQQHIERELNGYLEWICKAEEVIEARHLAVSKQLKHLVQQSTETRLRVFVRRFVKTQFFYWLVITLVFLNTVCVSIEHYGQPQWLDEFLYYAEWTFLGIFLFEMLFKMFGLGIGTYFQSSFNIFDFVVITGSLFEVIKGGSFGISVLRALRLLRIFKVTKYWTSLRNLVVSLMNSMRSIISLLFLLFLFILIFALLGMQLFGGEFNFPEGRPSTHFDTFPVALITVFQILTGEDWNEVMYLAIESQNMIYSIYFIILVLFGNYTLLNVFLAIAVDNLANAQELTAAEEAHEQE-------GQAKLTDACDAVCDVKNALNIQENKTIVPYSSLFIFSPKNRFRLFVHKIVCTKYFEMAIMVVISLSSISLAAEDPV-DESNPRNKYLNYLDYAFTAVFTVEMILKVIDMGVIIHPYCRDLWNIMDATVVICALVGFAFVNLSTIKSLRVLRVLRPLKTIKRIPKLKAVFDCVVNSLKNVFNILIVFILFQFIFAVIAVQLFKGKFFYCTDRTKRFEQDCQGTSFALNYDNTIHAMLTLFTVTTGEGWPGIRQASIDATEENQGPIPFNHIEVALFYVVYFIVFPFFFVNIFVALIIITFQEQGEAELAEGDLDKNQKQIDFALNARPVCRYIPIKYHIWKMVVSTPFEYFIMAMICLNTIILMMEPPAYRAVLRYLNSTLTAVFTVEAILKILAFGVRNYFKDGWNIFDFITVIGSITDALVTE--GGNFVSLGFLRLFRAARLIKLLRQGYTIRILLWTFVQSFKALPYVCLLIGMLFFIYAIVGMQVFGNIELNGTEINRHNNFQTFFNSIILLFRCATGEAWQEVTLACILNSECGTNFAYVYFTSFVFLSSFLMLNLFVAVIMDNFDYLTR

>Trichinella_pseudospiralis_Cav1_KRY90155.1

SLLCLGLKNPVRKLFISIVEWKPFEWLILCMICANCIALAVYQPFPAHDSDRKNAVLEQVEYIFIVVFTIECVMKVIAYGFLFHPGAYLRNGWNLLDFLIVVIGLISTALSTLNDVKALRAFRVLRPLRLVSGVPSLQVVLNSILRAMVPLFHIALLVLFVIIIYAIIGLELFCGKLHKTCVD--QWTGEHVPDPGPCGESFHCD-PGPNDGITNFDNFGLAMLTVFQCISLEGWTDVMYWVNDSVGEWPWIYFITLVILGSFFVLNLVLGVLSGEFSREKQQLEDDLKGYLDWITQAEDIDEEQEAMDREEFGADGEGGEEGRCRRSCRRIVKSQAFYWLVIVLVFLNTMVLTSEHYGQPEWLDHFQEIANLFFVVLFTLEMFLKMYSLGFVNYFVALFNRFDCFVVIGSILEFALMKPLGVSVLRSARLLRIFKVTKYWNSLRNLVASLLNSLRSIASLLLLLFLFIVIFALLGMQVFGGKFNPNMNKPRANFDTFVQALLTVFQILTGEDWNAVMYNGIAAFGVIVCIYFIVLFICGNYILLNVFLAIAVDNLADAESLTAAEKEEEGKRANEADPSGEVKLPFNEPTDAEDEHLASGDEDQIPDASSLFLFSSTNKVRIFCNKVINHSYFTNSVLVCILVSSAMLAAEDPL-QASSFRNEVLNYFDYFFTTVFTIEISLKVLVYGLILHKFCRNAFNLLDMLVVGVSLTSFGLKAISVVKILRVLRVLRPLRAINRAKGLKHVVQCVIVAVKTIGNIMLVTFMLEFMFAIIGVQIFKGSFFRCTDRARLTAEECKGTNYDFNFDNVQNAMVALFVVSTFEGWPDLLHVAMDSSDEGIGPQYNARVSVAIFFITFIVVIAFFMMNIFVGFVIVTFQSEGEREYENCELDKNQRKIEFALTAKPQRRYIPFQYKIWWFVTSQPFEYAIFIIIILNTLILGMMTRTFDDVLDTMNLIFTGIFAMEFILKVMAFRCKNYFGDAWNVFDFIIVLGSFIDIIYG-KPGSNIISINFFRLFRVMRLVKLLSRGEGIRTLLWTFMKSFQALPYVALLIVLLFFIYAVIGMQIFGKIALNSTEIHRNNNFQTFPSAVLVLFRSATGEAWQLIMLSCARGQPCGNDFAYPFFISFFMLCSFLIINLFVAVIMDNFDYLTR

>Trichinella_britovi_Cav3_KRY57796.1

ALRCLDQRTPIRYWCLRIVNNSWFEKISMMVILINCVTLGMYKPCDEDMNTTRCITLSVIDHLVFAFFAVEMIIKIIAMGF-YGPDTYMSDTWNRLDFFIVIAGCAEYIFKERMNLTAIRTVRVLRPLRAINRIPSMRILVNLLLDTLPMLGNVLLLCFFVFFIFGIIGVQLWAGLLRNRCLLLVRYYMPDETSMYICSLGHICNSHNPFQGSVSFDNIGFAWVSIYLVISLEGWSDIMYYVQDAHSFWDWIYFVLLIVIGAFFMINLCLVVIATQFASDSDGVYRAIVKYIAHLGRRAKRKLASSKRYDPRFALNYTESAPVFLKVCIKRFVDSDHFTRGILVAILINTLSMGVEYHNQPEELTIILEYSNVFFTALFSIEMALKIIADGPFTYVSNGFNLFDGGIVILSIMELLQGGNGGLSVLRTFRLLRILKLVRFMPALRYQLVVMLRTMDNVTVFFGLLCLFIFIFSLLGMELFAGRFCPACKCPRRNFDTIINAAFTVFQILTQEDWNVVLFNGMAQTSPWSALYFIALMTFGNYVLFNLLVAILVEGFQEKARLMQEEIDKTEIDPSTKAKPRKVSSQLSQQIYSSVNPHIFYPYCKTRMDASLFIFMPKNKFRAQCVYLSQQKWFDFCILTFIGINCITLAMERPGIPPNSLERLFLDISGYIFTVIFALEMFIKVVAKSLVLGDYFKNGWDFMDGTLVLISLSNLIFDIFSVVRVLRLLRALRPLRVINRAPGVKLVVQTLISSLQPIGNIVLICCTFFIIFGILGVQLFKGKMWHCVNVAVINKTDCLAINHRYNFDNLGQALMSLFVVSSKDGWVSIMYQGIDAVGVDMQPVVNYFEWRMLYFISFLLLVGFFVLNMFVGVVVENFHRCKEKEMKEKEREKKMRKLQKQLSRQPYWYYGPVRMYLHGIVTSKYFDLAISAVIGVNVITMAMMPPELTYALKVFNYFFTAIFTLEAILKVFALSIPRYLKDRWNQLDVIIVLLSIGGIVLEEILPINPTIMRVMRMLRIARVLKLLKMAKGIRSLLDTVMQALPQVGNLGLLFFLLFFIFAALGVELFGRLECSDDGLGEHAHFKNFGMAFLTLFRIATGDNWNGIMKDTLTSNCCVAIIAPVYFVVFVLMAQFVLVNVVVAVLMKHLEESSK

>Trichinella_britovi_Cav2_KRY49212.1

SLFIFTENNLVRRYAQISLTFTPFEYFILMTIIANCIVLALDQHLPHNDKMPLSLKLEATEPYFMGIFTIECLLKIIAFGFVMHKGSYLRSGWNILDFIVVMSGVISMLPFTTSDLRTLRAVRVLRPLKLVSGIPSLQVVLKSILCAMAPLLQIGLLVLFAIVIFAIIGLEFYSGAFHRIPVA-------IGNKPFPCTNAYHCP-IGPNYGITSFDNIAFAMLTVFQCITMEGWTNVMYYTNDSQGTFNWLYFIPLIILGSFFMLNLVLGVLSGEFARRQQHIERELNGYLEWICKAEEVIEARHLAVSKQLKHLVQQSTETRLRVFVRRFVKTQFFYWLVITLVFLNTVCVSIEHYGQPQWLDEFLYYAEWTFLGIFLFEMLFKMFGLGIGTYFQSSFNIFDFVVITGSLFEVIKGGSFGISVLRALRLLRIFKVTKYWTSLRNLVVSLMNSMRSIISLLFLLFLFILIFALLGMQLFGGEFNFPEGRPSTHFDTFPVALITVFQILTGEDWNEVMYLAIESQN---------------DTLLNVFLAIAVDNLANAQELTAAEEAHEQE-------GQAKLTDACDAVCDVKNTLNIQENKTIVPYSSLFIFSPKNRFRIFVHKIVCTKYFEMAIMVVICLSSISLAAEDPV-DESNPRNKYLNYLDYAFTAVFTIEMILKVIDMGVIIHPYCRDLWNIMDATVVICALVGFAFVNLSTIKSLRVLRVLRPLKTIKRIPKLKAVFDCVVNSLKNVFNILIVFILFQFIFAVIAVQLFKGKFFYCTDRTKRFEQDCQGTSFALNYDNTIHAMLTLFTVTTGEGWPGIRQASIDATEENQGPIPFNHIEVALFYVVYFIVFPFFFVNIFVALIIITFQEQGEAELAEGDLDKNQKQIDFALNARPVCRYIPIKYHIWKMVVSTPFEYFIMAMICLNTIILMMEPPAYRAVLRYLNSTLTAVFTVEAILKILAFGVRNYFKDGWNIFDFITVIGSITDALVTE--GGNFVSLGFLRLFRAARLIKLLRQGYTIRILLWTFVQSFKALPYVCLLIGMLFFIYAIVGMQVFGNIELNGTEINRHNNFQTFFNSIILLFRCATGEAWQEVTLACILNNECGTNFAYVYFTSFVFLSSFLMLNLFVAVIMDNFDYLTR

>Trichinella_britovi_Cav1_KRY55670.1

SLLCLGLKNPIRKLFISIVEWKPFEWLILCMICANCIALAVYQPFPAHDSDRKNAVLEQVEYIFIVVFTIECVMKVIAYGFLFHPGAYLRNGWNLLDFLIVVIGLISTALSTLNDVKALRAFRVLRPLRLVSGVPSLQVVLNSILRAMVPLFHIALLVLFVIIIYAIIGLELFCGKLHKTCVD--QWTGEHVPDPGPCGESFHCD-PGPNDGITNFDNFGLAMLTVFQCISLEGWTDVMYWVNDSVGEWPWIYFITLVILGSFFVLNLVLGVLSGEFSREKQQLEDDLKGYLDWITQAEDIDEEQEAMDREEFGADGEGGEEGRCRRSCRRIVKSQAFYWLVIVLVFLNTMVLTSEHYGQPEWLDHFQEIANLFFVVLFTLEMFLKMYSLGFVNYFVALFNRFDCFVVIGSILEFALMKPLGVSVLRSARLLRIFKVTKYWNSLRNLVASLLNSLRSIASLLLLLFLFIVIFALLGMQVFGGKFNPNMNKPRANFDTFVQALLTVFQILTGEDWNAVMYNGIAAFGVIVCIYFIVLFICGNYILLNVFLAIAVDNLADAESLTAAEKEEENKRANEADPSGEVKLPFNEPTDAEDEHLASGDDDQIPDASSLFLFSSTNKVRIFCNKVINHSYFTNSVLVCILVSSAMLAAEDPL-QASSFRNEVLNYFDYFFTTVFTIEISLKVLVYGLILHKFCRNAFNLLDMLVVGVSLTSFGLKAISVVKILRVLRVLRPLRAINRAKGLKHVVQCVIVAVKTIGNIMLVTFMLEFMFAIIGVQIFKGSFFRCTDRARLTAEECKGTNYDFNFDNVQNAMVALFVVSTFEGWPDLLHVAMDSSDEGIGPQYNARVSVAIFFITFIVVIAFFMMNIFVGFVIVTFQSEGEREYENCELDKNQRKIEFALTAKPQRRYIPFQYKIWWFVTSQPFEYAIFIIIILNTLILGMMTRTFDDVLDTMNLIFTGIFAMEFILKVMAFRCKNYFGDAWNVFDFIIVLGSFIDIIYGKSPGSNIISINFFRLFRVMRLVKLLSRGEGIRTLLWTFMKSFQALPYVALLIVLLFFIYAVIGMQIFGKIALNSTEIHRNNNFQTFPSAVLVLFRSATGEAWQLIMLSCARGQPCGNDFAYPFFISFFMLCSFLIINLFVAVIMDNFDYLTR

>Stylophora_pistillata_Cav3b_XP_022784214.1

SCFLISKDNKARRWMIQLIKWPWFERISMSVILINCLTLAMYNPLDKDCKSTRCQVLENVEIIVFAFFSAEMIIKMIAMGVTG-KKGYLQDKWNRLDCFIVIIGVIEKAVVRVNYLTIIRAFRVLRPLRAINKVPSIRILVTLLLDTLPMLGNVLLLSFLIFFVFGIIGVQLWQGKLRNRCFTKSQFYQPSFSNPFVCSLSTKCPDQNPFYNTTSFDNIGIAWIAIFQVITLEGWSDIMYFVQDAHSFWNWIYFVVLIVMGSFFLVNLCLVVITMQFQESPKLL-KTLMKYLQHFCNCKPQEPSKSLETKEADDYRDVEANPEKLRIICRKTIESKQFTISIMGAILLNMICMGLEHYEQPDRLTVALENINIIFVSIFGIEMVIKLLGYGVTSYLSEGQNVFDGLIVTVSVCEILLTKDASLSIFRSIRLLRIFKLVR---PVRYQLLVVVKTMTSVMTFFGLLFLFIFAFAILGMNLFGGKFIGKQVTSRSNFDDFLWAMVTVFQILTQENWNLVMYDGMRATSHWAALYFIALMAVGYYVLFNVLVAILVEGFTNSGTTPKNEPGASKNEDAPKPALHLEVSERERCNKAGGKTLEEIVSPTKRSDWSLFLFSPSNSFRRAVMAVCEHKYFDYVVLAFILISCVVLALEEPNIPRNSKKRQIIDIAMHILTVIFTIEMLMKIIAQGFLLGAYLKDGWNILDGILVLFSWIDVIITVLGALKVFRALRTLRPLRMIRRAPGLKLVVQTLLFSLKPIGNTVLIAAIFFVMFGILGVQIFKGTFYHCEDSEVKDKKDCRGQNRMYNFDNLFNALISLFVFSTRDGWVEIMHHGIDAVGIDKQPIKNYREWRLAYFIPFLMLGGFLVLNMIVGVVVENFQRCRERLEDEEQ-QRRRRKKNQEEVDRTYYEYSPWRRHIHDMCLHLYWDVAIAIVICLNVLCMSLMSKAFQRFVDTANYFFAAVFVLEVIFKFIAFGFVRFFKDRWNIIDLVIVILSLTGILIQSKVPINPTVARSLRVLRIIRVLKLVKLAKGVRSLLDTLFEALPQVANLGLLFLLLFFIYACLGIQLFGKLDCSSQGLGPHAHFKDFGTAMLTLFRIATGDNWNGILEDALESAECVRYIAQLFFVTFVLAAQFVLVNVVIAVLMKHLKESKE

>Stylophora_pistillata_Cav3a_XP_022794522.1

AFFFLHREKRPRRWFIRLVTWPYFERLSILVILINCVTLGLYDPFDPECQTQRCQTLDAMEKVIYTFFLAEMLCKWIAMGL-FGKLAYFGDAWNRLDCFIVAAGTFELLYNKGEYLSAVRAIRVLRPLRAINRVPSIRILVTLLLDTLPMLWNVLAICFFIFAIFGIVAVQLWRGALRGRCFLLTDFYIPSFDKPFVCAVNRECSGDNPSWGAIGFDNIFIAWVAIFQVITLEGWSDIMYYVQDAHGFWNWIYFVILIVIASYFMTNLCLVVITTQFQYGKDGCWVEILRYVEHLIRRFKRRRGGEMITTATAAVAINGKSAVRLRQVCRRSVDSKWFMYIIMAAIFLNTLSMGIEYHGQPTKMTHVLEILNHIFTAIFGVEMLLKLVGMGLYGYIKDPFNLFDGFIVIMSIVELFGVGDSNISVLRSFRLLRIFKLVRFLPALRRQLLVMIHTMDNVVTFLALLVLFIFTASILGMNLFGGKYMGIYSAARANFDDLFWALVTVFQVLTQEDWNTVMYDGMRATTKWAALYFILLMTIGNYILFNLLVAILVEGFANTGSLKSRSSSSRNNGDYETCRVARKTLSFGDTTTMVIANPAHAQRCIRRRNWSLYLFSPSNRFRQWNIALYKNKWFDRVILFFILLNCIAMALEKPGLKSDDKLKKAIDISMYIFLGIFTLEMMVKVIALGFWVGKYMRNSWNVMDGFLVLVSWIDVIVSILGVLRVFRALRTLRPLRVISRAPGLKIVVETLISSLKPIGNIVLIAATFFIIFGILGVQLFKGKFYHCKDSEVATKNDCQKENKEYNFDNLARALLTLFVFSTKDGWVTIMYDGIDAVGVDKQPIPNHNKWNVLFFVAFLLLAGFVVLNMLVGVVVENFQKCRDMIEKDRLAEKDKEREKLQSEADDEAEFPQPRRFFHQICTHGYFDLGISAVIFLNVICMAMQPEEMTLFLRYANYVFTAIFIVEGVLKIYALGFKKYIKERWNQLDLLIILLSIVGIVLEEKLPINPTIIRVMRVLRIARVLKLLKTAEGIRKLLDTVAEALPQVGNLGLLFLLMFFIFAALGMELFGQVECTDEGLDHHAHFKDFGFSMLTLFRVSTGDNWNGILKDIISTSGCAEHIAPIYFAIFVLATQFVLLNVVVAVLMKHLEDAKE

>Stylophora_pistillata_Cav2c_PFX33508.1

ALFCLPEDNPIRFYCKKIVESKKFEYFILLTIAVNCVVLMLDEPLPNGDTTKRNEQLEKSEKYFVIIYCIEAATKIIASGFLLHKDAYLRNGWNILDFVVVVVGLVGMISDPEISLKVLRAVRVLRPLKIVSGIPSLQVVMKSIARAMIPLLQILFLILFVIVIYAIVGLELLRGKFQWTCYNTTDGLDTKFITGRVCSVGRPCD-RGPNKGITTFDNIFLSMLTVFQCITMEGWTDIMYHSYDARDVITSIIYISLIIIGSFFMLNLVLGVLSGEFARSREKIERQVTAYTDWIGRAEDIIKKRYSLSDSIMHLIEDHGEMVLFRIHVRQMVKSQVFYWSVIVCVFLNTVLMSVEHHGQPDWLERFQAISEYVFLSIFIVEMLLKMYGLGPRVYFKSAFNRFDCAVVLGGIVEIVTNYSFGISVLRSLRLLRIFKFTRFWASLRNFVTSLLNSMRSILSLIFLLLLFIFIFALLGMQLFGGKFSERLEAPRTNFDNFLKAMLAVFQIMTGEDWNTVMNDGIVASGILSSLYFVSLVILGNYTLLNVFLAIAVDNLANAQAVTQDEKEEQLQPEAMRKKVKISQDANGYLEANGKFLNGNSREGGIIRKSSMFIFGPDNPIRQACHWVVNLRYFDDFILVVILLSSILLAVEDPV-NPEARRNKVIRYFDYGITGIFALEVLVKMIDLGVILHKYLRSGWNIIDAFVVACNIAALLLDQKDAIKSFRVLRVLRPLKAINKSKKLK------------------------------------GKFWYCNDRSKMTRETCRGTKHKFHFDNVPNAMLALFSSSTGEGWPQGMHNTIDATKEDHGPIKDYQIQMSLYYVCFVVVFSFFFLNMFVALIIVTFQEQGEKEMDGCELDRNQRDIQFAMTAKPRQRYMPCFYKVWKVVDSKPFEILIMATIVLNAIVLMVESSEYEKVLDYLNYAFTFVFLIEAVLKLIAFRQ-NYFRDFWNVFDFIIGVTTSVGMILEF-----------------------------------------KALPYVVILIGMLFFVYAVIGMQLFGRIDLSERQINHHNNFRSFLMALQVLFRASTGENWHKIMLDCFKSKTCGTVASVIYFCTFYFFCTFLMLNLFVAVIMDNFEYLTR

>Stylophora_pistillata_Cav2b_XP_022780503.1

ALFCLSETNPLRELSKSIVVSKVFEYFILLAIGANCIVLALNTPLPNNDRTDMAQQLEDAEYYFVAIFCVEALLKIMAFGFVLHPGSYLRNGWNILDFTVVVVGVISLPQVSAIDVKALRAVRVLRPLKLISGVPSLQVVMKSIVRAMVPLLQIALLVLFCILIYAIIGLDFLKDKFHTTCVN-KTTNLTASSNPKPCDKGRSCE-IGPNSGITLFDNIALSMLTVFQCITMEGWTSIMYDTFDAMDYLYACYYVSLIVIGSFFVLNLVLGVLSGEFARRQQQLNRQVDAYLSWIAKAAETTSQSQLVTSDVVSNPGRVVTRRKLSVKVGHMVKTQAFYWTVLVCVFLNTIVLAVEYHNQPRWLSNFQEWAEIVFLTLFFVEMILKIYGLGLHIYFNSSFNCFDCSIVISGFLDIIVQIKLGISVLRCVRLLRVFKMTRHWRSLRNLATSLVSSIKSIVSLIFLLFLFILIAALLGMQIFGGKF-TEEEIPKTNFDNFQNAMLAVFQILTGEDWNSVMYSGVIAYGIAVSLYFVLLVILGNYTLLNVFLAIAVDNLANAQILTEDEENEKLERERARAATSSDADEKKHKTRRIKRGKKEANGDTILKTKTFFIFGPENRFRRQCHRIVNLRHFDNFMLVIILLSSITIAIENPV-NDDAKLNRVLKYFDYVFTGIFALEVLIKVVDMGIILHKYFRDWWNIIDALVVSFNIASLILVGHSLIKALRVFRVLRPFKGVHKIKKLQAVFRCMWYSVKNVANILMITMLFLFIFAVMGVQLFKGKFQYCNDSSKRTKEECQGSNKDLNFDDVLQAMLTLYTSSTGEGWPSAMKTTMDTTEIDKGPIHNYSPGYALYYIAFVVVFSFFFLNIFVALIILTFQEEGEREIASCELDRNQRDIQFALTAKPAQRYMPLQYKVWVIVMSKPFDTFILVLIALNTGVLMSQDDQFTDILMYLNIAFTVLYMIEAGLKFFALRL-KYFRDYWNIFDFIVVLGGLMDVLVTVGTLGIGIDPSMFRLFRAARLIKLLRRGYTIRILLWTFLQSFKALPYVTLLIMLMFFMYAVIGMQLFGKIALDSTEINSKNNFRNLLQALQVLFRSATGEDWHKIMLACYEKYSCGTVGAILYFCTFIFLCMFLMLNLFVAVIMDNFEYLTR

>Stylophora_pistillata_Cav1_AAD11470.1

ALLCLSLGNPIRSAAINLVEWKPFDVMILITIFANCAALAAFEPLPEKDSSEINDNLEVAEYVFLAVFTMEAVLKIIAYGFLFHPGAYLRNGWNILDFVIVVVGLATILVKATLDVKALRAFRVLRPLRLVSGVPSLQVVLNSIIKALIPLFHIALLVVFVVIIYAIIGVELFMGRLHKTCYD-NVTGAESFEEPHPCSSGFQCD-KGPNHGITNFDNIGLACMTVFQCITLEGWTDVLYWINDAVGSWPWVYFVTLIIWGSFFVLNLVLGVLSGEFAREKQQVEDAYNGYLDWITQAEDIEKKTSSRQSRTEDIEMIDRNERRRQTELRKAVKTQAFYWIVIVVVFLNSLTLALEHYDQPDWLTKFLDIANKLFLGIFTIEMIVKMYCLGFHGYFASLFNRFDCLVVISSLLELAKQPPIGISVLRCIRLLRIFKVTRYWSSLSNLVASLLNSMRSIAGLLLLLSLFMLICSLLGMQIFGGKFNDDDEIPRSNFDSFWRALITVFQILTGEDWNAVMYDGIRAWGAIAILYFIFLVVVGNYILLNVFLAIAVDNLADAENLTEMEEEKKKKKEKAREKLDKSTQELHSTGTLNGNGVARTASHDMPPESALFIFSPTNIFRVVCYKIATNTYFVNFILCLIIVSSILLAAEDPL-NASAKRNQVLNYFDYFFTSVFTFEILVKFISYGLILHKFCRSAFNLLDLLVVSVSVISISLRQFSVVRILRVLRVLRPLRAINRAKGLKHVVQSVFVAVKTIGNIKLVTMLFQFLFAVIGVQLFKGTFFSCNDEKILTAEECQGRRHDFNFDNVGNAMLTLFTVMTFEGWPGILENSIDSTEVDKGPNQNNRPWVAIYYIIYIIIIAFFMVNIFVGFVIVTFQSEGREEFKGCELDKNQRQIEFALKAKPLKRYIPLQFHIWPVVTSQAFEYLIFAFIVCNTVVLMMEPKLYTRVLDGFNIGFTAVFLLECILKLIAFKPKNYFTDRWNLFDFIIVVGSIIDITMNEVSSEQMFAFGFFRLFRALRLVKLLNQGSGIKTLLWTFIKSFQALPYVALLIVMMFFIYAVIGMQMFGRIAINSTAINRNNNFQTFPQSLMVLFRSATGENWQQIMLACTPSGLCGSDFAYFYFVSFYSICSFLIINLFVAVIMDNFDYLTR

>Sturnus_vulgaris_Cav2.1_XP_014748708.1

SLFLFSEDNVVRKYAKKITEWPPFEYMILATIIANCIVLALEQHLPDEDKTPMSERLDDTEPYFIGIFCFEAGIKIIALGFAFHKGSYLRNGWNVMDFVVVLTGILASVG----DLRTLRAVRVLRPLKLVSGIPSLQVVLKSIMKAMIPLLQIGLLLFFAILIFAIIGLEFYMGKFHTTCFD---LVTDEIKVEVPCGTARICP-EGPNYGITQFDNILFAVLTVFQCITMEGWTDLLYYSNDASGTWNWLYFIPLIIIGSFFMLNLVLGVLSGEFARRQQQIERELNGYMEWISKAEEVISKTDLLSPEEGEEQLGDIAAMRMRFHIRRMVKTQAFYWTVLSLVALNTLCVAIVHYDQPDWLSDFLYYAEFIFLGLFMSEMFIKMYGLGTRPYFHSSFNCFDCAVIIGSIFEVIPGTSFGISVLRALRLLRIFKVTKYWASLRNLVVSLLNSMKSIISLLFLLFLFIVVFALLGMQLFGGQFNFDDGTPPTNFDTFPAAIMTVFQILTGEDWNAVMYDGIKSQGMVFSVYFIVLTLFGNYTLLNVFLAIAVDNLANAQELTKDEQEEEEAANQKLALGRREDKERRHRRRRETQPPPAPSGPTMVPYSSMFILSTTNPFRRLCHYIVNLRYFEMCILMVIAMSSIALAAEDPV-QPNAPRNNVLRYFDYVFTGVFTFEMVIKMVDLGLVLHQYFRDLWNILDFIVVSGALVAFAFTDINTIKSLRVLRVLRPLKTIKRLPKLKAVFDCVVNSLKNVLNILIVYMLFMFIFAVVAVQLFKGKFFYCTDESKEFEKDCRGKKYEFHYDNVLWALLTLFTVSTGEGWPQVLKHSVDATYENQGPSPGYRMEMSIFYVVYFVVFPFFFVNIFVALIIITFQEQGDKMMEEYSLEKNERAIDFAISAKPLTRHMPFQYRMWQFVVSPPFEYTIMAMIALNTIVLMMASDTYENVLKMFNNVFTSLFSLECLLKIMAFGVLNYFRDAWNIFDFVTVLGSITDILVTE--GNNFINLSFLRLFRAARLIKLLRQGYTIRILLWTFVQSFKALPYVCLLIAMLFFIYAIIGMQVFGNIGIEDSAITQHNNFRTFFQALMLLFRSATGEAWHEIMLSCLKEDECGNEFAYFYFVSFIFLCSFLMLNLFVAVIMDNFEYLTR

>Strongyloides_ratti_Cav3_XP_024510580.1

ALHCFSQTKIPRKWCLKMVTNPWFDRLTMIVIIINCITLGMYKPCEDGPNTYRCQLLSMIDHMIFIYFFLEMVIKVIALGF-TGQAAYLSDTWNRLDFFIVIAGIAEYLLQEYLNLTAIRTIRVLRPLRAVNRIPSMRILVNLLLDTLPMLGNVLLLCFFVFFIFGIIGVQLWAGLLRNRCTLLSRYYIPEDTSLYICSQGIKCNPMNPFQNSVSFDNIGFAWVAIFLVISLEGWTDIMYYVQDAHSFWNWIYFVLLIVIGAFFMINLCLVVIATQFAENNGDTYAALVRFISQTARRVKRHSDDLLEKKDKNSDNNNSICALEIRNIVKKFVICDHFTRGILVAILLNTLLMGVEYHQQPEWLTIILEYSNYFFTGLFAFEMLLKVFADGLFGYLADGFNLFDGGIVALSVLEIFQEGKGGLSVLRTFRLLRILKLVRFMPALRYQLVVMLRTMDNVTVFFGLLALFIFIFSILGMNLFGCKFCDSKKCERKNFDSLLWALITVFQILTQEDWNMVLFNGMAQTTPWAALYFVALMTFGNYVLFNLLVAILVEGFQEEEQLEEEAKKRAEEEEKERNKLENSRLRTNSWCGIQTLFNPQCPIHSKRKDYSLFLFSTKNKLRISCLKLTQKKWFDYTVLVFIGINCITLAMERPSIPPDSLERKFLTIIGYIFTIIFTLEMSLKVIANGCLFGRYFKDGWNILDGILVIISLINIIFEIFGVIRVLRLLRALRPLRVINRAPGVKLVVMTLISSLKPIGNIVLICCTFFIIFGILGVQLFKGMMYHCIDVSITTKNECLAVNHRYNFDNLGQALMSLFVLSSKDGWVSIMYQGIDAVGVDMQPIENHNEWRMIYFISFLLLVGFFVLNMFVGVVVENFHKCKEAEMREKARKRQQYEDHCALKKKPYWYYGPARMYTHNIVTSKYFDLAIAAVIGINVISMAMMPSGLRYVLKALNYFFTAVFTLEAAMKLYALGLKTFFMEKWNRLDMFIVILSIAGIIFEEELPINPTIIRVMRVLRIARVLKLLKMAKGIRSLLDTVGEALPQVGNLGSLFFLLFFIFAALGVELFGKLECSDDGLGEHAHFKNFGMAFLTLFRIATGDNWNGIMKDALDTNCCVPILAPCFFVVFVLISQFVLVNVVVAVLMKHLEESNK

>Strongyloides_ratti_Cav2_XP_024503488.1

SLFIFSEDNFIRKNAKAIIEWGPFEYFILLTIIGNCVVLAMEQHLPKNDKKPLSEMLERTEPYFMGIFCFECLIKIIAFGFILHKGSYLRSGWNIMDFIVVVSGVLTMLPFSPSDLRTLRAVRVLRPLKLVSGIPSLQVVLKSILCAMAPLLQIGLLVLFAIVIFAIIGLEFYSGIFHSACYNSDGEIENLSEKPFPCSNAYNCD-IGPNYGITSFDNIAFAMITVFQCITMEGWTSVMYYTNDSLGTYNWAYFIPLIVLGSFFMLNLVLGVLSGEFARRQQQIERELNGYLEWILTAEEVIKQQSTETEEELEEEEEEEDEENIRTSLRIIVKSQIFYWSVITLVFLNTACVASEHYGQPAWFTEFLKYAEYGFLGIFICEMLVKLFAMGYRTYFASKFNRFDCIVIVGSAFEVLKGGSFGISVLRALRLLRIFKLTSYWVSLRNLVRSLMNSMRSIISLLFLLFLFIVIFALLGMQLFGGKFNFPNMHPYTHFDTFPIALITVFQILTGEDWNEVMYLAIESQGMVYCIYFIVLVLFGNYTLLNVFLAIAVDNLANAQELTAAEE-----ADEKANEICEDSDDGEDENGDQCIDMEERDYYDMVPYSSMFIFSPTNCLRVFVHSFVSTKYFEMFVMFVICLSSIALSAEDPV-DEENPRNKVLQYMDYCFTGVFACEMFLKLIDQGVILHRYCRDFWNVLDGVVVVCALVAFSFANLNTIKSLRVLRVLRPLKTIKRIPKLKAVFDCVVNSLKNVFNILIVYFLFQFIFAVIAVQLFKGTFFYCTDSNKKFAHECHGKLRPFNYDNTLNAMLTLFVVTTGEGWPGIRQNSMDTTEEDQGPSPFFRVEMALFYVMFFIVFPFFFVNIFVALIIITFQEQGEAELSEGDLDKNQKQIDFALNARP-RSFMPMKYRIWRLVTSTPFEYFIMAMICCNTLILMMNSPGYEKVLRFFNTALTAVFTVESILKILAFGVRNYFKDGWNRFDFITVVGSITDALVTE--GGHFVSLGFLRLFRAARLIRLLQQGYTIRILLWTFVQSFKALPYVCLLIGMLFFIYAIVGMQVFGNIRLDPTEINRHNNFQSFFNSVILLFRCATGEAWQDIMLSCTKGATCGTNMSYAYFTSFVFLSSFLMLNLFVAVIMDNFDYLTR

>Strongyloides_ratti_Cav1_XP_024499702.1

SLMCLSVRNPIRRACIGIVEWRPFEWLILLMICANCIALAVYQPYPAQDSDTKNTILEQIEYLFIIVFTIECILKVIALGFLCHSGAYLRNAWNMLDFLIVVIGLVSTVLSRMNDVKALRAFRVLRPLRLVSGVPSLQVVLNAILRAMVPLLHIALLVMFVIIIFAIIGLELFCGKLHSTCVD-VNTGELAMKNPTPCGFAYHCE-QGPNNGITNFDNFGLAMLTVFQCVSLEGWTDVMYWVNDAVGEWPWIYFVTLVILGSFFVLNLVLGVLSGEFSREKQQLEEDLKGYLDWITQAEDIENEDGMDEEGEERTEDSRP---RCRRACRRLVKSQSFYWLVIILVLLNTLVLTSEHYGQSEWLDNFQTAANLFFVILFSMEMLLKMYSLGLTTYTTSQFNRFDCFVVISSILEFILMKPLGVSVLRSARLLRIFKVTKYWTSLRNLVSSLLNSLRSIMSLLLLLFLFIVIFALLGMQVFGGKFNPQQPKPRANFDTFIQSLLTVFQILTGEDWNTVMYNGIESFGVLVSIYFIVLFICGNYILLNVFLAIAVDNLADADSLTNAEKEEEQG-------DYEDEKYLENRGDDDQEEDEHAVIPEIPKASSLFILSHTNPFRVFCNKIVNHQYFTNSVLVCILVSSAMLAAEDPL-QAQSPRNLILNYFDYFFTTVFTIEISLKVIVFGLVIHKFCRNAFNLLDILVVAVSLISFVLKAISVVKILRVLRVLRPLRAINRAKGLKHVVQCVIVAVKTIGNIMLVTFMLQFMFAIIGVQLFKGTFFSCNDPSKMTEAECRGTKNDFNFDNVLDAMISLFVVSTFEGWPDLLYVAINSNEEDRGPVHNARQAVAFFFITFIVVIAFFMMNIFVGFVIVTFQNEGEREYENCELDKNQRKIEFALKAKPHRRYIPFQYRVWWFVTSQFFEYAIFIIILLNTTTLAMPPTQMDHILDVLNLIFTAVFAFEALFKIIALNPKNYFGDRWNAFDFVIVLGSFIDIIYGKSPGSNIISINFFRLFRVMRLVKLLSRGEGIRTLLWTFMKSFQALPYVALLIVLLFFIYAVIGMQVFGKVALDSTQIHRNNNFHTFPAAVLVLFRSATGEAWQEIMLACAPNALCGVNFAYPYFISFFMLCSFLVINLFVAVIMDNFDYLTR

>Strongylocentrotus_purpuratus_Cav2_XP_011662956.1

--------------------M-----------------ITQGVNFGGVNSASVCNLA----------FIIQSSEKS--------PG---------------LTG------------------------------PSVELP-------------------------------------------------------ETVK------SEGPQ--------------------------------------FVWVYFIPLIILGSFFMLNLVLGVLSGEFARRQQQIDKELMGYLEWICKAEEVMRRDAATRTEDLGNGKPSGADVRLRFAIRHAVKTQAFYWLVIVLVFLNTICVAIEHYNQPHWLEQFLYYAEIVFLCIFIMEMVIKLYGLGPGVYFQSAFNKFDCIVICASMFEVIKEESFGLSVLRALRLLRIFKVTRYWTNLRYLLISLVHSIQSIVSLVFLLFLFLVIFALLGMEFFGGDFNATQAKPSSNFDTFWIALITVFQILTGEDWNVVMYQGIKSQGMWASIYFVILVLFGNYTLLNVFLAIAVDNLANAQELTKLDQEEDEENRQAAANCPSCDNCHCPCVCECSCSCQMRKKANMVPYSALFIFSTTNPVRRFCHYIVTLRYFDTMIMVVIALSSIALAAEDPI-DPDNKRNKVLEYFDYIFTGIFTIEMVLKIMDMGLLLHKYMRDLWNILDAIVVVCALFAYGLLQLNTIKSLRVLRVLRPLKTIKRVPKLKAVFDCVVNSVKNVTNIAIVYLLFMFIFAVIGVQLFKGKFFHCSDLSKVTEEECQGDKYEFNYDNVALAILTLFTVSTGEGWPDVLKHSIEATEEGMGPTPYNRIEMSLYYVVYFIIFPFFFLNIFVALIIITFQEQGDQDFFDGEIDKNQKQIEFCITAKPVDRFMPFRFKVWKLVVSSAFEYFIMTLITLNTFTLMVQPEIYSAILKNLNIAFTVLFTIEAMLKLTAFGIRNYFKEGWNTFDFITVVGSVADVIIS-EVGDGFINLSVLRLFRAARLIKLLRQGSSIRILLWTFIQSFKALPWVCLLIWMLFFIYAIIGMQIFGNIAPIDQQINRHNNFGQFFSSLLLLFRCATGEAWQSIMMSCLEFDTCGSYFSYVYFVSFSFLCSFLMLNLFVAVIMDNFDYLTR

>Strongylocentrotus_purpuratus_Cav1_XP_011670134.1

SLFCLTLDNPMRRMCISIVEWKPFEYLILITIFANCIALATYTPFPKQDSNDVNRNLEYVEYAFLIIFLIEALLKICAQGFLFHPGAYLRNGWNILDFLIVAVGVISTILSLRNDVKALRAFRVFRPLRLVSGVPSLQVVLNSIFRAMVPLLHIALLVIFVIIIYAVIGLELFMEYMHKTCYF-KDTSIIAMDDPHPCGNGFRCT-EGPNDGITTFDNIGLAMLTVFQCITMEGWTDIMYDINDGAGWWPFLYFVSLIIIGSFFVLNLVLGVLSGEFSREKQQIEEDLRGYLDWITQAEDIDHVPKPISESDSDDKSEEMGSSRCRRLCRQAVKSQAFYWIVIIMVFFNTVILASEHYSQPAWLTDFQDFGNLCFVVIFTIEMIIKMYSLGLQGYFVSLFNRFDCFVVCSSIIEVVIIPPIGISVLRCVRLLRVFKATRYWTALRNLVASLLNSMRSIASLLLLLFLFILIFALLGMQVFGGHFNSTKLKPRSNFDSFFQSLFTVFQILTGEDWNEVMYDGIQAYGFLASTYFIILYICGNYILLNVFLAIAVDNLADAESLTALEKEKEEEKRNKSIRSKSQDSEENVNVIDEDDVKKSLKEPDIPKANSLFVFSNTNRFRRLCYNLVNHPYFTNVVLVLILISSTMLAAEDPL-DEDKRRNYILSLFDYGFTSIFTIEILLKVVAYGLVFHQFCRNSFNLLDLLVVTVAYISIIFDKISAVKTLRVLRVLRPLRAINRAKGLKHVVQCVFVAIKTIGNIMLVTLLLVFMFACIGVQLFRGRFFSCNDSSKLYEEDCQGSKNEFHFNDVGNAMLALFTVATFEGWPKLLYVAVDSTEDDKGPVHSSRMPVAVFFFAYIIVIAFFMVNIFVGFVIVTFQNEGEQEYKNCELDKNQRNLAFALKAKPVRKYIPKQHLVWKLVTSRAFEYFIFVLIMVNTIILAMQTEAYKNVLDYMNIVFTAVFTVEFLLKIIAYKPKNYFRDYWNAFDFIIVLGSIIDIMIDMANEKKQFSINFFRLFRVMRLIKLLSRGEGIRTLLWTFIKSFQALPYVALLIVMLFFVYAVIGMQMFGKIKLSLGALNRNNNFRTFPTAVMVLFRSATGEAWQQIMMACSDTEICGNNFAYVYFISFYSICSFLIINLFVAVIMDNFDYLTR

>Pseudopodoces_humilis_Cav1.4_XP_005533405.1

----MTGVVGGKTWCPPLPP-RPFDILILATIFANCVALGVYIPFPEDDSNTSNHNLEQVEYVFLIIFTVETFLKIIAYGLVLHPSAYIRNGWNLLDFVIVIVGLFSVILEQVSDVKALRAFRVLRPLRLVSGVPSLHIVLNSIMKAMVPLLHIALLVLFVIIIYAIIGLELFIGRMHKTCFF-IGSDLESEDDPSPCAFGRACL-EGPNGGITNFDNFFFAMLTVFQCITMEGWTDVLYWMQDAMGELPWIYFVSLVIFGSFFVLNLVLGVLSGEFSREKQQLEEDLRGYMDWITQAE---ASHSTDTHASLPGSETTSVNTLWRRRCRAAVKSVSFYWTVLLLVFLNTLTIASEHHGQPPWLTETQAYANKALLSLFAAEMVLKLYALGPSCYFASFFNRFDCFVVCGGVLETAAMEPLGISVLRCVRLLRVFKVTRHWASLSNLVGSLLNSMKSIASLLLLLFLFIIIFALLGMQLFGGRFSDETQTKRSTFDTFPQALLTVFQILTGEDWNAVMYDGIMAYGMLVCVYFVILFICGNYILLNVFLAIAVDNLADGDNINSGGDKKXAQ-------RGEPQPPGPQGGVEGDQPQEEEEEEEIPEGSAFFVLSSTNPLRVKCHALINHHIFTNLILVFIILSSISLAAEDPV-RAHSPRNHILGYFDYAFTSIFTVEILLKMTVFGGFLHKFLRNWFNLLDVLVVGVSLISFGMHAISVVKILRVLRVLRPLRAINRAKGLKHVVQCVFVAIRTIGNIMIVTTLLQFMFACIGVQLFKGKFYSCTDEAKHTPGECKGLNSDFNFDNVLAGMMALFTVSTFEGWPALLYKAIDANAENQGPIYNYRVEISIFFIVYIIVIAFFMMNIFVGFVIITFRAQGESEYRNCELDKNQRQVEYALKAQPLRRYIPTQYRVWAMVNSTAFEYIMFVLILLNTIALAVQSKPFNYVMDLLNMVFTGLFTVEMVLKIIAFKPRHYFCDAWNTFDALIVVGSVVDIAVTESEDSSRISITFFRLFRVMRLVKLLSKGEGIRTLLWTFVKSFQALPYVALLIAMIFFIYAVIGMQTFGKVALQDTQINRNNNFQTFPQAVLLLFRCATGEAWQEIMLASLEEFTCGSNFAIAYFISFFMLCAFLIINLFVAVIMDNFDYLTR

>Pristionchus_pacificus_Cav3_PDM63609.1

SLFFFKQAKAPRSWALQAVMSPWFDRVTMGVILINCITLGMFRPCEDGANTYRCQMLQLADHVIFAYFAFEMCVKVIAMGF-TGPAGYLSDTWNRLDFFIVIAGCAEYVLQEYVNLTAIRTIRVLRPLRAVNRIPSMRILINLLLDTLPMLGNVLLLCFFVFFIFGIIGVQLWAGLLRNRCTLSRFYIPEDTSMEYICSQGIRCNHKNPFQGSVSFDNIGFAWVAIFLVISLEGWTDIMYYVQDAHSFWNWIYFVLLIVIGAFFMINLCLVVIATQFAGGGDSVYAAMVRFIGQTFRRTKRRRQHSLPAIEERPDEENPSSENALRDRIRAFVVCDHFTRGILVAILVNTLSMGVEYHQQPLWLTKILDISNYFFTALFAFEMILKVIADGLFGYLADGFNLFDGGIVALSVLELFQEGKGGLSVLRTFRLLRILKLVRFMPALRYQLVVMLRTMDNVTVFFGLLVLFIFIFSILGMNLFGCKFCDLCYCDRSNYDNLLHATFTTFQILTQEDWNMVLFNGMSQTTPWAALYFVALMTFGNYVLFNLLVAILVEGFQEEEQLEEEARKKADEEEEKRKHNIQPRQRLHSWGGVHVHFNPNCPVHGGRTDYSLFMFSPKNWLRIKCLQITQKKWFDYTILFFIGINCVTLAMERPSIPPMSPERVFLDISGYCFTVIFSLEMLMKVISNGCIIGEYFKDGWNILDGILVIISLVNVLFEIFGVIRVLRLLRALRPLRVINRAPGVKLVVMTLISSLKPIGNIVLICCTFFIIFGILGVQLFKGVMYHCVDIAVTTKWDCLLVNHRYNFDNLGQALMSLFVLSSKDGWVSIMYQGIDAVGVDIQPIENYNEWRMIYFISFLLLVGFFVLNMFVGVVVENFHKCKEKEMREKAREKRIQKFKRKLKRQPYWEFGPTRLFLHQVVTSKYFDLAIAAVIGINVISMAMMPMGLKYVLRALNYFFTAVFTLEAGMKLTALGFKRFFRETWNRLDMFIVILSIAGIIFEEELPINPTIIRVMRVLRIARVLKLLKMAKGIRSLLDTVGEALPQVGNLGSLFFLLFFIFAALGVELFGKLECSEDGLGEHAHFKNFGMAFLTLFRIATGDNWNGIMKDALETNCCVPILAPCFFVIFVLISQFVLVNVVVAVLMKHLEESNK

>Pristionchus_pacificus_Cav1_PDM72512.1

SLFCLNLANPLRKACIAITEWRPFEWLILFMICANCIALAVYQPYPAQDSDLKNNILEQIEYLFIVVFTIECVLKVIAQGFLMHPGAYLRNAWNSLDFVIVVIGLVSTILARMNDVKALRAFRVLRPLRLVSGVPSLQVVLNAILRAMIPLFHIALLVLFVIVIYAIIGLELFCGKLHSTCVD-QNTQQFAMKEQTPCGTAFNCN-TGPNNGITNFDNFGLAMLTVFQCVSLEGWTDVMYWVNDAVGEWPWIYFVSLVILGSFFVLNLVLGVLSGEFSREKQQLEEDLKGYLDWITQAEDIEPKNGGLAVSVGDEENEEGVETRFRRACRRLVKSQTFYWLVIFLVFLNTLVLTTEHHNQPPWLDHFQTAGNLFFVILFSLEMLLKMYSLGFTSYTRSQFNRFDCFVVISSIIEFVLMKPLGVSVLRSARLLRIFKVTKYWASLRNLVASLLNSLRSIMSLLLLLFLFIVIFALLGMQVFGGKFNPQAPKPRANFDTFIQSLLTVFQILTGEDWNAVMYNGIESFGMLVCIYYIVLFICGNYILLNVFLAIAVDNLADADSLTNAEKEEEAH-------EMDEEEEMDEGLYDEDGMEKEERELDIPKASSLFVLSHTNPFRVFCNKIINHAYFTNAVLVCILVSSAMLAAEDPL-NSESDRNQVLNYFDYFFTSVFTVEISLKVVVYGLILHKFCRNAFNLLDILVVAVSLVSFLLKAISVVKILRVLRVLRPLRAINRAKGLKHVVQCVIVAVKTIGNIMLVTFMLQFMFAIIGVQLFKGTFYACNDQSKVTERDCRGSNNDFNFDNVANAMVSLFVVSTFEGWPDLLYVAINSNEEDHGPVYNSRQSVAVFFIAFIIVIAFFMMNIFVGFVIVTFQNEGEREYENCELDKNQRKIEFALKAKPHRRYIPFQYRVWWFVTSRAFEYVIFLIIVLNTLSLACPSESFDHVLDMLNLIFTGVFAFEAFFKIIALNPKNYFGDRWNAFDFIIVLGSFIDIIYGRSPGQNFISINFFRLFRVMRLVKLLSRGEGIRTLLWTFMKSFQALPYVALLIVLLFFIYAVIGMQMFGRVALDDTEIHRNNNFHTFPMAVLVLFRSATGEAWQLIMLSCSEIPMCGNDFAYPYFISFFMLCSFLVINLFVAVIMDNFDYLTR

>Porif_osccar_Cav_ManualEdit

--------------------M------------------------------------------------------------------------------------------------------------------------------------------------------------------------------------------------------GLAMLTVFQCMTTEGWSQILYWTQDAVGKWPWVYFITLLSIGSFFVLNLVLGVLSCEFSRDRQKVANDVKQFLDWI-------ERPQDVVTSTKGEEIQASDDTRFRPLVRRLVKSRIFYWLIIALVSANTISLAAHHYGEGSFLRSFDEKSYVVFLAIFVLEMMLKLYGLGVQGYFMSNFNRFDFCIVVTCCMEFFGVRKIGISVLRCIRLLRVFKVPSYWVSLKNLVASLLNSIRSIISLLFLLFLFILIFALLGMQIFGGRFNFDYGVPRPNFDSFVSSLLTVFQILTGEDWNEVMYLGVEAYGAVAVLYFVVLVVFGNYILLNVFLAIAVDNLADAQLLTKEEELHWMERKRSREELDGAQPDEIQSIGTTPTKVDEEPNTEMPDTRSCICISPRNRLRRLCHSVVCHRYFDLT---LILICCVAMAVEDPLDAENSTKNQVLQYFDYFFTFLFSLEMLAKILVFGFVCGKYLRSFYNILDFCVVLLSVSSFILKDLEVIKVLRVFRVLRPLRAINRTSRLKTAVRCMIASVRAMGKILLVTGLLQFAFAVIGMQLFKGTFFYCSDPSKMIQEECQGRKRDFNFDNVLEATKTLFPVATFEGWPTFLYWGIDSNKEDHGPIRDSRPAVALFYVIYIVVISFFMVNIFVGFVIVSFQRVGEEEFKDSELNKNQRALEVALTAKP-SRYRPTLFALWVVATSKKFELFTLVIIGINVLVMTLQSEFYGITLEYINIVFTSLYTVECILKLVAFTPKHYFRDRWNVFDFLIVLGSLIDTLLYSGDSYEGVNLNFFRVFRAFRLVKLLRKQKGIKTLFWTFFKSLKTLPYIGIIIALIFFIYAIVGMQVFGRIKLDSTEIFRHNNFQTFFQALLVLFRAATGENWQKIMTACASPGSCGNGFSYFYFVSFIVLCSFLIINLFLAVILDNFDYLTR

>Porif_haltub_Cav_c56633_g1_i5_m.43115_Partial2088

ILCGISKKNKWRKRVKTLVNSFGFFAFISFIVIVHFIALTLYLPYINDDYEDRNELLAYINVIFILFYIVEAGLRIFAAGFILHRSSYMRSWDNALDVFVILIGIATRIVWASSEFSSLMALRLLR---ILSYFKSVQFILRAIGRSVKPLVHLGWFIFCVMMFFALIGIELTQGNLHKGCFVDKFDSVKTVRVSYPCSDSHKCDSRGPYNGMVGFDNLPVAMLTVFQCMTLEGWTSVLYLYESSVPAIAWIYFIALIIFGSLLILNLFLGVLTSVFIKEKRKLNEELYGYREW-IRTAIEDLTKSHSETKRTSSHSVEVLDAQARKKVWKLVKSHAFFWFMIVIIQLNLIILLFSHYKAPIWLDSTIYNLNYFLTGIYILETLLKVFAMGFHNYFASNFNKLSFVLCCVNVLDIVIRIPISVNAIRALRLITIFKYTRYWHDMKAVVNAFANIWSTMLSLLGLLGVYLVLVALFGMMMFRTRLT-IPNLKHPNFNTFINAILLAFQLATTEDWHVVMYSTANSYINWVGLFYVSSIMIGGLLLANIFLAIAVDKLLENEDITKRVEQMEEQRREREKQNPIQVPGNSPILKPGSTPMVINDEMGIPAHSTFFIIAPDNRLRSWCFEVVRSKAFQSIAYTVTVISSLILAVESPI-PVSSGLYCTVIYFDFAVLVWFLIEMVLKVISLGFIVHRYLRSLFNIIDFLIIIVTMIPLVVYQIPRIYIWILVRVSRPIRII-RITGLYLVCRGMISSVKRMGHMLLIGVLCLLIFAIIGVQCFKGRFFYCSDFVTEYEYDCGGKEYRLNFDNIFWSMLTLFTMLTKEGWQDVMYSAIDSTGQDRGPKYNTSQESLIYFILYMIFVTFFMLNFIVGFIIVTFQRVGIKTYTDSGLDKNQRNLYIALTAH-YKKFVPFHKFLYAIITSTVFDTVIFILVFMNTAILAIMGETLQEAIYWINLGFTVLFTMEAILKLVILTPPNYFRSFSRVFEFFVVIGSILELILQNTDFLMENGFRYSHVISAFRVLRLTLVSKSTRLVFWTVIRSIEIFPWIGVLLVLVLYMFAVSGMQIFGQIDVIDTAIHQYNNFENFFQAMKVMIRTFTGENWEKIMLSCVTTTTCGSPAAYAFFPTFFILSSILILNLFVAIIIDNFDYYVR

>Porif_halamb_Cav

ILCCLPKKNRWRKRIKTLVNSRWFNWFIALLVVAHCIALATYQTYMAGDYKPRNINLIYITVVFEVLYIIEAGLRIVAAGLIMHPSSYLRKLSNVVDVIVIVISFV-FMLRPDNRIETLSAFISLRLLRVMLPIKTIRFLLRALAKSLLPLLYVAWMTFCIMVFFALVGVEIFRGGLHSACFVSQSLSPKQYQINFPCSKSHDCAEGGPHSHLVGFDNIGSALLTVFQCMTLEGWTSVLYLYESAYGHYVWLYFILLISFGSIFMLNLLLGVLTSVFIKQKRKIKEDLEGYRQWLKKGV---LQRQLTHYGSMDSSWSDSEEDRLRSVVRHIVKSNEFFWFMIVVVLTNFILLAADFYPSPSKWKLALLIINYLFTAVYILEFVLKFYSLGPKVYFYSKFNRIDFVFTMINILDILFQVSAAVNACRAIRLVSAFKYTRHWQGMKALTSTFVDLWLIILSVLILLGSFLFIASLVGMRLFGTRLS-AYGSLYPSFFDFLDAFLLVFQLSTTEDWHAIMYTAIKEDKFITAAYYVFVIILGGFIIVNIFLAIAIDKLSEVKSYRSEQREEQRREREEQLLLPPPPPPLLSSHSQSMTPPPADVSIAIPNHSSFFILAPDNRLRVRCYTTIKSKPFQTITLTVICLSSILLALETPLVAEIGGIFCTIIFFDVFCLLWFLLEFCLKIIALGVFLHKYFHCLFNIIDFFVIVTTIVPIYYHLPAIIDVILILRALRPLRIL-RVEGLYLVTKGLVASLRKMGYVFFVGVILLLIFAVIGIQVFKGRFFYCSDLVKVKEEDCRGNRRALHFDHIGWSFLSLYTMITKEGWQDIMYHAIDSTGINTGPMYHANRANLIYFVLFMVLVTFFLINLLVGFVIVTFQQVGMKSHKEAHFDRNQENLYLALTYSPKKRYIPCHKRLYRIAKLKFFSIIIYVAVLINIIILALMDPKLTQSIYWANVGFTVFFTLEALFKLIILTPAHYFRSSERSFEFLVVIGSILEFILQTSLDLGNVMWRYSHIVSSLRVLRLIQITRNTKLIVWTVLRSLEIFPWVGVFLILLLYTYAVIGMQVFGQIEPVPSAIHQYNNFENFPQALLVMIRCFTGENWERIMLTSANRTYCGSEFSYFFYPSFFFLSSILVLNLFVAIVIDNFDYLIR

>Porif_ampque_Cav_Aqu2.38198_001

ILCCLPKKNRWRKRIKTLVYSRYFNGVIGLLVIIHCIVLATYKSYSANDLKGYNKNLLYSTIAFLLVYIVEACLRIIADGLIMHPSAYLRKIPNLVDVFVIFVSFITPFGLVIGEVAALITLRLFR---VMLQLKTVRFLFRALGSSLFPLLYVAWVTFCIMMFFALVGLELFSGGLHKACYVTDTLVMRQFQVSFPCHDGHNCSPEGPHSGLVGFDNIGIGLLTVFQCITLEGWTSILYQYESVYGSFVWIYFISLISFGSIFMLNLLLGVLTSVFIKKSRKIKEDYDGYNEWMKRGGYCHENDDGDVDDDDDQEIDEEFNKRLRSVIRHVVKSHVFFWSMIAIISINFMFLSADFYPIDKEWISSLIIINYIFTGIYIIECFLKLYSLGPRRYFTSQFNRIDFLFTLINIIDILFSVSSIVNACRAIRLISAFKYTRYWKGMRAIISTFAGVGVVILSVMALLLSFIFTASLLGMRMFGTRL-FDITHLHPTFNDFVDSFLLVFQLTTTEDWNTIMYSGLTSDEFAIIIYYLYIIIIGAFNIVNIFLAIAIDKLSEEERLEQREEERKELEEQLNALLVAPPSPLDPPTDASVARLVRKSSLEIPAHSAFFILAPDNRLRVYFYNVVKSKPFQTVTFTIIILSSLLLTLEIPV-RSESSLLCTIFFFDIFCSVWFLLEFILKIISLGAIIHRYFHCLFNIIDFFVVVTTIFPLILHSYMYIEVILVFRVLRPLR-LWRVEGLFLVTKGLFSSLRRMGYVFFVGTVLLLIFAVIGVQLFKGRFFYCTDFVSHTEEECRGKKWDLHFDNIGWSFLSIYTMITKEGWQDIMYHAIDSHDVGEGPVYNFSRWALVYHVLFMVLVTFFLINLLVGFVIVTFQQNGIKRYDEANLDRNQRNLYFSLTSVP-RKYIPFHKRLYSVINSWLWRLVINILVAINVIILSCSPSKLEVTIYWINFGFTILFTLEAVFKIIILTPPHYFRSSERGFEFLLVIGSVLELTLQSLVRDSNSIWRYSQIASCLRVLRLVQISKNTRLIVWTVLRSLEIFPWVGVLLLAVLFTYAIVGMQVFGRIKPIDNAIHQYNNFANFPQALLVMIRCFTGENWEQIMLGSVQTTTCGSPFTYFFYPSFLLISSILVLNLFVAIIIDNFDYFVR

>Pogona_vitticeps_Cav3.3_XP_020665783.1

VFFCLKQTTSPRSWCIKMVCNPWFEFVSMMVILLNCVTLGMYQPCDDMDLSNRCQILQVFDDFIFIFFAMEMVLKMVALGI-FGKKCYLGDTWNRLDFFIVMAGMVEYSLDLQNNLSAIRTVRVLRPLKAINRVPSMRILVNLLLDTLPMLGNVLLLCFFVFFIFGIIGVQLWAGLLRNRCFMPPYYQPEEDDEMFICSLGHECCNANPHKGAINFDNIGYAWIVIFQVITLEGWVEIMYYVMDAHSFYNFIYFILLIIVGSFFMINLCLVVIATQFSMEPGDCYEEIFQYVCHIVRKAKRRRCRKHNSMDHMPQGLGQPIAVEVRVKLRGIVESKYFNRGIMIAILVNTISMGIEHHEQPEELTNILEICNVVFTSMFALEMILKLAAFGLFDYLRNPYNIFDSIIVIISIWEIVGQADGGLSVLRTFRLLRVLKLVRFMPALRRQLVVLMKTMDNVATFCMLLMLFIFIFSILGMHIFGCKFSGDTVPDRKNFDSLLWAIVTVFQILTQEDWNVVLYNGMASTSPWASLYFVALMTFGNYVLFNLLVAILVEGFQAGDSYSDDDQSSSNTEDFDKFQPSIPPGHRPARKAGVTTGPNEHQDCNLREDWSIYLFSPQNRFRILCQTIIAHKLFDYVVLAFIFLNCITIALERPQIEQGSTERIFLTVSNYIFTAIFVAEMTLKVVSLGLYFGEYLRSSWNILDGFLVFVSIIDIVVSILGVLRVLRLLRTLRPLRVISRAPGLKLVVETLISSLKPIGNIVLICCAFFIIFGILGVQLFKGKFYHCLDIRITNRSDCMAVHHKYNFDNLGQALMSLFVLASKDGWVNIMYNGLDAVAVDQQPVTNNNPWMLLYFISFLLIVSFFVLNMFVGVVVENFHKCRQHQEAEEARRREEKRRRLEKKRRPYYAYCSVRLLIHSVCTSHYLDIFITFIICLNVVTMSLQPMSLETALKYCNYMFTTVFVLEAVLKLVAFGLRRFFKDRWNQLDLAIVLLSVMGITLEEALPINPTIIRIMRVLRIARVLKLLKMATGMRALLDTVVQALPQVGNLGLLFMLLFFIYAALGVELFGKLVCNDEGMSRHATFENFGMAFLTLFQVSTGDNWNGIMKDTLDDRSCLQFISPLYFVSFVLTAQFVLINVVVAVLMKHLDDSNK

>Pogona_vitticeps_Cav3.2_XP_020633697.1

--------------------M----------------------------------------------------------------------------------------------V------------------------ISTMTR--------------------------LSGS-------------------------------------------------------------------DGVG----------------------------------------------W---------------------------------------------------------------VSQPEELTNALEISNIVFTSMFALEMVLKLLAFGIWGYIKNPYNIFDGIIVVISVWEIIGQSDGGLSVLRTFRLLRVLKLVRFMPALRRQLVVLMKTMDNVATFCMLLMLFIFIFSILGMHLFGCKFGGDTMPDRKNFDTLLWAIVTVFQILTQEDWNVVLYNGMASTSSWAALYFVALMTFGNYVLFNLLVAILVEGFQAGDSDTDEDKNFEEELERLKELELLQVPGIHPSLSLSPLRPTEYPDCNTHENWSLYLFSPQNRFRASCQKVIAHKMFDHVVLVFIFLNCITIALERPDIDPHSTERVFLSVSNYIFTAIFVAEMMVKVVALGFFSGEYLQSSWNVLDGVLVFVSIIDIIVSILGVLRVLRLLRTLRPLRVISRAPGLKLVVETLISSLRPIGNIVLICCAFFIIFGILGVQLFKGKFYHCEDIRVSTKADCTNVRRKYNFDNLGQALMSLFVLSSKDGWVNIMYDGLDAVGIDQQPSQNHNPWMLLYFISFLLIVSFFVLNMFVGVVVENFHKCRQHQEAEEARRREEKRRRLEKKRRPYYAYSPARRYIHTLCTSHYLDLFITFIIGVNVITMSMQPKSLDEALKYCNYVFTIVFVIEAVLKLVAFGFRRFFKDRWNQLDLAIVLLSIMGITLEEALPINPTIIRIMRVLRIARVLKLLKMATGMRALLDTVVQALPQVGNLGLLFMLLFFIYAALGVELFGKLDCSEEGLSRHATFNNFGMAFLTLFRVSTGDNWNGIMKDTLDDKHCLPIISPVYFVTFVLIAQFVLVNVVVAVLMKHLEESNK

>Pogona_vitticeps_Cav3.1_XP_020670763.1

--------------------M--------LVILLNCVTLGMFHPCEDMGDSPRCKILQSFDDFIFAFFAVEMVVKMIALGI-FGKKCYLGDTWNRLDFFIVLAGMLEYSLDLQN--SAVRTVRVLRPLRAINRVPSMRILVTLLLDTLPMLGNVLLLCFFVFFIFGIVGVQLWAGLLRNRCFLEPYYQTENEDESFICSQGLECTEHNPFKGAINFDNIGYAWIAIFQVITLEGWVDIMYFVMDAHSFYNFIYFILLIIVGSFFMINLCLVVIATQFSSEPGSCYDELLKYLVHVLRKATKQSPMQKILETQSTGPCQSSCKIVICETFRKIVDSKYFGRGIMIAILINTLSMGIEYHEQPEELTNALEISNIVFTSLFALEMLLKLLVYGPFGYIKNPYNIFDGIIVVISVWEIVGQQGGGLSVLRTFRLMRVLKLVRFMPALQRQLVVLMKTMDNVATFCMLLMLFIFIFSILGMHLFGCKFAGDTLPDRKNFDSLLWAIVTVFQILTQEDWNKVLYNGMASTSSWAALYFIALMTFGNYVLFNLLVAILVEGFQTGDSDSEGELNLEEDCGFKNSFLQVPSLYRTGSIHSSRSSASERQDCNERDSWSIYIFAPQSKFRMICSKIISHKMFDHIVLVIIFLNCITIAMERPKIDPHSAERIFLTLSNYIFTAIFLAEMTIKVVALGLCFGEYLRSSWNMLDGVLVLISVADILVSILGMLRVLRLLRTLRPLRVISRAQGLKLVVETLMSSLKPIGNIVVICCAFFIIFGILGVQLFKGKFFVCQDTRITNKSDCAEVRHKYNFDNLGQALMSLFVLASKDGWVDIMYDGLDAVGVDQQPSMNYNPWMLLYFISFLLIVAFFVLNMFVGVVVENFHKCRQHQEEEEAKRREEKRKRLEKKRRPYYSYSRFRLLIHQMCTTHYLDLFITGVIGLNVITMAMQPKVLDEALKICNYIFTIIFVMESVFKLVAFGFRRFFQDRWNQLDLAIVLLSIMGITLEESLPINPTIIRIMRVLRIARVLKLLKMAVGMRALLDTVMQALPQVGNLGLLFMLLFFIFAALGVELFGDLECDDEGLGRHATFRNFGMAFLTLFRVSTGDNWNGIMKDTLEESTCYTVISPIYFVSFVLTAQFVLVNVVIAVLMKHLEESNK

>Pogona_vitticeps_Cav2.3_XP_020645405.1

SLFIFGEDNIVRKYAKKLIDWPPFEYMILATIIANCIVLALEQHLPEDDKTPMSRRLEKTEPYFIGIFCFEAGIKIVALGFVFHKGSYLRNGWNVMDFIVVLSGILATAGTHFNDLRTLRAVRVLRPLKLVSGIPSLQIVLKSIMKAMVPLLQIGLLLFFAILMFAIIGLEFYSGKLHRACYMNNSGKLEEMDPPHPCG-VQGCP-IGPNDGITQFDNILFAVLTVFQCITMEGWTTVLYNTNDALGTWNWLYFIPLIIIGSFFVLNLVLGVLSGEFARRQQQIERELNGYRAWIDKAEEVMRSRTEAMNRDSSDERVDISAVLLRISVRHMVKSQVFYWLVLSIVALNTACVAIVHHDQPPWLTHLLYYAEFIFLGLFLLEMSLKMYGMGPRLYFHSSFNCFDCGVTVGSIFEVVPGTSFGISVLRALRLLRIFKVTKYWASLRNLVVSLMSSMKSIISLLFLLFLFIVVFALLGMQLFGGRFNFADGTPSANFDTFPAAIMTVFQILTGEDWNEVMYNGIRSQGMWSSIYFIILTLFGNYTLLNVFLAIAVDNLANAQELTKDEQEEEEAFNQKHALSEELKTTNEAPANDADFPVPPQETNSMVPHSSMFIFSTTNPIRRACHYIVNLRYFEMCILLVIAASSIALAAEDPV-LTNSDRNKVLRYFDYVFTGVFTFEMVIKMIDQGLILQDYFRDLWNILDFIVVVGALMAFALADIKTIKSLRVLRVLRPLKTIKRLPKLKAVFDCVVTSLKNVFNILIVYKLFMFIFAVIAVQLFKGKFFYCTDSSKDTEKDCIGKRHEFHYDNIIWALLTLFTVSTGEGWPQVLQHSVDVTEEDRGPSRSNRMEMSIFYVVYFVVFPFFFVNIFVALIIITFQEQGDKMMEECSLEKNERAIDFAISAKPLTRYMPFQYRVWHFVVSPSFEYTIMAMIALNTVVLMMAPYTYELALKYLNIAFTMVFSLECVLKIIAFGFLNYFRDTWNIFDFITVIGSITEIILTDLVNTSSFNMSFLKLFRAARLIKLLRQGYTIRILLWTFVQSFKALPYVCLLIAMLFFIYAIIGMQVFGNIKLDESHINRHNNFRSFLGSLMLLFRSATGEAWQEIMLSCLENEQCGTDLAYVYFVSFIFFCSFLMLNLFVAVIMDNFEYLTR

>Pogona_vitticeps_Cav2.2_XP_020668759.1

--------------------M------ILATIIANCIVLALEQHLPDGDKTPMSERLDDTEPYFIGIFCFEAGIKIMALGFVFHKGSYLRNGWNVMDFVVVLTGILATAG----DLRTLRAVRVLRPLKLVSGIPSLQVVLKSIMKAMVPLLQIGLLLFFAIVMFAIIGLEFYMGKFHKACFS----NETGERVGFPCGEARVCE-EGPNFGITNFDNILFAVLTVFQCITMEGWTDILYNTNDAAGMWNWLYFIPLIIIGSFFMLNLVLGVLSGEFARRQQQIERELNGYLEWIFKAEEVMSKNDLIHAEEGEDHFTDVCSVMFRFFIRRMVKAQSFYWMVLCVVALNTMCVAIVHYDQPEGLTTALYFAEFVFLGLFLTEMSLKMYGLGPRNYFHSSFNCFDFGVIVGSIFEVIPGTSFGISVLRALRLLRIFKVTKYWNSLRNLVVSLLNSMKSIISLLFLLFLFIVVFALLGMQLFGGQFHFNNETPTTNFDTFPTAILTVFQILTGEDWNAVMYQGIQSQGMFSSAYFIVLTLFGNYTLLNVFLAIAVDNLANAQELTKDEEEMEEATNQKLALPGNREGEPGSKGEGGEEPHRRHRMRQILPYSSMFILSPTNPIRRLCHYIVNMRHFEMVILFVIVLSSIALATEDPV-QAESPRNEALKYLDYIFTGVFTFEMVIKMIDLGLLLHPYFRDLWNILDFIVVSGALVAFAFSDINTIKSLRVLRVLRPLKTIKRLPKLKAVFDCVVNSLKNVLNILIVYMLFMFIFAVIAVQLFKGRFFYCTDESKDLEKDCRGKKYEFHYDNVLWALLTLFTVSTGEGWPTVLKHSVDATDENQGPSPGYRMEMSIFYVVYFVVFPFFFVNIFVALIIITFQEQGDKVMSECSLEKNERAIDFAISAKPLTRYMPFQYKMWKFVVSPPFEYFIMVMIALNTIVLMMAPEPYENMLKCLNIVFTSMFSLECVLKIIAFGALNYFRDAWNIFDFVTVLGSITDILVTEADTDNFINLSFLRLFRAARLIKLLRQGYTIRILLWTFVQSFKALPYVCLLIAMLFFIYAIIGMQVFGNIALNDTAINRHNNFQTFLQALMLLFRSATGEAWHEIMLACLSKNECGSDFAYFYFVSFIFLCSFLMLNLFVAVIMDNFEYLTR

>Pogona_vitticeps_Cav2.1_XP_020656028.1

SLFLFSEDNVVRKYAKKITEWPPFEYMILATIIANCIVLALEQHLPDNDKTPMSERLDDTEPYFIGIFCFEAGIKIIALGFAFHKGSYLRNGWNVMDFVVVLTGILATVG----DLRTLRAVRVLRPLKLVSGIPSLQVVLKSIMKAMIPLLQIGLLLFFAILIFAIIGLEFYMGKFHTACKD---IHTDEITQEAPCGTSRLCP-QGPNYGITQFDNILFAVLTVFQCITMEGWTDLLYDSNDASGAWNWLYFIPLIIIGSFFMLNLVLGVLSGEFARRQQQIERELNGYMEWISKAEEVISKTDLLNPDEADDPLADISSVRLRFYIRRIVKTQAFYWTVLSLVALNTLCVAIVHYNQPDWLSNFLYYAEFIFLGLFMSEMFIKMYGLGTRPYFHSSFNCFDCAVIIGSIFEVIPGTSFGISVLRALRLLRIFKVTKYWASLRNLVVSLLNSMKSIISLLFLLFLFIVVFALLGMQLFGGQFNFDEGTPPTNFDTFPAAIMTVFQILTGEDWNEVMYHGIQSQGMMFSIYFIVLTLFGNYTLLNVFLAIAVDNLANAQELTKDEQEEEEAANQKLALMKNNKLATNETSNSHPSGHPQNPTKGMPPYSSMFILSTTNPFRRLCHYIVNLHHFEMCILTVIVMSSIALAAEDPV-QPRAPRNNVLRYFDYVFTGVFTFEMVVKMIDLGLILHRYFRDLWNILDFIVVSGALVAFAFTDINTIKSLRVLRVLRPLKTIKRLPKLKAVFDCVVNSLKNVLNILIVYMLFMFIFAVVAVQLFKGKFFYCTDESKEFEKDCRGRKYEFHYDNVLWALLTLFTVSTGEGWPDVLKHSVDATYENQGPSPGYRMEMSIFYVVYFVVFPFFFVNIFVALIIITFQEQGDKMMEEYSLEKNERAIDFAISAKPLTRHMPFQYRMWQFVVSPPFEYTIMAMIALNTIVLMMATPVYDNLLKMFNIVFTSLFSLECILKIIAFGLLNYFRDAWNIFDFVTVLGSITDILVTEGDPNNFINLSFLRLFRAARLIKLLRQGYTIRILLWTFVQSFKALPXVCLLIAMLFFIYAIIGMQVFGNISIDYPAITEHNNFRTFFQALMLLFRSATGEAWHEIMLSCLEGNECGNEFAYFYFVSFIFLCSFLMLNLFVAVIMDNFEYLTR

>Pogona_vitticeps_Cav1.4_XP_020648615.1

ALFCLRLNNPIRRAAISIVEWKPFDILILMTIFANCVALGVYIPFPEDDSNVSNHNLEQVEYVFLIIFTVETFLKILAYGLVMHPSAYIRNGWNLLDFVIVVVGLFSVILEQVSDVKALRAFRVLRPLRLVSGVPSLHIVLNSIMKAMVPLLHIALLVLFVIIIYAIIGLELFIGRMHKTCFI-IGSDLEAEEDPSPCAFGRECT-EGPNGGITNFDNFFFAMLTVFQCITMEGWTDVLYWMQDAMGELPWLYFVSLVIFGSFFVLNLVLGVLSGEFSREKQQMEEDLKGYLDWIMQAEDIESETTSVNTENVGDEEHHQDNCLFRKKCRLAVKSVTFYWIVLILVFLNTLTIASEHYMQPDWLTQIQAYANKVLLSLFTLEMLVKMYSLGLQAYFVSFFNRFDCFVVCGGILETVIMEPLGISVLRCVRLLRIFKVTRHWASLSNLVASLLNSMKSIASLLLLLFLFIIIFSLLGMQLFGGKFNDETQTKRSTFDTFPQALLTVFQILTGEDWNAVMYDGIMAYGMLVCVYFIILFICGNYILLNVFLAIAVDNLADGDNINTSKEEEQKEKKRKKKK-VVKNKWPSRKSQSKSSFYSPAQTGEIPDGSAFFCLSKTNPLRVGCHKLIHHHIFTNLILVFIILSSISLAAEDPI-RAHSFRNIILGYFDYAFTSIFTVEILLKMTAYGAFLHQFCRNWFNLLDLLVVSVSLISFGIHAISVVKILRVLRVLRPLRAINRAKGLKHVVQCVFVAIRTIGNIMIVTTLLQFMFACIGVQLFKGKFYSCTDEARHTPKECKGLNSDFNFDNVLSGMMALFTVSTFEGWPALLYKAIDANAENHGPIYNYRVEISIFFIVYIIIIAFFMMNIFVGFVIITFRAQGEQEYKNCELDKNQRQVEYALKAQPLRRYIPYQYKFWYIVNSTGFEYIMFVLILLNTIALAVQSQPFNYVMDLLNMVFTGLFTIEMVLKIIAFKPKHYFVDAWNTFDALIVVGSVVDIAVTESEDSSRISITFFRLFRVMRLVKLLSKGEGIRTLLWTFIKSFQALPYVALLIAMIFFIYAVIGMQTFGKVAMQDTPINRNNNFQTFPQAVLLLFRCATGEAWQEIMLASLEEFTCGSNFAIVYFISFFMLCAFLIINLFVAVIMDNFDYLTR

>Pogona_vitticeps_Cav1.3_XP_020659553.1

ALFCLSLNNPIRRACISIVEWKPFDIFILLAIFANCVALAVYIPFPEDDSNSTNHNLEKVEYAFLIIFTIETFLKIIAYGLLLHPNAYVRNGWNLLDFVIVIVGLFSVILEQLTDVKALRAFRVLRPLRLVSGVPSLQVVLNSIIKAMVPLLHIALLVLFVIIIYAIIGLELFIGKMHKSCYF-DDTDILAEEDPAPCAFGRQCP-PGPNGGITNFDNFGYAMLTVFQCITMEGWTDVLYWVNDAIGEWPWIYFVSLIILGSFFVLNLVLGVLSGEFSREKQQLEEDLKGYLDWITQAEDIDSETESVNTENVGGDGENPPCCFNRRRCRAAVKSVSFYWLVIVLVFLNTLTISSEHYDQPEWLTQIQDIANKVLLALFTCEMLVKMYSLGLQSYFVSLFNRFDCFVVCGGIVETIIMSPLGISVFRCVRLLRIFKVTRHWTSLSNLVASLLNSMKSIASLLLLLFLFIIIFSLLGMQLFGGKFNDETQTKRSTFDNFPQALLTVFQILTGEDWNAVMYDGIMAYGMIVCVYFIILFICGNYILLNVFLAIAVDNLADAESLNTAQKEEAEEKEKKKAASKVTINDYGEGEDEDKDPYPPCDVPGIPEGSSFFLFSNTNPIRVGCHRLINHHIFTNLILVFIMLSSASLAAEDPI-RSHSFRNNILGYFDYAFTAIFTVEILLKLTVFGAFLHKFCRNYFNLLDLLVVGVSLVSFGIQAISVVKILRVLRVLRPLRAINRAKGLKHVVQCVFVAIRTIGNIMIVTTLLQFMFACIGVQLFKGKFYRCTDEAKQNPVECRGQNSDFNFDNVLTAMMALFTVSTFEGWPALLYKAIDSNGENIGPIYNYRVEISIFFIIYIIIIAFFMMNIFVGFVIVTFQEQGEQEYKNCELDKNQRQVEYALKARPLRRYIPYQYKFWYMVNSTVFEYIMFVLIMLNTLCLAMQSKLFNDAMDILNMVFTAVFTVEMVLKLIAFKPKGYFSDAWNTFDSLVVLGSIVDIVLSESEDSARISITFFRLFRVMRLVKLLSRGEGIRTLLWTFIKSFQALPYVALLIAMLFFIYAVIGMQVFGKVALKDSQINRNNNFQTFPQAVLLLFRCATGEAWQEIMLACMEEYTCGSNFAIIYFISFYMLCAFLIINLFVAVIMDNFDYLTR

>Pogona_vitticeps_Cav1.2_XP_020644644.1

ALLCLTLKNPIRRACISIVEWKPFEIIILLTIFANCVALAIYIPFPEDDSNATNSNLERVEYLFLIIFTVEAFLKVIAYGLLFHPNAYLRNGWNLLDFIIVVVGLFSAILEQATDVKALRAFRVLRPLRLVSGVPSLQVVLNSIIKAMVPLLHIALLVLFVIIIYAIIGLELFMGKMHKTCYVGILSDTPAEEEPSPCA-GRQCQ-EGPKHGITNFDNFAFAMLTVFQCITMEGWTDVLYWVNDAIGKWPWIYFVTLIIIGSFFVLNLVLGVLSGEFSREKQQLEEDLKGYLDWITQAEDIDNMSMPTSETES-VNTDNVTGGFCRRKCRAAVKSNVFYWLVIFLVFLNTLTIASEHYNQPDWLTEVQDTANKVLLALFTAEMLLKMYSLGLQAYFVSLFNRFDCFIVCGGILETIIMSPLGISVLRCVRLLRIFKITRYWNSLSNLVASLLNSVRSIASLLLLLFLFIIIFSLLGMQLFGGKFNDEMQTRRSTFDNFPQSLLTVFQILTGEDWNSVMYDGIMAYGMLVCIYFIILFICGNYILLNVFLAIAVDNLADAESLTSAQKEEEEEKERKKLANKGNTDEYQPNENEEKNAYPTTETPGMPEASAFFIFSPSNRFRVHCHRIVNDNIFTNLILFFILLSSISLAAEDPV-QHYSVRNQILFYFDIFFTVIFTIEIALKMTAYGAFLHKFCRNYFNILDLLVVSVSLISFGIQAINVVKILRVLRVLRPLRAINRAKGLKHVVQCVFVAIRTIGNIVIVTTLLQFMFACIGVQLFKGKLYSCSDSSKQTEAECKGENSKFDFDNVLTAMMALFTVSTFEGWPELLYRSIDSHMEDVGPIYNHRVEISIFFIIYIIIIAFFMMNIFVGFVIVTFQEQGEQEYKNCELDKNQRQVEYALKARPLRRYIPYQYKVWYVVNSTYFEYLMFVLILLNTICLAMQSCLFKEAMNILNMLFTGLFTVEMVLKLIAFKPKGYFSDPWNVFDFLIVIGSIIDVILSEAEENSRISITFFRLFRVMRLVKLLSRGEGIRTLLWTFIKSFQALPYVALLIVMLFFIYAVIGMQVFGKIALDDTDINRNNNFQTFPQAVLLLFRCATGEAWQEIMLACLGEHSCGSSFAVFYFISFYMLCAFLIINLFVAVIMDNFDYLTR

>Pogona_vitticeps_Cav1.1_XP_020660962.1

SLLCLTLQNPVRKACIAIVEWKPFETIVLLTIFANCVALALYLPMPEDDTNKMNSRLEKLEYFFLIVFAIEATLKIIAYGFLFHADAYLRNGWNVLDFTIVFLGVFTVILEKISDVKALRAFRVLRPLRLVSGIPSLQVVLNSIVKAMLPLFHIAVLVVFMLTIYAIMGQELFKGKMHKTCYYTDIIATVENEKPSPCTSGHQCT-PGPNNGITHFDNFGFAMLTVYQCISMEGWTQVLYWVNDAIGEWPWIYFVSLILLGSFFILNLILGVLSGEFTREKQQLDEDMKGYMDWIVHAEVMERGEGMMSEDEGGSETESLYELLFRRKCREVVKSRFFYWLVILIIALNTFSIASEHHNQPDWLTQAQDVANRVLLALFTVEMILKMYALGLRQYFMSLFNRFDCLVVCTGILEIISMSPLGISVLRCIRLLRLFKITKYWRSLNNLVASLLNSVRSIASLLTLLFLFMVIFALLGMQLFGGKFDDDVEIRRSTFDNFPQALITVFQVLTGEDWTSVMYNGIMSYGMLVCIYFIVLFVCGNYILLNVFLAIAVDNLAEAETLTSAQKAKAEEKKRKKLAAKLKVDEFESNVNEIKDPYPSADFPGMPESSAFFIFSPTNKIRVLCHRIVNATWFTNFILLFILLSSISLAAEDPI-RAESFRNKILGHFDTGFTTVFTVEIVLKMTAYGAFLHKFCRNYFNILDLLVVAVSLISMGLEAISVVKILRVLRVLRPLRAINRAKGLKHVVQCVFVAIKTIGNIVLVTFLLQFMFACIGVQLFKGKFFYCTDTTKITESECWGLQNEFHFDNVFSAMMSLFTVSTFEGWPKLLYRAIDTHTENMGPIYNYRMGIAIFFIIYLILIAFFMMNIFVGFVIVTFQEQGETEYKDCELDKNQRQVQYALKARPLRCYIPYQYQIWYLVTSSYFEYLMFFLIMLNTVCLGMQSETMNQVSDVLNVVFTILFTVEMIVKLIAFKAKGYFGDPWNVFDFLIVIGSIIDVILSQPDETGRISITFFRLFRVLRLVKLLSRGEGIRNLLWTFIKSFQALPHVALLIVMLFFVYAVIGMQMFGKIALVDTQINRNNNFQTFPQAVLLLFRCATGEAWQEIMLASYEEYSCGSGFAYFYFISFYMICAFLIINLFVAVIMDNFDYLTR

>Placo_triadh_Cav3_evg954676

VCWYLHKDQWPRKYFVKLSQWRWFERITIIVILINCVTLGSYNPAGQFKVDSTCQITSVVDNIIFGYFVVEMIIKMVALGV-FGKYAYFSSGWNRLDFIIVLTGCLEYLINEGEFLTIIRTVRVLRPLRAINRVPSMRLLVNLLLDTLPLLGNVLMLCFIVFSIFGIVGVQLWKGILRSRCTLLYRYYLPSYEDPYVCSLDLTCKGPNIFYDNISFDNFAMAFIAIFQVITLEAWVDIMYAIQDGHASIDWIYFVILILIGSLFLLNFTLVVMATQFSAESRTCYNETIRYIAYLIQQFYSRRQRSYSQNTTTTSQIYQESALRLRVRCCIFTKSQKFSLIVLFAILANTIVMAIEHHNQPTYQIQALEVCNIIFTIFFTLEMVFKLFALGLLRYAKDSFNVFDAIIVIVSLVEIA-TDGKGLSVLRSFRLLRIFKIVRFLPTLQRQMMVMAQTFDNVVIFLGLLFLFMFTFSILGMHLFGNRFCGPVVCSRKNFDSLLWAFVTVFQILTQEDWNVVMYDGMLARGKWAAIYFLALVTLGNYVLLNLLVAILVNGFQE---------QEKDEKNRKKDLLPNLVDPNEVQIFPCRVPIGPRDPPQRHRNHSLLILPKENRFRKWCKNVVKNPYFDRIILVVIIFNCVTLAMERPGIDPNSMERHFLNLMVIIFTFIFTSEMIIKVLALGLVTGDYLRNGWNVLDLLLVIISWVDLIITILGVLRILRGFRTLRPLRVINRAPGLKLVVQTLFSSLKAIGNIVIICVAFFVIFGILGVQLFSGKFYYCKATDVENKTQCVEVNRPYNFDDLVNASLSLFVISSKDGWMDITYHGIDARGVDLQPKKNHNVVVLLYFISFLLLVGFFVLNMFVGVVVENFHKCQERRQREKKRKANAKKVIIKKKVKEKKKHIPWRRKLYRFCVHRYFDITITIVIAVNIIFMATMSQAWEEVHKYANYFFTVVFTLEAVIHLVAFGVVYYFRDRWNIFDLLIVILSWTGIIIESNPTINPTIIRVMRLLRIVRILKLIKAAKGIRALLRTIMNAMPQVVNLGMLFFLLFFIFAALGIELFGRLDCTTNGLSQHANFKTFGMAMLTLFRIATGDNWQGILQDTLQQNCCSPILSSLFFVIFVLAAQYVLTNVVVAVLMKHLEESDE

>Placo_triadh_Cav2_evg1041627

SLCCLPANNLLRKYAKKLVDWTPFEYLVILTIVANCVVLAMDVPLPDNDSTEISLLLRNAEIGFLVVFCIEAALKIIAKGFFFHPQAYLRSGWNILDFLIVVVGLVNAFYINAADVKILRVVRVLRPLKLVSGMPSLQIVLRSLLTAMGPLFQISLLVLFVIVIYSIIGMEFFLGKFHLGCRD-PRTGQLLTQNHSPCNNGFRCVYDGPNYGITGFDNIFMALLTVFQCISLEGWTNLLYDTNNAVGTFTWFYFLTLIIWGSFFMLNLVLGVLSGEFAQENRKIERDFLGYLEWIGRAEDLIFDTRSEIFQFAEQDQVAELAEFLRLAIRRTVKSRPFFWIVILLVFLNAVTIASEHSGEPLWLKDFREATNIVFVALFTLELILKLYGLGAVFYFSSTFNCFDFAGVIASIAELKGGPKLGIRGFPCIRLLRIFEITKHWKSLSNLVASLISSLRSILSLLFLIGLCIMVFALLGMQLFGGRFNFAEGVPRSNFNDFGHAVLSVFQVLSGEDWNEVMYNGIRAYRYAVSLYFVVLVCLGNYTLLNVFLAIAVDNLTKAQEISKDEDEEIMLQKRLSIKNTSQQETRIDEINDIENPGNENDDEEILQVNSLFIFSPQNRFRRFCHYIVHLRHFENFMIAAIIISSGLLAVEDPM-NEDPVLNYVLRIFDSIFTGIFLIELILKVVDFGFILHRYCRNLWNILDMIVVVTAVTSFIYFNISAIKAIRTLRVLRPLKAIRTAKKLLACFQCMVNSLKNVLNVCIVMLLFLFMFAVIGVQLFKGKFHYCTDQTKHTKSQCRGEQHPYNFDDLPRAMLTLFTMSTAEGWPRILYWSIDATNENEGPMRDYNLAVALYFCIYIVVFPFFFINIFVALIIVTFQEEGDKDIANYQLNRNQRDIEFALNAKPIHRHMPYAYKVWRIVTSTPFEFIIMVLIIVNTIVLMMQSKDYKDMLQIINITVTILFTVEMLLKVIAFSPRNFIKEWWNIFDLIVVIGSWTDIIITYINGTSTVSISFFRLFRAGRLIKLLRKGYTIRVLLWTFLKSFQALPYVGLLIGMLFFISAVLGMQLFGQIQSDPTAIFRYNNFQTFTGALIVLVRCSTGENWPEVMLACLKFPDCGSYVAYPYFVIFVFLSTFLMLNLFVAVIMDNFKYLTR

>Placo_triadh_Cav1_evg1032956

VLFFLKTDNPIRQFATTVVEKKAFEYLILFTIFANCVALALYQPLPNNDNTLLNENMEKVEYVFLAIFTIESFLKIITYGFAIPSGAYLRNGWNILDFIIVIVGIINIIFTATSDVRALRAFRVLRPLRLVSGVPSLQVVMNAIMKAMVPLFHVAALVVFVIIIYATIGLELFNGVLHRACYH--NITKKLINDPRPCAAAHHCK-TGPNAGITSFDNIGLSMLTVFQCITMEGWTNIMYSINDAVGEWPWIYFVTLIILGSFFVLNLVLGVLSGEFSREKRQIEEDYRGYLEWIGKAEDLEQVDMEQQVAAEGVRFDDVEHGRSRRYCRLVVKSQTFYWLVILAVLLNTICLAVEHYEQQRVVTQFLSITNSVFVGLFTIEMSIKMYALGIEGYFMSLFNRFDFLVVLVSIIELIGAAALGLSVLRCVRLLRIFKITRYWNTLRNLVASLLNSMRSIASLLLLLFLFVLIFALLGMQIFGGRFNFNKSIPRSNFDSFWQSLLTVFQILTGEDWNEIMYNGIKALDILAVFYFIILVVVGNYILLNVFLAIAVDNLANAESLTEINERRNAKRKMAKEQYKMAVFSETESELSRVEELDSNSDCTIPKATSMFLFKSTNRFRCKIFDFVTNLYFSNIILIIIILSSITLAAEDPL-GKDKIRNQVLSYCDKTFTAIFCVEAAMKMIAFGVIMHEYFRNIFNILDMIVIAVSIADYTIANLKQLKVLRVLRVLRPLRALNRARGLKHVVQCVFVAIKTIWSIMLVTLLLVFMFAVIGVQLFKGRFYFCTDASKMDNSTCKGKESSFNFDNVPNAMMTLFTITTFEGWPSILYRAIDATEAGRGPSRNHQPLVAIYFVIYIIIVAFFMVNIFVGFVIVTFQTEGEQEYRNCDLDKNQRNIEFALKTKPQKRYIPLQLAVWKLATSLGFEYTIFGLITLNTMTLMMRPRVYNDVLEYLNIGFTVLFGLEAILKIVAFKPQNYFRDKWNVFDFVIVVGSIIDIVISESNVTVDFSVNFFRLFRAMRLVKLLSRGGGMRTLLWTFMKSFQALPYVGLLIVFVFFIYAVIGMQLFGTVLTSPSAITEYNNFHSFFSSVLVLFRCATGENWQLITLSCTSENSCGSNISYLYFSSFYVLSSFLVINLFVAVIMDNFDYLTR

>Paramecium_Cav1c_GSPATP00010323001

RKKILISTKLVSFYAQLVTTHPIFEVITLIMIVFNSVMLAIDDPT----TNVQSPFQNLTDLIFLAYYTFEAVLKIVAQGFIIPKKSYLRDTWNILDFSVIITAYIPYFL----NLNALRSFRVLRPLRTVSSIKALRTILLALFASIAQLRDAAVVLMFFYSIFAIAGVQLFSGYLKRRCIGEESGITWVSEEILFCADDNNCPIANPQNNLINFDTFGYAFLQVFIITTLEGWTQIQTAVMLTFSQFVVLYFIIVVLVGAFFLVNLTLAIIKLNFKYVPEREKVDTAGFITWIMNRRKHTGNSESDEDNENENNIDRDQNNVLQNQLLYFVQSGYFEASMNLAVALNTVILALDGLL-PDSTSEITNQFNFGFTILFTIELGLKMLGMGPRKYLRDTMNIFDAVIVALSLVELFGKSSLSVRIFRAFRVLRVTKLMRSLQFMGFLIKVLSNAFQSFMYIMILLLLFIFIFTLLGMAFFGGQ--LSKTPSRQSYDDIQSAFLVVFQVLTLENWNSILWDLLVQDVFITIPYLVFWIMIGNYVFLNLFLAILLENFEE--EYKNDKAGLDTN-------AVNSTSTLKSTMKTKKHTIAQQLENNGMCQYSLYLFSQQNIVRKICYRIVKDDKFETLIFTMIFLTSAKLVFDTYI-PDTGQLKEVSLDIDIFFAAFFGVEMCMKIIAFGFISQEYLRESWNVLDFFIVIASFIDVSVSNLSFVKILRLLRTLRPLRFITHNRSMKILVSALLQSINGIFNVAIVVILVWMMFAILGINLEKNKMSFCNDDEHYGVQECKEENRKTNFDNILNGMLTLFILSTLEGWPDMMYWFID--ADESGPIKAAQLQFSWYFIVFILFGSILLMNLFIGVILVNYH-LAEEASRDKILTQPQVDQKLIVHSSPLAMFSPFRAKIFTIIKHRYFDPTILMIIVLNIIIMGLSPILYDQVLTQFNTAFTFVFIGEAILKIIALGPVGYMRNSWNQFDFFVVCASILDLILS-FTGNSFISFLIFRVLRVTRLFRLIKSFEGLQKLIETAIYSLPAMLNVTALLFLVFFIFSILGVFLFGTIKS--WVIDDTNNFSDFHHSIELLFRCATGEDWYKVMFDTMQGYYCI------FFIIFIVIQQYIMLNLFILIILDQYEYFNS

>Paramecium_Cav1b_GSPATG00033414001+33415001

RKQILIFTKLISSYAQLITTHPLFELMTLLMIIFNSAMLALDDPTTDVQTSFQ----DLTDIIFLAYYTAEAVLKIVALGFIFPKKAYLKDTWNILDFSVIVTAYIPYFL----NLNALRSFRVLRPLRTVSSIKALRTILLALFASIAQLRDAVVVLIFFYSIFAIAGVSLFSGYLKRRCIGEMSGITWISDEILFCADDNNCPIANPQNDLVNFDTFGYSFLQVFIITTLEGWTQIQTAVMLTFSQYVVLYFIIVVIVGAFFLVNLTLAIIKLNFKYEPERYIVDTKGYTTWIMKRRDNSNVNENSDSDDEPEENNEKNNNVCQNHLLYFVQSSYFEAAMNLAVALNTVILALDGLL-PDSSANTLNQFNLGFTILFTIELGLKLIGMGPKNYISDTMNIFDAIIVALSLVELFGKSSLSVRIFRAFRVLRVTKLMRSLQFMGFLIKVLGNAFQSFMYIMVLLVLFIFIFTLLGMAFFGGQ--LSKTPSRQSYDDIQSAFLVVFQVLTLENWNSILWDLLIQDVFITVPYLVFWIMIGNYVFLNLFLAILLENFEE--EYKNDKAGLDTN-------AVNSTSTLKSQMKTKKATVAKNLENNGICQFSLYMFSQENIIRRICYRIVKDDKFETLIFFMIFLTSTKLVFDTYI-PDTGQLKETSLQIDIFFAVFFGVEMIMKIIAFGFVQQEYLRESWNILDFFIVIASFIDVSVSNLSFVKILRLLRTLRPLRFITHNRSMKILVSALLQSINGIFNVAIVVILVWMMFAILGINLEKNKMHYCDDDEHYGPEECAQANRKVNFDNILNGMLSLFILSTLEGWPDQMYWFID--ADESGPIKGAQLQFSWYFIVFILVGSILLMNLFIGVILVNYH-LAEEASRDKILTQPQVDQKLIVHANPLAMFSPFRAKVFIIIKHRYFDPTILMIIVCNIVTMGLSPIAYDSILQSLNTAFTFVFITEALLKIIALGPVGYMRNSWNQFDFFVVCASILDLILQFFLSAGPQLARVFRVLRVTRLFRLIKSFEGLQKLIETAIYSLPAMLNVTALLFLVFFIFSILGVFLFGSIRS--WAIDDVNNFSDFHHSFELLFRCSTGEDWYKVMFDTMQGYYCI------FFIIFIVIQQYIMLNLFILIILDQYEYFNS

>Paramecium_Cav1a_GSPATP00010323001

RKQILIFTKLISSYAQKITTHPLFELMTLLMIIFNSAMLAIDDPT----TNVQTSFQDLTDIIFLAYYTAEAVLKIVALGFILPKKAYLKDTWNILDFSVIVTAYIPYFL----NLNALRSFRVLRPLRTVSSIKALRTILLALFASIAQLRDAVVVLIFFYSIFAIAGVSLFSGYLKRRCIGEMSGITWISDEILFCADDNNCPIANPQNDLVNFDTFGYSFLQVFIITTLEGWTQIQTAVMLTFSQYVVLYFIIVVIVGAFFLVNLTLAIIKLNFKYDPQRPIVDTDGYISWIMNRRKNSENENSDSDDEPDEQNNEKNNNVCQNHLLYFVQSGYFEAAMNLAVALNTVILALDGLL-PDSSANTLQQFNLGFTILFTIELGLKVIGMGPKNYISDTMNVFDAVIVALSLVELFGKSSLSVRIFRAFRVLRVTKLMRSLQFMGFLIKVLGNAFQSFMYIMVLLLLFIFIFTLLGMAFFGGQ--LSKTPSRQSYDDIQSAFLVVFQVLTLENWNSILWDLLIQDVFITVPYLVFWIMIGNYVFLNLFLAILLENFEE--EYKNDKAGLDTN-------QVNSTSTLKSTMKTKKHTVAQQLENNGLCQFSLYLFSQENIVRRICYRIVKDDKFETLIFFMIFLTSTKLVFDTYI-PDTGKLKETSLQIDIFFAVFFGVEMIMKIIAFGFVQQEYLRESWNILDFFIVIASFIDVSVSNLSFVKILRLLRTLRPLRFITHNRSMKILVSALLQSINGIFNVAIVVILVWMMFAILGINLEKNKMHYCDDDEHYGPDECAKANRKVNFDNILNGMLTLFILSTLEGWPDQMYWFID--ADESGPIKGAQLQFSWYFIVFILIGSILLMNLFIGVILVNYH-LAEEASRDKILTQPQVDQKLIVHANPLAMFSPFRAKVFIIIKHRYFDPTILMIIVCNIVTMGLSPIAYDNALQSLNTAFTFVFITEALLKIIALGPVGYMRNSWNQFDFFVVCASILDLVLQ-FTGNSFISFLIFRVLRVTRLFRLIKSFEGLQKLIETAIYSLPAMLNVTALLFLVFFIFSILGVFLFGSIRS--WAIDDVNNFSDFHHSFELLFRCSTGEDWYKVMFDTMQGYYCI------FFIIFIVIQQYIMLNLFILIILDQYEYFNS

>Orbicella_faveolata_Cav3b_XP_020626608.1

SCFIISKDSKGRRWMIQLIKWPWFERISMFVILINCVTLAMYNPLDPQCISTRCQVLEHVEHVVFAFFFAEMVIKMLAMGV-TGKKGYLQDKWNRLDCFIVVIGLIEKVIKSGNYLTIMRAFRVLRPLRAINKVPSIRILVTLLLDTLPMLGNVLLLSFLIFFVFGIIGVQLWQGKLRNRCFTRSTFYQPSFYFPFVCSSGLECHGPNPFYDTTSFDNIGIAWIAIFQVITLEGWSDIMYFVQDAHSNWNWIYFVVLIVMGSFFLVNLCLVVITMQFQQEKEHVHHHHHIYHHHVANVPKIQGISSYNSSEHFDEDVAELSKNKLRTLCRKITESKQFTFVIMSAILLNMICMGLEHYQQPERLTVALETVNIIFVSIFAVEMIIKLLGFGVTAYVSQGQNVFDGFIVIVSVCEIL-QGNSALSIFRSIRLLRIFKLVR---PVRYQLLVVVKTMTSVMTFFGLLFLFIFAFAILGMNLFGGKFKGKQVVSRSNFDNFLWAMVSVFQILTQENWNQVMYDGMRSTRKWAALYFIALMAVGYYVLFNLLVAILVEGFTNGKSPKPKEETGGKTLGEPVVRDTALESERHSEASSKRTPELTWKIESIRSDWSLFLFSPSNRFRRFLTAVCGHKYFDYAVLFFILISCVVLALEEPNIPSDHQKRRIIDIAMLTLTIIFSLEMMIKIIAHGLVLGPYLKDGWNVLDGILVLFSWIDVIITVLGALRVFRALRTLRPLRMIRRAPGLKLVVQTLLYSLKPIGNTVLIAAIFFVMFGILGVQIFKGTFHYCERGHVTNKSECQNENHMYNFDNLPNALVSLFVFSTRDGWVEIMHNGIDAVGIDKQPIKNYREWRLAYFIPFLMLGGFLVLNMIVGVVVENFQRCREKLEDE---EKQRRR-------------------------------------------------------------------------------------------------------------------------------KLLEK-----------------------------------------------------------------------------------------------------------------------------

>Orbicella_faveolata_Cav3a_XP_020622701.1

AFFFLHREKRPRKWFIRLITWPYFERLSILVILVNCVTLGLYDPFDPECKTQRCQTLDAMEKVIYTFFLAEMLCKWIAMGL-FGKLAYFAESWNRLDCFIVAAGTFELLYNQGEEYSAVRAIRVLRPLRAINRVPSIRILVTLLLDTLPMLWNVLAICFFIFAIFGIVAVQLWRGALRGRCFLLTDFYVPSFDEPFICAMGRTCTGDNPSWGAIGFDNIFIAWVAIFQVITLEGWTDIMYYVQDAHGLWNWIYFVILIVIASYFMTNLCLVVITTQFQYGKDGCWVEILKYIEHLIRRFKRRPGGEMITTATAAVAINGKSAVRFRHMCRNSVDSKWFMYIIMGAIFLNTLSMGIEYHGQPLKMTEVLEILNYIFTAIFGFEMLLKLMGLGPYGYIKDPFNLFDGFIVVMSIVELFGGGDSSISVLRSFRLLRIFKLVRFLPALRRQLLVMIHTMDNVMTFLALLVLFIFTASILGMNLFGGKYMGVETAARANFDDLFWALVTVFQVLTQEDWNTVMYDGMRATSKWAALYFILLMTIGNYILFNLLVAILVEGFANSSSVKSRSYSIKHDGDYETCKVTRKTLSFGDTKTMVIANPAQAERCIKRRNWSLFLFSPSNRFRTWMVTVYRNKWFDRVVLVFILLNCIVMALERPDLPSDSTMKKVITICMYIFLAIFTLEMIIKVIALGFWIGRYMRSAWNVMDGFLVIVSWVDVIVSILGVLRVFRALRTLRPLRVISRAPGLKIVVETLISSLKPIGNIVLIAATFFIIFGILGVQLFKGKFHYCKEAEVITKEDCLHENKEYNFDNLAKALLTLFVFSTKDGWVTIMYDGIDAVGIDKQPIRNNNKWNVLFFVAFLLLAGFVVLNMLVGIVVENFQKCREKDRLAEKQKKKLKNVAFLLSAGKEGGFPQPRRFFHRICTHGYFDLGISAVIVLNVICMAMQPEDMSVFLKYANYVFTAIFIVEGVLKIYALRFRKYIKERWNQLDLFIILLSIVGIVLEEEVPINPTIIRVMRVLRIARAS-----------------------------------------------------------------------------------------------------IPSYIMVTQ--------PSLVENTNNVNN

>Orbicella_faveolata_Cav2c_XP_020613420.1

ALFCLPEDNPVRFYCKKIVESKKFEYFILLTIAANCVVLMLEEPLPNGDTTDRNKKLEESEKYFVIIYCIEAATKIIANGFLLHKDAYLRNGWNILDFVVVVVGLVGMISDLEISLKVLRAVRVLRPLKIVSGIPSLQVVMKSIARAMIPLLQILFLILFVIVIYAIVGLELLHGKFHLTCYDITGELDTKFTSPRVCSPGRPCE-SGPNKGISTFDNIFLSMLTVFQCITMEGWTDIMYHSYDARDVVTSIIYISLIIIGSFFMLNLVLGVLSGEFARSREKIERQVNAYTDWIGRAEDILLKKRYSLSDSIMHLIEDHGEMLLRIQVRHMVKSQVFYWSVIVCVFLNTVLMSVEHYGQPDWLEKFQEISEYVFLSIFIAEMLLKMYGLGPRVYFKSAFNRFDCAVVLGGIIEIVTDYSFGISVLRSLRLLRIFKFTRFWASLRNFVTSLLNSMRSILSLIFLLFLFIFIFALLGMQLFGGKFSERHDAPRTNFDNFLKAMLAVFQIMTGEDWNAVMYDGIVASKMLSSLYFVSLVILGNYTLLNVFLAIAVDNLANAQAVTQDEKEEQRQLEAMRKKKISQDGKFPQATNGQMSNGNSKENPDIIRKSSMFIFGPDNPIRQACHWVVNLRYFDDFILAVILISSVLLAIEDPV-HPDASRNKVIRYFDYGITGIFALEVLVKMIDLGVILHKYLRSGWNVIDAFVVGCNIAALLLDQKDAIKSFRVLRVLRPLKAINKSKKLKAVFECMMYSLKNVRNILLITLLFYFIFAVVGVQLFKGKFWYCTDLSKMTNATCQGKKHKFNFDNVPHAMLALFSSSTGEGWPQGMYHTVDATKEDQGPIKDYQIQMSLYYVCFVVVFSFFFLNMFVALIIVTFQEQGEKEMDGCELDRNQRDIQFAMTAKPRQRYMPCVYKVWKVVDSKPFEIFIMATIVLNAIVLMVASPQYERILINLNAAFTFVFLSEAILKLIAFRQ-NYFRDFWNVFDFIIVITTLVGVILELASDALPVDPSFFRLFRAARLVKLLRQGYTIRILLWTFLQSFKALPYVVILLGMLFFVYAVIGMQLFGRIRPSEKQINHHNNFRNFFMALQVLFRASTGENWHKIMLDCFENRSCGTVASIIYFCTFYFFCTFLMLNLFVAVIMDNFEYLTR

>Orbicella_faveolata_Cav2b_XP_020612369.1

ALFCLSETNPIRELSKSIVVSKVFEYFILLAIGANCIVLALNTPLPNNDRTDMAQQLEDAEYYFVGIFCVEALLKIMAFGFVLHPGSYLRNGWNILDFVVVVVGIIEVNKRLNLDVKALRAVRVLRPLKLISGVPSLQVVMKSIVRAMVPLLQILLLVLFCILIYAIIGLEFLKDKFHTTCFD---IETGARQTNKPCDAGRDCL-IGPNDGITTFDNIALAMLTVFQCITMEGWTSIMYKTFDAMDYLYACYYVSLIVIGSFFVLNLVLGVLSGEFARRQQQLNRQVDAYMSWIAKAAESASQSQLVTNDMVSNPGRVVARRRFSIKVGRMVKTQAFYWTVLVCVFLNTIVLAVEYYNQPKWLTEFQKYAEIVFLTFFFVEMVLKIYGLGFHVYFSSSFNCFDCAVVCSGFLDLIEGIKLGISVLRCLRLLRVFKVTRHWRSLRNLATSLVSSIKSIVSLIFLLFLFILIAALLGMQIFGGKFT---KQPQTNFDNFQNAMLAVFQILTGEDWNSVMYSGVMALGILASLYFVLLVILGNYTLLNVFLAIAVDNLANAQILTEDEENEKQERELNRARTRSEGGDRTRRIRRMRRGRNEVNGDTIIKTKTLFIFGPENRFRRLCHRIVNLRHFDNFMLVIIMLSSISIAIEDPV-NDNSKRNEVLMYFDYVFTAIFALEVIIKIVDVGVIFHKYFRDWWNVIDALVVSFNMASLILVGRSVIKALRVFRVLRPFKGVHKIKKLQAVFRCMWYSVKNVANILMITALFLFIFAVMGVQLFKGKFQNCNDESKLWKEECQGETKALNFDNVFKAMLTLYTSSTGEGWPSAMQATMATTEVDKGPIPNYSPGYALYYISFVVVFSFFFLNIFVALIILTFQQEGEREIASCELDRNQRDIQFALTAKPAQRYMPLQYKIWVLVMSKPFDTFILVLIALNTGVLMSQGEQFTDILMYLNIAFTILYIVEAGLKFIALRL-NYFRDYWNIFDFIVVLGGLLDVVVTVNTLGIGIDPSMFRLFRAARLIKLLRRGYTIRILLWTFLQSFKALPYVTLLIMLMFFMYAVIGMQLFGKIELDEGEINANNHFRNLLEALQVLFRSATGEDWHKIMRACYYGSTCGTVGAILYFCSFIFLCMFLMLNLFVAVIMDNFEYLTR

>Orbicella_faveolata_Cav2a_XP_020626975.1

SLFIFSKENFIRKICRTIVESKPFEYFILLTIFVNCILLAANTPLPNNDKSDLNQKLEDAEVYLLAIFCLEAVLKIIALGFVLHADSYLRNGWNVLDFVVVVTGLLSLPQLNVFSLKALRAARVLRPLKLVSGIPSLQVVMKSIMCAMVPLLQICLLVGFVIIIYAIIGLEFLVGRFHYVCNQTGRMEITDPDGPQICVAGTRCP-EGPNDGITSFDNIFAGMLTVFQVITNEGWTDIMYWTFDAYDYVFWIYYYSLVVIGSFFMLNLVLGVLSGEFARRTQKMERHLHGYIDWISKAEDLMRRRNLDRVEDGDVAMTSVVLTRWKIRVRQAVKHQAFYWTVLICVFLNTVITALQYYNQPAWLTQFQDVAEIVFISFFFCEMMLKLYGLGPQLYFKSQFNTFDCLVVSCGIIELIEGTSLGISVLRALRLLRLFKFTRYWSSLRNLVTSLLSSVRSILSLLFLLFLFIVIFALLGMQLFGAEFRGRNGNPRTNFDNFWNAALAVFQILTGEDWNAVMYEGVLSQGVW-CLYFVLLVVLGNYTLLNVFLAIAVDNLANAQQLSQDEEAEENAREERRREAGPTEDEFEEEADDGKPSFLSNLRNPMIDTWSLFLFPPGNPVRKACHWLVNLRHFDNFILVIILISSVLLALEDPV-DEESKRNEVLTYFDYVFTTVFAMEVLVKLIDYGAILHPYFRDAWNCIDALVVSCAIASLVMGSKKTVKVLRVLRVLRPLKAINKAKKLKAVFQCMVYSLKNVLNILIITILFLFIFSVIGVQLFQGKFFKCDDPSKMTKEECQGEGQEYNFDNVFYAMLSLFTSSTGEGWPALMQASIDTTAVDRGPIVDNKIEIALFYIFFVVVFSFFFINIFVALIILTFQEQGEKDQGDCELDRNQRDLHFAIVAKPSERFMPWQYRIWRIVDSSPFEFFIMILIALNTLILTMEPPLYREILDLFNTIFTFMFTGEAILKLFAFRT-NYFRDSWNVFDFIIVLGSLLDFALSRESDSMPFDPSLFRLFRAARLIKLLRQGYTIRILLWTFLQSFKALPYVGMLIGLLFFIYAVIGMQMFGQITIAEDQIASRNNFQSFPEAIQVLFRSATGENWQLIMLACTKEGKCGSAFAYLYFISFIFFCSFLLLNLFVAVIMDNFEYLTR

>Octopus_bimaculoides_Cav1_XP_014774811.1

SMFCLTLKNPLRKVCIRIVEWKAFEALILMTIFANCVALAIYTPYPESDTNEINMALENVEYVFLVIFTLECAMKIIAYGFVLHPGAYLRNGWNILDFIIVVIGVISTVLSFLNDVKALRAFRVLRPLRLVSRAPSLQVVLNSILRAMVPLLHIALLVIFVIIIYAIVGLELFSGKMHKTCFI-KNTEHLALEKPHPCGKGYSCH-IGPSDGITNFDNFGLAMLTVFQCITMEGWTGVLYNINDAMGSWPWIYFISLIIIGSFFVLNLVLGVLSGEFSREKQQIEEDLRGYLDWITQAEDIDHNKPSDTESSEKNEDLDS---RCRRLCRRLVKSQAFYWVVIVMVFLNTGVLTSEHYLQPSWLDQFQEVANLFFVVVFTCEMLLKMYSLGFESYFVSLFNRFDSFVVICSIVEVILIKPLGVSVLRCARLLRVFKATRYWTSLRNLVASLLNSMRSIASLLLLLFLFIVIFALLGMQLFGGKFNETQDKPRSNFDTFWQSLLTVFQILTGEDWNEVMYNGIKAYGVLVCLYFVILFICGNYILLNVFLAIAVDNLADAQSLTEIEQEKEEEKERSRSIEYRENTQIPSQEEEIPDEEHATEDEEIPKASALFVFSHDNRFRVFCHFVCNHNYFGNFVLACILISSAMLAAEDPL-DQSAERSKILNYFDYIFTSVFTIEIIIKLISYGLVLHKFCRSYFNLLDLLVVGVSLISINPSAISVVKILRVLRVLRPLRAINRAKGLKHVVQCVIVAVRTIGNIMLVTFLLQFMFAVIGVQLFKGTFHMCSDSSKRTMEECQGTNNDFNFDDVGKAMLTLFTVSTFEGWPNLLYISIDSHQEGMGPVYDNRPVVAIFYFVYIIVIAFFMVNIFVGFVIVTFQNEGEQEYKNCELDKNQRKIEFALKVKPARRYIPWQYKVWWFVTSQPFEYAVFALIMINTITLAMENKGYSDVLDYLNMIFTGLFTVEFMLKLAAFRFKNYFGDAWNVFDFIIVLGSFIDIIYTEAPGSPIISINFFRLFRVMRLVKLLSRGEGIRTLLWTFIKSFQVLIYFFFLFFFFFFAFSISIKXTFGRIALNETEIHRNNNFQTFPQAVLVLFRSATGEAWQEVMLSCVTNSTCGTNFAYPYFISFYILCSFLIINLFVAVIMDNFDYLTR

>Nemat_caeele_Cav3_CCD68017.1

ALRCFYQARPPRKWALQMVMSPWFDRITMAVIMINCVTLGMYRPCEDGPDTYRCQILDIIDNCIFVYFAFEMVIKIMALGF-YGPAAYMSDTWNRLDFFIVMAGIAEFVLHEYLNLTAIRTVRVLRPLRAVNRIPSMRILVNLLLDTLPMLGNVLLLCFFVFFIFGIVGVQLWAGLLRNRCVILTRFYIPEDTSLYICSQGVKCNQRNPFQGSVSFDNIGFAWVAIFLVISLEGWTDIMYYVQDAHSFWNWIYFVLLIVIGAFFMINLCLVVIATQFAKEEGDTYAAFVRFIGHTFRRTKRASRIEEKAEDEEDETTITRENGWFREKIQKFVICDHFTRGILVAILVNTLSMGVEYHQQPEILTVILEYSNLFFTALFALEMLLKIIASGLFGYLADGFNLFDGGIVALSVLELFQEGKGGLSVLRTFRLLRILKLVRFMPALRYQLVVMLRTMDNVTVFFGLLVLFIFIFSILGMNLFGCKFCLAKKCERKNFDTLLWALITVFQILTQEDWNMVLFNGMAQTNPWAALYFVALMTFGNYVLFNLLVAILVEGFQEEEQLEEDARAVEEEDERKRELVPYRRQRVHSWSGLCHHFNPNCPVHGNRTEFSLFLMGPKNPLRIKCLQTTQKKWFDYTVLFFIGINCITLAMERPSIPPDSFERQFLHISGYIFTVIFTGEMMMKVIANGCFIGQYFKDGWNILDGILVVISLINIAFEIFGVIRVLRLLRALRPLRVINRAPGVKLVVMTLISSLKPIGNIVLICCTFFIIFGILGVQLFKGMMYHCIEVGVTTKADCIEVNHRYNFDNLGQALMSLFVLSSKDGWVSIMYQGIDAVGVDVQPIENYNEWRMIYFISFLLLVGFFVLNMFVGVVVENFHKCKEKEMREKEKEKRLKRQKFEESMAPYYHYGHTRLFLHGIVTSKYFDLAIAAVIGINVISMAMMPMGLKYVLKALNYFFTAVFTLEAAMKLIALGFKRFFIEKWNRLDMFIVILSIAGIIFEEELPINPTIIRVMRVLRIARVLKLLKMAKGIRSLLDTVGEALPQVGNLGSLFFLLFFIFAALGVELFGKLECSEDGLGEHAHFKNFGMAFLTLFRIATGDNWNGIMKDALETNCCVPILAPCFFVIFVLISQFVLVNVVVAVLMKHLEESNK

>Nemat_caeele_Cav2_NP_001123176.1

SLFIFAEDNIIRRNAKAIIEWGPFEYFILLTIIGNCVVLSMEQHLPKNDKKALSEWLERTEPYFMGIFCLECVLKVIAFGFALHKGSYLRSGWNIMDFIVVVSGVVTMLPFSPADLRTLRAVRVLRPLKLVSGIPSLQVVLKSILCAMAPLLQIGLLVLFAIIIFAIIGLEFYSGAFHSACYNERGEIENVSERPMPCTNVYNCD-IGPNYGITSFDNIGFAMITVFQCITMEGWTTVMYYTNDSLGTYNWAYFIPLIVLGSFFMLNLVLGVLSGEFARRQQQIERELNGYLEWILTAEEVIKQQSTETEEDFEEDEDEMEEEQLRIQIRIMVKTQIFYWSVITLVFLNTCCVASEHYGQPQWFTDFLKYAEFVFLGIFVVEMLLKLFAMGSRTYFASKFNRFDCVVIVGSAAEVIYGGSFGISVMRALRLLRIFKLTSYWVSLRNLVRSLMNSMRSIISLLFLLFLFILIFALLGMQLFGGRFNFPTMHPYTHFDTFPVALITVFQILTGEDWNEVMYLAIESQGGWYSIYFIVLVLFGNYTLLNVFLAIAVDNLANAQELTAAEEADEKANEIEEE-GDHCTIDMEGKTAGDMCAVARAMDDLMVPYSSMFFLSPTNPFRVLIHSIVCTKYFEMMVMTVICLSSVSLAAEDPV-DEENPRNKVLQYMDYCFTGVFACEMLLKLIDQGILLHPYCRDFWNILDGIVVTCALFAFGFANLNTIKSLRVLRVLRPLKTIKRIPKLKAVFDCVVNSLKNVFNILIVYFLFQFIFAVIAVQLFNGKFFFCTDKNRKFANTCHGRLRPFNYDNTINAMLTLFVVTTGEGWPGIRQNSMDTTFEDQGPSPFFRVEVALFYVMFFIVFPFFFVNIFVALIIITFQEQGEAELSEGDLDKNQKQIDFALNARP-RSFMPTKYRIWRLVTSPPFEYFIMTMICCNTLILMMNPLFYEEILRLFNTALTAVFTVESILKILAFGVRNYFRDGWNRFDFVTVVGSITDALVTE--GGHFVSLGFLRLFRAARLIRLLQQGYTIRILLWTFVQSFKALPYVCLLIGMLFFIYAIVGMQVFGNIWLNATEINRHNNFQSFFNAVILLFRCATGEGWQDIMMAAVKGQTCGSNVSYAYFTSFVFLSSFLMLNLFVAVIMDNFDYLTR

>Nemat_caeele_Cav1_NP_001023079.1

SLLCLSLNNPIRKLCISIVEWKPFEFLILFMICANCIALAIYQPYPAQDSDYKNTLLETIEYVFIVVFTIECVLKIVAMGFMFHPSAYLRNAWNILDFIIVVIGLVSTILSKMSDVKALRAFRVLRPLRLVSGVPSLQVVLNAILRAMIPLLHIALLVLFVILIYAIIGLELFCGKLHSTCID-PATGQLAQKDPTPCGTAFKCQ-PGPNNGITNFDNFGLAMLTVFQCVSLEGWTDVMYWVNDAVGEWPWIYFVTLVILGSFFVLNLVLGVLSGEFSREKQQLEEDLKGYLDWITQAEDIETVVGEEADEEGEERVEDVRP-RCRRACRRLVKSQTFYWLVILLVLLNTLVLTSEHYGQSEWLDHFQTMANLFFVILFSMEMLLKMYSLGFTTYTTSQFNRFDCFVVISSILEFVLMKPLGVSVLRSARLLRIFKVTKYWTSLRNLVSSLLNSLRSIISLLLLLFLFIVIFALLGMQVFGGKFNPQQPKPRANFDTFVQALLTVFQILTGEDWNTVMYHGIESFGVIVCIYYIVLFICGNYILLNVFLAIAVDNLADADSLTNAEKEEEQQ-----------------EIEGEDEEFEEGEDEGIPKASSLFILSHTNSFRVFCNMVVNHSYFTNAVLFCILVSSAMLAAEDPL-QANSTRNMILNYFDYFFTSVFTVEITLKVIVFGLVFHKFCRNAFNLLDILVVAVSLTSFVLRAMSVVKILRVLRVLRPLRAINRAKGLKHVVQCVIVAVKTIGNIMLVTFMLQFMFAIIGVQLFKGTFFLCNDLSKMTEAECRGSNNDFNFDNVGDAMISLFVVSTFEGWPQLLYVAIDSNEEDKGPIHNSRQAVALFFIAFIIVIAFFMMNIFVGFVIVTFQNEGEREYENCELDKNQRKIEFALKAKPHRRYIPLQYRVWWFVTSRAFEYVIFLIIVMNTVSLACSSRGFEDFLDVFNLIFTGVFAFEAVLKIVALNPKNYISDRWNVFDLLVVVGSFIDITYGKPGGTNLISINFFRLFRVMRLVKLLSRGEGIRTLLWTFMKSFQALPYVALLIVLLFFIYAVIGMQFFGKVALDDTSIHRNNNFHSFPAAILVLFRSATGEAWQDIMLSCSNESRCGNNFAYPYFISFFMLCSFLVINLFVAVIMDNFDYLTR

>Mollu_lymsta_Cav3_AAO83843.2

TFYIFTQRNYIRFWCLRCITWPWFERISMFVIILNCVTLGMYQPCNDKEVTLRCRILEGFDHFIFAFFAVEMIIKMIAMGV-IGKETYLADSWNRLDCFIVVAGLAEYIVNKTISLSAIRTIRVLRPLRAINRIPSMRILVMLLLDTLPMLGNVLLLCFFVFFIFGIIGVQLWSGVLRHRCYMVSEFYKQKNKQEYICSPGILCNAPNPFEGAVSFDNIGLAWVAIFQVISLESWVIIMYHVQDAHSFWDWIYFVALIVIGSFFMINLCLVVIATQFSSEPGGCYSELLKLVAQVYRRVKRKASNMLLNVDYEPSKTQSLADKCLQKKLKTFVESNFFQRSILIAILLNTLSMGVEFHNQPDLLTTILEYSNVVFCVLFGTEMAFKISAYGLFGYISNGFNVFDGFIVILSIVELAQGGASGLSVLRTFRLLRILKLVRFMPALRRQLVVMLRTMDNVATFFALLVLFMFIFSILGMSLFGGTFCSLCSCDRANFDNLLWSLVTVFQVLTQEDWNTVLYNGMAKTSNWASLYFVALMTFGNYVLFNLLVAILVEGFSTKEELEDVDKEDEEEEEEKEKQSRQNSFTSHRTLNSLGSGNSKDDNKSERHEYAFYLLSNENRIRKFALHLISRKWFDNAVLFFIALNCITLAMERPDIPPDSIERYFLTYTNYIFTFVFALEMMIKVIGKGFFVGKYLKSGWNVMDGFLVIISLIDILISIFGILRVFRLLRTLRPLRVISRAPGLKLVVQTLLSSLRPIGNIVLICCTFFIIFGILGVQLFKGTFYHCKNILITNRSQCEAINQKYNFDNLGQALMALFVLASKDGWVQIMYTGLDAVGIDQQPIENYNEWRLIYFISFLLLVAFFVLNMFVGVVVENFHKCRESQEIEERAKRAAKREKLDKKRKPYWAYSHSRLLIHTVINSKYFDLAIAAVIGLNVITMAMMPEELEFALKIFNFFFTSVFILEAVMKIIALGFYRYIRDRWNQLDIMIVILSIVGIVLEEVIPINPTIIRVMRVLRIARVLKLLKMAKGIRALLDTVIQALPQVGNLGLLFFLLFFIFAALGVELFGRLDC--EGLGVHAHFRNFGMAFLTLFRVATGDNWNGIMKDTLLKNCCVPLIAPVYFVVFVLMAQFVLVNVVVAVLMKHLEYKYK

>Mollu_lymsta_Cav2_AAO83841.1

SLFIFSEENFIRKYAKIIIEWGPFEYMVLLTIIANCIVLALEEHLPSQDKTPLALQLDDTEVYFLGIFCVEAFLKIVALGFCLHKRSYLRNIWNIMDFIVVVTGFITLFAQGSSDLRTLRAVRVLRPLKLVSGIPSLQVVLKSIIRAMAPLLQVCLLVLFAIVIFAIIGLEFYVGVFHNACYKSEDDIDTGDEDDRPCLPAFQCQ-RGPNAGITSFDNIGYAMLTVFQCITMEGWTNVLYYTNDALGQFNFLYFIPLIILGSFFMLNLVLGVLSGEFARRQQQIERELNGYLEWICKAEEVIKMKQLKAEDNENDSEQNDNDLRFRYSIRRLVKSQLFYWIVIVLVFLNTASVASEHYNQPEWHVQFLYITEYAFLGLFIFEMSIKMYALGVRMYFQSSFNIFDCVVIVGSIVEVIRGSSFGISVLRALRLLRIFKVTRYWSSLRNLVISLLSSMRSILSLLFLLFLFIIVFALLGMQLFGGEMNFEEGRPSAHFDTFPIALLTVFQILTGEDWNEVMYNGIKSHGMFYSSYFIVLVLFGNYTLLNVFLAIAVDNLANAQELTAAEEEQEEEEAVRREESVDNVLETAAKTSSVTMPLANNTEEDMLPYSSMFIFGPTNPIRRFCHFVVNLRYFDLFIMIVICASSVALAAEDPV-IENSRRNEILNYFDFVFTGVFTIELVLKVIDLGVLLHPYIRDLWNILDATVVICALVAFVFKNLNTIKSLRVLRVLRPLKTINRVPKLKAVFDCVVNSLKNVSNILIVYILFQFIFAVIAVQLFKGRFFYCTDESKSTRDECQGLRQDFHYDNIMMAMLTLFTVTTGEGWPSVLKHSMDSTYEDRGPKPVYRMEMSLFYVVFFIVFPFFFVNIFVALIIITFQEQGENELMDQEMDKNQKQIDFAINAKPHCRFIPIKYKIWRLVQSSKFEYFVMTLITLNTIVLMMMSDNYKDVLAKLNEGFTVLFTLECLLKIIGLGPRNYFHDPWNVFDFTTVVGSIIDVLITE--SKRQVSFGFFRLFRAARLVKLLRQGYTIRLLLWTFFQSFKALPYVCLLILMLFFIYAIIGMQVFGSIKLDSTSINRHNNFRTFFSALTLLFRCATGEAWQQIMQSCLAESGCGTNIAYMYFVSFIFLCSFLMLNLFVAVIMDNFDYLTR

>Mollu_lymsta_Cav1_AAO83839.1

ALFCLTLKNPIRKFCIQVAEYKAFEFLVLITIFANCVALAIYTPYPMSDSNEVNSALDRIEYVFLVIFLLEGILKIIAYGFVMHQGAYLRNGWNALDFTIVVIGIISSILSFVQDVKALRAFRVLRPLRLVSRAPSLQVVLNSIVRAMVPLLHIALLVIFVIIIYAIIGLELFYGKLHNACYK--INSTEFSGDPRICGQGYSCD-EGPNYGITNFDNFGLAMLTVFQCITMEGWTTVLYDVNNALGEWPWIYFISLIIIGSFFVLNLVLGVLSGEFSREKQQLEEDLRGYLDWITQAEDIDITSDIDSEDKVEEGEN-----RYRRFCRRIVKSQAFYWGVIVLVFLNTVVLTSEHYKQPVWLDDFQAIANLFFVVLFTMEMLVKMYSLGFQGYFVSLFNRFDSFVVVCSILEVIVFPPLGISVLRCARLLRVFKATRYWSSLRNLVASLLNSMRSIASLLLLLFLFIVIFALLGMQLFGGKFNPQGEKPRSNFDTFWPSLLTVFQILTGEDWNAVMYDGIRAYDILVCLYFVVLFIVGNYILLNVFLAIAVDNLADAQSLTEIEEEKEEEKERTRSLDEDENNEESDEEEGTEGTETEEDEEGIPPFSSLFIFSATNKFRIICHKICNHSYFGNVVLACILISSAMLAAEDPL-RNESPRNQILNKFDYFFTSVFTIEIIIKIITYGLMLHKFCRSLFSILDLVVVAVSLISFPLDAISVVKILRVLRVLRPLRAINRAKGLKHVVQCVIVALRTIYNIMLVTFLLNFMFSVMGVQLFKGKFSMCTDESKLTEDECQGKKNDFNFDDVSNGMLTLFTVSTFEGWPGLLYKSIDSHAEGKGPIQNSKPAVAVFYFIFIIVIAFFMMNIFVGFVIVTFQNEGEQEYKNCELDKNQRKIEFALKVKPIRRYIPWQYKIWWFVTSQAFEYGIFVLIMINTVALAMQSASYSDALDYLNIIFTGVFTVEFVLKLAAFRFKNYFGDAWNVFDFIIVLGSFIDIIYAENPGKAFISINFFRLFRVMRLIKLLSRGEGIRTLLWTFIKSFQALPYVALLIVMLFFIYAVIGMQMFGRIKLDYTQIHPNNNFQTFPHAVLVLFRSATGESWQEIMLACSEANRCGNDFAYVYFISFYILCSFLIINLFVAVIMDNFDYLTR

>Lingula_anatina_Cav1_XP_023932094.1

ALFCLTLKNPIRRTCISIIEWKPFEALILLTIFANCVALAIYLPYPKGDSNEVNDALDNVEIVFLIIFAVEAILKIIAYGFLFHQGAYLRNGWNILDFTIVVIGTLSTVLSYMRDVKALRAFRVLRPLRLVSRAPSLQVVLNSIIMAMVPLLHIAMLVCFVIIIYAIVGLELFSGKMHKTCYH--SITDEIMDDPTPCGSSYMCD-EGPNSGITNFDNIGLAVLTVFQCVTMEGWTTVLYWINDAVGEWPWMYFISLIILGSFFVLNLVLGVLSGEFSREKQQLEEDLKGYLDWITQAEDIEPTHKANESQSEKTEELGSGEMRCRRTCRKIVKSQGFYWIVIVMVFFNTCVLTTEHYKQPEWLDEFQFIGNLFFVILFALEMLLKMYSLGFQGYFVSLFNRFDCFVVICSIVEVVVMPPLGVSVLRCARLLRVFKVTRYWSSLSNLVASLLNSMRSIASLLLLLFLFIVIFALLGMQLFGGKFNTDKSKPRSNFDDFFQSLLTVFQILTGEDWNEVMYDGINSYKVLVCLYFVILFIVGNYILLNVFLAIAVDNLADAEQLTEMEKEKEEEKERQRSIQILTGEDWNEVMYDGINSYKGVASPGIPNASSLFIFSPTNKIRIFCHQVCNHSYFTNIVLACILISSAMLAAEDPL-NADSERNQILNYFDYFFTSVFTVEIIIKVIAYGFFVHKYCRSIFNLLDLLVVSVSLISIFLEAFSVVKILRVLRVLRPLRAINRAKGLKHVVQCVIVAIKTIGNIMLVTFLLNFMFAVIGVQLFKGRFYHCTDESKMTEEECQGVNNPLNYDDVSQGLLTLFTVATFEGWPGLLYTSIDSNEESQGPIYNYRQAVAVFYIIFIIIIAFFMVNIFVGFVIVTFQNEGEQEYKGCELNKNQRKIEFALKAKPTRRYIPIQYKIWWFVTSRAFEYSIFGLILFNVMALAMMSDGYSQFLSVMNIIFTAAFTAECVLKLIAFKFKNYFGDAWNVFDFIIVLGSFIDIIYTKNEGEKMISVNFFRLFRVMRLVKLLSRGEGIRTLLWTFIKSFQALPYVALLIVMLFFIYAVIGMQVFGKIALDDTPMTRNNNFQTFPQAVLVLFRSATGEAWQDVMLGCIDKESCGSDFAYVYFISFYILCSFLIINLFVAVIMDNFDYLTR

>Hyalella_azteca_Cav3_XP_018025902.1

-----------------MITW--FERVSMMAILLNCVTLGMYQPCHDEEVTTRCKTLQIFDDLIFAFFAVEMLIKVMAMGF-HGKGTYLAESWNRLDFFIVVAGAVEYCLSVENNLSAIRTIRVLRPLRAINRIPSF-----------------------------------------------------NLQEIYPPT-------------------------SILSVISLEGWVDIMYYVQDGHSFWDWIYFVLLIVVS----MDVLLIVVSMDVLSDQPGCYAEILKYIAHLARKFKRKVCLSEKTTQTDPEPPEPLVSRAGRKKVQQLVNSNYFQRWILCAILVNTLSMGVEYHNQPEELTQIVETSNMIFSGVFAFEMFLKIISEGPFGYISNGFNLFDGVIVVLGIVEMCLADAAGLSVLRTFRLLRILKLVRFMPQLRRQLFVMLRTMDNVAIFFSLLVLFIFIFSVLGMYLFGGKFCSYCRCHRMHFNSLLWATVTVFQILTQEDWNVVLFNGMEKTSHWASLYFVALMTFGNYVLFNLLVAILVEGFSALQEQTEVQAAPENDEQEEKKLERTAVGNGERDAEDGGRHVEEETLNNERVDYSLYIFSETNWIRRVCKYLVSKKYFDSTVLFFIGLNCITLAMERPNIPPDSTERAFLSTCNDVFTVVFGLEMLIKVIAQGLLYGKYFSSGWNIMDGILVIVSVIDVLMSIFGILRVFRLLRALRPLRVINRAPGLKLVVQTLLSSLQPIGNIVLICCTFFIIFGILGVQLFKGAFFYCDGLDVETKADCLKVNRKYNFDNLGHALMSLFVLSSKDGWVNIMYTGLDAVGVDRQPKENYSQWRLLYFISFLLLVGFFVLNMFVGVVVENFHRCREEQEKEEKARRAAKRKKLEKKRKPYYAYSKGRLFVHNLVTSKYFDLAIAAVIGLNVVTMAMMP------------------------KIVARAIK---CPRWNQLDVVIVLLSVAGIVLEEIIPINPTIIRVMRVLRIARVLKLLKMAKGIRALLDTVMQALPQVGNLGLLFFLLFFIFAALGVELFGRLECDDQGLGEHAHFKNFGIAFLTLFRVATGDNWNGIMKDTILKNCCLPFVAPIFFVIFVLMAQFVLVNVVVAVLMKHLEESHK

>Hyalella_azteca_Cav2_XP_018019172.1

SLFILSDTNIFRKYTKFIIEWPPFEYAVLLTIIANCIVLALEEHLPLSDKTVLAQELEKTEPYFLFIFCIESSLKILALGFVLHPNSYLRNIWNIMDFVVVVTGFVTLSMQEDLDLRTLRAIRVLRPLKLVSGIPSLQVVLKSIIKAMAPLLQIGLLVLFAIVIFAIIGLDFYCGALHKTCYSDLDVIITEGEAATPCYAAFLCD-DGPNFGITSFDNIGFAMLTVFQCITQEGWTSILYWTNDSVGTFNIFYFVPLIVIGSFFMLNLVLGVLSGEFSRMRKLFTEDFLNYFEWIGRAEEVIGKSKSTDTDEEDNDIEEEDGQRFRFRIRRTVKQQWFYWFVIVLVFLNTACVASEHYAQPLWLADFLYYAEYAFLGLFLTEMLVKIYALGPRIYFESSFNRFDCVVISGSIFEVIYDESFGFSVLRALRLLRIFKVTNS------------------------------TYGLPGLFL--------------NFNRFAIALLTVFQILTGEDWNEVMYQGIKSQGMIYSMYFIILVVFGNYTLLNVFLAIAVDNLANAQELTAAEEEKEEEDKEKQQAQQSAAGDGPPTIDASKDADPDIDFADMLPYSSMFILSPTNPIRRAAHWVVNLRYFDFFIMVVISLSSMALASEDPV-EEGSKWNTYLTYFDYAFTGVFAVEMILKVVDLGVIFHPYLRDLWNIMDSVVVICAAVSFCFDNLSTIKSLRVLRVLRPLKTIKRVPKLKAVFDCVVNSLKNVFNILIVYILFQFIFAVIAVQLFNGKFFYCTDLSIMTKAECQGKQQSFHYDNVMFAMLTLFAVQTGEGWPQVLQNSMAATYENHGPQPHFAIEMSIFYIVYFIVFPFFFVNIFVALIIITFQEQGEAELQDGELDKNQKSIDFAIQAKPLERYMPLKYKTWKVVVSTPFEYLIMTLIVLNTILLMMQSKIYQRSLHYLNSFFTALFTIECMLKISAFGIRNYFKDNWNTFDFICVVGSIIDALVVEDSSNSYINLRFLRLFRAARLIKLLRQGDTIRILLWTFIQSFKALPYVCLLIVILFFIYAIIGMQVFGAILLDPTSYTRHNNFRNFWQGLMLLFRCATGESWQSIMLSCIKEETCGSNIAYAYFVSFIFFCSFLMLNLFVAVIMDNFDYLTR

>Heterololigo_bleekeri_Cav2_BAA13136.2

SLFIFSEENFIRKYAKIIIEWGPFEYMVLLTIIANCIVLALEEHLPNEDKTPLAVQLEATEFYFLGIFCVEALLKIVALGFALHKGSYLRNVWNIMDFVVVVTGFISIFP----DLRTLRAVRVLRPLKLVSGIPSLQVVLKSIIRAMAPLLQVCLLVLFAIVIFAIIGLEFYTGAFHKACFISEDNIEYGDEDTRPCANGYHCR-AGPNYGITSFDNMGFAMLTVFQCVTMEGWTQVLYYTDDAVGAYNWIYFVPLIVLGSFFMLNLVLGVLSGEFARRQQQIERELNGYLEWICKAEEVINKELTIGDEDNDGDLLSGINVHLRFTIRKCVKTQGFYWFVIILVFLNTLCVASEHYGQAEWHTEFLYVMEFAFLALFMSEMLIKMYGLGVRLYFQSSFNIFDCVVILVSIIEVIDGSSFGISTLRALRLLRMFKVTRYWSSLRNLVVSLLSSMRSIVSLLFLLFLFILIFALLGMQLFGGMMNFEEGRPPGHFDTFPIALLTVFQILTGEDWNEVMYSGIRARGMLYCSYFIILVLFGNYTLLNVFLAIAVDNLANAQELTAAEEIQEGVRQEQEAAEERNKIGADKDLKKDRDPLDNVSLHEMLPYSSMFIFGPTNPVRRFCHFVVNLRYFDLFIMIVICASSIALAAEDPV-NDESVNNQILNYFDYVFTGVFTIEMLLKIVDLGIILHPYCRDAWNILDATVVICALVAFAFGNLNTIKSLRVLRVLRPLKTINRIPKLKAVFDCVVNSLKNVSNILIVYLLFQFIFAVIAVQLFKGKFFYCTDMSKSNREECQGLRQDFHYDNVMFAMLTLFTVTTGEGWPMVLKNSMDSTSDDMGPKPGYRMEMAIYYVVFFIVFPFFFVNIFVALIIITFQEQGENELVDQDLDKNQKQIEFSIEAKPSCRYVPIKYKIWQVVVSPKFECVVMVLIALNTLVLMMSPTEYKLLLQNLNLAFSVLFTIECILKLMGFGIGNYFRDRWNMFDFIIVIGSIIDVVTTNLPSASSFRTGSFRLFRAARLVKLLRQGYTIRLLLWTFLQSFKALPYVCLLIAMLFFIYAIIGMQVFGNIRLDSTSINRHNNFRSFFYAVLLLFRCATGESWQQIMLSCLLDNSCGLDIAYIYFVTFIFLCSFLMLNLFVAVIMDNFDYLTR

>Halocynthia_roretzi_Cav1_BAA34927.2

ALLCLSLKNPIRKACMKIVDWRPFDVLILLTILANCVALAVYVPFPGDDSNRTNEILEKVEYIFLGIFTIEAILKIIAYGLFFHPNAYLRNGWNVIDFVIVVIGLVSIVLETANDKRSLRAFRVLRPLRLVSGVPSLEVVLNAIIRAMVPLLHIALLVIFVIIIYAVVGLELFKGKLHKTCYH-NEVAVLIMEDEKPCADGRHCS-AGPSKGIINFDTFYFAVITVFQCITMEGWTDVLYYMNDAVGLWPWIYFVSLIIIGSFFVMNLILGVLSGEFSREKQQTDEDMKGYMDWITQAEDLDEDRRSASNEQLNDADSEVSGLKTRRRCRTMVKSKSFYWLVIVLVFCNTLSLATEHYRQPPWLTLAQDLANKILLTLFTIEMLVKMYSLGMQQYFVSLFNRFDCFVVCGGIVELVIMEPLGISVLRCVRLLRIFKMTSSWNSLSNLVASLLNSIRSIASLLVLLFLFIIIFALLGMQMFGGRFSEQEDKIRSNFDTFLQALLTVFQILTGEDWNVVMYNGIEAYGLLTSVYFIVLFIGGNYILLNVFLAIAVDNLADAESLGAAQKEKEEEKKMKKTLYYTPDHDIRIEVTEASDTNSDKHLPELPEGSSFFILSNTNRLRVFCYDIVNYNWFNNAILACIILSSIALACEDPV-SAHSARNKVLEYFDYVFTGVFAVEIVLKMTAFGVFLHKFCRSYFNLLDLLVVAVSLVSMLSNKFSVVKILRVLSVLRPLRAINRAKGLKHVVQCVFVAISTIGNIMVITGLLQFMFACIGVQLFKGRLYYCTDQSKETKEECHGENSEFNYDNVMNAMLTLFVVATFEGWPGLLYKSIDSWSENHGPRYDARQAVALFYFVFIIVIAFFMMNIFVGFVIVTFQEQGEQEYKNCELDKNQRQLEYALKAKPVKRYIPWQYKVWFIVNSTYFEYFMLVLILLNTVCLAVQSKELTVILNHMNYVFTALFALEMIVKLVAYKPRGYLSDPWNVFDSLIVIGSIVDIVFSEHGNEKSFSINFFRLFRVLRLVKLLSRGEGIRTLLWTFIKSFPALPYVALLIIMLFFIYAVIGMQIFGKIKPNDSQINRNNNFQTFLQAVLLLFRCATGESWQEVMLACAAKFTCGNDFAYTYFLTFYMLCAFLIINLFVAVIMDNFDYLTR

>Haemonchus_contortus_Cav2_CDJ96819.1

SLFIFSEDNFIRRNAKAIIEWGPFEYFILLTIIGNCVVLAMEQHLPKNDKKPLSELLERTEPYFMGIFCLECVLKIVAFGFIAHKGSYLRSGWNIMDFIVVVSGVVTMLPVSPADLRTLRAVRVLRPLKLVSGIPSLQVVLKSILCAMAPLLQIGLLVLFAIVIFAIIGLEFYSGAFHSACYNDRGEIENVSEKPSPCTNVYNCD-IGPNYGITSFDNIAFAMITVFQCITMEGWTTVMYYTNDSLGTYNWAYFIPLIVLGSFFMLNLVLGVLSGEFARRQQQIERELNGYLEWIMAAEEVIKQQSTETEEDMEEEEEELDEEKARIQMRVIVKTQIFYWSVITLVFLNTACVASEHYGQPPWLTKFLQYAEYVFLGIFIMEVLLKLFAMGSRTYFASKFNRFDCIVIVGSAFEVIKGGSFGISVLRALRLLRIFKLTSYWVSLRNLVRSLMNSMRSIISLLFLLFLFILIFALLGMQLFGGKFNFPTMHPYTHFDTFPVALITVFQILTGEDWNEVMYLAIEAQGMVYCIYFIVLVLFGNYTLLNVFLAIAVDNLANAQELTAAEEADEKANEMDDSE-EEEPDGDHCAIDMDGNDQDDDEECEMVPYTSMFFLSPSNPLRVLVHSIVCTKYFEMMVMGVICLSSISLAAEDPV-DEENPRNKVLQYMDYCFTGVFACEMLLKLIDQGIILHPYCRDFWNILDGVVVTCALVAFGFANLNTIKSLRVLRVLRPLKTIKRIPKLKAVFDCVVNSLKNVFNILIVYFLFQFIFGVIAVQLFNGKFFYCTDKTKRFAYQCHGRLRPFNYDNTINAMLTLFVVTTGEGWPGIRQNSMDTTFEDQGPSPFYRVEVALFYVMFFIVFPFFFVNIFVALIIITFQEQGEAELSEGDLDKNQKQIDFALNARP-RSFMPIKYRIWRLVTSAPFEYFIMAMICCNTIILMMNSDFYEKVLRLFNTALTAVFTVESILKILAFGVRNYFRDGWNRFDFVTVVGSITDALVTE--GGHFVSLGFLRLFRAARLIRLLQQGYTIRILLWTFVQSFKALPYVCLLIGMLFFIYAIVGMQVFGNIWLNATEINRHNNFQSFFNSVILLFRCATGEGWQDIMMACGKGQTCGSNVSYAYFTSFVFLSSFLMLNLFVAVIMDNFDYLTR

>Gonium_pectorale_Cav_KXZ52368.1

--------------------M--------------------SSNRQDFEDTTLGSTLNKFEFLWVAIFTVEALLKIVSMGFALAPGTYLRDGWNVVDFMVVALGYIDIFTAG--NLTALRTVRVLRPLRAITRVRGMRVLVTTLLGSMPMLLDVFLLCAFTFFIYGLIAVQVFAGALRYRCGTHNVSYVVSDDDAELCGGGHVCP-GNPNYGMTSFDHILWAWLTIFQCITQEAWTDVMYFTSDSLSWWVWPFFVVLVVFGSLYIINLALAVIYAQFMAEVALKAEAAEG----VQREAERAGKAALAAAAGKPGKEGDRHSSPLRRAACRVAVGKRLEYTTMVVIILNTALMCINWFRMPASVENSANIVNYVFTMYFFLELLLKLFAFGVVRYFRDGMNIFDFIVVLISMVEFISVSGVGLSVLRAFRLLRIFRLARSWKELNFIIRALFRSVTSTTYLLLLTLLFLFISALMGMQLFGYKFCGKSDVPRARFDTVFWGIYTVFQLLTTENWNNIMYDSMRSTTPWGAVYYVAVLVIGTYLVFNLFVAILLDNFS-GSSLDTSARSDPALKQQEDQLFLRVSAATKASADGGRRELPQRDPSGFIRGRSLLLLDSNHWLRWRAARLVHDTRFETVVLVLIVASSVTLALDTPSLQRGSRLELAIRYCDYVFVGAFTLEALLKIITFGFFTGKYVRNGWNVLDFLVVLIGLTMIALENLQMLRVLRTLRALRPLRTASRYEELKVVVNALFAVIPAMGNVALVSLLFYLIFAILAVNLFKGQLYNCVDADTMTRQWCEAVNLRSNFDNVGSAVLTLFQLSTLELWVDIAFSAVDATGVDQQPLWNHQPQMLLFFGLFIVVCAFFILNLFVGVTLDKFMEMHEAQTARSVLITPQQTLRTSMDQRPAQPGPPWREGLYALATSNAFNNFIMGTITVNVLFMAMMSNSWQAVMSISNVIFTAIFAVEAALKLAAFGPTDYFRDKWNCFDFSVVLISMASIALDFSDTQNLSFMPVLRVLRVVRVIRIIRRAKGLQRLLVTLLYSLPALGNVGGVMLLFFFCFSVIGMNLFGGIKF--DFLNRHANFNNFPNAMLLLFRMITGESWNGIMHDCMLNDRCAPAAAVIYFPVFIILCAFIMLNLIVAVIVENMIMAGA

>Gekko_japonicus_Cav3.3_XP_015274652.1

VFFCLKQTTSPRNWCIKMVCNPWFECVSMMVILLNCVTLGMYQPCDDMDLSNRCKILQVFDDFIFIFFAMEMVLKMVALGI-FGKKCYLGDTWNRLDFFIVMAGMVEYSLDLQNNLSAIRTVRVLRPLKAINRVPSMRILVNLLLDTLPMLGNVLLLCFFVFFIFGIIGVQLWAGLLRNRCFMPPYYQPEEDDEMFICSLGHECCNSNPHKGAINFDNIGYAWIVIFQVITLEGWVEIMYYVMDAHSFYNFIYFILLIIVGSFFMINLCLVVIATQFSMEPGDCYEEIFQYVCHIVRKAKRRRCRKHNSVDYMPQNLGQPIAMEVRVKLRGIVESKYFNRGIMIAILVNTISMGIEHHEQPEELTNILEICNVVFTSMFALEMILKLSAFGLFDYLRNPYNIFDSIIVIISIWEIIGQADGGLSVLRTFRLLRVLKLVRFMPALRRQLVVLMKTMDNVATFCMLLMLFIFIFSILGMHIFGCKFSGDTVPDRKNFDSLLWAIVTVFQILTQEDWNVVLYNGMASTSPWASLYFVALMTFGNYVLFNLLVAILVEGFQAGDSYSDEDQSSSNIEEFDKCQPAVPSGHRPTRRIGVTTGPIEHQDCNLREDWSIYLFSPQNRFRILCQTIIAHKLFDYIVLAFIFLNCITIALERPQIEQRSTERIFLTVSNYIFTAIFVAEMTLKVVSLGLYFGDYLRSSWNILDGFLVLVSIIDIVVSILGVLRVLRLLRTLRPLRVISRAPGLKLVVETLISSLKPIGNIVLICCAFFIIFGILGVQLFKGKFYHCLDIRITNRSDCMAVHHKYNFDNLGQALMSLFVLASKDGWVNIMYNGLDAVAVDQQPVTNNNPWMLLYFISFLLIVSFFVLNMFVGVVVENFHKCRQHQEAEEARRREEKRRRLEKKRRPYYAYCSVRLLIHSVCTSHYLDIFITFIICLNVVTMSLQPMSLETALKYCNYMFTAVFVLEAVLKLVAFGLRRFFKDRWNQLDLAIVLLSVMGITLEEALPINPTIIRIMRVLRIARVLKLLKMATGMRALLDTVVQALPQVGNLGLLFMLLFFIYAALGVELFGKLVCNDEGMSRHATFENFGMAFLTLFQVSTGDNWNGIMKDTLDDRSCLQFISPLYFVSFVLTAQFVLINVVVAVLMKHLDDSNK

>Gekko_japonicus_Cav3.1_XP_015266211.1

VFFYLSQHSRPRSWCLRMVCNPWFERASMLVILLNCVTLGMFHPCEDTDDSPRCKILQSFDDFIFAFFAVEMIIKMIALGI-FGKKCYLGDTWNRLDFFIVIAGMLEYSLDLQN--SAVRTVRVLRPLRAINRVPSMRILVTLLLDTLPMLGNVLLLCFFVFFIFGIVGVQLWAGLLRNRCFLEPYYQTENEDESFICSQGLECTEHNPFKGAINFDNIGYAWIAIFQVITLEGWVDIMYFVMDAHSFYNFIYFILLIIVGSFFMINLCLVVIATQFSSEPGSCYDELLKYLIYILRKASKQSTMQKLLETQSTGPCQSSCKIVVCETFRKIVDSKYFGRGIMIAILINTLSMGIEYHEQPEELTNALEISNIVFTSLFALEMLLKLLVYGPFGYIKSPYNIFDGIIVVISVWEIVGQQGGGLSVLRTFRLMRVLKLVRFMPALQRQLVVLMKTMDNVATFCMLLMLFIFIFSILGMHLFGCKFAGDTLPDRKNFDSLLWAIVTVFQILTQEDWNKVLYNGMASTSSWAALYFIALMTFGNYVLFNLLVAILVEGFQTGDSDSDGELSLEEECGLKKNFLQVPSLYRTSSMHSSQTSASEHQDCNERDSWSVYIFAPQSKFRLVCNKIITHKMFDHVVLVIIFLNCITIAMERPKIEPHSAERIFLTLSNYIFTAIFLAEMTIKVVALGLCFGEYLRSSWNMLDGVLVLISIIDILVSILGMLRVLRLLRTLRPLRVISRAQGLKLVVETLMSSLKPIGNIVVICCAFFIIFGILGVQLFKGKFFVCQDTRITNKSDCAEVRHKYNFDNLGQALMSLFVLASKDGWVDIMYDGLDAVGVDQQPIMNYNP-------------------LFVGVVVENFHKCRQHQEEEEAKRREEKRRRLEKKRRPYYSYSRFRLLIHQMCTTHYLDLFITGVIGLNVITMAMQPKVLDEALKICNYIFTVIFVMESVFKLIAFGFRRFFQDRWNQLDLAIVLLSIMGITLEESLPINPTIIRIMRVLRIARVLKLLKMAVGMRALLDTVMQALPQVGNLGLLFMLLFFIFAALGVELFGDLECDAEGLGRHATFRNFGMAFLTLFRVSTGDNWNGIMKDTLEESTCYTVISPIYFVSFVLTAQFVLVNVVIAVLMKHLEESNK

>Gekko_japonicus_Cav2.2_XP_015276634.1

AWSWAGIHNVVQFFPLTIAS-TPFEYMILATIIANCIVLALEQHLPDGDKTPMSERLDDTEPYFIGIFCFEAGIKIIALGFVFHKGSYLRNGWNVMDFVVVLTGILATAG----DLRTLRAVRVLRPLKLVSGIPSLQVVLKSIMKAMVPLLQIGLLLFFAIVMFAIIGLEFYMGKFHKACFS----NETEERVEFPCGEARQCD-EGPNFGITNFDNILFAVLTVFQCITMEGWTDILYNTNDAAGTWNWLYFIPLIIIGSFFMLNLVLGVLSGEFARRQQQIERELNGYLEWIFKAEEVMSKNDLIHAEEGEDHFTDVCSVMFRFFIRRMVKAQSFYWIVLCVVALNTLCVASVHYDQPEGLTTALYFAEFVFLGLFLTEMSLKMYGLGPRNYFHSSFNCFDFGVIVGSIFEVIPGTSFGISVLRALRLLRIFKVTKYWNSLRNLVVSLLNSMKSIISLLFLLFLFIVVFALLGMQLFGGQFHFNDETPTTNFDTFPTAILTVFQILTGEDWNAVMYQGIESQGMFSSIYFIVLTLFGNYTLLNVFLAIAVDNLANAQELTKDEEEMEEATNQKLALPGNREGDPGSKGEGCEEPHRRHKTRHILPYSSMFVLSPTNPIRRLCHYIVNMRYFEMVILIVIALSSIALAAEDPV-QAESPRNEALKYLDYIFTGVFTFEMVIKMIDLGLILHPYFRDLWNILDFIVVSGALVAFAFSDINTIKSLRVLRVLRPLKTIKRLPKLKAVFDCVVNSLKNVLNILIVYMLFMFIFAVIAVQLFKGRFFYCTDESKDLEKDCRGNKYEFHYDNVLWALLTLFTVSTGEGWPTVLKHSVDATYENQGPSPGFRMEMSIFYVVYFVVFPFFFVNIFVALIIITFQEQGDKVMSECSLEKNERAIDFAISAKPLTRYMPFQYKMWKFVVSPPFEYFIMAMIALNTIVLMMAPAPYEDMLKCLNIVFTSMFSLECVLKIIAFGALNYFRDAWNIFDFVTVLGSITDILVTEADTDNFINLSFLRLFRAARLIKLLRQGYTIRILLWTFVQSFKALPYVCLLIAMLFFIYAIIGMQVFGNIALNDTSINRHNNFQTFLQALMLLFRSATGEAWHEIMLSCLSKNECGSDFAYFYFVSFIFLCSFLMLNLFVAVIMDNFEYLTR

>Gekko_japonicus_Cav2.1_XP_015274358.1

SLFLFSEDNVVRKYAKKITEWPPFEYMILATIIANCIVLALEQHLPDDDKTPMSERLDDTEPYFIGIFCFEAGIKIIALGFAFHKGSYLRNGWNVMDFVVVLTGILATVG----DLRTLRAVRVLRPLKLVSGIPSLQVVLKSIMKAMIPLLQIGLLLFFAILIFAIIGLEFYMGKFHTTCLD---IHTEEIKLEVPCGTARLCP-DGPNYGITQFDNILFAVLTVFQCITMEGWTDLLYYSNDASGAWNWLYFIPLIIIGSFFMLNLVLGVLSGEFARRQQQIERELNGYMEWISKAEEVIKSKTDLLTEEEEDPLADISSVRLRFYIRRMVKTQAFYWTVLSLVALNTLCVAIVHYKQPDWLSDFLYYAEFIFLGLFMSEMFIKMYGLGTRPYFHSSFNCFDCAVIIGSIFEVIPGTSFGISVLRALRLLRIFKVTKYWASLRNLVVSLLNSMKSIISLLFLLFLFIVVFALLGMQLFGGQFNFDQGTPPTNFDTFPAAIMTVFQILTGEDWNAVMYDGIQSQGMVFSIYFIVLTLFGNYTLLNVFLAIAVDNLANAQELTKDEQEEEEAANQKLALIEPPVAEDIDNMKNNKLATNETSDPHMPPYSSMFILSTTNPFRRLCHYIVNLRYFEITILMVIAMSSIALAAEDPV-QPSAPRNNVLRYFDYVFTGVFTFEMVVKMIDLGLVLHQYFRDLWNILDFIVVSGALVAFAFTDINTIKSLRVLRVLRPLKTIKRLPKLKAVFDCVVNSLKNVLNILIVYMLFMFIFAVVAVQLFKGKFFFCTDESKEFEKDCKGKKYDFHYDNVLWALLTLFTVSTGEGWPQVLKNSVDATYENQGPSPGYRMEMSIFYVVYFVVFPFFFVNIFVALIIITFQEQGDKMMEEYSLEKNERAIDFAISAKPLTRHMPFQYRMWQFVVSPPFEYTVMAMIALNTIVLMMASNLYEQVLKMFNIVFTALFSLECILKIIAFGLVNYFRDAWNIFDFVTVLGSITDILVTEGDPNNFISLSFLRLFRAARLIKLLRQGYTIRILLWTFVQSFKALPYVCLLIAMLFFIYAIIGMQVFGNIGIKDSAINEHNNFRTFFQALMLLFRSATGEAWHEIMLSCLKDNECGNEFAYFYFVSFIFLCSFLMLNLFVAVIMDNFEYLTR

>Gekko_japonicus_Cav1.3_XP_015274327.1

-------------------------MLVEQTVGGKTMSYSS----------------EKVEYAFLIIFTIETFLKIIAYGLLLHPNAYVRNGWNLLDFVIVIVGLFSVILEQLTDVKALRAFRVLRPLRLVSGVPSLQVVLNSIIKAMVPLLHIALLVLFVIIIYAIIGLELFIGKMHKSCYF-DDADILAEDDPAPCAFGRQCS-VGPNGGITNFDNFAFAMLTVFQCITMEGWTDVLYWVNDAIGEWPWIYFVSLIILGSFFVLNLVLGVLSGEFSREKQQLEEDLKGYLDWITQAEDIDATNKHASMQASETESVNTENVFNRRRCRAAVKSVSFYWLVIVLVFLNTLTISSEHYNQPDWLTQIQDIANKVLLALFTCEMLVKMYSLGLQSYFVSLFNRFDCFVVCGGIVETIIMSPLGISVFRCVRLLRIFKVTRHWTSLSNLVASLLNSMKSIASLLLLLFLFIIIFSLLGMQLFGGKFNDETQTKRSTFDNFPQALLTVFQILTGEDWNAVMYDGIMAYGMVVCIYFIILFICGNYILLNVFLAIAVDNLADAESLNTAQKEEAEEKERKKNATKVTINEYGEGEEEDKDPYPPCDVPVIPEGSSFFIFSNTNPIRVGCHRLINHHIFTNLILVFIMLSSASLAAEDPI-RSHSFRNNILGYADYVFTSMFTFEIILKLTVFGAFLHKFCRNYFNLLDLLVVGVSLVSFGIQAISVVKILRVLRVLRPLRAINRAKGLKHVVQCVFVAIRTIGNIMIVTTLLQFMFACIGVQLFKGKFYRCTDEAKQNPEECRGQNSDFNFDNVLAAMMALFTVSTFEGWPALLYKAIDSNGENIGPVYNYRVEISIFFIIYIIIIAFFMMNIFVGFVIVTFQEQGEQEYKNCELDKNQRQVEYALKARPLRRYIPYQYKFWYVVNSTGFEYIMFVLIMLNTLCLAMQSKLFNDAMDVLNMVFTAVFTIEMVLKLIAFKPKGYFSDAWNSFDSLVVLGSIVDIVLSENEDSARISITFFRLFRVMRLVKLLSRGEGIRTLLWTFIKSFQALPYVALLIAMLFFIYAVIGMQVFGKVALRDTQINRNNNFQTFLQAVLLLFRCATGEAWQEIMLACLEENTCGSNFAIIYFITFYMLCAFLIINLFVAVIMDNFDYLTR

>Gekko_japonicus_Cav1.2_XP_015278355.1

ALLCLTLKNPIRRACISIVEWXPFEIIILLTIFANCVALAIYIPFPEDDSNATNSNLERVEYLFLIIFTVEAFLKIITYGLLFHPNAYLRNGWNLLDFIIVVVGXVLAILEQATDVKALRAFRVLRPLRLVSGVPSLQVVLNSIIKAMVPLLHIALLVLFVIIIYAIIGLELFMGKMHKTCYLLGVTDTPAEEDPSPCAPGRQCQ-EGPKHGITNFDNFAFAMLTVFQCITMEGWTDVLYWMQDAMGELPWVYFVSLVIFGSFFVLNLVLGVLSGEFSREKQQLEEDLKGYLDWITQAEDIDSETESVNTDNVAGGDIEGENCFCRRKCRAAVKSNVFYWLVIFLVFLNTLTIASEHYNQSDWLTEVQDTANKVLLALFTAEMLLKMYSLGLQAYFVSLFNRFDCFIVCGGILETIIMSPLGISVLRCVRLLRIFKITRYWNSLSNLVASLLNSVRSIASLLLLLFLFIIIFSLLGMQLFGGKFNDEMQTRRSTFDNFPQSLLTVFQILTGEDWNSVMYDGIMAYGMLVCIYFIILFICGNYILLNVFLASTVDNVADAESLTSAQKEEEEEKERKKLANKSNVDEYQPNENEEKNPYPTTETPGMPDASAFFIFSPSNRFRVHCHRIVNNNIFTNLILFFILLSSISLAAEDPV-RHSSVRNQILFYFDIFFTVIFTIEIALKMTAYGAFLHKFCRNYFNILDLLVVSVSLISFGIQAINVVKILRVLRVLRPLRAINRAKGLKHVVQCVFVAIRTIGNIVIVTTLLQFMFACIGVQLFKGKLYSCSDSSKQTEAECKGENSKFDFDNVLTAMMALFTVSTFEGWPELLYRSIDSHLEDVGPIYNHRVEISIFFIIYIIIIAFFMMNIFVGFVIVTFQEQGEQEYKNCELDKNQRQVEYALKARPLRRYIPYQYKVWYVVNSTYFEYLIFILIMLNTICLAMQSCLFKEAMNILNMLFTGLFTVEMVLKLIAFKPKGYFSDPWNVFDFLIVIGSIIDVILSEAEENSRISITFFRLFRVMRLVKLLSRGEGIRTLLWTFIKSFQALPYVALLIVMLFFIYAVIGMQVFGKIALNDTEINRNNNFQTFPQAVLLLFRCATGEAWQEIMLACLDEYSCGSSFAIFYFISFYMLCAFLIINLFVAVIMDNFDYLTR

>Gekko_japonicus_Cav1.1_XP_015267320.1

SLFCLTLQNPVRKACIAIVEWKPFETIILLTIFANCVALAIYLPMPEDDSNKANSRLEKLEYFFLMVFAIEAVLKIIAYGFLFHADAYLRNGWNVLDFTIVFLGVFTVILERISDVKALRAFRVLRPLRLVSGIPSLQVVLNSIGKAILPLFHIAVLVVFMLIIYAIVGEELFKGKMHKTCYYTDIIATVENEEPSPCTNGRHCT-PGPNNGITHFDNFGFAMLTVYQCISMEGWTKVLYWVNDAIGEWPWIYFVSLILLGSFFILNLILGVLSGEFTREKQQLEEDMKGYMDWIVHAEVIDRGEGMMPSDEGGSETESLYEILFRRKCREVVKSRFFYWFVILIVSLNTLSIASEHHRQPGWLTHVQDIANRVLLALFTVEMILKMYALGLHQYFMSIFNRFDCLVVCTGILEIISMSPLGISVLRCIRLLRIFKITKYWTSLNNLVASLLNSVRSIISLLTLLFLYIVIFALLGMQLFGGRFDDDAEIRRSTFDNFPQALISVFQVLTGEDWTSIMYDGIMAYGILVCIYFIILFVCGNYILLNVFLAIAVDNLAEAETLTSAQKAKEEEKKRKKMAAKLKVDEFESNVNEIKDPYPSADFPGMPEASAFFIFSPTNKIRVLCHRIVNATWFTNFILLFILLSSISLAAEDPI-RAESFRNQILEYFDYVFTSVFTVEIVLKMTAYGAFLHKFCRNSFNILDLLVVAVSLISMGIQAISVVKILRVLRVLRPLRAINRAKGLKHVVQCVFVAIKTIGNIVLVSALLQFMFACIGVQLFKGKFYSCTDPLKITEEECRGFHNEFHFDNVLSAMMSLFTVSTFEGWPKLLYQAIDTHTEDMGPIYNYRMGIAIYFIVYLILIAFFMMNIFVGFVIVTFQEQGETEYKSCELDKNQRQVQYALKARPLKCYIPYQYQIWYVVTSSYFEYLMFFLITLNTICLGMQSETMNQVSDILNVVFTLLFTVEMILKLIAFKAKGYFGDPWNVFDFLIVIGSIIDVILSQDDDNGRISITFFRLFRVLRLVKLLSRAEGIRNLLWTFIKSFQALPHVALLIVMLFFIYAVIGMQMFGKVGLVDTQINRNNNFQTFPQAVLLLFRCATGEAWQEVLLASYEEYSCGTGFAYFYFISFYMICAFLIINLFVAVIMDNFDYLTR

>Gallus_gallus_Cav3.3_XP_015144216.1

VFFCLKQTTSPRSWCIKMVCNPWFECVSMMVILLNCVTLGMYQPCEDMDLSDRCKILQVFDDFIFIFFAMEMVLKMVALGI-FGKKCYLGDTWNRLDFFIVMAGMVEYSLDLQNNLSAIRTVRVLRPLKAINRVPSMRILVNLLLDTLPMLGNVLLLCFFVFFIFGIIGVQLWAGLLRNRCFMPPYYQPEEDDEMFICSLGHECCNANPHKGAINFDNIGYAWIVIFQVITLEGWVEIMYYVMDAHSFYNFIYFILLIIVGSFFMINLCLVVIATQFSMEPGDCYEEIFQYVCHIMRKAKRRICQQHNPLDCPPQGLVQPIAVEVRVKLRGIVDSKYFNRGIMIAILVNTISMGIEHHEQPEELTNILEISNVVFTSMFALEMILKLAAFGLFDYLRNPYNIFDSIIVIISIWEIIGQSDGGLSVLRTFRLLRVLKLVRFMPALRRQLVVLMKTMDNVATFCMLLMLFIFIFSILGMHIFGCKFSGDTVPDRKNFDSLLWAIVTVFQILTQEDWNVVLYNGMASTSSWAALYFVALMTFGNYVLFNLLVAILVEGFQAGDSYSEEDQSSSNMEELDQFQASIPSGHRGTCRPGTTSGGSEHQDCNLREDWSIYLFSPQNRFRLLCQTIIAHKLFDYVVLAFIFLNCITIALERPQIEHRSTERIFLTVSNYIFTAIFVAEMTLKVVSLGLYFGDYLRSSWNVLDGFLVFVSLIDIVVSILGVLRVLRLLRTLRPLRVISRAPGLKLVVETLISSLKPIGNIVLICCAFFIIFGILGVQLFKGKFYHCLDIRITNRSDCVAVHHKYNFDNLGQALMSLFVLASKDGWVNIMYNGLDAVAVDQQPVTNNNPWMLLYFISFLLIVSFFVLNMFVGVVVENFHKCRQHQEAEEARRREEKRRRLEKKRRPYYAYCPIRLLIHSVCTSHYLDIFITFIICLNVVTMSLQPVSLETALKYCNYLFTTVFVLEAVLKLVAFGLRRFFKDRWNQLDLAIVLLSIMGITLEEALPINPTIIRIMRVLRIARVLKLLKMATGMRALLDTVVQALPQVGNLGLLFMLLFFIYAALGVELFGKLVCNDEGMSRHATFENFGMAFLTLFQVSTGDNWNGIMKDTLDDRSCLQFISPLYFVSFVLTAQFVLINVVVAVLMKHLDDSNK

>Gallus_gallus_Cav3.2_XP_015149910.1

AFFCLKQTTRPRSWCLRLVCNPWFEHVSMLVILLNCVTLGMFQPCEDVEQSERCTILEAFDDFIFAFFAVEMVIKMVALGI-FGQKCYLGDTWNRLDFFIVMAGMMEYSL----SLSAIRTVRVLRPLRAINRVPSMRILVTLLLDTLPMLGNVLLLCFFVFFIFGIVGVQLWAGLLRNRCFLHPYYRTDEAEENFICSSKVECTDVNPHNGAINFDNIGYAWIAIFQVITLEGWVDIMYYVMDAHSFYNFIYFILLIIVGSFFMINLCLVVIATQFSSEPGSCYEELLKYICHIFRKVKRRRTPSRLSGLSVACPLPSPPGGAFGSKLKRIVESKYFNRGIMIAILINTLSMGIEYHEQPDELTNALEISNIVFTSMFALEMLLKLLAFGLFGYIKNPYNIFDGIIVVISVWEIIGQSDGGLSVLRTFRLLRVLKLVRFMPALRRQLVVLMKTMDNVATFCMLLMLFIFIFSILGMHLFGCKFSGDTVPDRKNFDSLLWAIVTVFQILTQEDWNVVLYNGMASTSSWAALYFVALMTFGNYVLFNLLVAILVEGFQAGDSDTDEDKNFDDDFEKLKDLLQLPPMRHSLSISPMAVLPAEYQDCNSHEDWSLYLFSPQNRFRVMCQKVIAHKMFDHVVLVFIFLNCITIALERPDIDPHSTERIFLSVSNYIFTAIFVAEMMVKVVALGFFSGEYLQSSWNVLDGVLVFVSIIDIIVSILGVLRVLRLLRTLRPLRVISRAPGLKLVVETLISSLRPIGNIVLICCAFFIIFGILGVQLFKGKFYYCDDVKITTKTDCTNVRRKYNFDNLGQALMSLFVLSSKDGWVNIMYDGLDAVGIDQQPIQNHNPWMLLYFISFLLIVSFFVLNMFVGVVVENFHKCRQHQEAEEARRREEKRRRLEKKRRPYYAYSPARKYIHTLCTSHYLDLFITFIIGVNVITMSMQPKSLDEALKYCNYVFTIVFVFEAVLKLVAFGFRRFFKDRWNQLDLAIVLLSIVGITLEEALPINPTIIRIMRVLRIARVLKLLKMATGMRALLDTVVQALPQVGNLGLLFMLLFFIYAALGVELFGKLDCSEEGLSRHATFTNFGMAFLTLFRVSTGDNWNGIMKDTLEDKHCLPVISPVYFVTFVLIAQFVLVNVVVAVLMKHLEESNK

>Gallus_gallus_Cav3.1_XP_015150965.1

VFFYLSQESRPRSWCLRLVCNPWFERVSMLVILLNCVTLGMFHPCEDIADSPRCRILQSFDDFIFAFFAVEMIVKMIALGI-FGKKCYLGDTWNRLDFFIVIAGMLEYSL--DLSFSAVRTVRVLRPLRAINRVPSMRILVTLLLDTLPMLGNVLLLCFFVFFIFGIVGVQLWAGLLRNRCFLERYYQTENEDENFICSQGLECTEHNPFKGAINFDNIGYAWIAIFQVITLEGWVDIMYFVMDAHSFYNFIYFILLIIVGSFFMINLCLVVIATQFSSEPGSCYDELLKYLVYIARKGSKQSVHVTDKGDGFRGPCPSSCKIVVCETFQKIVDSKYFGRGIMVAILINTLSMGIEYHEQPEELTNALEISNIVFTSLFALEMLLKVLVYGPFGYIKNPYNIFDGIIVVISVWEIVGQQGGGLSVLRTFRLMRVLKLVRFMPALQRQLVVLMKTMDNVATFCMLLMLFIFIFSILGMHLFGCKFAGDTLPDRKNFDSLLWAIVTVFQILTQEDWNKVLYNGMASTSSWAALYFIALMTFGNYVLFNLLVAILVEGFQTGDSDSEGELSLEEEGGLKKNLLQVPSLYRTSSMHSSRTSTSEHQDCNERDSWSIYVFAPHSRFRLMCNKIITHKMFDHIVLVIIFLNCITIAMERPKIEPHSAERIFLTLSNYIFTVIFLTEMTVKVVALGLCFGEYLKSSWNVLDGVLVLISVIDILVSILGMLRVLRLLRTLRPLRVISRAQGLKLVVETLMSSLKPIGNIVVICCAFFIIFGILGVQLFKGKFFICQDTRITNKSDCAEVRHKYNFDNLGQALMSLFVLASKDGWVDIMYNGLDAVGVDQQPVMNYNPWMLLYFISFLLIVAFFVLNMFVGVVVENFHKCRQHQEEEEAKRREEKRRRLEKKRRPYYSYSRFRFLIHQMCTSHYLDLFITGVIGLNVITMAMQPKVLDEALKICNYIFTVIFVLESVSKLIAFGFRRFFQDRWNQLDLAIVLLSIMGITLEESLPINPTIIRIMRVLRIARVLKLLKMAVGMRALLDTVMQALPQVGNLGLLFMLLFFIFAALGVELFGDLECDDEGLGRHATFRNFGMAFLTLFRVSTGDNWNGIMKDTLQESTCYTVISPIYFVSFVLTAQFVLVNVVIAVLMKHLEESNK

>Gallus_gallus_Cav2.3_XP_015145962.1

SLFLFGEDNIVRKYAKKLIDWPPFEYMILATIIANCIVLALEQHLPEDDKTPMSRRLEKTEPYFIGIFCFEAGIKIVALGFVFHKGSYLRNGWNVMDFIVVLSGILATAGTHFNDLRTLRAVRVLRPLKLVSGIPSLQIVLKSIMKAMVPLLQIGLLLFFAILMFAIIGLEFYSGKLHRACYTNNSGELEELDPPHPCG-VQGCP-IGPNDGITQFDNILFAVLTVFQCITMEGWTTVLYNTNDALGTWNWLYFIPLIIIGSFFVLNLVLGVLSGEFARRQQQIERELNGYRAWIDKAEEVMNRTDAMNRDSSDEHCVDISSVLLRISVRHMVKSQVFYWIVLSLVALNTACVAIVHHNQPAWLTHFLYYAEFLFLGLFLLEMSLKMYGMGPRLYFHSSFNCFDCGVTVGSIFEVVPGTSFGISVLRALRLLRIFKITKYWASLRNLVVSLMSSMKSIISLLFLLFLFIVVFALLGMQLFGGGFNFIDGTPSANFDTFPAAIMTVFQILTGEDWNEVMYNGIRSQGMWSSIYFIVLTLFGNYTLLNVFLAIAVDNLANAQELTKDEQEEEEAFNQKHALAPEPSRGMEGSLGEQDCSSPDTSEQAMVPHSSMFIFSTTNPVRRACHYIVNLRYFEMCILLVIAASSIALAAEDPV-LTNSDRNKVLRYFDYVFTGVFTFEMVIKMIDQGLILQDYFRDLWNILDFIVVVGALVAFALADIKTIKSLRVLRVLRPLKTIKRLPKLKAVFDCVVTSLKNVFNILIVYKLFMFIFAVIAVQLFKGKFFYCTDSSKDTEKDCIGKRHEFHYDNIIWALLTLFTVSTGEGWPQVLQHSVDVTEEDRGPSRSNRMEMSIFYVVYFVVFPFFFVNIFVAFIIITFQEQGDKMMEECSLEKNERAIDFAISAKPLTRYMPFQYRVWHFVVSPSFEYTIMAMIALNTVVLMMAPYTYELALKYLNIAFTMVFSLECVLKIIAFGFLNYFRDTWNIFDFITVIGSITEIILTDLVNTSSFNMSFLKLFRAARLIKLLRQGYTIRILLWTFVQSFKALPYVCLLIAMLFFIYAIIGMQVFGNIKLDESHINRHNNFRSFLGSLMLLFRSATGEAWQEIMLSCLENERCGTDLAYVYFVSFIFFCSFLMLNLFVAVIMDNFEYLTR

>Gallus_gallus_Cav2.2_XP_015134766.1

SLFIFSEDNVIRKYAKRITEWPPFEYMILATIIANCIVLALEQHLPDGDKTPMSERLDDTEPYFIGIFCFEAGIKIIALGFVFHKGSYLRNGWNVMDFVVVLTGILATAG----DLRTLRAVRVLRPLKLVSGIPSLQVVLKSIMKAMVPLLQIGLLLFFAIVMFAIIGLEFYMGKFHKTCFS----NETGEEVGFPCGEARQCE-QGPNYGITNFDNILFAVLTVFQCITMEGWTDILYNTNDAAGTWNWLYFIPLIIIGSFFMLNLVLGVLSGEFARRQQQIERELNGYLEWIFKAEEVMSKNDLIHAEEGEDHFTDICSVMFRFFIRRMVKAQSFYWIVLCVVALNTLCVAMVHYDQPEKLTTALYFAEFVFLGLFLTEMSLKMYGLGPRNYFHSSFNCFDFGVIVGSIFEVIPGTSFGISVLRALRLLRIFKVTKYWNSLRNLVVSLLNSMKSIISLLFLLFLFIVVFALLGMQLFGGQFNFRDETPTTNFDTFPAAILTVFQILTGEDWNAVMYHGIESQGMFSSIYFIVLTLFGNYTLLNVFLAIAVDNLANAQELTKDEEEMEEATNQKLALSGNREGEPGSKGENGEEPHRRHRFRSILPYSSMFILSPTNPIRRLFHYIVNLRYFEMVILIVIALSSIALAAEDPV-QAESPRNDALKYLDYIFTGVFTFEMVIKMIDLGLLLHPYFRDLWNILDFIVVSGALVAFAFSDINTIKSLRVLRVLRPLKTIKRLPKLKAVFDCVVNSLKNVLNILIVYMLFMFIFAVIAVQLFKGRFFYCTDESKELEKDCRGKKYEFHYDNVLWALLTLFTVSTGEGWPTVLKHSVDATYEEQGPSPGYRMEMSIFYVVYFVVFPFFFVNIFVALIIITFQEQGDKVMSECSLEKNERAIDFAISAKPLTRYMPFQYKMWKFVVSPPFEYFIMVMIALNTIVLMMAPEAYEEMLKCLNIVFTSMFSMECVLKIIAFGVLNYFRDAWNVFDFVTVLGSITDILVTEADTDNFINLSFLRLFRAARLIKLLRQGYTIRILLWTFVQSFKALPYVCLLIAMLFFIYAIIGMQVFGNIALNDTSINRHNNFRTFLQALMLLFRSATGEAWHEIMLSCLTKNECGSEFAYFYFVSFIFLCSFLMLNLFVAVIMDNFEYLTR

>Gallus_gallus_Cav2.1_tr_A0A2H4LIM4

SLFLFSEDNAVRKYAKRITEWPPFEYMILATIIANCIVLALEQHLPDDDKTPMSERLDDTEPYFIGIFCFEAGIKIIALGFAFHKGSYLRNGWNVMDFVVVLTGILATVG----DLRTLRAVRVLRPLKLVSGIPSLQVVLKSIMKAMIPLLQIGLLLFFAILIFAIIGLEFYMGKFHTTCFD----LVTNEIKVVPCGTARICP-EGPNYGITQFDNILFAVLTVFQCITMEGWTDLLYYSNDASGTWNWLYFIPLIIIGSFFMLNLVLGVLSGEFARRQQQIERELNGYMEWISKAEEVISKTDLLSPEDAEEQLADIASVRMRFYIRRVVKTQAFYWTVLSLVALNTLCVAIVHYDQPEWLSDFLYYAEFIFLGLFMSEMFIKMYGLGTRPYFHSSFNCFDCAVIIGSIFEVIPGTSFGISVLRALRLLRIFKVTKYWASLRNLVVSLLNSMKSIISLLFLLFLFIVVFALLGMQLFGGQFNFDTGTPPTNFDTFPAAIMTVFQILTGEDWNAVMYDGIKSQGMVFSVYFIVLTLFGNYTLLNVFLAIAVDNLANAQELTKDEQEEEEAANQKLALPHPAGPGPAAAARGGGHRQHEIPACPMVPYSSMFILSPTNPFRRLCHYIVNLRYFEMCILMVIAMSSIALAAEDPV-QPNAPRNNVLRYFDYVFTGVFTFEMVIKMVDLGLVLHQYFRDLWNILDFIVVSGALVAFAFTDINTIKSLRVLRVLRPLKTIKRLPKLKAVFDCVVNSLKNVLNILIVYMLFMFIFAVVAVQLFKGKFFYCTDESKEFEKDCRGKKYDFHYDNVLWALLTLFTVSTGEGWPQVLKHSVDATYENQGPSPGYRMEMSIFYVVYFVVFPFFFVNIFVALIIITFQEQGDKMMEEYSLEKNERAIDFAISAKPLTRHMPFQYRMWQFVVSPPFEYTIMAMIALNTIVLMMASDAYENVLKMFNNVFTSLFSLECLLKIMAFGVLNYFRDAWNVFDFVTVLGSITDILVTE--GNNFINLSFLRLFRAARLIKLLRQGYTIRILLWTFVQSFKALPYVCLLIAMLFFIYAIIGMQVFGNIGIEDSAITQHNNFRTFFQALMLLFRSATGEAWHEIMLSCLKEDECGNEFAYFYFVSFIFLCSFLMLNLFVAVI--------R

>Gallus_gallus_Cav1.3_XP_015148473.1

ALFCLSLNNPIRRACISLVEWKPFDIFILLSIFANCVALAVYIPFPEDDSNSTNHNLEKVEYAFLIIFTVETFLKIIAYGLLLHPNAYVRNGWNLLDFVIVVVGLFSVILEQLTDVKALRAFRVLRPLRLVSGVPSLQVVLNSIIKAMVPLLHIALLVLFVIIIYAIIGLELFIGKMHKSCFL-IDSDILVEEDPAPCAFGRQCV-VGPNGGITNFDNFAFAMLTVFQCITMEGWTDVLYWMNDAMGELPWVYFVSLVIFGSFFVLNLVLGVLSGEFSREKQQLEEDLKGYLDWITQAEDIDSETESVNTENVSGEGENPACCFNRRKCRAAVKSVTFYWLVIVLVFLNTLTISSEHYNQPDWLTQIQDIANKVLLALFTCEMLVKMYSLGLQAYFVSLFNRFDCFVVCGGIVETIIMSPLGISVFRCVRLLRIFKVTRHWASLSNLVASLLNSMKSIASLLLLLFLFIIIFSLLGMQLFGGKFNDETQTKRSTFDNFPQALLTVFQILTGEDWNAVMYDGIMAYGMIVCIYFIILFICGNYILLNVFLAIAVDNLADAESLNTAQKEEAEEKERKKNASKVTIAEYGEGEDEDKDPYPPCDVPVIPEGSAFFIFSSTNPIRVGCHRLINHHIFTNLILVFIMLSSVSLAAEDPI-RSHSFRNNILGYFDYAFTAIFTVEILLKMTAFGAFLHKFCRNYFNLLDLLVVGVSLVSFGIQAISVVKILRVLRVLRPLRAINRAKGLKHVVQCVFVAIRTIGNIMIVTTLLQFMFACIGVQLFKGKFYKCTDEAKQNPEECRGQNSDFNFDNVLSAMMALFTVSTFEGWPALLYKAIDSNGENVGPVYNYRVEISIFFIIYIIIIAFFMMNIFVGFVIVTFQEQGEQEYKNCELDKNQRQVEYALKARPLRRYIPYQYKFWYVVNSTGFEYIMFVLIMLNTLCLAMQSKLFNDAMDIMNMVFTGVFTVEMVLKLIAFKPKGYFSDAWNTFDSLIVIGSIVDVVLSESEDSARISITFFRLFRVMRLVKLLSRGEGIRTLLWTFIKSFQALPYVALLIAMLFFIYAVIGMQVFGKVAMRDNQINRNNNFQTFPQAVLLLFRCATGEAWQEIMLACLEEYTCGSNFAIIYFISFYMLCAFLIINLFVAVIMDNFDYLTR

>Gallus_gallus_Cav1.2_XP_015142138.1

ALLCLTLKNPIRRACISIVEWKPFEIIILLTIFANCVALAIYIPFPEDDSNATNSNLERVEYLFLIIFTVEAFLKVIAYGLLFHPNAYLRNGWNLLDFIIVVVGLFSAILEQATDVKALRAFRVLRPLRLVSGVPSLQVVLNSIIKAMVPLLHIALLVLFVIIIYAIIGLELFMGKMHKTCYHGGLIDTPAEDDPSPCAPGRQCQ-EGPKHGITNFDNFAFAMLTVFQCITMEGWTDVLYWVNDAIGDWPWIYFVTLIIIGSFFVLNLVLGVLSGEFSREKQQLEEDLKGYLDWITQAEDIDSETESVNTDNVPGTDIEGENCFCRRKCRAAVKSNVFYWLVIFLVFLNTLTIASEHYNQPDWLTEVQDTANKVLLALFTAEMLLKMYSLGLQAYFVSLFNRFDCFIVCGGILETIIMSPLGISVLRCVRLLRIFKITRYWNSLSNLVASLLNSVRSIASLLLLLFLFIIIFSLLGMQLFGGKFNDEMQTRRSTFDNFPQSLLTVFQILTGEDWNSVMYDGIMAYGMLVCIYFIILFICGNYILLNVFLAIAVDNLADAESLTSAQKEEEEEKERKKLATKINMDDYQPNENEEKSPYPTTEAPAMPDASAFFIFSPNNRFRVHCHRIVNDNIFTNLILFFILLSSISLAAEDPV-RHLSFRNQVLFYFDIVFTVIFTIEIALKMTAYGAFLHKFCRNYFNILDLLVVSVSLISFGIQAINVVKILRVLRVLRPLRAINRAKGLKHVVQCVFVAIRTIGNIVIVTTLLQFMFACIGVQLFKGKLYSCTDSSKQTEAECRGENSKFDFDNVLTAMMALFTVSTFEGWPELLYRSIDSHMEDVGPIYNHRVEISIFFIIYIIIIAFFMMNIFVGFVIVTFQEQGEQEYKNCELDKNQRQVEYALKARPLRRYIPYQYKVWYVVNSTYFEYLMFVLILLNTICLAMQSCMFKEAMNILNMLFTGLFTVEMVLKLIAFKPKGYFSDPWNVFDFLIVIGSIIDVILSEAEENSRISITFFRLFRVMRLVKLLSRGEGIRTLLWTFIKSFQALPYVALLIVMLFFIYAVIGMQVFGKIALNDTEINRNNNFQTFPQAVLLLFRCATGEAWQEIMLACLADHSCGSSFAVFYFISFYMLCAFLIINLFVAVIMDNFDYLTR

>Gallus_gallus_Cav1.1_NP_001292076.1

ALFCLTLQNPLRKACISIVEWKPFEIIILLTIFANCVALAIYQPMPEDDTNVANSSLEKLEYVFLIFFAIEAMLKIIAYGFLFHTDAYLRNGWNVLDFSIVSLGLVTMTLEQINDVKALRAFRVLRPLRLVSGVPSLQVVLNSIIKAMVPLLHIALLVLFMIIIYAIVGQELFKGKMHKTCYYTDVIATVGSEKPAPCTTGRHCS-PGPNNGITHFDNFGFAMLTVYQCITMEGWTEVLYWVNDAIGEWPWIYFVSLILLGSFFVLNLVLGVLSGEFTREKQQLEEDMKGYMDWITHAEVMDRGEGMMPLDEGGSETESLYEIMFRRKCRDVVKSKFFYWLVILLVALNTLSIASEHHFQPEWLTIVQDNANRVLLALFVAEMLLKMYALGLRQYFMSLFNRFDCFVVCAGVLEIITLSPLGISVLRCIRLLRIFKITRYWTSLSNLVASLLNSVRSIASLLLLLFLFIIVFALLGMQLFGGMYDEDMEVRRSTFDNFPQALISVFQILTGEDWNSIMYNGIMAYGMLVCIYFIILFVCGNYILLNVFLAIAVDNLAEAESLTSAQKAKAEERKRRKMSAKLKVDEFESNVNEIKDPYPSADFPGMPEASSFFIFSPTNKFRMLCHRIVNATWFTNFILLFILLSSISLAAEDPI-RAESFRNQILGYFDIGFTSVFTVEIVLKMTAYGAFLHKFCRNSFNILDLLVVAVSLISMGFETISVVKILRVLRVLRPLRAINRAKGLKHVVQCVFVAIKTIGNIVVVTTLLQFMFACIGVQLFKGKFYSCTDPSKLTEKECRGFHNAFHFNNVLSAMMSLFTVSTFEGWPELLYRAIDTNDENKGPIYNYRVEIAMFFIIYIILIAFFMMNIFVGFVIVTFQEQGESEYKNCELDKNQRQVQYALKARPLRRYIPYQYQIWYVVTSSYFEYLMFFLIMLNTICLGMQSAEMNHVSDILNVAFTVLFTLEMILKLMAFKAKGYFGDPWNVFDFLIVIGSIIDVILSEPDDNSRVSITFFRLFRVMRLVKLLSRGEGVRTLLWTFIKSFQALPYVALLIVMLFFIYAVIGMQMFGKIAMVDTQINRNNNFQTFPQAVLLLFRCATGEAWQEILLDCSEEYTCGTGFAYFYFISFYMLCAFLIINLFVAVIMDNFDYLTR

>Fungi_schpom_CCH1_NP_593894.1

ANETPGRNLPFSPILRQIYNHPLYNIFIFVVIVLHAVLLMIRSDDPHDK--------QTIDYLIIVIYTLEMLLKIYLFGFLYDSSSYLRHSWNRVDFVAIVALWISVIG----GIRLFSMIACLRLTRLLNITRKTETILKSLKESSTPLVQVVSFNAFFGVMIAILGVQFFKASLNRQCVWLGDYGDQYLPTGQFCGGGFICAVENPYSNTVSFDNFFNSLELIFVIMSSNGFTDIMYDIMDAEYFVSCLLFIISAYFLTLWLMSLVIAVVTSSFIKEQKSVDKHLIRN-----------------------------------KLCEKYLFYSNFIW--ISFIVAQFVTLCTQTYDQTSSTANRYLIFYACVDFLLAAEVILRFFAYDYRLFFRRYTNLVDIVLAVLNLVTLLRKNPVAFGWLSIFAIARIYRCILLIPYTRKIAKLLFSNFKQLLNLMLFLVIVLFIASLCAVRLFQDLPNGDSDDDAISFATTYESFLYMYQILTSENWTDVMF-AIQARLSWPGAFFTLWFLFSNNVVLSMFIAVIQVNFAPSEKMEQLKMYLARLLRNYNPFFVMRDDPFSQAYFKRVIGIRWEKDMNPKYNDVFWVIKPSNRIRRFCQRLVPYQWVYRVIQVFIYA-CILTAVATPIYERDHLLNDWFVWTEVAFATIFTIEAAIKIIADGFCITPYLRSTWNCIDFFVLVTLWINLLTS--LLSRAFRAFKALRVLRLINLTQTSQRMFHALISGFFKIFSAAVVSATLLIPFALWAKNVFGGLLYSCNDDNVLSASQCV-NPPDYDFDRFPHALLALFEIASIEGWVDIMRSVMDITGFNNQPQTNASSGNAMFFVLFNLVSMIYILTLFIAIIISNYAERTGSAFFTAE-QRAWLERRKIKSMRPSKRAIRLRGLCYDFAVQKIWRRTFTGLYIVHLLFLLTCPIAYTYVRNSIFLILSICYTINICVKVYGLSFYYFFHSFWNMFDVVVTLGSLTNIILAKNRSLTLLQTTLLVLVTV----HLIPKFDNFDQLSKTVVASLPSIFSLIATWIVLYITFAIAFNQIFGLTKL--LNGGPNKNFRSIRNALVLLFTMTFGEGWNDVMHDTIYNSDCGKPWAYGLFIAWNIISMYIFVNMFITVVFDNFSYIHT

>Fungi_saccer_CCH1_NP_011733.3

SLGVFSPTNPLRIKIARFLLHRRYSLLYNTLLTFYAILLAIRTYNPHN-VVFLYRFSNWTDYFIFILFTGNDIAKIIAFGFWFGPRAFARSSWNRIDLVSSVSFWLGMFLSIKSRIKPLAILRILRLVNVDTGMPSI---LRGLKYGIPQLVNVSSMLVYFWIFFGILGVQIFQGSFRRQCVWNPEDPTDTYQYDQFCGGGFLCPNANPYNGRISFDNIVNSMELVFVIMSANTFTDLMYYTMDSDEMAACLFFIVCIFVLTIWLLNLLIAVLVSSFESRKTGYARIVTGY--W------------------------------------GLAIYSHVEFIFVILIICDIGMRASVKVSTSANCNNILLKTDRGISIVLFIESLARLVLYNMWKFLTKPSYVYDFIISIITLVISCVEGVLGYAWLSIFHISRFYRVIISFNLTKKLWKQILSNGVMIWNLSSFYFFFTFLVAIIMAVYFEGVIPEEMADQPFGMYSLPNSFLSLFIIGSTENWTDILY-ALQKHSTFCSVFFIIWFLLSNSVILNIFIALISESM----EVKEEEKRPQQIKHYLKFVLTIGVPSLKRLRMFANNPFYKNSDVVPLFNYSYYFFSPQHRFRRFCQRLVGSRFFEDSVFVFIFALATILLIVTPLYRMHHKMGTWSSALDCAFIGAFSIEFIVKTVADGFIYSPYLRNPWNFIDFCVLISMWINLIAY---LSRIFKGLTALRALRCLTISNTARQTFNVMFDGLNKIFEAGLISLSLLFPFTVWGLSIFKGRLGTCNDGS-LGRADCYNQQPYLHLDSFASAFSSLYQIISLEGWVDLLENMMNSSGIGTPATVMGSAGNALFLVLFNFLSMVFILNLFVSFIVNNQARTTGSAYFTIE-EKAWLEQKLLSQAKP---AIPVRQFFYQLVEKKYYASFLQVVLYLHIIMLLSNPGNLIGYQGVYFMFSTSVFLIQEALHMCGEGPRLYFRQKWNSIRLSIIIIAFINAAF--HVPASHYWFHNIKGFFLLVIFFIIPQNDTLTELLETAMASLPPILSLTYTWGVLFLVYAIALNQIFGLTRL--SNTTDNINFRTVIKSMIVLFRCSFGEGWNYIMADTVTYTDCGETYAYLLLMSWNIISMYIFVNMFVSLIIGNFSYVYR

>Exaiptasia_pallida_Cav3b_XP_020912298.1

SCMFLGVRTLPRRWLLVVYSNRWFERVSMFVILVNCVTLAMDDPYDRQCESLRCQILQSIEHFIFAFFLLELVVKMIAMGV-WGKKGYMQEPWNRLDFVIIAVGTVEKLMEGSDYLTIIRAFRVLRPLRAINKVPSIRILVTLLLDTLPMLGNVLLLSFLIFFVFGIIAVQLWQGHLRNRCFIESLFYKPSFDKPYICSLDRTCTGPNPFYGTTSFDNIVIAWIAIFQVITLEGWSEIMYLVQDSHSFWNWIYFVILVVIGAFFLVNLCLVVITMQFQ-QAYSCFQTLKRYQEFCCSRQAFKTCGKRSNTEVVTNHSVHQRRHRLRNFFRRLAESKQFVHFIMISILVNMICMGLEHYKQPERLTAALEQSNIIFVSIFSLEMLINLMAYGFTGYLSQVQNVFDGFVVVVSVAELL--DYARLSVFRSIRLLRIFKLVR---PVRYQLLVILRTMNSVVTFFGLLFLFMFAFCILGMNLFGGKFFNEKVLRRSNFDSFLWAMVTVFQILTQENWNHVMYDGMRVTNNWAALYFIALMIVGYYVLFNLLVAILVEGFTNEDAQSEARRSSIKQGQTSKYSTTSATQESIREFVNNQVSHENKQDLKSRREWSLFMLSPDDRFRTTLLTICSHKIFDYVILVFILMSCVVLAMEGPGTDNNATVREIIDISMLLFTVVFTIEMMMKVVAMGFIIGSYLKDGWNVLDFILVIISWLDVIITVLGTLKVFRALRTLRPLRMIRRAPGLKLVVQTLLYSLKPIGNTVLIAAIFFAMFGILGVQLFKGKFYHCEDNHVIDRAQCLNVNRRYNFDNLFQALISLFVVSTRDGWVEVMHHGIDAVDIDMQPKVNHAEWCLVYFIPFLLLGGFLVINMIVGVVVENFQRCRDEEQQNSKKKRHKSETNYNEQDDSYSVYNVARRWIYHVCMHQYWDIAIAIIICVNVICMSMMAPTFIIFVETSNYFFTAVFVIEALLKVVALGWLRYIKDRWNIIDLLIVVLSVSGILLDDELPINPTVIRTLRVLRIVRVLKLVKLAKGVRSLLDTLFEALPQVANLGLLFLLLFFIYSCLGIQLFGSLECSYEGLGQHANFNSFGSSMLTLFRVATGDNWNGILKDTLRKNCCIKYTAPLYFVTFVLSAQFVLVNVVIAVLMKHLKESKE

>Exaiptasia_pallida_Cav3a_XP_020900273.1

AFYFLKRXKYPRLLFVKLVSWSYFERISIFVILCNCVTLGLYDPFDPDCLTQRCQILEKIERAIYAFFVVEMLCKWIAMGI-FGKLGYLSDNWNKLDCFIVAAGTFELFYDKGKYMTAVRAIRVLRPLRAINRVPSIRILVTLLLDTLPMLGNVLAMCSLIFSIFGIVGVQMWQGLLRNRCMLLSSYYTP-SKGGLVCSLGRECLGENPDHDTISFDNILIAWVAIFQVITLEGWSDIMYYIQDANSPWDFIYFVVLIVIGSYFMTNLCLVVITTQFQFGRDGCWVEILKYIAHVFRRTKRRINGQSASSDVAAHSFVSAASLKFRHALRDWVEGKMFMYFIMGAIFINTLSMGIEFYGQPQEMTDVLEILNYIFTGIFALEMLVKLIALGLYGYIKDAFNLFDGAIVIVSIVELFGDGNNNISVLRSFRLLRIFKIVRFLPALKRQLLVMIHTLDNVVTFLALLGIFIFTASILGMNLFGGKYSGVMVKSRANFDDLFWALVTVFQILTQEDWNVVMVDGMRATGKWAALYFILLMTIGNYILFNLLVAILVEGFANPDSTATIRQSAVVLSKTEYNQLSPSPSRKIGFFLNDRNEVFMHSLKRKRVDWSLYIFSPENSFRKWNVALYKNKWFDRIVLVFILLNCVVMAMENPSVKDDSTERQAIDICMYIFLGIFTLEMLIKIIAMGFWVGEYLRSGWNVMDGFLVVISWVDVIVTILGVLRVFRALRTLRPLRVISRAPGIKIVVETLISSLKPIGNIVLIAATFFMIFGILGVQLFKGKFYYCDSLEVDTRQQCEAKNKEYNFDNLAKALLTLFVFATKDGWVSIMHDGIDAVGIDKQPKENYARVNVLYFVAFLLLAGFVVLNMLVGVVVENFKKCREMERDEEEKKKKEKEREGDEASTHYSR---HRRFIYHTVTHAYFDLGIAVTIGLNVICMALQPKGLGDFLQSANYVFTAIFILEAILKIYALGVKRYFSDRWNHLDLVIVILSIAGIILEEDLPINPTIIRVMRVLRIARVLKLLKTAEGIRKLLDTVLQALPQVGNLGMLFLLLFFIFAALGIELFGGIDCKNEGMNEHAHFQRFDIAMLTLFRISTGDNWNGILKDTIPEVDCNEHVAPIYFAMFVLATQFVLLNVVVAVLMKHLEDAKD

>Exaiptasia_pallida_Cav2c_XP_020902316.1

SLLCLTERNPLRHYCKKIVESKAFEYFILVTIAVNCIVLMLDSPLPENDTTERNQKLETAETVFVVIYCVEAVTKIIALGFVFHPNAYLRNGWNILDFAVVVVGLIGFIWDSVEDTKVLRAARVLRPLKIVSGIPSLQVVMKTIWRAMVPLLQILVLILFVIVIYAIVGLELLKGRFHYTCYNLNQEAIDKTYSPKICGGGRPCP-PGPNKGITNFDNIFLSMLTVFQCITMEGWTDIMYHSYDARDLLTSTIYISLIMIGSFFMLNLVLGVLSGEFARRQQQMQRMFDDYLEWINKAEDIM----LKKRYSLSDSIMHLIEDLLRGHIRSLVRSSVFYWGVLLCVFLNTVIMLTEHYDQPYWLDKTQEISEKVFLGIFIVEMLLKLYGLEPRNYFNSAFNKFDFIVVLSGIAELFTKMSFGSSVLRSLRLLRVFKFTRVWSSLRNLVTSLLKSMRSILSLIFLLLLFIFIFALLGMQLFGGKFSKRLDAPRTNFDSFVKAMLAVFQIMTGEDWNTVMYDGIEAAGILASLYFVXLVIIGDYTLLNVFLAIAVDXLANAQALTRLEEEDERQREQNKKRVKISEDSPSVSSRNGRISRQDTTDNEIIRKSSMFIFGPDNPVRRFCHWLVNLRYFDTFILFIILISSVLLAFEDPV-NSDSQRNRILGYFDYVITAIFALEVVAKMIDLGVIVHKYLRDWWNVIDAFVVVCNITALILNLEDAIKSFRVLRVLRPLKAINKLKNLKTVFQCMIFSLKNVRNILIITGLFYFIFAVIGVQLFKGKFFYCTDPSKKFEKDCKGKNVDFNFDNVPYAMLTLFSSSTGEGWPIAMYHTIDATNVDEGPRRDNNIHMSIYYVCFVVVFSFFFLNIFVALIIVTFQEQGEKEMVGCELNRNQRDIQFAMTAKPRQRYMPCFFKVWRVVDSKPFEIFIMTMIVLNALVLMMASPQYDKYLEYINMAFTFVFLLEAILKLIAFKL-NYFRDYWNIFDFIIVVATLVEVLIELSDSKRDIDPSFFRLFRAARLVKLLRQGYTIRILLWTFFQSFKALPYVVILIALLFFVYAVIGMQLFGRIKQVPNKINIHNHFQSFLQSLLVLFRSATGENWHLVMLACFVGKDCGTMASIIYFCTFYFFCSFLMLNLFVAVIMDNFEYLTR

>Exaiptasia_pallida_Cav2b_XP_020906771.1

SLYCLSTTNPARRLCRFIVDSKAFEYFILLNIVANCVVLAMNKPLPKNDKIEMAVDLEKAELYFLAIFCIEAFLKIVAFGFVLHPGSYLRNLWNVLDFIVVLVGIFSLEQLPWNDVKALRAVRVLRPLKLISGVPSLQVVMKSIGRAMVPLLQIALLVLFVIVIYAIIGLDFLIGKFHYTCVKTTXGVKWLSKTNQPCDGGRKCN-VGPNHGITSFDNIALSMITVFQCITMEGWTEIMYLTFQAIDYLYSIYFATLIVIGSFFMLNLVLGVLSGEFARRQEQLEREVAGYIEWISKAEEVMSQAHVISNDLVVNPMAVLTKPQVRIKLRKAVKSQWFYWTILLCVFLNTVSLATEHNNQPPWLGEFQEWAERIFLIIFIVEMILKMYSLGMRIYFSSSFNIFDCVVVCSGIIDTILNLNLGISVLRCLRLLRIFKVTRHWTSLRNLATSLISSIKSIISLIFLLFLFILISALLGMQVFGGKF-SEKSTPRTNFDNFPNAMLTVFQILTGEDWNAVMYNGMMAYGGVVSLYFVFLVIIGNYTLLNVFLAIAVDNLANAQILTEDEENEKKEREIKRAKKRRVREQRGGRLGRGRGGRGKSQSDGIIKTRALFIFGPDNCFRRLCHRIVCLPHFDNFMLIVIMLSSLVIAIENPV-YDYAEINQYLWYFDCVFTAIFVFEVVVKVIDMGLIIHKYLRSVWNIIDFIVVICNLASLILSGSSAIRALRVMRVLRPFKSVHKIKKLQAVFRCMWFSVKNVANIGMITVLFLFIFAVMGVQLFNGKFSYCTDEAKLTQEECQGKTRVFNFDDVGHAMLTLYTSSTGEGWPTAMHYTMDTTEKGRGPVQDSNTPYAIYYVSFVVVFSFFFLNIFVALIILTFQDLGEKEILNCELDRNQRDVHFALTAKPVQLYMPFQYYVWMLVTSKPFEILIMVLISLNTIVLMMQHKDYKSVCSKLNIAFTALFVVEAALKLIAFRL-NYFRDYWNDFDFIVVLGGLADIILTFDREDIPIDPSMFRLFRAARLIKLLRQGYTIRILLWTFFRSFKALPYVTLLILLMFFMYAVIGMQLFGKIDIHQSALTEFNNFRHMPMAVQVLFRSATGENWHEIMRSCFLSEDCGTPLAVIYFCSFIFLCMFLMLNLFVAVIMDNFEYLTR

>Exaiptasia_pallida_Cav2a_XP_020910418.1

---------------------------------------------------------------MLIV-------------------KYF-------------------------SL------------------LGLQVVMKSIMCAMLPLLQICLLVGFVIVIYAIIGLEFLCGKFHYACFNNGSAPTIAGDDPELCDPGKGCE-DGPNDGITSFDNIFAGCLTVFQVITNEGWTDIMYWTFRVYDLAFWLYYYSLVIIGSFFMLNLVLGVLSGEFARRTKQMERMLHGYLDWISKAEDLMRKRNLDRVEDGDVAMTSQVLTRWRIRVRQVVKHQAFYWSVIICVFLNTVITACQHYQQPTWLTQFQEKAELIFISFFFLEMCLKLYGLGPQLYFKSQFNTFDCVVVWCGILELIEGIELGISVLRALRLLRLFKYTRYWSSLRNLVTSLLSSVRSILSLLFLLFLFIVIFALLGMQIFGSQFRGRDENPRTNFDDFWNAFLAVFQILTGEDWNAVMYDGVLSAGALYSLYFVALVVLGNYVLLNVFLAIAVDNLANAQQLSADEAEEEEAREERKRERNGNARRDDDDMGPNEDEFEEVEESGIINTWSLFLFPPGNPVRKFCHWLVNLRHFDNVILVIILISSGLLAAEDPV-VENSERNKILTYFDYVFTTIFAMEVVVKLIDYGAILHPYFRDAWNCIDALVVSCAIASLVLSSKKTVKVLRVLRVLRPLKAINKAKKLKAVFQCMVFSLRNVLNILIITVLFLFIFSVIGVQLFQGKFFTCSDKSKMTEQECRGANQEYNFDNVFKAMLALFASSTGEGWPALMQASIDTTGIDRGPIVDNKVEIFLFYIFFVIVFSFFFLNIFVALIILTFQEQGEKEQGDCELDRNQRDLHFAIVAKPSERFMPIQYRVWKLVDCRPFEFTIMTLIALNTLILMMEPKEYRYWLNLFNTIFTFMFTTEAILKLIAFRQ-NYFRDSWNVFDFVVVLGSLLDFILDKDGQKLPFDPSLFRLFRAARLVKLLRQGYTIRILLWTFLQSFKVRTLINISMSIHFFLF-----QLFGQIYRGDEQISVENNFQSFTQAIQVLFRSATGENWHVIMLACRTGETCGSDITYLYFISFIFFCSFLLLNLFVAVIMDNFEYLTR

>Exaiptasia_pallida_Cav1_XP_020903719.1

AVFCLTLGNPIRSAAISLVEWKPFDVMILITIFANCAALAAYQPLPEQDSSSVNEELEVAEYVFLAVFTLEALLKIIAYGFVMHPGAYLRNGWNILDFVIVVVGLATILVKALNDVKALRAFRVLRPLRLVSGVPSLQVVLNSIIKALIPLFHIALLVVFVVIIYAIIGVELFMGKLHKTCYD-NVTGQMAFDEAHPCSTGYACS-DGPNFGITNFDNIGLACLTVFQCITLEGWTDVMYSINDAVGSWPWLYFVTLIIWGSFFVLNLILGVLSGEFAREKQLIDDAYHGYLEWIAQAEDIERKLSTKRRKEDGELAENEENVRLRRFLRKAVKTQAFYWTVIVVVFLNSLTLALEHYNQPEFLTQFLDKANKLFLALFTLEMLVKMYCLGFHVYFASLFNRFDCLVVVSSLLELADQRPIGISVLRCVRLLRIFKVTRYWSSLSNLVASLLNSMRSIAGLLLLLSLFMLIFSLLGMQIFGGKFNDDTEVPRSNFDSFWRAIVTVFQILTGEDWNAIMYIGILSWGVIPILYFIFLVIVGNYILLNVFLAIAVDNLADAESLTEMENEKKKKEEEKKELEKKKKEEEKKELESSRSSLSESPTKGMPLESSMFIFSSTNCFRILCHKFVTNIYFVNFILILIIVSSALLAAEDPL-NANSKRNQILNYFDYFFTTAFTIEITVKIIAYGVFLHKFCRSAFNLLDALVVAVSIISIALRQISTVRILRVLRVLRPLRAINRAKGLKHVVQCVFVAVKTIGNIMIVTVLFNFLFAVIGVQLFKGTFFHCTDGGKITQEECQGKHRDFNFDNVLNAMLTLFTVMTFEGWPGILENSMDSTDVDQGPFLNNRPWVAVYYVIYIIIIAFFMVNIFVGFVIVTFQNEGEAEYEDCELDKNQGT------------FFH--------CTDG---------------------KITQEECQG-----------------------QYL-----------------EF----KGPGLSNPVTQQREWK------------------------------------------------------------HRDFNFDNVLNAMLTLFTVMTFEGWPGILENSMDVDQGPRPWVAVYYVIYIIIIAFFMVNIFVGFVIVTFQNEGE

>Echin_acapla_Cav3_XP_022091428.1

VFYCLSQTSKPRVWCIQLVCWPWFERISMAVIILNCITLGMYEPCEKECTSTRCVVLEGFDHFIFAFFAAEMVVKILALGV-RGKSGYFFETWNRLDCFIVVAGITDYTMQALENLTAIRTIRVLRPLRAINRIPSLRILVMLLLDTLPMLGNVLMLCFFVFFIFGIIGVQLWKGLLRNRCFLVTTFYTLPDMYQYVCSFEQECNDVNPFLGSISFDNILYAWVAIFQVITLESWVEIQYYLQDVHSQWVWIYFMLLIVIGAFFLINLCLVVIATQFSSEPGSCYDEILKYIGHLVRKVKRKPPSPCLSIHSAGCPVHQPCPVKVSDKIGEIVESKYFMRSILICILVNTLSMGIEFHNQPDELTEALEISNRIFTSLFALEMLLKLMAYGFVGYIRNGFNVFDGIIVIVSVVEIVQQGGGGLSVLRTFRLLRILKLVRFMPALRRQLLIMLKTMDNVATFFSLLSLFIFIFSILGMHLFGCNFCGVKVCDRKNFDNLLWALVTVFQILTQEDWNIVLYNGMHNNSAWAALYFIALMTFGNYVLFNLLVAILVEGFAAAASLEEDEGYSGEEHEKKEEGDSRRTSVASCNGDSRRTSYISCNGGSTRIEYSLYLLSPHNRFRRRLQSLIAHRWFDYVILLIIMINCITLAMERPDIRDDSVERTFLSISNYIFTGIFTFEMVVKVLAKGLFIGEYLYSGWNIMDGSLVVISWIDITISIFGILRVFRLLRTLRPLRVISRAPGLKLVVQTLLSSLRPIGNIVIICCTFFVIFGILGIQLFKGTFYYCTSVKVMNKTHCLQVNQQYNFDNLGQALMALFVLASKDGWVEIMYNGIDAVGVDKQPKENNNEWLILFFISFILIVGFFVLNMFVGVVVENFHKCREQQAAEETARRQAKRRKMEKARQPYYIYSRSRRFIHNCVINKYFDLGVAAVIFINVISMALMSKTLQDILRYLNYFFTVVFILEAVLKIAALGFKRYIKDRWNQLDMIIIILSIVGIFLEEIIPINPTIIRIMRVLRIARVLKIMKTMKGLRELLAVLMGAIPQVGNLGLLFFLLFFIFAALGVELFGRLDCTKQGLGRHASFKNFFIAFLTLFRIATADNWNGIMKDTLTYNCCASILAPIYFVCFVLMAQFVLVNVVVAVLMKHLEESHK

>Echin_acapla_Cav2_XP_022085274.1

SLFIFSEQNFIRRCAKWLTEWPPFEYLVLATIIANCVVLALEVHLPMQDKTPMSQELENTEIYFLAIFCLEACIKITALGLVLHEGSYLRNGWNLMDFVVVVTGFVTFIGGLVEDLRTLRAIRVLRPLKLVSGIPSLQVVLKAILKAMAPLLQIGLLILFVIIIFAITGMEFFQGKFHYTCFETGKISETEGEDPQVCGKGRECP-EGPNFGITNFDNMLFAMLTVFQCITMEGWTDIMYNCNDSEGYFVWLYFIPLIILGSFFMLNLILGVLSGEFARRQQQLDKELNGYLEWICKAEEVMRAISGTIEDNFDLETLDKAKLRLRFQVRHMVKTQAFYWLVIVLVFLNTVCVAIEHYQQPEWLTQFLNHAEYVFLGIFITEMAIKMYGLSPAVYFKSAFNKFDCMVILASLFEVIQGGSFGLSVLRALRLLRIFKVTRYWSPMRYLIISLVHSIRSIVSLVFLLFLFIIIFALLGMQLFGGSFNSESPKPANNFDIFPIALMTVFQILTGEDWNMVMYYGVVSKGMVYSLYFVILVLFGNYTLLNVFLAIAVDNLANAQEMTKLDQEDEEEAQAARENLNVHGIPWPCPSWGPVDCPGMDAICNMVPFSSLFIFSTTNPVRRFCHYIVNLRYFDFLIMVAIGLSSLTLAMEDPV-NTGSQYNQVLEYFDYAFTTIFTIEMILKIIDMGLLFHKYCRDFWNILDSTVVICALVAFGVTNLNTIKALRVFRVLRPLKTIKRVPKLKAVFDCVVNSVKNVTNIAVVYGLFMFIFSVIGVQLYKGRFYHCTDPSKHTRDECMGQLYPFNYDNVGSALLTLFTVSTGEGWPDVLKHSIDATEEGRGPEPYNNLQMALFYVVYFIIFPFFFLNIFVALIIITFQEQGDQDVQDGEIDKNQKQMEFCIHAKP-TDFVPVKYKIWKLVVSQPFEYFIMSLIALNTIALMMAEQTYLDTLKYLNIAFTVLFTIEAILKLIGFGPRNYFRVSWNTFDFITVIGSIADAIISEVGVDDFINLSVLRLFRAARLIKLLRQGSSIRILLWTFVQSFKLVSFVFFLTFMLFFIYAIIGMQIFGTVNIDDTAITRHNNFSNFFLAIIMLFRCATGESWQSIMLACQEKYGCGSSLSVAYFVSFIFFSSFLMLNLFVAVIMDNFDYLTR

>Echin_acapla_Cav1_XP_022081221.1

ALFCLTLDNPVRRMCISIVEWKPFEYLILLTIFANCFALAIYTPFPHEDTNTTNKNLENVEYIFLFIFTLEAMLKIVAMGFLFHSGAYLRNAWNFLDFIIVIIGVVSTILSHTADVKALRAFRVLRPLRLVSGVPSLQVVLNSIVRAMVPLLHIALLVIFVILIYAVIGLELFIGKLHRTCWIEDGVRRHVEEEPHPCGDGFNCS-EGPNEGITNFDNIGQAMLTVFQCITMEGWTDVLYNVNYAIEWWPWFYFVTLILLGSFFVLNLILGVLSGEFSREKKQIEEDLKGYLDWIMQAEDIDKVPKPMSESDSSEKSEELGSTRCRRLCRQAVKSQAFYWVVIIMVFLNTIILASEHYRQPKWLMDFQDIGNLLFVVIFTVEMLIKMYSLGLQGYFVSLFNRFDCFVVCSSMVEVVVIQPIGISVLRCVRLLRVFKVTRYWASLRNLVASLLNSMRSIASLLLLLFLFILIFALLGMQVFGGRFNKTQDKPRSNFDNFWQSLFTVFQILTGEDWNEVMYDGIAAYGIIASSYFIILYIWGNYILLNVFLAIAVDNLADAESLTALEKEREEEKRNKSIRIEDGEKQVHIEEDEAKKALKEPGDEDMPKESSLFVLGPENRFRKGCYFVCTHNYFSNVVLLLILISSIMLAAEDPI-DKNKTLNFILNCFDYGFTVAFTIEILLKVISFGLVIHKFCRNFFNLLDLLVVTVSYISIALQAISAVKILRVLRVLRPLRAINRAKGLKHVVQCVFVAIKTIGNIMMVMLLLVFMFACIGVQLFSGKFYSCTDLSKMTEEHCHGKLNEFSFNNVGSGMLALFTICTFEGWPQLLYVAIDASEPDHGPIRNNQLGVAIFFFIYIIVVAFFMVNIFVGFVIVTFQNEGEQEFKNCELDKNQRQLEFALKARP-KKYIPKQLKVWKIVTSRPFEYLIFVLIMVNTIVLAMQSDEYSEVLDRVNIVFTAIFLLECILKIIAFKIKNYVRDLWNLFDFVIVVGSIIDIILSEMDSESRFSINFFRLFRVMRLVKLLSKGEGIRTLLWTFIKSFQALPYVALLIVMLFFVYAVIGMQLFGKIALTTGPINRNNNFQSFVAALLVLFRSATGEAWQQIMLACAVGETCGNDFAYIYFLTFYSFCSFLVINLFVAVIMDNFDYLTR

>Diploscapter_pachys_Cav3_PAV61762.1

VLHCFYQSRPPRSWALKLVMSPWFDRITMAVILINCITLGMYRPCDDGPSTYRCHILDIVDNCIFVYFTLEMMIKVIALGL-VGPTGYLADTWNRLDFFIVIAGIVEVVLADVMNLTAIRTVRVLRPLRAVNRIPSMRILVNLLLDTLPMLGNVLLLCFFVFFIFGIVGVQLWAGLLRSRCIILTRYYIPEDTSLYICSQGVKCNERNPFQGSVSFDNIGFAWISIFLVISLEGWTDIMYYVQDAHSFWNWIYFVLLIVIGAFFMINLCLVVIATQFAVEGGSVYAAIVRMISYTFRRAKRQMATLSRIEERMEEEEDDEKPNWFRFHVKRFVDSDHFSRSILVAILVNTLSMGVEHHQQPEIFTTILEYSNLFFTGLFAFEMLLKVIAYGLFGYLADGFNLFDGGIVALSVLELFQDGKGGLSVLRTFRLLRILKLVRFMPALRVF----------------------------GMILFGRKFCGECICDRMNYDTFMHATLT---ILTQEDWNMVLFNGMSQTNPWAALYFVALMTFGNYVLFNLLVAILVEGFQESKEEEKRQQEEEARKHALEEEGRKPRERTHSWSGVGHIFNENCPIHNRRSENSLFIFSPKNPLRVKFITLTQKKWFDYTILLFIGINCVTLAMERPSIPPYSIERKFLDISGYIFTIIFTVEMLMKVIANGCVFGDYFKDGWNVLDGILVIISLINVGFEIFGVIRVLRLLRALRPLRVINRAPGVKLVVMTLISSLKPIGNIVLICCTFFVIFGILGVQLFKGMMYHCTDIAVTTKEECLQVNHRYNFDNLGQALMSLFVLSSKDGWVSIMYQGIDAVGVDIQPIENYNEWRMIYFISFLLLVGFFVLNMFVGVVVENFHKCKEKEMREKAKEKQLQRQELARRERPYYAYGRTRLFLHGIVTSKYFDLAIAAVIGINVISMAMMPVVLKYVLKALNYFFTAVFTLEAIMKLVALGFRRFLKEKWNQLDMFIVILSIAGIIFEEELPINPTIIRVMRVLRIARVLKLLKMAKGIRSLLDTVGEALPQVGNLGSLFFLLFFIFAALGVELFGRLECSEDGLGEHAHFKNFGMAFLTLFRIATGDNWNGVLKDALESNCCVPILAPCFFVVFVLISQFVLVNVVVAVLMKHLEESNK

>Diploscapter_pachys_Cav1_PAV74346.1

-------RKPLRQA--NVVE-RPFEFLILLMICLNCIALAVTQPYPAQDSDSINAKLERVENVFIVVFTIECILKVIAYGFMFHPSAYLRNAWNVLDFIIVVIGIISTVLAGMHDVKALRAFRVLRPLRLVSGVPSLQVVLNAILRAMIPLLHIALLVLFVILIYAIIGLELFCGKLKQTCID-PATGQLAMKDPTPCGNSFQCI-TGPNFGITNFDNFGLAMLTVFQCVSLEGWTEVMYWVNDAVGEWPWIYFVSLVILGSFFVLNLVLGVLSGEFSREKQQLEEDLKGYLDWITQAEDIEPQTQQQQGEEQDEEGEERPDERCRRSCRRLVKSQTFYWLVILLVFLNTLVLTSEHYGQSDWLDEFQNWANLFFVILFTMEMFLKMYSLGLTTYTTSQFNRFDCFVVISSILEFILMKPLGVSVLRSARLLRIFKVTKYWTSLRNLVSSLLNSLRSIMSLLLLLFLFIVIFALLGMQVFGGKFNPQAPKPRANFDTFIQALLT---ILTGEDWNTVMYNGINSFGVVVSIYYIVLFICGNYILLNVFLAIAVDNLADADSLTNAEKEEEQQ------------------ENEEGAELDEHEGDDIPKASSLFILSHTNPFRVFCNKVVNHSYFTNSVLVCILVSSAMLAAEDPL-KADSPRNLILNKFDYFFTTVFTIEITLKVVFFGLILHKFCRNAFNLLDIIVVAVSIISPILKAFSVVKILRVLRVLRPLRAINRAKGLKHVVQCVIVAVKTIGNIMLVTFMLQFMFAIIGVQLFKGTFFYCNDISKMTEAECRGQNNDFNFDHVGNAMISLFVVSTFEGWPDLLYIAIDSNEENKGPIHNNRQAVALFFIAFIIVIAFFMMNIFVGFVIVTFQNEGEREYENCELDKNQRKIEFALKAKPHRRYIPFQYRVWWFVTSRAFEYLIFLIIVLNTVALACSSQSFNDILDKLNLCFTSVFAFEAFFKIIALNPKNYFGDRWNAFDFIIVLGSFIDITWSNPDNKSLISINFFRLFRVMRLVKLLSRGEGIRTLLWTFMKSFQALPYVALLIVLLFFIYAVIGMQVFGKIALDDTEIHRNNNFQTFFSAVLVLFRSATGEAWQQIMLSCSPGQKCGTNFAYPYFISFFMLCSFLVINLFVAVIMDNFDYLTR

>Daphnia_pulex_Cav1_EFX89598.1

ALLCLSLTNPLRKLCISVVEWKPFEYLILLTIFANCVALAVYTPYPNGDSNITNAYLEKVEYVFLVIFTIECVMKIIAYGFVAHSGAYLRNTWNLLDFTIVVIGAVSTALSTMMDVKALRAFRVLRPLRLVSGVPSLQVVLNSILKAMVPLLHIALLVIFVIIIYAIIGLELFSGKLHTTCYD--PETGDMMKDPHPCSNGFDCR-EGPNDGITNFDNFGLAMLTVFQCVTLEGWTDVLYQIEDAMGSWQWIYFISMVIIGAFFVMNLILGVLSGEFSREKQQIEEDLRGYLDWITQAEDIEQDNSVDKTEESSNA-------RMRRACRKAVKSQAFYWLIIILVFLNTGVLATEHYQQPEWLDFFQDVTNIFFIVLFAFEMLLKMYSLGFQGYFVSLFNRFDCFVVISSIVEVVLMPPLGVSVLRCVRLLRVFKVTKYWRSLSNLVASLLNSIQSIASLLLLLFLFMVIFALLGMQVFGGKFNENQELPRSNFDSFWQSLLTVFQILTGEDWNAVMYDGIKAYGILACIYFIILFICGNYILLNVFLAIAVDNLADAESLTAIEKEEADQAEAKAEAEDNGLGDDQGSYDDDNNGEEEEAEVGIPPYTSFFILSHSNRFRVFCHWFCNHNYFGNIILACILISSAMLAAEDPL-SADTDRNKILNHFDYFFTTVFTIEIALKVIAYGLVLHKFCRSAFNLLDLLVVCVSLISFGFSAISVVKILRVLRVLRPLRAINRAKGLKHVVQCVIVAVKTIGNIMLVTCLLEFMFAVIGVQLFKGKFFKCNDPSKMTRDDCIGENNDFHFDDVGKAMLTLFTVSTFEGWPGLLYVSIDSHTEDMGPMHNYRPMIACFYFIYIIIIAFFMVNIFVGFVIVTFQNEGEQEYRNCELDKNQRNIEFALKAKPFRRYIPFQYKVWWMVTSQPFEYVLFTLISMNTITLGMQPEVYTQALDILNMIFTSVFALEFLLKLAAFRFKNYFSDPWNVFDFVIVLGSFIDIVYSQVPDSNLISINFFRLFRVMRLVKLLSRGEGIRTLLWTFMKSFQALPYVALLIVMLFFIYAVIGMQVFGKIALDDTAIHRNNNFQTFPQAVLVLFRSATGEAWQDVMLGCSSTDHCGTEFAIPYFISFYVLCSFLIINLFVAVIMDNFDYLTR

>Daphnia_magna_Cav1_KZS09199.1

ALFCLSLTNPVRKLCISVVDYTPFEYLILLTIFANCVALAVYTPYPNGDSNITNAYLEKVEYVFLVIFTIECVMKIIAYGFVAHSGAYLRNTWNFLDFTIVVIGAVSTALSTMMDVKALRAFRVLRPLRLVSGVPSLQVVLNSILKAMVPLLHIALLVIFVIIIYAIIGLELFSGKLHTTCYDTG----DMMKDPHPCSNGFDCAE-GPNDGITNFDNFGLAMLTVFQCVTLEGWTDVLYQIEDAMGSWQWIYFISMVIIGAFFVMNLILGVLSGEFSREKQQIEEDLRGYLDWITQAEDIEQDNSVDKTEESSNA-------RMRRACRKAVKSQAFYWLIIILVFLNTGVLATEHYQQPEWLDFFQDVTNIFFIVLFAFEMLLKMYSLGFQGYFVSLFNRFDCFVVISSIVEVVLMPPLGVSVLRCVRLLRVFKVTKYWRSLSNLVASLLNSIQSIASLLLLLFLFMVIFALLGMQVFGGKFNENQELPRSNFDSFWQSLLTVFQILTGEDWNVVMYDGIKAYGILACIYFIILFICGNYILLNVFLAIAVDNLADAESLTAIEKEEADQAGAKAEADDEGLGDDQGSYDDDNNGEEEEAEVGIPPYTSFFILSHSNRFRVFCHWFCNHNYFGNIILACILISSAMLAAEDPL-SADTDRNKILNHFDYFFTTVFTIEIALKVIAYGLVFHKFCRSAFNLLDLLVVCVSLISFGFSAISVVKILRVLRVLRPLRAINRAKGLKHVVQCVIVAVKTIGNIMLVTCLLEFMFAVIGVQLFKGKFFKCNDPSKMTRDDCIGENNDFHFDDVGKAMLTLFTVSTFEGWPGLLYVSIDSHTEDMGPMHNYRPMIACFYFIYIIIIAFFMVNIFVGFVIVTFQNEGEQEYRNCELDKNQRNIEFALKAKPFRRYIPFQYKVWWMVTSQPFEYVIFTLIITNTITLGMQPEVYTQALDVLNMIFTSVFALEFVLKLAAFRFKNYFSDPWNVFDFVIVLGSFIDIVYSQNPDSNLISINFFRLFRVMRLVKLLSRGEGIRTLLWTFMKSFQALPYVALLIVMLFFIYAVIGMQVFGKIALDNTAIHRNNNFQTFPQAVLVLFRSATGEAWQDVMLGCSATEHCGTEFAIPYFISFYVLCSFLIINLFVAVIMDNFDYLTR

>Danio_rerio_Cav3.3_XP_021329632.1

-------MNVGLNSCFL----TWFERISIMVILLNCVTLGMYQPCENIDTSERCQVLQAFDAFIYIFFALEMVVKMVALGI-FGRRCYLGDTWNRLDFFIVMAGMVEYSLDLQNNFSAIRTVRVLRPLKAINRVPSMRILVNLLLDTLPMLGNVLLLCFFVFFIFGIIGVQLWAGLLRNRCYPTPYYQPEEDDERFICSLGRECCHTNPHKGAINFDNIGYAWIVIFQVITLEGWVEIMYYVMDAHSFYNFIYFIFLIIIGSFFMINLCLVVIATQFSAEPGDCYEELFQLVCHILRKARRRQTSHCIHETKLDHSSENAVSLEIRVKLWGIVESKYFNRGIMIAILINTISMGIEHHNQPDELTNVLEICNIVFTSMFTLEMILKLTAFGFFEYLRNPYNIFDGIIVIISVCEIIGQSDGGLSVLRTFRLLRVIKLVRFMPALRRQLVVLMKTMDNVATFCMLLMLFIFIFSILGMHIFGCKFSGDTVPDRKNFDSLLWAIVTVFQILTQEDWNMVLYNGMASTSPLAALYFVALMTFGNYVLFNLLVAILVEGFQAGDSYSDDDRSSCNLEETEKKDPPLHPRQRKKSFSGGLGSVGEHQDCNTREDWSVFLFSPQNKFRLLCQSIIAHKLFDYVVLAFIFSNCITVALERPKILQGSLERLFLTVSNYIFTAIFVGEMTLKVVSMGLYIGEYLRSSWNILDGFLVFVSLIDIVVSILGVLRVLRLLRTLRPLRVISRAPGLKLVVETLITSLKPIGNIVLICCAFFIIFGILGVQLFKGKFYYCLDVKITNKSDCLLVHHKYNFDNLGQALMSLFVLASKDGWVNIMYHGLDAVAVDQQPITNNNPWMLLYFISFLLIVSFFVLNMFVGVVVENFHKCRQHQEVEEAKRREEKRRRMEKKRRPYYAYSHVRLMIHTLCTSHYLDIFITFIICVNVVTMSLQPHSLEIALKYCNYFFTSTFVLEAVLKLIAFGFRRFFKDRWNQLDLAIVLLSVMGITLEEALPINPTIIRIMRVLRIARVLKLLKMATGMRALLDTVVQALPQVGNLGLLFMLLFFIYAALGVELFGELVCNEEGMSRHATFENFGMAFLTLFQVSTGDNWNGIMKDTLGEYTCNQFISPLYFVSFVLTAQFVLINVVVAVLMKHLDDSNK

>Danio_rerio_Cav3.2_XP_009297960.1

----------------------------MLVILLNCVTLGMYQPCEDLKQSEWCIVLQAFDDCIFAFFAVEMVIKMIALGI-FGINGYLGDTWNRLDFFIVMAGMMEYSL--DGSLSAIRTVRVLRPLRAINRVPSMRILVTLLLDTLPMLGNVLLLCFFVFFIFGIVGVQLWAGLLRNRCFMSLYYENEDGEDNFICSSGVECTDLNPHKGAVNFDNIGYAWIAIFQVITLEGWVDIMYYVMDAHSFYNFIYFILLIIVGSFFMINLCLVVIATQFASEPGSCYEEMLKYVSHLYRKVKRRQQIEAHSHSRVLLQMAGVVPTDFRERLTRIVDSKYFNRGIMIAILINTLSMGIEYHEQPEELTNILEISNIVFTSMFVLEMLFKLLAFGIFGYIRNPYNIFDGVIVVISVWEIIGHADGGLSVLRTFRLLRVLKLVRFLPALRRQLLVLMKTMDNVATFCMLLMLFIFTFSILGMHLFGCKFSGDTIPDRKNFDSLLWAIVTVFQILTQEDWNVVLYNGMASTSPWAALYFVALMTFGNYVLFNLLVAILVEGFQAGDSDGDEEKTSVNSEEKMEQLSGGSLQHPSSLDIPELPQMGEYLDCNDHEEWSLYLFSPHNKFRMMCQKLISHKMFDYVVLVFIFLNCITIALERPH--IQQSERLFLLVSNYVFTVIFVAEMTVKVVALGFYSGNYLKSTWNVLDGVLVFVSLIDILVSIFGILRVLRLLRTLRPLRVISRAPGLKLVVETLITSLRPIGNIVLICCAFFIVFGILGVQLFKGKFFHCEDTRITNKSDCLQIRRKYNFDNLGQALMSLFVLSCKDGWVNIMYDGLDAVGVDQQPERNHNPWMLLYFISFLLIVSFFVLNMFVGVVVENFHKCRDQEEVEARLRRSKENGSEALRR-PYYAYSPARLYIHTLCTNHYLDLFITGIICINVVTMSIQPSYLDEALKYCNYVFTIIFIIEALLKLVAFGIRRFFKDRWNQLDLAIVLLSIMGITLEEALPINPTIIRIMRVLRIARVLKLLKMATGMRSLLDTVMQALPQVGNLGLLFMLLFFIYAALGVELFGKLECTEEGLSPHATFENFGMAFLTLFRVSTGDNWNGIMKDTLSETQCLPWVSPIYFVTFVLMAQFVLVNVVVAVLMKHLEESNK

>Danio_rerio_Cav3.1_XP_021336020.1

VFFYLKQTTRPRSWCLKMVCNPWFERASMLVILLNCVTLGMFHPCEDSHDSERCKILEDFDDFIFAFFAVEMVIKMVALGI-FGKKCYLGDTWNRLDFFIVLAGMLEYSL----SFSAVRTVRVLRPLRAINRVPSMRILVTLLLDTLPMLGNVLLLCFFVFFIFGIVGVQLWAGLLRNRCFLRKYYHTENDDENFICSQGRQCQEANPFKGAINFDNIGYAWIAIFQVITLEGWVDIMYFVMDAHSFYNFIYFILLIIVGSFFMINLCLVVIATQFSSEPGSCYDELLKYLVHVIRKGTRQPPGKVISKDALMNCSSTNIDTLVCGTFRKIVDSKYFGQGIMIAILINTLSMGIEYHEQPDELTNALEISNIVFTSLFVLEMLLKLLVYGPFGYIKNPYNIFDGIIVVISVWEIVGQQGGGLSVLRTFRLMRVLKLVRFMPALQRQLVVLMKTMDNVATFCMLLMLFIFIFSILGMHLFGCKFGGDTLPDRKNFDSLLWAIVTVFQILTQEDWNKVLYNGMASTSPVAALYFIALMTFGNYVLFNLLVAILVEGFQTGDSDSEADISLEDVSGHKKDMFDLHPDTLQVPYPHRSGSIHSARPPNQRVTWSLYLFPPESRFRVTCNKIITHKMFDHVVLVIIFLNCITIAMERPRIDPSSAERIFLTLSNYIFTAIFVTEMTIKVVALGFCFGEYLKSSWNILDGMLVMISVIDILVSILGMLRVLRLLRTLRPLRVISRAPGLKLVVETLMSSLKPIGNIVVICCAFFIIFGILGVQLFKGKFFVCHDTRITNKSDCLLVRHKYNFDNLGQALMSLFVLASKDGWVDIMYDGLDAVGVDQQPVMNYNPWMLLYFISFLLIVAFFVLNMFVGVVVENFHKCRRNQEEEEAKRREAKRKRRDKKRRPFYSYCPTRRLIYNMCKSQYLDLFITIVIALNVITMSMQPKVLSDSLKICNYIFTVIFVLESIFKLVAFGFRRFFKDRWNQLDLAIVLLSIMGITLEESLPINPTIIRIMRVLRITRVLKLLKMAVGMRALLDTVMQALPQVGNLGLLFMLLFFIFAALGVELFGDLICDEEGLGRYATFKNFGMAFLLLFRVSTGDNWNGIMKDTLDTGICYTVVSPIYFVSFVLTAQFVLVNVVIAVLMKHLEESNK

>Danio_rerio_Cav2.3_XP_017206695.2

SLFIFAENNMIRKYAKRIIEWPPFEYMILATIIANCIVLSLEQHLPGEDKTPMSKRLEKTEPYFIGIFCFEAGIKLVALGFVFHKGSYLRNGWNVMDFIVVLSGILATAGSHMNDLRTLRAVRVLRPLKLVSGIPSLQIVLKSIIKAMVPLLQIGLLLFFAILMFAIIGLEFYSGKLHHTCLAILDNETVDSSEVFACG-VRKCP-IGPNDGITQFDNILFAVLTVFQCITMEGWTAVLYNTNDALGTWNWIYFIPLIIIGSFFVLNLVLGVLSGEFARRQQQVERELNGYRAWIDRAEEVMSKKNARRRGPGEEKYSEISTVMLRFSIRRMVKTDSFYWIVLSLVALNTISVSIVHHNQPEWLTVIQYYTEFVFLGLFLAEMFLKMYGLGFRLYFHSSFNCFDCGVIVGSIFEVVPGVSFGISVLRALRLLRIFKITKYWSSLRNLVVSLMSSMKSIISLLFLLFLFIVVFALLGMQLFGGRFIFEDYTP-TNFDTFPASIMTVFQILTGEDWNEVMYNGIRSQGMWSSIYFIVLTLFGNYTLLNVFLAIAVDNLANAQELTKEEEEEEEESERRRRPTEEAQSKHMMNMCDPERQIENQEECQPERPDSMFIFKSKNPIRRICHYVVTLRYFEMTILLVIVASSIALAAEDPV-CTSSERNKVLRYFDYVFTGVFTFEMIIKMIDQGLILHDYFRDMWNILDFIVVVGALIAFALTDIKTIKSLRVLRVLRPLKTIKRLPKLKAVFDCVVTSLKNVFNILIVYKLFMFIFAVIAVQLFKGKFFYCTDGSMGTQKECQGRRHEFHYDNVLWALLTLFTVSTGEGWPQVLQHSVDVTEEDHGPSRGNRMEMSIFYVIYFVVFPFFFVNIFVALIIITFQEQGDKMMEECNLEKNERAIDFAISAKPLTRYMPLQYRLWHFVVSPSFEYTVLVMIALNTVVLMMAPTAYDIVLKHLNTAFTVLFSLECILKIMAFGFMNYFRDTWNIFDFITVLGSITEIVVD-LQSVNTINMSFLKLFRAARLIKLLRQGYTIRILLWTFVQSFKALPYVCLLIAMLFFIYAIIGMQVFGNIKLNDSHINQHNNFKTFFGALMLLFRSATGESWQEIMLSCLHEGGCGTDFAYFYFVSFIFFSSFLMLNLFVAVIMDNFEYLTR

>Danio_rerio_Cav2.2_XP_021331856.1

SLFIFSEDNIIRKYAKKITEWPPFEYMILATIIANCIVLGLEQHLPALDKTPMSKRLDDTEPYFIGIFCFEAGIKIIALGFAFHKGSYLRNGWNVMDFVVVLTGILTIVG----DLRTLRAVRVLRPLKLVSGIPSLQVVLKSIMKAMVPLLQIGLLLFFAILMFAIIGLDFYMGKFHRTCFR---TDTGEQVDEFPCGLAWTCE-IGPNFGITNFDNILFAVLTVFQCITMEGWVDILYNANDASGTWNWLYFIPLIIIGSFFMLNLVLGVLSGEFARRQQQIERELTGYLEWICKAEEVMSKNDLINAEEGEDHFTDISSVRFRFFIRRMVKAQSFYWTVLCIVGLNTLCVAIVHYDQPEWLTYALYLAEFVFLGLFLIEMSLKMYGLGPRTYFHSSFNCFDFGVIVGSIFEVIPGASFGISVLRALRLLRIFKVTKYWNSLRNLVVSLLNSMKSIISLLFLLFLFIVVFALLGMQLFGGQFNFEDETPTTNFDTFPAAILTVFQILTGEDWNAVMYHGIESQGMFCSVYFIVLTLFGNYTLLNVFLAIAVDNLANAQELTKDEEEQEEAISKKLALSGTREGERCSRGENGGNGGRRRHRPRILPFSSMFIFNPTNPVRRLCHYIVSLRYFEMCILVVIAMSSIALAAEDPV-QANAPRNNVLKYLDYVFTGVFTFEMVIKMVDLGLILHPYFRDLWNILDFIVVSGALVAFAFSDISTIKSLRVLRVLRPLKTIKRLPKLKAVFDCVVNSLKNVLNILIVYILFMFIFAVIAVQLFKGKFFYCTDESKGLEKDCRGKKYEFHYDNVLWAFLTLFTVSTGEGWPTVLKHSVDATFEDQGPSPGYRIEMSIFYVVYFVVFPFFFVNIFVALIIITFQEQGDKVLSECSLEKNERAIDFAINAKPLTRYMPFQYRLWKFVVSPPFEYSIMIMIALNTVVLMMAPDFYEAMLKYLNIVFTVLFSLECILKIIAFGPLNYLKDAWNVFDFVTVLGSITDILVTENTTERQLNFSFLRLFRAARLIKLLRQGYTIRILLWTFVQSFKALPYVCLLIAMLFFIYAIIGMQVFGNIDLNDTAINRHNNFRTFLQALMLLFRSATGEAWHDIMLSCLLGKECGSDFAYFYFVSFIFLCSFLMLNLFVAVIMDNFEYLTR

>Danio_rerio_Cav2.1_XP_021330116.1

SLFIFSEDNFVRKYAKKITEWPPFEYMILATIIANCIVLALEQHLPDGDKTPMSERLEDTEPYFIGIFCFESGIKILALGFAFHKGSYLRNGWNVMDFVVVLTGILSTVG----DLRTLRAVRVLRPLKLVSGIPSLQVVLKSIMKAMIPLLQIGLLLFFAILMFAIIGLEFYMGKFHTTCFD---KITDEIREEFPCGEARICP-LGPNYGITQFDNILFAVLTVFQCITMEGWTDLLYYSNDAAGAWNWMYFIPLIIIGSFFMLNLVLGVLSGEFAKRQQQIERELNGYLEWICKAEEVISKTDLLDAEDGDGDIGSPF--RLRFFIRRIVKTQAFYWTVLCLVGLNTLCVAVVHYDQPETLSDFLYFAEFIFLGIFMSEMCIKMYGLGTRPYFHSSFNCFDCIVICGSIFEVLPGTSFGISVLRALRLLRIFKVTKYWASLRNLVVSLLNSMKSIISLLFLLFLFIVVFALLGMQLFGGQFNFEAGTPPTNFDTFPAAIMTVFQILTGEDWNMVMYDGIESQGMVFSVFFIVLTLFGNYTLLNVFLAIAVDNLANAQELTKDEQEEEQAANKKMALLSRQDSQYSEDLDNAMNNKLATQPHDMPPYTSMFILTTTNPFRRLCHYIVTLRYFEMCILLVIAMSSIALAAEDPV-WPESPRNNVLRYFDYVFTGVFTFEMLIKMVDLGLVLHQYFRDLWNILDFIVVSGALVAFAFTDISTIKSLRVLRVLRPLKTIKRLPKLKAVFDCVVNSLKNVLNILIVYMLFMFIFAVVAVQLFKGRFFYCTDESKEFERDCRGKKYDFHYDNVLWALLTLFTVSTGEGWPQVLKHSVDATYENQGPSPGYRMEMSIFYVVYFVVFPFFFVNIFVALIIITFQEQGDKMMEDYSLEKNERAIDFAINAKPLTRHMPFQYRMWEFVVSPPFEYTIMALIALNTIVLMMASLTYEDVLKYLNIVFTSLFSMECILKIIAFGALNYFKDAWNIFDCVTVLGSITDILVT-ELGNNFINLSFLRLFRAARLIKLLRQGETIRILLWTFVQSFKALPYVCLLIAMLFFIYAIIGMQLFGNIKINSSAITQHNNFRTFFQALMLLFRSATGEAWHDIMLSCLPKPECGSEFAYLYFVSFIFLCSFLMLNLFVAVIMDNFEYLTR

>Danio_rerio_Cav1.3_XP_021335256.1

ALFCLNLNNPIRRACISLVEWKPFDIFILIAIFANCMALAVYVPFPEDDSNSTNHDLETVEYAFLIIFTIETFLKIIAYGLVMHQNAYVRNGWNMLDFVIVVIGLFSVVLEVLTDVKALRAFRVLRPLRLVSGVPSLQVVLNSIIKAMVPLLHIALLVLFVIIIYAIIGLELFIGKMHASCYF--QGTDILEDEPAPCAVGRTCP-QGPNGGITNFDNFMFAMLTVFQCITMEGWTDVLYWMNDAMGELPWVYFVSLVIFGSFFVLNLVLGVLSGEFSREKQQLEEDLKGYLDWITQAEDIDSETESMNTENEKGEDEKATCCLCRRNCRLAVKSVPFYWLVIILVFLNTLTISSEHYNQPMWLTQVQDVANKVLLAMFTCEMLVKMYSLGLQAYFVSLFNRFDCFVVCGGITETIIMSPLGISVFRCVRLLRIFKVTRHWASLSNLVASLLNSMKSIASLLLLLFLFIIIFSLLGMQVFGGKFNDETQTKRSTFDNFPQALLTVFQILTGEDWNAVMYDGIMAYGMIVCIYFIILFICGNYILLNVFLAIAVDNLADAESLNTDDTKKPDE--------------IDEIEDEAKAGEEDEKDNAIPEGSAFFIFSNTNPVRVACHKLINHHIFTNLILVFIMLSSASLAAEDPI-RNFSARNIILGYFDYAFTAIFTVEIVLKMTTYGAFLHKFCRNYFNLLDLLVVGVSLVSFGIQAISVVKILRVLRVLRPLRAINRAKGLKHVVQCVFVAIRTIGNIMIVTTLLQFMFACIGVQLFKGKFYRCNDEAKSSPEECKGHNSDFNFDNVLMAMMALFTVSTFEGWPALLYKAIDSNRENMGPIYNYRVEISIFFIIYIIIIAFFMMNIFVGFVIVTFQEQGEKEYKNCELDKNQRQVEYALKARPLRRYIPYQYKFWYVVNSTGFEYIMFVLILLNTICLAVQSELFNYVMDILNMVFTAVFTVEMVLKLIAFKPRHYFTDAWNTFDALIVVGSVVDIAITETEDSARISITFFRLFRVMRLVKLLSRGEGIRTLLWTFIKSFQALPYVALLIAMLFFIYAVIGMQVFGKIAMVDTQINRNNNFQTFPQAVLLLFRCATGEAWQEIMLACMEEMTCGSSFAIIYFITFYMLCAFLIINLFVAVIMDNFDYLTR

>Danio_rerio_Cav1.2_XP_009298610.1

ALLCLTLKNPIRRACINIVEWKPFEIIILMTIFANCVALAVYIPFPEDDSNATNSNLERVEYLFLIIFTVEAFLKVIAYGLLCHPNAYLRNGWNLLDFIIVVVGLFSAILEQATDVKALRAFRVLRPLRLVSGVPSLQVVLNSIIKAMVPLLHIALLVLFVIIIYAIIGLELFMGKMHRTCFFDGHKGHIAEEKPAPCAPGRHCS-EGPNDGITNFDNFAFAMLTVFQCITMEGWTDVLYWMQDAMGELPWVYFVSLVIFGSFFVLNLVLGVLSGEFSREKQQLEEDLKGYLDWITQAEDIDSENESVNTDNAPAGDMEGETCLCRRKCRAAVKSNVFYWLVIFLVFLNTLTIASEHHQQPEWLTNVQDIANKVLLALFTGEMLLKMYSLGLQAYFVSLFNRFDSFVVCGGILETIIMSPLGISVLRCVRLLRIFKITRYWNSLSNLVASLLNSVRSIASLLLLLFLFIIIFSLLGMQLFGGKFN--DETRRSTFDNFPQSLLTVFQILTGEDWNSVMYDGIMAYGMLVCIYFIILFICGNYILLNVFLAIAVDNLADAESLTSAQKEEEEEKERKKLATKINIDEYTGEDNEEKNPYPVNDFPAMPQAKAFFIFSPSNKFRVLCHKIVNHNIFTNLILFFILLSSISLAAEDPV-KNDSFRNQILGYADYVFTGIFTIEIILKMTAYGAFLHKFCRNYFNILDLVVVSVSLISSGIQAINVVKILRVLRVLRPLRAINRAKGLKHVVQCVFVAIRTIGNIVIVTSLLQFMFACIGVQLFKGKFFYCTDTSKQTQAECRGENSDFNFDDVLQGMMALFAVSTFEGWPGLLYRAIDSHAEDVGPIYNYRVVISIFFIIYIIIIAFFMMNIFVGFVIVTFQEQGEQEYKNCELDKNQRQVEYALKARPLRRYIPYQYKVWYVVNSTYFEYLMFTLILLNTICLAMQSQSFNKAMNILNMLFTGLFTVEMILKLIAFKPRGYFSDPWNVFDFLIVIGSIIDVILSETEDNARISITFFRLFRVMRLVKLLSRGEGIRTLLWTFIKSFQALPYVALLIVMLFFIYAVIGMQMFGKIALRDSQINRNNNFQTFPQAVLLLFRCATGEAWQEIMLACSSSEDCGSHFAIFYFVSFYMLCAFLIINLFVAVIMDNFDYLTR

>Danio_rerio_Cav1.1_NP_001139622.1

SLLFLTLKNPFRKACINIVEWKPFEIIILLTIFANCVALAVFMPMPEEDTNNTNSNLESLEYIFLIIFTMECFLKIVAYGFLFHADAYLRNCWNILDFVIVTMGLFTVVVDFINDMKALRAFRVLRPLRLVSGVPSLQVVMSSILKSMLPLFHISLLVFFMVTIYAIIGLELFKCKMHKTCYHTDIIATGDDAQAAPCAQGRRCT-PGPNNGITHFDNLGFSMLTVYQCITTQGWTDVLYWVNDAIGEWPWLYFVTLILLGSFFILNLVLGVLCGEFTRERQQFDEDLKGYMEWITQAEVMDGLLPLQDGSETETLYELDILNFFRRKCRVWVKSKLFYWLVILLVFFNTLAIATEHHQQPDSLTNFQDNTNKALLSLFAVEMFLKMYAMGLPSYFMSLFNRFDCFVVSVGILELIVMSVMGISVLRCIRLLRIIKITRHWTTLSNLVASLLNSVRSIASLLLLLFLFIVIFALLGMQVFGGKFNPDDRVRRSNFDNFPQALITVFQILTGEGWNYVMYDGIMAHGILVSIYFIILFICGNYILLNVFLAIAVDNLAEAESLTSAQKEKAEEKKRKRLLAKLKVDEFESNVNEIKDPFPPADFPGMPEASAFFLFGPQNKFRKLCHRIINATTFTNIILLFILLSSISLAAEDPI-DPMSFRNQVLAYADYVFTSVFTAEIVLKMTTYGAFLHKFCRNSFNILDLIVVGVSLLSMGMEAISVVKILRVLRVLRPLRAINRAKGLKHVVQCVFVAIKTIGNIVLVTMLLDFMFACIGVQLFKGKFLYCTDPLKMTAEECQGMNSDLNFDNVLNGMLALFTVSTFEGWPDLLYKAIDSNLENMGPVYNNHIEISIFFIVYLILIAFFMMNIFVGFVIVTFQEQGEQEYKNCELDKNQRQVQYALKARPLRCYIPYQYQVWYIVTSCYFEYLMFLLIMLNTMCLGMQSDHITDLADTLNVIFTVLFTVEMILKLGAFKAKGYFGDPWNVFDFVIVVGSIVDVILSEDSENMSVSITLFRLFRVMRLVKLLNRFEGIRNLLWTFIKSFQALPYVALLIVMLFFIYAVIGMQVFGKIALLDTIINRNNNFQTFPQAVLLLFRCATGEGWHEIMLGCLEEYTCGSGFAILYFMSFYMLCAFLIINLFVAVIMDNFDYLTR

>Cyanea_capillata_Cav1_AAC63050.1

ALFCLTLDNPVRSAAITIVDWKPFDFFILASIFANCAALAAYEPLPAGDMTSTNQDLEVAEYFFLAVFAIEGLLKIIAYGFILHPGAYLRNGWNILDFSIVVIGFASMIFEEYLDVKALRAFRVLRPLRLVSGVPSLQVVMNSIVKAMLPLFHIALLVVFVIIIYAIIGVELFTGKLHQTCYD-NITNLPASSEPKPCSTGRQCP-EGPNYGITNFDNVALAALTVFQCTTLEGWTDVLYDINNVSGGWPWIYFVTLITFGSFFVLNLILGVLSGEFAREKHMLDDAVKGYLDWINQASDIERKLSSRRASGISHASSGIYNIRLRRQCRGIVKSQTFYWMVIIAVFLNSLVLAVEHYDQPDYITMFLDRANYFFLGLFTFEMLLKIYCLGIYGYLNSLFNRFDCLVVLSSLLEVAGWPPIGISVLRCVRLLRIFKVTRYWESLSNLVQSLVNSIKSIGSLLLLLSLFILIFSLLGMQIFGGRFNDEQAPPRTNFDSFWRSLITVFQILTGEDWNAVMYVGIQSWGIIAIIYFVALVIVGNYILLNVFLAIAVDNLADAENMTKVNEEEKRKKKDAKLMLSGSEGPEDVELGNPKSKNGTLRHMGMPPESSFFIFSANNKLRYLCYRLAVNKIFINSILVLIIMSSVALAAEDPI-GRDVLRNKILGYFDIFFTAMFTFEVTVKMIAFGVILHKFCRSFFNQLDLVIVAVSWAAIMLSATSVVRILRVLRVLRPLRAINRAKGLKHVVQCVFLAIKSIGNIMIVTLLFQFLFAVIGIQLFKGTFFYCTDRSKMTAEECKGEKHTFRFDNVFQAYLSLFVVMTFEGWPSILEHSIDSTTVNQGPKFNNRPFVAIYYVIYIIIIAFFMINIFVGFVIVTFQNEGEEEFADCELDKNQRKVEYVLTVKPTYRFVPFQYHIWRVVTSRLFEYMIFGFILGNTIVLAAASKLYERVLDGFNIGFTAVFLLECVLKLMAFNAKNYFRDPWNIFDFVIVVGSIADIIIGEISKDGGIKVNFFRLFRALRLVKLLSQGDGIRTLLWTFMKSFQALPFVGLLILLLFFIYAVIGMQVFGTIRLDSTVINSNNNFQTFPQALIVLFRSATGENWQQIMMACVPSKTCGTDFAYLYFMSFYMICSFLIINLFVAVIMDNFDYLTR

>Cteno_mnelei_Cav_evg1108101

ALCCLYESHPLRKLCIYIARSRPFEIFIYLAIFANCVSLAMYKPMPKSDVDETSSVLDTLEYIFIGIFWVESIVKIIAQGFMCHHGSYLRSFWNVLDFFIVIVGTLSVMSLDTGGIKALRALRVLRPLKFIAGCRPLQVVMNSILMALIPLMNVAVLILFFIIFCAIMGMELFRGQFNQACAD--MHTEEIPDNYHPCDEGFMCP-EGPNEGITTFNHIGLASLTVFQIITLEGWTDIMYWSMDAIGELPPLFYCNIVIWGAFFMLNLVLGVLAGEFSKRNDDYKKELEGYELWIREGEKISGGKNSYDGNHLEDNFDDVSNILLRERAAKIVKSKSMYWVVMFLVTMNTISVSLKWYGMPEEMDKNMKHVEFVFTFIFIIEMTIRMYGLGIEQYFKSKFNTFDFTVIFASLFEQFDGADMGLTVFRSLRLLRLFKVTRYWTSLRNLAAALLNSLKSIVSLLFLLFLFLLIFALLGMQLFGGHFVFAEQVPRVNFDSFLTALLAVFQMLTGEDWNVIMYNAINALGGICSIYFIVFMVIMNYTLLNVFLAIAVDGLADFEMMNDAEKEDEKKEEAEKDALRNRAQAYRADYPDHADTPISIEDSDIVPHRSFFIFSPTNPLRRFCHWIVNLRHFDNFILFIILLSCIMQMLQNPK-DLESDNNRFIEYLEYGVTVIFCLEMILRVIDLGFVIHPYLRDPWNVLDAVVVLVSILRVAVK--TVKRLTNVMLSMRSLKSINRIQKLKNVCLCLVKSVGSILNLLLIAILLTFMFSVMGVHMLEGKFFFCTDSSKKTAPECMGQKHFFRFDDVQLAMLTLFTVATFEGWPDVMTNMMDASLKDHGPIENANPALCIYIVVYLVVMSFFMINIFVGFVIVTFQEKGEKDNETSILDRNKRSIEFSLKVKPQNKYIPLRYKLWQVVSSSIYTNLIMFIIFINTIQLMMMSRDYETGLETVNAILVAVFTVDIILKLAAYGVKQYFGGSWNIFDFVVLVGSFIDIMVSTWYTDNFMISRLVKMFRAARLIKLLNYGGEMRTLLFVFLKALKSLPHVMFLVMLTFYIYAIIGMQIFGKVIADPYVITRYNNFESFENAMILLFRCSTGEAWHAIMRD-LKDTNCGSSFSLIYFCTFVITSQFLVVNLFVSVIIDNFDYLTR

>Cteno_horcal_Cav

ALCCLYDHHLLRRVCNAIARSRPFEIFIYLAIFANCVSLAMYKPMPKGDVDETSSVLDRLEKFFIGIFWVESIVKIVAQGFMCHHGSYLRSFWNVLDFFIVIVGTLSVMDLDTGGIKALRALRVLRPLKFIAGCRPLQVVMNSILMALIPLMNVAVLILFFIIFTAIMGMEFFRGQFNHACAD--MHTQQIPDNFHPCD-GFVCP-LGPNEGITTFDHIGLACLTVFQIITLEGWTDIMYWSMDAIGQLPPFFYFNIVIWGAFFMLNLVLGVLAGEFSKRNNDFRNELEGYEAWVKEGERICEALKYRGKGVDENHLHDDTSSRLRERAAKIVKSPAMYWIVMFLVTMNTISVSLKWYGMPEEMNRRMKSVEFVFTFIFIIEMTIRMYGLGVEQYFKSKFNTFDFTVIFASLFEQFDGADMGLTVFRSLRLLRLFKVTRYWTDLRSLAAALLNSLKSIISLLFLLFLFLLIFALLGMQLFGGHFVFAEQVPRVNFDSFLTALLAVFQMLTGEDWNVIMYNAINALGGIASIYFIVFMVIMNYTLLNVFLAIAVDGLADFDMVSDAEKEHEKKEEAEKVRDFQSADNLEHSIDIDRHAIEESEDEGIVPHPSFFIFSPTNPLRRFCHWIVNMRHFDNFILFIILLSCIMQMLQNPK-DLESDNNRFIEYLEYGVTVIFCLEMILRVIDLGFVIHPYLRDPWNVLDAVVVLVSILRVAVKASTVKRLTNVMLSMRSLKSINRIQKLKNVCLCLIKSVGSILNLLLIAILLTFMFSVMGVHMLEGKFFYCTDSSKKTEDECKGDSHFFRFDNVQSAMLTLFTVATFEGWPDVMTNMMDASQKDMGPIENTNPALCIYIVVYLVVMSFFMINIFVGFVIVTFQEKGEKDNDSELLDRNKRSIEFSLKVKPQNKYIPLRYKLWQIVSSNFYTNLIMFIIFINTIQLMMMSDEYEYALEIVNLMLVAVFTIDILLKLAAYGVKQYFGSSWNVFDFVVLVGSFVDILVSTWYKNNFMIVRLVKMFRAARLIKLLNYGGEMRTLLFVFLKALKSLPHVMFLIMLTFYIYAVIGMQIFGMVEANPYTITRYNNFASFQMAMIILFRCSTGEAWHAIMRD-LPDSNCGSKFSEAYFCTFVITSQFLVVNLFVSVIIDNFDYLTR

>Cteno_berova_Cav

ALCCLYETAPLRKLCIYIARSRPFEIFIYLAIFANCVSLAMYKPMPKGDVDETSSVLDTLELFFIGIFWVESIVKIIAQGFLCHHGSYLRSFWNVLDFFIVIVGTLSVMNLDTGGIKALRALRVLRPLKFIAGCRPLQVVMNSILMALIPLMNVAVLILFFIIFCAIMGMELFRGQFNLACAD--MHTGDIPDNYHPCDEGFECP-EGPNEGITTFNHIGLASLTVFQIITLEGWTDIMYWSMDAIGALPPLFYCNIVIWGAFFMLNLVLGVLAGEFSKRNDDYKKELEGYELWIKEGEKIVEGNHLEDNFDDTSHLSDPERGILREKTARIVKSKSMYWVVMFLVTMNTISVSLKWYDMPPEMNSNMKNVEFVFTFIFIIEMTIRMYGLGIEQYFKSKFNTFDFTVIFASLFEQFNGADMGLTVFRSLRLLRLFKVTRYWTSLRNLASALLNSLKSIVSLLFLLFLFLLIFALLGMQLFGGHFVFAEQVPRVNFDSFLTALLAVFQMLTGEDWNVIMYNAISALGGICSIYFIVFMVIMNYTLLNVFLAIAVDGLADFEMMNDAEKEDEKKEEAEKDALRNRAQAYRADYPDHADTPISIEDSDIVPYTSFFIFTPTNPLRRFCHWIVNLRHFDNFILFIILLSCIMQMLQNPK-DLESDNNRFIEYLEYGVTVIFCMEMILRVIDLGFVIHPYLRDPWNVLDAVVVLVSILRVAVKAQTVKRLTNVMLSMRSLKSINRIQKLKNVCLCLIKSVGSILNLLLIAILLTFMFSVMGVHMLEGKFFYCTDSSKKTQLECMGKKHFFRFDDVQLAMLTLFTVATFEGWPDVMTNMMDASEKDRGPVENANPALCIYIVVYLVVMSFFMINIFVGFVIVTFQEKGEKDNEGTMLDRNKRSIEFSLKVKPQNKYIPIRFKLWQVVSSSIYTNLIMFIIFINTIQLMMMSSKYEEGLEMVNAILVAVFTVDIILKLAAYGVKQYFGGSWNIFDFVVLVGSFIDIMVSNWYTDNFMISRLVKMFRAARLIKLLNYGGEMRTLLFVFLKALKSLPHVMFLVMLTFYIYAIIGMQIFGKVEADNYTITRYNNFASFQNAMILLFRCSTGEAWHAIMRDTPGDGTCGSSFSKIYFCTFVITSQFLVVNLFVSVIIDNFDYLTR

>Crocodylus_porosus_Cav3.2_XP_019397684.1

VFFCLQQATRPRSWCLRLVCNPWFEHVSMLVILLNCVTLGMFQPCEDVKKSERCTILEAFDHFIFAFFAVEMVIKMVALGI-FGQKCYLGDTWNRLDFFIVMAGMLEYSLDGHNSLSAIRTVRVLRPLRAINRVPSMRILVTLLLDTLPMLGNVLLLCFFVFFIFGIVGVQLWAGLLRNRCFFHPYYQPDDGEDNFICSSKVECTDVNPHNGAINFDNIGYAWIAIFQVITLEGWVDIMYYVMDAHSFYNFIYFILLIIVGSFFMINLCLVVIATQFSSEPGSCYEELLKYICHIFRKVKRRRASSRLSGLNVPCPLPSPQASAFGNKLKKIVESKYFNRGIMIAILINTLSMGIEYHEQPDELTNALEISNIVFTSMFALEMLLKLLAFGIFGYIKNPYNIFDGIIVIISVWEIIGQSDGGLSVLRTFRLLRVLKLVRFMPALRRQLVVLMKTMDNVATFCMLLMLFIFIFSILGMHLFGCKFSGDTVPDRKNFDSLLWAIVTVFQILTQEDWNVVLYNGMASTSSWAALYFVALMTFGNYVLFNLLVAILVEGFQAGDSDTDEDKNFDEDFEKLKDLLQLPSMRHSLSINPMAMLPTEYQDCNNHEDWSLYLFSPQNRFRAMCQKVIAHKMFDHVVLVFIFLNCITIALERPDIDPHSTERIFLSVSNYIFTAIFVAEMMVKVVALGFFSGEYLQSSWNVLDGVLVFVSIIDIIVSILGVLRVLRLLRTLRPLRVISRAPGLKLVVETLISSLRPIGNIVLICCAFFIIFGILGVQLFKGKFYSCEDTKITTKADCTNVRRKYNFDNLGQALMSLFVLSSKDGWVNIMYDGLDAVGIDQQVATKD------------------ILRFFPGIQCESAWEVIEETLAVGSYRAGDRELGFSLSTEPYYAYSPARRYIHTLCTSHYLDLFITFIIGVNVITMSMQPKSLDEALKYCNYVFTIVFVFEAVLKLVAFGFRRFFKDRWNQLDLAIVLLSIMGITLEEALPINPTIIRIMRVLRIARVLKLLKMATGMRALLDTVVQALPQVGNLGLLFMLLFFIYAALGVELFGKLDCSEEGLSRHATFTNFGMAFLTLFRVSTGDNWNGIMKDTLREDKHCPVISPVYFVTFVLIAQFVLVNVVVAVLMKHLEESNK

>Crocodylus_porosus_Cav3.1_XP_019398025.1

SFPASAWKGLLVSLTERLCTW--FERVSMLVILLNCVTLGMFHPCEDTAGSPRCRILQSFDDFIFAFFAVEMIVKMIALGI-FGKKCYLGDTWNRLDFFIVIAGMLEYSL--DLSFSAVRTVRVLRPLRAINRVPSMRILVTLLLDTLPMLGNVLLLCFFVFFIFGIVGVQLWAGLLRNRCFLERYYQTENEDENFICSQGLECTEHNPFKGAINFDNIGYAWIAIFQVITLEGWVDIMYFVMDAHSFYNFIYFILLIIVGSFFMINLCLVVIATQFSSEPGSCYDELLKYLVYVTRKASKQSTMHKLLENQSTGACQSSCKIVVCETFRKIVDSKYFGRGIMIAILINTLSMGIEYHEQPEELTNALEISNIVFTSLFALEMLLKVLVYGPFGYIKNPYNIFDGIIVVISVWEIVGQQGGGLSVLRTFRLMRVLKLVRFMPALQRQLVVLMKTMDNVATFCMLLMLFIFIFSILGMHLFGCKFAGDTLPDRKNFDSLLWAIVTVFQILTQEDWNKVLYNGMASTSSWAALYFIALMTFGNYVLFNLLVAILVEGFQTGESDSEGDVSLEEEGGLKKHLLQVPSLYRTSSMYSSRTSASEHQDCNERDSWSIYIFAPHSKFRLMCNKIITHKMFDHVVLVIIFLNCITIAMERPKIEPHSAERIFLTLSNYIFTVIFLAEMTVKVVALGLCFGEYLKSSWNVLDGVLVLISVIDILVSILGMLRVLRLLRTLRPLRVISRAQGLKLVVETLMSSLKPIGNIVVICCAFFIIFGILGVQLFKGKFFVCQDTRITNKSDCAEVRHKYNFDNLGQALMSLFVLASKDGWVDIMYDGLDAVGVDQQPVMNYNPWMLLYFISFLLIVAFFVLNMFVGVVVENFHKCRQHQEEEEAKRREEKRRRLEKKRRPYYSYSRFRLLIHQMCTSHYLDLFITGVIGLNVITMAMQPKVLDEALKICNYIFTVIFVLESVFKLIAFGFRRFFQDRWNQLDLAIVLLSIMGITLEESLPINPTIIRIMRVLRIARVLKLLKMAVGMRALLDTVMQALPQVGNLGLLFMLLFFIFAALGVELFGDLECDDEGLGRHATFRNFGMAFLTLFRVSTGDNWNGIMKDTLQESTCYTVISPIYFVSFVLTAQFVLVNVVIAVLMKHLEESNK

>Crocodylus_porosus_Cav2.3_XP_019386495.1

SLFLFGEDNVVRKYAKKLIDWPPFEYMILATIIANCIVLALEQHLPGDDKTPMSRRLEKTEPYFIGIFCFEAGIKIVALGFVFHKGSYLRNGWNVMDFIVVLSGILATAGTHFNDLRTLRAVRVLRPLKLVSGIPSLQIVLKSIMKAMVPLLQIGLLLFFAILMFAIIGLEFYSGKLHRACYANNSGELEELDPPHPCG-VQGCP-IGPNDGITQFDNILFAVLTVFQCITMEGWTTVLYNTNDALGTWNWLYFIPLIIIGSFFVLNLVLGVLSGEFARRQQQIERELNGYRAWIDKAEEVMNRTEAMNRDSSDEHCVDISSVLLRISVRHMVKSQVFYWIVLSLVALNTACVAIVHHNQPLWLTHLLYYAEFLFLGLFLLEMSLKMYGMGPRLYFHSSFNCFDCGVTVGSIFEVVPGTSFGISVLRALRLLRIFKITKYWASLRNLVVSLMSSMKSIISLLFLLFLFIVVFALLGMQLFGGRFNFMDGTPSANFDTFPAAIMTVFQILTGEDWNEVMYNGIRSQGMWSSIYFIVLTLFGNYTLLNVFLAIAVDNLANAQELTKDEQEEEEAFNQKHALKDRSPPKVPHELNQGNNVSLTEQDCSMVPHSSMFIFSTTNPVRRACHYIVNLRYFEMCILLVIAASSIALAAEDPV-LTNSDRNKVLRYFDYVFTGVFTFEMVIKMIDQGLILQDYFRDLWNILDFIVVVGALVAFALADIKTIKSLRVLRVLRPLKTIKRLPKLKAVFDCVVTSLKNVFNILIVYKLFMFIFAVIAVQLFKGKFFYCTDSSKDTEKDCIGKRHEFHYDNIIWALLTLFTVSTGEGWPQVLQHSVDVTEEDRGPSRSNRMEMSIFYVVYFVVFPFFFVNIFVALIIITFQEQGDKMMEECSLEKNERAIDFAISAKPLTRYMPFQYRVWHFVVSPSFEYTIMAMIALNTVVLMMAPYTYELALKYLNIAFTMVFSLECVLKIIAFGFLNYFRDTWNIFDFITVIGSITEIILTDLVNTSGFNMSFLKLFRAARLIKLLRQGYTIRILLWTFVQSFKALPYVCLLIAMLFFIYAIIGMQVFGNIKLDESHINRHNNFRSFLGSLMLLFRSATGEAWQEIMLSCLENERCGTDLAYVYFVSFIFFCSFLMLNLFVAVIMDNFEYLTR

>Crocodylus_porosus_Cav2.2_XP_019396155.1

SLFIFSEDNVIRKYAKRITEWPPFEYMILATIIANCIVLALEQHLPDGDKTPMSERLDDTEPYFIGIFCFEAGIKIIALGFVFHKGSYLRNGWNVMDFVVVLTGILATAG----DLRTLRAVRVLRPLKLVSGIPSLQVVLKSIMKAMVPLLQIGLLLFFAIVMFAIIGLEFYMGKFHKTCFS----NETGEEVGFPCGEARQCE-QGPNYGITNFDNILFSVLTVFQCITMEGWTDILYNTNDAAGTWNWLYFIPLIIIGSFFMLNLVLGVLSGEFARRQQQIERELNGYLEWIFKAEEVMSKNDLIHAEEGEDHFTDICSVMFRFFIRRMVKAQSFYWIVLCVVTLNTLCVAMVHYAQPEKLTTALYFAEFVFLGLFLTEMSLKMYGLGPRNYFHSSFNCFDFGVIVGSIFEVIPGTSFGISVLRALRLLRIFKVTKYWNSLRNLVVSLLNSMKSIISLLFLLFLFIVVFALLGMQLFGGQFNFQDETPTTNFDTFPAAILTVFQILTGEDWNAVMYHGIESQGMFSCVYFIVLTLFGNYTLLNVFLAIAVDNLANAQELTKDEEEMEEATNQKLALSGNREGDPGSKGENGEEPHRRHRIRNILPYSSMFILSPTNPIRRLFHYIVNLRYFEMVILIVIALSSIALAAEDPV-QAESPRNDALKYLDYIFTGVFTFEMVIKMIDLGLLLHPYFRDLWNILDFIVVSGALVAFAFSDINTIKSLRVLRVLRPLKTIKRLPKLKAVFDCVVNSLKNVLNILIVYMLFMFIFAVIAVQLFKGRFFYCTDESKELEKDCRGKKYEFHYDNVLWALLTLFTVSTGEGWPTVLKHSVDATYEEQGPSPGYRMEMSIFYVVYFVVFPFFFVNIFVALIIITFQEQGDKVMSECSLEKNERAIDFAISAKPLTRYMPFQYKMWKFVVSPPFEYFIMVMIALNTIVLMMAPEAYEEMLKCLNIVFTSMFSMECVLKIIAFGVLNYFRDAWNVFDFVTVLGSITDILVTEADTDNFINLSFLRLFRAARLIKLLRQGYTIRILLWTFVQSFKALPYVCLLIAMLFFIYAIIGMQVFGNIALDDSSINRHNNFRTFLQALMLLFRSATGEGWHEIMLSCLTKDECGSDFAYFYFVSFIFLCSFLMLNLFVAVIMDNFEYLTR

>Crocodylus_porosus_Cav1.3_XP_019392560.1

--------MLKNALGHRILSW--------------------YN--------------EKVEYAFLIIFTIETFLKIIAYGLLLHPNAYVRNGWNLLDFVIVIVGLFSVILEQLTDVKALRAFRVLRPLRLVSGVPSLQVVLNSIIKAMVPLLHIALLVLFVIIIYAIIGLELFIGKMHKSCFL-IGTDILVEEDPAPCAFGRQCS-VGPNGGITNFDNFAFAMLTVFQCITMEGWTDVLYWMNDAMGELPWVYFVSLVIFGSFFVLNLVLGVLSGEFSREKQQLEEDLKGYLDWITQAEDIDSETESVNTENVSGEGENPICCFNRRRCRAAVKSVSFYWLVIVLVFLNTLTISSEHYNQPNWLTQIQDIANKVLLALFTCEMLVKMYSLGLQAYFVSLFNRFDCFVVCGGIVETIIMSPLGISVFRCVRLLRIFKVTRHWTSLSNLVASLLNSMKSIASLLLLLFLFIIIFSLLGMQLFGGKFNDETQTKRSTFDNFPQALLTVFQILTGEDWNAVMYDGIMAYGMIVCIYFIILFICGNYILLNVFLAIAVDNLADAESLNTAQKEEAEEKERKKNASKVTIAEYGEGEDEDKDPYPPCDVPVIPEGSAFFIFSSTNPIRVGCHRLINHHIFTNLILVFIMLSSISLAAEDPI-RSHSFRNNILGYFDYAFTAIFTVEILLKMTTFGAFLHKFCRNYFNLLDLLVVGVSLVSFGIQAISVVKILRVLRVLRPLRAINRAKGLKHVVQCVFVAIRTIGNIMIVTTLLQFMFACIGVQLFKGKFYRCSDEAKQNPEECRGQNSDFNFDNVLSAMMALFTVSTFEGWPALLYKAIDSNGENVGPIYNYRVEISIFFIIYIIIIAFFMMNIFVGFVIVTFQEQGEQEYKNCELDKNQRQVEYALKARPLRRYIPYQYKFWYVVNSTGFEYIMFVLIMLNTLCLAMQSKLFNDAMDILNMVFTGVFTVEMVLKVIAFKPKHYFTDAWNTFDALIVVGSVVDIAITESEDSARISITFFRLFRVMRLVKLLSRGEGIRTLLWTFIKSFQALPYVALLIAMLFFIYAVIGMQVFGKVAMRDNQINRNNNFQTFPQAVLLLFRCATGEAWQEIMLACLEEYTCGSNFAIIYFISFYMLCAFLIINLFVAVIMDNFDYLTR

>Crocodylus_porosus_Cav1.2_XP_019400097.1

ALLCLTLKNPIRRACISIVEWKPFEIIILLTIFANCVALAIYIPFPEDDSNATNSNLERVEYLFLIIFTVEAFLKVIAYGLLFHPNAYLRNGWNLLDFIIVVVGLFSAILEQATDVKALRAFRVLRPLRLVSGVPSLQVVLNSIIKAMVPLLHIALLVLFVIIIYAIIGLELFMGKMHKTCYHGALADTPAEEDPSPCAPGRQCQ-EGPKHGITNFDNFAFAMLTVFQCITMEGWTDVLYWVNDAIGDWPWIYFVTLIIIGSFFVLNLVLGVLSGEFSREKQQLEEDLKGYLDWITQAEDIDSETESVNTDNVAGADIEGENCFCRRKCRAAVKSTVFYWLVIFLVFLNTLTIASEHYHQSNWLTEVQDTANKVLLALFTAEMLLKMYSLGLQAYFVSLFNRFDCFIVCGGILETIIMPPLGISVLRCVRLLRIFKITRYWNSLSNLVASLLNSVRSIASLLLLLFLFIIIFSLLGMQLFGGKFNDEMQTRRSTFDNFPQSLLTVFQILTGEDWNSVMYDGIMAYGMLVCIYFIILFICGNYILLNVFLAIAVDNLADAESLTSAQKEEEEEKERKKLATKINVEDYQPNENEEKNPYPTTEAPGMPDASAFFIFSPSNRFRVHCHRIVNDNIFTNLILFFILLSSISLAAEDPV-RHYSFRNQILFYFDIVFTVIFTIEIALKMTAYGAFLHKFCRNYFNILDLLVVSVSLISFGIQAINVVKILRVLRVLRPLRAINRAKGLKHVVQCVFVAIRTIGNIVIVTTLLQFMFACIGVQLFKGKLKSCSDSSKQTPAECKGENSKFDFDNVLNAMMALFTVSTFEGWPELLYRSIDSHMEDVGPIYNHRVEISIFFIIYIIIIAFFMMNIFVGFVIVTFQEQGEQEYKNCELDKNQRQVEYALKARPLRRYIPYQYKVWYVVNSTYFEYLMFILILLNTICLAMQSCMFKEAMNILNMLFTGLFTVEMVLKLIAFKPKGYFSDPWNVFDFLIVIGSIIDVILSEAEENSRISITFFRLFRVMRLVKLLSRGEGIRTLLWTFIKSFQALPYVALLIVMLFFIYAVIGMQVFGKIALNDTGINRNNNFQTFPQAVLLLFRCATGEAWQEIMLACLEDQSCGSSFAIFYFISFYMLCAFLIINLFVAVIMDNFDYLTR

>Crocodylus_porosus_Cav1.1_XP_019401893.1

ALFCLTLQNPLRKACISIVEWKPFEIIILLTIFANCVALAVYLPMPEDDTNATNSRLEKIEYVFLIIFTIEAMLKIIAYGFLFHTDAYLRSGWNVLDFAIVSLGLFTVTLEQISDVKALRAFRVLRPLRLVSGVPSLQVVLNSIIKAMVPLLHIALLVLFMIIIYAIVGQELFKGKMHKTCYYTDIIATVGSEKPAPCTSGRHCT-PGPNNGITHFDNFGFAMLTVYQCITMEGWTEVLYWVNDAIGEWPWIYFVSLILLGSFFILNLVLGVLSGEFTREKQQLEEDLKGYMDWITHAEVMDRGEGMLPLDEGSSETESLYEILFRRKCREVVKSKFFYWLVILLVALNTLSIASEHHMQPDWLTHVQDNANRVLLSLFAAEMLLKMYALGLRQYFMSLFNRFDCFVVCAGILETILMSPLGISVLRCIRLLRIFKITKYWTSLSNLVASLLNSIRSIASLLLLLFLFIVIFSLLGMQLFGGKYDEDMEVRRSTFDNFPQALISVFQVLTGEDWNSIMYNGIMAYGMLVCIYFIILFVCGNYILLNVFLAIAVDNLAEAESLTSAQKAKAEERKRRKMSAKLKVDEFESNVNEIKDPYPSADFPGMPEASAFFIFSPTNKFRILCHRIVNATWFTNFILLFILLSSISLAAEDPI-RAESFRNQILGYFDIGFTSVFTVEIVLKMTAYGAFLHKFCRNSFNILDLLVVAVSLISMGIQTISVVKILRVLRVLRPLRAINRAKGLKHVVQCVFVAIKTIGNIVIVTTLLQFMFACIGVQLFKGKFNSCTDPSKITERECRGQHNDFHFDNVLSAMMSLFTVSTFEGWPQLLYKAIDTHTEDMGPIYNYRVEMAIFFIIYIILIAFFMMNIFVGFVIVTFQEQGESEYKNCELDKNQRQVQYALKARPLRRYIPYQYQIWYVVTSSYFEYLMFFLILLNTICLGMQSDEMNHASDILNVTFTILFTVEMFVKLMAFKAKGYFGDPWNVFDFLIVIGSIIDVILSESEDNSRVSITFFRLFRVLRLVKLLSRGEGVRTLLWTFIKSFQALPYVALLIVMLFFIYAVIGMQMFGKVAMVDTQINRNNNFQTFPQAVLLLFRCATGEAWQEILLASYEEYTCGTGFAYFYFISFYMLCAFLIINLFVAVIMDNFDYLTR

>Crassostrea_gigas_Cav3_XP_011439125.1

SLRCLPQTNKLRFICLKIITWTWFERISMAVILLNCITLGMYQPCSDLETTTRCKILEKFDHFIFAFFAAEMIIKIIAMGL-YGKYTYLDDSWNRLDCFIVIAGAIEYGVDSENSLSAIRTIRVLRPLRAINRIPSMRILVMLLLDTLPMLWNVLLLCFFVFFIFGIIGVQLWAGVLRNRCFLSPYYVPKSMPPGFICSKGIKCNDKNPFQGGVSFDNIGLAWVAIFQVISLESWVNIMYYVQDAHSFWDWIYFVALIVIGSFFMINLCLVVIATQFSSEPGGCYAEILKYLAHLWRRAKRKSCHKLAANSLLHVDSGSRVRINFQQVMKTLVEHRFFQRGILTAILINTLSMSVEYHNQPRELTEAVEYSNMVFSILFAVEMFFKLCAYGIIGYIQDGFNVFDGLIVILSMIEFTQGGASGLSVLRTFRLLRILKLVRFMPALRRQLVVMLRTMDNVATFFALLVLFIFIFSVLGMNLFGGKFCMDCQCERANFNSLLWSVVTVFQVLTQEDWNTVLYNGMETTSSWASLYFIALMTLGNYVLFNLLVAILVEGFSTDRSLLSETNNNEEKSENIKLKKRPVPSPRNSFKIRNNSCSSSQSNCNTRHEYALYLLHPNNRLRRVCHHLMAQRWFDNTVLFFIALNCITLAMERPNIPPDSVEREFLNYSNYVFTFVFSVEMMIKVLAKGLVIGQYLKSGWNVMDGFLVGISLVDILISIFGILRVFRLLRTLRPLRVISRAPGLKLVVQTLLSSLRPIGNIVIICCTFFIIFGILGVQLFKGTFYYCKTVRIKNKTQCLMINQKYNFDDLGQALMALFVLASKDGWVSIMYTGLDAVGIDQQPIENYNEWRLIYFISFLLLVGFFVLNMFVGVVVENFHKCRQDQEKEERERRTAKR-RYKMEQKPYWAYSPSRLIIHQVVNSKYFDLAIAGVIGLNVITMAMMPEELEFALKIFNYFFTSVFIIEATLKILALGFMRYIKDRWNQLDIFIVILSIVGIILEEVIPINPTIIRVMRVLRIARVLKLLKMAKGIRALLDTVIQALPQVGNLGLLFFLLFFIFAALGVELFGRLECSEEGLGEHAHFRDFAMAFLTLFRVATGDNWNGIMKDTLLTNCCVSFIAPVFFVVFVLMAQFVLVNVVVAVLMKHLEYKHK

>Crassostrea_gigas_Cav2_XP_019920407.1

SLFIFSEENFIRKYAKIIIEWGPFEYMVLLTIIANCIVLALEEHLPKDDKTPLAVQLEETEIYFVVIFLVEALLKIVALGFVLHKGAYLRNIWNIMDFVVVVTGIITMAA----DLRTLRAVRVLRPLKLVSGIPSLQVVLKSIIRAMTPLLQVCLLVIFAIIIFAIVGLEFYSGAFKNACFKEDDIYIGDEANIRPCSSAFKCQ-VGPYFGITNFDNIAYAMLTVFQCITMEGWTEVLYYTNDAIGYINWLYFYPLIILGSFFMLNLVLGVLSGEFARRQQQIDMELSGYLEWICKAEEVIKEPSDENNEDNDSDLLSDINIHFRFSLRRLVKSQPFYWTVIVLVFLNTVCTASEHYGQPKWHEEFLYYTEFVFLGLFIFEMLIKMYGLGVRIYFQSSFNIFDCGVIIVSIIEVIDGASFGISTLRALRLLRVFKVTRYWSSLRNLVVSLLSSMRSIVSLLFLLFLFILIFALLGMQLFGGEMNFDDGRPPAHFDTFPIALLTVFQILTGADWNEVMYNGIRAHGMFYSIYFIILVVFGNYTLLNVFLAIAVDNLTNAQEMTAAEEEEEVGRKEHLEEVTKKSNISQNQFDNDGLDTENNDLVNMLPYSSMFIFGPKNPIRRFCHFVVNLRYFDLFIMIVICASSFALATEEPV-NEDAFRNKILNYFDYVFTIVFTVEMILKVIDLGVFLHPYCRNLWNILDATVVICAVVAFFFDNLNTIKSMRVLRVLRPLKTINRVPKLKAVFDCVVNSLKNVANILIVYMLFQLIFAVIAVQLFKGKFFYCTDESKSTEEECRGLRRDFHYDNLFEAMLTLFTVTTGEGWPGILHNSMDSTYEDQGPKPGNRMEMAIFYVVFFIVFPFFFVNIFVALIIITFQDQGEAELEDAQLDKNQKQIDFAVNARPTSRYMPIKYKIWRLVVSTKFEYFVMTLIALNTIVLMMMSARYKDILKYLNMGFTIMFSIECTLKLIGCGK-NYFHDPWNVFDFITVVGSIIDVLVNEGSTYSSFNVGVFRLFRAARLIKLLRQGYTIRLLLWTFLQSFKALPYVCLLILMLFFIYAIIGMQVFGNIKLDSTDLNRHNNFRNFLYALMLLFRCATGENWQAIMIACLPPKTCGSAIAYVYFVSFMFLSSFLMLNLFVAVIMDNFDYLTR

>Crassostrea_gigas_Cav1_XP_011452714.1

ALFCLTLENPIRKLCIRIVEWKAFEYLILLTIFANCVALAVFQPFPNLDSNEVNLALERVEYVFLVIFTLEAIMKIIAYGFMLHSGAYLRNGWNILDFIIVVIGIITPVFSLFNDVKALRAFRVLRPLRLVSRAPSLQVVLNAIVRAMVPLLHIALLVIFVIFIYAIIGLELFSGSMHETCFDKQTNSVMTLSDVHPCGKGFSCP-AGPNDGITNFDNFGLAMLTVFQCITLEGWTDVLYNINDSLGSWPWTYFISLIIIGSFFVLNLVLGVLSGEFSREKKQLEEDLRGYLDWITQAEDIDHKNQEIPSVKTEDVES-----RCRRMCRKLVKSQAFYWTVIVMVFLNTLVLTSEHHKQPQWLDSFQAIANLFFVILFTLEMLLKMYSLGLQGYFVSLFNRFDSLVVLFSIIEVIVLPPLGVSVLRCARLLRVFKATRYWSSLRNLVASLLNSMRSIASLLLLLFLFIVICALLGMQLFGGKFNNTEDKPRSNFDTFWQSLLTVFQILTGEDWNMVMYDGINSYGLITCLYFVILFICGNYILLNVFLAIAVDNLADAQSLTEDEAEKEDEKERIRSLDTCASDTSSYDLLEGETAVEDEVDEEIPKASSLFILQPSNKFRIICHKICNHPYFGNIVLACILISSGMLAAEDPL-QSQSKRNEILNYFDIFFTSVFTVEIIIKVITYGLIVHKFCRSFFNILDFTVVGVSIISFVLDAISVVKILRVLRVLRPLRAINRAKGLKHVVQCVIVAIRTIYNIMLVTFLLQFMFAVIGVQLFKGRFFSCSDKSKLTESECRGTNNPLNYDNVPEAMLTLFTVSTFEGWPTLLYKSIDANEENNGPIHNNQPIVAVFYFIFIIVIAFFMMNIFVGFVIVTFQNEGEQEYKNCELDKNQRKIEFALKTRPTRRYIPWQYKIWWFVTSRAFEYGIFTLIILNTVILAMQSAAYSDALDYLNMIFTGVFTIEFILKLMAFRFRNYFGDPWNVFDFIIVLGSFIDIIYTENPGQGIISINFFRLFRVMRLVKLLSRGEGIRTLLWTFIKSFQALPYVALLIVMLFFIYAVIGMQLFGKISTDDSQIHRNNNFQTFPQAVLVLFRSATGEAWQDIMLSCVPSQTCGNDVAYFYFISFYMLCSFLIINLFVAVIMDNFDYLTR

>Cnida_nemvec_Cav3b_NVE7616

--------------------M---------------------------------------------------VVKMVAMGV-FGSHGYLQDTWNKLDFVIIVMGVIEKSLQGSDYLTIIRAFRVLRPLRAINKVPSIRILVTLLLDTLPMLGNVLLLSFLVFFVFGIIGVQLWQGKLRNRCFAQSVFYKPSFIQPFVCSGGRQCTGPNPFWGTISFDNIAIAWMVIFQVITLEGWSDVMYLVQDTHSFWNWIYFIILIVIGAFFLVNLCLVVITMQFQRCQGEPKEHIHHHHHHIYHHHHFHTRARWGSSEDLTQDLKRDLQLRFRAFCRRMAESRHFSRLIMIAILLNMICMGLEHHNQPQALTVTLEKTNIVFVTIFVLEMIINVISFGIMGYLSQLQNIFDGFVVVLSVTELL--GYARLSVFRSIRLLRIFKLVR---PVRYQLLVVIRTMTSVVTFFGLLFLFMFAFAILGMNLFGGEFYNISVPARTNFDSFLWAMVTVFQILTQENWNQVMFKGMRATSYWAALYFIALMTVGYYVLFNLLVAILVEGFTSGCAISQDTAEASRQRHHNQADDNVDDEKITEKLELETPEGNIPEACKVRHDWSFFMFAPDNRFRQRMRAVCSHRAFDYVILVFIIFSCAVLAIEAPDIAEQGLKRQIIDISMLVFTIIFTIEMLIKLVAMGLVLGPYLRDGWDVLDGFLVMVSWIDIIVTVLGTLRVFRALRTLRPLRVIRRAPGLKLVVQTLLYSLKPIGNTVLIAAIFFVMFGILGVQLFKGKFYYCEDSHVISKQECANVNRRYNFDDLLQALISLFVVSTKDGWVEIMHHGIDAVDVDVQPIVNYAEWRLVYFIPFLLLGGFLVLNMIVGVVVENFQRCRDEEEQARPHRKGKKANQMQAEDSLYYEYGPLRLKIHFICTHRNWDITIAAIICINVICMSLMPQSYEVFVETTNYFFTSVFVIEVVVKVIALGFVRYPKDRWNLIDLAIVLLSVTGIVLELDHLFNPTVIRTLRVLRITRVLKLVKLAKGVRSLLDTLFEALPQVANLGLLFFLLFFIYSCLGIQLFGSLECSHQGFNRHAHFRDFGTAMLTLFRIATGDNWNGILKDTLSKNCCLRYTSPLYFVTFVLAAQFVLVNVVIAVLMKHLKESKE

>Cnida_nemvec_Cav3a_NVE5017

--------------------M---------------------------------------------------LCKWLAMGV-FGKQGYLAENWNKLDCFIVAAGTFELCYDQGKYMTAVRAIRVLRPLRAINRVPSIRILVTLLLDTLPMLGNVLAMCSLIFSIFGIVGVQMWQGVLRSRCVAISAFYQPGGSSDLVCSLGRECGGINPLYDAVSFDNILIAWVAIFQVITLEGWTDIMYYIQDAHGFWNFIYFVVLIVIGSYFMTNLCLVVITTQFQYGREGCWLEILKWIGHVYRHTKRRLIGEMLTTATATVSVNGQPAIKFRDGCASSVDSKWFMYVIMGAIFVNTLTMGIEYYGQPQKMTDVLEIFNYIFTAIFGIEMIMKLIGLGFYGYIKDAFNIFDGTIVIISVVELFGDDDSGISVLRSFRLLRVFKLVRFLPALRRQLLVMIHTMDNVMTFLALLVIFMFTASILGMNLFGGKYRGVMETSRANFDDLFWAIVTVFQVLTQEDWNIVMYDGMRATSKWAGLYFILLMTIGNYILFNLLVAILVEGFASAPESTASLRPTPCPRTEYQLAVTEKPNPFCPPAGRTQVFVNSRTPSEKRADWSLYVFAPDNRFRMLNKELYQNKWFDRTVLLFILLNCVVMALEGPSVLPGSLERRVIDICMYVFLGIFTIEMMVKVIALGLWIGPYLRSGWNVMDGFLVVISWVDVIVTILGVLRVFRALRTLRPLRVISRAPGLKIVVETLISSLKPIGNIVLIAATFFIIFGILGVQLFKGKFHYCTDATVTTKTECLEVNREYNFDNLAKALLTLFVFSTKDGWVNIMYDGIDAVGIDKQPIRNHARYNVIYFVGFLLLAGFVVLNMLVGVVVENFQKCREMEKKIDEEKKNKKEMRREIAEDALSEYPRPRKFIHAVCTHGYFDLGIAAVIALNVLCMALQPDGLTAFLKCANYVFTAVFILEAILKIFALGIKRYIKDRWNQLDMLIVILSIVGIALEEELPINPTIIRVMRVLRIARVLKLLKTAEGIRKLLDTVAQALPQVGNLGMLFLLMFFIFSALGIELFGKIDCEKQGMDEHANFRSFGIAMLTLFRISTGDNWNGILKDTIPEIDCSEHVAPIYFAVFVLATQFVLLNVVVAVLMKHLE-EAK

>Cnida_nemvec_Cav2c

ALFCLKENNKLRVNCKKIVDSKFFETSILLIIAANCIVLILDTPLPKGDSTDLNKTLEQAEYVFVVIYCLESALKIIAQGFLFHEQAYLRNGWNILDFAVVVVGLVGMVWDLDGSLKVLRAVRVLRPLKIVSGIPSLQVVMKTIWRAMIPLLQILLLIIFVIVIYAIIGLELLKGKFHSTCYDANDRMEKGYLFPKICSNGRQCS-PGPNKGITNFDNIFLSMLTVFQCITMEGWTDIMYHSYDARDVVTSIIYISLIIIGSFFMLNLVLGVLSGEFARHQQQMERQLSGYMDWIARAEDIMRKRTSLSDSLAHLVEDQKMFLLLRRKVKVMVRSQIFYWAVLVCVFLNTVLMSFEHYGQPDWLERTQTIAEKVFLGIFIAEMLLKLYGLGATQYFKSSFNRFDFVVVLSGIVEMFLHISFGSSVLRSLRLLRIFKFTRFWSSLRNLVTSLLSSMRSILSLIFLLLLFIFIFALLGMQLFGGRFSTTLDAPRTNFDNFVKAMLAVFQIMTGEDWNTVMYNGIEAAVILGSLYFVALVIVGNYTLLNVFLAIAVDNLANAQALTRDEEQEVRMREQIKKRQNGRISRQDTQDNALTDPVTDREDEDIIRKSSMFIFGPDNPIRRLCHWVVNLRYFDTFILFIILISSVLLVFEDPV-STNSQTNTILGYCDYVITAIFGLEVLFKVIDLGVILHKYFRDAWNVIDAFVVACNIAALVLNVQEAIKSFRVLRVLRPVKAINKSKKLKTVFQCMVYSLKNVRFILLINLLFYYIFAVIGVQLFKGKFFYCTDMSKMQKSECKGTRWHFNFDDVPSAMLTLFSASTGEGWPTAMYHTVDATHVDRGPRRDNNIQMSIYMVCIVVIFSFFFLNIFVALIIVTFQEQGEKEMVGCELDRNQRDIQFAMTARPRQRYMPCFYKVWCVVDSKPFEIFIMTMIVLNAIVLMMATQEFNNIIEYVNMAFTFVFLFEAILKLIAFKL-NYFRDYWNVFDFIIVVTTLVGVLLELTKNQLDIDPSFFRLFRAARLVKLLRQGYTIRILLWTFLQSFKALPYVVMLIAMLFFVYAVIGMQLFGRIAK--REINIHNNFQSFFQALLVLFRAATGENWHLVMLACFGGKACGTAASIIYFITFYFFCSFLMLNLFVAVIMDNFEYLTR

>Cnida_nemvec_Cav2b

----------------------------------------MNKPLPMDDKIEIAKDLEHAEVYFVAIFCIEATLKILALGFVLHPGSYLRNGWNILDFVVVVIGVISLPNVDIKDIKSLRAVRVLRPVKLISGVPSLQVVMKSIGRAMVPLLQIALLVLFVIVIYAIIGLEFLMKQFHTTCFRNSSGVMEMTPGPSPCDKGRQCS-VGPNNGISTFDNIALSMLTVFQCITMEGWTGIMYSTFEAIDYLYSVYFVSLIVIGSFFMLNLVLGVLSGEFARRQQQLDRQVAGYLDWITKADEILRESISTVTADTESQAQVISQDRLKLLIRKSVKSQWFYWTVLICVFLNTISLATEHYNQPIWLDEFQDKAEKVFLAIFTLEMVLKMYSLGFDVYFSSSFNVFDCVVVCSGLVEAVQKINLGLSVLRCVRLLRVFKVTRHWKSLRNLATSLVSSIKSIMSLIFLLFLFILISALLGMQIFGGKFIGETKNPRTNFDNFPNAMLTVFQILTGEDWNSVMSYGIMAYGLVVSLYFVLLVIVGNYTLLNVFLAIAVDNLANAQILTQDEEQEEELKERKKSEGSTRKRRVRERRGGRRRTTDKKDSKQVIRTRTLFIFGPDNGFRKLCHSIVNLAHFDTAMLVVIGLSSLTIAAEDPL-RDDAPRNNILWYFDCVFTAIFAFEVVVKVVDLGLILHKYLRNTWNIIDAIVVICNIASLVLSAASWIKALRVVRVLRPFKSVHKIKKLQAVFRCMWFSVKNVANILMITGLFLLIFAVIGVQLFNGKFWKCTDEAKLYEKECQGEKIKLNFDNVGEAMLTLYTSSTGEGWPTAMHRTMDTTEKDKGPIQDYSTEYAIFYVSFVVVFSFFFLNIFVALIILTFQDLGEKEISNCELDRNQRDIHFALSAKPAQFYMPFQYKVWLVASSRPLDIFIMVLIALNSVVLMMQSDQYEKACQYLNIAFTSMFTLEAAIKITALRL-NYFRDYWNLFDFFIVLGGLLDMAFTIGKSYMPIDPSMFRLFRAARLIKLLRQGYTIRILLWTFLRSFKALPYVTLLIMLQFFMYAVIGMQLFGKIALDDTEINSHNNFRDIFQALQVLFRAATGEDWHLVMTACFISTGCGTPLAIIYFCSFIFLCMFLMLNLFVAVIMDNFEYLTR

>Cnida_nemvec_Cav2a

--------------------M-----------IAYCSLLSL-------------------------------------------PQ--------------LNVGINASSL------KALRAARVLRPLKLVSGIPSLQVVMKSIMCAMLPLLQICLLVGFVIIIYAIIGLEFLCGSFHYACHDNSTGVPTLPDEPTICGLGFACE-VGPNDGITSFDNIFAGCLTVFQVITNEGWTDIMYWTFNATDYYFWLYYYSLVIIGSFFMLNLVLGVLSGEFARRTKQMERQLNGYIDWISKAEDIMRKRNLDRVEDGDVAMTAQVLTRWRIRVRQIVKHQAFYWAVIICVILNTVITACQHYGQPDWFTQFQDTAEIIFITFFFTEMLFKLYGLGPQLYFKSQFNTFDCVVVCCGIIELVQGTELGISVLRALRILRLFKFTRYWSSLRNLVTSLLSSVRSIVSLLFLLFLFIVIFALLGMQLFGAQFKGRTGNPRTNFDDFWNAFLAVFQILTGEDWNAVMYDGVLSQGLGASLYFVMLVVLGNYVLLNVFLAIAVDNLANAQQLSADEEEDEKEREERKQQGRNGNARRMDDDMPDENEFEEVEEGNIINTWSLFLFPPGNPVRIGCHYVVNLRHFDNVILVIILISSVLLAAEDPV-VEDSYQNQILTYFDYVFTTIFAFEVIFKLIDYGAILHPYFRDAWNCIDALVVSCAIASLVLGSKKTVKVLRVLRVLRPLKAINKAKKLKAVFQCMLYSLRNVLNILIITVLFLIIFSVIGVQLFQGKFFSCNDRSKMTEAECKGALADYNFNTVYYAMLALFTSSTGEGWPALMQASIDTTKVDQGPVVDNKIEIFLFYIFFVIVFSFFFLNIFVALIILTFQEQGEKEQGDCELDRNQRDLHFAMVAKPSERFMPIQYRVWKIVDSRPFEYFIMTLIALNTLILMMEPTLYRYYLDLFNSIFTFMFTAEALLKLIAFRT-NYFRDNWNVFDFVVVLGSLLDFVLDKEGNEMPFDPSLFRLFRAARLIKLLRQGYTIRILLWTFLQSFKALPYVGMLIGLLFFIYAVIGMQLFGQIGKDSTAISGDNHFQSFFPAIQVLFRSATGENWHVIMLACTADETCGSNVAYVYFVSFIFFCSFLLLNLFVAVIMDNFEYLTR

>Cnida_nemvec_Cav1

ALFCLTLGNPIRSTAISLVEWRPFDVMILITIFANCAALAAFQPLPEQDSSLINEELEVAEFVFLGIFTMESVLKIIAYGFVMHPGAYLRNGWNILDFVIVVVGLATIIVKLYTDVKALRAFRVLRPLRLVSGVPSLQVVLNSIIKALIPLFHIALLVVFVVIIYAIIGVELFMGKLHSTCYD-NVTGQPTFDESHPCSTGYSCS-EGPNYGITNFDNIGLACLTVFQCITLEGWTDVMYSINDAIGSWPWLYFVTLIIWGSFFVLNLVLGVLSGEFAREKQLVDDAYHGYLDWISQAEDIERKPSFRRRKENDDISKNKENQRLRRSCRKAVKTQWFYWTVIVFVFLNSLTLALEHYNQPEFLTQFLDKANKLFLALFTLEMVVKMYCLGFHGYFASLFNRFDCLVVISSLLELGDQRPIGISMLRCVRLLRIFKVTRYWSSLSNLVASLLNSMRSIMGLLLLLSLFMVIFSLLGMQIFGGKFNGDEDVPRSNFDSFWRALVTVFQILTGEDWNAVMYTGIQSWGAIPILYFIFLVVVGNYILLNVFLAIAVDNLADAESLTEMEEEKKKEREEELEMLHSNGNSIPRALSSDAESPSLAEDSKMPPESSLFIFSNTNCFRVVCHRIATNSYFVNFILLLIIVSSCMLAAEDPL-NSNSKRNQVLNYFDYFFTAVFTIEITIKIIAYGVILHKFCRSAFNLLDFLVVAVSIVSIALRQISVVRILRVLRVLRPLRAINRAKGLKHVVQCVFVAVKTIGNIMLVTVLFNFLFAVIGVQLFKGTFFSCTDAEKITKRECQGQPQTFNFNDVPQAMLTLFTVMTFEGWPGILESSMDSTDVDEGPFLNNRPWVAIYYVIYIIIIAFFMINIFVGFVIVTFQNEGEEEFKDCELDKNQRKVEFALKARPTRRYIPLQFHVWRVVTSQPFEYLIFAFITGNTILLMMEPKLYTRVLDGFNIGFTSVFLLECILKLFAFKPKNYFLDRWNLFDFVVVVGSVVDITMNEVSSEQMFAFGFFRLFRALRLVKLLNQGSGIKTLLWTFIKSFQALPYVGLLIIMTFFIYAVVGMQMFGRIAIDPTQINRNNNFQTFPQSLMVLFRSATGENWQLIMLACTTDGLCGTDFAYAYFCSFYAICSFLIINLFVAVIMDNFDYLTR

>Chrysemys_picta_bellii_Cav2.3_XP_008164802.1

SLFLFGEDNMVRKYAKKLIDWPPFEYMILATIIANCIVLALEQHLPEDDKTPMSRRLEKTEPYFIGIFCFEAGIKIVALGFVFHKGSYLRNGWNVMDFIVVLSGILATAGTHFNDLRTLRAVRVLRPLKLVSGIPSLQIVLKSIMKAMVPLLQIGLLLFFAILMFAIIGLEFYSGKLHRACYTNNSGELEELDPPHPCG-VQGCP-IGPNDGITQFDNILFAVLTVFQCITMEGWTTVLYNTNDALGTWNWLYFIPLIIIGSFFVLNLVLGVLSGEFARRQQQIERELNGYRAWIDKAEEVMNRTEAVNRDSSDEHCVDISSVLLRISVRHMVKSQVFYWTVLSLVALNTACVAIVHHNQPPWLTHLLYYAEFLFLGLFLLEMSLKMYGMGPRLYFHSSFNCFDCGVTVGSIFEVVPGTSFGISVLRALRLLRIFKITKYWASLRNLVVSLMSSMKSIISLLFLLFLFIVVFALLGMQLFGGRFNFMDGTPSANFDTFPAAIMTVFQILTGEDWNEVMYNGIRSQGMWSSIYFIILTLFGNYTLLNVFLAIAVDNLANAQELTKDEQEEEEAFNQKHALQEMSPLKVPHELNEEKNTSLTEQDCSMVPHSSMFIFSTTNPVRRACHYIVNLRYFEMCILLVIAASSIALAAEDPV-LTNSDRNKVLRYFDYVFTGVFTFEMIIKMIDQGLILQDYFRDLWNILDFIVVVGALVAFALADIKTIKSLRVLRVLRPLKTIKRLPKLKAVFDCVVTSLKNVFNILIVYKLFMFIFAVIAVQLFKGKFFYCTDSSKDTEKDCIGKRHEFHYDNIIWALLTLFTVSTGEGWPQVLQHSVDVTEEDRGPSRSNRMEMSIFYVVYFVVFPFFFVNIFVALIIITFQEQGDKMMEECSLEKNERAIDFAISAKPLTRYMPFQYRVWHFVVSPSFEYTIMAMIALNTVVLMMAPYTYELALKYLNIAFTMVFSLECVLKIIAFGFLNYFRDTWNIFDFITVIGSITEIILTDLVNTSSFNMSFLKLFRAARLIKLLRQGYTIRILLWTFVQSFKALPYVCLLIAMLFFIYAIIGMQVFGNIKLDESHINRHNNFRSFLASLMLLFRSATGEAWQEIMLSCLENEHCGTDLAYVYFVSFIFFCSFLMLNLFVAVIMDNFEYLTR

>Chrysemys_picta_bellii_Cav2.2_XP_008172570.1

SLFIFSEDNVIRKYAKRITEWPPFEYMILATIIANCIVLALEQHLPEDDKTPMSERLDDTEPYFIGIFCFEAGIKIIALGFVFHKGSYLRNGWNVMDFVVVLTGILATAG----DLRTLRAVRVLRPLKLVSGIPSLQVVLKSIMKAMVPLLQIGLLLFFAIVMFAIIGLEFYMGKFHKTCFS----NETGEEVGFPCGEARQCE-PGPNYGITNFDNILFAVLTVFQCITMEGWTDILYNTNDAAGTWNWLYFIPLIIIGSFFMLNLVLGVLSGEFARRQQQIERELNGYLEWIFKAEEVMSKNDLIHAEEGEDHFTDICSVMFRFFIRRMVKAQSFYWIVLCVVTLNTLCVAMVHYDQPEGLTTALYFAEFVFLGLFLTEMSLKMYGLGPRNYFHSSFNCFDFGVIVGSIFEVIPGTSFGISVLRALRLLRIFKVTKYWNSLRNLVVSLLNSMKSIISLLFLLFLFIVVFALLGMQLFGGQFNFQDETPTTNFDTFPAAILTVFQILTGEDWNAVMYHGIVSQGMFSSIYFIVLTLFGNYTLLNVFLAIAVDNLANAQELTKDEEEMEEATNQKLALSGNREGDPGLKGENGEEPHRRYKIRHILPYSSMFILSPTNPIRRLFHYIVNMRYFEMVILIVIALSSIALAAEDPV-QAESPRNDALKYLDYIFTGVFTFEMVIKMIDLGLLLHPYFRDLWNILDFIVVSGALVAFAFSDINTIKSLRVLRVLRPLKTIKRLPKLKAVFDCVVNSLKNVLNILIVYMLFMFIFAVIAVQLFKGKFFYCTDESKELEKDCRGKKYEFHYDNVLWALLTLFTVSTGEGWPTVLKHSVDATYEEQGPSPGYRMEMSIFYVVYFVVFPFFFVNIFVALIIITFQEQGDKVMSECSLEKNERAIDFAISARPLXRYMPFQYKMWKFVVSPPFEYFIMVMIALNTIVLMMAPDAYEEMLKCLNIVFTSMFSMECVLKIIAFGVLNYFRDAWNVFDFVTVLGSITDILVTEAETDNFINLSFLRLFRAARLIKLLRQGYTIRILLWTFVQSFKALPYVCLLIAMLFFIYAIIGMQVFGNIALDDTSINRHNNFRTFLQALMLLFRSATGEAWHEIMLSCLTKNECGSDFAYFYFVSFIFLCSFLMLNLFVAVIMDNFEYLTR

>Chrysemys_picta_bellii_Cav2.1_XP_008161515.1

SLFLFSEDNVVRKYAKKITEWPPFEYMILATIIANCIVLALEQHLPDEDKTPMSERLDDTEPYFIGIFCFEAGIKIIALGFAFHKGSYLRNGWNVMDFVVVLTGILAKVG----DLRTLRAVRVLRPLKLVSGIPSLQVVLKSIMKAMIPLLQIGLLLFFAILIFAIIGLEFYMGKFHTTCFD----SVTGEIKVVPCGTARMCP-EGPNYGITQFDNILFAVLTVFQCITMEGWTELLYYSNDASGTWNWLYFIPLIIIGSFFMLNLVLGVLSGEFARRQQQIERELNGYMEWISKAEEVISKTDLLNPEEADDQLADISSVRMRFYIRRMVKTQAFYWTVLSLVALNTLCVAIVHYSQPDWLSDFLYYAEFIFLGLFMSEMFIKMYGLGTRPYFHSSFNCFDCAVIIGSIFEVIPGTSFGISVLRALRLLRIFKITKYWASLRNLVVSLLNSMKSIISLLFLLFLFIVVFALLGMQLFGGQFNFDDGTPSTNFDTFPAAIMTVFQILTGEDWNMVMYDGIKSQGMVYSVYFIVLTLFGNYTLLNVFLAIAVDNLANAQELTKDEQEEEEVANQKLALSKRDDRERRHRRRKENQGPSPVPGPNMVPYSSMFILSTTNPFRRLCHYIVNLRYFEMCILMVIAMSSIALAAEDPV-QPNATRNNVLRYFDYVFTGVFTFEMVIKMVDLGLVLHQYFRDLWNILDFIVVSGALVAFAFTDINTIKSLRVLRVLRPLKTIKRLPKLKAVFDCVVNSLKNVLNILIVYMLFMFIFAVVAVQLFKGKFFYCTDESKEFEKDCRGKKYEFHYDNVLWALLTLFTVSTGEGWPQVLKHSVDATYENQGPSPGYRMEMSIFYVVYFVVFPFFFVNIFVALIIITFQEQGDKMMEEYSLEKNERAIDFAISAKPLTRHMPFQYRMWQFVVSPPFEYTIMAMIALNTIVLMMASTAYEDVLKMFNHVFTSLFSLECLLKIMAFGVLNYFRDAWNIFDFVTVLGSITDILVTE--GNNFINLSFLRLFRAARLIKLLRQGYTIRILLWTFVQSFKALPYVCLLIAMLFFIYAIIGMQVFGNIGIEESAITEHNNFRTFFQALMLLFRSATGEAWHEIMLACLKEHECGNEFAYFYFVSFIFLCSFLMLNLFVAVIMDNFEYLTR

>Chrysemys_picta_bellii_Cav1.3_XP_023956012.1

ALFCLSLNNPIRRACISLVEWKPFDIFILLAIFANCVALAVYIPFPEDDSNSTNHNLEKVEYAFLIIFTIETFLKIIAYGLLLHPNAYVRNGWNLLDFVIVVVGLFSVILEQLTDVKALRAFRVLRPLRLVSGVPSLQVVLNSIIKAMVPLLHIALLVLFVIIIYAIIGLELFIGKMHKSCFL-VDSDILVEDDPAPCAFGRQCA-VGPNGGITNFDNFAFAMLTVFQCITMEGWTDVLYWVNDAIGEWPWIYFVSLIILGSFFVLNLVLGVLSGEFSREKQQLEEDLKGYLDWITQAEDIDSETESVNTENVSGEGESPACCFNRRKCRAAVKSVSFYWLVIVLVFLNTLTISSEHYNQPDWLTQVQDTANKVLLALFTCEMLIKMYSLGLQAYFVSLFNRFDCFVVCGGIVETIIMSPLGISVFRCVRLLRIFKVTRHWTSLSNLVASLLNSMKSIASLLLLLFLFIIIFSLLGMQLFGGKFNDETQTKRSTFDNFPQALLTVFQILTGEDWNAVMYDGIMAYGMVVCIYFIILFICGNYILLNVFLAIAVDNLADAESLNTAQKEEAEEKQRKKNANKVTIAEYREGEDEDKDPYPPCDVPVIPEGSAFFIFSSTNPIRVGCHRLINHHIFTNLILVFIMLSSVSLAAEDPI-RSHSFRNNILGYFDYAFTAIFTVEILLKMTAFGAFLHKFCRNYFNLLDLLVVGVSLVSFGIQAISVVKILRVLRVLRPLRAINRAKGLKHVVQCVFVAIRTIGNIMIVTTLLQFMFACIGVQLFKGKFYRCTDEAKQNPEDCRGQNSDFNFDNVLSAMMALFTVSTFEGWPALLYKAIDSNAENIGPVYNYRVEISIFFIIYIIIIAFFMMNIFVGFVIVTFQEQGEQEYKNCELDKNQRQVEYALKARPLRRYIPYQYKFWYMVNSTGFEYIMFVLIMLNTLCLAMQSKLFNDAMDILNMVFTGVFTVEMVLKLIAFKPKGYFSDAWNAFDSLIVIGSIVDVVLSESEDSARISITFFRLFRVMRLVKLLSRGEGIRTLLWTFIKSFQALPYVALLIAMLFFIYAVIGMQVFGKVAMRDNQINRNNNFQTFPQAVLLLFRCATGEAWQEIMLACLEEFTCGSNFSIIYFISFYMLCAFLIINLFVAVIMDNFDYLTR

>Chrysemys_picta_bellii_Cav1.2_XP_008162493.1

ALLCLTLKNPIRRACISIVEWKPFEIIILLTIFANCVALAIYIPFPEDDSNATNSNLERVEYLFLIIFTVEAFLKVIAYGLLFHPNAYLRNGWNLLDFIIVVVGLFSAILEQATDVKALRAFRVLRPLRLVSGVPSLQVVLNSIIKAMVPLLHIALLVLFVIIIYAIIGLELFMGKMHKTCYF-LQGGLPAEEEASPCAPGRQCQ-EGPKHGITNFDNFAFAMLTVFQCITMEGWTDVLYWVNDAIGDWPWIYFVTLIIIGSFFVLNLVLGVLSGEFSREKQQLEEDLKGYLDWITQAEDIDSETESVNTDNVAGADIEGENCFCRRKCRAAVKSNIFYWLVIFLVFLNTLTIASEHYNQPYWLTEVQDTANKVLLALFTAEMLLKMYSLGLQAYFVSLFNRFDCFIVCGGILETIIMSPLGISVLRCVRLLRIFKITRYWNSLSNLVASLLNSVRSIASLLLLLFLFIIIFSLLGMQLFGGKFNDEMQTKRSTFDNFPQSLLTVFQILTGEDWNSVMYDGIMAYGMLVCIYFIILFICGNYILLNVFLAIAVDNLADAESLTSAQKEEEEEKERKKLATKINVDDYQPNENEEKNPYPTTEAPAMPDASAFFIFSPTNRYLRSCYAAEHYSFRNQILFYFDIVFTVIFTIEIAL--------KILGNADYVFTSIFTLEIILKMTAYGAFLHKFCRNYFNILDLLVVSVSLISFGIQAINVVKILRVLRVLRPLRAINRAKGLKHVVQCVFVAIRTIGNIVIVTTLLQFMFACIGVQLFKGKLYSCSDSSKQTEAECRGENSKFNFDNVLTAMMALFTVSTFEGWPELLYKSIDSHMEDVGPIYNHRVEISIFFIIYIIIIAFFMMNIFVGFVIVTFQEQGEQEYKNCELDKNQRQVEYALKARPLRRYIPYQYKVWYVVNSTYFEYLMFVLILLNTICLAMQSCPFKQAMNILNMLFTGLFTVEMILKLIAFKPKGYFSDPWNVFDFLIVIGSIIDVILSEAEENSRISITFFRLFRVMRLVKLLSRGEGIRTLLWTFIKSFQALPYVALLIVMLFFIYAVIGMQVFGKIALNDSYINRNNNFQTFPQAVLLLFRCATGEGWQDIMLDCLGDHSCGSSFSIFYFISFYMLCAFLIINLFVAVIMDNFDYLTR

>Chord_homsap_Cav3.3_NP_066919.2

AFFCLRQTTSPRNWCIKMVCNPWFECVSMLVILLNCVTLGMYQPCDDMDLSDRCKILQVFDDFIFIFFAMEMVLKMVALGI-FGKKCYLGDTWNRLDFFIVMAGMVEYSLDLQNNLSAIRTVRVLRPLKAINRVPSMRILVNLLLDTLPMLGNVLLLCFFVFFIFGIIGVQLWAGLLRNRCFLPPYYQPEEDDEMFICSLGRECCSANPHKGAINFDNIGYAWIVIFQVITLEGWVEIMYYVMDAHSFYNFIYFILLIIVGSFFMINLCLVVIATQFSAEPGDCYEEIFQYVCHILRKAKRRLCPQHSPLDATPHTLVQPIPAETRAKLRGIVDSKYFNRGIMMAILVNTVSMGIEHHEQPEELTNILEICNVVFTSMFALEMILKLAAFGLFDYLRNPYNIFDSIIVIISIWEIVGQADGGLSVLRTFRLLRVLKLVRFMPALRRQLVVLMKTMDNVATFCMLLMLFIFIFSILGMHIFGCKFSGDTVPDRKNFDSLLWAIVTVFQILTQEDWNVVLYNGMASTSPWASLYFVALMTFGNYVLFNLLVAILVEGFQAGDSYSDEDQSSSNIEEFDKLQLVPAVGAHPRAAWRAAGPAPGHEDCNVREDWSVYLFSPENRFRVLCQTIIAHKLFDYVVLAFIFLNCITIALERPQIEAGSTERIFLTVSNYIFTAIFVGEMTLKVVSLGLYFGEYLRSSWNVLDGFLVFVSIIDIVVSILGVLRVLRLLRTLRPLRVISRAPGLKLVVETLISSLKPIGNIVLICCAFFIIFGILGVQLFKGKFYHCLDTRITNRSDCMAVHHKYNFDNLGQALMSLFVLASKDGWVNIMYNGLDAVAVDQQPVTNHNPWMLLYFISFLLIVSFFVLNMFVGVVVENFHKCRQHQEAEEARRREEKRRRLEKKRRPYYAYCHTRLLIHSMCTSHYLDIFITFIICLNVVTMSLQPTSLETALKYCNYMFTTVFVLEAVLKLVAFGLRRFFKDRWNQLDLAIVLLSVMGITLEEALPINPTIIRIMRVLRIARVLKLLKMATGMRALLDTVVQALPQVGNLGLLFMLLFFIYAALGVELFGKLVCNDEGMSRHATFENFGMAFLTLFQVSTGDNWNGIMKDTLDERSCLQFVSPLYFVSFVLTAQFVLINVVVAVLMKHLDDSNK

>Chord_homsap_Cav3.2_NP_066921.2

VFFCLGQTTRPRSWCLRLVCNPWFEHVSMLVIMLNCVTLGMFRPCEDVEGSERCNILEAFDAFIFAFFAVEMVIKMVALGL-FGQKCYLGDTWNRLDFFIVVAGMMEYSL----SLSAIRTVRVLRPLRAINRVPSMRILVTLLLDTLPMLGNVLLLCFFVFFIFGIVGVQLWAGLLRNRCFLRPYYQTEEGEENFICSSRMPCTDSNPHNGAINFDNIGYAWIAIFQVITLEGWVDIMYYVMDAHSFYNFIYFILLIIVGSFFMINLCLVVIATQFSSEPGSCYEELLKYVGHIFRKVKRRQAPGHLSGLSVPCPLPSPPAGTFSGKLRRIVDSKYFSRGIMMAILVNTLSMGVEYHEQPEELTNALEISNIVFTSMFALEMLLKLLACGPLGYIRNPYNIFDGIIVVISVWEIVGQADGGLSVLRTFRLLRVLKLVRFLPALRRQLVVLVKTMDNVATFCTLLMLFIFIFSILGMHLFGCKFSGDTVPDRKNFDSLLWAIVTVFQILTQEDWNVVLYNGMASTSSWAALYFVALMTFGNYVLFNLLVAILVEGFQAGDSDTDEDKHFEEDFHKLRELLRRAESLDPRPLRPAALPPTKCRDRDSREAWALYLFSPQNRFRVSCQKVITHKMFDHVVLVFIFLNCVTIALERPDIDPGSTERVFLSVSNYIFTAIFVAEMMVKVVALGLLSGEYLQSSWNLLDGLLVLVSLVDIVVAILGVLRVLRLLRTLRPLRVISRAPGLKLVVETLISSLRPIGNIVLICCAFFIIFGILGVQLFKGKFYYCEDTRISTKAQCRAVRRKYNFDNLGQALMSLFVLSSKDGWVNIMYDGLDAVGVDQQPVQNHNPWMLLYFISFLLIVSFFVLNMFVGVVVENFHKCRQHQEAEEARRREEKRRRLERRRRPYYAYSPTRRSIHSLCTSHYLDLFITFIICVNVITMSMQPKSLDEALKYCNYVFTIVFVFEAALKLVAFGFRRFFKDRWNQLDLAIVLLSLMGITLEEALPINPTIIRIMRVLRIARVLKLLKMATGMRALLDTVVQALPQVGNLGLLFMLLFFIYAALGVELFGRLECSEEGLSRHATFSNFGMAFLTLFRVSTGDNWNGIMKDTLEDKHCLPALSPVYFVTFVLVAQFVLVNVVVAVLMKHLEESNK

>Chord_homsap_Cav3.1_NP_061496.2

VFFYLSQDSRPRSWCLRTVCNPWFERISMLVILLNCVTLGMFRPCEDIADSQRCRILQAFDDFIFAFFAVEMVVKMVALGI-FGKKCYLGDTWNRLDFFIVIAGMLEYSL--DLSFSAVRTVRVLRPLRAINRVPSMRILVTLLLDTLPMLGNVLLLCFFVFFIFGIVGVQLWAGLLRNRCFLERYYQTENEDESFICSQGPPCGEHNPFKGAINFDNIGYAWIAIFQVITLEGWVDIMYFVMDAHSFYNFIYFILLIIVGSFFMINLCLVVIATQFSSEPGSCYEELLKYLVYILRKAARRSSMHKLLETQSTGACQSSCKILICDTFRKIVDSKYFGRGIMIAILVNTLSMGIEYHEQPEELTNALEISNIVFTSLFALEMLLKLLVYGPFGYIKNPYNIFDGVIVVISVWEIVGQQGGGLSVLRTFRLMRVLKLVRFLPALQRQLVVLMKTMDNVATFCMLLMLFIFIFSILGMHLFGCKFAGDTLPDRKNFDSLLWAIVTVFQILTQEDWNKVLYNGMASTSSWAALYFIALMTFGNYVLFNLLVAILVEGFQAGDSESEPDFSLDGDGDRKKCLPDTLQVPGLHRTASGRGSASEHQDCNERDSWSAYIFPPQSRFRLLCHRIITHKMFDHVVLVIIFLNCITIAMERPKIDPHSAERIFLTLSNYIFTAVFLAEMTVKVVALGWCFGEYLRSSWNVLDGLLVLISVIDILVSILGMLRVLRLLRTLRPLRVISRAQGLKLVVETLMSSLKPIGNIVVICCAFFIIFGILGVQLFKGKFFVCQDTRITNKSDCAEVRHKYNFDNLGQALMSLFVLASKDGWVDIMYDGLDAVGVDQQPIMNHNPWMLLYFISFLLIVAFFVLNMFVGVVVENFHKCRQHQEEEEARRREEKRRRLEKKRRPYYSYSRFRLLVHHLCTSHYLDLFITGVIGLNVVTMAMQPQILDEALKICNYIFTVIFVLESVFKLVAFGFRRFFQDRWNQLDLAIVLLSIMGITLEESLPINPTIIRIMRVLRIARVLKLLKMAVGMRALLDTVMQALPQVGNLGLLFMLLFFIFAALGVELFGDLECDEEGLGRHATFRNFGMAFLTLFRVSTGDNWNGIMKDTLQESTCYTVISPIYFVSFVLTAQFVLVNVVIAVLMKHLEESNK

>Chord_homsap_Cav2.3_NP_001192222.1

SLFIFGEDNIVRKYAKKLIDWPPFEYMILATIIANCIVLALEQHLPEDDKTPMSRRLEKTEPYFIGIFCFEAGIKIVALGFIFHKGSYLRNGWNVMDFIVVLSGILATAGTHFNDLRTLRAVRVLRPLKLVSGIPSLQIVLKSIMKAMVPLLQIGLLLFFAILMFAIIGLEFYSGKLHRACFMNNSGILEGFDPPHPCG-VQGCP-IGPNDGITQFDNILFAVLTVFQCITMEGWTTVLYNTNDALGTWNWLYFIPLIIIGSFFVLNLVLGVLSGEFARRQQQIERELNGYRAWIDKAEEVMSRTEAMTRDSSDEHCVDISSVLLRISIRHMVKSQVFYWIVLSLVALNTACVAIVHHNQPQWLTHLLYYAEFLFLGLFLLEMSLKMYGMGPRLYFHSSFNCFDFGVTVGSIFEVVPGTSFGISVLRALRLLRIFKITKYWASLRNLVVSLMSSMKSIISLLFLLFLFIVVFALLGMQLFGGRFNFNDGTPSANFDTFPAAIMTVFQILTGEDWNEVMYNGIRSQGMWSAIYFIVLTLFGNYTLLNVFLAIAVDNLANAQELTKDEQEEEEAFNQKHALADTPLVLPHPELEVGKHVVLTEQEPEMVPHSSMFIFSTTNPIRRACHYIVNLRYFEMCILLVIAASSIALAAEDPV-LTNSERNKVLRYFDYVFTGVFTFEMVIKMIDQGLILQDYFRDLWNILDFVVVVGALVAFALADIKTIKSLRVLRVLRPLKTIKRLPKLKAVFDCVVTSLKNVFNILIVYKLFMFIFAVIAVQLFKGKFFYCTDSSKDTEKECIGKRHEFHYDNIIWALLTLFTVSTGEGWPQVLQHSVDVTEEDRGPSRSNRMEMSIFYVVYFVVFPFFFVNIFVALIIITFQEQGDKMMEECSLEKNERAIDFAISAKPLTRYMPFQYRVWHFVVSPSFEYTIMAMIALNTVVLMMAPCTYELALKYLNIAFTMVFSLECVLKVIAFGFLNYFRDTWNIFDFITVIGSITEIILTDLVNTSGFNMSFLKLFRAARLIKLLRQGYTIRILLWTFVQSFKALPYVCLLIAMLFFIYAIIGMQVFGNIKLDESHINRHNNFRSFFGSLMLLFRSATGEAWQEIMLSCLENERCGTDLAYVYFVSFIFFCSFLMLNLFVAVIMDNFEYLTR

>Chord_homsap_Cav2.2_NP_000709.1

SLFVFSEDNVVRKYAKRITEWPPFEYMILATIIANCIVLALEQHLPDGDKTPMSERLDDTEPYFIGIFCFEAGIKIIALGFVFHKGSYLRNGWNVMDFVVVLTGILATAG----DLRTLRAVRVLRPLKLVSGIPSLQVVLKSIMKAMVPLLQIGLLLFFAILMFAIIGLEFYMGKFHKACFP---NSTDAEPVGFPCGKARLCE-PGPNFGITNFDNILFAILTVFQCITMEGWTDILYNTNDAAGTWNWLYFIPLIIIGSFFMLNLVLGVLSGEFARRQQQIERELNGYLEWIFKAEEVMSRNDLIHAEEGEDRFADLCAVMFRFFIRRMVKAQSFYWVVLCVVALNTLCVAMVHYNQPRRLTTTLYFAEFVFLGLFLTEMSLKMYGLGPRSYFRSSFNCFDFGVIVGSVFEVVPGSSFGISVLRALRLLRIFKVTKYWSSLRNLVVSLLNSMKSIISLLFLLFLFIVVFALLGMQLFGGQFNFQDETPTTNFDTFPAAILTVFQILTGEDWNAVMYHGIESQGMFSSFYFIVLTLFGNYTLLNVFLAIAVDNLANAQELTKDEEEMEEAANQKLALEKEAEIVEADKEKELRNHQPREPHCDIVPYSSMFCLSPTNLLRRFCHYIVTMRYFEVVILVVIALSSIALAAEDPV-RTDSPRNNALKYLDYIFTGVFTFEMVIKMIDLGLLLHPYFRDLWNILDFIVVSGALVAFAFSDINTIKSLRVLRVLRPLKTIKRLPKLKAVFDCVVNSLKNVLNILIVYMLFMFIFAVIAVQLFKGKFFYCTDESKELERDCRGKKYDFHYDNVLWALLTLFTVSTGEGWPMVLKHSVDATYEEQGPSPGYRMELSIFYVVYFVVFPFFFVNIFVALIIITFQEQGDKVMSECSLEKNERAIDFAISAKPLTRYMPFQYKTWTFVVSPPFEYFIMAMIALNTVVLMMAPYEYELMLKCLNIVFTSMFSMECVLKIIAFGVLNYFRDAWNVFDFVTVLGSITDILVTEAETNNFINLSFLRLFRAARLIKLLRQGYTIRILLWTFVQSFKALPYVCLLIAMLFFIYAIIGMQVFGNIALDDTSINRHNNFRTFLQALMLLFRSATGEAWHEIMLSCLNATECGSDFAYFYFVSFIFLCSFLMLNLFVAVIMDNFEYLTR

>Chord_homsap_Cav2.1_O00555

SLFLFSEDNVVRKYAKKITEWPPFEYMILATIIANCIVLALEQHLPDDDKTPMSERLDDTEPYFIGIFCFEAGIKIIALGFAFHKGSYLRNGWNVMDFVVVLTGILATVG----DLRTLRAVRVLRPLKLVSGIPSLQVVLKSIMKAMIPLLQIGLLLFFAILIFAIIGLEFYMGKFHTTCFE-EGTDDIQGESPAPCGTARTCP-EGPNNGITQFDNILFAVLTVFQCITMEGWTDLLYNSNDASGTWNWLYFIPLIIIGSFFMLNLVLGVLSGEFARRQQQIERELNGYMEWISKAEEVISKTDLLNPEEAEDQLADIASVRMRFYIRRMVKTQAFYWTVLSLVALNTLCVAIVHYNQPEWLSDFLYYAEFIFLGLFMSEMFIKMYGLGTRPYFHSSFNCFDCGVIIGSIFEVIPGTSFGISVLRALRLLRIFKVTKYWASLRNLVVSLLNSMKSIISLLFLLFLFIVVFALLGMQLFGGQFNFDEGTPPTNFDTFPAAIMTVFQILTGEDWNEVMYDGIKSQGMVFSIYFIVLTLFGNYTLLNVFLAIAVDNLANAQELTKDEQEEEEAANQKLALLGRQDPPLAEDIDNMKNNKLATAESAMPPYSSMFILSTTNPLRRLCHYILNLRYFEMCILMVIAMSSIALAAEDPV-QPNAPRNNVLRYFDYVFTGVFTFEMVIKMIDLGLVLHQYFRDLWNILDFIVVSGALVAFAFTDINTIKSLRVLRVLRPLKTIKRLPKLKAVFDCVVNSLKNVFNILIVYMLFMFIFAVVAVQLFKGKFFHCTDESKEFEKDCRGKKYEFHYDNVLWALLTLFTVSTGEGWPQVLKHSVDATFENQGPSPGYRMEMSIFYVVYFVVFPFFFVNIFVALIIITFQEQGDKMMEEYSLEKNERAIDFAISAKPLTRHMPFQYRMWQFVVSPPFEYTIMAMIALNTIVLMMASVAYENALRVFNIVFTSLFSLECVLKVMAFGILNYFRDAWNIFDFVTVLGSITDILVTE--GNNFINLSFLRLFRAARLIKLLRQGYTIRILLWTFVQSFKALPYVCLLIAMLFFIYAIIGMQVFGNIGIDVFQITEHNNFRTFFQALMLLFRSATGEAWHNIMLSCLLTRECGNEFAYFYFVSFIFLCSFLMLNLFVAVIMDNFEYLTR

>Chord_homsap_Cav1.4_NP_005174.2

ALFCLTLANPLRRSCISIVEWKPFDILILLTIFANCVALGVYIPFPEDDSNTANHNLEQVEYVFLVIFTVETVLKIVAYGLVLHPSAYIRNGWNLLDFIIVVVGLFSVLLEQGPDVKALRAFRVLRPLRLVSGVPSLHIVLNSIMKALVPLLHIALLVLFVIIIYAIIGLELFLGRMHKTCYF-LGSDMEAEEDPSPCASGRACT-PGPNGGITNFDNFFFAMLTVFQCVTMEGWTDVLYWMQDAMGELPWVYFVSLVIFGSFFVLNLVLGVLSGEFSREKQQMEEDLRGYLDWITQAEELDTRSTHSTSSHASLPASDTGSMVLRARCRRAVKSNACYWAVLLLVFLNTLTIASEHHGQPVWLTQIQEYANKVLLCLFTVEMLLKLYGLGPSAYVSSFFNRFDCFVVCGGILETTAMQPLGISVLRCVRLLRIFKVTRHWASLSNLVASLLNSMKSIASLLLLLFLFIIIFSLLGMQLFGGKFNDQTHTKRSTFDTFPQALLTVFQILTGEDWNVVMYDGIMAYGMLVCIYFIILFICGNYILLNVFLAIAVDNLASGD-------GTAKDKGGEKSNGLVPGVEKEEEEGARREGADMEEEEEIPEGSAFFCLSQTNPLRKGCHTLIHHHVFTNLILVFIILSSVSLAAEDPI-RAHSFRNHILGYFDYAFTSIFTVEILLKMTVFGAFLHRFCRSWFNMLDLLVVSVSLISFGIHAISVVKILRVLRVLRPLRAINRAKGLKHVVQCVFVAIRTIGNIMIVTTLLQFMFACIGVQLFKGKFYTCTDEAKHTPQECKGVNSDFNFDNVLSAMMALFTVSTFEGWPALLYKAIDAYAEDHGPIYNYRVEISVFFIVYIIIIAFFMMNIFVGFVIITFRAQGEQEYQNCELDKNQRQVEYALKAQPLRRYIPHQYRVWATVNSAAFEYLMFLLILLNTVALAMQTAPFNYAMDILNMVFTGLFTIEMVLKIIAFKPKHYFTDAWNTFDALIVVGSIVDIAVTESEDSSRISITFFRLFRVMRLVKLLSKGEGIRTLLWTFIKSFQALPYVALLIAMIFFIYAVIGMQMFGKVALQDTQINRNNNFQTFPQAVLLLFRCATGEAWQEIMLASLEEFTCGSNFAIAYFISFFMLCAFLIINLFVAVIMDNFDYLTR

>Chord_homsap_Cav1.3_NP_001122312.1

ALFCLSLNNPIRRACISIVEWKPFDIFILLAIFANCVALAIYIPFPEDDSNSTNHNLEKVEYAFLIIFTVETFLKIIAYGLLLHPNAYVRNGWNLLDFVIVIVGLFSVILEQLTDVKALRAFRVLRPLRLVSGVPSLQVVLNSIIKAMVPLLHIALLVLFVIIIYAIIGLELFIGKMHKTCFF-ADSDIVAEEDPAPCAFGRQCT-VGPNGGITNFDNFAFAMLTVFQCITMEGWTDVLYWMNDAMGELPWVYFVSLVIFGSFFVLNLVLGVLSGEFSREKQQLEEDLKGYLDWITQAEDIDSETESVNTENVSGEGENRGCCFNRRRCRAAVKSVTFYWLVIVLVFLNTLTISSEHYNQPDWLTQIQDIANKVLLALFTCEMLVKMYSLGLQAYFVSLFNRFDCFVVCGGITETIIMSPLGISVFRCVRLLRIFKVTRHWTSLSNLVASLLNSMKSIASLLLLLFLFIIIFSLLGMQLFGGKFNDETQTKRSTFDNFPQALLTVFQILTGEDWNAVMYDGIMAYGMIVCIYFIILFICGNYILLNVFLAIAVDNLADAESLNTAQKEEAEEKERKKIADNKVTIDDYREEDEDKDPYPPCDVPVIPEGSAFFILSKTNPIRVGCHKLINHHIFTNLILVFIMLSSAALAAEDPI-RSHSFRNTILGYFDYAFTAIFTVEILLKMTTFGAFLHKFCRNYFNLLDMLVVGVSLVSFGIQAISVVKILRVLRVLRPLRAINRAKGLKHVVQCVFVAIRTIGNIMIVTTLLQFMFACIGVQLFKGKFYRCTDEAKSNPEECRGQNSDFNFDNVLSAMMALFTVSTFEGWPALLYKAIDSNGENIGPIYNHRVEISIFFIIYIIIVAFFMMNIFVGFVIVTFQEQGEKEYKNCELDKNQRQVEYALKARPLRRYIPYQYKFWYVVNSSPFEYMMFVLIMLNTLCLAMQSKMFNDAMDILNMVFTGVFTVEMVLKVIAFKPKGYFSDAWNTFDSLIVIGSIIDVALSESEESNRISITFFRLFRVMRLVKLLSRGEGIRTLLWTFIKSFQALPYVALLIAMLFFIYAVIGMQMFGKVAMRDNQINRNNNFQTFPQAVLLLFRCATGEAWQEIMLACLEEYTCGSNFAIVYFISFYMLCAFLIINLFVAVIMDNFDYLTR

>Chord_homsap_Cav1.2_Q13936

ALLCLTLKNPIRRACISIVEWKPFEIIILLTIFANCVALAIYIPFPEDDSNATNSNLERVEYLFLIIFTVEAFLKVIAYGLLFHPNAYLRNGWNLLDFIIVVVGLFSAILEQATDVKALRAFRVLRPLRLVSGVPSLQVVLNSIIKAMVPLLHIALLVLFVIIIYAIIGLELFMGKMHKTCYNEGIADVPAEDDPSPCALGRQCQ-DGPKHGITNFDNFAFAMLTVFQCITMEGWTDVLYWVNDAVGDWPWIYFVTLIIIGSFFVLNLVLGVLSGEFSREKQQLEEDLKGYLDWITQAEDIDSETESVNTENVAGGDIEGENCFCRRKCRAAVKSNVFYWLVIFLVFLNTLTIASEHYNQPNWLTEVQDTANKALLALFTAEMLLKMYSLGLQAYFVSLFNRFDCFVVCGGILETIIMSPLGISVLRCVRLLRIFKITRYWNSLSNLVASLLNSVRSIASLLLLLFLFIIIFSLLGMQLFGGKFNDEMQTRRSTFDNFPQSLLTVFQILTGEDWNSVMYDGIMAYGMLVCIYFIILFICGNYILLNVFLAIAVDNLADAESLTSAQKEEEEEKERKKLATKINMDDLQPNENEDKSPYPNPETTGMPEASAFFIFSSNNRFRLQCHRIVNDTIFTNLILFFILLSSISLAAEDPV-QHTSFRNHILFYFDIVFTTIFTIEIALKMTAYGAFLHKFCRNYFNILDLLVVSVSLISFGIQAINVVKILRVLRVLRPLRAINRAKGLKHVVQCVFVAIRTIGNIVIVTTLLQFMFACIGVQLFKGKLYTCSDSSKQTEAECKGENSKFDFDNVLAAMMALFTVSTFEGWPELLYRSIDSHTEDKGPIYNYRVEISIFFIIYIIIIAFFMMNIFVGFVIVTFQEQGEQEYKNCELDKNQRQVEYALKARPLRRYIPHQYKVWYVVNSTYFEYLMFVLILLNTICLAMQSCLFKIAMNILNMLFTGLFTVEMILKLIAFKPKGYFSDPWNVFDFLIVIGSIIDVILSEAEENSRISITFFRLFRVMRLVKLLSRGEGIRTLLWTFIKSFQALPYVALLIVMLFFIYAVIGMQVFGKIALNDTEINRNNNFQTFPQAVLLLFRCATGEAWQDIMLACMGETPCGSSFAVFYFISFYMLCAFLIINLFVAVIMDNFDYLTR

>Chord_homsap_Cav1.1_NP_000060.2

ALFCLTLENPLRKACISIVEWKPFETIILLTIFANCVALAVYLPMPEDDNNSLNLGLEKLEYFFLIVFSIEAAMKIIAYGFLFHQDAYLRSGWNVLDFTIVFLGVFTVILEQVNDVKALRAFRVLRPLRLVSGVPSLQVVLNSIFKAMLPLFHIALLVLFMVIIYAIIGLELFKGKMHKTCYFTDIVATVENEEPSPCARGRRCT-PGPNHGITHFDNFGFSMLTVYQCITMEGWTDVLYWVNDAIGEWPWIYFVTLILLGSFFILNLVLGVLSGEFTREKQQLDEDLRGYMSWITQGEVMDGKLSLDEGGSDTESLYEIAGLIFRWKCHDIVKSKVFYWLVILIVALNTLSIASEHHNQPLWLTRLQDIANRVLLSLFTTEMLMKMYGLGLRQYFMSIFNRFDCFVVCSGILEILAMTPLGISVLRCIRLLRIFKITKYWTSLSNLVASLLNSIRSIASLLLLLFLFIVIFALLGMQLFGGRYDEDTEVRRSNFDNFPQALISVFQVLTGEDWTSMMYNGIMAYGMLVCIYFIILFVCGNYILLNVFLAIAVDNLAEAESLTSAQKAKAEEKKRRKMSAKLKIDEFESNVNEVKDPYPSADFPGIPEASSFFIFSPTNKIRVLCHRIVNATWFTNFILLFILLSSAALAAEDPI-RADSMRNQILKHFDIGFTSVFTVEIVLKMTTYGAFLHKFCRNYFNMLDLLVVAVSLISMGLEAISVVKILRVLRVLRPLRAINRAKGLKHVVQCMFVAISTIGNIVLVTTLLQFMFACIGVQLFKGKFFRCTDLSKMTEEECRGVHSDFHFDNVLSAMMSLFTVSTFEGWPQLLYKAIDSNAEDVGPIYNNRVEMAIFFIIYIILIAFFMMNIFVGFVIVTFQEQGETEYKNCELDKNQRQVQYALKARPLRCYIPYQYQVWYIVTSSYFEYLMFALIMLNTICLGMQSEQMNHISDILNVAFTIIFTLEMILKLMAFKARGYFGDPWNVFDFLIVIGSIIDVILSEPDESARISSAFFRLFRVMRLIKLLSRAEGVRTLLWTFIKSFQALPYVALLIVMLFFIYAVIGMQMFGKIALVDTQINRNNNFQTFPQAVLLLFRCATGEAWQEILLACSEEYTCGTNFAYYYFISFYMLCAFLVINLFVAVIMDNFDYLTR

>Chord_cioint_Cav3_XP_018667817.1

VLLSLKQTSVPRIWCLRMVANPWFERVSMLVILINCVTLGLYQPCQHRTESERCMVLEMFDHFVFAFFALEMLIKMLAMGV-WGKLGYLGEAWNRLDFFIVLCGMLEYTLQMEDNFTSVRTVRVLRPLRAINRVPNMRILVMLLLDTLPMLGNVLMLCSFVFFIFGVVAVQLWEGTLRQRCFLPPYFTLDDSDDEAICSLNVTCSDQNPYLGSINFDNIMYAWVAIFQVISLEGWVDIMYYLMDGYSFYSFIYFILLIVIGSFFMINLCLVVIATQFSSPPGNCYEEMLKYISHLYRRNKRKQRSPSLCSDRRPSAEINVNPGKFQTQTKVVVDSNYFNRGIMVAILINTLSMGIEHHNQPTGLTEVLEISNVVFTTLFALEMLSKIVAYGFAGYIKNLYNVFDALIVIISVWEIAQNSGGGLSVLRTFRLLRVLKLVRFMPALQRQLVVLMKTMDNVATFMMLLTLFIFIFSILGMHLFGCDFCGRTECDRKNFDSLLWAFVTVFQILTQEDWNIVLYNGMAATSPFAAIYFVTLMTIGNYVLFSLLVAILVEGFQATEQTSQDEEDSDAEEPAILAEASFENEPEQEIEIKTPNVITESQDSSRKKDWSLYLFSPELKFRKAVQKITEHKLFDYMILLLIFGNCITIALERPSLKEEDHERKVIDGFNNVFTFVFLLELILKVIASGFYIGHYLKSGWNVLDFFLVASSLIDVIMTLLGILRVFRLLRALRPLRVISRAPGLKLVVQTLISSLKPIGNIVLICCAFFLIFGILGVQVLKGKFYYCDDLRVTNKTDCLLVNRRYNFDDVGQALMSLFVISSKDGWVEIMYHGIDATGIDQQPIRNSNPWMLLYFVSFLLIVGFFVLNMFVGVVVENFHRCREEHELEEQKRREERRLDYKRSMRPSFQNTKTRRALHAFCLNKYFEIGVSIVIGINIFTMAAQPKVLDQALKIANYFFTAVFVLEAILKLIALGVRRYFRDKWNQVDMIIVILSLVGIAVEASLLINPTIIRVMRVLRIARVLKLLKVSKGIRSLLETVANALPQVGNLGLLFLLLFFIFAALGVELFGTLSCDENGLSRHASFSNFGIALLTLFRISTGDNWNGIMKDVMVTNCCGSIISPIYFVLFVMTAQFVLVNVVVAVLMKQLEDNRA

>Chord_cioint_Cav2_XP_018670105.1

SLFVFGVDNVVRKLAKRIIEWPPFEYLILATIVANCIVLALEEHLAAGDKTPRTIRLEGTEPYFLGIFIVEAAVKILALGFVLHKDSYLRYGWNIMDFTVVVTGCVTYFDTSLG--QTLRAVRVLRPLKLVSGIPSLQVVLKSIMKAMVPLLQIAVLLLFFIVVCSIVGLELYMGRFHRTCYS--NSTNKIIRDNQICSEKYVCEPLGPNHGITTFDNIIFSMMTVFQCITMEGWTDILYFADDATGIYNWAYFIMLIIVGSFFMLNLVLGVLSGEFARKKQQMDRELDGYLEWMQKAEEVIRKSKVDLLDSAESSFTDISATRFRVKCRHLVKSPVFYWIVLFLVLLNTAFLSSVHYKQPKWWEDFLYYAEFVFLGLFSGEILLKVYGLGPRTYFRSSFNIFDFVVIIGSIFEIIPDASFGISVLRALRLLRVFKFTSAWSGLRNLVVSLMSSLRSIVSLIFLLFLFLVVFALLGMQIFGGRFSFKDGKPNSNFDSFPSAILTVFQILTGEDWNMVMYNGVEAKGLWWSLYFIFLVMFGNYTLLNVFLAIAVDNLANAQELTKDEEEEKDNKEAQKALIIGGESRFLSRQNSLRVRNTPMRELGVLPYSSMFIFSPTNPIRLACHYIVNLKYFETTILVIIILSSLTLATEDPV-TKDSQRNNVLKYFDYIFTAVFTFEMVMKMIDLGLVLHPYFHSLWNILDFVVVCSALVGFALTDLGVIKSLRVLRVLRPLKTIKRLPKLKAVFMCVVNAFRNVATILIVYMLFMFIFAVIAVELFKGKFFYCTDPMINVESECKGLQHDFHYDNVLYSFLTLFVISTGEGWPEVLWHSIGSTYEDKGPERGFRMEVSIFYIVFFVVFPFFFVNIFVAFIIITFQEEGDKAMSNCSLEKNERAVDFAISSKPLTRFMPLQYHVWKVVVSPVFEWIIMALIVLNTVVLMLQSPNYENILQYCNMAFTAIFTIECIIKMAAFKPINYFRDSWNIFDFITVVGSIADVTITLHDMSGFINLSFLRLFRAARLIKLLRQGETIRILLWTFVQSIKALPYVCLLIAMLFFIYSIIGMQLFGNIQLDPSAINHHNNFTHIFQALMLLFRCATGEAWQSVMLACVKENGCGSVISYIYFTSFIFFCSFLMLNLFVAVIMDNFEYLTR

>Chord_cioint_Cav1_XP_018667169.1

SLLCLSLKNPFRKACLKIVEWRPFDVLILLTIFANCCALAIYVPFPGEDSNATNEILEKVEYVFLAIFTVESFMKIIAFGFAFHPNAYLRNGWNILDFIIVIVGLISIVFEMADDKRALRAFRVLRPLRLVSGVPSLQVVLNAIIRAMLPLLHIALLVMFVIIIYAVVGLELFKGKLHKTCYFTGMTDVIANEDPQPCAGGRHCP-DGPANGIINFDTFYFSFITVFQCITMEGWTEVLYYTNDAMGHLPWMYFVSLIIVGSFFVMNLILGVLSGEFSREKQQLDEDVRGYMEWITQAEDIDDMDEKRQGDNEDGSSDVTA--KTRRKCRLMVKSQTFYWLVIVLVFFNTLSLATEHYQQPDWLTSVQEISNKVLLGIFTLEMLLKMYALGMQVYFVSLFNRFDCFVVCGGIVEMVVMEPLGISVLRCVRLLRIFKVTRYWSSLSNLVASLLNSIRSIAGLLLLLFLFIVIFSLLGMQLFGGRFNEGDQKIRSNFDTFLQALLTVFQILTGEDWNVVMYYGIRAYGLITSIYFIILFVCGNYILLNVFLAIAVDNLADAESLNVAQKEKEEEQKRKKTMSSLQTDEIDHEIRIEVTEASETNSDRMPQATSFFIFTPTNPFRKWCHFIANNNIFNNGIFVCIMLSSVALACEDPI-DSKSELNEVLKYFDYVFTGIFTVEIILKMVAYGVILHKFCRNSFNLLDLLVVGVSLISIFGNGFSVVKILRVLRVLRPLRAINRAKGLKHVVQCVIVAISTIGNIFIITTLLQFMFACIGVQLFKGRLYGCTDESKSTREECKGVNNDFNYDNVLNAMLTLFVVATFEGWPALLYKSIDSWKEGVGPKYDARPAVALFYFIYIIVIAFFMMNIFVGFVIVTFQEQGEQEYRNCELDKNQRQVEYALKAKPTRRYIPWQYKAWFVVNSTYFEYFMLVLILLNTVCLAIQDAGLTRILNHMNLVFTTLFTIEMIFKLIAFKPRGYISDPWNIFDFLVVIGSIVDILLSKTGGDKSFSINFFRLFRVMRLVKLLSRGEGIRTLLWTFIKSFQALPYVALLIVLLFFIYAVIGMQVFGKVKPIDEQINRNNNFQTFIQSVLLLFRCATGESWQEVMLAASDKFACGSDFSYTYFLTFYMLCAFLIINLFVAVIMDNFDYLTR

>Choan_salros_Cav_XP_004995501.1

RTWLAQAGATVRDACAALSRSFIFEQAVLLVILFNCITLALYDTSDSTCSTRRCKILEVCELVVTVAFTVEMLIRMIATGVR----QYFGSGWNRFDAIIIVFGFVDFIPTVSGGTTLVRLVRILRPMRMVTRFQSLQLLVALLLDIIPMLGSLAILTLFLFCTFGLVGVQMWKGMLRQRCYDVPSTHATPPWTTYICGADFTCPAPNPFAGAVSFDNIAIACNTVFQVITLETWGNIMAAVQKAHSFWAFIYFVLLIFAGSWFALNLVLVVIATQFKRPRTRREEVVADFRRW-LKMDIVHQLPKRIATRTIPRPASTAMRTQIRRACARIARHPRFSNFVVICIFVNMAVMSLEHMGQPHALEEFNRITNAVFTVVFAGEMVIKIVGLGPIAYLINKANVFDFVIVLISLSEF-GTGTNGLTVLRSFRLLRVFTALQVLPTMRRQLAVMIKTLDSVLTFLFLLGLFVFMAAIAGMRLFGQRLEDGTGVPRANFSTFWNAVILVFQVLTTEDWTLVMYKAAHATSPTACIYFVLVLVLGTYILFNLFIAILVEGFATSPGLGQAQQERQEQREQQGDIITPEEERINEDDNRPPLFVRVCPGCN----WRLRVA----RARACVRRVLQNPRCEAAMFVLILMSALTLLWETP--RAGEKRKRILQGVYMFFNTAFLIEVLCRVFAEGFLPRKFLASGWNILDGFVVLTSWISMGLEVLRTIRVLRALRALRPLRLVHRMPKLKQTVGTLFTSVRPLTNVLLIEIIFFLIFAILGVQLFKGSLHACTNTTVTDKASCIAEAFQYNFDNLAMALLTLFVVSTRDGWVLVMDRATDAVGPGRQPQRNANPLAALYFVAFVLVVGYFVINMFVGVLVENFQDEGNKDDKGGDIEDEEKEVALTLNARPRRRGHTFRLACCALVHHPTFDLVTSSLNASNVVLMMAQPAWVGHMLSTAELFFTACFTLEVIVKLVASGARVFAQSAWNKLDLFVVVTSIAGIVVEWGLPVNPTVLRVLRILRVARVFRLAKMTRGMRSLLATVTQSASQVGSLALLLLLLFYISAAIAVELFGRMSCSVSGLSSHTNFRNLGMAMLALFQVATGDNWTGILSDALEIGCCAPHIASLFLVTFVILAQFILLNVVVAVLMKHLT----

>Choan_salros_Cav_XP_004989719.1

--------------------------------------------MTNDDLSSRET--DSLEYVFLAIFTLEALLKIIATGFLFCPPSYLRNKWNILDFIIVAVGLIGVVVEQSGDVKALRALRVLRPLRLITSVQSLQIVLNSILLSIPALADVAMLLGFLIVIYAIIGLEFYRGVLNHQCFLNNTPYFLAPDTA-PCDPGRVCSGSSPNSNITAFDHAG-SFLKFDAAESMAEHEQNFFNSSAGADDDD----DDDIDDDATTVLVLGLSLPSRDFVAEQGEDRDKLMSFLAH--------QGVQVKSLREYNEILT-----QMLSKLGAVVKSRWFNLVVTFMVLVNTVLLAVQTDAGATDEAAFATIVEASFVGLFVLEMLVKLAGLRPHMYFESKFNRFDLTVVLLSLLELIGLRSIGISALRSLRLLRIFRMKQYWEDINDFVVSLLNSIASIVSLLLLILIYMVIVALLGMQIFGGRFDFEDPKPRINFDDFFSALLTVFIVIVGDDWNSVMYNGILAYNGWAIVFFCVVVILGMFVLLNVFLAIAVKSLDDARDLKAARDEHKERWKAEAAVDRRRHRQYANPLVGAAEQEKEEVELQVANNKSLFCLGPRNSFRKFCNNIAYDNRFESVILLLILISSALLAAEDPV-NLDAQINKDLETADIFFTSVFSLEMALKIVALGF----YITDPWNDLDAVVVLASVVSLAISDAAVVRVLRVFRVLRPLRAIKRAPGLRKVVSCMVVSIKTIGNVFIVTFLLTFIYAIIGVQSFKECFGRCNDPDVMFKSQCNGSTPYFNFDNVGKGMLTLFTVSTLEGWIDVMNNAIDCTAENRQPERNNNPVAALFFVTYVILVAFFMLNIFVGYVIITFSSEGESYEAVDGLDKNQRKLAFCLNAQPIRVHRPAQISIFRFVSSKHFEWFIMAAIIGNSIVLLMMPSDYEMGLQLCNIVFTGIFTVEALLKLFALNPTGYFHDSWNFFDFIIVVGSLVDVFLSASSGDSGVNIGFLRLFRVARLLKLVSRGKGMKRLLWTFAKSFQSLPYVAALIMMLFFVYAVIGMQLFARTGFREGDINEHNNFRDFFGALLLLFRCATGENWQNMMRDHLEPGVCGSVVAVPFFCTFLVLCSFLILNLFVAVIMDNFEYLTQ

>Chlamydomonas_eustigma_Cav_GAX86028.1

ALLFLQVNSFIRKPCIRLVRW--------------------------------------------MFFDALMLLRMV---FVFGKYTYLRDGWNILDFIVVVMGVLELTSLGNY--TFIRSFRALRPLRAITKIASLKIIVESLFRSLPMLGDVMILAMFYFSVFGIFCTELFKGQLYGRCGAMNVSYVVSTTAAQVCKGGYACP-GNPDGGYRNFDNILITWVQLYQHMTWQDWSYIMYATQAAMSWWTWPLHIFLVIVGGLLLANLALAVIFLHFSKSAKMLAVELNV-VTDI-------LRQPILDFPPGPSLQKPLVVTQFRDLNYTICYSTWFLHLTTFMIVSNAIVLAIYWYEMPQEWVTGTTNANIAFSCYFVLEMLIKIIGMGPRQYAADSFNIFDFFVTLLGVVDMSGVSSPGLSVFRTFRLLRVMRLARSWTGLNRIIQVLLSSLVSVGWLTVLLFMYIFITGLLGMAFFGFKFDGQSTTFMPNYDNIFQAMLSTFIILTSDNWDSNMKIMVLTQSPWPAFYTIITMTLGIFTVLNLFLAILLNNLDDVVQSSNVERVQDEETRYIEDLSKTRTESFWPRKINRVSPLSQGQYPVTLEGRSLFMFAPTNQVRKFLLLVTSNVHFEYAMLFLISLSSLELCFDDASSVPGTTKFAALRALDVFFTITFGLEALMKIFTYGLFNGKYLRNPWNILDMFVVIVNVLVLALDYIIWLRAFRALRALRPLRVASQLDGIRVIVMAMAKSLPAMGEIFLVGALFFYIFAVLGVNLMCGLFLGCYSQGNRTWCEADGNGQLARFDNLIMALWVLFWMTSLENWSPIMIQAMDITSLDDQPVFNNNIYITFYFIVFIVIGVYFIMNLVIGVAISTFGKMREQLGRSALLTEAQQEATLQLTKK-YKR--PFRLSVYKMVMTERFEKIMMCIIIANLLPLFMESDTWAAGLGVVNVVFTALYVIEMILKWISIGACAYFKDKWCLFDFLVVVVSVMGVIIDYVLHDNLTILTVLRSLRVLRIFKIIPKARGLKMMMTTLLWSLPALMNVATVLLLFMYTFAIIAMNIFGNMKW--GEIDLYANFESFPTAMFTLFRMQTGENWNYVMTACMQNNRCSPAVAAIFFTLYMALCTYLVLELVVAVIIENIEYQSQ

>Caenorhabditis_nigoni_Cav3_PIC19332.1

ALRCFYQARPPRKWALQMVMSPWFDRITMAVILINCVTLGMYRPCEDGPDTYRCQILDIIDNCIFVYFAIEMVIKIMALGF-CGPAAYMSDTWNRLDFFIVMAGIAEFVLHEYLNLTAIRTVRVLRPLRAVNRIPSMRILVNLLLDTLPMLGNVLLLCFFVFFIFGIVGVQLWAGLLRNRCVILTRFYIPEDTSLYICSQGIKCNQRNPFQGSVSFDNIGFAWVAIFLVISLEGWTDIMYYVQDAHSFWNWIYFVLLIVIGAFFMINLCLVVIATQFAEEGGDTYAAIVRFIGHTFRRTKRASRIEEKAEDEEDEIATPPEIKWFRDKVRKFVVCDHFTRGILVAILVNTLSMGVEYHQQPEILTVILEYSNLFFTALFALEMLLKIIASGFFGYLADGFNLFDGGIVALSVLELFQEGKGGLSVLRTFRLLRILKLVRFMPALRYQLVVMLRTMDNVTVFFGLLVLFIFIFSILGMNLFGCKFCLAKKCERKNFDTLLWALITVFQILTQEDWNMVLFNGMAQTNPWAALYFVALMTFGNYVLFNLLVAILVEGFQEEEQLEEEARAVEEEDERKRELVPYRRQRVHSWSGLCHHFNPNCPVHGNRTEFSLFLMGPKNPLRIKCLQTTQKKWFDYTILFFIGINCITLAMERPSIPPDSFERRFLQVSGYIFTVIFTGEMMMKVIANGCFIGQYFKDGWNILDGILVVISLINVAFEIFGVIRVLRLLRALRPLRVINRAPGVKLVVMTLISSLKPIGNIVLICCTFFIIFGILGVQLFKGMMYHCIEVGVTTKVDCLKVNHRYNFDNLGQALMSLFVLSSKDGWVSIMYQGIDAVGVDVQPIENYNEWRMIYFISFLLLVGFFVLNMFVGVVVENFHKCKEREMREKEKEKRLKRQKFEESMAPYYHYGHTRLFLHGIVTSKYFDLAIAAVIGINVISMAMMPMGLKYVLKALNYFFTAVFTLEAAMKLIALGFKRFFIEKWNRLDMFIVILSIAGIIFEEELPINPTIIRVMRVLRIARVLKLLKMAKGIRSLLDTVGEALPQVGNLGSLFFLLFFIFAALGVELFGKLECSEDGLGEHAHFKNFGMAFLTLFRIATGDNWNGIMKDALETNCCVPILAPCFFVIFVLISQFVLVNVVVAVLMKHLEESNK

>Caenorhabditis_nigoni_Cav2_PIC16379.1

SLFIFAEDNIIRRNAKAIIEWGPFEYFILLTIIGNCVVLSMEQHLPKNDKKALSEWLERTEPYFMGIFCLECILKVIAFGFALHKGSYLRSGWNVMDFIVVVSGVVTMLPFGSTDLRTLRAVRVLRPLKLVSGIPSLQVVLKSILCAMAPLLQIGLLVLFAIIIFAIIGLEFYSGAFHSACYNERGEIENVSEKPMPCTNVYNCD-IGPNYGITSFDNIGFAMITVFQCITMEGWTTVMYYTNDSLGTYNWAYFIPLIVLGSFFMLNLVLGVLSGEFARRQQQIERELNGYLEWILTAEEVIKQQSTETEEDFEEDEDEMEEEQLRIQIRIMVKTQIFYWSVITLVFLNTCCVASEHYGQPQWFTDFLKYAEFVFLGIFVVEMLLKLFAMGSRTYFASKFNRFDCVVIVGSAAEVIYGGSFGISVMRALRLLRIFKLTSYWVSLRNLVRSLMNSMRSIISLLFLLFLFILIFALLGMQLFGGRFNFPTMHPYTHFDTFPVALITVFQILTGEDWNEVMYLAIESQGGWYSIYFIVLVLFGNYTLLNVFLAIAVDNLANAQELTAAEEADEKANEIEEESGDHCTIDMEGKTAGDMCAVARAMDEMMVPYSSMFFLSPTNPFRVLIHSIVCTKYFEMMVMTVICLSSVSLAAEDPV-DEENPRNKVLQYMDYCFTGVFACEMLLKLIDQGILLHPYCRDFWNILDGIVVTCALFAFGFANLNTIKSLRVLRVLRPLKTIKRIPKLKAVFDCVVNSLKNVFNILIVYFLFQFIFAVIAVQLFNGKFFFCTDKNRKFAHTCHGRLRPFNYDNTINAMLTLFVVTTGEGWPGIRQNSMDTTFEDQGPSPFFRVEVALFYVMFFIVFPFFFVNIFVALIIITFQEQGEAELSEGDLDKNQKQIDFALNARP-RSFMPTKYRIWRLVTSPPFEYFIMTMICCNTLILMMNPLFYEEILRLFNTALTAVFTVESILKILAFGVRNYFRDGWNRFDFVTVVGSITDALVTE--GGHFVSLGFLRLFRAARLIRLLQQGYTIRILLWTFVQSFKALPYVCLLIGMLFFIYAIVGMQVFGNIWLNATEINRHNNFQSFFNAVILLFRCATGEGWQDIMMAAVKGQTCGSNVSYAYFTSFVFLSSFLMLNLFVAVIMDNFDYLTR

>Caenorhabditis_nigoni_Cav1_PIC34019.1

SLLCLSLNNPIRKLCISIVEWKPFEFLILFMICANCIALAIYQPYPAQDSDYKNTLLETIEYVFIVVFTIECVLKIVAMGFLFHPSAYLRNAWNILDFIIVVIGLVSTILSKMSDVKALRAFRVLRPLRLVSGVPSLQVVLNAILRAMIPLLHIALLVLFVILIYAIIGLELFCGKLHSTCID-PATGQLAQKDPTPCGNAFKCR-PGPNNGITNFDNFGLAMLTVFQCVSLEGWTDVMYWVNDAVGEWPWIYFVTLVILGSFFVLNLVLGVLSGEFSREKQQLEEDLKGYLDWITQAEDIEAVTGEEVDEEGEERVEDVRP-RCRRACRRLVKSQTFYWLVILLVLLNTLVLTSEHYGQSEWLDHFQTMANLFFVILFSMEMLLKMYSLGFTTYTTSQFNRFDCFVVISSILEFVLMKPLGVSVLRSARLLRIFKVTKYWTSLRNLVSSLLNSLRSIISLLLLLFLFIVIFALLGMQVFGGKFNPQQPKPRANFDTFVQALLTVFQILTGEDWNTVMYHGIESFGVIVCIYYIVLFICGNYILLNVFLAIAVDNLADADSLTNAEKEEEQQ-------------EIEGEDEEFDEGEEEGDEHGIPKASSLFILSHTNSFRVFCNMVVNHSYFTNAVLFCILVSSAMLAAEDPL-QANSTRNMVLNYFDYFFTSVFTVEITLKVIVFGLVFHKFCRNAFNLLDILVVAVSLTSFVLRAMSVVKILRVLRVLRPLRAINRAKGLKHVVQCVIVAVKTIGNIMLVTFMLQFMFAIIGVQLFKGTFFLCNDLSKMTEAECRGSNNDFNFDNVGDAMVSLFVVSTFEGWPQLLYVAIDSNEEDKGPVHNSRQAVALFFIAFIIVIAFFMMNIFVGFVIVTFQNEGEREYENCELDKNQRKIEFALKAKPHRRYIPLQYRVWWFVTSRAFEYVIFLIIVMNTVSLACSSRGFEDFLDVFNLIFTGVFAFEAVLKIVALNPKNYISDRWNVFDLLVVVGSFIDITYGKPGGTNLISINFFRLFRVMRLVKLLSRGEGIRTLLWTFMKSFQALPYVALLIVLLFFIYAVIGMQFFGKVALDDTSIHRNNNFHSFPAAILVLFRSATGEAWQDIMLSCSNESRCGNNFAYPYFISFFMLCSFLVINLFVAVIMDNFDYLTR

>Biomphalaria_glabrata_Cav3_XP_013096433.1

----------------------------MFVIILNCVTLGMYQPCNDECDTFRCKLLENFDHFIFAFFAVEMGIKMIAMGV-AGKDTYLADSWNRLDCFIVVAGLAEYIVNKTISLSAIRTVRVLRPLRAINRIPSMRILVMLLLDTLPMLGNVLLLCFFVFFIFGIIGVQLWSGVLRRRCFLVPLYYIKSPKIEYICSPGIACNGENPYQGAVSFDNIGLAWVAIFQVISLESWVNIMYFVQDAHSFWDWIYFVALIVIGSFFMINLCLVVIATQFSSEPAGCYTELLKLIAQMYRRVKRKASNMLLNVDLEPSKTQSIADKGMQKRLKYFVESNFFQRSILVAILLNTLSMGVEYHNQPEELTIVLEYSNIVFCVMFGTEMTFKIFAYGLFGYISNGFNVFDGFIVILSIVELAENGASGLSVLRTFRLLRILKLVRFMPALRRQLVVMLRTMDNVATFFALLVLFMFIFSILGMNLFGGSFCSLCKCDRANFDNLLWSLVTVFQVLTQEDWNTVLYNGMAKTSTWASLYFVALMTFGNYVLFNLLVAILVEGFSTKEETDAADKDEEDEEEKEKQRSRQNSFTSHRTVNSLGSGNSKDDKSNERHEYAFYLLSNENSLRKLAHHLISRKWFDNTVLIFIALNCITLAMERPDIPPESVERHFLIYTNYVFTFVFTIEMLIKVTAKGFFIGKYFKSGWNVMDGFLVIISLIDIFISIFGILRVFRLLRTLRPLRVISRAPGLKLVVQTLLSSLRPIGNIVLICCTFFIIFGILGVQLFKGTFYYCRNVTVTNRNQCLEVNQKYNFDNLGQALMALFVLASKDGWVQIMYTGLDAVGVDKQPIENYNEWRLIYFISFLLLVAFFVLNMFVGVVVENFHKCRESQEIEERAKRAAKREKLDKKRKPYWAYSHSRLLIHTVINSKYFDLAIAAVIGLNVITMAMMPEELTFALKIFNFFFTSVFILESVMKIIALGFLRYIKDRWNQLDILIVILSVVGIVLEEFIPINPTIIRVMRVLRIARVLKLLKMAKGIRALLDTVIQALPQVGNLGLLFFLLFFIFAALGVELFGRLDCDREGLGKHAHFKNFGMAFLTLFRVATGDNWNGIMKDTLLKDCCVPLIAPVYFVVFVLMAQFVLVNVVVAVLMKHLEYKYK

>Arthr_dromel_Cav3_ABW09342.1

SIRALTQYTRPRNWCLMLITNPWFERVSILVILLNCITLGMYQPCVDDAVTNRCKILQIFDDIIFAFFALEMTIKMVAMGI-CGKNTYLADSWNRLDFFIVLAGLLEYVMHVENNLTAIRTIRVLRPLRAINRIPSMRILVMLLLDTLPMLGNVLLLCFFVFFIFGIIGVQLWEGILRQRCSLLSQYYEFSKDQDYICSTSLVCNGENPFQGTISFDNIGMAWVAIFLVISLEGWTDIMYYVQDAHSFWDWIYFVLLIVIGSFFMINLCLVVIATQFSSEPATCYAEIVKYIAHLWRRFKRRHTAEALRAHHKPRSVPTGQNQWIRRYIRRLVEHKYFQQGILLAILINTLSMGIEYHNQPPELTAIVETSNVVFSGIFAVEMLLKVVAEGPFRYIANGFNVFDGIIVILSAIEICGGGGSGLSVLRTFRLLRILKLVRFMPNLRRQLFVMLRTMDNVAVFFSLLVLFIFIFSILGMYLFGGKFCPQCECDRKHFNNILWATVTVFQILTQEDWNVVLFNGMEKTSHWAALYFVALMTFGNYVLFNLLVAILVEGFSSESADSDRDRERDRDRDRERDRRRASACIFNSQVYQNLNQPPKLRPGSEREDYSLYIFPEDNRFRQICTWFVNQKWFDNVVLLFIALNCITLAMERPNIPPSSTERLFLATANYVFTVVFTVEMFIKVVATGMFYGHYFTSGWNIMDGSLVTISIIDLLMSIFGILRVFRLLRSLRPLRVINRAPGLKLVVQTLLSSLRPIGNIVLICCTFFIIFGILGVQLFKGTFYYCENIKVRNADECRRTNRKYNFDDLGKALMSLFVLSSRDGWVNIMYTGLDAVGVDQQPIVNYNEWRLLYFIAFILLVGFFVLNMFVGVVVENFHRCREEQEKEEKIRRAAKRLQMEKKRRPYYTYSPTRMFVHNVVTSKYFDLAIAAVIGLNVVTMAMMPSGLKYALKIFNYFFTAVFILEANMKLVALGWKLYLKDRWNQLDVGIVLLSIVGIVLEEIIPINPTIIRVMRVLRIARVLKLLKMANGIRALLDTVMQALPQVGNLGLLFFLLFFIFAALGVELFGRLECSDQGLGEHAHFANFGMAFLTLFRVATGDNWNGIMKDTLVRNCCVSVIAPIFFVIFVLMAQFVLVNVVVAVLMKHLEESHK

>Arthr_dromel_Cav2_AFH07350.1

SLFILTEDNPIRKYTRFIIEWPPFEYAVLLTIIANCVVLALEEHLPGGDKTVLAQKLEKTEAYFLCIFCVEASLKILALGLVLHKHSYLRNIWNIMDFFVVVTGFMTQYPQIGPDLRTLRAIRVLRPLKLVSGIPSLQVVLKSIIKAMAPLLQIGLLVLFAIVIFAIIGLEFYSGALHKTCYSDPNKLVKEGESETPCNTSFVCN-EGPNSGITSFDNIGFAMLTVFQCITMEGWTAILYWTNDALGAFNWIYFVPLIVIGSFFMLNLVLGVLSGEFSRFRAMFQTAMVSYLDWITQAEEVIGKSKSTDTEEEEAEEDYGDDGRFRFWIRHTVKTQWFYWFVIVLVFLNTVCVAVEHYGQPSFLTEFLYYAEFIFLGLFMSEMFIKMYALGPRIYFESSFNRFDCVVISGSIFEVIKGGSFGLSVLRALRLLRIFKVTKYWSSLRNLVISLLNSMRSIISLLFLLFLFILIFALLGMQLFGGQFNLPGGTPETNFNTFPIALLTVFQILTGEDWNEVMYQGIISQGMIYSIYFIVLVLFGNYTLLNVFLAIAVDNLANAQELTAAEEEQVEEDKEKQLQALQADGVHVENGDGAVAPSKSKGKKKMLPYSSMFILSPTNPIRRGAHWVVNLPYFDFFIMVVISMSSIALAAEDPV-RENSRRNKILNYFDYAFTGVFTIEMLLKIVDLGVILHPYLREFWNIMDAVVVICAAVSFGFDNLSTIKSLRVLRVLRPLKTIKRVPKLKAVFDCVVNSLKNVVNILIVYILFQFIFSVIGVQLFNGKFFYCTDESKHTSAECQGKPRAFHYDNVAAAMLTLFAVQTGEGWPQVLQHSMAATYEDRGPIQNFRIEMSIFYIVYFIVFPFFFVNIFVALIIITFQEQGEAELQDGEIDKNQKSIDFTIGARPLERYMPFKYKVWRIVVSTPFEYFIMMLIVFNTLLLMMQGDMYEKSLKYINMGFTGMFSVETVLKIIGFGVKNFFKDPWNIFDLITVLGSIVDALWMEGHDSNSINVGFLRLFRAARLIKLLRQGYTIRILLWTFVQSFKALPYVCLLIAMLFFIYAIIGMQVFGNIKLGTNSITRHNNFQSFIQGVMLLFRCATGEAWPNIMLACLPGEYCGSTLAYAYFVSFIFFCSFLMLNLFVAVIMDNFDYLTR

>Arthr_dromel_Cav1_AAF53504.1

ALFCLSVKNPLRALCIRIVEWKPFEFLILLTIFANCIALAVYTPYPGSDSNVTNQTLEKVEYVFLVIFTAECVMKILAYGFVLHNGAYLRNGWNLLDFTIVVIGAISTALSQLMDVKALRAFRVLRPLRLVSGVPSLQVVLNSILKAMVPLFHIALLVLFVIIIYAIIGLELFSGKLHKACRD---EITGEYEENRPCGVGYQCP-DGPNDGITNFDNFGLAMLTVFQCVTLEGWTDVLYSIQDAMGDWQWMYFISMVILGAFFVMNLILGVLSGEFSREKQQIEEDLRGYLDWITQAEDIEQPNEMDSTENLGEEMPEVQMTRMRRACRKAVKSQAFYWLIIVLVFLNTGVLATEHYGQLDWLDNFQEYTNVFFIGLFTCEMLLKMYSLGFQGYFVSLFNRFDCFVVIGSITETLMMPPLGVSVLRCVRLLRVFKVTKYWRSLSNLVASLLNSIQSIASLLLLLFLFIVIFALLGMQVFGGKFNGKEEKYRMNFDCFWQALLTVFQIMTGEDWNAVMYVGINAYGALACIYFIILFICGNYILLNVFLAIAVDNLADADSLSEVEKEEEPHDESAQKKIDMEQQELDDEDKMDHETLSDEEVREIPPGTSFFLFSQTNRFRVFCHWLCNHSNFGNIILCCIMFSSAMLAAENPL-RANDDLNKVLNKFDYFFTAVFTIELILKLISYGFVLHDFCRSAFNLLDLLVVCVSLISLVSSAISVVKILRVLRVLRPLRAINRAKGLKHVVQCVIVAVKTIGNIVLVTCLLQFMFAVIGVQLFKGKFFKCTDGSKMTQDECYGSNNRFHFDDVAKGMLTLFTVSTFEGWPGLLYVSIDSNKENGGPIHNFRPIVAAYYIIYIIIIAFFMVNIFVGFVIVTFQNEGEQEYKNCDLDKNQRNIEFALKAKPVRRYIPIQYKVWWFVTSSSFEYTIFILIMINTVTLAMQPLWYTELLDALNMIFTAVFALEFVFKLAAFRFKNYFGDAWNVFDFIIVLGSFIDIVYSESAGSNLISINFFRLFRVMRLVKLLSKGEGIRTLLWTFIKSFQALPYVALLIVLLFFIYAVVGMQVFGKIALDGNAITANNNFQTFQQAVLVLFRSATGEAWQEIMMSCSPGEPCGSSIAYPYFISFYVLCSFLIINLFVAVIMDNFDYLTR

>Arthr_apimel_Cav3_NP_001314887.1

ALRYLDQNTRPRNWCLALITNPWFERVSMMVILLNCITLGMYQPCVDDQVTNRCKILQMFDDIIFAFFSLEMTIKMVAMGI-YGKGTYLADSWNRLDFFIVIAGALEYCLNVENNLSAIRTIRVLRPLRAINRIPSMRILVMLLLDTLPMLGNVLLLCFFVFFIFGIVGVQLWEGILRQRCFLDDLEKYFEYQGQYICSRNVVCNGNNPFQGTISFDNIGLAWVAIFLVISLEGWTDIMYYVQDAHSFWDWIYFVLLIVIGSFFMINLCLVVIATQFSSEPTTCYAEIVKYIAHLWRRGKRREAMTCQELLALSGALSAALPTCIRRLIKKLVEHKYFQQGILLAILINTLSMGIEYHNQPEQLTIVVEVSNIVFSAVFAVEMLLKIIAEGPFGYISNGFNVFDGVVVVLSVVEICRGGSSGLSVLRTFRLLRILKLVRFLPNLRRQLFVMLRTMDNVAVFFSLLVLFIFIFSILGMYLFGGKFCPLCRCDRKHFNDIVWALVTVFQILTQEDWNVVLFNGMQKTSHWAALYFVALMTFGNYVLFNLLVAILVEGFSSQRRLAAKETGIGSDDGSSRISRRNSLRENENVQSPTRKTLPLDEVPMERDDYSLYIFPPNNRFRVLCRLLVDQRWFDNVVLFFIGLNCITLAMERPNIPPDSGERLFLSTANYIFTGVFAVEMFIKVVASGMLYGSYFTSGWNIMDGVLVIISIIDLSMSIFGILRVFRLLRSLRPLRVINRAPGLKLVVQTLLSSLRPIGNIVLICCTFFVIFGILGVQLFKGAFYYCEDIKVRNKTDCLALNRKYNFDDLGKALMSLFVLSSRDGWVNIMYTGLDAVGVDQQPIENYSEWRLLYFIAFILLVGFFVLNMFVGVVVENFHRCREEQEKEERVRRAAKRLQMEKKRRPYYTYSKSRLFVHNVVTSKYFDLAIAAVIGLNVVTMAMMPKALTYALKIFNYFFTAVFILESFMKLLALGLHLYLKDKWNQLDVGIVILSVVGIVLEEIIPINPTIIRVMRVLRIARVLKLLKMAKGIRALLDTVMQALPQVGNLGLLFFLLFFIFAALGVELFGRLECSDQGLGEHAHFSNFGMAFLTLFRVATGDNWNGIMKDTLVKNCCVTIIAPIFFVIFVLMAQFVLVNVVVAVLMKHLEESHK

>Arthr_apimel_Cav2_XP_016766516.1

SLFILSEDNCIRKHTRFIIEWPPFEYAVLLTIIANCVVLALEEHLPKQDKTILAQKLEATEIYFLGIFCVEASLKILALGFVLHRGSYLRNIWNIMDFFVVVTGFITAFSQGIEDLRTLRAIRVLRPLKLVSGIPSLQVVLKSIIKAMAPLLQIGLLVLFAIVIFAIIGLEFYSGTLHKTCYSDINVIVKEGEQASPCNTAHVCD-EGPNFGITSFDNIGFAMLTVFQCITMEGWTAILYWTNDALGTYNWIYFIPLIVLGSFFMLNLVLGVLSGEFARRQQQLEHELYCYLNWICKAEEVIGKSKSTDTEEEEGDDDQDDGFRFRYWIRKSVKSQKFYWFVIVLVFFNTVCVAVEHYGQPQWLTDFLYFAEFVFLALFMLEMFIKVYALGPRTYFDSSFNRFDCVVISGSIFEVIKSGSFGLSVLRALRLLRIFKVTKYWKSLRNLVISLLSSMRSIISLLFLLFLFILIFALLGMQLFGGQFNFDSGTPPTNFNTFPIALLTVFQILTGEDWNEVMYQGIESQGMIYSLYFIVLVLFGNYTLLNVFLAIAVDNLANAQELSAAENEEEEEDKQKQAQVEICPPSPNQNFKDGKGGKQSSEEEKMLPYSSMFILSPTNPVRRAAHWVVNLRYFDFFIMVVISLSSIALAAEDPV-WEDSPRNEVLNYFDYAFTGVFTVEMILKIIDLGIILHPYLREFWNIMDAVVVICAAVSFAFDNLSTIKSLRVLRVLRPLKTIKRVPKLKAVFDCVVNSLKNVINILIVYILFQFIFAVIAVQLFNGKFFYCSDESKYTQQDCQGQSQFFHYDNVMAAMLTLFAVQTGEGWPQILQNSMAATYEDKGPIQNFRIEMSIFYIVYFIVFPFFFVNIFVALIIITFQEQGEAELQDGEIDKNQKSIDFTIQARPLERYMPVKYKIWRIVVSTPFEYFIMGLIVLNTVLLMMQSDAYKNTLKYMNMCFTGMFTVECILKIAAFGVRNFFKDAWNTFDFITVIGSIVDALVIEEKKENFINVGFLRLFRAARLIKLLRQGYTIRILLWTFVQSFKALPYVCLLIAMLFFIYAIIGMQVFGNIALDATSITKHNNFQSFIQGLMLLFRCATGEAWPNIMLSCVQEGGCGSNIAYAYFVSFIFFCSFLMLNLFVAVIMDNFDYLTR

>Arthr_apimel_Cav1_XP_016766333.1

TLFCLPLKNPLRKMCIDVVEWKPFEWLILMTIFANCIALAVYTPYPYGDSNLTNQYLEKIEYIFLVIFTVECVMKIIAYGFVAHPGAYLRNGWNILDFSIVVIGMVSTVLSVLMDVKALRAFRVLRPLRLVSGVPSLQVVLNSILRAMIPLLHIALLVLFVIIIYAIIGLELFSGKMHKTCRH--NMTDAIMDDPVPCGPGYQCD-EGPNWGITNFDNFGLAMLTVFQCVTLEGWTEVLYNIEDAMGSWQWIYFISMVILGAFFVMNLILGVLSGEFSREKQQIEDDLRGYLDWITQAEDIEQQSEMESTDQLEGDEE-----RMRRACRKAVKSQVFYWLIIVLVFLNTGVLATEHYNQPHWLDDFQEITNMFFIALFTMEMMLKMYSLGFQGYFVSLFNRFDCFVVIGSITEMIVMPPLGVSVLRCVRLLRVFKVTKYWRSLSNLVASLLNSIQSIASLLLLLFLFIVIFALLGMQVFGGKFNVLENKPRHNFDSFWQSLLTVFQILTGEDWNAVMYDGIRAYGMLACFYFIILFICGNYILLNVFLAIAVDNLADAESLTAIEKEAEEEAEKNKSHNTHAKVRLNIESDEEVEEEEEVEHNEIPAGSAFFIFSQTNRIRIFCHWLCNHSTFGNVILVCIMISSAMLAAEDPL-RASSSRNLVLQKFDYFFTTVFTIEICLKMISYGFIIHEFCRSAFNLLDLLVVCSSLISMSFSAFSVVKVLRVLRVLRPLRAINRAKGLKHVVQCVIVAVKTIGNIVLVTSLLQFVFAVVGVQLFKGKFFYCTDASKMTKEECQGCQQRFHFDDVAKAMLTLFTVSTFEGWPSLLDYSIDSNKEDHGPIHNFRPIVAAYYIIYIIIIAFFMVNIFVGFVIVTFQNEGEQEYKNCELDKNQRNIEFALKAKPVRRYIPIQYKVWWFVTSQPFEYTIFTLIMINTVTLAMQPEIYTQALDVLNMIFTAVFALEFIFKLAAFRFKNYFGDAWNVFDFIIVLGSFIDIVYSENPGSTIISINFFRLFRVMRLVKLLSRGEGIRTLLWTFIKSFQALPYVALLIIMLFFIYAVIGMQVFGKIAIDDTSINRNNNFQSFPQAVLVLFRSATGESWQEIMMDCSNTNGCGSDIAFPYFISFYVLCSFLIINLFVAVIMDNFDYLTR

>Aplysia_californica_Cav2_AVD53847.1

SLFIFSEENFIRKYAKIIIEWGPFEYMVLLTIIANCIVLALEEHLPEMDKTPLALQLDDTEVYFLGIFCVEAFLKIVALGFCLHKGSYLRNVWNIMDFIVVVTGFITLFASSGSDLRTLRAVRVLRPLKLVSGIPSLQVVLKSILRAMAPLLQVCLLVLFAIVIFAIIGLEFYVGVFHSACFRTEDDIDLGDEISWPCDAAFRCQ-VGPNDGITSFDHIGYAMLTVFQCITMEGWTTVLYYTNDALGWFNYLYFIPLIIVGSFFMLNLVLGVLSGEFARRQQQIERELNGYLEWICKAEEVIQLKGEDTDNDNEQNDDDLLAERLRYSIRRLVKSQLFYWIVIVLVLLNTISVASEHYNQPEWFVDFLYITEYAFLGLFIFEMSLKMYALGVRLYFQSSFNIFDCVVIVGSIFEVIKQDSFGFSVLRALRLLRIFKVTRYWASMRNLVISLLSSMRSILSLLFLLFLFILIFALLGMQLFGGKMNFEDGRPSAHFDTFPIALLTVFQILTGEDWNEVMYDGIRAHGMLASSYFIVLVLFGNYTLLNVFLAIAVDNLANAQELTAAEEEQEEEEAVRREEVENVDSGPPTKTSSVTMPLAKNAEEDMLPYSSMFIFYPTNPIRQFCHFVVNLRYFDLFIMIVICASSVALAAEDPV-VAVSGRNDILNYFDFVFTGVFTIELILKVIDLGIILHPYIRDLWNILDATVVICALVAFAFNNLNTIKSLRVLRVLRPLKTINRVPKLKAVFDCVVNSLKNVSNILIVYILFQFIFAVIAVQLFKGRFFYCTDESKSTRDDCQGLRQDFHYDDLANAMLTLFTVTTGEGWPSVLKHSMDSTQENRGPKPGSRMEMAIFYVVFFIVFPFFFVNIFVALIIITFQEQGENELMDQEMDKNQKQIDFAINAKPLCRFMPVKYKIWKLVQSPKFEYFIMTLITLNTIVLMMREKGEDRILHLINTVFTSLYGLGFLLKLCAYGK-NYFHDPWNVFDLITVIGSIIDVVISE--SMGRVSFGFFRLFRAARLVKLLRQGYTIRLLLWTFFQSFKALPYVCLLILMLFFIFAIIGMQVFGSIKLDSTEITRHNNFRTFFSALTLLFRRATGEAWQQIMWSCLAESGCGSNIAYIYFVSFIFLCSFLMLNLFVAVIMDNFDYLTR

>Anolis_carolinensis_Cav2.3_XP_008107103.1

SLFIFGEDNIVRKYAKKLIDWPPFEYMILATIIANCIVLALEQHLPEDDKTPMSRRLEKTEPYFIGIFCFEAGIKIVALGFVFHKGSYLRNGWNVMDFIVVLSGILATAGTHFNDLRTLRAVRVLRPLKLVSGIPSLQIVLKSIMKAMVPLLQIGLLLFFAILMFAIIGLEFYSGKLHRACYVNNSGELQELDPPHPCG-VQGCP-IGPNDGITQFDNILFAVLTVFQCITMEGWTTVLYNTNDALGTWNWLYFIPLIIIGSFFVLNLVLGVLSGEFARRQQQIERELNGYRAWIDKAEEVMRNRTEVMNRDSSDERVDISSVLLRISVRHMVKSQVFYWLVLSIVALNTACVAIVHHNQPPWLTHLLYYAEFIFLGLFLLEMSLKMYGMGPRLYFHSSFNCFDCGVTVGSIFEVVPGTSFGISVMRALRLLRIFKVTKYWASLRNLVVSLMSSMKSIISLLFLLFLFIVVFALLGMQLFGGRFNFMDGTPSANFDTFPAAIMTVFQILTGEDWNEVMYNGIRSQGMWSSIYFIVLTLFGNYTLLNVFLAIAVDNLANAQELTKDEQEEEEAFNQKHALHETSPSEGHLDLNQSKDVSPVEQDGNMVPHSSMFIFSTTNPIRRACHYIVNLRYFEMCILLVIAASSIALAAEDPV-LTNSERNKVLRYFDYVFTGVFTFEMVIKMIDQGLILQDYFRDLWNILDFIVVVGALMAFALADIKTIKSLRVLRVLRPLKTIKRLPKLKAVFDCVVTSLKNVFNILIVYKLFMFIFAVIAVQLFKGKFFYCTDSSKDTQKDCIGKRHEFHYDNIIWALLTLFTVSTGEGWPQVLQHSVDVTEEDRGPSRSNRMEMSIFYVVYFVVFPFFFVNIFVALIIITFQEQGDKMMEECSLEKNERAIDFAISAKPLTRYMPFQYRVWHFVVSPSFEYTIMAMIALNTIVLMMAPYTYELALKYLNIAFTMVFSLECVLKIIAFGFLNYFRDTWNIFDFITVIGSITEIILTDLVNTSSFNMSFLKLFRAARLIKLLRQGYTIRILLWTFVQSFKALPYVCLLIAMLFFIYAIIGMQVFGNIKLDESHINRHNNFRSFLGSLMLLFRSATGEAWQEIMLSCLGSEQCGTDLAYVYFVSFIFFCSFLMLNLFVAVIMDNFEYLTR

>Anolis_carolinensis_Cav1.4_XP_016846283.1

ALFCLRLNNPIRRAAISIVEWKPFDILILMTIFANCVALGVYIPFPEDDSNVANHNLEQVEYIFLIIFTVETFLKILAYGLVMHPSAYIRNGWNLLDFVIVVVGLFSVILEQFSDVKALRAFRVLRPLRLVSGVPSLHIVLNSIMKAMVPLLHIALLVLFVIIIYAIIGLELFIGRMHKTCFI-IGSDLEAEEDPSPCAFGRECT-EGPNGGITNFDNFFFAMLTVFQCITMEGWTDVLYWMQDAMGELPWLYFVSLVIFGSFFVLNLVLGVLSGEFSREKQQMEEDLQGYLDWIMQAEDIERHSSSTDTHSNFLIFLHMSPYLLRKRCRLAVKSVSFYWMVLILVFLNTLTIASEHYNQPDWLTQIQAYANKVLLSLFTLEMLLKMYSLGLQAYFVSFFNRFDCFVVCGGILETVIMQPLGISVLRCVRLLRIFKVTRHWASLSNLVASLLNSMKSIASLLLLLFLFIIIFSLLGMQLFGGKFNDETQTKRSTFDTFPQALLTVFQILTGEDWNTVMYDGIMAYGMLVCVYFIILFICGNYILLNVFLAIAVDNLADGDNINTSKNKEAPAEGEQSNETEEKDVKLECEEEEEEEEAEEGSEEAIPDGSSFFCLSKTNPLRVGCHKLIHHHIFTNLILVFIILSSISLAAEDPI-RAHSFRNN--------------------MTAFGGFLHQFCRNWFNLLDLLVVSVSLISFGIHAISVVKILRVLRVLRPLRAINRAKGLKHVVQCVFVAIRTIGNIMIVTTLLQFMFACIGVQLFKGKFYSCTDEAKHTPNECKGLNSDFNFDNVLSGMMALFTVSTFEGWPALLYKAIDANAENEGPIYNYRVEISIFFIIYIIIIAFFMMNIFVGFVIITFRAQGEQEYKNCELDKNQRQVEYALKAQPLRRYIPYQYKFWYMVNSTGFEYIMFVLILLNTIALAVQSQPFNYVMDLLNMVFTGLFTVEMVLKIIAFKPKHYFCDAWNTFDALIVVGSVVDIAVTESEDSSRISITFFRLFRVMRLVKLLSKGEGIRTLLWTFIKSFQALPYVALLIAMIFFIYAVIGMQTFGKVAMQDTPINRNNNFQTFPQAVLLLFRCATGEAWQEIMLASLEEFTCGSNFAIVYFISFFMLCAFLIINLFVAVIMDNFDYLTR

>Alligator_mississippiensis_Cav3.3_XP_019344978.1

--------------------M--------MVILLNCVTLGMYQPCEDMDLSDRCKILQVFDDFIFIFFAMEMVLKMVALGI-FGKKCYLGDTWNRLDFFIVMAGMVEYSLDLQNNLSAIRTVRVLRPLKAINRVPSMRILVNLLLDTLPMLGNVLLLCFFVFFIFGIIGVQLWAGLLRNRCFMPPYYQPEEDDEMFICSLGHECCNTNPHKGAINFDNIGYAWIVIFQVITLEGWVEIMYYVMDAHSFYNFIYFILLIIVGSFFMINLCLVVIATQFSMEPGDCYEEIFQYICHIVRKAKRRHCQKHNSLDYTPQALVQPIAVEVRVKLRGIVESKYFNRGIMIAILVNTISMGIEHHEQPEELTNILEISNVVFTSMFALEMILKLAAFGLFDYLRNPYNIFDSIIVIISIWEIIGQSDGGLSVLRTFRLLRVLKLVRFMPALRRQLVVLMKTMDNVATFCMLLMLFIFIFSILGMHIFGCKFSGDTVPDRKNFDSLLWAIVTVFQILTQEDWNVVLYNGMASTSPWASLYFVALMTFGNYVLFNLLVAILVEGFQAGDSYSDEDQSSSNAEELDRFQASVPPSHRATRKVGVTSGTTEHQDCNLREDWSIYLFSPQNRFRILCQTIIAHKLFDYIVLAFIFLNCITIALERPQIEHRSTERIFLSVSNYIFTAIFVAEMTLKVVSLGLYFGDYLRSSWNILDGFLVFVSLIDIVVSILGVLRVLRLLRTLRPLRVISRAPGLKLVVETLISSLKPIGNIVLICCAFFIIFGILGVQLFKGKFYHCLDIRITNRSDCVAVHHKYNFDNLGQALMSLFVLASKDGWVNIMYNGLDAVAVDQQPVTNNNPWMLLYFISFLLIVSFFVLNMFVGVVVENFHKCRQHQEAEEARRREEKRRRLEKKRRPYYAYCPVRLLIHSVCTSHYLDIFITFIICLNVVTMSLQPVSLETALKYCNYMFTTVFVLEAVLKLVAFGLRRFFKDRWNQLDLAIVLLSVMGITLEEALPINPTIIRIMRVLRIARVLKLLKMATGMRALLDTVVQALPQVGNLGLLFMLLFFIYAALGVELFGKLVCNDEGMSRHATFENFGMAFLTLFQVSTGDNWNGIMKDTLDDRSCLQFISPLYFVSFVLTAQFVLINVVVAVLMKHLDDSNK

>Alligator_mississippiensis_Cav2.1_XP_019354952.1

SLFLFSEDNVVRKYAKKITEWPPFEYMILATIIANCIVLALEQHLPDEDKTPMSERLDDTEPYFIGIFCFEAGIKIIALGFAFHKGSYLRNGWNVMDFVVVLTGILAKVG----DLRTLRAVRVLRPLKLVSGIPSLQVVLKSIMKAMIPLLQIGLLLFFAILIFAIIGLEFYMGKFHTTCFD---SVTGEIKDRVPCGMARTCP-EGPNYGITQFDNILFAVLTVFQCITMEGWTDLLYNSNDASGTWNWLYFIPLIIIGSFFMLNLVLGVLSGEFARRQQQIERELNGYMEWISKAEEVISKTDLLNPEEADDQLADIASVRMRFYIRRMVKTQAFYWTVLSLVALNTLCVAVVHYSQPDWLSDFLYYAEFIFLGLFMSEMFIKMYGLGTRPYFHSSFNCFDCAVIIGSIFEVVPGTSFGISVLRALRLLRIFKVTKYWASLRNLVVSLLNSMKSIISLLFLLFLFIVVFALLGMQLFGGQFNFDTGTPATNFDTFPAAIMTVFQILTGEDWNMVMYDGIKSQGMVFSVYFIVLTLFGNYTLLNVFLAIAVDNLANAQELTKDEQEEEEAATQKLALGPHLSTTRPIQQDMGRPEPPVAEDIDMVPYSSMFILSTTNPFRRLCHYIVNLRYFEMCILMVIAMSSIALAAEDPV-QPNAPRNNVTPFLDLVFTGVFTFEMVIKMVDLGLVLHQYFRDLWNILDFIVVSGALVAFAFTDINTIKSLRVLRVLRPLKTIKRLPKLKAVFDCVVNSLKNVLNILIVYMLFMFIFAVVAVQLFKGKFFYCTDESKEFENDCRGKKYEFHYDNVLWALLTLFTVSTGEGWPQVLKHSVDATYENQGPSPGYRMEMSIFYVVYFVVFPFFFVNIFVALIIITFQEQGDKMMEEYSLEKNERAIDFAISAKPLTRHMPFQYRMWQFVVSPPFEYTIMAMIALNTVVLMMASDAYENVLKMFNNVFTSLFSLECLLKIMAFGVLNYFRDAWNIFDFVTVLGSITDILVTE--GNNFINLSFLRLFRAARLIKLLRQGYTIRILLWTFVQSFKALPYVCLLIAMLFFIYAIIGMQVFGNIGIKDSAITEHNNFRTFFQALMLLFRSATGEGWQEIMLSCLMEHECGNEFAYFYFVSFIFLCSFLMLNLFVAVIMDNFEYLTR

>Algae_chlrei_CAV_XP_001701475.1

SLLFLRKSNFVRKWCVYTTHTRVFEWTILAAIIANCVTLAVSSNRQDFDETPLGRTLVNLEYLWVAIFTTEALLKIVAMGFVLAPGTYLRDGWNIVDFTVVALGFVDIFSSG--NLTALRTVRVLRPLRAITRIRGMRILVTTMIAALPMLIDVFALCAFTFFIFGLVAVQLFSGRMTHRCAVRNVTYVVQDEESDGCSGGRACP-GNPNYGITSFDHILWAWLTIFQMITQEGWTDIMYFTSDTITWWVWPFFVALVAFGSFYIINLALAVLFMQFSSGRQGSDGQRQG-----DKAGAGGSESSMSDVGSEEDERLRPTMAPFRRRLYRVAASRWLELFTMSLIILNTVLMCINWFEMPESVERATNYINYVFTVYFLVEMILKMTAFGLVRYFRDGMNIFDCLVVVISVTEMVSVSGLGLSVLRAFRLLRIFRLARSWKELNLIIRAIFKSITSTTYLLLLMLLFMFIASLMGMQLFGYKFCGKSEVPRANFDNIFWSMYTVFQILTMENWNNIMYDGMRSTTPWCAAYFVAVVLIGTYLVFNLFVAILLDNFSSGSSLKADGQAAGSGKHKSRRSPVDDEGRPGSSHSGSQPPRPSLRPVSQIKGRSLLLLDPEHWVRWRAARMVHHTHFETVILGLIVLSSITLALDSPGLDPDSQLAQALRYLDYIFLGAFTLEAALKIITFGFFTGKYIRNGWNVLDFIIVLAGYALLAVEDLKMLRILRTLRALRPLRAASRYEGLKLVVNTLFAVLPAMADVALVCALFYIIFSILAVNLFKGQLYNCIDADTLERGMCEAVNPVANFDNVAISMLTLFQIATLELWVDIMFTAVDVAGVGKQPLWNNHPVVILFFILVVIVCCFFVLNLFIGVTLDKFTELQQAQTASSVFVTPQQQLRTGMTSRPARFEPAWRAGLYDFVMGSVFEEFILITIVVNVLFMAMMSPQWQACMTYTNLIFTCVFVIEAALKIMAFGAFAYFRDRWNAFDFFVVVISVASVVLDFSGTQNLSFMPVLRVLRVVRVVRLIRRAVGMRRLLLTLVQSLPALGNVGGVMVLFFFVYAVIGMNLFGGIKF--DYISRHANFNNFGKAMLLLFRMITGESWSGVMQDCMHQNQCSPWAAVVYFPTFIVLCGFILLNLVIAIILENMITSEN

>Aedes_aegypti_Cav3_XP_021706157.1

SLKYLSQDTKPRIWCLQLITNPWFERVSMLVILLNCVTLGMYQPCVDDAVTNRCKILQIFDDIIFAFFSLEMTIKIVAMGA-WGKGTYLADSWNRLDFFIVLAGALEYCLQVENNLTAIRTIRVLRPLRAINRIPSMRILVMLLLDTLPMLGNVLLLCFFVFFIFGIVGVQLWEGILRQRCVIDISFYYEFSKEQYICSKPLVCNGNNPFQGTISFDNIGLAWVAIFLVISLEGWTDIMYYVQDAHSFWDWIYFVLLIVIGSFFMINLCLVVIATQFSSEPTTCYAEIVKYIAHLYRRLKRRQTMATVFNDYSDLCMTDAMTCCIRVYIKKLVEHKYFQQGILLAILINTLSMGIEYHNQPEELTAIVETSNIVFSGIFAVEMVLKVIAEGPFGYVANGFNVFDGVIVILSVVELGGQGSSGLSVLRTFRLLRILKLVRFMPNLRRQLFVMLRTMDNVAIFFSLLILFIFIFSILGMYLFGGKFCPLCECDRKHFNNILWATVTVFQILTQEDWNVVLFNGMEKTSHWAALYFVTLMTFGNYVLFNLLVAILVEGFSSYDSFSESTTSAEELRKIRDIKIDSGPGSIASNISAPSYSKYIYNDKNERDLYTLYIFPEDNRFRQICSWFVNQKWFDNVILLFIALNCITLAMERPNIPPTCTERYFLSTANYVFTVVFAVEMFIKVVATGMFYGRYFTSGWNIMDGSLVIISIVDLLMSIFGILRVFRLLRSLRPLRVINRAPGLKLVVQTLLSSLRPIGNIVLICCTFFIIFGILGVQLFKGTFYYCENIKVKNKTECLSVNRKYNFDDLGKALMSLFVLSSRDGWVNIMYTGLDAVGVDQQPVVNYNEWRLLYFIAFILLVGFFVLNMFVGVVVENFHRCREEQEKEEKIRRAAKRLQMEKKRRPYYTYSPLRMFVHNVVTSKYFDLAIAAVIGLNVVTMAMMPRALEYALKIFNYFFTAVFILEAIMKLVALGLKIYMKDKWNQLDVAIVILSIVGIVLEEIIPINPTIIRVMRVLRIARVLKLLKMAKGIRALLDTVMQALPQVGNLGLLFFLLFFIFAALGVELFGRLECSEQGLGEHAHFANFGMAFLTLFRVATGDNWNGIMKDTLVKNCCVTIIAPIFFVIFVLMAQFVLVNVVVAVLMKHLEESHK

>Aedes_aegypti_Cav2_XP_021710362.1

SLFILSEDNIIRKYTRFIIEWPPFEYAVLLTIIANCVVLALEEHLPHGDKTLLAQKLEKTEAYFLGIFCVEASLKILALGFVMHKHSYLRNIWNIMDFFVVVTGFITLFPQEGPDLRTLRAIRVLRPLKLVSGIPSLQVVLKSIIKAMAPLLQIGLLVLFAIVIFAIIGLEFYSGALHRSCYSDISQIVKEGEFPTPCNAAYVCN-EGPNFGITSFDNIGFAMLTVFQCITMEGWTAILYWTNDALGTFNWIYFVPLIVLGSFFMLNLVLGVLSGEFARRQQQLEKELNGYVEWICKAEEVI--RNLGKSKSTDTEEDDPDDDRFRFWIRHTVKTQWFYWFVIVLVFFNTVCVAVEHYGQPNWLTQFLYYAEYVFLGLFMMEMWIKMYALGPRIYFESSFNRFDCVVISGSIFEVVKGGSFGLSVLRALRLLRIFKVTKYWSSLRNLVISLLNSMRSIISLLFLLFLFILIFALLGMQLFGGQFNLPDGTPPTNFNTFPIALLTVFQILTGEDWNEVMYQGIESQGMIYSLYFIILVLFGNYTLLNVFLAIAVDNLANAQELTAAEEEQMEENKEKQQMPRVEVSSPSPTRGNGSKANKKEEDKEMLPYSSMFVLSPTNPIRCAAHWVVNLRYFDFFIMVVISLSSIALAAEDPV-EEDSPRNKILNFFDYAFTGVFTIEMLLKIVDLGVILHPYLREFWNIMDAVVVICAAVSFGFDNLSTIKSLRVLRVLRPLKTIKRVPKLKAVFDCVVNSLKNVINILIVYILFQFIFAVIAVQLFNGKFFYCTDDSKHTSEECKGKTQSFHYDNVATAMLTLFAVQTGEGWPQVLQNSMAATYEDKGPIQNFRIEMSIFYIVYFIVFPFFFVNIFVALIIITFQEQGEAELQDGEIDKNQKSIDFTIGARPLERYMPFKYKVWRIVVSTPFEYFIMMLIVFNTLLLMMQGKEFEKSLKYLNMGFTGMFSVETILKIIGFGVKNFFKDPWNIFDFITVIGSIIDAVL--ELGENSFNVGFLRLFRAARLIKLLRQGYTIRILLWTFVQSFKALPYVCLLIAMLFFIYAIIGMQVFGNIELEPSAITRHNNFRSFVQGLMLLFRCATGESWPNIMLACLSNETCGSTLAYAYFVSFIFFCSFLMLNLFVAVIMDNFDYLTR

>Aedes_aegypti_Cav1_XP_021699870.1

ALFCLTLKNPLRKLCIDIVEWKPFEYLILLTIFANCVALAVYTPFPNSDSNTTNAALEKIEYIFLVIFTAECVMKLIAYGFIMHPGSYLRNGWNILDFTIVVIGMISTALSNLMDVKALRAFRVLRPLRLVSGVPSLQVVLNSILRAMVPLLHIALLVLFVIIIYAIIGLELFSGKLHKTCFHLFLFLDEIMDDPHPCGDGFQCA-AGPNFGITNFDNFGLSMLTVFQCVTLEGWTDMLYYIQDAMGTWQWVYFISMVILGAFFVMNLILGVLSGEFSREKQQIEEDLRGYLDWITQAEDIDTANEIDSSDHMGEEG------RLRRACRKAVKSQAFYWLIIVLVFLNTGVLATEHYQQPPWLDDFQEYTNMFFVALFTMEMLLKMYSLGFQGYFVSLFNRFDCFVVIGSIGEMVIMPPLGVSVLRCVRLLRVFKVTKYWQSLSNLVASLLNSIQSIASLLLLLFLFIVIFALLGMQVFGGKFNSETDKPRSNFDSFVQSLLTVFQILTGEDWNAVMYDGIQAYGILASIYFIILFICGNYILLNVFLAIAVDNLADADSLTTVEKEEGEGEEGADVEEMNISEDYEHNGSETKMTLPDDDEGYIPPGSSFFIMSQTNRFRVFCHWLCNHSTFGNIILVCIMFSSAMLAAEDPL-NANSERNQILNYFDYFFTTVFTIELLLKVISYGFLFHDFCRSAFNLLDLLVVCVSLISMFFSAISVIKILRVLRVLRPLRAINRAKGLKHVVQCVIVAVKTIGNIVLVTCLLQFMFAVIGVQLYKGKFFSCSDGSKMQESECHGSRNRFHFDDVSKAMLTLFTVSTFEGWPGLLYVSIDSHEEDSGPIHNFRPIVAAYYIIYIIIIAFFMVNIFVGFVIVTFQNEGEQEYKNCDLDKNQRNIEFALKAKPIRRYIPIQYKVWWFVTSQPFEYMIFILIMINTITLSMQPEIYTEVLDLLNLIFTAVFALEFVFKLAAFRFKNYFGDAWNVFDFIIVLGSFIDIVYSEKGGSSIISINFFRLFRVMRLVKLLARGEGIRTLLWTFIKSFQALPYVALLIVMLFFIYAVIGMQVFGKIAMDDTSIHRNNNFQTFPQAVLVLFRSATGEAWQDIMLDCSSPEGCGSSIAFPYFISFYVLCSFLIINLFVAVIMDNFDYLTR

>Acropora_digitifera_Cav3b_XP_015766817.1

--------------------M--------FVIFVNCVILAMYNPLDENCTETRCRVLENVEHFVFAFFCAEIVIKMVAMGV-TGNRGYLQDKWNRLDCLIVMIGLIEKVISNNNYLTIIRAFRVLRPLRAINKVPSIRILVTLLLDTLPMLGNVLLLSFLIFFVFGIISVQLWQGKLRNRCFTRSTFYKPSFDEPFVCSLDTNCSGPNPFYDLVSFDNIGIAWIVIFQVITLEGWSDIMYFVQDAHSSLSWIYFVVLIVVGSFFLVNLCLVVITMQFQSEPLRAVMKFAKNLCWCRTEDEPQSPQSANVSSYSPSEMLDQNMSSLRALCRKVAASKHFTFFIMVVILLNMICMAPEHYDQPEFLTDAMEITNKIFVCIFSFEMVIKLLGDGIAMYLSSGQNVFDGIIVIVSVCEILSRQTSALSVFRSIRLLRIFKLVR---PVRYQLLVVIKTMTSVMTFFGLLFLFIFAFAILGMNLFGGKFEGKSVTSRSNFDSFLWAMVSVFQILTQENWNLVMYDGMRTTNKWAALYFTALMAIGYYVLFNLLVAILVEGFTNGSALS----VFRSIRLLRIFKGEPKQTPKTEEQKGQNSNAGKKDTPKTRQNWSLYLFAPSNRFRCRMASVCEHKYFDYVVLMFILISCIVLAMEEPN--ILPKKRQIIDATMLLLTVIFTCEMMMKDVTPSV------------------LCLLTSF------------------HHRMIRRAPGLKLVVQTLLYSLKPIGNTVLIAAIFFVMFGILGVQIFKGKFHYCKNDNVTNKAECEQENRMYNFDNLAQALISLFVFSTRDGWVKIMHNGIDAVGIDQQPITNYAEWRLVYFIPFLMLGGFLVLNMIVGVVVENFQRCRERLEDEEKQKKRKKLVKTEKQEDNYSEYSKWRLRILGICLHPYWDVSIAIVILVNVICMSLMPKSLEVFVEIANYFFTGVFVLEVVVKFIAFGFARYFSDRWNLVDLVIVVLSLAGIIIESKIMINPTVARSLRVLRFIRVLKLVKLAKGVRSLLDTLFEALPQVANLGLLFLLLFFIYSCLGIQLFGNLKCCPQGLGPHAHFRDFGTAMLTLFRIATGDNWNGILEDTLEEYCCVKYTAPLFFFTFVLAAQFVLVNVVIAVLMKHLKESKE

>Acropora_digitifera_Cav3a_XP_015765864.1

ACFFLRSESPPRSWFIRLVTWPYFERVSIFVILLNCVTLGLYDPFDPECSSQRCQTLDTMEKIIYAFFLVEMLCKWMAMGL-FGKMSYFADPWNRLDCFIVAAGTFELLYDKGEYLSAVRAIRVLRPLRAINRVPSIRILVTLLLDTLPMLWNVLAICFFIFAIFGIVAVQLWQGALRGRCFMINEFFNYNVSDMFVCELGLQCSGDNPAWGAIGFDNIFIAWVAIFQVITLEGWADIMYFVQDAHGFWNWIYFVILIVIASYFLTNLCLVVITTQFQYGKDGCWVEILKYIEHVCRRLKRRSQGEMITTATAAVSLNGKSAVRFRRYCRRSVDSKWFMYIIMGSIFLNTLSMGIEYHGQSSI---------------------VELFGEG------------DSS----------------ISVLRSFRLLRIFKLVRFLPALRRQLLVMIHTMDNVVTFLALLALFIFTASVLGMNLFGGKYTGGKVTARANFDDLFWALVTVFQVLTQEDWNTVMYDGMRATTKWAALYFILLMTIGNYILFNLLVAILVEGFANQPASTWSIKSKQSKQDQDHVTGVLTQEDWNTVMYDGMRATTKWAALY--GSLALFVLQCLERFRKLMIATYSNKWFDRVVLVFILLNCVVMALERPDLPKDSELQKIIDICMYIFLGIFTLEMFIKVMALGLWVGPYLRSSWNVMDGFLVVVSWIDVIVTILGVLRVFRALRTLRPLRVISRAPGLKIVVETLISSLKPIGNIVLIAATFFIIFGILGVQLFKGKFYHCKGASVETRAECTSVNKEYNFDNLARALLTLFVFSTKDGWVTIMYDGLDSVGVDKQPKRNNNKWNVLYFVAFLLLAGFVVLNMLVGVVVENFQKCRDIIEKDRQVEKEKEKQERARKRQKLAEFLQPRRFFHRICTHGYFDLGISAVIVLNVICMAMQPQEMRDFLKYANYVFTAVFVMEGILKIFALGFKKYIKERWNQLDLIIILLSIVGIVLEESLPINPTIIRVMRVLRIARVLKLLKTAEGIRKLLDTVAEALPQVGNLGLLFLLMFFIFAALGMELFGQINC-DEGLDNHAHFRNFGFAMLTLFRVSTGDNWNGILKDIIPPPGCAEHIAPIYFAFFVLVTQFVLLNVVVAVLMKHLE--DA

>Acropora_digitifera_Cav2a_XP_015773841.1

SLFIFSKDNLIRKICRTIVESKPFEYFILLTIFVNCILLAANKPLPKEDKSDLNVELEKAEIYLLAIFCLEAALKIVALGFLLHSDSYLRNGWNVLDFVVVVTGLLSLPELNIGSLKALRAARVLRPLKLVSGIPSLQVVMKSIMCAMVPLLQICLLVGFVVIIYAIIGLEFLNGKFHYVCHNNETGKIENPDTPQICDPGRSCK-EGPNDGITSFDNIFAGMLTVFQVITNEGWTDIMYWTFDAADYAFWIYYCSLVIIGSFFMLNLVLGVLSGEFARRTQKMERHLHGYIDWISKAEDLMRRRNLDRVEDGDIAMTAQVLTRWKIRIRQIVKHQAFYWVVLVCVFLNTLITALQHYRQPEWLTQFQDIAEIVFISFFFCEMTMKLYGLGPQLYFKSQFNTFDCVVVCCGILELIQGISLGISVLRALRLLRLFKFTRYWSSLRNLVTSLLSSVRSILSLLFLLFLFIVIFALLGMQIFGARFRGRKGNPRTNFDDFANAALAVFQILTGEDWNAVMYDGVLSYDALWAIYFVLLVVLGNYVLLNVFLAIAVDNLANAQQLSQDEEAEETEREERKKNLGPNEDEFESEMDDRPSFISNLRNPGIIDTWSLFLFPPGNPVRKACHWLVNLRYFDNTILVIILISSVLLALEDPV-VEGSYRNRVLTYFDYVFTTIFALEVIVKLIDYGAILHPYFRDAWNCIDCLVVCCAVASLVMGSKKIVKVLRVLRVLRPLKAINKAKKLKAVFQCMVYSLKNVLNILIITILFLFIFSVIGVQLFQGKFFYCDDASKMTEEECQGGKHEYNFDNVLHAMLALFTSSTGEGWPALMQNSIDATEVDKGPITDNKIEIALFYIFFVVVFSFFFINIFVALIILTFQEQGEKDQGDCELDRNQRDLHFAIVAKPSERFMPWQYRIWRIVDSRPFEYFIMLLIALNTLILTMEPKLYRDILDIFNTIFTFMFTAEAILKLFAFRL-NYFRDGWNVFDFIIVLGSLLDFLQQGGAKQMPFDPSLFRLFRAARLIKLLRQGYTIRILLWTFLQSFKALPYVGMLIGLLFFIYAVIGMQMFGQISKESAQISSQNNFQSFPQAIQVLFRSATGENWQLIMLACAPPETCGSSFTYVYFITFIFFCSFLLLNLFVAVIMDNFEYLTR

>Acropora_digitifera_Cav1_XP_015778662.1

ALLCLSLGNPIRSAAINLVEWKPFDVMILITIFANCAALAAYEPLPGRDSSEVNEGLEIAEYVFLAIFTLEAILKIIAYGFFFHSGAYLRNGWNILDFVIVVVGXVNLVTIWELDNIALACMTVFQVLNLVLGVSSLSNLVASLLNSMRSIAGLLLLLSLFMLICSLLGMQIFGGSIPLPCKDCLFQGAEAMENPHPCSSGFHCN-KGPNYGITNFDNIALACMTVFQCITLEGWTDVLYMINDAVGSWPWIYFVTLIIWGSFFVLNLVLGVLSGEFAREKQQVEDAYNGYLDWITQAEDIERR-ASRHSRIDDIEMIDKNERRTRRELRKAVKTQAFYWIVIVVVFLNSLTLALEHYGQPHFLTIFLEASASVFREVTKHNLLLPVSTCPLRDLFLVLFLVTCCIVVVSSLLELASQRPIGISVLRCIRLLRIFKVTRYWSSLSNLVASLLNSMRSIAGLLLLLSLFMLICSLLGMQIFGGRFSDGEDVPRSNFDSFWKALITVFQILTGEDWNTVMYDGIRSWGILAILYFIFLVVVGNYILLNVFLAIAVDNLADAENLTEMEEEKKRRKXKAKEKLDKSNQELHSAGNLNGNAVAQTASHSMPPESALFIF-----------------------------SSTNIQSNSP----------VLNYFDYFFTSVFTLEILIKFVAYGLILHKFCRSAFNLLDLLVVSVSVISISLKQFSVVRILRVLRVLRPLRAINRAKGLKHVVQCVFVAVKTIWNIMLVTMLFNFLFAVIGVQLWKGTFFYCTDQKKRFEDECKGKRRDFNFDNVGNAMLTLFTVMTFEGWPGILYNSIDSTEVDEGPLQNNRPWVAVYYIIYIIIIAFFMVNIFVGFVIVTFQSEGEEEFKDCELDKNQ-------------------------VSKRSF---------------------------------------------------NYFIDRWNLFDFIIVVGSIIDITMNEVSSEQMFAFGFFRLFRALRLVKLLNQGSGIKTLLWTFIKSFQALPYVALLIVMMFFIYAVIGMQMFGRIALDPTAINRNNNFQTFPHSLMVLFRSATGENWQEIMLSCTPSGLCGSDFAYFYFVSFYSICSFLIINLFVAVIMDNFDYLTR
